## Supplementary material for "Game Bird Carcasses are Less Persistent Than Raptor Carcasses, But Can Predict Raptor Persistence Dynamics": S1 Appendix

**S1 Table. Full meta-analysis dataset.** Table includes all raw data from the data curation effort (in addition to the data collected during our Field Trials). Trials are aggregated by study (the trial placement effort during a period up to a year, at a single facility), predominant habitat at the facility, Region, and season.

| **Species** | **Class** | **Season** | **State** | **County** | **USFWS Region** | **Habitat** | **Study ID** | **Date Placed** | **Last Known Present (Days Since Placed)** | **First Known Absent (Days Since Placed)** | **Used in Scalar Model** | **Analysis Group** |
| --- | --- | --- | --- | --- | --- | --- | --- | --- | --- | --- | --- | --- |
| red-tailed hawk | Raptor | winter | CA | Kern | 8 | shrub/scrub | R8CA-ss-5 | 2017-12-11T00:00:00Z | 21.0152778 | 29.20625 | 1 | R8CA-ss-0 |
| red-tailed hawk | Raptor | winter | CA | Kern | 8 | shrub/scrub | R8CA-ss-5 | 2017-12-11T00:00:00Z | 84.0229167 | 91.0958333 | 1 | R8CA-ss-0 |
| red-tailed hawk | Raptor | winter | CA | Kern | 8 | shrub/scrub | R8CA-ss-5 | 2017-12-11T00:00:00Z | 91.2409722 | NA | 1 | R8CA-ss-0 |
| red-tailed hawk | Raptor | winter | CA | Kern | 8 | shrub/scrub | R8CA-ss-5 | 2017-12-11T00:00:00Z | 91.1493056 | NA | 1 | R8CA-ss-0 |
| barn owl | Raptor | winter | CA | Kern | 8 | shrub/scrub | R8CA-ss-5 | 2017-12-11T00:00:00Z | 20.9152778 | 29.1402778 | 1 | R8CA-ss-0 |
| red-tailed hawk | Raptor | spring | CA | Kern | 8 | shrub/scrub | R8CA-ss-6 | 2018-05-14T00:00:00Z | 91.0944444 | NA | 0 | R8CA-ss-6 |
| osprey | Raptor | spring | CA | Kern | 8 | shrub/scrub | R8CA-ss-6 | 2018-05-14T00:00:00Z | 90.7840278 | NA | 0 | R8CA-ss-6 |
| red-shouldered hawk | Raptor | spring | CA | Kern | 8 | shrub/scrub | R8CA-ss-6 | 2018-05-14T00:00:00Z | 90.7729167 | NA | 0 | R8CA-ss-6 |
| red-tailed hawk | Raptor | spring | CA | Kern | 8 | shrub/scrub | R8CA-ss-6 | 2018-05-14T00:00:00Z | 90.84375 | NA | 0 | R8CA-ss-6 |
| turkey vulture | Raptor | spring | CA | Kern | 8 | shrub/scrub | R8CA-ss-6 | 2018-05-14T00:00:00Z | 50.0277778 | 57.0166667 | 0 | R8CA-ss-6 |
| great horned owl | Raptor | summer | CA | Kern | 8 | shrub/scrub | R8CA-ss-6 | 2018-07-23T00:00:00Z | 91.9423611 | NA | 0 | R8CA-ss-6 |
| red-tailed hawk | Raptor | summer | CA | Kern | 8 | shrub/scrub | R8CA-ss-6 | 2018-07-23T00:00:00Z | 92.0451389 | NA | 0 | R8CA-ss-6 |
| barn owl | Raptor | summer | CA | Kern | 8 | shrub/scrub | R8CA-ss-6 | 2018-07-23T00:00:00Z | 91.9756944 | NA | 0 | R8CA-ss-6 |
| red-tailed hawk | Raptor | summer | CA | Kern | 8 | shrub/scrub | R8CA-ss-6 | 2018-07-23T00:00:00Z | 91.9902778 | NA | 0 | R8CA-ss-6 |
| red-tailed hawk | Raptor | summer | CA | Kern | 8 | shrub/scrub | R8CA-ss-6 | 2018-07-23T00:00:00Z | 91.9104167 | NA | 0 | R8CA-ss-6 |
| peregrine falcon | Raptor | summer | CA | Kern | 8 | shrub/scrub | R8CA-ss-6 | 2018-07-23T00:00:00Z | 42.9527778 | 49.0951389 | 0 | R8CA-ss-6 |
| osprey | Raptor | spring | CA | Kern | 8 | shrub/scrub | R8CA-ss-6 | 2018-04-02T00:00:00Z | 92.1611111 | NA | 0 | R8CA-ss-6 |
| red-shouldered hawk | Raptor | spring | CA | Kern | 8 | shrub/scrub | R8CA-ss-6 | 2018-04-02T00:00:00Z | 92.1243056 | NA | 0 | R8CA-ss-6 |
| red-tailed hawk | Raptor | spring | CA | Kern | 8 | shrub/scrub | R8CA-ss-6 | 2018-04-02T00:00:00Z | 92.1270833 | NA | 0 | R8CA-ss-6 |
| Cooper's hawk | Raptor | spring | CA | Kern | 8 | shrub/scrub | R8CA-ss-6 | 2018-04-02T00:00:00Z | 92.1256944 | NA | 0 | R8CA-ss-6 |
| great horned owl | Raptor | spring | CA | Kern | 8 | shrub/scrub | R8CA-ss-6 | 2018-04-02T00:00:00Z | 92.1097222 | NA | 0 | R8CA-ss-6 |
| great horned owl | Raptor | winter | CA | Kern | 8 | shrub/scrub | R8CA-ss-6 | 2018-11-26T00:00:00Z | 90.9805556 | NA | 0 | R8CA-ss-6 |
| red-tailed hawk | Raptor | winter | CA | Kern | 8 | shrub/scrub | R8CA-ss-6 | 2018-11-26T00:00:00Z | 90.9520833 | NA | 0 | R8CA-ss-6 |
| turkey vulture | Raptor | winter | CA | Kern | 8 | shrub/scrub | R8CA-ss-6 | 2018-11-26T00:00:00Z | 76.95625 | 85.1902778 | 0 | R8CA-ss-6 |
| red-tailed hawk | Raptor | winter | CA | Kern | 8 | shrub/scrub | R8CA-ss-6 | 2018-11-26T00:00:00Z | 50.825 | 55.8520833 | 0 | R8CA-ss-6 |
| red-tailed hawk | Raptor | winter | CA | Kern | 8 | shrub/scrub | R8CA-ss-6 | 2018-11-26T00:00:00Z | 42.8958333 | 50.9194444 | 0 | R8CA-ss-6 |
| turkey vulture | Raptor | winter | CA | Kern | 8 | shrub/scrub | R8CA-ss-6 | 2018-11-26T00:00:00Z | 9.875 | 13.8833333 | 0 | R8CA-ss-6 |
| turkey vulture | Raptor | winter | CA | Kern | 8 | shrub/scrub | R8CA-ss-6 | 2018-11-26T00:00:00Z | 9.9791667 | 13.9854167 | 0 | R8CA-ss-6 |
| great horned owl | Raptor | winter | CA | Kern | 8 | shrub/scrub | R8CA-ss-6 | 2018-11-26T00:00:00Z | 4.0354167 | 6.9423611 | 0 | R8CA-ss-6 |
| mallard | Game Bird | spring | CA | Kern | 8 | shrub/scrub | R8CA-ss-7 | 2014-05-05T00:00:00Z | 37 | NA | 0 | R8CA-ss-7 |
| mallard | Game Bird | spring | CA | Kern | 8 | shrub/scrub | R8CA-ss-7 | 2014-05-05T00:00:00Z | 37 | NA | 0 | R8CA-ss-7 |
| mallard | Game Bird | spring | CA | Kern | 8 | shrub/scrub | R8CA-ss-7 | 2014-05-05T00:00:00Z | 37 | NA | 0 | R8CA-ss-7 |
| mallard | Game Bird | summer | CA | Kern | 8 | shrub/scrub | R8CA-ss-7 | 2014-07-29T00:00:00Z | 40 | NA | 0 | R8CA-ss-7 |
| mallard | Game Bird | summer | CA | Kern | 8 | shrub/scrub | R8CA-ss-7 | 2014-07-29T00:00:00Z | 40 | NA | 0 | R8CA-ss-7 |
| mallard | Game Bird | summer | CA | Kern | 8 | shrub/scrub | R8CA-ss-7 | 2014-07-29T00:00:00Z | 4 | 7 | 0 | R8CA-ss-7 |
| mallard | Game Bird | fall | CA | Kern | 8 | shrub/scrub | R8CA-ss-7 | 2014-09-22T00:00:00Z | 1 | 1 | 0 | R8CA-ss-7 |
| mallard | Game Bird | fall | CA | Kern | 8 | shrub/scrub | R8CA-ss-7 | 2014-09-22T00:00:00Z | 25 | 40 | 0 | R8CA-ss-7 |
| mallard | Game Bird | fall | CA | Kern | 8 | shrub/scrub | R8CA-ss-7 | 2014-09-22T00:00:00Z | 25 | 40 | 0 | R8CA-ss-7 |
| mallard | Game Bird | fall | CA | Kern | 8 | shrub/scrub | R8CA-ss-7 | 2014-09-22T00:00:00Z | 1 | 1 | 0 | R8CA-ss-7 |
| red-tailed hawk | Raptor | fall | CA | Kern | 8 | shrub/scrub | R8CA-ss-7 | 2014-10-31T00:00:00Z | 4 | 7 | 1 | R8CA-ss-0 |
| mallard | Game Bird | fall | CA | Kern | 8 | shrub/scrub | R8CA-ss-7 | 2014-11-14T00:00:00Z | 40 | NA | 0 | R8CA-ss-7 |
| mallard | Game Bird | fall | CA | Kern | 8 | shrub/scrub | R8CA-ss-7 | 2014-11-14T00:00:00Z | 40 | NA | 0 | R8CA-ss-7 |
| mallard | Game Bird | winter | CA | Kern | 8 | shrub/scrub | R8CA-ss-7 | 2014-12-31T00:00:00Z | 18 | NA | 0 | R8CA-ss-7 |
| mallard | Game Bird | winter | CA | Kern | 8 | shrub/scrub | R8CA-ss-7 | 2014-12-31T00:00:00Z | 1 | 1 | 0 | R8CA-ss-7 |
| mallard | Game Bird | winter | CA | Kern | 8 | shrub/scrub | R8CA-ss-7 | 2015-01-20T00:00:00Z | 40 | NA | 0 | R8CA-ss-7 |
| mallard | Game Bird | winter | CA | Kern | 8 | shrub/scrub | R8CA-ss-7 | 2015-01-20T00:00:00Z | 25 | 40 | 0 | R8CA-ss-7 |
| red-tailed hawk | Raptor | winter | CA | Kern | 8 | shrub/scrub | R8CA-ss-7 | 2015-03-08T00:00:00Z | 40 | NA | 1 | R8CA-ss-0 |
| mallard | Game Bird | spring | CA | Kern | 8 | shrub/scrub | R8CA-ss-3 | 2015-05-05T00:00:00Z | 1 | 2 | 1 | R8CA-ss-0 |
| mallard | Game Bird | summer | CA | Kern | 8 | shrub/scrub | R8CA-ss-3 | 2015-07-10T00:00:00Z | 10 | 14 | 1 | R8CA-ss-0 |
| mallard | Game Bird | winter | CA | Kern | 8 | shrub/scrub | R8CA-ss-3 | 2015-12-09T00:00:00Z | 12 | NA | 1 | R8CA-ss-0 |
| mallard | Game Bird | winter | CA | Kern | 8 | shrub/scrub | R8CA-ss-3 | 2015-12-09T00:00:00Z | 2 | 6 | 1 | R8CA-ss-0 |
| mallard | Game Bird | spring | CA | Kern | 8 | shrub/scrub | R8CA-ss-3 | 2016-03-16T00:00:00Z | 15 | NA | 1 | R8CA-ss-0 |
| mallard | Game Bird | spring | CA | Kern | 8 | shrub/scrub | R8CA-ss-3 | 2016-03-16T00:00:00Z | 15 | NA | 1 | R8CA-ss-0 |
| mallard | Game Bird | winter | CA | Kern | 8 | shrub/scrub | R8CA-ss-4 | 2016-12-12T00:00:00Z | 10 | NA | 1 | R8CA-ss-0 |
| mallard | Game Bird | winter | CA | Kern | 8 | shrub/scrub | R8CA-ss-4 | 2016-12-12T00:00:00Z | 10 | NA | 1 | R8CA-ss-0 |
| mallard | Game Bird | winter | CA | Kern | 8 | shrub/scrub | R8CA-ss-4 | 2016-12-12T00:00:00Z | 10 | NA | 1 | R8CA-ss-0 |
| mallard | Game Bird | spring | CA | Kern | 8 | shrub/scrub | R8CA-ss-4 | 2016-05-09T00:00:00Z | 16 | NA | 1 | R8CA-ss-0 |
| mallard | Game Bird | spring | CA | Kern | 8 | shrub/scrub | R8CA-ss-4 | 2016-05-09T00:00:00Z | 16 | NA | 1 | R8CA-ss-0 |
| mallard | Game Bird | spring | CA | Kern | 8 | shrub/scrub | R8CA-ss-4 | 2016-05-09T00:00:00Z | 15 | NA | 1 | R8CA-ss-0 |
| mallard | Game Bird | winter | CA | Kern | 8 | shrub/scrub | R8CA-ss-4 | 2016-12-12T00:00:00Z | 10 | NA | 1 | R8CA-ss-0 |
| mallard | Game Bird | winter | CA | Kern | 8 | shrub/scrub | R8CA-ss-4 | 2016-12-12T00:00:00Z | 10 | NA | 1 | R8CA-ss-0 |
| mallard | Game Bird | winter | CA | Kern | 8 | shrub/scrub | R8CA-ss-4 | 2016-12-12T00:00:00Z | 1 | 1 | 1 | R8CA-ss-0 |
| mallard | Game Bird | winter | CA | Kern | 8 | shrub/scrub | R8CA-ss-4 | 2016-12-12T00:00:00Z | 4 | 10 | 1 | R8CA-ss-0 |
| mallard | Game Bird | summer | CA | Kern | 8 | shrub/scrub | R8CA-ss-4 | 2016-07-05T00:00:00Z | 16 | NA | 1 | R8CA-ss-0 |
| mallard | Game Bird | summer | CA | Kern | 8 | shrub/scrub | R8CA-ss-4 | 2016-07-05T00:00:00Z | 16 | NA | 1 | R8CA-ss-0 |
| mallard | Game Bird | summer | CA | Kern | 8 | shrub/scrub | R8CA-ss-4 | 2016-07-05T00:00:00Z | 16 | NA | 1 | R8CA-ss-0 |
| ring-necked pheasant | Game Bird | spring | CA | Imperial | 8 | shrub/scrub | R8CA-ss-9 | 2018-04-23T00:00:00Z | 30.9791667 | 42.0881944 | 1 | R8CA-ss-9 |
| ring-necked pheasant | Game Bird | spring | CA | Imperial | 8 | shrub/scrub | R8CA-ss-9 | 2018-04-23T00:00:00Z | 4.0270833 | 6.9458333 | 1 | R8CA-ss-9 |
| ring-necked pheasant | Game Bird | spring | CA | Imperial | 8 | shrub/scrub | R8CA-ss-9 | 2018-04-23T00:00:00Z | 18.0736111 | 23.8881944 | 1 | R8CA-ss-9 |
| ring-necked pheasant | Game Bird | spring | CA | Imperial | 8 | shrub/scrub | R8CA-ss-9 | 2018-04-23T00:00:00Z | 6.93125 | 10.0159722 | 1 | R8CA-ss-9 |
| mallard | Game Bird | spring | CA | Imperial | 8 | shrub/scrub | R8CA-ss-9 | 2018-04-23T00:00:00Z | 83 | 92.9618056 | 1 | R8CA-ss-9 |
| ring-necked pheasant | Game Bird | spring | CA | Imperial | 8 | shrub/scrub | R8CA-ss-9 | 2018-04-23T00:00:00Z | 6.9131944 | 9.9993056 | 1 | R8CA-ss-9 |
| ring-necked pheasant | Game Bird | spring | CA | Imperial | 8 | shrub/scrub | R8CA-ss-9 | 2018-04-23T00:00:00Z | 6.8993056 | 9.9840278 | 1 | R8CA-ss-9 |
| ring-necked pheasant | Game Bird | spring | CA | Imperial | 8 | shrub/scrub | R8CA-ss-9 | 2018-04-23T00:00:00Z | 3.98125 | 6.9006944 | 1 | R8CA-ss-9 |
| mallard | Game Bird | spring | CA | Imperial | 8 | shrub/scrub | R8CA-ss-9 | 2018-04-23T00:00:00Z | 62.9236111 | 72.8402778 | 1 | R8CA-ss-9 |
| ring-necked pheasant | Game Bird | spring | CA | Imperial | 8 | shrub/scrub | R8CA-ss-9 | 2018-04-23T00:00:00Z | 3.96875 | 6.8861111 | 1 | R8CA-ss-9 |
| ring-necked pheasant | Game Bird | spring | CA | Imperial | 8 | shrub/scrub | R8CA-ss-9 | 2018-04-23T00:00:00Z | 30.9791667 | 42.0881944 | 1 | R8CA-ss-9 |
| ring-necked pheasant | Game Bird | spring | CA | Imperial | 8 | shrub/scrub | R8CA-ss-9 | 2018-04-23T00:00:00Z | 4.0270833 | 6.9458333 | 1 | R8CA-ss-9 |
| ring-necked pheasant | Game Bird | spring | CA | Imperial | 8 | shrub/scrub | R8CA-ss-9 | 2018-04-23T00:00:00Z | 18.0736111 | 23.8881944 | 1 | R8CA-ss-9 |
| ring-necked pheasant | Game Bird | spring | CA | Imperial | 8 | shrub/scrub | R8CA-ss-9 | 2018-04-23T00:00:00Z | 6.93125 | 10.0159722 | 1 | R8CA-ss-9 |
| mallard | Game Bird | spring | CA | Imperial | 8 | shrub/scrub | R8CA-ss-9 | 2018-04-23T00:00:00Z | 51.9777778 | NA | 1 | R8CA-ss-9 |
| ring-necked pheasant | Game Bird | spring | CA | Imperial | 8 | shrub/scrub | R8CA-ss-9 | 2018-04-23T00:00:00Z | 6.9131944 | 9.9993056 | 1 | R8CA-ss-9 |
| ring-necked pheasant | Game Bird | spring | CA | Imperial | 8 | shrub/scrub | R8CA-ss-9 | 2018-04-23T00:00:00Z | 6.8993056 | 9.9840278 | 1 | R8CA-ss-9 |
| ring-necked pheasant | Game Bird | spring | CA | Imperial | 8 | shrub/scrub | R8CA-ss-9 | 2018-04-23T00:00:00Z | 3.98125 | 6.9006944 | 1 | R8CA-ss-9 |
| mallard | Game Bird | spring | CA | Imperial | 8 | shrub/scrub | R8CA-ss-9 | 2018-04-23T00:00:00Z | 51.9541667 | NA | 1 | R8CA-ss-9 |
| ring-necked pheasant | Game Bird | spring | CA | Imperial | 8 | shrub/scrub | R8CA-ss-9 | 2018-04-23T00:00:00Z | 3.96875 | 6.8861111 | 1 | R8CA-ss-9 |
| ring-necked pheasant | Game Bird | summer | CA | Imperial | 8 | shrub/scrub | R8CA-ss-9 | 2018-05-14T00:00:00Z | 50.0243056 | 60.0201389 | 1 | R8CA-ss-9 |
| ring-necked pheasant | Game Bird | summer | CA | Imperial | 8 | shrub/scrub | R8CA-ss-9 | 2018-05-14T00:00:00Z | 10.1104167 | 15.1055556 | 1 | R8CA-ss-9 |
| ring-necked pheasant | Game Bird | summer | CA | Imperial | 8 | shrub/scrub | R8CA-ss-9 | 2018-05-14T00:00:00Z | 50.0256944 | 60.0201389 | 1 | R8CA-ss-9 |
| ring-necked pheasant | Game Bird | summer | CA | Imperial | 8 | shrub/scrub | R8CA-ss-9 | 2018-05-14T00:00:00Z | 50.0201389 | 60.0236111 | 1 | R8CA-ss-9 |
| ring-necked pheasant | Game Bird | summer | CA | Imperial | 8 | shrub/scrub | R8CA-ss-9 | 2018-05-14T00:00:00Z | 50.0180556 | 60.0215278 | 1 | R8CA-ss-9 |
| ring-necked pheasant | Game Bird | summer | CA | Imperial | 8 | shrub/scrub | R8CA-ss-9 | 2018-05-14T00:00:00Z | 18.0583333 | 24.0881944 | 1 | R8CA-ss-9 |
| ring-necked pheasant | Game Bird | summer | CA | Imperial | 8 | shrub/scrub | R8CA-ss-9 | 2018-05-14T00:00:00Z | 60.0236111 | 70.0708333 | 1 | R8CA-ss-9 |
| ring-necked pheasant | Game Bird | summer | CA | Imperial | 8 | shrub/scrub | R8CA-ss-9 | 2018-05-14T00:00:00Z | 15.1118056 | 18.0604167 | 1 | R8CA-ss-9 |
| ring-necked pheasant | Game Bird | summer | CA | Imperial | 8 | shrub/scrub | R8CA-ss-9 | 2018-05-14T00:00:00Z | 2.9666667 | 4.01875 | 1 | R8CA-ss-9 |
| ring-necked pheasant | Game Bird | summer | CA | Imperial | 8 | shrub/scrub | R8CA-ss-9 | 2018-05-14T00:00:00Z | 2.9638889 | 4.0166667 | 1 | R8CA-ss-9 |
| ring-necked pheasant | Game Bird | summer | CA | Imperial | 8 | shrub/scrub | R8CA-ss-9 | 2018-08-06T00:00:00Z | 1.0020833 | 1.0020833 | 1 | R8CA-ss-9 |
| ring-necked pheasant | Game Bird | summer | CA | Imperial | 8 | shrub/scrub | R8CA-ss-9 | 2018-08-06T00:00:00Z | 0.9881944 | 0.9881944 | 1 | R8CA-ss-9 |
| ring-necked pheasant | Game Bird | summer | CA | Imperial | 8 | shrub/scrub | R8CA-ss-9 | 2018-08-06T00:00:00Z | 0.9895833 | 1.9354167 | 1 | R8CA-ss-9 |
| ring-necked pheasant | Game Bird | summer | CA | Imperial | 8 | shrub/scrub | R8CA-ss-9 | 2018-08-06T00:00:00Z | 2.8375 | 3.8770833 | 1 | R8CA-ss-9 |
| ring-necked pheasant | Game Bird | summer | CA | Imperial | 8 | shrub/scrub | R8CA-ss-9 | 2018-08-06T00:00:00Z | 1.0006944 | 1.9409722 | 1 | R8CA-ss-9 |
| ring-necked pheasant | Game Bird | summer | CA | Imperial | 8 | shrub/scrub | R8CA-ss-9 | 2018-08-06T00:00:00Z | 0.9993056 | 1.9402778 | 1 | R8CA-ss-9 |
| ring-necked pheasant | Game Bird | summer | CA | Imperial | 8 | shrub/scrub | R8CA-ss-9 | 2018-07-09T00:00:00Z | 98.8666667 | 109.0027778 | 1 | R8CA-ss-9 |
| ring-necked pheasant | Game Bird | summer | CA | Imperial | 8 | shrub/scrub | R8CA-ss-9 | 2018-07-09T00:00:00Z | 49.9826389 | 59.8041667 | 1 | R8CA-ss-9 |
| ring-necked pheasant | Game Bird | summer | CA | Imperial | 8 | shrub/scrub | R8CA-ss-9 | 2018-07-09T00:00:00Z | 49.9756944 | 59.8041667 | 1 | R8CA-ss-9 |
| ring-necked pheasant | Game Bird | summer | CA | Imperial | 8 | shrub/scrub | R8CA-ss-9 | 2018-07-09T00:00:00Z | 59.8090278 | 70.2291667 | 1 | R8CA-ss-9 |
| ring-necked pheasant | Game Bird | summer | CA | Imperial | 8 | shrub/scrub | R8CA-ss-9 | 2018-07-09T00:00:00Z | 30.0576389 | 39.8173611 | 1 | R8CA-ss-9 |
| ring-necked pheasant | Game Bird | summer | CA | Imperial | 8 | shrub/scrub | R8CA-ss-9 | 2018-07-09T00:00:00Z | 1.9597222 | 2.9472222 | 1 | R8CA-ss-9 |
| ring-necked pheasant | Game Bird | summer | CA | Imperial | 8 | shrub/scrub | R8CA-ss-9 | 2018-07-09T00:00:00Z | 2.95 | 4.1076389 | 1 | R8CA-ss-9 |
| turkey vulture | Raptor | fall | CA | Imperial | 8 | shrub/scrub | R8CA-ss-9 | 2018-09-17T00:00:00Z | 69.9902778 | 80.0430556 | 1 | R8CA-ss-9 |
| Cooper's hawk | Raptor | fall | CA | Imperial | 8 | shrub/scrub | R8CA-ss-9 | 2018-09-17T00:00:00Z | 38.9840278 | 50.05625 | 1 | R8CA-ss-9 |
| red-tailed hawk | Raptor | fall | CA | Imperial | 8 | shrub/scrub | R8CA-ss-9 | 2018-09-17T00:00:00Z | 80.0416667 | 92.0298611 | 1 | R8CA-ss-9 |
| red-tailed hawk | Raptor | fall | CA | Imperial | 8 | shrub/scrub | R8CA-ss-9 | 2018-09-17T00:00:00Z | 0.89375 | 0.89375 | 1 | R8CA-ss-9 |
| red-shouldered hawk | Raptor | fall | CA | Imperial | 8 | shrub/scrub | R8CA-ss-9 | 2018-09-17T00:00:00Z | 69.8659722 | 79.9458333 | 1 | R8CA-ss-9 |
| barn owl | Raptor | fall | CA | Imperial | 8 | shrub/scrub | R8CA-ss-9 | 2018-09-17T00:00:00Z | 79.9583333 | 91.9361111 | 1 | R8CA-ss-9 |
| red-shouldered hawk | Raptor | fall | CA | Imperial | 8 | shrub/scrub | R8CA-ss-9 | 2018-09-17T00:00:00Z | 69.8284722 | 79.9270833 | 1 | R8CA-ss-9 |
| barred owl | Raptor | winter | CA | Imperial | 8 | shrub/scrub | R8CA-ss-9 | 2019-01-07T00:00:00Z | 14.20625 | 21.2125 | 1 | R8CA-ss-9 |
| red-shouldered hawk | Raptor | winter | CA | Imperial | 8 | shrub/scrub | R8CA-ss-9 | 2019-01-07T00:00:00Z | 33.1222222 | NA | 1 | R8CA-ss-9 |
| Swainson's hawk | Raptor | winter | CA | Imperial | 8 | shrub/scrub | R8CA-ss-9 | 2019-01-07T00:00:00Z | 73.1541667 | NA | 1 | R8CA-ss-9 |
| red-shouldered hawk | Raptor | winter | CA | Imperial | 8 | shrub/scrub | R8CA-ss-9 | 2019-01-07T00:00:00Z | 73.1465278 | NA | 1 | R8CA-ss-9 |
| turkey vulture | Raptor | winter | CA | Imperial | 8 | shrub/scrub | R8CA-ss-9 | 2019-01-07T00:00:00Z | 8.01875 | 11.1076389 | 1 | R8CA-ss-9 |
| ring-necked pheasant | Game Bird | winter | CA | Imperial | 8 | shrub/scrub | R8CA-ss-9 | 2019-01-07T00:00:00Z | 7.9972222 | 11.0861111 | 1 | R8CA-ss-9 |
| ring-necked pheasant | Game Bird | winter | CA | Imperial | 8 | shrub/scrub | R8CA-ss-9 | 2019-01-07T00:00:00Z | 42.08125 | 52.1180556 | 1 | R8CA-ss-9 |
| ring-necked pheasant | Game Bird | winter | CA | Imperial | 8 | shrub/scrub | R8CA-ss-9 | 2019-01-07T00:00:00Z | 11.0638889 | 14.0729167 | 1 | R8CA-ss-9 |
| barn owl | Raptor | winter | CA | Imperial | 8 | shrub/scrub | R8CA-ss-9 | 2019-02-18T00:00:00Z | 31.1868056 | NA | 1 | R8CA-ss-9 |
| great horned owl | Raptor | winter | CA | Imperial | 8 | shrub/scrub | R8CA-ss-9 | 2019-02-18T00:00:00Z | 3.9840278 | 7.2222222 | 1 | R8CA-ss-9 |
| red-shouldered hawk | Raptor | winter | CA | Imperial | 8 | shrub/scrub | R8CA-ss-9 | 2019-02-18T00:00:00Z | 31.1451389 | NA | 1 | R8CA-ss-9 |
| barn owl | Raptor | winter | CA | Imperial | 8 | shrub/scrub | R8CA-ss-9 | 2019-02-18T00:00:00Z | 31.1152778 | NA | 1 | R8CA-ss-9 |
| red-shouldered hawk | Raptor | winter | CA | Imperial | 8 | shrub/scrub | R8CA-ss-9 | 2019-02-18T00:00:00Z | 31.0972222 | NA | 1 | R8CA-ss-9 |
| barn owl | Raptor | winter | CA | Imperial | 8 | shrub/scrub | R8CA-ss-9 | 2019-02-18T00:00:00Z | 31.0722222 | NA | 1 | R8CA-ss-9 |
| red-shouldered hawk | Raptor | winter | CA | Imperial | 8 | shrub/scrub | R8CA-ss-9 | 2019-02-18T00:00:00Z | 31.1041667 | NA | 1 | R8CA-ss-9 |
| red-tailed hawk | Raptor | winter | CA | Imperial | 8 | shrub/scrub | R8CA-ss-9 | 2019-02-18T00:00:00Z | 31.0791667 | NA | 1 | R8CA-ss-9 |
| Cooper's hawk | Raptor | winter | CA | Imperial | 8 | shrub/scrub | R8CA-ss-9 | 2019-02-18T00:00:00Z | 13.9118056 | 28.0618056 | 1 | R8CA-ss-9 |
| red-tailed hawk | Raptor | winter | CA | Imperial | 8 | shrub/scrub | R8CA-ss-9 | 2019-02-18T00:00:00Z | 31.0506944 | NA | 1 | R8CA-ss-9 |
| mallard | Game Bird | summer | NV | White Pine | 8 | shrub/scrub | R8NV-ss-7 | 2017-08-15T00:00:00Z | 41.13125 | NA | 1 | R8NV-ss-7 |
| ring-necked pheasant | Game Bird | fall | NV | White Pine | 8 | shrub/scrub | R8NV-ss-7 | 2017-10-09T00:00:00Z | 2.1465278 | 3.2111111 | 1 | R8NV-ss-7 |
| mallard | Game Bird | fall | NV | White Pine | 8 | shrub/scrub | R8NV-ss-7 | 2017-10-09T00:00:00Z | 14.0409722 | 39.0270833 | 1 | R8NV-ss-7 |
| mallard | Game Bird | summer | NV | White Pine | 8 | shrub/scrub | R8NV-ss-7 | 2017-08-15T00:00:00Z | 41.0138889 | NA | 1 | R8NV-ss-7 |
| mallard | Game Bird | winter | NV | White Pine | 8 | shrub/scrub | R8NV-ss-7 | 2017-12-11T00:00:00Z | 7.0409722 | 17.0493056 | 1 | R8NV-ss-7 |
| ring-necked pheasant | Game Bird | winter | NV | White Pine | 8 | shrub/scrub | R8NV-ss-7 | 2017-12-11T00:00:00Z | 1.0881944 | 1.0881944 | 1 | R8NV-ss-7 |
| barn owl | Raptor | spring | NV | White Pine | 8 | shrub/scrub | R8NV-ss-7 | 2017-04-25T00:00:00Z | 120.1826389 | NA | 1 | R8NV-ss-7 |
| red-tailed hawk | Raptor | spring | NV | White Pine | 8 | shrub/scrub | R8NV-ss-7 | 2017-04-25T00:00:00Z | 120.1777778 | NA | 1 | R8NV-ss-7 |
| red-tailed hawk | Raptor | spring | NV | White Pine | 8 | shrub/scrub | R8NV-ss-7 | 2017-04-25T00:00:00Z | 120.1368056 | NA | 1 | R8NV-ss-7 |
| barred owl | Raptor | spring | NV | White Pine | 8 | shrub/scrub | R8NV-ss-7 | 2017-04-25T00:00:00Z | 120.1215278 | NA | 1 | R8NV-ss-7 |
| red-tailed hawk | Raptor | spring | NV | White Pine | 8 | shrub/scrub | R8NV-ss-7 | 2017-04-25T00:00:00Z | 120.1111111 | NA | 1 | R8NV-ss-7 |
| barn owl | Raptor | spring | NV | White Pine | 8 | shrub/scrub | R8NV-ss-7 | 2017-04-25T00:00:00Z | 71.0673611 | 79.89375 | 1 | R8NV-ss-7 |
| red-tailed hawk | Raptor | spring | NV | White Pine | 8 | shrub/scrub | R8NV-ss-7 | 2017-04-25T00:00:00Z | 120.06875 | NA | 1 | R8NV-ss-7 |
| barn owl | Raptor | spring | NV | White Pine | 8 | shrub/scrub | R8NV-ss-7 | 2017-04-25T00:00:00Z | 120.0534722 | NA | 1 | R8NV-ss-7 |
| Swainson's hawk | Raptor | spring | NV | White Pine | 8 | shrub/scrub | R8NV-ss-7 | 2017-04-25T00:00:00Z | 120.0388889 | NA | 1 | R8NV-ss-7 |
| great horned owl | Raptor | spring | NV | White Pine | 8 | shrub/scrub | R8NV-ss-7 | 2017-04-25T00:00:00Z | 119.8854167 | NA | 1 | R8NV-ss-7 |
| great horned owl | Raptor | spring | NV | White Pine | 8 | shrub/scrub | R8NV-ss-7 | 2017-04-25T00:00:00Z | 120.0083333 | NA | 1 | R8NV-ss-7 |
| peregrine falcon | Raptor | spring | NV | White Pine | 8 | shrub/scrub | R8NV-ss-7 | 2017-04-25T00:00:00Z | 119.9 | NA | 1 | R8NV-ss-7 |
| mallard | Game Bird | spring | NV | White Pine | 8 | shrub/scrub | R8NV-ss-7 | 2017-04-17T00:00:00Z | 39.1361111 | NA | 1 | R8NV-ss-7 |
| mallard | Game Bird | winter | NV | White Pine | 8 | shrub/scrub | R8NV-ss-7 | 2017-02-28T00:00:00Z | 41 | NA | 1 | R8NV-ss-7 |
| ring-necked pheasant | Game Bird | winter | NV | White Pine | 8 | shrub/scrub | R8NV-ss-7 | 2017-02-28T00:00:00Z | 3 | 7 | 1 | R8NV-ss-7 |
| mallard | Game Bird | winter | NV | White Pine | 8 | shrub/scrub | R8NV-ss-7 | 2017-02-28T00:00:00Z | 14 | NA | 1 | R8NV-ss-7 |
| mallard | Game Bird | spring | NV | White Pine | 8 | shrub/scrub | R8NV-ss-7 | 2017-04-03T00:00:00Z | 39 | NA | 1 | R8NV-ss-7 |
| ring-necked pheasant | Game Bird | spring | NV | White Pine | 8 | shrub/scrub | R8NV-ss-7 | 2017-04-03T00:00:00Z | 1 | 2 | 1 | R8NV-ss-7 |
| mallard | Game Bird | spring | NV | White Pine | 8 | shrub/scrub | R8NV-ss-7 | 2017-04-03T00:00:00Z | 10 | 14 | 1 | R8NV-ss-7 |
| mallard | Game Bird | summer | NV | White Pine | 8 | shrub/scrub | R8NV-ss-7 | 2017-06-12T00:00:00Z | 39.0618056 | NA | 1 | R8NV-ss-7 |
| ring-necked pheasant | Game Bird | summer | NV | White Pine | 8 | shrub/scrub | R8NV-ss-7 | 2017-06-12T00:00:00Z | 14.1069444 | 38.9958333 | 1 | R8NV-ss-7 |
| ring-necked pheasant | Game Bird | summer | NV | White Pine | 8 | shrub/scrub | R8NV-ss-7 | 2017-07-24T00:00:00Z | 41.9388889 | NA | 1 | R8NV-ss-7 |
| mallard | Game Bird | summer | NV | White Pine | 8 | shrub/scrub | R8NV-ss-7 | 2017-07-24T00:00:00Z | 14.0513889 | 38.9173611 | 1 | R8NV-ss-7 |
| red-tailed hawk | Raptor | summer | NV | White Pine | 8 | shrub/scrub | R8NV-ss-7 | 2017-09-11T00:00:00Z | 108.2131944 | NA | 1 | R8NV-ss-7 |
| red-tailed hawk | Raptor | summer | NV | White Pine | 8 | shrub/scrub | R8NV-ss-7 | 2017-09-11T00:00:00Z | 108.2125 | NA | 1 | R8NV-ss-7 |
| red-tailed hawk | Raptor | summer | NV | White Pine | 8 | shrub/scrub | R8NV-ss-7 | 2017-09-11T00:00:00Z | 108.2125 | NA | 1 | R8NV-ss-7 |
| red-tailed hawk | Raptor | summer | NV | White Pine | 8 | shrub/scrub | R8NV-ss-7 | 2017-09-11T00:00:00Z | 14.1152778 | 31.1166667 | 1 | R8NV-ss-7 |
| red-tailed hawk | Raptor | summer | NV | White Pine | 8 | shrub/scrub | R8NV-ss-7 | 2017-09-11T00:00:00Z | 108.2013889 | NA | 1 | R8NV-ss-7 |
| red-tailed hawk | Raptor | summer | NV | White Pine | 8 | shrub/scrub | R8NV-ss-7 | 2017-09-11T00:00:00Z | 108.1986111 | NA | 1 | R8NV-ss-7 |
| red-tailed hawk | Raptor | summer | NV | White Pine | 8 | shrub/scrub | R8NV-ss-7 | 2017-09-11T00:00:00Z | 108.125 | NA | 1 | R8NV-ss-7 |
| red-tailed hawk | Raptor | summer | NV | White Pine | 8 | shrub/scrub | R8NV-ss-7 | 2017-09-11T00:00:00Z | 109.0006944 | NA | 1 | R8NV-ss-7 |
| red-tailed hawk | Raptor | summer | NV | White Pine | 8 | shrub/scrub | R8NV-ss-7 | 2017-09-11T00:00:00Z | 109.0944444 | NA | 1 | R8NV-ss-7 |
| red-tailed hawk | Raptor | summer | NV | White Pine | 8 | shrub/scrub | R8NV-ss-7 | 2017-09-11T00:00:00Z | 108.10625 | NA | 1 | R8NV-ss-7 |
| red-tailed hawk | Raptor | summer | NV | White Pine | 8 | shrub/scrub | R8NV-ss-7 | 2017-09-11T00:00:00Z | 108.1104167 | NA | 1 | R8NV-ss-7 |
| red-tailed hawk | Raptor | summer | NV | White Pine | 8 | shrub/scrub | R8NV-ss-7 | 2017-09-11T00:00:00Z | 52.0756944 | 62.9486111 | 1 | R8NV-ss-7 |
| red-tailed hawk | Raptor | summer | NV | White Pine | 8 | shrub/scrub | R8NV-ss-7 | 2017-09-11T00:00:00Z | 107.9229167 | NA | 1 | R8NV-ss-7 |
| red-tailed hawk | Raptor | summer | NV | White Pine | 8 | shrub/scrub | R8NV-ss-7 | 2017-09-11T00:00:00Z | 52.0680556 | 62.9479167 | 1 | R8NV-ss-7 |
| red-tailed hawk | Raptor | summer | NV | White Pine | 8 | shrub/scrub | R8NV-ss-7 | 2017-09-11T00:00:00Z | 107.9152778 | NA | 1 | R8NV-ss-7 |
| ring-necked pheasant | Game Bird | summer | NV | White PIne | 8 | shrub/scrub | R8NV-ss-8 | 2019-06-18T00:00:00Z | 41.0895833 | 55.9173611 | 1 | R8NV-ss-8 |
| ring-necked pheasant | Game Bird | summer | NV | White PIne | 8 | shrub/scrub | R8NV-ss-8 | 2019-08-19T00:00:00Z | 1.9715278 | 2.9729167 | 1 | R8NV-ss-8 |
| ring-necked pheasant | Game Bird | summer | NV | White PIne | 8 | shrub/scrub | R8NV-ss-8 | 2019-06-18T00:00:00Z | 20.1354167 | 27.0798611 | 1 | R8NV-ss-8 |
| ring-necked pheasant | Game Bird | summer | NV | White PIne | 8 | shrub/scrub | R8NV-ss-8 | 2019-06-18T00:00:00Z | 34.1652778 | 41.1298611 | 1 | R8NV-ss-8 |
| ring-necked pheasant | Game Bird | summer | NV | White PIne | 8 | shrub/scrub | R8NV-ss-8 | 2019-06-18T00:00:00Z | 15.8854167 | 20.1104167 | 1 | R8NV-ss-8 |
| ring-necked pheasant | Game Bird | summer | NV | White PIne | 8 | shrub/scrub | R8NV-ss-8 | 2019-06-18T00:00:00Z | 7.1645833 | 15.8409722 | 1 | R8NV-ss-8 |
| ring-necked pheasant | Game Bird | summer | NV | White PIne | 8 | shrub/scrub | R8NV-ss-8 | 2019-06-18T00:00:00Z | 2.8236111 | 37.1597222 | 1 | R8NV-ss-8 |
| ring-necked pheasant | Game Bird | summer | NV | White PIne | 8 | shrub/scrub | R8NV-ss-8 | 2019-06-18T00:00:00Z | 27.0152778 | 34.10625 | 1 | R8NV-ss-8 |
| ring-necked pheasant | Game Bird | summer | NV | White PIne | 8 | shrub/scrub | R8NV-ss-8 | 2019-06-18T00:00:00Z | 14.0993056 | 20.0555556 | 1 | R8NV-ss-8 |
| ring-necked pheasant | Game Bird | summer | NV | White PIne | 8 | shrub/scrub | R8NV-ss-8 | 2019-06-18T00:00:00Z | 7.1472222 | 14.1069444 | 1 | R8NV-ss-8 |
| ring-necked pheasant | Game Bird | summer | NV | White PIne | 8 | shrub/scrub | R8NV-ss-8 | 2019-06-18T00:00:00Z | 20.0354167 | 26.9875 | 1 | R8NV-ss-8 |
| ring-necked pheasant | Game Bird | summer | NV | White PIne | 8 | shrub/scrub | R8NV-ss-8 | 2019-08-19T00:00:00Z | 3.06875 | 70.0631944 | 1 | R8NV-ss-8 |
| ring-necked pheasant | Game Bird | summer | NV | White PIne | 8 | shrub/scrub | R8NV-ss-8 | 2019-08-19T00:00:00Z | 3.0569444 | 9.0583333 | 1 | R8NV-ss-8 |
| ring-necked pheasant | Game Bird | summer | NV | White PIne | 8 | shrub/scrub | R8NV-ss-8 | 2019-08-19T00:00:00Z | 17.9354167 | NA | 1 | R8NV-ss-8 |
| ring-necked pheasant | Game Bird | summer | NV | White PIne | 8 | shrub/scrub | R8NV-ss-8 | 2019-08-19T00:00:00Z | 18.0097222 | 24.1020833 | 1 | R8NV-ss-8 |
| ring-necked pheasant | Game Bird | fall | NV | White PIne | 8 | shrub/scrub | R8NV-ss-8 | 2019-09-10T00:00:00Z | 7.1951389 | 14.1631944 | 1 | R8NV-ss-8 |
| ring-necked pheasant | Game Bird | fall | NV | White PIne | 8 | shrub/scrub | R8NV-ss-8 | 2019-09-10T00:00:00Z | 7.2013889 | 14.1076389 | 1 | R8NV-ss-8 |
| ring-necked pheasant | Game Bird | fall | NV | White PIne | 8 | shrub/scrub | R8NV-ss-8 | 2019-09-10T00:00:00Z | 22.9340278 | 29.0729167 | 1 | R8NV-ss-8 |
| ring-necked pheasant | Game Bird | fall | NV | White PIne | 8 | shrub/scrub | R8NV-ss-8 | 2019-09-10T00:00:00Z | 29.0986111 | 41.0569444 | 1 | R8NV-ss-8 |
| ring-necked pheasant | Game Bird | fall | NV | White PIne | 8 | shrub/scrub | R8NV-ss-8 | 2019-09-10T00:00:00Z | 41.0048611 | NA | 1 | R8NV-ss-8 |
| barn owl | Raptor | fall | NV | White PIne | 8 | shrub/scrub | R8NV-ss-8 | 2019-10-21T00:00:00Z | 99.9465278 | NA | 1 | R8NV-ss-8 |
| Cooper's hawk | Raptor | fall | NV | White PIne | 8 | shrub/scrub | R8NV-ss-8 | 2019-10-21T00:00:00Z | 56.7479167 | 99.7979167 | 1 | R8NV-ss-8 |
| Cooper's hawk | Raptor | fall | NV | White PIne | 8 | shrub/scrub | R8NV-ss-8 | 2019-10-21T00:00:00Z | 99.8034722 | NA | 1 | R8NV-ss-8 |
| Cooper's hawk | Raptor | fall | NV | White PIne | 8 | shrub/scrub | R8NV-ss-8 | 2019-10-21T00:00:00Z | 27.8506944 | 56.9736111 | 1 | R8NV-ss-8 |
| barn owl | Raptor | fall | NV | White PIne | 8 | shrub/scrub | R8NV-ss-8 | 2019-10-21T00:00:00Z | 27.84375 | 57.6361111 | 1 | R8NV-ss-8 |
| Cooper's hawk | Raptor | fall | NV | White PIne | 8 | shrub/scrub | R8NV-ss-8 | 2019-10-21T00:00:00Z | 27.8326389 | 57.8159722 | 1 | R8NV-ss-8 |
| barn owl | Raptor | fall | NV | White PIne | 8 | shrub/scrub | R8NV-ss-8 | 2019-10-21T00:00:00Z | 99.7895833 | NA | 1 | R8NV-ss-8 |
| northern harrier | Raptor | fall | NV | White PIne | 8 | shrub/scrub | R8NV-ss-8 | 2019-10-21T00:00:00Z | 99.7791667 | NA | 1 | R8NV-ss-8 |
| Cooper's hawk | Raptor | fall | NV | White PIne | 8 | shrub/scrub | R8NV-ss-8 | 2019-10-21T00:00:00Z | 99.7881944 | NA | 1 | R8NV-ss-8 |
| Cooper's hawk | Raptor | fall | NV | White PIne | 8 | shrub/scrub | R8NV-ss-8 | 2019-10-21T00:00:00Z | 99.7840278 | NA | 1 | R8NV-ss-8 |
| Swainson's hawk | Raptor | winter | NV | White PIne | 8 | shrub/scrub | R8NV-ss-8 | 2020-02-11T00:00:00Z | 71.8895833 | NA | 1 | R8NV-ss-8 |
| great horned owl | Raptor | winter | NV | White PIne | 8 | shrub/scrub | R8NV-ss-8 | 2020-02-11T00:00:00Z | 13.9881944 | 43.9805556 | 1 | R8NV-ss-8 |
| great horned owl | Raptor | winter | NV | White PIne | 8 | shrub/scrub | R8NV-ss-8 | 2020-02-11T00:00:00Z | 3.7909722 | 13.975 | 1 | R8NV-ss-8 |
| red-tailed hawk | Raptor | winter | NV | White PIne | 8 | shrub/scrub | R8NV-ss-8 | 2020-02-11T00:00:00Z | 13.9340278 | 44.0111111 | 1 | R8NV-ss-8 |
| great horned owl | Raptor | winter | NV | White PIne | 8 | shrub/scrub | R8NV-ss-8 | 2020-02-11T00:00:00Z | 13.9805556 | 43.9784722 | 1 | R8NV-ss-8 |
| red-tailed hawk | Raptor | winter | NV | White PIne | 8 | shrub/scrub | R8NV-ss-8 | 2020-02-11T00:00:00Z | 13.9090278 | 43.9909722 | 1 | R8NV-ss-8 |
| great horned owl | Raptor | winter | NV | White PIne | 8 | shrub/scrub | R8NV-ss-8 | 2020-02-11T00:00:00Z | 43.9881944 | 70.9986111 | 1 | R8NV-ss-8 |
| Swainson's hawk | Raptor | winter | NV | White PIne | 8 | shrub/scrub | R8NV-ss-8 | 2020-02-11T00:00:00Z | 70.7152778 | NA | 1 | R8NV-ss-8 |
| red-tailed hawk | Raptor | winter | NV | White PIne | 8 | shrub/scrub | R8NV-ss-8 | 2020-02-11T00:00:00Z | 43.9576389 | 69.0944444 | 1 | R8NV-ss-8 |
| great horned owl | Raptor | winter | NV | White PIne | 8 | shrub/scrub | R8NV-ss-8 | 2020-02-11T00:00:00Z | 43.9486111 | 69.0534722 | 1 | R8NV-ss-8 |
| northern harrier | Raptor | winter | WY | Converse | 6 | shrub/scrub | R6WY-ss-9 | 2021-01-06T00:00:00Z | 1.9631944 | 5.0340278 | 0 | R6WY-ss-9 |
| great horned owl | Raptor | winter | WY | Converse | 6 | shrub/scrub | R6WY-ss-9 | 2021-01-06T00:00:00Z | 1.95625 | 5.0326389 | 0 | R6WY-ss-9 |
| great horned owl | Raptor | winter | WY | Converse | 6 | shrub/scrub | R6WY-ss-9 | 2021-01-06T00:00:00Z | 11.9965278 | 16.0305556 | 0 | R6WY-ss-9 |
| red-tailed hawk | Raptor | winter | WY | Converse | 6 | shrub/scrub | R6WY-ss-9 | 2021-01-06T00:00:00Z | 125.9298611 | NA | 0 | R6WY-ss-9 |
| great horned owl | Raptor | winter | WY | Converse | 6 | shrub/scrub | R6WY-ss-9 | 2021-01-06T00:00:00Z | 75.025 | 84.9236111 | 0 | R6WY-ss-9 |
| red-tailed hawk | Raptor | winter | WY | Converse | 6 | shrub/scrub | R6WY-ss-9 | 2021-01-06T00:00:00Z | 63.8590278 | 75 | 0 | R6WY-ss-9 |
| red-tailed hawk | Raptor | winter | WY | Converse | 6 | shrub/scrub | R6WY-ss-9 | 2021-01-06T00:00:00Z | 126.8465278 | NA | 0 | R6WY-ss-9 |
| red-tailed hawk | Raptor | winter | WY | Converse | 6 | shrub/scrub | R6WY-ss-9 | 2021-01-06T00:00:00Z | 126.8479167 | NA | 0 | R6WY-ss-9 |
| red-tailed hawk | Raptor | winter | WY | Converse | 6 | shrub/scrub | R6WY-ss-9 | 2021-01-06T00:00:00Z | 126.8402778 | NA | 0 | R6WY-ss-9 |
| red-tailed hawk | Raptor | winter | WY | Converse | 6 | shrub/scrub | R6WY-ss-9 | 2021-01-06T00:00:00Z | 74.9673611 | 84.8618056 | 0 | R6WY-ss-9 |
| red-tailed hawk | Raptor | spring | WY | Converse | 6 | shrub/scrub | R6WY-ss-9 | 2020-05-04T00:00:00Z | 123.0347222 | NA | 0 | R6WY-ss-9 |
| great horned owl | Raptor | spring | WY | Converse | 6 | shrub/scrub | R6WY-ss-9 | 2020-05-04T00:00:00Z | 113.1638889 | NA | 0 | R6WY-ss-9 |
| red-tailed hawk | Raptor | spring | WY | Converse | 6 | shrub/scrub | R6WY-ss-9 | 2020-05-04T00:00:00Z | 123.0298611 | NA | 0 | R6WY-ss-9 |
| red-tailed hawk | Raptor | spring | WY | Converse | 6 | shrub/scrub | R6WY-ss-9 | 2020-05-04T00:00:00Z | 102.9548611 | 113.1486111 | 0 | R6WY-ss-9 |
| Cooper's hawk | Raptor | spring | WY | Converse | 6 | shrub/scrub | R6WY-ss-9 | 2020-05-04T00:00:00Z | 102.9465278 | 113.1409722 | 0 | R6WY-ss-9 |
| red-tailed hawk | Raptor | spring | WY | Converse | 6 | shrub/scrub | R6WY-ss-9 | 2020-05-04T00:00:00Z | 102.9444444 | 113.1368056 | 0 | R6WY-ss-9 |
| red-tailed hawk | Raptor | spring | WY | Converse | 6 | shrub/scrub | R6WY-ss-9 | 2020-05-04T00:00:00Z | 73.1680556 | 83.1409722 | 0 | R6WY-ss-9 |
| red-tailed hawk | Raptor | spring | WY | Converse | 6 | shrub/scrub | R6WY-ss-9 | 2020-05-04T00:00:00Z | 123.0118056 | NA | 0 | R6WY-ss-9 |
| barn owl | Raptor | spring | WY | Converse | 6 | shrub/scrub | R6WY-ss-9 | 2020-05-04T00:00:00Z | 83.1319444 | 93.1208333 | 0 | R6WY-ss-9 |
| red-tailed hawk | Raptor | summer | WY | Converse | 6 | shrub/scrub | R6WY-ss-9 | 2020-07-13T00:00:00Z | 126.1131944 | NA | 0 | R6WY-ss-9 |
| red-tailed hawk | Raptor | spring | WY | Converse | 6 | shrub/scrub | R6WY-ss-9 | 2020-05-04T00:00:00Z | 73.1923611 | 83.1270833 | 0 | R6WY-ss-9 |
| red-tailed hawk | Raptor | summer | WY | Converse | 6 | shrub/scrub | R6WY-ss-9 | 2020-07-13T00:00:00Z | 126.1125 | NA | 0 | R6WY-ss-9 |
| turkey vulture | Raptor | summer | WY | Converse | 6 | shrub/scrub | R6WY-ss-9 | 2020-07-13T00:00:00Z | 126.1097222 | NA | 0 | R6WY-ss-9 |
| barn owl | Raptor | summer | WY | Converse | 6 | shrub/scrub | R6WY-ss-9 | 2020-07-13T00:00:00Z | 53.0979167 | 63.1868056 | 0 | R6WY-ss-9 |
| turkey vulture | Raptor | summer | WY | Converse | 6 | shrub/scrub | R6WY-ss-9 | 2020-07-13T00:00:00Z | 126.1 | NA | 0 | R6WY-ss-9 |
| great horned owl | Raptor | summer | WY | Converse | 6 | shrub/scrub | R6WY-ss-9 | 2020-07-13T00:00:00Z | 53.0798611 | 63.1701389 | 0 | R6WY-ss-9 |
| red-tailed hawk | Raptor | summer | WY | Converse | 6 | shrub/scrub | R6WY-ss-9 | 2020-07-13T00:00:00Z | 53.0729167 | 63.15625 | 0 | R6WY-ss-9 |
| turkey vulture | Raptor | summer | WY | Converse | 6 | shrub/scrub | R6WY-ss-9 | 2020-07-13T00:00:00Z | 63.1597222 | 73.0770833 | 0 | R6WY-ss-9 |
| turkey vulture | Raptor | summer | WY | Converse | 6 | shrub/scrub | R6WY-ss-9 | 2020-07-13T00:00:00Z | 126.0666667 | NA | 0 | R6WY-ss-9 |
| Swainson's hawk | Raptor | fall | WY | Converse | 6 | shrub/scrub | R6WY-ss-9 | 2020-10-15T00:00:00Z | 22.0326389 | 31.9888889 | 0 | R6WY-ss-9 |
| Swainson's hawk | Raptor | fall | WY | Converse | 6 | shrub/scrub | R6WY-ss-9 | 2020-10-15T00:00:00Z | 31.9875 | 45.9013889 | 0 | R6WY-ss-9 |
| Swainson's hawk | Raptor | fall | WY | Converse | 6 | shrub/scrub | R6WY-ss-9 | 2020-10-15T00:00:00Z | 4.0076389 | 15.0576389 | 0 | R6WY-ss-9 |
| Swainson's hawk | Raptor | fall | WY | Converse | 6 | shrub/scrub | R6WY-ss-9 | 2020-10-15T00:00:00Z | 2.0166667 | 4.0027778 | 0 | R6WY-ss-9 |
| Swainson's hawk | Raptor | fall | WY | Converse | 6 | shrub/scrub | R6WY-ss-9 | 2020-10-15T00:00:00Z | 2.0159722 | 3.9986111 | 0 | R6WY-ss-9 |
| Swainson's hawk | Raptor | fall | WY | Converse | 6 | shrub/scrub | R6WY-ss-9 | 2020-10-15T00:00:00Z | 2.0145833 | 3.9965278 | 0 | R6WY-ss-9 |
| Swainson's hawk | Raptor | fall | WY | Converse | 6 | shrub/scrub | R6WY-ss-9 | 2020-09-14T00:00:00Z | 13.8715278 | 20.9826389 | 0 | R6WY-ss-9 |
| Swainson's hawk | Raptor | fall | WY | Converse | 6 | shrub/scrub | R6WY-ss-9 | 2020-09-14T00:00:00Z | 20.9986111 | 30.8333333 | 0 | R6WY-ss-9 |
| red-tailed hawk | Raptor | fall | WY | Converse | 6 | shrub/scrub | R6WY-ss-9 | 2020-09-14T00:00:00Z | 119.1006944 | 128.8805556 | 0 | R6WY-ss-9 |
| ferruginous hawk | Raptor | fall | WY | Converse | 6 | shrub/scrub | R6WY-ss-9 | 2020-09-14T00:00:00Z | 128.9006944 | NA | 0 | R6WY-ss-9 |
| turkey vulture | Raptor | spring | WY | Converse | 6 | shrub/scrub | R6WY-ss-8 | 2019-04-15T00:00:00Z | 137.0104167 | NA | 0 | R6WY-ss-8 |
| red-tailed hawk | Raptor | spring | WY | Converse | 6 | shrub/scrub | R6WY-ss-8 | 2019-04-15T00:00:00Z | 136.9979167 | NA | 0 | R6WY-ss-8 |
| red-tailed hawk | Raptor | spring | WY | Converse | 6 | shrub/scrub | R6WY-ss-8 | 2019-04-15T00:00:00Z | 136.9833333 | NA | 0 | R6WY-ss-8 |
| red-tailed hawk | Raptor | spring | WY | Converse | 6 | shrub/scrub | R6WY-ss-8 | 2019-04-15T00:00:00Z | 136.975 | NA | 0 | R6WY-ss-8 |
| red-tailed hawk | Raptor | spring | WY | Converse | 6 | shrub/scrub | R6WY-ss-8 | 2019-04-15T00:00:00Z | 137.0215278 | NA | 0 | R6WY-ss-8 |
| red-tailed hawk | Raptor | spring | WY | Converse | 6 | shrub/scrub | R6WY-ss-8 | 2019-04-24T00:00:00Z | 128.0354167 | NA | 0 | R6WY-ss-8 |
| red-tailed hawk | Raptor | spring | WY | Converse | 6 | shrub/scrub | R6WY-ss-8 | 2019-04-24T00:00:00Z | 128.0541667 | NA | 0 | R6WY-ss-8 |
| red-tailed hawk | Raptor | spring | WY | Converse | 6 | shrub/scrub | R6WY-ss-8 | 2019-04-24T00:00:00Z | 128.0590278 | NA | 0 | R6WY-ss-8 |
| red-tailed hawk | Raptor | spring | WY | Converse | 6 | shrub/scrub | R6WY-ss-8 | 2019-04-24T00:00:00Z | 128.0333333 | NA | 0 | R6WY-ss-8 |
| red-tailed hawk | Raptor | spring | WY | Converse | 6 | shrub/scrub | R6WY-ss-8 | 2019-04-24T00:00:00Z | 128.0444444 | NA | 0 | R6WY-ss-8 |
| red-tailed hawk | Raptor | summer | WY | Converse | 6 | shrub/scrub | R6WY-ss-8 | 2019-06-06T00:00:00Z | 124.1263889 | NA | 0 | R6WY-ss-8 |
| red-tailed hawk | Raptor | summer | WY | Converse | 6 | shrub/scrub | R6WY-ss-8 | 2019-06-06T00:00:00Z | 124.1131944 | NA | 0 | R6WY-ss-8 |
| red-tailed hawk | Raptor | summer | WY | Converse | 6 | shrub/scrub | R6WY-ss-8 | 2019-06-06T00:00:00Z | 123.8777778 | NA | 0 | R6WY-ss-8 |
| Swainson's hawk | Raptor | summer | WY | Converse | 6 | shrub/scrub | R6WY-ss-8 | 2019-06-06T00:00:00Z | 123.9340278 | NA | 0 | R6WY-ss-8 |
| red-tailed hawk | Raptor | summer | WY | Converse | 6 | shrub/scrub | R6WY-ss-8 | 2019-06-06T00:00:00Z | 124.0256944 | NA | 0 | R6WY-ss-8 |
| northern goshawk | Raptor | summer | WY | Converse | 6 | shrub/scrub | R6WY-ss-8 | 2019-06-24T00:00:00Z | 136.0902778 | NA | 0 | R6WY-ss-8 |
| northern harrier | Raptor | summer | WY | Converse | 6 | shrub/scrub | R6WY-ss-8 | 2019-06-24T00:00:00Z | 136.14375 | NA | 0 | R6WY-ss-8 |
| northern harrier | Raptor | summer | WY | Converse | 6 | shrub/scrub | R6WY-ss-8 | 2019-06-24T00:00:00Z | 136.1354167 | NA | 0 | R6WY-ss-8 |
| northern harrier | Raptor | summer | WY | Converse | 6 | shrub/scrub | R6WY-ss-8 | 2019-06-24T00:00:00Z | 136.1277778 | NA | 0 | R6WY-ss-8 |
| northern harrier | Raptor | summer | WY | Converse | 6 | shrub/scrub | R6WY-ss-8 | 2019-06-24T00:00:00Z | 136.1215278 | NA | 0 | R6WY-ss-8 |
| red-tailed hawk | Raptor | fall | WY | Converse | 6 | shrub/scrub | R6WY-ss-8 | 2019-10-23T00:00:00Z | 122.9444444 | NA | 0 | R6WY-ss-8 |
| red-tailed hawk | Raptor | fall | WY | Converse | 6 | shrub/scrub | R6WY-ss-8 | 2019-10-23T00:00:00Z | 70.9604167 | 79.0861111 | 0 | R6WY-ss-8 |
| red-tailed hawk | Raptor | fall | WY | Converse | 6 | shrub/scrub | R6WY-ss-8 | 2019-10-23T00:00:00Z | 59.9215278 | 70.9006944 | 0 | R6WY-ss-8 |
| great horned owl | Raptor | fall | WY | Converse | 6 | shrub/scrub | R6WY-ss-8 | 2019-10-23T00:00:00Z | 1.9506944 | 14.0680556 | 0 | R6WY-ss-8 |
| red-tailed hawk | Raptor | fall | WY | Converse | 6 | shrub/scrub | R6WY-ss-8 | 2019-10-23T00:00:00Z | 91.05625 | 102.9868056 | 0 | R6WY-ss-8 |
| red-tailed hawk | Raptor | fall | WY | Converse | 6 | shrub/scrub | R6WY-ss-8 | 2019-11-11T00:00:00Z | 122 | NA | 0 | R6WY-ss-8 |
| red-tailed hawk | Raptor | fall | WY | Converse | 6 | shrub/scrub | R6WY-ss-8 | 2019-11-11T00:00:00Z | 121.99375 | NA | 0 | R6WY-ss-8 |
| red-tailed hawk | Raptor | fall | WY | Converse | 6 | shrub/scrub | R6WY-ss-8 | 2019-11-11T00:00:00Z | 121.9604167 | NA | 0 | R6WY-ss-8 |
| red-tailed hawk | Raptor | fall | WY | Converse | 6 | shrub/scrub | R6WY-ss-8 | 2019-11-11T00:00:00Z | 121.9409722 | NA | 0 | R6WY-ss-8 |
| red-tailed hawk | Raptor | fall | WY | Converse | 6 | shrub/scrub | R6WY-ss-8 | 2019-11-11T00:00:00Z | 40.9048611 | 49.9826389 | 0 | R6WY-ss-8 |
| red-tailed hawk | Raptor | winter | WY | Converse | 6 | shrub/scrub | R6WY-ss-8 | 2020-01-06T00:00:00Z | 131.03125 | NA | 0 | R6WY-ss-8 |
| red-tailed hawk | Raptor | winter | WY | Converse | 6 | shrub/scrub | R6WY-ss-8 | 2020-01-06T00:00:00Z | 110.8416667 | 121.0277778 | 0 | R6WY-ss-8 |
| red-tailed hawk | Raptor | winter | WY | Converse | 6 | shrub/scrub | R6WY-ss-8 | 2020-01-06T00:00:00Z | 121.01875 | NA | 0 | R6WY-ss-8 |
| red-tailed hawk | Raptor | winter | WY | Converse | 6 | shrub/scrub | R6WY-ss-8 | 2020-01-06T00:00:00Z | 121.00625 | NA | 0 | R6WY-ss-8 |
| red-tailed hawk | Raptor | winter | WY | Converse | 6 | shrub/scrub | R6WY-ss-8 | 2020-01-06T00:00:00Z | 121 | NA | 0 | R6WY-ss-8 |
| red-tailed hawk | Raptor | spring | WY | Converse | 6 | shrub/scrub | R6WY-ss-4 | 2018-05-22T00:00:00Z | 122.0201389 | NA | 0 | R6WY-ss-4 |
| red-tailed hawk | Raptor | spring | WY | Converse | 6 | shrub/scrub | R6WY-ss-4 | 2018-05-22T00:00:00Z | 122.0243056 | NA | 0 | R6WY-ss-4 |
| red-tailed hawk | Raptor | spring | WY | Converse | 6 | shrub/scrub | R6WY-ss-4 | 2018-05-22T00:00:00Z | 122.0194444 | NA | 0 | R6WY-ss-4 |
| red-tailed hawk | Raptor | spring | WY | Converse | 6 | shrub/scrub | R6WY-ss-4 | 2018-05-22T00:00:00Z | 122.0159722 | NA | 0 | R6WY-ss-4 |
| red-tailed hawk | Raptor | spring | WY | Converse | 6 | shrub/scrub | R6WY-ss-4 | 2018-05-22T00:00:00Z | 122.0229167 | NA | 0 | R6WY-ss-4 |
| red-tailed hawk | Raptor | summer | WY | Converse | 6 | shrub/scrub | R6WY-ss-4 | 2018-07-09T00:00:00Z | 118.875 | NA | 0 | R6WY-ss-4 |
| red-tailed hawk | Raptor | summer | WY | Converse | 6 | shrub/scrub | R6WY-ss-4 | 2018-07-09T00:00:00Z | 118.8680556 | NA | 0 | R6WY-ss-4 |
| red-tailed hawk | Raptor | summer | WY | Converse | 6 | shrub/scrub | R6WY-ss-4 | 2018-07-09T00:00:00Z | 118.8861111 | NA | 0 | R6WY-ss-4 |
| red-tailed hawk | Raptor | summer | WY | Converse | 6 | shrub/scrub | R6WY-ss-4 | 2018-07-09T00:00:00Z | 118.8944444 | NA | 0 | R6WY-ss-4 |
| red-tailed hawk | Raptor | summer | WY | Converse | 6 | shrub/scrub | R6WY-ss-4 | 2018-07-09T00:00:00Z | 118.8618056 | NA | 0 | R6WY-ss-4 |
| red-tailed hawk | Raptor | summer | WY | Converse | 6 | shrub/scrub | R6WY-ss-4 | 2018-07-26T00:00:00Z | 113.0520833 | NA | 0 | R6WY-ss-4 |
| northern harrier | Raptor | summer | WY | Converse | 6 | shrub/scrub | R6WY-ss-4 | 2018-07-26T00:00:00Z | 112.9534722 | 133.8611111 | 0 | R6WY-ss-4 |
| Cooper's hawk | Raptor | summer | WY | Converse | 6 | shrub/scrub | R6WY-ss-4 | 2018-07-26T00:00:00Z | 113.0388889 | NA | 0 | R6WY-ss-4 |
| turkey vulture | Raptor | summer | WY | Converse | 6 | shrub/scrub | R6WY-ss-4 | 2018-07-26T00:00:00Z | 133.8541667 | NA | 0 | R6WY-ss-4 |
| turkey vulture | Raptor | summer | WY | Converse | 6 | shrub/scrub | R6WY-ss-4 | 2018-07-26T00:00:00Z | 113.03125 | NA | 0 | R6WY-ss-4 |
| red-tailed hawk | Raptor | fall | WY | Converse | 6 | shrub/scrub | R6WY-ss-4 | 2018-09-10T00:00:00Z | 55.8840278 | NA | 0 | R6WY-ss-4 |
| Swainson's hawk | Raptor | fall | WY | Converse | 6 | shrub/scrub | R6WY-ss-4 | 2018-09-10T00:00:00Z | 126.8854167 | NA | 0 | R6WY-ss-4 |
| red-tailed hawk | Raptor | fall | WY | Converse | 6 | shrub/scrub | R6WY-ss-4 | 2018-09-10T00:00:00Z | 55.8743056 | 67.0722222 | 0 | R6WY-ss-4 |
| red-tailed hawk | Raptor | fall | WY | Converse | 6 | shrub/scrub | R6WY-ss-4 | 2018-09-10T00:00:00Z | 126.8708333 | NA | 0 | R6WY-ss-4 |
| Swainson's hawk | Raptor | fall | WY | Converse | 6 | shrub/scrub | R6WY-ss-4 | 2018-09-10T00:00:00Z | 126.8763889 | NA | 0 | R6WY-ss-4 |
| Cooper's hawk | Raptor | fall | WY | Converse | 6 | shrub/scrub | R6WY-ss-4 | 2018-09-24T00:00:00Z | 123.8319444 | NA | 0 | R6WY-ss-4 |
| Swainson's hawk | Raptor | fall | WY | Converse | 6 | shrub/scrub | R6WY-ss-4 | 2018-09-24T00:00:00Z | 123.8458333 | NA | 0 | R6WY-ss-4 |
| Swainson's hawk | Raptor | fall | WY | Converse | 6 | shrub/scrub | R6WY-ss-4 | 2018-09-24T00:00:00Z | 123.8423611 | NA | 0 | R6WY-ss-4 |
| red-tailed hawk | Raptor | fall | WY | Converse | 6 | shrub/scrub | R6WY-ss-4 | 2018-09-24T00:00:00Z | 53.0326389 | NA | 0 | R6WY-ss-4 |
| barred owl | Raptor | fall | WY | Converse | 6 | shrub/scrub | R6WY-ss-4 | 2018-09-24T00:00:00Z | 123.8388889 | NA | 0 | R6WY-ss-4 |
| prairie falcon | Raptor | winter | WY | Converse | 6 | shrub/scrub | R6WY-ss-4 | 2019-02-12T00:00:00Z | 111.0840278 | NA | 0 | R6WY-ss-4 |
| red-tailed hawk | Raptor | winter | WY | Converse | 6 | shrub/scrub | R6WY-ss-4 | 2019-02-25T00:00:00Z | 126.0166667 | NA | 0 | R6WY-ss-4 |
| turkey vulture | Raptor | winter | WY | Converse | 6 | shrub/scrub | R6WY-ss-4 | 2019-02-25T00:00:00Z | 126.0201389 | NA | 0 | R6WY-ss-4 |
| red-tailed hawk | Raptor | winter | WY | Converse | 6 | shrub/scrub | R6WY-ss-4 | 2019-02-25T00:00:00Z | 126.0208333 | NA | 0 | R6WY-ss-4 |
| turkey vulture | Raptor | winter | WY | Converse | 6 | shrub/scrub | R6WY-ss-4 | 2019-02-25T00:00:00Z | 126.0229167 | NA | 0 | R6WY-ss-4 |
| great horned owl | Raptor | winter | WY | Converse | 6 | shrub/scrub | R6WY-ss-4 | 2019-02-25T00:00:00Z | 126.0277778 | NA | 0 | R6WY-ss-4 |
| ferruginous hawk | Raptor | winter | WY | Converse | 6 | shrub/scrub | R6WY-ss-4 | 2019-02-25T00:00:00Z | 126.0270833 | NA | 0 | R6WY-ss-4 |
| red-tailed hawk | Raptor | winter | WY | Converse | 6 | shrub/scrub | R6WY-ss-4 | 2019-02-25T00:00:00Z | 126.0243056 | NA | 0 | R6WY-ss-4 |
| great horned owl | Raptor | winter | WY | Converse | 6 | shrub/scrub | R6WY-ss-4 | 2019-02-25T00:00:00Z | 126.0138889 | NA | 0 | R6WY-ss-4 |
| red-tailed hawk | Raptor | winter | WY | Converse | 6 | shrub/scrub | R6WY-ss-4 | 2019-02-25T00:00:00Z | 126.0263889 | NA | 0 | R6WY-ss-4 |
| turkey vulture | Raptor | summer | WY | Converse | 6 | shrub/scrub | R6WY-ss-3 | 2017-08-08T00:00:00Z | 118.9673611 | NA | 0 | R6WY-ss-3 |
| turkey vulture | Raptor | summer | WY | Converse | 6 | shrub/scrub | R6WY-ss-3 | 2017-08-08T00:00:00Z | 118.98125 | NA | 0 | R6WY-ss-3 |
| rough-legged hawk | Raptor | summer | WY | Converse | 6 | shrub/scrub | R6WY-ss-3 | 2017-08-08T00:00:00Z | 118.9777778 | NA | 0 | R6WY-ss-3 |
| red-tailed hawk | Raptor | summer | WY | Converse | 6 | shrub/scrub | R6WY-ss-3 | 2017-08-08T00:00:00Z | 103.7743056 | NA | 0 | R6WY-ss-3 |
| Cooper's hawk | Raptor | fall | WY | Converse | 6 | shrub/scrub | R6WY-ss-3 | 2017-10-13T00:00:00Z | 30.9576389 | 109 | 0 | R6WY-ss-3 |
| turkey vulture | Raptor | fall | WY | Converse | 6 | shrub/scrub | R6WY-ss-3 | 2017-10-13T00:00:00Z | 10.8340278 | 109 | 0 | R6WY-ss-3 |
| red-tailed hawk | Raptor | fall | WY | Converse | 6 | shrub/scrub | R6WY-ss-3 | 2017-10-13T00:00:00Z | 124.8618056 | NA | 0 | R6WY-ss-3 |
| red-tailed hawk | Raptor | fall | WY | Converse | 6 | shrub/scrub | R6WY-ss-3 | 2017-10-13T00:00:00Z | 124.8527778 | NA | 0 | R6WY-ss-3 |
| red-tailed hawk | Raptor | fall | WY | Converse | 6 | shrub/scrub | R6WY-ss-3 | 2017-10-13T00:00:00Z | 124.8506944 | NA | 0 | R6WY-ss-3 |
| red-tailed hawk | Raptor | winter | WY | Converse | 6 | shrub/scrub | R6WY-ss-3 | 2018-02-12T00:00:00Z | 109.0756944 | 119.9145833 | 0 | R6WY-ss-3 |
| Cooper's hawk | Raptor | winter | WY | Converse | 6 | shrub/scrub | R6WY-ss-3 | 2018-02-12T00:00:00Z | 1.0444444 | NA | 0 | R6WY-ss-3 |
| red-tailed hawk | Raptor | winter | WY | Converse | 6 | shrub/scrub | R6WY-ss-3 | 2018-02-12T00:00:00Z | 119.8916667 | NA | 0 | R6WY-ss-3 |
| red-tailed hawk | Raptor | winter | WY | Converse | 6 | shrub/scrub | R6WY-ss-3 | 2018-02-12T00:00:00Z | 119.8854167 | NA | 0 | R6WY-ss-3 |
| Cooper's hawk | Raptor | winter | WY | Converse | 6 | shrub/scrub | R6WY-ss-3 | 2018-02-12T00:00:00Z | 119.8770833 | NA | 0 | R6WY-ss-3 |
| red-tailed hawk | Raptor | spring | WY | Converse | 6 | shrub/scrub | R6WY-ss-3 | 2018-03-05T00:00:00Z | 37.9118056 | NA | 0 | R6WY-ss-3 |
| Swainson's hawk | Raptor | spring | WY | Converse | 6 | shrub/scrub | R6WY-ss-3 | 2018-03-05T00:00:00Z | 134.8611111 | NA | 0 | R6WY-ss-3 |
| great horned owl | Raptor | spring | WY | Converse | 6 | shrub/scrub | R6WY-ss-3 | 2018-03-05T00:00:00Z | 134.85 | NA | 0 | R6WY-ss-3 |
| great horned owl | Raptor | spring | WY | Converse | 6 | shrub/scrub | R6WY-ss-3 | 2018-03-05T00:00:00Z | 134.8472222 | NA | 0 | R6WY-ss-3 |
| great horned owl | Raptor | spring | WY | Converse | 6 | shrub/scrub | R6WY-ss-3 | 2018-03-05T00:00:00Z | 135.3236111 | NA | 0 | R6WY-ss-3 |
| red-tailed hawk | Raptor | spring | WY | Converse | 6 | shrub/scrub | R6WY-ss-3 | 2018-04-23T00:00:00Z | 122.8916667 | NA | 0 | R6WY-ss-3 |
| red-tailed hawk | Raptor | spring | WY | Converse | 6 | shrub/scrub | R6WY-ss-3 | 2018-04-23T00:00:00Z | 122.8895833 | NA | 0 | R6WY-ss-3 |
| red-tailed hawk | Raptor | spring | WY | Converse | 6 | shrub/scrub | R6WY-ss-3 | 2018-04-23T00:00:00Z | 122.8888889 | NA | 0 | R6WY-ss-3 |
| red-tailed hawk | Raptor | spring | WY | Converse | 6 | shrub/scrub | R6WY-ss-3 | 2018-04-23T00:00:00Z | 122.8854167 | NA | 0 | R6WY-ss-3 |
| turkey vulture | Raptor | spring | WY | Converse | 6 | shrub/scrub | R6WY-ss-3 | 2018-04-23T00:00:00Z | 122.8722222 | NA | 0 | R6WY-ss-3 |
| Swainson's hawk | Raptor | summer | WY | Converse | 6 | shrub/scrub | R6WY-ss-2 | 2016-06-21T00:00:00Z | 90 | 101 | 0 | R6WY-ss-0 |
| Swainson's hawk | Raptor | summer | WY | Converse | 6 | shrub/scrub | R6WY-ss-2 | 2016-06-21T00:00:00Z | 121 | NA | 0 | R6WY-ss-0 |
| rough-legged hawk | Raptor | summer | WY | Converse | 6 | shrub/scrub | R6WY-ss-2 | 2016-06-21T00:00:00Z | 121 | NA | 0 | R6WY-ss-0 |
| red-tailed hawk | Raptor | winter | WY | Converse | 6 | shrub/scrub | R6WY-ss-2 | 2017-02-20T00:00:00Z | 123 | NA | 0 | R6WY-ss-0 |
| red-tailed hawk | Raptor | winter | WY | Converse | 6 | shrub/scrub | R6WY-ss-2 | 2017-02-20T00:00:00Z | 123 | NA | 0 | R6WY-ss-0 |
| red-tailed hawk | Raptor | winter | WY | Converse | 6 | shrub/scrub | R6WY-ss-2 | 2017-02-20T00:00:00Z | 123 | NA | 0 | R6WY-ss-0 |
| ferruginous hawk | Raptor | winter | WY | Converse | 6 | shrub/scrub | R6WY-ss-2 | 2017-02-20T00:00:00Z | 123 | NA | 0 | R6WY-ss-0 |
| great horned owl | Raptor | winter | WY | Converse | 6 | shrub/scrub | R6WY-ss-1 | 2015-12-11T00:00:00Z | 372 | NA | 0 | R6WY-ss-0 |
| Swainson's hawk | Raptor | winter | WY | Converse | 6 | shrub/scrub | R6WY-ss-1 | 2015-12-11T00:00:00Z | 371 | NA | 0 | R6WY-ss-0 |
| Swainson's hawk | Raptor | winter | WY | Converse | 6 | shrub/scrub | R6WY-ss-1 | 2015-12-11T00:00:00Z | 119 | NA | 0 | R6WY-ss-0 |
| Swainson's hawk | Raptor | winter | WY | Converse | 6 | shrub/scrub | R6WY-ss-1 | 2015-12-11T00:00:00Z | 384 | NA | 0 | R6WY-ss-0 |
| northern harrier | Raptor | spring | WY | Converse | 6 | shrub/scrub | R6WY-ss-1 | 2016-04-25T00:00:00Z | 121 | NA | 0 | R6WY-ss-0 |
| short-eared owl | Raptor | spring | WY | Converse | 6 | shrub/scrub | R6WY-ss-1 | 2016-04-25T00:00:00Z | 121 | NA | 0 | R6WY-ss-0 |
| Swainson's hawk | Raptor | spring | WY | Converse | 6 | shrub/scrub | R6WY-ss-1 | 2016-04-25T00:00:00Z | 121 | NA | 0 | R6WY-ss-0 |
| great horned owl | Raptor | spring | WY | Converse | 6 | shrub/scrub | R6WY-ss-1 | 2016-04-25T00:00:00Z | 121 | NA | 0 | R6WY-ss-0 |
| red-tailed hawk | Raptor | fall | SD | Bon Homme, Charles Mix, Hutchinson | 6 | grassland | R6SD-g-1 | 2020-10-12T00:00:00Z | 7.075 | 22.1666667 | 0 | R6SD-g-1 |
| red-tailed hawk | Raptor | fall | SD | Bon Homme, Charles Mix, Hutchinson | 6 | grassland | R6SD-g-1 | 2020-10-20T00:00:00Z | 39.9291667 | NA | 0 | R6SD-g-1 |
| ring-necked pheasant | Game Bird | summer | NE | Holt | 6 | grassland | R6NE-g-2 | 2018-06-06T00:00:00Z | 4.2083333 | 6.9986111 | 0 | R6NE-g-2 |
| ring-necked pheasant | Game Bird | summer | NE | Holt | 6 | grassland | R6NE-g-2 | 2018-06-06T00:00:00Z | 10.2229167 | 14.2506944 | 0 | R6NE-g-2 |
| ring-necked pheasant | Game Bird | summer | NE | Holt | 6 | grassland | R6NE-g-2 | 2018-06-07T00:00:00Z | 3.14375 | 4.0569444 | 0 | R6NE-g-2 |
| ring-necked pheasant | Game Bird | summer | NE | Holt | 6 | grassland | R6NE-g-2 | 2018-06-07T00:00:00Z | 1.0284722 | 1.0284722 | 0 | R6NE-g-2 |
| ring-necked pheasant | Game Bird | summer | NE | Holt | 6 | grassland | R6NE-g-2 | 2018-06-12T00:00:00Z | 8.125 | 10.8569444 | 0 | R6NE-g-2 |
| ring-necked pheasant | Game Bird | summer | NE | Holt | 6 | grassland | R6NE-g-2 | 2018-06-14T00:00:00Z | 4.1041667 | 7.2291667 | 0 | R6NE-g-2 |
| ring-necked pheasant | Game Bird | summer | NE | Holt | 6 | grassland | R6NE-g-2 | 2018-06-14T00:00:00Z | 29.6923611 | NA | 0 | R6NE-g-2 |
| ring-necked pheasant | Game Bird | summer | NE | Holt | 6 | grassland | R6NE-g-2 | 2018-06-18T00:00:00Z | 4.3118056 | 7.8729167 | 0 | R6NE-g-2 |
| ring-necked pheasant | Game Bird | summer | NE | Holt | 6 | grassland | R6NE-g-2 | 2018-06-18T00:00:00Z | 30.04375 | NA | 0 | R6NE-g-2 |
| ring-necked pheasant | Game Bird | summer | NE | Holt | 6 | grassland | R6NE-g-2 | 2018-06-18T00:00:00Z | 4.63125 | 7.6493056 | 0 | R6NE-g-2 |
| ring-necked pheasant | Game Bird | fall | NE | Holt | 6 | grassland | R6NE-g-2 | 2018-09-05T00:00:00Z | 0.8701389 | 0.8701389 | 0 | R6NE-g-2 |
| ring-necked pheasant | Game Bird | fall | NE | Holt | 6 | grassland | R6NE-g-2 | 2018-09-05T00:00:00Z | 29.2486111 | NA | 0 | R6NE-g-2 |
| ring-necked pheasant | Game Bird | fall | NE | Holt | 6 | grassland | R6NE-g-2 | 2018-09-05T00:00:00Z | 4.2145833 | 6.9048611 | 0 | R6NE-g-2 |
| ring-necked pheasant | Game Bird | fall | NE | Holt | 6 | grassland | R6NE-g-2 | 2018-09-05T00:00:00Z | 28.9368056 | NA | 0 | R6NE-g-2 |
| ring-necked pheasant | Game Bird | fall | NE | Holt | 6 | grassland | R6NE-g-2 | 2018-09-24T00:00:00Z | 7.0944444 | 10.2625 | 0 | R6NE-g-2 |
| ring-necked pheasant | Game Bird | fall | NE | Holt | 6 | grassland | R6NE-g-2 | 2018-09-24T00:00:00Z | 29.0805556 | NA | 0 | R6NE-g-2 |
| ring-necked pheasant | Game Bird | fall | NE | Holt | 6 | grassland | R6NE-g-2 | 2018-09-24T00:00:00Z | 7.06875 | 9.8784722 | 0 | R6NE-g-2 |
| ring-necked pheasant | Game Bird | fall | NE | Holt | 6 | grassland | R6NE-g-2 | 2018-09-24T00:00:00Z | 29.1111111 | NA | 0 | R6NE-g-2 |
| ring-necked pheasant | Game Bird | fall | NE | Holt | 6 | grassland | R6NE-g-2 | 2018-10-09T00:00:00Z | 29.6493056 | NA | 0 | R6NE-g-2 |
| ring-necked pheasant | Game Bird | fall | NE | Holt | 6 | grassland | R6NE-g-2 | 2018-10-09T00:00:00Z | 3.6729167 | 6.7534722 | 0 | R6NE-g-2 |
| ring-necked pheasant | Game Bird | winter | NE | Holt | 6 | grassland | R6NE-g-2 | 2019-01-07T00:00:00Z | 4.0423611 | 7.2416667 | 0 | R6NE-g-2 |
| ring-necked pheasant | Game Bird | winter | NE | Holt | 6 | grassland | R6NE-g-2 | 2019-01-07T00:00:00Z | 21.1479167 | 30.1277778 | 0 | R6NE-g-2 |
| ring-necked pheasant | Game Bird | winter | NE | Holt | 6 | grassland | R6NE-g-2 | 2019-01-07T00:00:00Z | 14.0131944 | 21.1256944 | 0 | R6NE-g-2 |
| ring-necked pheasant | Game Bird | winter | NE | Holt | 6 | grassland | R6NE-g-2 | 2019-01-07T00:00:00Z | 7 | 9.91875 | 0 | R6NE-g-2 |
| ring-necked pheasant | Game Bird | winter | NE | Holt | 6 | grassland | R6NE-g-2 | 2019-02-04T00:00:00Z | 2.2 | 29.9555556 | 0 | R6NE-g-2 |
| ring-necked pheasant | Game Bird | winter | NE | Holt | 6 | grassland | R6NE-g-2 | 2019-02-05T00:00:00Z | 7.1291667 | 9.8541667 | 0 | R6NE-g-2 |
| ring-necked pheasant | Game Bird | winter | NE | Holt | 6 | grassland | R6NE-g-2 | 2019-02-05T00:00:00Z | 14.1479167 | 29.9861111 | 0 | R6NE-g-2 |
| ring-necked pheasant | Game Bird | winter | NE | Holt | 6 | grassland | R6NE-g-2 | 2019-02-05T00:00:00Z | 29.9930556 | NA | 0 | R6NE-g-2 |
| ring-necked pheasant | Game Bird | winter | NE | Holt | 6 | grassland | R6NE-g-2 | 2019-02-11T00:00:00Z | 21.0354167 | NA | 0 | R6NE-g-2 |
| ring-necked pheasant | Game Bird | winter | NE | Holt | 6 | grassland | R6NE-g-2 | 2019-02-11T00:00:00Z | 36.3236111 | NA | 0 | R6NE-g-2 |
| red-tailed hawk | Raptor | spring | NE | Holt | 6 | grassland | R6NE-g-2 | 2019-04-03T00:00:00Z | 3.8993056 | 13.9576389 | 0 | R6NE-g-2 |
| ring-necked pheasant | Game Bird | spring | NE | Holt | 6 | grassland | R6NE-g-2 | 2019-04-01T00:00:00Z | 20.9722222 | 29.9777778 | 0 | R6NE-g-2 |
| ring-necked pheasant | Game Bird | spring | NE | Holt | 6 | grassland | R6NE-g-2 | 2019-04-01T00:00:00Z | 14.3145833 | 20.9298611 | 0 | R6NE-g-2 |
| ring-necked pheasant | Game Bird | spring | NE | Holt | 6 | grassland | R6NE-g-2 | 2019-04-01T00:00:00Z | 6.9444444 | 13.98125 | 0 | R6NE-g-2 |
| ring-necked pheasant | Game Bird | spring | NE | Holt | 6 | grassland | R6NE-g-2 | 2019-04-01T00:00:00Z | 3.9805556 | 6.95 | 0 | R6NE-g-2 |
| ring-necked pheasant | Game Bird | spring | NE | Holt | 6 | grassland | R6NE-g-2 | 2019-04-01T00:00:00Z | 3.0263889 | 4.0381944 | 0 | R6NE-g-2 |
| ring-necked pheasant | Game Bird | spring | NE | Holt | 6 | grassland | R6NE-g-2 | 2019-04-08T00:00:00Z | 13.9909722 | 20.9395833 | 0 | R6NE-g-2 |
| ring-necked pheasant | Game Bird | spring | NE | Holt | 6 | grassland | R6NE-g-2 | 2019-04-08T00:00:00Z | 9.8104167 | 14.0840278 | 0 | R6NE-g-2 |
| ring-necked pheasant | Game Bird | spring | NE | Holt | 6 | grassland | R6NE-g-2 | 2019-04-08T00:00:00Z | 14.1451389 | 20.9694444 | 0 | R6NE-g-2 |
| ring-necked pheasant | Game Bird | spring | NE | Holt | 6 | grassland | R6NE-g-2 | 2019-04-08T00:00:00Z | 2.0375 | 10.0090278 | 0 | R6NE-g-2 |
| ring-necked pheasant | Game Bird | spring | NE | Holt | 6 | grassland | R6NE-g-2 | 2019-04-08T00:00:00Z | 14.4145833 | 31.0131944 | 0 | R6NE-g-2 |
| red-tailed hawk | Raptor | spring | WY | Converse | 6 | grassland | R6WY-g-11 | 2020-05-18T00:00:00Z | 83.9305556 | 147.9027778 | 0 | R6WY-g-11 |
| red-tailed hawk | Raptor | spring | WY | Converse | 6 | grassland | R6WY-g-11 | 2020-05-18T00:00:00Z | 62.8576389 | 72.9041667 | 0 | R6WY-g-11 |
| red-tailed hawk | Raptor | spring | WY | Converse | 6 | grassland | R6WY-g-11 | 2020-05-18T00:00:00Z | 83.9270833 | NA | 0 | R6WY-g-11 |
| red-tailed hawk | Raptor | spring | WY | Converse | 6 | grassland | R6WY-g-11 | 2020-05-18T00:00:00Z | 83.9138889 | NA | 0 | R6WY-g-11 |
| red-tailed hawk | Raptor | spring | WY | Converse | 6 | grassland | R6WY-g-11 | 2020-05-18T00:00:00Z | 83.9104167 | NA | 0 | R6WY-g-11 |
| barn owl | Raptor | spring | WY | Converse | 6 | grassland | R6WY-g-11 | 2020-05-18T00:00:00Z | 83.8958333 | NA | 0 | R6WY-g-11 |
| red-tailed hawk | Raptor | spring | WY | Converse | 6 | grassland | R6WY-g-11 | 2020-05-18T00:00:00Z | 83.8986111 | NA | 0 | R6WY-g-11 |
| red-tailed hawk | Raptor | spring | WY | Converse | 6 | grassland | R6WY-g-11 | 2020-05-18T00:00:00Z | 72.9402778 | NA | 0 | R6WY-g-11 |
| red-tailed hawk | Raptor | spring | WY | Converse | 6 | grassland | R6WY-g-11 | 2020-05-18T00:00:00Z | 83.89375 | NA | 0 | R6WY-g-11 |
| Swainson's hawk | Raptor | fall | WY | Converse | 6 | grassland | R6WY-g-11 | 2020-10-19T00:00:00Z | 20.9972222 | 41.94375 | 0 | R6WY-g-11 |
| red-tailed hawk | Raptor | spring | WY | Converse | 6 | grassland | R6WY-g-11 | 2020-05-18T00:00:00Z | 72.9319444 | 83.8847222 | 0 | R6WY-g-11 |
| Swainson's hawk | Raptor | fall | WY | Converse | 6 | grassland | R6WY-g-11 | 2020-10-19T00:00:00Z | 20.9944444 | 41.9409722 | 0 | R6WY-g-11 |
| Swainson's hawk | Raptor | fall | WY | Converse | 6 | grassland | R6WY-g-11 | 2020-10-19T00:00:00Z | 20.9881944 | 41.9548611 | 0 | R6WY-g-11 |
| Swainson's hawk | Raptor | fall | WY | Converse | 6 | grassland | R6WY-g-11 | 2020-10-19T00:00:00Z | 20.9861111 | 41.9402778 | 0 | R6WY-g-11 |
| Swainson's hawk | Raptor | fall | WY | Converse | 6 | grassland | R6WY-g-11 | 2020-10-19T00:00:00Z | 20.9868056 | 41.9395833 | 0 | R6WY-g-11 |
| Swainson's hawk | Raptor | fall | WY | Converse | 6 | grassland | R6WY-g-11 | 2020-10-19T00:00:00Z | 1.9847222 | 6.9152778 | 0 | R6WY-g-11 |
| Swainson's hawk | Raptor | fall | WY | Converse | 6 | grassland | R6WY-g-11 | 2020-10-19T00:00:00Z | 1.9805556 | 3.9673611 | 0 | R6WY-g-11 |
| Swainson's hawk | Raptor | fall | WY | Converse | 6 | grassland | R6WY-g-11 | 2020-10-19T00:00:00Z | 20.9722222 | 41.9270833 | 0 | R6WY-g-11 |
| Swainson's hawk | Raptor | fall | WY | Converse | 6 | grassland | R6WY-g-11 | 2020-10-19T00:00:00Z | 20.9763889 | 41.9284722 | 0 | R6WY-g-11 |
| Swainson's hawk | Raptor | fall | WY | Converse | 6 | grassland | R6WY-g-11 | 2020-10-19T00:00:00Z | 20.9729167 | 41.925 | 0 | R6WY-g-11 |
| Swainson's hawk | Raptor | summer | WY | Converse | 6 | grassland | R6WY-g-11 | 2020-07-20T00:00:00Z | 21.04375 | 91.0215278 | 0 | R6WY-g-11 |
| Swainson's hawk | Raptor | summer | WY | Converse | 6 | grassland | R6WY-g-11 | 2020-07-20T00:00:00Z | 21.0381944 | 91.0166667 | 0 | R6WY-g-11 |
| red-tailed hawk | Raptor | summer | WY | Converse | 6 | grassland | R6WY-g-11 | 2020-07-20T00:00:00Z | 7.0597222 | 10.0430556 | 0 | R6WY-g-11 |
| Swainson's hawk | Raptor | summer | WY | Converse | 6 | grassland | R6WY-g-11 | 2020-07-20T00:00:00Z | 133.0409722 | NA | 0 | R6WY-g-11 |
| Swainson's hawk | Raptor | summer | WY | Converse | 6 | grassland | R6WY-g-11 | 2020-07-20T00:00:00Z | 133.0159722 | NA | 0 | R6WY-g-11 |
| barn owl | Raptor | summer | WY | Converse | 6 | grassland | R6WY-g-11 | 2020-07-20T00:00:00Z | 91.0597222 | 101.0076389 | 0 | R6WY-g-11 |
| red-tailed hawk | Raptor | winter | WY | Converse | 6 | grassland | R6WY-g-11 | 2021-01-12T00:00:00Z | 0.9979167 | 2.0048611 | 0 | R6WY-g-11 |
| prairie falcon | Raptor | summer | WY | Converse | 6 | grassland | R6WY-g-11 | 2020-07-20T00:00:00Z | 21.0055556 | 91.0583333 | 0 | R6WY-g-11 |
| great horned owl | Raptor | winter | WY | Converse | 6 | grassland | R6WY-g-11 | 2021-01-12T00:00:00Z | 59.0402778 | 70.1027778 | 0 | R6WY-g-11 |
| red-tailed hawk | Raptor | winter | WY | Converse | 6 | grassland | R6WY-g-11 | 2021-01-12T00:00:00Z | 13.9305556 | 20.0055556 | 0 | R6WY-g-11 |
| red-tailed hawk | Raptor | winter | WY | Converse | 6 | grassland | R6WY-g-11 | 2021-01-12T00:00:00Z | 30.0152778 | 39.9854167 | 0 | R6WY-g-11 |
| red-tailed hawk | Raptor | winter | WY | Converse | 6 | grassland | R6WY-g-11 | 2021-01-12T00:00:00Z | 59.0277778 | 70.0631944 | 0 | R6WY-g-11 |
| red-tailed hawk | Raptor | winter | WY | Converse | 6 | grassland | R6WY-g-11 | 2021-01-12T00:00:00Z | 59.0166667 | 69.0527778 | 0 | R6WY-g-11 |
| red-tailed hawk | Raptor | winter | WY | Converse | 6 | grassland | R6WY-g-11 | 2021-01-12T00:00:00Z | 29.9791667 | 39.9694444 | 0 | R6WY-g-11 |
| red-tailed hawk | Raptor | winter | WY | Converse | 6 | grassland | R6WY-g-11 | 2021-01-12T00:00:00Z | 89.9951389 | 100.0423611 | 0 | R6WY-g-11 |
| red-tailed hawk | Raptor | winter | WY | Converse | 6 | grassland | R6WY-g-11 | 2021-01-12T00:00:00Z | 119.94375 | NA | 0 | R6WY-g-11 |
| red-tailed hawk | Raptor | winter | WY | Converse | 6 | grassland | R6WY-g-11 | 2021-01-12T00:00:00Z | 2.9736111 | 6.8736111 | 0 | R6WY-g-11 |
| great horned owl | Raptor | spring | WY | Converse | 6 | grassland | R6WY-g-10 | 2019-04-15T00:00:00Z | 136.7416667 | NA | 0 | R6WY-g-10 |
| red-tailed hawk | Raptor | spring | WY | Converse | 6 | grassland | R6WY-g-10 | 2019-04-15T00:00:00Z | 136.7409722 | NA | 0 | R6WY-g-10 |
| red-tailed hawk | Raptor | spring | WY | Converse | 6 | grassland | R6WY-g-10 | 2019-04-15T00:00:00Z | 136.7416667 | NA | 0 | R6WY-g-10 |
| red-tailed hawk | Raptor | spring | WY | Converse | 6 | grassland | R6WY-g-10 | 2019-04-15T00:00:00Z | 136.7520833 | NA | 0 | R6WY-g-10 |
| red-tailed hawk | Raptor | spring | WY | Converse | 6 | grassland | R6WY-g-10 | 2019-04-24T00:00:00Z | 127.8006944 | NA | 0 | R6WY-g-10 |
| red-tailed hawk | Raptor | spring | WY | Converse | 6 | grassland | R6WY-g-10 | 2019-04-24T00:00:00Z | 127.7993056 | NA | 0 | R6WY-g-10 |
| red-tailed hawk | Raptor | spring | WY | Converse | 6 | grassland | R6WY-g-10 | 2019-04-24T00:00:00Z | 127.7993056 | NA | 0 | R6WY-g-10 |
| red-tailed hawk | Raptor | spring | WY | Converse | 6 | grassland | R6WY-g-10 | 2019-04-24T00:00:00Z | 127.8006944 | NA | 0 | R6WY-g-10 |
| red-tailed hawk | Raptor | spring | WY | Converse | 6 | grassland | R6WY-g-10 | 2019-04-24T00:00:00Z | 127.7972222 | NA | 0 | R6WY-g-10 |
| red-tailed hawk | Raptor | spring | WY | Converse | 6 | grassland | R6WY-g-10 | 2019-04-24T00:00:00Z | 127.7944444 | NA | 0 | R6WY-g-10 |
| northern harrier | Raptor | summer | WY | Converse | 6 | grassland | R6WY-g-10 | 2019-06-24T00:00:00Z | 134.8944444 | NA | 0 | R6WY-g-10 |
| northern harrier | Raptor | summer | WY | Converse | 6 | grassland | R6WY-g-10 | 2019-06-24T00:00:00Z | 134.9326389 | NA | 0 | R6WY-g-10 |
| turkey vulture | Raptor | summer | WY | Converse | 6 | grassland | R6WY-g-10 | 2019-06-24T00:00:00Z | 134.9513889 | NA | 0 | R6WY-g-10 |
| northern harrier | Raptor | summer | WY | Converse | 6 | grassland | R6WY-g-10 | 2019-06-24T00:00:00Z | 1.9986111 | NA | 0 | R6WY-g-10 |
| northern harrier | Raptor | summer | WY | Converse | 6 | grassland | R6WY-g-10 | 2019-06-24T00:00:00Z | 134.9305556 | NA | 0 | R6WY-g-10 |
| northern harrier | Raptor | summer | WY | Converse | 6 | grassland | R6WY-g-10 | 2019-06-24T00:00:00Z | 134.9416667 | NA | 0 | R6WY-g-10 |
| turkey vulture | Raptor | summer | WY | Converse | 6 | grassland | R6WY-g-10 | 2019-06-24T00:00:00Z | 134.9833333 | NA | 0 | R6WY-g-10 |
| turkey vulture | Raptor | summer | WY | Converse | 6 | grassland | R6WY-g-10 | 2019-06-24T00:00:00Z | 135.0069444 | NA | 0 | R6WY-g-10 |
| turkey vulture | Raptor | summer | WY | Converse | 6 | grassland | R6WY-g-10 | 2019-06-24T00:00:00Z | 135.0458333 | NA | 0 | R6WY-g-10 |
| northern harrier | Raptor | summer | WY | Converse | 6 | grassland | R6WY-g-10 | 2019-06-24T00:00:00Z | 134.8631944 | NA | 0 | R6WY-g-10 |
| turkey vulture | Raptor | fall | WY | Converse | 6 | grassland | R6WY-g-10 | 2019-10-21T00:00:00Z | 124.9347222 | NA | 0 | R6WY-g-10 |
| northern harrier | Raptor | fall | WY | Converse | 6 | grassland | R6WY-g-10 | 2019-10-21T00:00:00Z | 13.9652778 | 20.9333333 | 0 | R6WY-g-10 |
| northern harrier | Raptor | fall | WY | Converse | 6 | grassland | R6WY-g-10 | 2019-10-21T00:00:00Z | 3.9625 | 13.9840278 | 0 | R6WY-g-10 |
| northern harrier | Raptor | fall | WY | Converse | 6 | grassland | R6WY-g-10 | 2019-10-21T00:00:00Z | 3.94375 | 6.9854167 | 0 | R6WY-g-10 |
| northern harrier | Raptor | fall | WY | Converse | 6 | grassland | R6WY-g-10 | 2019-10-21T00:00:00Z | 3.9520833 | 9.9597222 | 0 | R6WY-g-10 |
| red-tailed hawk | Raptor | winter | WY | Converse | 6 | grassland | R6WY-g-10 | 2019-11-12T00:00:00Z | 120.8923611 | NA | 0 | R6WY-g-10 |
| red-tailed hawk | Raptor | winter | WY | Converse | 6 | grassland | R6WY-g-10 | 2019-11-12T00:00:00Z | 120.8777778 | NA | 0 | R6WY-g-10 |
| red-tailed hawk | Raptor | winter | WY | Converse | 6 | grassland | R6WY-g-10 | 2019-11-12T00:00:00Z | 120.8701389 | NA | 0 | R6WY-g-10 |
| turkey vulture | Raptor | winter | WY | Converse | 6 | grassland | R6WY-g-10 | 2019-11-12T00:00:00Z | 9.9972222 | 110.9548611 | 0 | R6WY-g-10 |
| red-tailed hawk | Raptor | winter | WY | Converse | 6 | grassland | R6WY-g-10 | 2019-11-12T00:00:00Z | 10.0236111 | 68.9055556 | 0 | R6WY-g-10 |
| northern harrier | Raptor | winter | WY | Converse | 6 | grassland | R6WY-g-10 | 2020-01-14T00:00:00Z | 123.0284722 | NA | 0 | R6WY-g-10 |
| red-tailed hawk | Raptor | winter | WY | Converse | 6 | grassland | R6WY-g-10 | 2020-01-14T00:00:00Z | 123.0319444 | NA | 0 | R6WY-g-10 |
| red-tailed hawk | Raptor | winter | WY | Converse | 6 | grassland | R6WY-g-10 | 2020-01-14T00:00:00Z | 122.9923611 | NA | 0 | R6WY-g-10 |
| red-tailed hawk | Raptor | winter | WY | Converse | 6 | grassland | R6WY-g-10 | 2020-01-14T00:00:00Z | 122.9840278 | NA | 0 | R6WY-g-10 |
| red-tailed hawk | Raptor | winter | WY | Converse | 6 | grassland | R6WY-g-10 | 2020-01-14T00:00:00Z | 122.9798611 | NA | 0 | R6WY-g-10 |
| red-tailed hawk | Raptor | spring | WY | Converse | 6 | grassland | R6WY-g-9 | 2018-05-29T00:00:00Z | 128.1097222 | NA | 0 | R6WY-g-9 |
| red-tailed hawk | Raptor | spring | WY | Converse | 6 | grassland | R6WY-g-9 | 2018-05-29T00:00:00Z | 128.0951389 | NA | 0 | R6WY-g-9 |
| red-tailed hawk | Raptor | spring | WY | Converse | 6 | grassland | R6WY-g-9 | 2018-05-29T00:00:00Z | 128.0291667 | NA | 0 | R6WY-g-9 |
| Cooper's hawk | Raptor | spring | WY | Converse | 6 | grassland | R6WY-g-9 | 2018-05-29T00:00:00Z | 128.0201389 | NA | 0 | R6WY-g-9 |
| red-tailed hawk | Raptor | spring | WY | Converse | 6 | grassland | R6WY-g-9 | 2018-05-29T00:00:00Z | 128.0041667 | NA | 0 | R6WY-g-9 |
| red-tailed hawk | Raptor | summer | WY | Converse | 6 | grassland | R6WY-g-9 | 2018-07-10T00:00:00Z | 15.8493056 | NA | 0 | R6WY-g-9 |
| red-tailed hawk | Raptor | summer | WY | Converse | 6 | grassland | R6WY-g-9 | 2018-07-10T00:00:00Z | 108.0236111 | NA | 0 | R6WY-g-9 |
| red-tailed hawk | Raptor | summer | WY | Converse | 6 | grassland | R6WY-g-9 | 2018-07-10T00:00:00Z | 108.0305556 | NA | 0 | R6WY-g-9 |
| red-tailed hawk | Raptor | summer | WY | Converse | 6 | grassland | R6WY-g-9 | 2018-07-10T00:00:00Z | 68.95 | NA | 0 | R6WY-g-9 |
| red-tailed hawk | Raptor | summer | WY | Converse | 6 | grassland | R6WY-g-9 | 2018-07-10T00:00:00Z | 108.0291667 | NA | 0 | R6WY-g-9 |
| Cooper's hawk | Raptor | summer | WY | Converse | 6 | grassland | R6WY-g-9 | 2018-07-30T00:00:00Z | 53.0416667 | NA | 0 | R6WY-g-9 |
| turkey vulture | Raptor | summer | WY | Converse | 6 | grassland | R6WY-g-9 | 2018-07-30T00:00:00Z | 130.0611111 | NA | 0 | R6WY-g-9 |
| northern harrier | Raptor | summer | WY | Converse | 6 | grassland | R6WY-g-9 | 2018-07-30T00:00:00Z | 108.9993056 | NA | 0 | R6WY-g-9 |
| Cooper's hawk | Raptor | summer | WY | Converse | 6 | grassland | R6WY-g-9 | 2018-07-30T00:00:00Z | 23.8222222 | NA | 0 | R6WY-g-9 |
| red-tailed hawk | Raptor | fall | WY | Converse | 6 | grassland | R6WY-g-9 | 2018-09-11T00:00:00Z | 65.8465278 | NA | 0 | R6WY-g-9 |
| rough-legged hawk | Raptor | summer | WY | Converse | 6 | grassland | R6WY-g-9 | 2018-07-30T00:00:00Z | 109.0291667 | NA | 0 | R6WY-g-9 |
| Swainson's hawk | Raptor | fall | WY | Converse | 6 | grassland | R6WY-g-9 | 2018-09-11T00:00:00Z | 136.8784722 | NA | 0 | R6WY-g-9 |
| Cooper's hawk | Raptor | fall | WY | Converse | 6 | grassland | R6WY-g-9 | 2018-09-11T00:00:00Z | 54.8965278 | NA | 0 | R6WY-g-9 |
| red-tailed hawk | Raptor | fall | WY | Converse | 6 | grassland | R6WY-g-9 | 2018-09-11T00:00:00Z | 136.8715278 | NA | 0 | R6WY-g-9 |
| Cooper's hawk | Raptor | fall | WY | Converse | 6 | grassland | R6WY-g-9 | 2018-09-25T00:00:00Z | 1.8138889 | NA | 0 | R6WY-g-9 |
| Cooper's hawk | Raptor | fall | WY | Converse | 6 | grassland | R6WY-g-9 | 2018-09-25T00:00:00Z | 1.8027778 | NA | 0 | R6WY-g-9 |
| Cooper's hawk | Raptor | fall | WY | Converse | 6 | grassland | R6WY-g-9 | 2018-09-25T00:00:00Z | 1.8319444 | NA | 0 | R6WY-g-9 |
| Cooper's hawk | Raptor | fall | WY | Converse | 6 | grassland | R6WY-g-9 | 2018-09-25T00:00:00Z | 1.84375 | NA | 0 | R6WY-g-9 |
| Cooper's hawk | Raptor | fall | WY | Converse | 6 | grassland | R6WY-g-9 | 2018-09-25T00:00:00Z | 1.8555556 | NA | 0 | R6WY-g-9 |
| ferruginous hawk | Raptor | fall | WY | Converse | 6 | grassland | R6WY-g-9 | 2018-09-11T00:00:00Z | 9.0111111 | NA | 0 | R6WY-g-9 |
| ferruginous hawk | Raptor | winter | WY | Converse | 6 | grassland | R6WY-g-9 | 2019-02-25T00:00:00Z | 122.8180556 | NA | 0 | R6WY-g-9 |
| turkey vulture | Raptor | winter | WY | Converse | 6 | grassland | R6WY-g-9 | 2019-02-25T00:00:00Z | 122.7833333 | NA | 0 | R6WY-g-9 |
| red-tailed hawk | Raptor | winter | WY | Converse | 6 | grassland | R6WY-g-9 | 2019-02-25T00:00:00Z | 78.7951389 | NA | 0 | R6WY-g-9 |
| red-tailed hawk | Raptor | winter | WY | Converse | 6 | grassland | R6WY-g-9 | 2019-02-25T00:00:00Z | 9.8020833 | 22.8305556 | 0 | R6WY-g-9 |
| Cooper's hawk | Raptor | winter | WY | Converse | 6 | grassland | R6WY-g-9 | 2019-02-12T00:00:00Z | 6.8166667 | 13.0944444 | 0 | R6WY-g-9 |
| Cooper's hawk | Raptor | winter | WY | Converse | 6 | grassland | R6WY-g-9 | 2019-02-12T00:00:00Z | 1.0604167 | NA | 0 | R6WY-g-9 |
| Cooper's hawk | Raptor | winter | WY | Converse | 6 | grassland | R6WY-g-9 | 2019-02-12T00:00:00Z | 6.8173611 | 13.1048611 | 0 | R6WY-g-9 |
| red-tailed hawk | Raptor | winter | WY | Converse | 6 | grassland | R6WY-g-9 | 2019-02-25T00:00:00Z | 65.7826389 | NA | 0 | R6WY-g-9 |
| great horned owl | Raptor | winter | WY | Converse | 6 | grassland | R6WY-g-9 | 2019-02-25T00:00:00Z | 78.8034722 | NA | 0 | R6WY-g-9 |
| red-tailed hawk | Raptor | winter | WY | Converse | 6 | grassland | R6WY-g-9 | 2019-02-25T00:00:00Z | 2.0277778 | NA | 0 | R6WY-g-9 |
| red-tailed hawk | Raptor | summer | WY | Converse | 6 | grassland | R6WY-g-8 | 2017-08-28T00:00:00Z | 154.1722222 | NA | 0 | R6WY-g-8 |
| red-tailed hawk | Raptor | summer | WY | Converse | 6 | grassland | R6WY-g-8 | 2017-08-28T00:00:00Z | 154.1243056 | NA | 0 | R6WY-g-8 |
| red-tailed hawk | Raptor | summer | WY | Converse | 6 | grassland | R6WY-g-8 | 2017-08-28T00:00:00Z | 30.0743056 | 104.90625 | 0 | R6WY-g-8 |
| red-tailed hawk | Raptor | summer | WY | Converse | 6 | grassland | R6WY-g-8 | 2017-08-28T00:00:00Z | 153.9729167 | NA | 0 | R6WY-g-8 |
| Swainson's hawk | Raptor | summer | WY | Converse | 6 | grassland | R6WY-g-8 | 2017-08-28T00:00:00Z | 104.8770833 | NA | 0 | R6WY-g-8 |
| ferruginous hawk | Raptor | fall | WY | Converse | 6 | grassland | R6WY-g-8 | 2017-10-13T00:00:00Z | 122.6819444 | NA | 0 | R6WY-g-8 |
| red-tailed hawk | Raptor | fall | WY | Converse | 6 | grassland | R6WY-g-8 | 2017-10-13T00:00:00Z | 122.6868056 | NA | 0 | R6WY-g-8 |
| red-tailed hawk | Raptor | fall | WY | Converse | 6 | grassland | R6WY-g-8 | 2017-10-13T00:00:00Z | 122.6888889 | NA | 0 | R6WY-g-8 |
| red-tailed hawk | Raptor | fall | WY | Converse | 6 | grassland | R6WY-g-8 | 2017-10-13T00:00:00Z | 122.6861111 | NA | 0 | R6WY-g-8 |
| red-tailed hawk | Raptor | fall | WY | Converse | 6 | grassland | R6WY-g-8 | 2017-10-13T00:00:00Z | 122.6819444 | NA | 0 | R6WY-g-8 |
| Swainson's hawk | Raptor | winter | WY | Converse | 6 | grassland | R6WY-g-8 | 2018-02-12T00:00:00Z | 37.0291667 | 52.9166667 | 0 | R6WY-g-8 |
| ferruginous hawk | Raptor | winter | WY | Converse | 6 | grassland | R6WY-g-8 | 2018-02-12T00:00:00Z | 3.7569444 | 27.7826389 | 0 | R6WY-g-8 |
| red-tailed hawk | Raptor | winter | WY | Converse | 6 | grassland | R6WY-g-8 | 2018-02-12T00:00:00Z | 79.7694444 | 91.7763889 | 0 | R6WY-g-8 |
| red-tailed hawk | Raptor | winter | WY | Converse | 6 | grassland | R6WY-g-8 | 2018-02-12T00:00:00Z | 91.7756944 | NA | 0 | R6WY-g-8 |
| red-tailed hawk | Raptor | winter | WY | Converse | 6 | grassland | R6WY-g-8 | 2018-02-12T00:00:00Z | 108.8506944 | NA | 0 | R6WY-g-8 |
| red-tailed hawk | Raptor | winter | WY | Converse | 6 | grassland | R6WY-g-8 | 2018-03-05T00:00:00Z | 127.7256944 | NA | 0 | R6WY-g-8 |
| ferruginous hawk | Raptor | winter | WY | Converse | 6 | grassland | R6WY-g-8 | 2018-03-05T00:00:00Z | 127.7222222 | NA | 0 | R6WY-g-8 |
| turkey vulture | Raptor | winter | WY | Converse | 6 | grassland | R6WY-g-8 | 2018-03-05T00:00:00Z | 127.7208333 | NA | 0 | R6WY-g-8 |
| Swainson's hawk | Raptor | winter | WY | Converse | 6 | grassland | R6WY-g-8 | 2018-03-05T00:00:00Z | 127.7159722 | NA | 0 | R6WY-g-8 |
| osprey | Raptor | winter | WY | Converse | 6 | grassland | R6WY-g-8 | 2018-03-05T00:00:00Z | 15.7493056 | 31.8194444 | 0 | R6WY-g-8 |
| red-tailed hawk | Raptor | spring | WY | Converse | 6 | grassland | R6WY-g-8 | 2018-04-23T00:00:00Z | 22.0479167 | NA | 0 | R6WY-g-8 |
| red-tailed hawk | Raptor | spring | WY | Converse | 6 | grassland | R6WY-g-8 | 2018-04-23T00:00:00Z | 121.8333333 | NA | 0 | R6WY-g-8 |
| red-tailed hawk | Raptor | spring | WY | Converse | 6 | grassland | R6WY-g-8 | 2018-04-23T00:00:00Z | 121.83125 | NA | 0 | R6WY-g-8 |
| red-tailed hawk | Raptor | spring | WY | Converse | 6 | grassland | R6WY-g-8 | 2018-04-23T00:00:00Z | 121.8277778 | NA | 0 | R6WY-g-8 |
| red-tailed hawk | Raptor | spring | WY | Converse | 6 | grassland | R6WY-g-8 | 2018-04-23T00:00:00Z | 121.8263889 | NA | 0 | R6WY-g-8 |
| Swainson's hawk | Raptor | summer | WY | Converse | 6 | grassland | R6WY-g-7 | 2016-06-06T00:00:00Z | 112 | 121 | 1 | R6WY-g-0 |
| red-tailed hawk | Raptor | summer | WY | Converse | 6 | grassland | R6WY-g-7 | 2016-06-06T00:00:00Z | 112 | 121 | 1 | R6WY-g-0 |
| Swainson's hawk | Raptor | summer | WY | Converse | 6 | grassland | R6WY-g-7 | 2016-06-06T00:00:00Z | 121 | NA | 1 | R6WY-g-0 |
| ferruginous hawk | Raptor | winter | WY | Converse | 6 | grassland | R6WY-g-7 | 2017-02-13T00:00:00Z | 120 | NA | 1 | R6WY-g-0 |
| great horned owl | Raptor | winter | WY | Converse | 6 | grassland | R6WY-g-7 | 2017-02-13T00:00:00Z | 120 | NA | 1 | R6WY-g-0 |
| red-tailed hawk | Raptor | winter | WY | Converse | 6 | grassland | R6WY-g-7 | 2017-02-13T00:00:00Z | 120 | NA | 1 | R6WY-g-0 |
| red-tailed hawk | Raptor | winter | WY | Converse | 6 | grassland | R6WY-g-7 | 2017-02-13T00:00:00Z | 120 | NA | 1 | R6WY-g-0 |
| Swainson's hawk | Raptor | winter | WY | Converse | 6 | grassland | R6WY-g-6 | 2015-12-07T00:00:00Z | 121 | NA | 1 | R6WY-g-0 |
| Swainson's hawk | Raptor | winter | WY | Converse | 6 | grassland | R6WY-g-6 | 2015-12-07T00:00:00Z | 370 | NA | 1 | R6WY-g-0 |
| great horned owl | Raptor | winter | WY | Converse | 6 | grassland | R6WY-g-6 | 2015-12-17T00:00:00Z | 111 | NA | 1 | R6WY-g-0 |
| Swainson's hawk | Raptor | winter | WY | Converse | 6 | grassland | R6WY-g-6 | 2015-12-07T00:00:00Z | 388 | NA | 1 | R6WY-g-0 |
| Swainson's hawk | Raptor | spring | WY | Converse | 6 | grassland | R6WY-g-6 | 2016-05-09T00:00:00Z | 119 | NA | 1 | R6WY-g-0 |
| red-tailed hawk | Raptor | spring | WY | Converse | 6 | grassland | R6WY-g-6 | 2016-05-09T00:00:00Z | 119 | NA | 1 | R6WY-g-0 |
| ferruginous hawk | Raptor | spring | WY | Converse | 6 | grassland | R6WY-g-6 | 2016-05-09T00:00:00Z | 120 | NA | 1 | R6WY-g-0 |
| Swainson's hawk | Raptor | spring | WY | Converse | 6 | grassland | R6WY-g-6 | 2016-05-09T00:00:00Z | 119 | NA | 1 | R6WY-g-0 |
| ferruginous hawk | Raptor | spring | WY | Converse | 6 | grassland | R6WY-g-4 | 2015-04-13T00:00:00Z | 119 | NA | 0 | R6WY-g-4 |
| red-tailed hawk | Raptor | summer | WY | Converse | 6 | grassland | R6WY-g-4 | 2014-09-22T00:00:00Z | 120 | NA | 0 | R6WY-g-4 |
| Swainson's hawk | Raptor | summer | WY | Converse | 6 | grassland | R6WY-g-4 | 2014-09-22T00:00:00Z | 21 | 50 | 0 | R6WY-g-4 |
| Swainson's hawk | Raptor | summer | WY | Converse | 6 | grassland | R6WY-g-4 | 2014-09-22T00:00:00Z | 30 | 50 | 0 | R6WY-g-4 |
| Swainson's hawk | Raptor | summer | WY | Converse | 6 | grassland | R6WY-g-4 | 2014-09-22T00:00:00Z | 21 | 50 | 0 | R6WY-g-4 |
| ferruginous hawk | Raptor | summer | WY | Converse | 6 | grassland | R6WY-g-4 | 2014-09-22T00:00:00Z | 21 | 50 | 0 | R6WY-g-4 |
| red-tailed hawk | Raptor | summer | WY | Converse | 6 | grassland | R6WY-g-4 | 2014-09-22T00:00:00Z | 14 | 50 | 0 | R6WY-g-4 |
| Swainson's hawk | Raptor | summer | WY | Converse | 6 | grassland | R6WY-g-4 | 2014-09-22T00:00:00Z | 14 | 50 | 0 | R6WY-g-4 |
| Swainson's hawk | Raptor | summer | WY | Converse | 6 | grassland | R6WY-g-4 | 2014-09-22T00:00:00Z | 21 | 50 | 0 | R6WY-g-4 |
| red-tailed hawk | Raptor | summer | WY | Converse | 6 | grassland | R6WY-g-4 | 2014-09-22T00:00:00Z | 21 | 50 | 0 | R6WY-g-4 |
| Swainson's hawk | Raptor | summer | WY | Converse | 6 | grassland | R6WY-g-4 | 2014-09-22T00:00:00Z | 21 | 50 | 0 | R6WY-g-4 |
| ferruginous hawk | Raptor | spring | WY | Converse | 6 | grassland | R6WY-g-4 | 2015-04-13T00:00:00Z | 119 | NA | 0 | R6WY-g-4 |
| red-tailed hawk | Raptor | spring | WY | Converse | 6 | grassland | R6WY-g-4 | 2015-04-13T00:00:00Z | 119 | NA | 0 | R6WY-g-4 |
| ferruginous hawk | Raptor | spring | WY | Converse | 6 | grassland | R6WY-g-4 | 2015-04-13T00:00:00Z | 119 | NA | 0 | R6WY-g-4 |
| ferruginous hawk | Raptor | spring | WY | Converse | 6 | grassland | R6WY-g-4 | 2015-04-13T00:00:00Z | 119 | NA | 0 | R6WY-g-4 |
| red-tailed hawk | Raptor | spring | WY | Converse | 6 | grassland | R6WY-g-4 | 2015-05-11T00:00:00Z | 121 | NA | 0 | R6WY-g-4 |
| red-tailed hawk | Raptor | spring | WY | Converse | 6 | grassland | R6WY-g-4 | 2015-05-11T00:00:00Z | 121 | NA | 0 | R6WY-g-4 |
| red-tailed hawk | Raptor | spring | WY | Converse | 6 | grassland | R6WY-g-4 | 2015-05-11T00:00:00Z | 121 | NA | 0 | R6WY-g-4 |
| red-tailed hawk | Raptor | spring | WY | Converse | 6 | grassland | R6WY-g-4 | 2015-05-11T00:00:00Z | 121 | NA | 0 | R6WY-g-4 |
| rough-legged hawk | Raptor | spring | WY | Converse | 6 | grassland | R6WY-g-4 | 2015-05-11T00:00:00Z | 121 | NA | 0 | R6WY-g-4 |
| ring-necked pheasant | Game Bird | summer | KS | Sumner | 6 | cropland | R6KS-c-1 | 2016-08-15T00:00:00Z | 1 | 1 | 1 | R6KS-c-1 |
| ring-necked pheasant | Game Bird | summer | KS | Sumner | 6 | cropland | R6KS-c-1 | 2016-08-15T00:00:00Z | 2 | 3 | 1 | R6KS-c-1 |
| ring-necked pheasant | Game Bird | summer | KS | Sumner | 6 | cropland | R6KS-c-1 | 2016-08-15T00:00:00Z | 3 | 4 | 1 | R6KS-c-1 |
| ring-necked pheasant | Game Bird | summer | KS | Sumner | 6 | cropland | R6KS-c-1 | 2016-08-15T00:00:00Z | 1 | 1 | 1 | R6KS-c-1 |
| ring-necked pheasant | Game Bird | summer | KS | Sumner | 6 | cropland | R6KS-c-1 | 2016-08-15T00:00:00Z | 10 | 18 | 1 | R6KS-c-1 |
| ring-necked pheasant | Game Bird | summer | KS | Sumner | 6 | cropland | R6KS-c-1 | 2016-08-16T00:00:00Z | 3 | 6 | 1 | R6KS-c-1 |
| ring-necked pheasant | Game Bird | summer | KS | Sumner | 6 | cropland | R6KS-c-1 | 2016-08-16T00:00:00Z | 1 | 2 | 1 | R6KS-c-1 |
| ring-necked pheasant | Game Bird | summer | KS | Sumner | 6 | cropland | R6KS-c-1 | 2016-08-22T00:00:00Z | 2 | 3 | 1 | R6KS-c-1 |
| ring-necked pheasant | Game Bird | summer | KS | Sumner | 6 | cropland | R6KS-c-1 | 2016-08-22T00:00:00Z | 4 | 7 | 1 | R6KS-c-1 |
| ring-necked pheasant | Game Bird | summer | KS | Sumner | 6 | cropland | R6KS-c-1 | 2016-08-22T00:00:00Z | 3 | 4 | 1 | R6KS-c-1 |
| ring-necked pheasant | Game Bird | summer | KS | Sumner | 6 | cropland | R6KS-c-1 | 2016-08-22T00:00:00Z | 1 | 1 | 1 | R6KS-c-1 |
| ring-necked pheasant | Game Bird | summer | KS | Sumner | 6 | cropland | R6KS-c-1 | 2016-08-22T00:00:00Z | 10 | 21 | 1 | R6KS-c-1 |
| ring-necked pheasant | Game Bird | summer | KS | Sumner | 6 | cropland | R6KS-c-1 | 2016-08-22T00:00:00Z | 2 | 7 | 1 | R6KS-c-1 |
| ring-necked pheasant | Game Bird | summer | KS | Sumner | 6 | cropland | R6KS-c-1 | 2016-08-22T00:00:00Z | 2 | 7 | 1 | R6KS-c-1 |
| ring-necked pheasant | Game Bird | summer | KS | Sumner | 6 | cropland | R6KS-c-1 | 2016-08-22T00:00:00Z | 30 | NA | 1 | R6KS-c-1 |
| ring-necked pheasant | Game Bird | fall | KS | Sumner | 6 | cropland | R6KS-c-1 | 2016-10-10T00:00:00Z | 3 | 4 | 1 | R6KS-c-1 |
| ring-necked pheasant | Game Bird | fall | KS | Sumner | 6 | cropland | R6KS-c-1 | 2016-10-10T00:00:00Z | 4 | 7 | 1 | R6KS-c-1 |
| ring-necked pheasant | Game Bird | fall | KS | Sumner | 6 | cropland | R6KS-c-1 | 2016-10-10T00:00:00Z | 4 | 8 | 1 | R6KS-c-1 |
| ring-necked pheasant | Game Bird | fall | KS | Sumner | 6 | cropland | R6KS-c-1 | 2016-10-10T00:00:00Z | 4 | 7 | 1 | R6KS-c-1 |
| ring-necked pheasant | Game Bird | fall | KS | Sumner | 6 | cropland | R6KS-c-1 | 2016-10-10T00:00:00Z | 2 | 3 | 1 | R6KS-c-1 |
| ring-necked pheasant | Game Bird | fall | KS | Sumner | 6 | cropland | R6KS-c-1 | 2016-10-10T00:00:00Z | 10 | 21 | 1 | R6KS-c-1 |
| ring-necked pheasant | Game Bird | fall | KS | Sumner | 6 | cropland | R6KS-c-1 | 2016-10-10T00:00:00Z | 1 | 2 | 1 | R6KS-c-1 |
| ring-necked pheasant | Game Bird | fall | KS | Sumner | 6 | cropland | R6KS-c-1 | 2016-10-10T00:00:00Z | 1 | 2 | 1 | R6KS-c-1 |
| ring-necked pheasant | Game Bird | fall | KS | Sumner | 6 | cropland | R6KS-c-1 | 2016-10-10T00:00:00Z | 7 | 11 | 1 | R6KS-c-1 |
| ring-necked pheasant | Game Bird | fall | KS | Sumner | 6 | cropland | R6KS-c-1 | 2016-10-10T00:00:00Z | 7 | 10 | 1 | R6KS-c-1 |
| ring-necked pheasant | Game Bird | fall | KS | Sumner | 6 | cropland | R6KS-c-1 | 2016-10-17T00:00:00Z | 7 | 10 | 1 | R6KS-c-1 |
| ring-necked pheasant | Game Bird | fall | KS | Sumner | 6 | cropland | R6KS-c-1 | 2016-10-17T00:00:00Z | 7 | 10 | 1 | R6KS-c-1 |
| ring-necked pheasant | Game Bird | fall | KS | Sumner | 6 | cropland | R6KS-c-1 | 2016-10-17T00:00:00Z | 7 | 10 | 1 | R6KS-c-1 |
| ring-necked pheasant | Game Bird | fall | KS | Sumner | 6 | cropland | R6KS-c-1 | 2016-10-17T00:00:00Z | 4 | 7 | 1 | R6KS-c-1 |
| ring-necked pheasant | Game Bird | fall | KS | Sumner | 6 | cropland | R6KS-c-1 | 2016-10-17T00:00:00Z | 1 | 1 | 1 | R6KS-c-1 |
| turkey vulture | Raptor | fall | KS | Sumner | 6 | cropland | R6KS-c-1 | 2016-10-20T00:00:00Z | 82 | NA | 1 | R6KS-c-1 |
| turkey vulture | Raptor | winter | KS | Sumner | 6 | cropland | R6KS-c-1 | 2016-11-28T00:00:00Z | 50 | 56 | 1 | R6KS-c-1 |
| northern harrier | Raptor | winter | KS | Sumner | 6 | cropland | R6KS-c-1 | 2017-01-05T00:00:00Z | 5 | 20 | 1 | R6KS-c-1 |
| red-tailed hawk | Raptor | winter | KS | Sumner | 6 | cropland | R6KS-c-1 | 2017-02-07T00:00:00Z | 16 | 23 | 1 | R6KS-c-1 |
| ring-necked pheasant | Game Bird | winter | KS | Sumner | 6 | cropland | R6KS-c-1 | 2016-11-28T00:00:00Z | 4 | 7 | 1 | R6KS-c-1 |
| ring-necked pheasant | Game Bird | winter | KS | Sumner | 6 | cropland | R6KS-c-1 | 2016-11-28T00:00:00Z | 2 | 3 | 1 | R6KS-c-1 |
| ring-necked pheasant | Game Bird | winter | KS | Sumner | 6 | cropland | R6KS-c-1 | 2017-01-09T00:00:00Z | 4 | 7 | 1 | R6KS-c-1 |
| ring-necked pheasant | Game Bird | winter | KS | Sumner | 6 | cropland | R6KS-c-1 | 2017-01-09T00:00:00Z | 3 | 4 | 1 | R6KS-c-1 |
| ring-necked pheasant | Game Bird | winter | KS | Sumner | 6 | cropland | R6KS-c-1 | 2017-01-09T00:00:00Z | 4 | 7 | 1 | R6KS-c-1 |
| ring-necked pheasant | Game Bird | winter | KS | Sumner | 6 | cropland | R6KS-c-1 | 2017-01-09T00:00:00Z | 7 | 10 | 1 | R6KS-c-1 |
| ring-necked pheasant | Game Bird | winter | KS | Sumner | 6 | cropland | R6KS-c-1 | 2017-01-09T00:00:00Z | 10 | 21 | 1 | R6KS-c-1 |
| ring-necked pheasant | Game Bird | winter | KS | Sumner | 6 | cropland | R6KS-c-1 | 2017-01-23T00:00:00Z | 4 | 7 | 1 | R6KS-c-1 |
| ring-necked pheasant | Game Bird | winter | KS | Sumner | 6 | cropland | R6KS-c-1 | 2017-01-23T00:00:00Z | 1 | 1 | 1 | R6KS-c-1 |
| ring-necked pheasant | Game Bird | winter | KS | Sumner | 6 | cropland | R6KS-c-1 | 2017-01-23T00:00:00Z | 30 | NA | 1 | R6KS-c-1 |
| ring-necked pheasant | Game Bird | winter | KS | Sumner | 6 | cropland | R6KS-c-1 | 2017-01-23T00:00:00Z | 21 | 30 | 1 | R6KS-c-1 |
| ring-necked pheasant | Game Bird | winter | KS | Sumner | 6 | cropland | R6KS-c-1 | 2017-01-23T00:00:00Z | 2 | 3 | 1 | R6KS-c-1 |
| ring-necked pheasant | Game Bird | winter | KS | Sumner | 6 | cropland | R6KS-c-1 | 2017-01-23T00:00:00Z | 1 | 2 | 1 | R6KS-c-1 |
| ring-necked pheasant | Game Bird | winter | KS | Sumner | 6 | cropland | R6KS-c-1 | 2017-01-23T00:00:00Z | 10 | 21 | 1 | R6KS-c-1 |
| ring-necked pheasant | Game Bird | winter | KS | Sumner | 6 | cropland | R6KS-c-1 | 2017-01-23T00:00:00Z | 10 | 21 | 1 | R6KS-c-1 |
| ring-necked pheasant | Game Bird | spring | KS | Sumner | 6 | cropland | R6KS-c-1 | 2017-03-20T00:00:00Z | 30 | NA | 1 | R6KS-c-1 |
| ring-necked pheasant | Game Bird | spring | KS | Sumner | 6 | cropland | R6KS-c-1 | 2017-03-20T00:00:00Z | 2 | 3 | 1 | R6KS-c-1 |
| ring-necked pheasant | Game Bird | spring | KS | Sumner | 6 | cropland | R6KS-c-1 | 2017-03-20T00:00:00Z | 4 | 7 | 1 | R6KS-c-1 |
| ring-necked pheasant | Game Bird | spring | KS | Sumner | 6 | cropland | R6KS-c-1 | 2017-03-20T00:00:00Z | 30 | NA | 1 | R6KS-c-1 |
| ring-necked pheasant | Game Bird | spring | KS | Sumner | 6 | cropland | R6KS-c-1 | 2017-03-27T00:00:00Z | 4 | 10 | 1 | R6KS-c-1 |
| ring-necked pheasant | Game Bird | spring | KS | Sumner | 6 | cropland | R6KS-c-1 | 2017-03-27T00:00:00Z | 4 | 10 | 1 | R6KS-c-1 |
| ring-necked pheasant | Game Bird | spring | KS | Sumner | 6 | cropland | R6KS-c-1 | 2017-03-27T00:00:00Z | 7 | 10 | 1 | R6KS-c-1 |
| ring-necked pheasant | Game Bird | spring | KS | Sumner | 6 | cropland | R6KS-c-1 | 2017-03-27T00:00:00Z | 7 | 10 | 1 | R6KS-c-1 |
| ring-necked pheasant | Game Bird | spring | KS | Sumner | 6 | cropland | R6KS-c-1 | 2017-03-27T00:00:00Z | 1 | 2 | 1 | R6KS-c-1 |
| ring-necked pheasant | Game Bird | spring | KS | Sumner | 6 | cropland | R6KS-c-1 | 2017-03-27T00:00:00Z | 2 | 4 | 1 | R6KS-c-1 |
| ring-necked pheasant | Game Bird | spring | KS | Sumner | 6 | cropland | R6KS-c-1 | 2017-04-03T00:00:00Z | 10 | 21 | 1 | R6KS-c-1 |
| ring-necked pheasant | Game Bird | spring | KS | Sumner | 6 | cropland | R6KS-c-1 | 2017-04-03T00:00:00Z | 4 | 7 | 1 | R6KS-c-1 |
| ring-necked pheasant | Game Bird | spring | KS | Sumner | 6 | cropland | R6KS-c-1 | 2017-04-03T00:00:00Z | 10 | 21 | 1 | R6KS-c-1 |
| ring-necked pheasant | Game Bird | spring | KS | Sumner | 6 | cropland | R6KS-c-1 | 2017-04-03T00:00:00Z | 2 | 3 | 1 | R6KS-c-1 |
| ring-necked pheasant | Game Bird | spring | KS | Sumner | 6 | cropland | R6KS-c-1 | 2017-04-03T00:00:00Z | 3 | 4 | 1 | R6KS-c-1 |
| turkey vulture | Raptor | spring | KS | Sumner | 6 | cropland | R6KS-c-1 | 2017-05-02T00:00:00Z | 28 | NA | 1 | R6KS-c-1 |
| Cooper's hawk | Raptor | winter | TX | Randall and Castro | 2 | grassland | R2TX-g-1 | 2018-02-12T00:00:00Z | 4.1569444 | 7.1680556 | 0 | R2TX-g-1 |
| Swainson's hawk | Raptor | winter | TX | Randall and Castro | 2 | grassland | R2TX-g-1 | 2018-02-12T00:00:00Z | 63.8520833 | NA | 0 | R2TX-g-1 |
| osprey | Raptor | winter | TX | Randall and Castro | 2 | grassland | R2TX-g-1 | 2018-02-12T00:00:00Z | 63.86875 | NA | 0 | R2TX-g-1 |
| barn owl | Raptor | winter | TX | Randall and Castro | 2 | grassland | R2TX-g-1 | 2018-02-12T00:00:00Z | 63.89375 | NA | 0 | R2TX-g-1 |
| great horned owl | Raptor | winter | TX | Randall and Castro | 2 | grassland | R2TX-g-1 | 2018-02-12T00:00:00Z | 63.9194444 | NA | 0 | R2TX-g-1 |
| great horned owl | Raptor | winter | TX | Randall and Castro | 2 | grassland | R2TX-g-1 | 2018-02-12T00:00:00Z | 63.9506944 | NA | 0 | R2TX-g-1 |
| red-tailed hawk | Raptor | winter | TX | Randall and Castro | 2 | grassland | R2TX-g-1 | 2018-02-12T00:00:00Z | 63.9798611 | NA | 0 | R2TX-g-1 |
| barn owl | Raptor | winter | TX | Randall and Castro | 2 | grassland | R2TX-g-1 | 2018-02-12T00:00:00Z | 63.7569444 | NA | 0 | R2TX-g-1 |
| Swainson's hawk | Raptor | winter | TX | Randall and Castro | 2 | grassland | R2TX-g-1 | 2018-02-12T00:00:00Z | 63.8 | NA | 0 | R2TX-g-1 |
| great horned owl | Raptor | winter | TX | Randall and Castro | 2 | grassland | R2TX-g-1 | 2018-02-12T00:00:00Z | 63.8333333 | NA | 0 | R2TX-g-1 |
| barn owl | Raptor | winter | TX | Randall and Castro | 2 | grassland | R2TX-g-1 | 2018-02-12T00:00:00Z | 63.8729167 | NA | 0 | R2TX-g-1 |
| red-tailed hawk | Raptor | winter | TX | Randall and Castro | 2 | grassland | R2TX-g-1 | 2018-02-12T00:00:00Z | 63.8694444 | NA | 0 | R2TX-g-1 |
| red-tailed hawk | Raptor | winter | TX | Randall and Castro | 2 | grassland | R2TX-g-1 | 2018-02-12T00:00:00Z | 63.9173611 | NA | 0 | R2TX-g-1 |
| great horned owl | Raptor | winter | TX | Randall and Castro | 2 | grassland | R2TX-g-1 | 2018-02-12T00:00:00Z | 63.9506944 | NA | 0 | R2TX-g-1 |
| great horned owl | Raptor | winter | TX | Randall and Castro | 2 | grassland | R2TX-g-1 | 2018-02-12T00:00:00Z | 63.9715278 | NA | 0 | R2TX-g-1 |
| Swainson's hawk | Raptor | fall | TX | Randall and Castro | 2 | grassland | R2TX-g-2 | 2018-10-04T00:00:00Z | 63.2569444 | NA | 0 | R2TX-g-2 |
| red-tailed hawk | Raptor | fall | TX | Randall and Castro | 2 | grassland | R2TX-g-2 | 2018-10-04T00:00:00Z | 21.2694444 | 28.2590278 | 0 | R2TX-g-2 |
| great horned owl | Raptor | fall | TX | Randall and Castro | 2 | grassland | R2TX-g-2 | 2018-10-04T00:00:00Z | 63.2104167 | NA | 0 | R2TX-g-2 |
| red-tailed hawk | Raptor | fall | TX | Randall and Castro | 2 | grassland | R2TX-g-2 | 2018-10-04T00:00:00Z | 14.2222222 | 20.8402778 | 0 | R2TX-g-2 |
| osprey | Raptor | fall | TX | Randall and Castro | 2 | grassland | R2TX-g-2 | 2018-10-04T00:00:00Z | 62.1819444 | NA | 0 | R2TX-g-2 |
| red-tailed hawk | Raptor | fall | TX | Randall and Castro | 2 | grassland | R2TX-g-2 | 2018-10-04T00:00:00Z | 3.0215278 | 6.7881944 | 0 | R2TX-g-2 |
| red-tailed hawk | Raptor | fall | TX | Randall and Castro | 2 | grassland | R2TX-g-2 | 2018-10-04T00:00:00Z | 2.9861111 | 6.7965278 | 0 | R2TX-g-2 |
| red-tailed hawk | Raptor | fall | TX | Randall and Castro | 2 | grassland | R2TX-g-2 | 2018-10-04T00:00:00Z | 63.1284722 | NA | 0 | R2TX-g-2 |
| red-tailed hawk | Raptor | fall | TX | Randall and Castro | 2 | grassland | R2TX-g-2 | 2018-10-04T00:00:00Z | 63.1215278 | NA | 0 | R2TX-g-2 |
| red-tailed hawk | Raptor | fall | TX | Randall and Castro | 2 | grassland | R2TX-g-2 | 2018-10-04T00:00:00Z | 0.9152778 | 2.0791667 | 0 | R2TX-g-2 |
| red-tailed hawk | Raptor | winter | TX | Randall and Castro | 2 | grassland | R2TX-g-2 | 2018-12-10T00:00:00Z | 2.9201389 | 3.8763889 | 0 | R2TX-g-2 |
| Swainson's hawk | Raptor | winter | TX | Randall and Castro | 2 | grassland | R2TX-g-2 | 2018-12-10T00:00:00Z | 10.0152778 | 14.1875 | 0 | R2TX-g-2 |
| red-tailed hawk | Raptor | winter | TX | Randall and Castro | 2 | grassland | R2TX-g-2 | 2018-12-10T00:00:00Z | 62.8 | NA | 0 | R2TX-g-2 |
| Swainson's hawk | Raptor | winter | TX | Randall and Castro | 2 | grassland | R2TX-g-2 | 2018-12-10T00:00:00Z | 62.7756944 | NA | 0 | R2TX-g-2 |
| Swainson's hawk | Raptor | winter | TX | Randall and Castro | 2 | grassland | R2TX-g-2 | 2018-12-10T00:00:00Z | 62.7854167 | NA | 0 | R2TX-g-2 |
| red-tailed hawk | Raptor | winter | TX | Randall and Castro | 2 | grassland | R2TX-g-2 | 2018-12-10T00:00:00Z | 63.1277778 | NA | 0 | R2TX-g-2 |
| red-tailed hawk | Raptor | winter | TX | Randall and Castro | 2 | grassland | R2TX-g-2 | 2018-12-10T00:00:00Z | 21.1076389 | 28.1006944 | 0 | R2TX-g-2 |
| red-tailed hawk | Raptor | winter | TX | Randall and Castro | 2 | grassland | R2TX-g-2 | 2018-12-10T00:00:00Z | 42.0881944 | 49.0958333 | 0 | R2TX-g-2 |
| red-tailed hawk | Raptor | winter | TX | Randall and Castro | 2 | grassland | R2TX-g-2 | 2018-12-10T00:00:00Z | 10.0763889 | 14.0909722 | 0 | R2TX-g-2 |
| red-tailed hawk | Raptor | winter | TX | Randall and Castro | 2 | grassland | R2TX-g-2 | 2018-12-10T00:00:00Z | 63.0784722 | NA | 0 | R2TX-g-2 |
| ring-necked pheasant | Game Bird | fall | OK | Kay | 2 | grassland | R2OK-g-1 | 2017-09-21T00:00:00Z | 0.9340278 | 0.9340278 | 1 | R2OK-g-0 |
| ring-necked pheasant | Game Bird | fall | OK | Kay | 2 | grassland | R2OK-g-1 | 2017-09-21T00:00:00Z | 0.9319444 | 0.9319444 | 1 | R2OK-g-0 |
| ring-necked pheasant | Game Bird | fall | OK | Kay | 2 | grassland | R2OK-g-1 | 2017-09-21T00:00:00Z | 2.0277778 | 2.9159722 | 1 | R2OK-g-0 |
| ring-necked pheasant | Game Bird | fall | OK | Kay | 2 | grassland | R2OK-g-1 | 2017-09-21T00:00:00Z | 0.93125 | 2.025 | 1 | R2OK-g-0 |
| ring-necked pheasant | Game Bird | fall | OK | Kay | 2 | grassland | R2OK-g-1 | 2017-09-21T00:00:00Z | 0.93125 | 2.025 | 1 | R2OK-g-0 |
| ring-necked pheasant | Game Bird | fall | OK | Kay | 2 | grassland | R2OK-g-1 | 2017-10-09T00:00:00Z | 3.8222222 | 6.8708333 | 1 | R2OK-g-0 |
| ring-necked pheasant | Game Bird | fall | OK | Kay | 2 | grassland | R2OK-g-1 | 2017-09-21T00:00:00Z | 0.9722222 | 0.9722222 | 1 | R2OK-g-0 |
| ring-necked pheasant | Game Bird | fall | OK | Kay | 2 | grassland | R2OK-g-1 | 2017-09-21T00:00:00Z | 2.0638889 | 2.9597222 | 1 | R2OK-g-0 |
| ring-necked pheasant | Game Bird | fall | OK | Kay | 2 | grassland | R2OK-g-1 | 2017-09-21T00:00:00Z | 0.9465278 | 0.9465278 | 1 | R2OK-g-0 |
| ring-necked pheasant | Game Bird | fall | OK | Kay | 2 | grassland | R2OK-g-1 | 2017-09-21T00:00:00Z | 0.9465278 | 0.9465278 | 1 | R2OK-g-0 |
| ring-necked pheasant | Game Bird | fall | OK | Kay | 2 | grassland | R2OK-g-1 | 2017-09-21T00:00:00Z | 0.9416667 | 0.9416667 | 1 | R2OK-g-0 |
| ring-necked pheasant | Game Bird | fall | OK | Kay | 2 | grassland | R2OK-g-1 | 2017-09-21T00:00:00Z | 0.9444444 | 0.9444444 | 1 | R2OK-g-0 |
| ring-necked pheasant | Game Bird | fall | OK | Kay | 2 | grassland | R2OK-g-1 | 2017-09-21T00:00:00Z | 0.8506944 | 2.0388889 | 1 | R2OK-g-0 |
| ring-necked pheasant | Game Bird | fall | OK | Kay | 2 | grassland | R2OK-g-1 | 2017-09-21T00:00:00Z | 0.9368056 | 0.9368056 | 1 | R2OK-g-0 |
| ring-necked pheasant | Game Bird | fall | OK | Kay | 2 | grassland | R2OK-g-1 | 2017-09-21T00:00:00Z | 0.9354167 | 2.0368056 | 1 | R2OK-g-0 |
| ring-necked pheasant | Game Bird | fall | OK | Kay | 2 | grassland | R2OK-g-1 | 2017-10-09T00:00:00Z | 1.8645833 | 2.96875 | 1 | R2OK-g-0 |
| ring-necked pheasant | Game Bird | fall | OK | Kay | 2 | grassland | R2OK-g-1 | 2017-10-09T00:00:00Z | 0.8444444 | 1.8347222 | 1 | R2OK-g-0 |
| ring-necked pheasant | Game Bird | fall | OK | Kay | 2 | grassland | R2OK-g-1 | 2017-10-09T00:00:00Z | 0.7729167 | 0.7729167 | 1 | R2OK-g-0 |
| ring-necked pheasant | Game Bird | fall | OK | Kay | 2 | grassland | R2OK-g-1 | 2017-10-09T00:00:00Z | 3.8409722 | 6.8597222 | 1 | R2OK-g-0 |
| ring-necked pheasant | Game Bird | fall | OK | Kay | 2 | grassland | R2OK-g-1 | 2017-10-09T00:00:00Z | 3.7701389 | 6.9020833 | 1 | R2OK-g-0 |
| ring-necked pheasant | Game Bird | winter | OK | Kay | 2 | grassland | R2OK-g-1 | 2017-11-27T00:00:00Z | 4.0152778 | 6.9611111 | 1 | R2OK-g-0 |
| ring-necked pheasant | Game Bird | winter | OK | Kay | 2 | grassland | R2OK-g-1 | 2017-11-27T00:00:00Z | 1.0284722 | 2.0131944 | 1 | R2OK-g-0 |
| ring-necked pheasant | Game Bird | winter | OK | Kay | 2 | grassland | R2OK-g-1 | 2017-11-27T00:00:00Z | 1.0284722 | 1.0284722 | 1 | R2OK-g-0 |
| ring-necked pheasant | Game Bird | winter | OK | Kay | 2 | grassland | R2OK-g-1 | 2017-11-27T00:00:00Z | 3.9888889 | 6.9409722 | 1 | R2OK-g-0 |
| ring-necked pheasant | Game Bird | winter | OK | Kay | 2 | grassland | R2OK-g-1 | 2017-11-27T00:00:00Z | 6.9388889 | 9.9270833 | 1 | R2OK-g-0 |
| ring-necked pheasant | Game Bird | winter | OK | Kay | 2 | grassland | R2OK-g-1 | 2017-11-27T00:00:00Z | 1.0236111 | 1.9993056 | 1 | R2OK-g-0 |
| ring-necked pheasant | Game Bird | winter | OK | Kay | 2 | grassland | R2OK-g-1 | 2017-11-27T00:00:00Z | 1.0236111 | 2.0041667 | 1 | R2OK-g-0 |
| ring-necked pheasant | Game Bird | winter | OK | Kay | 2 | grassland | R2OK-g-1 | 2017-11-27T00:00:00Z | 9.9298611 | 13.925 | 1 | R2OK-g-0 |
| ring-necked pheasant | Game Bird | winter | OK | Kay | 2 | grassland | R2OK-g-1 | 2017-11-27T00:00:00Z | 1.025 | 2.0048611 | 1 | R2OK-g-0 |
| ring-necked pheasant | Game Bird | winter | OK | Kay | 2 | grassland | R2OK-g-1 | 2017-11-27T00:00:00Z | 13.9104167 | 20.8888889 | 1 | R2OK-g-0 |
| mallard | Game Bird | winter | OK | Kay | 2 | grassland | R2OK-g-1 | 2018-01-08T00:00:00Z | 3.9680556 | 9.9833333 | 1 | R2OK-g-0 |
| mallard | Game Bird | winter | OK | Kay | 2 | grassland | R2OK-g-1 | 2018-01-08T00:00:00Z | 13.9611111 | 20.9520833 | 1 | R2OK-g-0 |
| mallard | Game Bird | winter | OK | Kay | 2 | grassland | R2OK-g-1 | 2018-01-08T00:00:00Z | 0.9888889 | 0.9888889 | 1 | R2OK-g-0 |
| mallard | Game Bird | winter | OK | Kay | 2 | grassland | R2OK-g-1 | 2018-01-08T00:00:00Z | 1.9916667 | 3.0090278 | 1 | R2OK-g-0 |
| mallard | Game Bird | winter | OK | Kay | 2 | grassland | R2OK-g-1 | 2018-01-08T00:00:00Z | 3.9715278 | 9.9847222 | 1 | R2OK-g-0 |
| mallard | Game Bird | winter | OK | Kay | 2 | grassland | R2OK-g-1 | 2018-01-08T00:00:00Z | 2.9993056 | 3.9652778 | 1 | R2OK-g-0 |
| mallard | Game Bird | winter | OK | Kay | 2 | grassland | R2OK-g-1 | 2018-01-08T00:00:00Z | 0.9819444 | 0.9819444 | 1 | R2OK-g-0 |
| mallard | Game Bird | winter | OK | Kay | 2 | grassland | R2OK-g-1 | 2018-01-08T00:00:00Z | 0.9791667 | 1.9729167 | 1 | R2OK-g-0 |
| mallard | Game Bird | winter | OK | Kay | 2 | grassland | R2OK-g-1 | 2018-01-08T00:00:00Z | 1.9701389 | 2.9833333 | 1 | R2OK-g-0 |
| mallard | Game Bird | winter | OK | Kay | 2 | grassland | R2OK-g-1 | 2018-01-08T00:00:00Z | 3.9388889 | 9.9486111 | 1 | R2OK-g-0 |
| mallard | Game Bird | spring | OK | Kay | 2 | grassland | R2OK-g-1 | 2018-03-05T00:00:00Z | 0.9868056 | 0.9868056 | 1 | R2OK-g-0 |
| ring-necked pheasant | Game Bird | spring | OK | Kay | 2 | grassland | R2OK-g-1 | 2018-03-05T00:00:00Z | 3.9763889 | 6.9590278 | 1 | R2OK-g-0 |
| ring-necked pheasant | Game Bird | spring | OK | Kay | 2 | grassland | R2OK-g-1 | 2018-03-05T00:00:00Z | 0.9819444 | 0.9819444 | 1 | R2OK-g-0 |
| ring-necked pheasant | Game Bird | spring | OK | Kay | 2 | grassland | R2OK-g-1 | 2018-03-05T00:00:00Z | 0.9791667 | 1.9854167 | 1 | R2OK-g-0 |
| ring-necked pheasant | Game Bird | spring | OK | Kay | 2 | grassland | R2OK-g-1 | 2018-03-05T00:00:00Z | 2.9625 | 3.9513889 | 1 | R2OK-g-0 |
| ring-necked pheasant | Game Bird | spring | OK | Kay | 2 | grassland | R2OK-g-1 | 2018-03-05T00:00:00Z | 6.9229167 | 9.9041667 | 1 | R2OK-g-0 |
| ring-necked pheasant | Game Bird | spring | OK | Kay | 2 | grassland | R2OK-g-1 | 2018-03-05T00:00:00Z | 0.9722222 | 0.9722222 | 1 | R2OK-g-0 |
| ring-necked pheasant | Game Bird | spring | OK | Kay | 2 | grassland | R2OK-g-1 | 2018-03-05T00:00:00Z | 13.8819444 | 21.0694444 | 1 | R2OK-g-0 |
| ring-necked pheasant | Game Bird | spring | OK | Kay | 2 | grassland | R2OK-g-1 | 2018-03-05T00:00:00Z | 6.9097222 | 9.9145833 | 1 | R2OK-g-0 |
| ring-necked pheasant | Game Bird | spring | OK | Kay | 2 | grassland | R2OK-g-1 | 2018-03-05T00:00:00Z | 2.9375 | 3.9270833 | 1 | R2OK-g-0 |
| mallard | Game Bird | spring | OK | Kay | 2 | grassland | R2OK-g-1 | 2018-04-30T00:00:00Z | 0.8409722 | 0.8409722 | 1 | R2OK-g-0 |
| mallard | Game Bird | spring | OK | Kay | 2 | grassland | R2OK-g-1 | 2018-04-30T00:00:00Z | 0.8652778 | 0.8652778 | 1 | R2OK-g-0 |
| Cooper's hawk | Raptor | spring | OK | Kay | 2 | grassland | R2OK-g-1 | 2018-04-30T00:00:00Z | 2.8493056 | 3.7451389 | 1 | R2OK-g-0 |
| mallard | Game Bird | spring | OK | Kay | 2 | grassland | R2OK-g-1 | 2018-04-30T00:00:00Z | 0.8715278 | 0.8715278 | 1 | R2OK-g-0 |
| barn owl | Raptor | spring | OK | Kay | 2 | grassland | R2OK-g-1 | 2018-04-30T00:00:00Z | 6.7263889 | 9.7458333 | 1 | R2OK-g-0 |
| Cooper's hawk | Raptor | spring | OK | Kay | 2 | grassland | R2OK-g-1 | 2018-04-30T00:00:00Z | 9.825 | 13.8604167 | 1 | R2OK-g-0 |
| red-tailed hawk | Raptor | spring | OK | Kay | 2 | grassland | R2OK-g-1 | 2018-04-30T00:00:00Z | 9.7194444 | 13.7493056 | 1 | R2OK-g-0 |
| red-tailed hawk | Raptor | spring | OK | Kay | 2 | grassland | R2OK-g-1 | 2018-04-30T00:00:00Z | 20.7798611 | 29.8881944 | 1 | R2OK-g-0 |
| red-tailed hawk | Raptor | spring | OK | Kay | 2 | grassland | R2OK-g-1 | 2018-04-30T00:00:00Z | 29.9159722 | NA | 1 | R2OK-g-0 |
| ring-necked pheasant | Game Bird | summer | OK | Kay | 2 | grassland | R2OK-g-1 | 2018-06-11T00:00:00Z | 1.3 | 2.2215278 | 1 | R2OK-g-0 |
| ring-necked pheasant | Game Bird | summer | OK | Kay | 2 | grassland | R2OK-g-1 | 2018-06-11T00:00:00Z | 1.2888889 | 2.2263889 | 1 | R2OK-g-0 |
| ring-necked pheasant | Game Bird | summer | OK | Kay | 2 | grassland | R2OK-g-1 | 2018-06-11T00:00:00Z | 2.0138889 | 3.2111111 | 1 | R2OK-g-0 |
| ring-necked pheasant | Game Bird | summer | OK | Kay | 2 | grassland | R2OK-g-1 | 2018-06-11T00:00:00Z | 2.0395833 | 3.1777778 | 1 | R2OK-g-0 |
| ring-necked pheasant | Game Bird | summer | OK | Kay | 2 | grassland | R2OK-g-1 | 2018-06-11T00:00:00Z | 1.1576389 | 2.1777778 | 1 | R2OK-g-0 |
| ring-necked pheasant | Game Bird | summer | OK | Kay | 2 | grassland | R2OK-g-1 | 2018-06-11T00:00:00Z | 3.0284722 | 4.0194444 | 1 | R2OK-g-0 |
| ring-necked pheasant | Game Bird | summer | OK | Kay | 2 | grassland | R2OK-g-1 | 2018-06-11T00:00:00Z | 3.025 | 4.0166667 | 1 | R2OK-g-0 |
| ring-necked pheasant | Game Bird | summer | OK | Kay | 2 | grassland | R2OK-g-1 | 2018-06-11T00:00:00Z | 0.9590278 | 0.9590278 | 1 | R2OK-g-0 |
| ring-necked pheasant | Game Bird | summer | OK | Kay | 2 | grassland | R2OK-g-1 | 2018-06-11T00:00:00Z | 2.9986111 | 3.875 | 1 | R2OK-g-0 |
| ring-necked pheasant | Game Bird | summer | OK | Kay | 2 | grassland | R2OK-g-1 | 2018-07-16T00:00:00Z | 2.2680556 | 3.0270833 | 1 | R2OK-g-0 |
| ring-necked pheasant | Game Bird | summer | OK | Kay | 2 | grassland | R2OK-g-1 | 2018-07-16T00:00:00Z | 2.2638889 | 3.0194444 | 1 | R2OK-g-0 |
| ring-necked pheasant | Game Bird | summer | OK | Kay | 2 | grassland | R2OK-g-1 | 2018-07-16T00:00:00Z | 1.1826389 | 1.1826389 | 1 | R2OK-g-0 |
| ring-necked pheasant | Game Bird | summer | OK | Kay | 2 | grassland | R2OK-g-1 | 2018-07-16T00:00:00Z | 0.925 | 1.9909722 | 1 | R2OK-g-0 |
| ring-necked pheasant | Game Bird | summer | OK | Kay | 2 | grassland | R2OK-g-1 | 2018-07-16T00:00:00Z | 0.9090278 | 1.9541667 | 1 | R2OK-g-0 |
| ring-necked pheasant | Game Bird | summer | OK | Kay | 2 | grassland | R2OK-g-1 | 2018-07-16T00:00:00Z | 1.9277778 | 2.9805556 | 1 | R2OK-g-0 |
| ring-necked pheasant | Game Bird | summer | OK | Kay | 2 | grassland | R2OK-g-1 | 2018-07-16T00:00:00Z | 3.7270833 | 6.9326389 | 1 | R2OK-g-0 |
| ring-necked pheasant | Game Bird | summer | OK | Kay | 2 | grassland | R2OK-g-1 | 2018-07-16T00:00:00Z | 0.7159722 | 0.7159722 | 1 | R2OK-g-0 |
| ring-necked pheasant | Game Bird | summer | OK | Kay | 2 | grassland | R2OK-g-1 | 2018-07-16T00:00:00Z | 1.6548611 | 2.9472222 | 1 | R2OK-g-0 |
| ring-necked pheasant | Game Bird | summer | OK | Kay | 2 | grassland | R2OK-g-1 | 2018-07-16T00:00:00Z | 1.6527778 | 2.9444444 | 1 | R2OK-g-0 |
| ring-necked pheasant | Game Bird | summer | OK | Kay | 2 | grassland | R2OK-g-1 | 2018-07-16T00:00:00Z | 2.9458333 | 3.6333333 | 1 | R2OK-g-0 |
| ring-necked pheasant | Game Bird | spring | OK | Garfield, Kay, Noble | 2 | grassland | R2OK-g-2 | 2018-04-30T00:00:00Z | 0.9611111 | 2.3118056 | 1 | R2OK-g-0 |
| ring-necked pheasant | Game Bird | spring | OK | Garfield, Kay, Noble | 2 | grassland | R2OK-g-2 | 2018-04-30T00:00:00Z | 2.2819444 | 3.2118056 | 1 | R2OK-g-0 |
| ring-necked pheasant | Game Bird | spring | OK | Garfield, Kay, Noble | 2 | grassland | R2OK-g-2 | 2018-04-30T00:00:00Z | 4.2270833 | 6.8902778 | 1 | R2OK-g-0 |
| ring-necked pheasant | Game Bird | spring | OK | Garfield, Kay, Noble | 2 | grassland | R2OK-g-2 | 2018-04-30T00:00:00Z | 3.1590278 | 4.2229167 | 1 | R2OK-g-0 |
| ring-necked pheasant | Game Bird | spring | OK | Garfield, Kay, Noble | 2 | grassland | R2OK-g-2 | 2018-04-30T00:00:00Z | 3.05 | 4.1458333 | 1 | R2OK-g-0 |
| Swainson's hawk | Raptor | winter | OK | Garfield, Kay, Noble | 2 | grassland | R2OK-g-3 | 2018-11-06T00:00:00Z | 14.0326389 | 21.0555556 | 1 | R2OK-g-0 |
| red-tailed hawk | Raptor | winter | OK | Garfield, Kay, Noble | 2 | grassland | R2OK-g-3 | 2018-11-06T00:00:00Z | 30.3 | 44.9930556 | 1 | R2OK-g-0 |
| Swainson's hawk | Raptor | winter | OK | Garfield, Kay, Noble | 2 | grassland | R2OK-g-3 | 2018-11-06T00:00:00Z | 13.9291667 | 21.025 | 1 | R2OK-g-0 |
| barn owl | Raptor | winter | OK | Garfield, Kay, Noble | 2 | grassland | R2OK-g-3 | 2018-11-06T00:00:00Z | 13.9618056 | 20.9458333 | 1 | R2OK-g-0 |
| Swainson's hawk | Raptor | winter | OK | Garfield, Kay, Noble | 2 | grassland | R2OK-g-3 | 2018-11-06T00:00:00Z | 6.9708333 | 13.9083333 | 1 | R2OK-g-0 |
| great horned owl | Raptor | winter | OK | Garfield, Kay, Noble | 2 | grassland | R2OK-g-3 | 2018-11-06T00:00:00Z | 6.8680556 | 13.8104167 | 1 | R2OK-g-0 |
| great horned owl | Raptor | winter | OK | Garfield, Kay, Noble | 2 | grassland | R2OK-g-3 | 2018-11-06T00:00:00Z | 20.7652778 | 29.7930556 | 1 | R2OK-g-0 |
| red-tailed hawk | Raptor | winter | OK | Garfield, Kay, Noble | 2 | grassland | R2OK-g-3 | 2018-11-06T00:00:00Z | 6.6625 | 6.6625 | 1 | R2OK-g-0 |
| red-tailed hawk | Raptor | winter | OK | Garfield, Kay, Noble | 2 | grassland | R2OK-g-3 | 2018-12-06T00:00:00Z | 22.0791667 | 30.2 | 1 | R2OK-g-0 |
| Swainson's hawk | Raptor | winter | OK | Garfield, Kay, Noble | 2 | grassland | R2OK-g-3 | 2018-12-06T00:00:00Z | 13.9166667 | 21.9604167 | 1 | R2OK-g-0 |
| red-tailed hawk | Raptor | winter | OK | Garfield, Kay, Noble | 2 | grassland | R2OK-g-3 | 2018-12-06T00:00:00Z | 6.9909722 | 6.9909722 | 1 | R2OK-g-0 |
| Swainson's hawk | Raptor | winter | OK | Garfield, Kay, Noble | 2 | grassland | R2OK-g-3 | 2018-12-06T00:00:00Z | 13.8944444 | 21.8909722 | 1 | R2OK-g-0 |
| red-tailed hawk | Raptor | winter | OK | Garfield, Kay, Noble | 2 | grassland | R2OK-g-3 | 2018-12-06T00:00:00Z | 21.7486111 | 29.9694444 | 1 | R2OK-g-0 |
| red-tailed hawk | Raptor | winter | OK | Garfield, Kay, Noble | 2 | grassland | R2OK-g-3 | 2018-12-03T00:00:00Z | 10.0708333 | 10.0708333 | 1 | R2OK-g-0 |
| red-tailed hawk | Raptor | winter | OK | Garfield, Kay, Noble | 2 | grassland | R2OK-g-3 | 2018-12-04T00:00:00Z | 24.0215278 | 32.04375 | 1 | R2OK-g-0 |
| red-tailed hawk | Raptor | winter | OK | Garfield, Kay, Noble | 2 | grassland | R2OK-g-3 | 2018-12-04T00:00:00Z | 47.0305556 | NA | 1 | R2OK-g-0 |
| northern harrier | Raptor | winter | OK | Garfield, Kay, Noble | 2 | grassland | R2OK-g-3 | 2018-12-04T00:00:00Z | 8.7659722 | 8.7659722 | 1 | R2OK-g-0 |
| Swainson's hawk | Raptor | winter | OK | Garfield, Kay, Noble | 2 | grassland | R2OK-g-3 | 2019-01-07T00:00:00Z | 7.2041667 | 7.2041667 | 1 | R2OK-g-0 |
| great horned owl | Raptor | winter | OK | Garfield, Kay, Noble | 2 | grassland | R2OK-g-3 | 2019-01-07T00:00:00Z | 14.0013889 | 20.9805556 | 1 | R2OK-g-0 |
| barn owl | Raptor | winter | OK | Garfield, Kay, Noble | 2 | grassland | R2OK-g-3 | 2019-01-07T00:00:00Z | 29.9395833 | 44.9555556 | 1 | R2OK-g-0 |
| Swainson's hawk | Raptor | winter | OK | Garfield, Kay, Noble | 2 | grassland | R2OK-g-3 | 2019-01-07T00:00:00Z | 7.2826389 | 7.2826389 | 1 | R2OK-g-0 |
| red-tailed hawk | Raptor | winter | OK | Garfield, Kay, Noble | 2 | grassland | R2OK-g-4 | 2019-02-11T00:00:00Z | 31.0333333 | 47.1201389 | 1 | R2OK-g-4 |
| Swainson's hawk | Raptor | winter | OK | Garfield, Kay, Noble | 2 | grassland | R2OK-g-4 | 2019-02-11T00:00:00Z | 30.9805556 | 47.0743056 | 1 | R2OK-g-4 |
| great horned owl | Raptor | winter | OK | Garfield, Kay, Noble | 2 | grassland | R2OK-g-4 | 2019-02-11T00:00:00Z | 22.3083333 | 30.9944444 | 1 | R2OK-g-4 |
| red-tailed hawk | Raptor | winter | OK | Garfield, Kay, Noble | 2 | grassland | R2OK-g-4 | 2019-02-11T00:00:00Z | 47.0069444 | NA | 1 | R2OK-g-4 |
| red-tailed hawk | Raptor | spring | OK | Garfield, Kay, Noble | 2 | grassland | R2OK-g-4 | 2019-03-14T00:00:00Z | 29.9263889 | 44.8958333 | 1 | R2OK-g-4 |
| red-tailed hawk | Raptor | spring | OK | Garfield, Kay, Noble | 2 | grassland | R2OK-g-4 | 2019-03-14T00:00:00Z | 44.8145833 | NA | 1 | R2OK-g-4 |
| barn owl | Raptor | spring | OK | Garfield, Kay, Noble | 2 | grassland | R2OK-g-4 | 2019-03-14T00:00:00Z | 6.8145833 | 9.7951389 | 1 | R2OK-g-4 |
| great horned owl | Raptor | spring | OK | Garfield, Kay, Noble | 2 | grassland | R2OK-g-4 | 2019-04-08T00:00:00Z | 30.1659722 | NA | 1 | R2OK-g-4 |
| barn owl | Raptor | spring | OK | Garfield, Kay, Noble | 2 | grassland | R2OK-g-4 | 2019-04-08T00:00:00Z | 1.8972222 | 2.8833333 | 1 | R2OK-g-4 |
| red-tailed hawk | Raptor | spring | OK | Garfield, Kay, Noble | 2 | grassland | R2OK-g-4 | 2019-04-08T00:00:00Z | 30.14375 | NA | 1 | R2OK-g-4 |
| barn owl | Raptor | spring | OK | Garfield, Kay, Noble | 2 | grassland | R2OK-g-4 | 2019-05-01T00:00:00Z | 47.2375 | NA | 1 | R2OK-g-4 |
| red-tailed hawk | Raptor | spring | OK | Garfield, Kay, Noble | 2 | grassland | R2OK-g-4 | 2019-05-01T00:00:00Z | 47.1777778 | NA | 1 | R2OK-g-4 |
| barn owl | Raptor | spring | OK | Garfield, Kay, Noble | 2 | grassland | R2OK-g-4 | 2019-05-01T00:00:00Z | 46.9145833 | NA | 1 | R2OK-g-4 |
| barn owl | Raptor | summer | OK | Garfield, Kay, Noble | 2 | grassland | R2OK-g-4 | 2019-06-07T00:00:00Z | 45.3715278 | NA | 1 | R2OK-g-4 |
| red-tailed hawk | Raptor | summer | OK | Garfield, Kay, Noble | 2 | grassland | R2OK-g-4 | 2019-06-07T00:00:00Z | 45.2256944 | NA | 1 | R2OK-g-4 |
| barn owl | Raptor | summer | OK | Garfield, Kay, Noble | 2 | grassland | R2OK-g-4 | 2019-06-07T00:00:00Z | 12.8229167 | 23.76875 | 1 | R2OK-g-4 |
| great horned owl | Raptor | summer | OK | Garfield, Kay, Noble | 2 | grassland | R2OK-g-4 | 2019-07-10T00:00:00Z | 10.1625 | NA | 1 | R2OK-g-4 |
| red-tailed hawk | Raptor | summer | OK | Garfield, Kay, Noble | 2 | grassland | R2OK-g-4 | 2019-07-10T00:00:00Z | 30.1444444 | 43.1583333 | 1 | R2OK-g-4 |
| turkey vulture | Raptor | summer | OK | Garfield, Kay, Noble | 2 | grassland | R2OK-g-4 | 2019-08-19T00:00:00Z | 20.1125 | 29.9770833 | 1 | R2OK-g-4 |
| turkey vulture | Raptor | fall | OK | Garfield, Kay, Noble | 2 | grassland | R2OK-g-4 | 2019-10-07T00:00:00Z | 105.8 | NA | 1 | R2OK-g-4 |
| red-tailed hawk | Raptor | winter | OK | Garfield, Kay, Noble | 2 | grassland | R2OK-g-4 | 2019-11-13T00:00:00Z | 0.9083333 | 7.7826389 | 1 | R2OK-g-4 |
| red-tailed hawk | Raptor | winter | OK | Garfield, Kay, Noble | 2 | grassland | R2OK-g-4 | 2019-11-22T00:00:00Z | 12.7993056 | 24.8013889 | 1 | R2OK-g-4 |
| barn owl | Raptor | winter | OK | Garfield, Kay, Noble | 2 | grassland | R2OK-g-4 | 2019-12-03T00:00:00Z | 13.9534722 | 19.9451389 | 1 | R2OK-g-4 |
| great horned owl | Raptor | winter | OK | Garfield, Kay, Noble | 2 | grassland | R2OK-g-4 | 2019-12-03T00:00:00Z | 17.9381944 | 27.9770833 | 1 | R2OK-g-4 |
| red-tailed hawk | Raptor | winter | OK | Garfield, Kay, Noble | 2 | grassland | R2OK-g-4 | 2019-12-03T00:00:00Z | 5.9743056 | 13.9555556 | 1 | R2OK-g-4 |
| barred owl | Raptor | winter | OK | Garfield, Kay, Noble | 2 | grassland | R2OK-g-4 | 2019-12-03T00:00:00Z | 5.9895833 | 13.9319444 | 1 | R2OK-g-4 |
| red-tailed hawk | Raptor | winter | OK | Garfield, Kay, Noble | 2 | grassland | R2OK-g-4 | 2019-12-03T00:00:00Z | 13.9569444 | 19.86875 | 1 | R2OK-g-4 |
| red-tailed hawk | Raptor | winter | OK | Garfield, Kay, Noble | 2 | grassland | R2OK-g-4 | 2019-12-03T00:00:00Z | 40.8569444 | 51.9729167 | 1 | R2OK-g-4 |
| Cooper's hawk | Raptor | winter | OK | Garfield, Kay, Noble | 2 | grassland | R2OK-g-4 | 2019-12-03T00:00:00Z | 0.9972222 | 0.9972222 | 1 | R2OK-g-4 |
| red-tailed hawk | Raptor | winter | OK | Garfield, Kay, Noble | 2 | grassland | R2OK-g-4 | 2019-12-03T00:00:00Z | 1.0152778 | 1.0152778 | 1 | R2OK-g-4 |
| prairie falcon | Raptor | winter | OK | Garfield, Kay, Noble | 2 | grassland | R2OK-g-4 | 2019-12-03T00:00:00Z | 1.0069444 | 1.9430556 | 1 | R2OK-g-4 |
| red-tailed hawk | Raptor | winter | OK | Garfield, Kay, Noble | 2 | grassland | R2OK-g-4 | 2019-12-03T00:00:00Z | 19.8034722 | 27.7944444 | 1 | R2OK-g-4 |
| barred owl | Raptor | winter | OK | Garfield, Kay, Noble | 2 | grassland | R2OK-g-4 | 2020-01-06T00:00:00Z | 2.88125 | 6.9506944 | 1 | R2OK-g-4 |
| red-tailed hawk | Raptor | winter | OK | Garfield, Kay, Noble | 2 | grassland | R2OK-g-4 | 2020-01-06T00:00:00Z | 6.9569444 | 13.8791667 | 1 | R2OK-g-4 |
| red-tailed hawk | Raptor | winter | OK | Garfield, Kay, Noble | 2 | grassland | R2OK-g-4 | 2020-01-06T00:00:00Z | 2.0194444 | 2.8701389 | 1 | R2OK-g-4 |
| Cooper's hawk | Raptor | winter | OK | Garfield, Kay, Noble | 2 | grassland | R2OK-g-4 | 2020-01-06T00:00:00Z | 2.0138889 | 2.8576389 | 1 | R2OK-g-4 |
| northern harrier | Raptor | winter | OK | Garfield, Kay, Noble | 2 | grassland | R2OK-g-4 | 2020-01-09T00:00:00Z | 12.1541667 | 18.1222222 | 1 | R2OK-g-4 |
| barn owl | Raptor | winter | OK | Garfield, Kay, Noble | 2 | grassland | R2OK-g-4 | 2020-01-06T00:00:00Z | 48.9784722 | NA | 1 | R2OK-g-4 |
| red-tailed hawk | Raptor | winter | OK | Garfield, Kay, Noble | 2 | grassland | R2OK-g-4 | 2020-01-06T00:00:00Z | 44.8625 | NA | 1 | R2OK-g-4 |
| red-tailed hawk | Raptor | winter | OK | Garfield, Kay, Noble | 2 | grassland | R2OK-g-4 | 2020-01-13T00:00:00Z | 41.9118056 | NA | 1 | R2OK-g-4 |
| red-tailed hawk | Raptor | winter | OK | Garfield, Kay, Noble | 2 | grassland | R2OK-g-4 | 2020-01-21T00:00:00Z | 12.86875 | 22.9388889 | 1 | R2OK-g-4 |
| ring-necked pheasant | Game Bird | fall | TX | Concho | 2 | shrub/scrub | R2TX-ss-1 | 2018-10-15T00:00:00Z | 2.2069444 | 2.9819444 | 0 | R2TX-ss-1 |
| ring-necked pheasant | Game Bird | fall | TX | Concho | 2 | shrub/scrub | R2TX-ss-1 | 2018-10-15T00:00:00Z | 7.0472222 | 10.2472222 | 0 | R2TX-ss-1 |
| ring-necked pheasant | Game Bird | fall | TX | Concho | 2 | shrub/scrub | R2TX-ss-1 | 2018-10-16T00:00:00Z | 1.1013889 | 1.1013889 | 0 | R2TX-ss-1 |
| ring-necked pheasant | Game Bird | fall | TX | Concho | 2 | shrub/scrub | R2TX-ss-1 | 2018-10-15T00:00:00Z | 2.2319444 | 2.99375 | 0 | R2TX-ss-1 |
| ring-necked pheasant | Game Bird | fall | TX | Concho | 2 | shrub/scrub | R2TX-ss-1 | 2018-10-16T00:00:00Z | 1.1055556 | 1.9930556 | 0 | R2TX-ss-1 |
| ring-necked pheasant | Game Bird | fall | TX | Concho | 2 | shrub/scrub | R2TX-ss-1 | 2018-10-16T00:00:00Z | 1.0909722 | 2.0270833 | 0 | R2TX-ss-1 |
| ring-necked pheasant | Game Bird | fall | TX | Concho | 2 | shrub/scrub | R2TX-ss-1 | 2018-10-16T00:00:00Z | 1.0875 | 2.0201389 | 0 | R2TX-ss-1 |
| ring-necked pheasant | Game Bird | fall | TX | Concho | 2 | shrub/scrub | R2TX-ss-1 | 2018-10-16T00:00:00Z | 1.16875 | 2.0236111 | 0 | R2TX-ss-1 |
| ring-necked pheasant | Game Bird | winter | TX | Concho | 2 | shrub/scrub | R2TX-ss-1 | 2018-11-27T00:00:00Z | 5.9868056 | 8.0111111 | 0 | R2TX-ss-1 |
| ring-necked pheasant | Game Bird | winter | TX | Concho | 2 | shrub/scrub | R2TX-ss-1 | 2018-11-27T00:00:00Z | 1.9694444 | 2.9305556 | 0 | R2TX-ss-1 |
| ring-necked pheasant | Game Bird | winter | TX | Concho | 2 | shrub/scrub | R2TX-ss-1 | 2018-11-27T00:00:00Z | 5.9729167 | 8.0034722 | 0 | R2TX-ss-1 |
| ring-necked pheasant | Game Bird | winter | TX | Concho | 2 | shrub/scrub | R2TX-ss-1 | 2018-11-27T00:00:00Z | 1.0777778 | 1.9493056 | 0 | R2TX-ss-1 |
| ring-necked pheasant | Game Bird | winter | TX | Concho | 2 | shrub/scrub | R2TX-ss-1 | 2018-11-28T00:00:00Z | 1.0930556 | 2.0729167 | 0 | R2TX-ss-1 |
| ring-necked pheasant | Game Bird | winter | TX | Concho | 2 | shrub/scrub | R2TX-ss-1 | 2018-11-28T00:00:00Z | 14.1756944 | 21.0097222 | 0 | R2TX-ss-1 |
| ring-necked pheasant | Game Bird | winter | TX | Concho | 2 | shrub/scrub | R2TX-ss-1 | 2018-11-28T00:00:00Z | 1.0652778 | 1.0652778 | 0 | R2TX-ss-1 |
| ring-necked pheasant | Game Bird | winter | TX | Concho | 2 | shrub/scrub | R2TX-ss-1 | 2018-11-28T00:00:00Z | 5.0215278 | 7.05 | 0 | R2TX-ss-1 |
| ring-necked pheasant | Game Bird | winter | TX | Concho | 2 | shrub/scrub | R2TX-ss-1 | 2018-11-28T00:00:00Z | 1.0583333 | 1.0583333 | 0 | R2TX-ss-1 |
| ring-necked pheasant | Game Bird | winter | TX | Concho | 2 | shrub/scrub | R2TX-ss-1 | 2018-11-28T00:00:00Z | 1.0527778 | 1.0527778 | 0 | R2TX-ss-1 |
| ring-necked pheasant | Game Bird | winter | TX | Concho | 2 | shrub/scrub | R2TX-ss-1 | 2018-11-28T00:00:00Z | 2.0118056 | 5.0020833 | 0 | R2TX-ss-1 |
| ring-necked pheasant | Game Bird | winter | TX | Concho | 2 | shrub/scrub | R2TX-ss-1 | 2019-01-07T00:00:00Z | 14.0340278 | 28.0229167 | 0 | R2TX-ss-1 |
| ring-necked pheasant | Game Bird | winter | TX | Concho | 2 | shrub/scrub | R2TX-ss-1 | 2019-01-07T00:00:00Z | 1.0708333 | 3.0173611 | 0 | R2TX-ss-1 |
| ring-necked pheasant | Game Bird | winter | TX | Concho | 2 | shrub/scrub | R2TX-ss-1 | 2019-01-08T00:00:00Z | 2.0368056 | 3.0236111 | 0 | R2TX-ss-1 |
| ring-necked pheasant | Game Bird | winter | TX | Concho | 2 | shrub/scrub | R2TX-ss-1 | 2019-01-08T00:00:00Z | 6.1270833 | 9.0555556 | 0 | R2TX-ss-1 |
| ring-necked pheasant | Game Bird | spring | TX | Concho | 2 | shrub/scrub | R2TX-ss-1 | 2019-03-12T00:00:00Z | 0.9388889 | 0.9388889 | 0 | R2TX-ss-1 |
| ring-necked pheasant | Game Bird | spring | TX | Concho | 2 | shrub/scrub | R2TX-ss-1 | 2019-03-12T00:00:00Z | 0.90625 | 0.90625 | 0 | R2TX-ss-1 |
| ring-necked pheasant | Game Bird | spring | TX | Concho | 2 | shrub/scrub | R2TX-ss-1 | 2019-03-12T00:00:00Z | 0.8840278 | 1.9375 | 0 | R2TX-ss-1 |
| ring-necked pheasant | Game Bird | spring | TX | Concho | 2 | shrub/scrub | R2TX-ss-1 | 2019-03-12T00:00:00Z | 0.8819444 | 0.8819444 | 0 | R2TX-ss-1 |
| ring-necked pheasant | Game Bird | spring | TX | Concho | 2 | shrub/scrub | R2TX-ss-1 | 2019-03-12T00:00:00Z | 0.875 | 1.9284722 | 0 | R2TX-ss-1 |
| ring-necked pheasant | Game Bird | spring | TX | Concho | 2 | shrub/scrub | R2TX-ss-1 | 2019-03-12T00:00:00Z | 7.0777778 | 9.875 | 0 | R2TX-ss-1 |
| ring-necked duck | Game Bird | summer | TX | Concho | 2 | shrub/scrub | R2TX-ss-1 | 2019-06-03T00:00:00Z | 1.9694444 | 2.9326389 | 0 | R2TX-ss-1 |
| turkey vulture | Raptor | summer | TX | Concho | 2 | shrub/scrub | R2TX-ss-1 | 2019-06-03T00:00:00Z | 29.9354167 | NA | 0 | R2TX-ss-0 |
| ring-necked pheasant | Game Bird | summer | TX | Concho | 2 | shrub/scrub | R2TX-ss-1 | 2019-07-29T00:00:00Z | 29.9013889 | NA | 0 | R2TX-ss-1 |
| ring-necked pheasant | Game Bird | summer | TX | Concho | 2 | shrub/scrub | R2TX-ss-1 | 2019-07-29T00:00:00Z | 1.2048611 | 1.9868056 | 0 | R2TX-ss-1 |
| ring-necked pheasant | Game Bird | summer | TX | Concho | 2 | shrub/scrub | R2TX-ss-1 | 2019-07-29T00:00:00Z | 1.9590278 | 3.0166667 | 0 | R2TX-ss-1 |
| ring-necked pheasant | Game Bird | summer | TX | Concho | 2 | shrub/scrub | R2TX-ss-1 | 2019-07-29T00:00:00Z | 1.13125 | 1.13125 | 0 | R2TX-ss-1 |
| ring-necked pheasant | Game Bird | summer | TX | Concho | 2 | shrub/scrub | R2TX-ss-1 | 2019-07-29T00:00:00Z | 1.9319444 | 2.9875 | 0 | R2TX-ss-1 |
| ring-necked pheasant | Game Bird | summer | TX | Concho | 2 | shrub/scrub | R2TX-ss-1 | 2019-07-29T00:00:00Z | 15.0361111 | 20.9576389 | 0 | R2TX-ss-1 |
| ring-necked pheasant | Game Bird | summer | TX | Concho | 2 | shrub/scrub | R2TX-ss-1 | 2019-07-29T00:00:00Z | 0.99375 | 1.8923611 | 0 | R2TX-ss-1 |
| ring-necked pheasant | Game Bird | summer | TX | Concho | 2 | shrub/scrub | R2TX-ss-1 | 2019-07-29T00:00:00Z | 15.0027778 | 20.9506944 | 0 | R2TX-ss-1 |
| ring-necked pheasant | Game Bird | summer | TX | Concho | 2 | shrub/scrub | R2TX-ss-1 | 2019-07-29T00:00:00Z | 29.8666667 | NA | 0 | R2TX-ss-1 |
| ring-necked pheasant | Game Bird | fall | TX | Irion and Reagan | 2 | shrub/scrub | R2TX-ss-2 | 2018-10-15T00:00:00Z | 29.9833333 | NA | 0 | R2TX-ss-2 |
| ring-necked pheasant | Game Bird | fall | TX | Irion and Reagan | 2 | shrub/scrub | R2TX-ss-2 | 2018-10-15T00:00:00Z | 20.8666667 | 29.9729167 | 0 | R2TX-ss-2 |
| ring-necked pheasant | Game Bird | fall | TX | Irion and Reagan | 2 | shrub/scrub | R2TX-ss-2 | 2018-10-15T00:00:00Z | 6.8298611 | 9.9034722 | 0 | R2TX-ss-2 |
| ring-necked pheasant | Game Bird | fall | TX | Irion and Reagan | 2 | shrub/scrub | R2TX-ss-2 | 2018-10-15T00:00:00Z | 21.8916667 | 29.9423611 | 0 | R2TX-ss-2 |
| ring-necked pheasant | Game Bird | fall | TX | Irion and Reagan | 2 | shrub/scrub | R2TX-ss-2 | 2018-10-15T00:00:00Z | 3.9263889 | 6.7972222 | 0 | R2TX-ss-2 |
| red-tailed hawk | Raptor | winter | TX | Irion and Reagan | 2 | shrub/scrub | R2TX-ss-2 | 2019-02-04T00:00:00Z | 30.0479167 | NA | 0 | R2TX-ss-0 |
| ring-necked pheasant | Game Bird | fall | TX | Irion and Reagan | 2 | shrub/scrub | R2TX-ss-2 | 2018-09-10T00:00:00Z | 1.6604167 | 2.9777778 | 0 | R2TX-ss-2 |
| ring-necked pheasant | Game Bird | fall | TX | Irion and Reagan | 2 | shrub/scrub | R2TX-ss-2 | 2018-09-10T00:00:00Z | 9.9763889 | 13.9555556 | 0 | R2TX-ss-2 |
| ring-necked pheasant | Game Bird | fall | TX | Irion and Reagan | 2 | shrub/scrub | R2TX-ss-2 | 2018-09-10T00:00:00Z | 0.9548611 | 0.9548611 | 0 | R2TX-ss-2 |
| ring-necked pheasant | Game Bird | fall | TX | Irion and Reagan | 2 | shrub/scrub | R2TX-ss-2 | 2018-09-10T00:00:00Z | 29.825 | NA | 0 | R2TX-ss-2 |
| ring-necked pheasant | Game Bird | fall | TX | Irion and Reagan | 2 | shrub/scrub | R2TX-ss-2 | 2018-09-10T00:00:00Z | 29.9298611 | NA | 0 | R2TX-ss-2 |
| ring-necked pheasant | Game Bird | fall | TX | Irion and Reagan | 2 | shrub/scrub | R2TX-ss-2 | 2018-09-10T00:00:00Z | 30.0270833 | NA | 0 | R2TX-ss-2 |
| ring-necked pheasant | Game Bird | fall | TX | Irion and Reagan | 2 | shrub/scrub | R2TX-ss-2 | 2018-09-10T00:00:00Z | 4.0375 | 6.8472222 | 0 | R2TX-ss-2 |
| ring-necked pheasant | Game Bird | fall | TX | Irion and Reagan | 2 | shrub/scrub | R2TX-ss-2 | 2018-09-10T00:00:00Z | 2.9694444 | 3.9743056 | 0 | R2TX-ss-2 |
| ring-necked pheasant | Game Bird | fall | TX | Irion and Reagan | 2 | shrub/scrub | R2TX-ss-2 | 2018-09-10T00:00:00Z | 20.9722222 | 29.8652778 | 0 | R2TX-ss-2 |
| ring-necked pheasant | Game Bird | fall | TX | Irion and Reagan | 2 | shrub/scrub | R2TX-ss-2 | 2018-09-10T00:00:00Z | 3.7902778 | 6.6263889 | 0 | R2TX-ss-2 |
| ring-necked pheasant | Game Bird | fall | TX | Irion and Reagan | 2 | shrub/scrub | R2TX-ss-2 | 2018-10-08T00:00:00Z | 3.9423611 | 7.0138889 | 0 | R2TX-ss-2 |
| ring-necked pheasant | Game Bird | fall | TX | Irion and Reagan | 2 | shrub/scrub | R2TX-ss-2 | 2018-10-08T00:00:00Z | 2.9416667 | 3.9256944 | 0 | R2TX-ss-2 |
| ring-necked pheasant | Game Bird | fall | TX | Irion and Reagan | 2 | shrub/scrub | R2TX-ss-2 | 2018-10-08T00:00:00Z | 21.0118056 | 30.9916667 | 0 | R2TX-ss-2 |
| ring-necked pheasant | Game Bird | fall | TX | Irion and Reagan | 2 | shrub/scrub | R2TX-ss-2 | 2018-10-08T00:00:00Z | 30.9597222 | NA | 0 | R2TX-ss-2 |
| ring-necked pheasant | Game Bird | fall | TX | Irion and Reagan | 2 | shrub/scrub | R2TX-ss-2 | 2018-10-15T00:00:00Z | 3.9416667 | 6.8027778 | 0 | R2TX-ss-2 |
| turkey vulture | Raptor | summer | TX | Irion and Reagan | 2 | shrub/scrub | R2TX-ss-2 | 2019-06-24T00:00:00Z | 29.8979167 | NA | 0 | R2TX-ss-0 |
| black vulture | Raptor | summer | TX | Irion and Reagan | 2 | shrub/scrub | R2TX-ss-2 | 2019-06-24T00:00:00Z | 29.9 | NA | 0 | R2TX-ss-0 |
| turkey vulture | Raptor | summer | TX | Irion and Reagan | 2 | shrub/scrub | R2TX-ss-2 | 2019-06-24T00:00:00Z | 29.9131944 | NA | 0 | R2TX-ss-0 |
| red-tailed hawk | Raptor | summer | TX | Irion and Reagan | 2 | shrub/scrub | R2TX-ss-2 | 2019-06-24T00:00:00Z | 29.9423611 | NA | 0 | R2TX-ss-0 |
| turkey vulture | Raptor | summer | TX | Irion and Reagan | 2 | shrub/scrub | R2TX-ss-2 | 2019-06-24T00:00:00Z | 29.9305556 | NA | 0 | R2TX-ss-0 |
| turkey vulture | Raptor | summer | TX | Irion and Reagan | 2 | shrub/scrub | R2TX-ss-2 | 2019-06-24T00:00:00Z | 29.91875 | NA | 0 | R2TX-ss-0 |
| Swainson's hawk | Raptor | fall | WA | Klickitat | 1 | grassland | R1WA-g-1 | 2018-10-17T00:00:00Z | 56.9673611 | 85.0993056 | 0 | R1WA-g-1 |
| Swainson's hawk | Raptor | fall | WA | Klickitat | 1 | grassland | R1WA-g-1 | 2018-10-17T00:00:00Z | 40.9722222 | 56.9930556 | 0 | R1WA-g-1 |
| Swainson's hawk | Raptor | fall | WA | Klickitat | 1 | grassland | R1WA-g-1 | 2018-10-17T00:00:00Z | 95.0625 | NA | 0 | R1WA-g-1 |
| Swainson's hawk | Raptor | fall | WA | Klickitat | 1 | grassland | R1WA-g-1 | 2018-10-17T00:00:00Z | 94.9861111 | NA | 0 | R1WA-g-1 |
| red-tailed hawk | Raptor | fall | WA | Klickitat | 1 | grassland | R1WA-g-1 | 2018-10-17T00:00:00Z | 57.0659722 | 85.1875 | 0 | R1WA-g-1 |
| red-tailed hawk | Raptor | winter | WA | Klickitat | 1 | grassland | R1WA-g-1 | 2018-12-03T00:00:00Z | 3.09375 | NA | 0 | R1WA-g-1 |
| Swainson's hawk | Raptor | winter | WA | Klickitat | 1 | grassland | R1WA-g-1 | 2018-12-03T00:00:00Z | 3.1006944 | NA | 0 | R1WA-g-1 |
| red-tailed hawk | Raptor | spring | WA | Klickitat | 1 | grassland | R1WA-g-1 | 2020-04-21T00:00:00Z | 56.1333333 | NA | 0 | R1WA-g-1 |
| Cooper's hawk | Raptor | spring | WA | Klickitat | 1 | grassland | R1WA-g-1 | 2020-04-21T00:00:00Z | 2.9333333 | 2.9333333 | 0 | R1WA-g-1 |
| red-tailed hawk | Raptor | spring | WA | Klickitat | 1 | grassland | R1WA-g-1 | 2020-04-21T00:00:00Z | 56.0895833 | NA | 0 | R1WA-g-1 |
| Cooper's hawk | Raptor | spring | WA | Klickitat | 1 | grassland | R1WA-g-1 | 2020-04-21T00:00:00Z | 2.925 | 7.2416667 | 0 | R1WA-g-1 |
| red-tailed hawk | Raptor | summer | WA | Klickitat | 1 | grassland | R1WA-g-1 | 2020-06-23T00:00:00Z | 58.0555556 | NA | 0 | R1WA-g-1 |
| red-tailed hawk | Raptor | summer | WA | Klickitat | 1 | grassland | R1WA-g-1 | 2020-06-23T00:00:00Z | 58.0625 | NA | 0 | R1WA-g-1 |
| red-tailed hawk | Raptor | summer | WA | Klickitat | 1 | grassland | R1WA-g-1 | 2020-06-23T00:00:00Z | 58.0451389 | NA | 0 | R1WA-g-1 |
| red-tailed hawk | Raptor | summer | WA | Klickitat | 1 | grassland | R1WA-g-1 | 2020-06-23T00:00:00Z | 14.3819444 | 21.1909722 | 0 | R1WA-g-1 |
| red-tailed hawk | Raptor | summer | WA | Klickitat | 1 | grassland | R1WA-g-1 | 2020-06-23T00:00:00Z | 58.0277778 | NA | 0 | R1WA-g-1 |
| red-tailed hawk | Raptor | spring | WA | Klickitat | 1 | grassland | R1WA-g-1 | 2020-04-21T00:00:00Z | 56.1333333 | NA | 0 | R1WA-g-1 |
| great horned owl | Raptor | spring | WA | Klickitat | 1 | grassland | R1WA-g-1 | 2020-04-21T00:00:00Z | 14.1965278 | 21.2090278 | 0 | R1WA-g-1 |
| barn owl | Raptor | spring | WA | Klickitat | 1 | grassland | R1WA-g-1 | 2020-04-21T00:00:00Z | 59.1166667 | NA | 0 | R1WA-g-1 |
| Cooper's hawk | Raptor | spring | WA | Klickitat | 1 | grassland | R1WA-g-1 | 2020-04-21T00:00:00Z | 2.9333333 | 2.9333333 | 0 | R1WA-g-1 |
| red-tailed hawk | Raptor | spring | WA | Klickitat | 1 | grassland | R1WA-g-1 | 2020-04-21T00:00:00Z | 56.0895833 | NA | 0 | R1WA-g-1 |
| great horned owl | Raptor | spring | WA | Klickitat | 1 | grassland | R1WA-g-1 | 2020-04-21T00:00:00Z | 56.0868056 | NA | 0 | R1WA-g-1 |
| Cooper's hawk | Raptor | spring | WA | Klickitat | 1 | grassland | R1WA-g-1 | 2020-04-21T00:00:00Z | 2.925 | 7.2416667 | 0 | R1WA-g-1 |
| barn owl | Raptor | spring | WA | Klickitat | 1 | grassland | R1WA-g-1 | 2020-04-21T00:00:00Z | 21.1604167 | 28.8479167 | 0 | R1WA-g-1 |
| great horned owl | Raptor | summer | WA | Klickitat | 1 | grassland | R1WA-g-1 | 2020-06-23T00:00:00Z | 58.0479167 | NA | 0 | R1WA-g-1 |
| red-tailed hawk | Raptor | summer | WA | Klickitat | 1 | grassland | R1WA-g-1 | 2020-06-23T00:00:00Z | 58.0555556 | NA | 0 | R1WA-g-1 |
| red-tailed hawk | Raptor | summer | WA | Klickitat | 1 | grassland | R1WA-g-1 | 2020-06-23T00:00:00Z | 58.0625 | NA | 0 | R1WA-g-1 |
| barn owl | Raptor | summer | WA | Klickitat | 1 | grassland | R1WA-g-1 | 2020-06-23T00:00:00Z | 49.375 | 58.0451389 | 0 | R1WA-g-1 |
| red-tailed hawk | Raptor | summer | WA | Klickitat | 1 | grassland | R1WA-g-1 | 2020-06-23T00:00:00Z | 58.0451389 | NA | 0 | R1WA-g-1 |
| great horned owl | Raptor | summer | WA | Klickitat | 1 | grassland | R1WA-g-1 | 2020-06-23T00:00:00Z | 58.0381944 | NA | 0 | R1WA-g-1 |
| red-tailed hawk | Raptor | summer | WA | Klickitat | 1 | grassland | R1WA-g-1 | 2020-06-23T00:00:00Z | 14.3819444 | 21.1909722 | 0 | R1WA-g-1 |
| red-tailed hawk | Raptor | summer | WA | Klickitat | 1 | grassland | R1WA-g-1 | 2020-06-23T00:00:00Z | 58.0277778 | NA | 0 | R1WA-g-1 |
| barred owl | Raptor | winter | WA | Kittitas, Garfield, Columbia | 1 | shrub/scrub | R1WA-ss-3 | 2020-01-03T00:00:00Z | 89.7881944 | NA | 0 | R1WA-ss-3 |
| barred owl | Raptor | winter | WA | Kittitas, Garfield, Columbia | 1 | shrub/scrub | R1WA-ss-3 | 2020-01-03T00:00:00Z | 90.0027778 | NA | 0 | R1WA-ss-3 |
| barred owl | Raptor | winter | WA | Kittitas, Garfield, Columbia | 1 | shrub/scrub | R1WA-ss-3 | 2020-01-03T00:00:00Z | 89.8555556 | NA | 0 | R1WA-ss-3 |
| barred owl | Raptor | winter | WA | Kittitas, Garfield, Columbia | 1 | shrub/scrub | R1WA-ss-3 | 2020-01-03T00:00:00Z | 3.4826389 | 3.4826389 | 0 | R1WA-ss-3 |
| barred owl | Raptor | winter | WA | Kittitas, Garfield, Columbia | 1 | shrub/scrub | R1WA-ss-3 | 2020-01-03T00:00:00Z | 89.7972222 | NA | 0 | R1WA-ss-3 |
| barred owl | Raptor | winter | WA | Kittitas, Garfield, Columbia | 1 | shrub/scrub | R1WA-ss-3 | 2020-01-03T00:00:00Z | 74.9131944 | 89.8020833 | 0 | R1WA-ss-3 |
| barred owl | Raptor | winter | WA | Kittitas, Garfield, Columbia | 1 | shrub/scrub | R1WA-ss-3 | 2020-01-03T00:00:00Z | 5.3520833 | NA | 0 | R1WA-ss-3 |
| barred owl | Raptor | winter | WA | Kittitas, Garfield, Columbia | 1 | shrub/scrub | R1WA-ss-3 | 2020-01-03T00:00:00Z | 5.0034722 | 10.1041667 | 0 | R1WA-ss-3 |
| barred owl | Raptor | winter | WA | Kittitas, Garfield, Columbia | 1 | shrub/scrub | R1WA-ss-3 | 2020-01-03T00:00:00Z | 89.8020833 | NA | 0 | R1WA-ss-3 |
| barred owl | Raptor | winter | WA | Kittitas, Garfield, Columbia | 1 | shrub/scrub | R1WA-ss-3 | 2020-01-03T00:00:00Z | 89.7993056 | NA | 0 | R1WA-ss-3 |
| barred owl | Raptor | winter | WA | Kittitas, Garfield, Columbia | 1 | shrub/scrub | R1WA-ss-3 | 2020-01-03T00:00:00Z | 5.3236111 | 10.3979167 | 0 | R1WA-ss-3 |
| barred owl | Raptor | winter | WA | Kittitas, Garfield, Columbia | 1 | shrub/scrub | R1WA-ss-3 | 2020-01-03T00:00:00Z | 31.8701389 | 47.7506944 | 0 | R1WA-ss-3 |
| barred owl | Raptor | winter | WA | Kittitas, Garfield, Columbia | 1 | shrub/scrub | R1WA-ss-3 | 2020-01-03T00:00:00Z | 5.0034722 | 9.9069444 | 0 | R1WA-ss-3 |
| barred owl | Raptor | winter | WA | Kittitas, Garfield, Columbia | 1 | shrub/scrub | R1WA-ss-3 | 2020-01-03T00:00:00Z | 74.8881944 | 89.8819444 | 0 | R1WA-ss-3 |
| barred owl | Raptor | winter | WA | Kittitas, Garfield, Columbia | 1 | shrub/scrub | R1WA-ss-3 | 2020-01-03T00:00:00Z | 5.33125 | 10.4291667 | 0 | R1WA-ss-3 |
| barred owl | Raptor | winter | WA | Kittitas, Garfield, Columbia | 1 | shrub/scrub | R1WA-ss-3 | 2020-01-03T00:00:00Z | 20.8784722 | 31.8604167 | 0 | R1WA-ss-3 |
| barred owl | Raptor | winter | WA | Kittitas, Garfield, Columbia | 1 | shrub/scrub | R1WA-ss-3 | 2020-01-03T00:00:00Z | 3.0493056 | 3.0493056 | 0 | R1WA-ss-3 |
| barred owl | Raptor | winter | WA | Kittitas, Garfield, Columbia | 1 | shrub/scrub | R1WA-ss-3 | 2020-01-03T00:00:00Z | 59.8631944 | 74.8868056 | 0 | R1WA-ss-3 |
| barred owl | Raptor | winter | WA | Kittitas, Garfield, Columbia | 1 | shrub/scrub | R1WA-ss-3 | 2020-01-03T00:00:00Z | 89.8833333 | NA | 0 | R1WA-ss-3 |
| barred owl | Raptor | winter | WA | Kittitas, Garfield, Columbia | 1 | shrub/scrub | R1WA-ss-3 | 2020-01-03T00:00:00Z | 89.9506944 | NA | 0 | R1WA-ss-3 |
| barred owl | Raptor | fall | WA | Kittitas, Garfield, Columbia | 1 | shrub/scrub | R1WA-ss-3 | 2020-09-03T00:00:00Z | 89.9784722 | NA | 0 | R1WA-ss-3 |
| barred owl | Raptor | fall | WA | Kittitas, Garfield, Columbia | 1 | shrub/scrub | R1WA-ss-3 | 2020-09-03T00:00:00Z | 42.9402778 | 59.99375 | 0 | R1WA-ss-3 |
| barred owl | Raptor | fall | WA | Kittitas, Garfield, Columbia | 1 | shrub/scrub | R1WA-ss-3 | 2020-09-03T00:00:00Z | 90.0583333 | NA | 0 | R1WA-ss-3 |
| barred owl | Raptor | fall | WA | Kittitas, Garfield, Columbia | 1 | shrub/scrub | R1WA-ss-3 | 2020-09-03T00:00:00Z | 90.0979167 | NA | 0 | R1WA-ss-3 |
| barred owl | Raptor | fall | WA | Kittitas, Garfield, Columbia | 1 | shrub/scrub | R1WA-ss-3 | 2020-09-03T00:00:00Z | 89.8479167 | NA | 0 | R1WA-ss-3 |
| barred owl | Raptor | fall | WA | Kittitas, Garfield, Columbia | 1 | shrub/scrub | R1WA-ss-3 | 2020-09-03T00:00:00Z | 12.0680556 | 20.0597222 | 0 | R1WA-ss-3 |
| barred owl | Raptor | fall | WA | Kittitas, Garfield, Columbia | 1 | shrub/scrub | R1WA-ss-3 | 2020-09-03T00:00:00Z | 90.0625 | NA | 0 | R1WA-ss-3 |
| barred owl | Raptor | fall | WA | Kittitas, Garfield, Columbia | 1 | shrub/scrub | R1WA-ss-3 | 2020-09-03T00:00:00Z | 29.1194444 | 42.95625 | 0 | R1WA-ss-3 |
| barred owl | Raptor | fall | WA | Kittitas, Garfield, Columbia | 1 | shrub/scrub | R1WA-ss-3 | 2020-09-03T00:00:00Z | 29.0777778 | 42.9680556 | 0 | R1WA-ss-3 |
| barred owl | Raptor | fall | WA | Kittitas, Garfield, Columbia | 1 | shrub/scrub | R1WA-ss-3 | 2020-09-03T00:00:00Z | 5.1166667 | 12.0326389 | 0 | R1WA-ss-3 |
| barred owl | Raptor | fall | WA | Kittitas, Garfield, Columbia | 1 | shrub/scrub | R1WA-ss-3 | 2020-09-03T00:00:00Z | 5.2347222 | 12.0520833 | 0 | R1WA-ss-3 |
| barred owl | Raptor | fall | WA | Kittitas, Garfield, Columbia | 1 | shrub/scrub | R1WA-ss-3 | 2020-09-03T00:00:00Z | 1.0277778 | 5.2090278 | 0 | R1WA-ss-3 |
| barred owl | Raptor | fall | WA | Kittitas, Garfield, Columbia | 1 | shrub/scrub | R1WA-ss-3 | 2020-09-03T00:00:00Z | 29.0541667 | 42.9326389 | 0 | R1WA-ss-3 |
| barred owl | Raptor | fall | WA | Kittitas, Garfield, Columbia | 1 | shrub/scrub | R1WA-ss-3 | 2020-09-03T00:00:00Z | 60.0847222 | 75.0347222 | 0 | R1WA-ss-3 |
| barred owl | Raptor | fall | WA | Kittitas, Garfield, Columbia | 1 | shrub/scrub | R1WA-ss-3 | 2020-09-03T00:00:00Z | 89.8270833 | NA | 0 | R1WA-ss-3 |
| barred owl | Raptor | fall | WA | Kittitas, Garfield, Columbia | 1 | shrub/scrub | R1WA-ss-3 | 2020-09-03T00:00:00Z | 12.0590278 | 20.0541667 | 0 | R1WA-ss-3 |
| barred owl | Raptor | fall | WA | Kittitas, Garfield, Columbia | 1 | shrub/scrub | R1WA-ss-3 | 2020-09-03T00:00:00Z | 1.0284722 | 1.0284722 | 0 | R1WA-ss-3 |
| barred owl | Raptor | fall | WA | Kittitas, Garfield, Columbia | 1 | shrub/scrub | R1WA-ss-3 | 2020-09-03T00:00:00Z | 5.0034722 | 12.0472222 | 0 | R1WA-ss-3 |
| barred owl | Raptor | fall | WA | Kittitas, Garfield, Columbia | 1 | shrub/scrub | R1WA-ss-3 | 2020-09-03T00:00:00Z | 90.0798611 | NA | 0 | R1WA-ss-3 |
| barred owl | Raptor | fall | WA | Kittitas, Garfield, Columbia | 1 | shrub/scrub | R1WA-ss-3 | 2020-09-03T00:00:00Z | 4.9333333 | 12.0458333 | 0 | R1WA-ss-3 |
| barred owl | Raptor | summer | WA | Lewis and Thurston | 1 | forest | R1WA-f-1 | 2020-07-20T00:00:00Z | 3.8875 | 7.0465278 | 0 | R1WA-f-1 |
| barred owl | Raptor | summer | WA | Lewis and Thurston | 1 | forest | R1WA-f-1 | 2020-07-20T00:00:00Z | 1.95625 | 2.8965278 | 0 | R1WA-f-1 |
| barred owl | Raptor | summer | WA | Lewis and Thurston | 1 | forest | R1WA-f-1 | 2020-07-20T00:00:00Z | 69.7715278 | 79.7611111 | 0 | R1WA-f-1 |
| barred owl | Raptor | summer | WA | Lewis and Thurston | 1 | forest | R1WA-f-1 | 2020-07-20T00:00:00Z | 89.8506944 | NA | 0 | R1WA-f-1 |
| barred owl | Raptor | summer | WA | Lewis and Thurston | 1 | forest | R1WA-f-1 | 2020-07-20T00:00:00Z | 89.8777778 | NA | 0 | R1WA-f-1 |
| barred owl | Raptor | summer | WA | Lewis and Thurston | 1 | forest | R1WA-f-1 | 2020-07-20T00:00:00Z | 89.9020833 | NA | 0 | R1WA-f-1 |
| barred owl | Raptor | summer | WA | Lewis and Thurston | 1 | forest | R1WA-f-1 | 2020-07-20T00:00:00Z | 39.8361111 | 55.9131944 | 0 | R1WA-f-1 |
| barred owl | Raptor | summer | WA | Lewis and Thurston | 1 | forest | R1WA-f-1 | 2020-07-20T00:00:00Z | 29.8013889 | 39.8159722 | 0 | R1WA-f-1 |
| barred owl | Raptor | summer | WA | Lewis and Thurston | 1 | forest | R1WA-f-1 | 2020-07-20T00:00:00Z | 89.9618056 | NA | 0 | R1WA-f-1 |
| barred owl | Raptor | summer | WA | Lewis and Thurston | 1 | forest | R1WA-f-1 | 2020-07-20T00:00:00Z | 59.8645833 | 69.8333333 | 0 | R1WA-f-1 |
| barred owl | Raptor | summer | WA | Lewis and Thurston | 1 | forest | R1WA-f-1 | 2020-07-20T00:00:00Z | 29.8215278 | 39.8451389 | 0 | R1WA-f-1 |
| barred owl | Raptor | summer | WA | Lewis and Thurston | 1 | forest | R1WA-f-1 | 2020-07-20T00:00:00Z | 29.9784722 | 40.0034722 | 0 | R1WA-f-1 |
| barred owl | Raptor | summer | WA | Lewis and Thurston | 1 | forest | R1WA-f-1 | 2020-07-20T00:00:00Z | 20.00625 | 29.9291667 | 0 | R1WA-f-1 |
| mallard | Game Bird | spring | WA | Garfield | 1 | cropland | R1WA-c-2 | 2017-04-17T00:00:00Z | 3.9694444 | 6.9541667 | 0 | R1WA-c-2 |
| mallard | Game Bird | spring | WA | Garfield | 1 | cropland | R1WA-c-2 | 2017-04-17T00:00:00Z | 0.9319444 | 1.95625 | 0 | R1WA-c-2 |
| ring-necked pheasant | Game Bird | spring | WA | Garfield | 1 | cropland | R1WA-c-2 | 2017-04-17T00:00:00Z | 9.9131944 | 14.0340278 | 0 | R1WA-c-2 |
| ring-necked pheasant | Game Bird | spring | WA | Garfield | 1 | cropland | R1WA-c-2 | 2017-05-01T00:00:00Z | 3.89375 | 6.8875 | 0 | R1WA-c-2 |
| mallard | Game Bird | spring | WA | Garfield | 1 | cropland | R1WA-c-2 | 2017-04-17T00:00:00Z | 29.9263889 | NA | 0 | R1WA-c-2 |
| ring-necked pheasant | Game Bird | spring | WA | Garfield | 1 | cropland | R1WA-c-2 | 2017-04-17T00:00:00Z | 29.8916667 | NA | 0 | R1WA-c-2 |
| ring-necked pheasant | Game Bird | spring | WA | Garfield | 1 | cropland | R1WA-c-2 | 2017-04-17T00:00:00Z | 13.8826389 | 20.15625 | 0 | R1WA-c-2 |
| mallard | Game Bird | spring | WA | Garfield | 1 | cropland | R1WA-c-2 | 2017-05-01T00:00:00Z | 29.7645833 | NA | 0 | R1WA-c-2 |
| ring-necked pheasant | Game Bird | spring | WA | Garfield | 1 | cropland | R1WA-c-2 | 2017-05-01T00:00:00Z | 29.8166667 | NA | 0 | R1WA-c-2 |
| mallard | Game Bird | spring | WA | Garfield | 1 | cropland | R1WA-c-2 | 2017-05-01T00:00:00Z | 29.8583333 | NA | 0 | R1WA-c-2 |
| ring-necked pheasant | Game Bird | summer | WA | Garfield | 1 | cropland | R1WA-c-2 | 2017-06-12T00:00:00Z | 3.9965278 | 6.7888889 | 0 | R1WA-c-2 |
| mallard | Game Bird | summer | WA | Garfield | 1 | cropland | R1WA-c-2 | 2017-06-28T00:00:00Z | 30.1333333 | NA | 0 | R1WA-c-2 |
| ring-necked pheasant | Game Bird | fall | WA | Garfield | 1 | cropland | R1WA-c-2 | 2017-08-14T00:00:00Z | 30.1555556 | NA | 0 | R1WA-c-2 |
| mallard | Game Bird | fall | WA | Garfield | 1 | cropland | R1WA-c-2 | 2017-10-01T00:00:00Z | 1.7722222 | 2.7527778 | 0 | R1WA-c-2 |
| ring-necked pheasant | Game Bird | winter | WA | Garfield | 1 | cropland | R1WA-c-2 | 2017-03-27T00:00:00Z | 13.9611111 | 20.2194444 | 0 | R1WA-c-2 |
| ring-necked pheasant | Game Bird | winter | WA | Garfield | 1 | cropland | R1WA-c-2 | 2017-03-27T00:00:00Z | 1.9520833 | 2.8979167 | 0 | R1WA-c-2 |
| ring-necked pheasant | Game Bird | winter | WA | Garfield | 1 | cropland | R1WA-c-2 | 2017-03-27T00:00:00Z | 3.9208333 | 6.9277778 | 0 | R1WA-c-2 |
| mallard | Game Bird | winter | WA | Garfield | 1 | cropland | R1WA-c-2 | 2017-03-27T00:00:00Z | 1.8944444 | 2.8791667 | 0 | R1WA-c-2 |
| ring-necked pheasant | Game Bird | winter | WA | Garfield | 1 | cropland | R1WA-c-2 | 2017-03-27T00:00:00Z | 30.1347222 | NA | 0 | R1WA-c-2 |
| ring-necked pheasant | Game Bird | winter | WA | Garfield | 1 | cropland | R1WA-c-2 | 2017-03-27T00:00:00Z | 20.0965278 | 30.1055556 | 0 | R1WA-c-2 |
| ring-necked pheasant | Game Bird | winter | WA | Garfield | 1 | cropland | R1WA-c-2 | 2017-03-27T00:00:00Z | 30.3548611 | NA | 0 | R1WA-c-2 |
| ring-necked pheasant | Game Bird | winter | WA | Garfield | 1 | cropland | R1WA-c-2 | 2017-03-27T00:00:00Z | 30.2041667 | NA | 0 | R1WA-c-2 |
| mallard | Game Bird | winter | WA | Garfield | 1 | cropland | R1WA-c-2 | 2017-03-27T00:00:00Z | 30.1395833 | NA | 0 | R1WA-c-2 |
| ring-necked pheasant | Game Bird | winter | WA | Garfield | 1 | cropland | R1WA-c-2 | 2017-03-27T00:00:00Z | 1.1819444 | 1.9680556 | 0 | R1WA-c-2 |
| ring-necked pheasant | Game Bird | winter | WA | Garfield | 1 | cropland | R1WA-c-2 | 2017-03-27T00:00:00Z | 3.9020833 | 6.9 | 0 | R1WA-c-2 |
| Cooper's hawk | Raptor | winter | WA | Garfield | 1 | cropland | R1WA-c-3 | 2017-01-23T00:00:00Z | 7 | 10 | 0 | R1WA-c-3 |
| osprey | Raptor | winter | WA | Garfield | 1 | cropland | R1WA-c-3 | 2017-01-23T00:00:00Z | 50 | 60 | 0 | R1WA-c-3 |
| Cooper's hawk | Raptor | winter | WA | Garfield | 1 | cropland | R1WA-c-3 | 2017-01-23T00:00:00Z | 1 | 1 | 0 | R1WA-c-3 |
| Cooper's hawk | Raptor | winter | WA | Garfield | 1 | cropland | R1WA-c-3 | 2017-01-23T00:00:00Z | 110 | NA | 0 | R1WA-c-3 |
| barn owl | Raptor | winter | WA | Garfield | 1 | cropland | R1WA-c-3 | 2017-01-23T00:00:00Z | 110 | NA | 0 | R1WA-c-3 |
| Cooper's hawk | Raptor | winter | WA | Garfield | 1 | cropland | R1WA-c-3 | 2017-01-23T00:00:00Z | 120 | NA | 0 | R1WA-c-3 |
| Cooper's hawk | Raptor | winter | WA | Garfield | 1 | cropland | R1WA-c-3 | 2017-01-23T00:00:00Z | 120 | NA | 0 | R1WA-c-3 |
| turkey vulture | Raptor | winter | WA | Garfield | 1 | cropland | R1WA-c-3 | 2017-01-23T00:00:00Z | 120 | NA | 0 | R1WA-c-3 |
| Cooper's hawk | Raptor | winter | WA | Garfield | 1 | cropland | R1WA-c-3 | 2017-01-23T00:00:00Z | 120 | NA | 0 | R1WA-c-3 |
| barn owl | Raptor | winter | WA | Garfield | 1 | cropland | R1WA-c-3 | 2017-01-23T00:00:00Z | 110 | NA | 0 | R1WA-c-3 |
| red-tailed hawk | Raptor | winter | WA | Garfield | 1 | cropland | R1WA-c-3 | 2017-01-23T00:00:00Z | 110 | NA | 0 | R1WA-c-3 |
| Cooper's hawk | Raptor | winter | WA | Garfield | 1 | cropland | R1WA-c-3 | 2017-01-23T00:00:00Z | 14 | 20 | 0 | R1WA-c-3 |
| red-tailed hawk | Raptor | winter | WA | Garfield | 1 | cropland | R1WA-c-3 | 2017-01-23T00:00:00Z | 120 | NA | 0 | R1WA-c-3 |
| short-eared owl | Raptor | winter | WA | Garfield | 1 | cropland | R1WA-c-3 | 2017-01-23T00:00:00Z | 30 | 40 | 0 | R1WA-c-3 |
| Swainson's hawk | Raptor | winter | WA | Garfield | 1 | cropland | R1WA-c-3 | 2017-01-23T00:00:00Z | 120 | NA | 0 | R1WA-c-3 |
| Swainson's hawk | Raptor | winter | WA | Garfield | 1 | cropland | R1WA-c-3 | 2017-01-23T00:00:00Z | 80 | 90 | 0 | R1WA-c-3 |
| Cooper's hawk | Raptor | winter | WA | Garfield | 1 | cropland | R1WA-c-3 | 2017-01-23T00:00:00Z | 4 | 7 | 0 | R1WA-c-3 |
| Cooper's hawk | Raptor | winter | WA | Garfield | 1 | cropland | R1WA-c-3 | 2017-01-23T00:00:00Z | 110 | NA | 0 | R1WA-c-3 |
| Cooper's hawk | Raptor | winter | WA | Garfield | 1 | cropland | R1WA-c-3 | 2017-01-23T00:00:00Z | 20 | 30 | 0 | R1WA-c-3 |
| red-tailed hawk | Raptor | winter | WA | Garfield | 1 | cropland | R1WA-c-3 | 2017-01-23T00:00:00Z | 120 | NA | 0 | R1WA-c-3 |
| Cooper's hawk | Raptor | winter | WA | Garfield | 1 | cropland | R1WA-c-3 | 2017-01-23T00:00:00Z | 7 | 10 | 0 | R1WA-c-3 |
| Cooper's hawk | Raptor | winter | WA | Garfield | 1 | cropland | R1WA-c-3 | 2017-01-23T00:00:00Z | 1 | 1 | 0 | R1WA-c-3 |
| osprey | Raptor | winter | WA | Garfield | 1 | cropland | R1WA-c-3 | 2017-01-23T00:00:00Z | 120 | NA | 0 | R1WA-c-3 |
| Cooper's hawk | Raptor | winter | WA | Garfield | 1 | cropland | R1WA-c-3 | 2017-01-23T00:00:00Z | 7 | 10 | 0 | R1WA-c-3 |
| red-tailed hawk | Raptor | winter | WA | Garfield | 1 | cropland | R1WA-c-3 | 2017-01-23T00:00:00Z | 1 | 1 | 0 | R1WA-c-3 |
| Cooper's hawk | Raptor | winter | WA | Garfield | 1 | cropland | R1WA-c-3 | 2017-01-23T00:00:00Z | 2 | 4 | 0 | R1WA-c-3 |
| barn owl | Raptor | winter | WA | Garfield | 1 | cropland | R1WA-c-3 | 2017-01-23T00:00:00Z | 7 | 10 | 0 | R1WA-c-3 |
| red-tailed hawk | Raptor | winter | WA | Garfield | 1 | cropland | R1WA-c-3 | 2017-01-23T00:00:00Z | 80 | 90 | 0 | R1WA-c-3 |
| great horned owl | Raptor | spring | WA | Garfield | 1 | cropland | R1WA-c-3 | 2017-05-29T00:00:00Z | 119.8729167 | NA | 0 | R1WA-c-3 |
| Cooper's hawk | Raptor | spring | WA | Garfield | 1 | cropland | R1WA-c-3 | 2017-05-29T00:00:00Z | 119.875 | NA | 0 | R1WA-c-3 |
| barn owl | Raptor | spring | WA | Garfield | 1 | cropland | R1WA-c-3 | 2017-05-29T00:00:00Z | 3.1041667 | 4.0770833 | 0 | R1WA-c-3 |
| prairie falcon | Raptor | spring | WA | Garfield | 1 | cropland | R1WA-c-3 | 2017-05-29T00:00:00Z | 4.0701389 | 6.9333333 | 0 | R1WA-c-3 |
| great horned owl | Raptor | spring | WA | Garfield | 1 | cropland | R1WA-c-3 | 2017-05-29T00:00:00Z | 119.8555556 | NA | 0 | R1WA-c-3 |
| red-tailed hawk | Raptor | spring | WA | Garfield | 1 | cropland | R1WA-c-3 | 2017-05-29T00:00:00Z | 119.8569444 | NA | 0 | R1WA-c-3 |
| barn owl | Raptor | spring | WA | Garfield | 1 | cropland | R1WA-c-3 | 2017-05-29T00:00:00Z | 119.8618056 | NA | 0 | R1WA-c-3 |
| red-tailed hawk | Raptor | spring | WA | Garfield | 1 | cropland | R1WA-c-3 | 2017-05-29T00:00:00Z | 119.8638889 | NA | 0 | R1WA-c-3 |
| Swainson's hawk | Raptor | spring | WA | Garfield | 1 | cropland | R1WA-c-3 | 2017-05-29T00:00:00Z | 119.7763889 | NA | 0 | R1WA-c-3 |
| rough-legged hawk | Raptor | spring | WA | Garfield | 1 | cropland | R1WA-c-3 | 2017-05-29T00:00:00Z | 119.8506944 | NA | 0 | R1WA-c-3 |
| Cooper's hawk | Raptor | spring | WA | Garfield | 1 | cropland | R1WA-c-3 | 2017-05-29T00:00:00Z | 109.8798611 | 119.9638889 | 0 | R1WA-c-3 |
| great horned owl | Raptor | spring | WA | Garfield | 1 | cropland | R1WA-c-3 | 2017-05-29T00:00:00Z | 119.9520833 | NA | 0 | R1WA-c-3 |
| red-tailed hawk | Raptor | spring | WA | Garfield | 1 | cropland | R1WA-c-3 | 2017-05-29T00:00:00Z | 119.9993056 | NA | 0 | R1WA-c-3 |
| Cooper's hawk | Raptor | spring | WA | Garfield | 1 | cropland | R1WA-c-3 | 2017-05-29T00:00:00Z | 120.0381944 | NA | 0 | R1WA-c-3 |
| barn owl | Raptor | spring | WA | Garfield | 1 | cropland | R1WA-c-3 | 2017-05-29T00:00:00Z | 119.7736111 | NA | 0 | R1WA-c-3 |
| barn owl | Raptor | spring | WA | Garfield | 1 | cropland | R1WA-c-3 | 2017-05-29T00:00:00Z | 119.85 | NA | 0 | R1WA-c-3 |
| ferruginous hawk | Raptor | spring | WA | Garfield | 1 | cropland | R1WA-c-3 | 2017-05-29T00:00:00Z | 120 | NA | 0 | R1WA-c-3 |
| short-eared owl | Raptor | spring | WA | Garfield | 1 | cropland | R1WA-c-3 | 2017-05-29T00:00:00Z | 119.9277778 | NA | 0 | R1WA-c-3 |
| great horned owl | Raptor | spring | WA | Garfield | 1 | cropland | R1WA-c-3 | 2017-05-29T00:00:00Z | 119.8854167 | NA | 0 | R1WA-c-3 |
| rough-legged hawk | Raptor | spring | WA | Garfield | 1 | cropland | R1WA-c-3 | 2017-05-29T00:00:00Z | 119.8729167 | NA | 0 | R1WA-c-3 |
| red-tailed hawk | Raptor | spring | WA | Garfield | 1 | cropland | R1WA-c-3 | 2017-05-29T00:00:00Z | 119.8548611 | NA | 0 | R1WA-c-3 |
| barn owl | Raptor | spring | WA | Garfield | 1 | cropland | R1WA-c-3 | 2017-05-29T00:00:00Z | 119.8229167 | NA | 0 | R1WA-c-3 |
| red-tailed hawk | Raptor | spring | WA | Garfield | 1 | cropland | R1WA-c-3 | 2017-05-29T00:00:00Z | 119.8243056 | NA | 0 | R1WA-c-3 |
| Cooper's hawk | Raptor | spring | WA | Garfield | 1 | cropland | R1WA-c-3 | 2017-05-29T00:00:00Z | 119.81875 | NA | 0 | R1WA-c-3 |
| prairie falcon | Raptor | spring | WA | Garfield | 1 | cropland | R1WA-c-3 | 2017-05-29T00:00:00Z | 69.7902778 | 79.8055556 | 0 | R1WA-c-3 |
| great horned owl | Raptor | fall | WA | Garfield | 1 | cropland | R1WA-c-3 | 2017-10-01T00:00:00Z | 3.8402778 | 6.8875 | 0 | R1WA-c-3 |
| northern goshawk | Raptor | fall | WA | Garfield | 1 | cropland | R1WA-c-3 | 2017-10-01T00:00:00Z | 6.8875 | 9.81875 | 0 | R1WA-c-3 |
| great horned owl | Raptor | fall | WA | Garfield | 1 | cropland | R1WA-c-3 | 2017-10-01T00:00:00Z | 1.6145833 | 2.6652778 | 0 | R1WA-c-3 |
| northern goshawk | Raptor | fall | WA | Garfield | 1 | cropland | R1WA-c-3 | 2017-10-01T00:00:00Z | 9.6666667 | 13.9333333 | 0 | R1WA-c-3 |
| northern harrier | Raptor | fall | WA | Garfield | 1 | cropland | R1WA-c-3 | 2017-10-01T00:00:00Z | 14.0319444 | 19.8138889 | 0 | R1WA-c-3 |
| great horned owl | Raptor | fall | WA | Garfield | 1 | cropland | R1WA-c-3 | 2017-10-01T00:00:00Z | 19.8166667 | 29.95 | 0 | R1WA-c-3 |
| broad-winged hawk | Raptor | fall | WA | Garfield | 1 | cropland | R1WA-c-3 | 2017-10-01T00:00:00Z | 49.7631944 | 59.8895833 | 0 | R1WA-c-3 |
| northern goshawk | Raptor | fall | WA | Garfield | 1 | cropland | R1WA-c-3 | 2017-10-01T00:00:00Z | 19.825 | 29.8805556 | 0 | R1WA-c-3 |
| Swainson's hawk | Raptor | fall | WA | Garfield | 1 | cropland | R1WA-c-3 | 2017-10-01T00:00:00Z | 49.8680556 | 60.2069444 | 0 | R1WA-c-3 |
| great horned owl | Raptor | fall | WA | Garfield | 1 | cropland | R1WA-c-3 | 2017-10-01T00:00:00Z | 29.8965278 | 39.5965278 | 0 | R1WA-c-3 |
| northern goshawk | Raptor | fall | WA | Garfield | 1 | cropland | R1WA-c-3 | 2017-10-01T00:00:00Z | 49.7604167 | 59.9166667 | 0 | R1WA-c-3 |
| northern goshawk | Raptor | fall | WA | Garfield | 1 | cropland | R1WA-c-3 | 2017-10-01T00:00:00Z | 3.8090278 | 6.8875 | 0 | R1WA-c-3 |
| barn owl | Raptor | fall | WA | Garfield | 1 | cropland | R1WA-c-3 | 2017-10-01T00:00:00Z | 29.9444444 | 39.7555556 | 0 | R1WA-c-3 |
| barn owl | Raptor | fall | WA | Garfield | 1 | cropland | R1WA-c-3 | 2017-10-01T00:00:00Z | 6.8819444 | 9.7923611 | 0 | R1WA-c-3 |
| northern goshawk | Raptor | fall | WA | Garfield | 1 | cropland | R1WA-c-3 | 2017-10-01T00:00:00Z | 69.9625 | NA | 0 | R1WA-c-3 |
| Swainson's hawk | Raptor | fall | WA | Garfield | 1 | cropland | R1WA-c-3 | 2017-10-01T00:00:00Z | 2.7520833 | 3.8076389 | 0 | R1WA-c-3 |
| great horned owl | Raptor | fall | WA | Garfield | 1 | cropland | R1WA-c-3 | 2017-10-01T00:00:00Z | 69.9909722 | NA | 0 | R1WA-c-3 |
| barn owl | Raptor | fall | WA | Garfield | 1 | cropland | R1WA-c-3 | 2017-10-01T00:00:00Z | 3.6208333 | 6.8270833 | 0 | R1WA-c-3 |
| northern goshawk | Raptor | fall | WA | Garfield | 1 | cropland | R1WA-c-3 | 2017-10-01T00:00:00Z | 69.9409722 | NA | 0 | R1WA-c-3 |
| barn owl | Raptor | fall | WA | Garfield | 1 | cropland | R1WA-c-3 | 2017-10-01T00:00:00Z | 69.9381944 | NA | 0 | R1WA-c-3 |
| northern goshawk | Raptor | fall | WA | Garfield | 1 | cropland | R1WA-c-3 | 2017-10-01T00:00:00Z | 29.8513889 | 39.5951389 | 0 | R1WA-c-3 |
| red-tailed hawk | Raptor | spring | WA | Columbia | 1 | cropland | R1WA-c-4 | 2018-03-12T00:00:00Z | 87.1069444 | NA | 1 | R1WA-c-4 |
| mallard | Game Bird | spring | WA | Columbia | 1 | cropland | R1WA-c-4 | 2018-03-12T00:00:00Z | 2.2770833 | 3.15625 | 1 | R1WA-c-4 |
| barn owl | Raptor | spring | WA | Columbia | 1 | cropland | R1WA-c-4 | 2018-03-12T00:00:00Z | 87.06875 | NA | 1 | R1WA-c-4 |
| mallard | Game Bird | spring | WA | Columbia | 1 | cropland | R1WA-c-4 | 2018-03-12T00:00:00Z | 6.90625 | 10.1541667 | 1 | R1WA-c-4 |
| red-tailed hawk | Raptor | spring | WA | Columbia | 1 | cropland | R1WA-c-4 | 2018-03-12T00:00:00Z | 42.0027778 | 49.1652778 | 1 | R1WA-c-4 |
| ring-necked pheasant | Game Bird | spring | WA | Columbia | 1 | cropland | R1WA-c-4 | 2018-03-12T00:00:00Z | 86.9888889 | NA | 1 | R1WA-c-4 |
| Swainson's hawk | Raptor | spring | WA | Columbia | 1 | cropland | R1WA-c-4 | 2018-03-12T00:00:00Z | 86.9354167 | NA | 1 | R1WA-c-4 |
| Cooper's hawk | Raptor | spring | WA | Columbia | 1 | cropland | R1WA-c-4 | 2018-03-12T00:00:00Z | 6.8277778 | 10.0381944 | 1 | R1WA-c-4 |
| ring-necked pheasant | Game Bird | spring | WA | Columbia | 1 | cropland | R1WA-c-4 | 2018-03-12T00:00:00Z | 14.075 | 21.1465278 | 1 | R1WA-c-4 |
| barn owl | Raptor | summer | WA | Columbia | 1 | cropland | R1WA-c-4 | 2018-05-14T00:00:00Z | 87.0888889 | NA | 1 | R1WA-c-4 |
| Swainson's hawk | Raptor | summer | WA | Columbia | 1 | cropland | R1WA-c-4 | 2018-05-14T00:00:00Z | 87.00625 | NA | 1 | R1WA-c-4 |
| red-tailed hawk | Raptor | summer | WA | Columbia | 1 | cropland | R1WA-c-4 | 2018-05-14T00:00:00Z | 86.9805556 | NA | 1 | R1WA-c-4 |
| mallard | Game Bird | summer | WA | Columbia | 1 | cropland | R1WA-c-4 | 2018-05-14T00:00:00Z | 34.8590278 | 41.8965278 | 1 | R1WA-c-4 |
| great horned owl | Raptor | summer | WA | Columbia | 1 | cropland | R1WA-c-4 | 2018-05-14T00:00:00Z | 21.1909722 | 27.9006944 | 1 | R1WA-c-4 |
| Swainson's hawk | Raptor | summer | WA | Columbia | 1 | cropland | R1WA-c-4 | 2018-05-14T00:00:00Z | 21.1944444 | 27.9041667 | 1 | R1WA-c-4 |
| ring-necked pheasant | Game Bird | summer | WA | Columbia | 1 | cropland | R1WA-c-4 | 2018-05-14T00:00:00Z | 0.9527778 | 1.9055556 | 1 | R1WA-c-4 |
| ring-necked pheasant | Game Bird | summer | WA | Columbia | 1 | cropland | R1WA-c-4 | 2018-05-14T00:00:00Z | 0.9381944 | 0.9381944 | 1 | R1WA-c-4 |
| ring-necked pheasant | Game Bird | summer | WA | Columbia | 1 | cropland | R1WA-c-4 | 2018-05-14T00:00:00Z | 3.9680556 | 6.8125 | 1 | R1WA-c-4 |
| mallard | Game Bird | summer | WA | Columbia | 1 | cropland | R1WA-c-4 | 2018-05-14T00:00:00Z | 1.9201389 | 4.0201389 | 1 | R1WA-c-4 |
| great horned owl | Raptor | fall | WA | Columbia | 1 | cropland | R1WA-c-4 | 2018-08-13T00:00:00Z | 87.1881944 | NA | 1 | R1WA-c-4 |
| rough-legged hawk | Raptor | fall | WA | Columbia | 1 | cropland | R1WA-c-4 | 2018-08-13T00:00:00Z | 56.1444444 | 63.3090278 | 1 | R1WA-c-4 |
| ring-necked pheasant | Game Bird | fall | WA | Columbia | 1 | cropland | R1WA-c-4 | 2018-08-13T00:00:00Z | 48.9888889 | 55.96875 | 1 | R1WA-c-4 |
| rough-legged hawk | Raptor | fall | WA | Columbia | 1 | cropland | R1WA-c-4 | 2018-08-13T00:00:00Z | 14.1173611 | 21.1215278 | 1 | R1WA-c-4 |
| osprey | Raptor | fall | WA | Columbia | 1 | cropland | R1WA-c-4 | 2018-08-13T00:00:00Z | 20.9715278 | 28.0611111 | 1 | R1WA-c-4 |
| ring-necked pheasant | Game Bird | fall | WA | Columbia | 1 | cropland | R1WA-c-4 | 2018-08-13T00:00:00Z | 7.2236111 | 10.1541667 | 1 | R1WA-c-4 |
| barn owl | Raptor | fall | WA | Columbia | 1 | cropland | R1WA-c-4 | 2018-08-13T00:00:00Z | 3.8104167 | 7.2145833 | 1 | R1WA-c-4 |
| mallard | Game Bird | fall | WA | Columbia | 1 | cropland | R1WA-c-4 | 2018-08-13T00:00:00Z | 2.8645833 | 3.8152778 | 1 | R1WA-c-4 |
| mallard | Game Bird | fall | WA | Columbia | 1 | cropland | R1WA-c-4 | 2018-08-13T00:00:00Z | 0.9381944 | 1.9770833 | 1 | R1WA-c-4 |
| ring-necked pheasant | Game Bird | winter | WA | Columbia | 1 | cropland | R1WA-c-4 | 2018-10-29T00:00:00Z | 21.0486111 | 28.0388889 | 1 | R1WA-c-4 |
| great horned owl | Raptor | winter | WA | Columbia | 1 | cropland | R1WA-c-4 | 2018-10-29T00:00:00Z | 21.0881944 | 28.1520833 | 1 | R1WA-c-4 |
| ring-necked pheasant | Game Bird | winter | WA | Columbia | 1 | cropland | R1WA-c-4 | 2018-10-29T00:00:00Z | 4.0951389 | 6.9222222 | 1 | R1WA-c-4 |
| great horned owl | Raptor | winter | WA | Columbia | 1 | cropland | R1WA-c-4 | 2018-10-29T00:00:00Z | 6.9256944 | 10.1069444 | 1 | R1WA-c-4 |
| barn owl | Raptor | winter | WA | Columbia | 1 | cropland | R1WA-c-4 | 2018-10-29T00:00:00Z | 1.2361111 | 2.0201389 | 1 | R1WA-c-4 |
| great horned owl | Raptor | winter | WA | Columbia | 1 | cropland | R1WA-c-4 | 2018-10-29T00:00:00Z | 6.9208333 | 9.9951389 | 1 | R1WA-c-4 |
| ring-necked pheasant | Game Bird | winter | WA | Columbia | 1 | cropland | R1WA-c-4 | 2018-10-29T00:00:00Z | 2.7798611 | 3.8076389 | 1 | R1WA-c-4 |
| mallard | Game Bird | winter | WA | Columbia | 1 | cropland | R1WA-c-4 | 2018-10-29T00:00:00Z | 1.1847222 | 1.8152778 | 1 | R1WA-c-4 |
| red-tailed hawk | Raptor | winter | WA | Columbia | 1 | cropland | R1WA-c-4 | 2018-10-29T00:00:00Z | 41.9715278 | 48.7493056 | 1 | R1WA-c-4 |
| red-tailed hawk | Raptor | winter | IA | Adair | 3 | cropland | R3IA-c-1 | 2020-12-07T00:00:00Z | 0.9784722 | 0.9784722 | 0 | R3IA-c-1 |
| red-tailed hawk | Raptor | winter | IA | Adair | 3 | cropland | R3IA-c-1 | 2020-12-07T00:00:00Z | 29.9590278 | 60.0423611 | 0 | R3IA-c-1 |
| red-tailed hawk | Raptor | winter | IA | Adair | 3 | cropland | R3IA-c-1 | 2020-12-07T00:00:00Z | 2.8666667 | 4.1201389 | 0 | R3IA-c-1 |
| red-tailed hawk | Raptor | winter | IA | Adair | 3 | cropland | R3IA-c-1 | 2020-12-07T00:00:00Z | 9.9305556 | 13.9861111 | 0 | R3IA-c-1 |
| red-tailed hawk | Raptor | winter | IA | Adair | 3 | cropland | R3IA-c-1 | 2020-12-07T00:00:00Z | 4.2201389 | 13.9763889 | 0 | R3IA-c-1 |
| red-tailed hawk | Raptor | winter | IA | Adair | 3 | cropland | R3IA-c-1 | 2021-02-22T00:00:00Z | 9.9819444 | 14.0451389 | 0 | R3IA-c-1 |
| red-tailed hawk | Raptor | winter | IA | Adair | 3 | cropland | R3IA-c-1 | 2021-02-22T00:00:00Z | 2.8173611 | 3.8923611 | 0 | R3IA-c-1 |
| red-tailed hawk | Raptor | winter | IA | Adair | 3 | cropland | R3IA-c-1 | 2021-02-22T00:00:00Z | 1.0875 | 1.8923611 | 0 | R3IA-c-1 |
| ring-necked pheasant | Game Bird | winter | IA | Boone and Greene | 3 | cropland | R3IA-c-2 | 2019-01-31T00:00:00Z | 6.9416667 | 29.7694444 | 0 | R3IA-c-2 |
| red-tailed hawk | Raptor | spring | IA | Boone and Greene | 3 | cropland | R3IA-c-2 | 2018-03-19T00:00:00Z | 19.8840278 | 30.7729167 | 0 | R3IA-c-2 |
| red-tailed hawk | Raptor | spring | IA | Boone and Greene | 3 | cropland | R3IA-c-2 | 2018-03-19T00:00:00Z | 14.2 | 29.9236111 | 0 | R3IA-c-2 |
| red-tailed hawk | Raptor | spring | IA | Boone and Greene | 3 | cropland | R3IA-c-2 | 2018-03-19T00:00:00Z | 29.9194444 | 39.8201389 | 0 | R3IA-c-2 |
| red-tailed hawk | Raptor | spring | IA | Boone and Greene | 3 | cropland | R3IA-c-2 | 2018-03-19T00:00:00Z | 13.9979167 | 19.9201389 | 0 | R3IA-c-2 |
| red-tailed hawk | Raptor | spring | IA | Boone and Greene | 3 | cropland | R3IA-c-2 | 2018-03-19T00:00:00Z | 59.7611111 | NA | 0 | R3IA-c-2 |
| red-tailed hawk | Raptor | spring | IA | Boone and Greene | 3 | cropland | R3IA-c-2 | 2018-03-26T00:00:00Z | 7.0631944 | 10.1243056 | 0 | R3IA-c-2 |
| red-tailed hawk | Raptor | spring | IA | Boone and Greene | 3 | cropland | R3IA-c-2 | 2018-03-26T00:00:00Z | 10.0923611 | 13.9861111 | 0 | R3IA-c-2 |
| red-tailed hawk | Raptor | spring | IA | Boone and Greene | 3 | cropland | R3IA-c-2 | 2018-03-26T00:00:00Z | 60.1458333 | NA | 0 | R3IA-c-2 |
| red-tailed hawk | Raptor | spring | IA | Boone and Greene | 3 | cropland | R3IA-c-2 | 2018-03-26T00:00:00Z | 60.3902778 | NA | 0 | R3IA-c-2 |
| red-tailed hawk | Raptor | spring | IA | Boone and Greene | 3 | cropland | R3IA-c-2 | 2018-03-26T00:00:00Z | 9.975 | 19.7506944 | 0 | R3IA-c-2 |
| red-tailed hawk | Raptor | winter | IA | Boone and Greene | 3 | cropland | R3IA-c-2 | 2018-11-20T00:00:00Z | 49.7895833 | 60.03125 | 0 | R3IA-c-2 |
| red-tailed hawk | Raptor | winter | IA | Boone and Greene | 3 | cropland | R3IA-c-2 | 2018-11-20T00:00:00Z | 49.7763889 | 60.0180556 | 0 | R3IA-c-2 |
| red-tailed hawk | Raptor | winter | IA | Boone and Greene | 3 | cropland | R3IA-c-2 | 2018-11-20T00:00:00Z | 2.7909722 | 6.9375 | 0 | R3IA-c-2 |
| red-tailed hawk | Raptor | winter | IA | Boone and Greene | 3 | cropland | R3IA-c-2 | 2018-11-20T00:00:00Z | 0.8055556 | 2.7243056 | 0 | R3IA-c-2 |
| red-tailed hawk | Raptor | winter | IA | Boone and Greene | 3 | cropland | R3IA-c-2 | 2018-11-20T00:00:00Z | 2.6951389 | 6.8194444 | 0 | R3IA-c-2 |
| red-tailed hawk | Raptor | winter | IA | Boone and Greene | 3 | cropland | R3IA-c-2 | 2018-12-10T00:00:00Z | 59.8888889 | NA | 0 | R3IA-c-2 |
| red-tailed hawk | Raptor | winter | IA | Boone and Greene | 3 | cropland | R3IA-c-2 | 2018-12-10T00:00:00Z | 29.7458333 | 39.9659722 | 0 | R3IA-c-2 |
| red-tailed hawk | Raptor | winter | IA | Boone and Greene | 3 | cropland | R3IA-c-2 | 2018-12-10T00:00:00Z | 19.9840278 | 29.7118056 | 0 | R3IA-c-2 |
| red-tailed hawk | Raptor | winter | IA | Boone and Greene | 3 | cropland | R3IA-c-2 | 2018-12-10T00:00:00Z | 20.0472222 | 29.71875 | 0 | R3IA-c-2 |
| red-tailed hawk | Raptor | winter | IA | Boone and Greene | 3 | cropland | R3IA-c-2 | 2018-12-10T00:00:00Z | 59.9840278 | NA | 0 | R3IA-c-2 |
| red-tailed hawk | Raptor | winter | IA | Boone and Greene | 3 | cropland | R3IA-c-2 | 2019-01-08T00:00:00Z | 30.0451389 | 63.9381944 | 0 | R3IA-c-2 |
| red-tailed hawk | Raptor | winter | IA | Boone and Greene | 3 | cropland | R3IA-c-2 | 2019-01-08T00:00:00Z | 10.0833333 | 62.8972222 | 0 | R3IA-c-2 |
| red-tailed hawk | Raptor | winter | IA | Boone and Greene | 3 | cropland | R3IA-c-2 | 2019-01-08T00:00:00Z | 2.9673611 | 6.9708333 | 0 | R3IA-c-2 |
| red-tailed hawk | Raptor | winter | IA | Boone and Greene | 3 | cropland | R3IA-c-2 | 2019-01-08T00:00:00Z | 61.8722222 | NA | 0 | R3IA-c-2 |
| red-tailed hawk | Raptor | winter | IA | Boone and Greene | 3 | cropland | R3IA-c-2 | 2019-01-08T00:00:00Z | 10.0583333 | 63.9236111 | 0 | R3IA-c-2 |
| red-tailed hawk | Raptor | spring | IA | Greene | 3 | cropland | R3IA-c-3 | 2019-03-18T00:00:00Z | 59.8076389 | NA | 0 | R3IA-c-3 |
| red-tailed hawk | Raptor | spring | IA | Greene | 3 | cropland | R3IA-c-3 | 2019-03-18T00:00:00Z | 59.9847222 | NA | 0 | R3IA-c-3 |
| red-tailed hawk | Raptor | spring | IA | Greene | 3 | cropland | R3IA-c-3 | 2019-03-18T00:00:00Z | 59.9395833 | NA | 0 | R3IA-c-3 |
| red-tailed hawk | Raptor | spring | IA | Greene | 3 | cropland | R3IA-c-3 | 2019-03-18T00:00:00Z | 59.9041667 | NA | 0 | R3IA-c-3 |
| red-tailed hawk | Raptor | spring | IA | Greene | 3 | cropland | R3IA-c-3 | 2019-03-18T00:00:00Z | 59.8597222 | NA | 0 | R3IA-c-3 |
| short-eared owl | Raptor | spring | IA | Greene | 3 | cropland | R3IA-c-3 | 2019-03-22T00:00:00Z | 9.9458333 | 14.0298611 | 0 | R3IA-c-3 |
| red-tailed hawk | Raptor | spring | IA | Greene | 3 | cropland | R3IA-c-3 | 2019-03-25T00:00:00Z | 59.9534722 | NA | 0 | R3IA-c-3 |
| red-tailed hawk | Raptor | spring | IA | Greene | 3 | cropland | R3IA-c-3 | 2019-03-25T00:00:00Z | 6.9701389 | 10.1930556 | 0 | R3IA-c-3 |
| red-tailed hawk | Raptor | spring | IA | Greene | 3 | cropland | R3IA-c-3 | 2019-03-25T00:00:00Z | 59.9020833 | NA | 0 | R3IA-c-3 |
| red-tailed hawk | Raptor | spring | IA | Greene | 3 | cropland | R3IA-c-3 | 2019-03-25T00:00:00Z | 59.9104167 | NA | 0 | R3IA-c-3 |
| red-tailed hawk | Raptor | spring | IA | Greene | 3 | cropland | R3IA-c-3 | 2019-03-25T00:00:00Z | 2.94375 | 6.9597222 | 0 | R3IA-c-3 |
| red-tailed hawk | Raptor | winter | IA | Greene | 3 | cropland | R3IA-c-3 | 2019-11-26T00:00:00Z | 60.0423611 | NA | 0 | R3IA-c-3 |
| red-tailed hawk | Raptor | winter | IA | Greene | 3 | cropland | R3IA-c-3 | 2019-11-26T00:00:00Z | 41.1173611 | 60.0444444 | 0 | R3IA-c-3 |
| red-tailed hawk | Raptor | winter | IA | Greene | 3 | cropland | R3IA-c-3 | 2019-11-26T00:00:00Z | 60.04375 | NA | 0 | R3IA-c-3 |
| red-tailed hawk | Raptor | winter | IA | Greene | 3 | cropland | R3IA-c-3 | 2019-11-26T00:00:00Z | 10.1777778 | 20.3333333 | 0 | R3IA-c-3 |
| red-tailed hawk | Raptor | winter | IA | Greene | 3 | cropland | R3IA-c-3 | 2019-11-26T00:00:00Z | 41.2569444 | 60.0375 | 0 | R3IA-c-3 |
| red-tailed hawk | Raptor | winter | IA | Greene | 3 | cropland | R3IA-c-3 | 2019-12-10T00:00:00Z | 60.1118056 | NA | 0 | R3IA-c-3 |
| red-tailed hawk | Raptor | winter | IA | Greene | 3 | cropland | R3IA-c-3 | 2019-12-10T00:00:00Z | 60.1118056 | NA | 0 | R3IA-c-3 |
| red-tailed hawk | Raptor | winter | IA | Greene | 3 | cropland | R3IA-c-3 | 2019-12-10T00:00:00Z | 60.0895833 | NA | 0 | R3IA-c-3 |
| red-tailed hawk | Raptor | winter | IA | Greene | 3 | cropland | R3IA-c-3 | 2019-12-10T00:00:00Z | 60.0777778 | NA | 0 | R3IA-c-3 |
| red-tailed hawk | Raptor | winter | IA | Greene | 3 | cropland | R3IA-c-3 | 2019-12-10T00:00:00Z | 60.0569444 | NA | 0 | R3IA-c-3 |
| red-tailed hawk | Raptor | winter | IA | Greene | 3 | cropland | R3IA-c-3 | 2020-01-07T00:00:00Z | 59.8583333 | NA | 0 | R3IA-c-3 |
| red-tailed hawk | Raptor | winter | IA | Greene | 3 | cropland | R3IA-c-3 | 2020-01-07T00:00:00Z | 59.8465278 | NA | 0 | R3IA-c-3 |
| red-tailed hawk | Raptor | winter | IA | Greene | 3 | cropland | R3IA-c-3 | 2020-01-07T00:00:00Z | 59.8340278 | NA | 0 | R3IA-c-3 |
| red-tailed hawk | Raptor | winter | IA | Greene | 3 | cropland | R3IA-c-3 | 2020-01-07T00:00:00Z | 0.79375 | 0.79375 | 0 | R3IA-c-3 |
| red-tailed hawk | Raptor | winter | IA | Greene | 3 | cropland | R3IA-c-3 | 2020-01-07T00:00:00Z | 2.9930556 | 43.0930556 | 0 | R3IA-c-3 |
| great horned owl | Raptor | winter | MI | St. Clair | 3 | cropland | R3MI-c-1 | 2017-12-06T00:00:00Z | 49.975 | 68.1451389 | 1 | R3MI-c-1 |
| ring-necked pheasant | Game Bird | winter | MI | St. Clair | 3 | cropland | R3MI-c-1 | 2017-12-06T00:00:00Z | 0.9972222 | 3.9465278 | 1 | R3MI-c-1 |
| ring-necked pheasant | Game Bird | winter | MI | St. Clair | 3 | cropland | R3MI-c-1 | 2017-12-06T00:00:00Z | 3.9263889 | 37.9541667 | 1 | R3MI-c-1 |
| red-tailed hawk | Raptor | winter | MI | St. Clair | 3 | cropland | R3MI-c-1 | 2017-12-06T00:00:00Z | 0.9645833 | 3.9159722 | 1 | R3MI-c-1 |
| osprey | Raptor | winter | MI | St. Clair | 3 | cropland | R3MI-c-1 | 2017-12-06T00:00:00Z | 14.2048611 | 37.9944444 | 1 | R3MI-c-1 |
| ring-necked pheasant | Game Bird | winter | MI | St. Clair | 3 | cropland | R3MI-c-1 | 2017-12-06T00:00:00Z | 1.0270833 | 1.0270833 | 1 | R3MI-c-1 |
| great horned owl | Raptor | winter | MI | St. Clair | 3 | cropland | R3MI-c-1 | 2017-12-06T00:00:00Z | 1.0645833 | 4.0041667 | 1 | R3MI-c-1 |
| ring-necked pheasant | Game Bird | winter | MI | St. Clair | 3 | cropland | R3MI-c-1 | 2017-12-06T00:00:00Z | 1.0472222 | 3.9888889 | 1 | R3MI-c-1 |
| ring-necked pheasant | Game Bird | winter | MI | St. Clair | 3 | cropland | R3MI-c-1 | 2017-12-06T00:00:00Z | 1.1395833 | 4.08125 | 1 | R3MI-c-1 |
| barn owl | Raptor | winter | MI | St. Clair | 3 | cropland | R3MI-c-1 | 2017-12-06T00:00:00Z | 50.0986111 | 68.2701389 | 1 | R3MI-c-1 |
| ring-necked pheasant | Game Bird | winter | MI | St. Clair | 3 | cropland | R3MI-c-1 | 2017-12-06T00:00:00Z | 4.0333333 | 38.0541667 | 1 | R3MI-c-1 |
| ring-necked pheasant | Game Bird | winter | MI | St. Clair | 3 | cropland | R3MI-c-1 | 2017-12-06T00:00:00Z | 1.0923611 | 1.0923611 | 1 | R3MI-c-1 |
| ring-necked pheasant | Game Bird | winter | MI | St. Clair | 3 | cropland | R3MI-c-1 | 2017-12-12T00:00:00Z | 31.9423611 | 62.1097222 | 1 | R3MI-c-1 |
| red-tailed hawk | Raptor | winter | MI | St. Clair | 3 | cropland | R3MI-c-1 | 2017-12-12T00:00:00Z | 8.1527778 | 31.9430556 | 1 | R3MI-c-1 |
| ring-necked pheasant | Game Bird | winter | MI | St. Clair | 3 | cropland | R3MI-c-1 | 2017-12-12T00:00:00Z | 6.1625 | 31.9409722 | 1 | R3MI-c-1 |
| red-tailed hawk | Raptor | winter | MI | St. Clair | 3 | cropland | R3MI-c-1 | 2017-12-12T00:00:00Z | 31.9361111 | 31.9361111 | 1 | R3MI-c-1 |
| ring-necked pheasant | Game Bird | winter | MI | St. Clair | 3 | cropland | R3MI-c-1 | 2017-12-12T00:00:00Z | 6.1673611 | 6.1673611 | 1 | R3MI-c-1 |
| ring-necked pheasant | Game Bird | winter | MI | St. Clair | 3 | cropland | R3MI-c-1 | 2017-12-12T00:00:00Z | 62.0979167 | 62.0979167 | 1 | R3MI-c-1 |
| ring-necked pheasant | Game Bird | winter | MI | St. Clair | 3 | cropland | R3MI-c-1 | 2017-12-12T00:00:00Z | 43.9381944 | 62.1020833 | 1 | R3MI-c-1 |
| great horned owl | Raptor | winter | MI | St. Clair | 3 | cropland | R3MI-c-1 | 2017-12-12T00:00:00Z | 43.9277778 | 62.0909722 | 1 | R3MI-c-1 |
| ring-necked pheasant | Game Bird | winter | MI | St. Clair | 3 | cropland | R3MI-c-1 | 2017-12-12T00:00:00Z | 6.1631944 | 6.1631944 | 1 | R3MI-c-1 |
| ferruginous hawk | Raptor | winter | MI | St. Clair | 3 | cropland | R3MI-c-1 | 2017-12-12T00:00:00Z | 43.9194444 | 62.0861111 | 1 | R3MI-c-1 |
| ring-necked pheasant | Game Bird | winter | MI | St. Clair | 3 | cropland | R3MI-c-1 | 2017-12-12T00:00:00Z | 8.1291667 | 31.9159722 | 1 | R3MI-c-1 |
| great horned owl | Raptor | winter | MI | St. Clair | 3 | cropland | R3MI-c-1 | 2017-12-12T00:00:00Z | 43.9173611 | 62.0833333 | 1 | R3MI-c-1 |
| ring-necked pheasant | Game Bird | spring | MI | St. Clair | 3 | cropland | R3MI-c-1 | 2018-04-02T00:00:00Z | 7.0958333 | 21.1159722 | 1 | R3MI-c-1 |
| red-tailed hawk | Raptor | spring | MI | St. Clair | 3 | cropland | R3MI-c-1 | 2018-04-02T00:00:00Z | 7.0965278 | 14.0680556 | 1 | R3MI-c-1 |
| ring-necked pheasant | Game Bird | spring | MI | St. Clair | 3 | cropland | R3MI-c-1 | 2018-04-23T00:00:00Z | 6.9777778 | 13.9451389 | 1 | R3MI-c-1 |
| Swainson's hawk | Raptor | summer | MI | St. Clair | 3 | cropland | R3MI-c-1 | 2018-06-25T00:00:00Z | 60.0694444 | NA | 1 | R3MI-c-1 |
| Swainson's hawk | Raptor | fall | MI | St. Clair | 3 | cropland | R3MI-c-1 | 2018-08-24T00:00:00Z | 2.9958333 | 2.9958333 | 1 | R3MI-c-1 |
| Swainson's hawk | Raptor | spring | MI | St. Clair | 3 | cropland | R3MI-c-1 | 2018-04-23T00:00:00Z | 62.9173611 | NA | 1 | R3MI-c-1 |
| barn owl | Raptor | fall | MI | St. Clair | 3 | cropland | R3MI-c-1 | 2018-08-27T00:00:00Z | 63.0118056 | NA | 1 | R3MI-c-1 |
| snowy owl | Raptor | spring | IA | Ida | 3 | cropland | R3IA-c-4 | 2020-03-16T00:00:00Z | 1.9409722 | 7.0826389 | 0 | R3IA-c-4 |
| snowy owl | Raptor | spring | IA | Ida | 3 | cropland | R3IA-c-4 | 2020-03-16T00:00:00Z | 4.1201389 | 9.9756944 | 0 | R3IA-c-4 |
| red-tailed hawk | Raptor | spring | IA | Ida | 3 | cropland | R3IA-c-4 | 2020-03-16T00:00:00Z | 59.9708333 | NA | 0 | R3IA-c-4 |
| snowy owl | Raptor | spring | IA | Ida | 3 | cropland | R3IA-c-4 | 2020-03-16T00:00:00Z | 59.96875 | NA | 0 | R3IA-c-4 |
| snowy owl | Raptor | spring | IA | Ida | 3 | cropland | R3IA-c-4 | 2020-03-16T00:00:00Z | 14.0131944 | 19.91875 | 0 | R3IA-c-4 |
| red-tailed hawk | Raptor | spring | IA | Ida | 3 | cropland | R3IA-c-4 | 2020-03-23T00:00:00Z | 59.85625 | NA | 0 | R3IA-c-4 |
| red-tailed hawk | Raptor | spring | IA | Ida | 3 | cropland | R3IA-c-4 | 2020-03-23T00:00:00Z | 59.8840278 | NA | 0 | R3IA-c-4 |
| red-tailed hawk | Raptor | spring | IA | Ida | 3 | cropland | R3IA-c-4 | 2020-03-23T00:00:00Z | 59.8680556 | NA | 0 | R3IA-c-4 |
| red-tailed hawk | Raptor | spring | IA | Ida | 3 | cropland | R3IA-c-4 | 2020-03-23T00:00:00Z | 31.1722222 | 43.125 | 0 | R3IA-c-4 |
| red-tailed hawk | Raptor | spring | IA | Ida | 3 | cropland | R3IA-c-4 | 2020-03-23T00:00:00Z | 14.1881944 | 21.1763889 | 0 | R3IA-c-4 |
| red-tailed hawk | Raptor | winter | IA | Ida | 3 | cropland | R3IA-c-4 | 2020-12-07T00:00:00Z | 60.30625 | NA | 0 | R3IA-c-4 |
| red-tailed hawk | Raptor | winter | IA | Ida | 3 | cropland | R3IA-c-4 | 2020-12-07T00:00:00Z | 30.0194444 | 50.3569444 | 0 | R3IA-c-4 |
| red-tailed hawk | Raptor | winter | IA | Ida | 3 | cropland | R3IA-c-4 | 2020-12-07T00:00:00Z | 30.01875 | 60.3770833 | 0 | R3IA-c-4 |
| red-tailed hawk | Raptor | winter | IA | Ida | 3 | cropland | R3IA-c-4 | 2020-12-07T00:00:00Z | 40.3340278 | 60.3756944 | 0 | R3IA-c-4 |
| red-tailed hawk | Raptor | winter | IA | Ida | 3 | cropland | R3IA-c-4 | 2020-12-07T00:00:00Z | 20.2375 | 30.0104167 | 0 | R3IA-c-4 |
| red-tailed hawk | Raptor | winter | IA | Ida | 3 | cropland | R3IA-c-4 | 2021-01-11T00:00:00Z | 2.9263889 | 9.9743056 | 0 | R3IA-c-4 |
| red-tailed hawk | Raptor | winter | IA | Ida | 3 | cropland | R3IA-c-4 | 2021-01-11T00:00:00Z | 49.9291667 | 59.9277778 | 0 | R3IA-c-4 |
| red-tailed hawk | Raptor | winter | IA | Ida | 3 | cropland | R3IA-c-4 | 2021-01-11T00:00:00Z | 59.90625 | NA | 0 | R3IA-c-4 |
| red-tailed hawk | Raptor | winter | IA | Ida | 3 | cropland | R3IA-c-4 | 2021-01-11T00:00:00Z | 2.9152778 | 6.9493056 | 0 | R3IA-c-4 |
| red-tailed hawk | Raptor | winter | IA | Ida | 3 | cropland | R3IA-c-4 | 2021-01-11T00:00:00Z | 2.9104167 | 13.9201389 | 0 | R3IA-c-4 |
| red-tailed hawk | Raptor | winter | IA | Grundy | 3 | cropland | R3IA-c-5 | 2019-11-26T00:00:00Z | 7.0472222 | 9.9701389 | 0 | R3IA-c-5 |
| red-tailed hawk | Raptor | winter | IA | Grundy | 3 | cropland | R3IA-c-5 | 2019-11-26T00:00:00Z | 1.0055556 | 3.0368056 | 0 | R3IA-c-5 |
| red-tailed hawk | Raptor | winter | IA | Grundy | 3 | cropland | R3IA-c-5 | 2019-11-26T00:00:00Z | 59.9784722 | NA | 0 | R3IA-c-5 |
| red-tailed hawk | Raptor | winter | IA | Grundy | 3 | cropland | R3IA-c-5 | 2019-11-26T00:00:00Z | 3.0354167 | 7.0472222 | 0 | R3IA-c-5 |
| red-tailed hawk | Raptor | winter | IA | Grundy | 3 | cropland | R3IA-c-5 | 2019-11-26T00:00:00Z | 1.0041667 | 3.0361111 | 0 | R3IA-c-5 |
| red-tailed hawk | Raptor | spring | IA | Grundy | 3 | cropland | R3IA-c-5 | 2020-03-16T00:00:00Z | 14.1270833 | 20.0902778 | 0 | R3IA-c-5 |
| red-tailed hawk | Raptor | spring | IA | Grundy | 3 | cropland | R3IA-c-5 | 2020-03-16T00:00:00Z | 3.9569444 | 7.0215278 | 0 | R3IA-c-5 |
| red-tailed hawk | Raptor | spring | IA | Grundy | 3 | cropland | R3IA-c-5 | 2020-03-16T00:00:00Z | 30.2798611 | 39.9465278 | 0 | R3IA-c-5 |
| red-tailed hawk | Raptor | spring | IA | Grundy | 3 | cropland | R3IA-c-5 | 2020-03-16T00:00:00Z | 1.0888889 | 1.0888889 | 0 | R3IA-c-5 |
| red-tailed hawk | Raptor | spring | IA | Grundy | 3 | cropland | R3IA-c-5 | 2020-03-16T00:00:00Z | 14.1118056 | 20.075 | 0 | R3IA-c-5 |
| red-tailed hawk | Raptor | winter | IA | Grundy | 3 | cropland | R3IA-c-5 | 2020-01-07T00:00:00Z | 3.0166667 | 7.0020833 | 0 | R3IA-c-5 |
| red-tailed hawk | Raptor | winter | IA | Grundy | 3 | cropland | R3IA-c-5 | 2020-01-07T00:00:00Z | 20.0069444 | 29.9763889 | 0 | R3IA-c-5 |
| red-tailed hawk | Raptor | winter | IA | Grundy | 3 | cropland | R3IA-c-5 | 2020-01-07T00:00:00Z | 1.0097222 | 1.0097222 | 0 | R3IA-c-5 |
| red-tailed hawk | Raptor | winter | IA | Grundy | 3 | cropland | R3IA-c-5 | 2020-01-07T00:00:00Z | 1.0111111 | 3.0145833 | 0 | R3IA-c-5 |
| red-tailed hawk | Raptor | winter | IA | Grundy | 3 | cropland | R3IA-c-5 | 2020-01-07T00:00:00Z | 13.9902778 | 19.9611111 | 0 | R3IA-c-5 |
| red-tailed hawk | Raptor | spring | IA | Grundy | 3 | cropland | R3IA-c-5 | 2020-03-23T00:00:00Z | 1.8979167 | 2.9375 | 0 | R3IA-c-5 |
| red-tailed hawk | Raptor | spring | IA | Grundy | 3 | cropland | R3IA-c-5 | 2020-03-23T00:00:00Z | 10.0381944 | 13.9583333 | 0 | R3IA-c-5 |
| red-tailed hawk | Raptor | spring | IA | Grundy | 3 | cropland | R3IA-c-5 | 2020-03-23T00:00:00Z | 30.1055556 | 39.9597222 | 0 | R3IA-c-5 |
| red-tailed hawk | Raptor | spring | IA | Grundy | 3 | cropland | R3IA-c-5 | 2020-03-23T00:00:00Z | 3.9243056 | 7.1131944 | 0 | R3IA-c-5 |
| red-tailed hawk | Raptor | spring | IA | Grundy | 3 | cropland | R3IA-c-5 | 2020-03-23T00:00:00Z | 1.01875 | 1.8909722 | 0 | R3IA-c-5 |
| ring-necked pheasant | Game Bird | winter | IA | Webster | 3 | cropland | R3IA-c-6 | 2015-01-12T00:00:00Z | 4 | 7 | 0 | R3IA-c-6 |
| ring-necked pheasant | Game Bird | winter | IA | Webster | 3 | cropland | R3IA-c-6 | 2015-01-12T00:00:00Z | 14 | 22 | 0 | R3IA-c-6 |
| ring-necked pheasant | Game Bird | winter | IA | Webster | 3 | cropland | R3IA-c-6 | 2015-02-09T00:00:00Z | 30 | NA | 0 | R3IA-c-6 |
| ring-necked pheasant | Game Bird | winter | IA | Webster | 3 | cropland | R3IA-c-6 | 2015-02-09T00:00:00Z | 30 | NA | 0 | R3IA-c-6 |
| ring-necked pheasant | Game Bird | winter | IA | Webster | 3 | cropland | R3IA-c-6 | 2015-02-09T00:00:00Z | 14 | 21 | 0 | R3IA-c-6 |
| ring-necked pheasant | Game Bird | winter | IA | Webster | 3 | cropland | R3IA-c-6 | 2015-03-16T00:00:00Z | 4 | 7 | 0 | R3IA-c-6 |
| ring-necked pheasant | Game Bird | winter | IA | Webster | 3 | cropland | R3IA-c-6 | 2015-03-16T00:00:00Z | 7 | 9 | 0 | R3IA-c-6 |
| ring-necked pheasant | Game Bird | spring | IA | Webster | 3 | cropland | R3IA-c-6 | 2015-04-13T00:00:00Z | 4 | 7 | 0 | R3IA-c-6 |
| ring-necked pheasant | Game Bird | spring | IA | Webster | 3 | cropland | R3IA-c-6 | 2015-04-13T00:00:00Z | 10 | 15 | 0 | R3IA-c-6 |
| ring-necked pheasant | Game Bird | spring | IA | Webster | 3 | cropland | R3IA-c-6 | 2015-04-13T00:00:00Z | 4 | 7 | 0 | R3IA-c-6 |
| ring-necked pheasant | Game Bird | summer | IA | Webster | 3 | cropland | R3IA-c-6 | 2015-05-18T00:00:00Z | 1 | 1 | 0 | R3IA-c-6 |
| ring-necked pheasant | Game Bird | summer | IA | Webster | 3 | cropland | R3IA-c-6 | 2015-05-18T00:00:00Z | 10 | 14 | 0 | R3IA-c-6 |
| mallard | Game Bird | summer | IA | Webster | 3 | cropland | R3IA-c-6 | 2015-06-15T00:00:00Z | 2 | 3 | 0 | R3IA-c-6 |
| ring-necked pheasant | Game Bird | fall | IA | Webster | 3 | cropland | R3IA-c-6 | 2015-07-27T00:00:00Z | 21 | 30 | 0 | R3IA-c-6 |
| ring-necked pheasant | Game Bird | fall | IA | Webster | 3 | cropland | R3IA-c-6 | 2015-09-28T00:00:00Z | 1 | 1 | 0 | R3IA-c-6 |
| ring-necked pheasant | Game Bird | fall | IA | Webster | 3 | cropland | R3IA-c-6 | 2015-08-17T00:00:00Z | 4 | 7 | 0 | R3IA-c-6 |
| ring-necked pheasant | Game Bird | fall | IA | Webster | 3 | cropland | R3IA-c-6 | 2015-08-17T00:00:00Z | 4 | 7 | 0 | R3IA-c-6 |
| ring-necked pheasant | Game Bird | fall | IA | Webster | 3 | cropland | R3IA-c-6 | 2015-09-14T00:00:00Z | 21 | 30 | 0 | R3IA-c-6 |
| ring-necked pheasant | Game Bird | fall | IA | Webster | 3 | cropland | R3IA-c-6 | 2015-09-14T00:00:00Z | 3 | 4 | 0 | R3IA-c-6 |
| ring-necked pheasant | Game Bird | fall | IA | Webster | 3 | cropland | R3IA-c-6 | 2015-10-12T00:00:00Z | 3 | 4 | 0 | R3IA-c-6 |
| ring-necked pheasant | Game Bird | fall | IA | Webster | 3 | cropland | R3IA-c-6 | 2015-10-12T00:00:00Z | 29 | NA | 0 | R3IA-c-6 |
| ring-necked pheasant | Game Bird | winter | IA | Madison | 3 | cropland | R3IA-c-7 | 2015-03-16T00:00:00Z | 7 | 10 | 0 | R3IA-c-7 |
| ring-necked pheasant | Game Bird | winter | IA | Madison | 3 | cropland | R3IA-c-7 | 2015-03-16T00:00:00Z | 14 | 24 | 0 | R3IA-c-7 |
| ring-necked pheasant | Game Bird | winter | IA | Madison | 3 | cropland | R3IA-c-7 | 2015-03-16T00:00:00Z | 34 | NA | 0 | R3IA-c-7 |
| ring-necked pheasant | Game Bird | spring | IA | Madison | 3 | cropland | R3IA-c-7 | 2015-03-30T00:00:00Z | 30 | NA | 0 | R3IA-c-7 |
| ring-necked pheasant | Game Bird | spring | IA | Madison | 3 | cropland | R3IA-c-7 | 2015-03-30T00:00:00Z | 4 | 7 | 0 | R3IA-c-7 |
| ring-necked pheasant | Game Bird | summer | IA | Madison | 3 | cropland | R3IA-c-7 | 2015-05-18T00:00:00Z | 10 | 14 | 0 | R3IA-c-7 |
| ring-necked pheasant | Game Bird | summer | IA | Madison | 3 | cropland | R3IA-c-7 | 2015-05-18T00:00:00Z | 4 | 7 | 0 | R3IA-c-7 |
| ring-necked pheasant | Game Bird | summer | IA | Madison | 3 | cropland | R3IA-c-7 | 2015-05-18T00:00:00Z | 1 | 1 | 0 | R3IA-c-7 |
| ring-necked pheasant | Game Bird | winter | IA | Madison | 3 | cropland | R3IA-c-7 | 2015-01-09T00:00:00Z | 14 | 20 | 0 | R3IA-c-7 |
| ring-necked pheasant | Game Bird | winter | IA | Madison | 3 | cropland | R3IA-c-7 | 2015-01-09T00:00:00Z | 30 | NA | 0 | R3IA-c-7 |
| ring-necked pheasant | Game Bird | winter | IA | Madison | 3 | cropland | R3IA-c-7 | 2015-02-10T00:00:00Z | 3 | 4 | 0 | R3IA-c-7 |
| ring-necked pheasant | Game Bird | winter | IA | Madison | 3 | cropland | R3IA-c-7 | 2015-02-10T00:00:00Z | 7 | 10 | 0 | R3IA-c-7 |
| ring-necked pheasant | Game Bird | winter | IA | Madison | 3 | cropland | R3IA-c-7 | 2015-02-10T00:00:00Z | 30 | NA | 0 | R3IA-c-7 |
| ring-necked pheasant | Game Bird | fall | IA | Madison | 3 | cropland | R3IA-c-7 | 2015-07-28T00:00:00Z | 1 | 1 | 0 | R3IA-c-7 |
| ring-necked pheasant | Game Bird | fall | IA | Madison | 3 | cropland | R3IA-c-7 | 2015-09-27T00:00:00Z | 5 | 8 | 0 | R3IA-c-7 |
| ring-necked pheasant | Game Bird | winter | IA | Adair/Adams/Cass | 3 | cropland | R3IA-c-8 | 2015-01-09T00:00:00Z | 7 | 10 | 1 | R3IA-c-8 |
| ring-necked pheasant | Game Bird | winter | IA | Adair/Adams/Cass | 3 | cropland | R3IA-c-8 | 2015-01-09T00:00:00Z | 20 | NA | 1 | R3IA-c-8 |
| mallard | Game Bird | winter | IA | Adair/Adams/Cass | 3 | cropland | R3IA-c-8 | 2015-02-12T00:00:00Z | 10 | 14 | 1 | R3IA-c-8 |
| mallard | Game Bird | winter | IA | Adair/Adams/Cass | 3 | cropland | R3IA-c-8 | 2015-02-12T00:00:00Z | 4 | 7 | 1 | R3IA-c-8 |
| mallard | Game Bird | spring | IA | Adair/Adams/Cass | 3 | cropland | R3IA-c-8 | 2015-03-16T00:00:00Z | 1 | 1 | 1 | R3IA-c-8 |
| mallard | Game Bird | spring | IA | Adair/Adams/Cass | 3 | cropland | R3IA-c-8 | 2015-03-16T00:00:00Z | 7 | 10 | 1 | R3IA-c-8 |
| mallard | Game Bird | spring | IA | Adair/Adams/Cass | 3 | cropland | R3IA-c-8 | 2015-03-16T00:00:00Z | 7 | 10 | 1 | R3IA-c-8 |
| red-tailed hawk | Raptor | spring | IA | Adair/Adams/Cass | 3 | cropland | R3IA-c-8 | 2015-03-25T00:00:00Z | 30 | NA | 1 | R3IA-c-0 |
| ring-necked pheasant | Game Bird | summer | IA | Adair/Adams/Cass | 3 | cropland | R3IA-c-8 | 2015-05-18T00:00:00Z | 14 | 21 | 1 | R3IA-c-8 |
| ring-necked pheasant | Game Bird | summer | IA | Adair/Adams/Cass | 3 | cropland | R3IA-c-8 | 2015-06-15T00:00:00Z | 1 | 1 | 1 | R3IA-c-8 |
| ring-necked pheasant | Game Bird | summer | IA | Adair/Adams/Cass | 3 | cropland | R3IA-c-8 | 2015-06-15T00:00:00Z | 14 | 22 | 1 | R3IA-c-8 |
| ring-necked pheasant | Game Bird | fall | IA | Adair/Adams/Cass | 3 | cropland | R3IA-c-8 | 2015-07-27T00:00:00Z | 31 | NA | 1 | R3IA-c-8 |
| ring-necked pheasant | Game Bird | fall | IA | Adair/Adams/Cass | 3 | cropland | R3IA-c-8 | 2015-09-28T00:00:00Z | 30 | NA | 1 | R3IA-c-8 |
| ring-necked pheasant | Game Bird | winter | IA | Adair/Cass | 3 | cropland | R3IA-c-11 | 2015-01-12T00:00:00Z | 7 | 10 | 1 | R3IA-c-11 |
| ring-necked pheasant | Game Bird | winter | IA | Adair/Cass | 3 | cropland | R3IA-c-11 | 2015-01-12T00:00:00Z | 7 | 10 | 1 | R3IA-c-11 |
| ring-necked pheasant | Game Bird | winter | IA | Adair/Cass | 3 | cropland | R3IA-c-11 | 2015-01-12T00:00:00Z | 35 | NA | 1 | R3IA-c-11 |
| ring-necked pheasant | Game Bird | winter | IA | Adair/Cass | 3 | cropland | R3IA-c-11 | 2015-02-15T00:00:00Z | 30 | NA | 1 | R3IA-c-11 |
| ring-necked pheasant | Game Bird | winter | IA | Adair/Cass | 3 | cropland | R3IA-c-11 | 2015-02-15T00:00:00Z | 14 | 20 | 1 | R3IA-c-11 |
| Cooper's hawk | Raptor | winter | IA | Adair/Cass | 3 | cropland | R3IA-c-11 | 2015-02-23T00:00:00Z | 10 | 14 | 1 | R3IA-c-0 |
| ring-necked pheasant | Game Bird | spring | IA | Adair/Cass | 3 | cropland | R3IA-c-11 | 2015-03-16T00:00:00Z | 14 | 20 | 1 | R3IA-c-11 |
| ring-necked pheasant | Game Bird | spring | IA | Adair/Cass | 3 | cropland | R3IA-c-11 | 2015-03-16T00:00:00Z | 20 | 30 | 1 | R3IA-c-11 |
| ring-necked pheasant | Game Bird | spring | IA | Adair/Cass | 3 | cropland | R3IA-c-11 | 2015-03-16T00:00:00Z | 4 | 7 | 1 | R3IA-c-11 |
| ring-necked pheasant | Game Bird | spring | IA | Adair/Cass | 3 | cropland | R3IA-c-11 | 2015-03-30T00:00:00Z | 30 | NA | 1 | R3IA-c-11 |
| ring-necked pheasant | Game Bird | spring | IA | Adair/Cass | 3 | cropland | R3IA-c-11 | 2015-03-30T00:00:00Z | 1 | 1 | 1 | R3IA-c-11 |
| ring-necked pheasant | Game Bird | summer | IA | Adair/Cass | 3 | cropland | R3IA-c-11 | 2015-05-18T00:00:00Z | 21 | 30 | 1 | R3IA-c-11 |
| ring-necked pheasant | Game Bird | summer | IA | Adair/Cass | 3 | cropland | R3IA-c-11 | 2015-06-15T00:00:00Z | 2 | 3 | 1 | R3IA-c-11 |
| ring-necked pheasant | Game Bird | summer | IA | Adair/Cass | 3 | cropland | R3IA-c-11 | 2015-06-15T00:00:00Z | 22 | 29 | 1 | R3IA-c-11 |
| ring-necked pheasant | Game Bird | fall | IA | Adair/Cass | 3 | cropland | R3IA-c-11 | 2015-07-28T00:00:00Z | 1 | 1 | 1 | R3IA-c-11 |
| ring-necked pheasant | Game Bird | fall | IA | Adair/Cass | 3 | cropland | R3IA-c-11 | 2015-09-28T00:00:00Z | 1 | 1 | 1 | R3IA-c-11 |
| ring-necked pheasant | Game Bird | winter | IA | Adair/Cass | 3 | cropland | R3IA-c-11 | 2015-12-01T00:00:00Z | 2 | 3 | 1 | R3IA-c-11 |
| ring-necked pheasant | Game Bird | winter | IA | Adair/Cass | 3 | cropland | R3IA-c-11 | 2015-12-01T00:00:00Z | 10 | 15 | 1 | R3IA-c-11 |
| ring-necked pheasant | Game Bird | winter | IA | Adair/Cass | 3 | cropland | R3IA-c-11 | 2015-12-14T00:00:00Z | 4 | 7 | 1 | R3IA-c-11 |
| ring-necked pheasant | Game Bird | winter | IA | Adair/Cass | 3 | cropland | R3IA-c-11 | 2015-12-14T00:00:00Z | 2 | 3 | 1 | R3IA-c-11 |
| red-tailed hawk | Raptor | winter | IA | Adair/Cass | 3 | cropland | R3IA-c-11 | 2015-12-29T00:00:00Z | 30 | NA | 1 | R3IA-c-0 |
| ring-necked pheasant | Game Bird | winter | IA | Adair/Cass | 3 | cropland | R3IA-c-11 | 2015-12-29T00:00:00Z | 7 | 10 | 1 | R3IA-c-11 |
| ring-necked pheasant | Game Bird | winter | IA | Adair/Cass | 3 | cropland | R3IA-c-11 | 2016-01-11T00:00:00Z | 4 | 7 | 1 | R3IA-c-11 |
| ring-necked pheasant | Game Bird | winter | IA | Adair/Cass | 3 | cropland | R3IA-c-11 | 2016-01-11T00:00:00Z | 3 | 4 | 1 | R3IA-c-11 |
| ring-necked pheasant | Game Bird | winter | IA | Adair/Cass | 3 | cropland | R3IA-c-11 | 2016-01-25T00:00:00Z | 2 | 3 | 1 | R3IA-c-11 |
| ring-necked pheasant | Game Bird | winter | IA | Adair/Cass | 3 | cropland | R3IA-c-11 | 2016-01-25T00:00:00Z | 2 | 3 | 1 | R3IA-c-11 |
| red-tailed hawk | Raptor | winter | IA | Adair/Cass | 3 | cropland | R3IA-c-11 | 2015-12-29T00:00:00Z | 60 | NA | 1 | R3IA-c-0 |
| red-tailed hawk | Raptor | winter | IA | Adair/Cass | 3 | cropland | R3IA-c-11 | 2015-12-29T00:00:00Z | 14 | 20 | 1 | R3IA-c-0 |
| ring-necked pheasant | Game Bird | winter | IA | Adair/Cass | 3 | cropland | R3IA-c-11 | 2016-02-09T00:00:00Z | 30 | NA | 1 | R3IA-c-11 |
| ring-necked pheasant | Game Bird | summer | IA | Adams | 3 | cropland | R3IA-c-12 | 2016-05-16T00:00:00Z | 3 | 4 | 1 | R3IA-c-12 |
| ring-necked pheasant | Game Bird | summer | IA | Adams | 3 | cropland | R3IA-c-12 | 2016-05-31T00:00:00Z | 14 | 21 | 1 | R3IA-c-12 |
| ring-necked pheasant | Game Bird | summer | IA | Adams | 3 | cropland | R3IA-c-12 | 2016-06-13T00:00:00Z | 4 | 7 | 1 | R3IA-c-12 |
| red-tailed hawk | Raptor | fall | IA | Adams | 3 | cropland | R3IA-c-12 | 2016-07-18T00:00:00Z | 51 | NA | 1 | R3IA-c-0 |
| ring-necked pheasant | Game Bird | fall | IA | Adams | 3 | cropland | R3IA-c-12 | 2016-08-02T00:00:00Z | 9 | 10 | 1 | R3IA-c-12 |
| ring-necked pheasant | Game Bird | fall | IA | Adams | 3 | cropland | R3IA-c-12 | 2016-08-15T00:00:00Z | 1 | 2 | 1 | R3IA-c-12 |
| red-tailed hawk | Raptor | fall | IA | Adams | 3 | cropland | R3IA-c-12 | 2016-08-15T00:00:00Z | 2 | 3 | 1 | R3IA-c-0 |
| red-tailed hawk | Raptor | fall | IA | Adams | 3 | cropland | R3IA-c-12 | 2016-09-12T00:00:00Z | 3 | 4 | 1 | R3IA-c-0 |
| ring-necked pheasant | Game Bird | fall | IA | Adams | 3 | cropland | R3IA-c-12 | 2016-09-26T00:00:00Z | 1 | 1 | 1 | R3IA-c-12 |
| red-tailed hawk | Raptor | fall | IA | Adams | 3 | cropland | R3IA-c-12 | 2016-09-26T00:00:00Z | 21 | 30 | 1 | R3IA-c-0 |
| ring-necked pheasant | Game Bird | winter | IA | Adams | 3 | cropland | R3IA-c-12 | 2016-11-28T00:00:00Z | 3 | 7 | 1 | R3IA-c-12 |
| ring-necked pheasant | Game Bird | winter | IA | Adams | 3 | cropland | R3IA-c-12 | 2016-11-28T00:00:00Z | 2 | 3 | 1 | R3IA-c-12 |
| ring-necked pheasant | Game Bird | winter | IA | Adams | 3 | cropland | R3IA-c-12 | 2016-12-12T00:00:00Z | 3 | 4 | 1 | R3IA-c-12 |
| ring-necked pheasant | Game Bird | winter | IA | Adams | 3 | cropland | R3IA-c-12 | 2016-12-12T00:00:00Z | 1 | 2 | 1 | R3IA-c-12 |
| ring-necked pheasant | Game Bird | winter | IA | Adams | 3 | cropland | R3IA-c-12 | 2017-01-09T00:00:00Z | 4 | 10 | 1 | R3IA-c-12 |
| mallard | Game Bird | winter | IA | Adams | 3 | cropland | R3IA-c-12 | 2017-01-09T00:00:00Z | 1 | 2 | 1 | R3IA-c-12 |
| ring-necked pheasant | Game Bird | winter | IA | Adams | 3 | cropland | R3IA-c-12 | 2017-01-23T00:00:00Z | 2 | 3 | 1 | R3IA-c-12 |
| ring-necked pheasant | Game Bird | winter | IA | Adams | 3 | cropland | R3IA-c-12 | 2017-01-23T00:00:00Z | 1 | 2 | 1 | R3IA-c-12 |
| ring-necked pheasant | Game Bird | spring | IA | Adams | 3 | cropland | R3IA-c-12 | 2017-03-27T00:00:00Z | 3.8965278 | 7.0243056 | 1 | R3IA-c-12 |
| ring-necked pheasant | Game Bird | spring | IA | Adams | 3 | cropland | R3IA-c-12 | 2017-03-27T00:00:00Z | 2.13125 | 2.9465278 | 1 | R3IA-c-12 |
| ring-necked pheasant | Game Bird | spring | IA | Adams | 3 | cropland | R3IA-c-12 | 2017-03-27T00:00:00Z | 2.9284722 | 3.7930556 | 1 | R3IA-c-12 |
| ring-necked pheasant | Game Bird | winter | IA | Carroll | 3 | cropland | R3IA-c-13 | 2015-01-12T00:00:00Z | 7 | 10 | 1 | R3IA-c-13 |
| ring-necked pheasant | Game Bird | winter | IA | Carroll | 3 | cropland | R3IA-c-13 | 2015-01-12T00:00:00Z | 7 | 10 | 1 | R3IA-c-13 |
| ring-necked pheasant | Game Bird | winter | IA | Carroll | 3 | cropland | R3IA-c-13 | 2015-01-12T00:00:00Z | 30 | NA | 1 | R3IA-c-13 |
| ring-necked pheasant | Game Bird | winter | IA | Carroll | 3 | cropland | R3IA-c-13 | 2015-02-09T00:00:00Z | 21 | 30 | 1 | R3IA-c-13 |
| ring-necked pheasant | Game Bird | winter | IA | Carroll | 3 | cropland | R3IA-c-13 | 2015-02-09T00:00:00Z | 21 | 30 | 1 | R3IA-c-13 |
| ring-necked pheasant | Game Bird | spring | IA | Carroll | 3 | cropland | R3IA-c-13 | 2015-03-16T00:00:00Z | 3 | 4 | 1 | R3IA-c-13 |
| ring-necked pheasant | Game Bird | spring | IA | Carroll | 3 | cropland | R3IA-c-13 | 2015-03-16T00:00:00Z | 14 | 21 | 1 | R3IA-c-13 |
| ring-necked pheasant | Game Bird | spring | IA | Carroll | 3 | cropland | R3IA-c-13 | 2015-03-17T00:00:00Z | 3 | 4 | 1 | R3IA-c-13 |
| ring-necked pheasant | Game Bird | spring | IA | Carroll | 3 | cropland | R3IA-c-13 | 2015-04-13T00:00:00Z | 1 | 2 | 1 | R3IA-c-13 |
| ring-necked pheasant | Game Bird | spring | IA | Carroll | 3 | cropland | R3IA-c-13 | 2015-04-13T00:00:00Z | 21 | 30 | 1 | R3IA-c-13 |
| ring-necked pheasant | Game Bird | summer | IA | Carroll | 3 | cropland | R3IA-c-13 | 2015-05-18T00:00:00Z | 1 | 2 | 1 | R3IA-c-13 |
| ring-necked pheasant | Game Bird | summer | IA | Carroll | 3 | cropland | R3IA-c-13 | 2015-06-15T00:00:00Z | 3 | 4 | 1 | R3IA-c-13 |
| ring-necked pheasant | Game Bird | summer | IA | Carroll | 3 | cropland | R3IA-c-13 | 2015-06-15T00:00:00Z | 1 | 1 | 1 | R3IA-c-13 |
| ring-necked pheasant | Game Bird | summer | IA | Carroll | 3 | cropland | R3IA-c-13 | 2015-07-27T00:00:00Z | 10 | 14 | 1 | R3IA-c-13 |
| ring-necked pheasant | Game Bird | summer | IA | Carroll | 3 | cropland | R3IA-c-13 | 2015-08-17T00:00:00Z | 1 | 1 | 1 | R3IA-c-13 |
| ring-necked pheasant | Game Bird | summer | IA | Carroll | 3 | cropland | R3IA-c-13 | 2015-09-14T00:00:00Z | 3 | 4 | 1 | R3IA-c-13 |
| ring-necked pheasant | Game Bird | summer | IA | Carroll | 3 | cropland | R3IA-c-13 | 2015-09-14T00:00:00Z | 20 | 30 | 1 | R3IA-c-13 |
| ring-necked pheasant | Game Bird | fall | IA | Carroll | 3 | cropland | R3IA-c-13 | 2015-09-28T00:00:00Z | 1 | 1 | 1 | R3IA-c-13 |
| ring-necked pheasant | Game Bird | fall | IA | Carroll | 3 | cropland | R3IA-c-13 | 2015-10-12T00:00:00Z | 2 | 3 | 1 | R3IA-c-13 |
| ring-necked pheasant | Game Bird | fall | IA | Carroll | 3 | cropland | R3IA-c-13 | 2015-10-12T00:00:00Z | 1 | 1 | 1 | R3IA-c-13 |
| ring-necked pheasant | Game Bird | fall | IA | Carroll | 3 | cropland | R3IA-c-13 | 2015-10-12T00:00:00Z | 14 | 21 | 1 | R3IA-c-13 |
| ring-necked pheasant | Game Bird | winter | IA | Carroll | 3 | cropland | R3IA-c-13 | 2015-12-01T00:00:00Z | 4 | 7 | 1 | R3IA-c-13 |
| ring-necked pheasant | Game Bird | winter | IA | Carroll | 3 | cropland | R3IA-c-13 | 2015-12-14T00:00:00Z | 15 | 22 | 1 | R3IA-c-13 |
| mallard | Game Bird | winter | IA | Carroll | 3 | cropland | R3IA-c-13 | 2015-12-14T00:00:00Z | 15 | NA | 1 | R3IA-c-13 |
| red-tailed hawk | Raptor | winter | IA | Carroll | 3 | cropland | R3IA-c-13 | 2015-12-29T00:00:00Z | 62 | NA | 1 | R3IA-c-0 |
| red-tailed hawk | Raptor | winter | IA | Carroll | 3 | cropland | R3IA-c-13 | 2015-12-29T00:00:00Z | 42 | 56 | 1 | R3IA-c-0 |
| ring-necked pheasant | Game Bird | winter | IA | Carroll | 3 | cropland | R3IA-c-13 | 2015-12-29T00:00:00Z | 2 | 14 | 1 | R3IA-c-13 |
| ring-necked pheasant | Game Bird | winter | IA | Carroll | 3 | cropland | R3IA-c-13 | 2015-12-29T00:00:00Z | 7 | 20 | 1 | R3IA-c-13 |
| ring-necked pheasant | Game Bird | winter | IA | Carroll | 3 | cropland | R3IA-c-13 | 2016-01-11T00:00:00Z | 7 | 10 | 1 | R3IA-c-13 |
| red-tailed hawk | Raptor | winter | IA | Carroll | 3 | cropland | R3IA-c-13 | 2016-01-18T00:00:00Z | 59 | NA | 1 | R3IA-c-0 |
| red-tailed hawk | Raptor | winter | IA | Carroll | 3 | cropland | R3IA-c-13 | 2016-01-18T00:00:00Z | 59 | NA | 1 | R3IA-c-0 |
| red-tailed hawk | Raptor | winter | IA | Carroll | 3 | cropland | R3IA-c-13 | 2016-01-26T00:00:00Z | 60 | NA | 1 | R3IA-c-0 |
| ring-necked pheasant | Game Bird | winter | IA | Carroll | 3 | cropland | R3IA-c-13 | 2016-01-26T00:00:00Z | 14 | 30 | 1 | R3IA-c-13 |
| ring-necked pheasant | Game Bird | winter | IA | Carroll | 3 | cropland | R3IA-c-13 | 2016-01-26T00:00:00Z | 1 | 2 | 1 | R3IA-c-13 |
| ring-necked pheasant | Game Bird | winter | IA | Carroll | 3 | cropland | R3IA-c-13 | 2016-02-09T00:00:00Z | 7 | 10 | 1 | R3IA-c-13 |
| ring-necked pheasant | Game Bird | winter | IA | Carroll | 3 | cropland | R3IA-c-13 | 2016-02-09T00:00:00Z | 7 | 10 | 1 | R3IA-c-13 |
| ring-necked pheasant | Game Bird | winter | IA | Hamilton/Wright | 3 | cropland | R3IA-c-14 | 2015-12-01T00:00:00Z | 14 | NA | 1 | R3IA-c-14 |
| ring-necked pheasant | Game Bird | winter | IA | Hamilton/Wright | 3 | cropland | R3IA-c-14 | 2015-12-01T00:00:00Z | 10 | 14 | 1 | R3IA-c-14 |
| ring-necked pheasant | Game Bird | winter | IA | Hamilton/Wright | 3 | cropland | R3IA-c-14 | 2015-12-14T00:00:00Z | 1 | 2 | 1 | R3IA-c-14 |
| ring-necked pheasant | Game Bird | winter | IA | Hamilton/Wright | 3 | cropland | R3IA-c-14 | 2015-12-14T00:00:00Z | 9 | NA | 1 | R3IA-c-14 |
| ring-necked pheasant | Game Bird | winter | IA | Hamilton/Wright | 3 | cropland | R3IA-c-14 | 2015-12-14T00:00:00Z | 7 | NA | 1 | R3IA-c-14 |
| ring-necked pheasant | Game Bird | winter | IA | Hamilton/Wright | 3 | cropland | R3IA-c-14 | 2016-01-04T00:00:00Z | 3 | 10 | 1 | R3IA-c-14 |
| ring-necked pheasant | Game Bird | winter | IA | Hamilton/Wright | 3 | cropland | R3IA-c-14 | 2016-01-04T00:00:00Z | 7 | 10 | 1 | R3IA-c-14 |
| ring-necked pheasant | Game Bird | winter | IA | Hamilton/Wright | 3 | cropland | R3IA-c-14 | 2016-01-26T00:00:00Z | 29 | NA | 1 | R3IA-c-14 |
| ring-necked pheasant | Game Bird | winter | IA | Hamilton/Wright | 3 | cropland | R3IA-c-14 | 2016-01-26T00:00:00Z | 6 | 29 | 1 | R3IA-c-14 |
| ring-necked pheasant | Game Bird | winter | IA | Hamilton/Wright | 3 | cropland | R3IA-c-14 | 2016-01-26T00:00:00Z | 29 | NA | 1 | R3IA-c-14 |
| ring-necked pheasant | Game Bird | winter | IA | Hamilton/Wright | 3 | cropland | R3IA-c-14 | 2016-02-09T00:00:00Z | 30 | NA | 1 | R3IA-c-14 |
| mallard | Game Bird | spring | IA | Hamilton/Wright | 3 | cropland | R3IA-c-14 | 2016-03-22T00:00:00Z | 10 | 14 | 1 | R3IA-c-14 |
| ring-necked pheasant | Game Bird | spring | IA | Hamilton/Wright | 3 | cropland | R3IA-c-14 | 2016-03-28T00:00:00Z | 20 | 30 | 1 | R3IA-c-14 |
| ring-necked pheasant | Game Bird | spring | IA | Hamilton/Wright | 3 | cropland | R3IA-c-14 | 2016-03-28T00:00:00Z | 3 | 4 | 1 | R3IA-c-14 |
| ring-necked pheasant | Game Bird | spring | IA | Hamilton/Wright | 3 | cropland | R3IA-c-14 | 2016-03-28T00:00:00Z | 30 | NA | 1 | R3IA-c-14 |
| ring-necked pheasant | Game Bird | spring | IA | Hamilton/Wright | 3 | cropland | R3IA-c-14 | 2016-03-28T00:00:00Z | 30 | NA | 1 | R3IA-c-14 |
| ring-necked pheasant | Game Bird | spring | IA | Hamilton/Wright | 3 | cropland | R3IA-c-14 | 2016-04-04T00:00:00Z | 20 | 30 | 1 | R3IA-c-14 |
| ring-necked pheasant | Game Bird | spring | IA | Hamilton/Wright | 3 | cropland | R3IA-c-14 | 2016-04-04T00:00:00Z | 1 | 2 | 1 | R3IA-c-14 |
| ring-necked pheasant | Game Bird | spring | IA | Hamilton/Wright | 3 | cropland | R3IA-c-14 | 2016-04-04T00:00:00Z | 10 | 14 | 1 | R3IA-c-14 |
| red-tailed hawk | Raptor | spring | IA | Hamilton/Wright | 3 | cropland | R3IA-c-14 | 2016-04-11T00:00:00Z | 30 | NA | 1 | R3IA-c-14 |
| red-tailed hawk | Raptor | spring | IA | Hamilton/Wright | 3 | cropland | R3IA-c-14 | 2016-04-11T00:00:00Z | 30 | NA | 1 | R3IA-c-14 |
| red-tailed hawk | Raptor | spring | IA | Hamilton/Wright | 3 | cropland | R3IA-c-14 | 2016-04-11T00:00:00Z | 14 | 20 | 1 | R3IA-c-14 |
| ring-necked pheasant | Game Bird | spring | IA | Hamilton/Wright | 3 | cropland | R3IA-c-14 | 2016-04-11T00:00:00Z | 30 | NA | 1 | R3IA-c-14 |
| red-tailed hawk | Raptor | spring | IA | Hamilton/Wright | 3 | cropland | R3IA-c-14 | 2016-04-11T00:00:00Z | 30 | NA | 1 | R3IA-c-14 |
| red-tailed hawk | Raptor | spring | IA | Hamilton/Wright | 3 | cropland | R3IA-c-14 | 2016-04-11T00:00:00Z | 30 | NA | 1 | R3IA-c-14 |
| ring-necked pheasant | Game Bird | spring | IA | Hamilton/Wright | 3 | cropland | R3IA-c-14 | 2016-04-11T00:00:00Z | 30 | NA | 1 | R3IA-c-14 |
| red-tailed hawk | Raptor | spring | IA | Hamilton/Wright | 3 | cropland | R3IA-c-14 | 2016-04-15T00:00:00Z | 21 | 30 | 1 | R3IA-c-14 |
| ring-necked pheasant | Game Bird | spring | IA | Hamilton/Wright | 3 | cropland | R3IA-c-14 | 2016-04-15T00:00:00Z | 21 | 30 | 1 | R3IA-c-14 |
| ring-necked pheasant | Game Bird | spring | IA | Hamilton/Wright | 3 | cropland | R3IA-c-14 | 2016-04-15T00:00:00Z | 30 | NA | 1 | R3IA-c-14 |
| ring-necked pheasant | Game Bird | summer | IA | Hamilton/Wright | 3 | cropland | R3IA-c-14 | 2016-05-24T00:00:00Z | 4 | 10 | 1 | R3IA-c-14 |
| ring-necked pheasant | Game Bird | summer | IA | Hamilton/Wright | 3 | cropland | R3IA-c-14 | 2016-06-08T00:00:00Z | 15 | 20 | 1 | R3IA-c-14 |
| ring-necked pheasant | Game Bird | fall | IA | Hamilton/Wright | 3 | cropland | R3IA-c-14 | 2016-08-04T00:00:00Z | 4 | 20 | 1 | R3IA-c-14 |
| ring-necked pheasant | Game Bird | fall | IA | Hamilton/Wright | 3 | cropland | R3IA-c-14 | 2016-10-11T00:00:00Z | 10 | 14 | 1 | R3IA-c-14 |
| ring-necked pheasant | Game Bird | fall | IA | Hamilton/Wright | 3 | cropland | R3IA-c-14 | 2016-10-14T00:00:00Z | 7 | 10 | 1 | R3IA-c-14 |
| ring-necked pheasant | Game Bird | winter | IA | Hamilton/Wright | 3 | cropland | R3IA-c-14 | 2016-11-28T00:00:00Z | 30 | NA | 1 | R3IA-c-14 |
| ring-necked pheasant | Game Bird | winter | IA | Hamilton/Wright | 3 | cropland | R3IA-c-14 | 2016-11-28T00:00:00Z | 30 | NA | 1 | R3IA-c-14 |
| red-tailed hawk | Raptor | winter | IA | Hamilton/Wright | 3 | cropland | R3IA-c-14 | 2016-12-05T00:00:00Z | 46 | 50 | 1 | R3IA-c-14 |
| red-tailed hawk | Raptor | winter | IA | Hamilton/Wright | 3 | cropland | R3IA-c-14 | 2016-12-05T00:00:00Z | 60 | NA | 1 | R3IA-c-14 |
| red-tailed hawk | Raptor | winter | IA | Hamilton/Wright | 3 | cropland | R3IA-c-14 | 2016-12-05T00:00:00Z | 60 | NA | 1 | R3IA-c-14 |
| ring-necked pheasant | Game Bird | winter | IA | Hamilton/Wright | 3 | cropland | R3IA-c-14 | 2016-12-12T00:00:00Z | 30 | NA | 1 | R3IA-c-14 |
| ring-necked pheasant | Game Bird | winter | IA | Hamilton/Wright | 3 | cropland | R3IA-c-14 | 2016-12-12T00:00:00Z | 21 | 30 | 1 | R3IA-c-14 |
| ring-necked pheasant | Game Bird | winter | IA | Hamilton/Wright | 3 | cropland | R3IA-c-14 | 2017-01-09T00:00:00Z | 2 | 2 | 1 | R3IA-c-14 |
| ring-necked pheasant | Game Bird | winter | IA | Hamilton/Wright | 3 | cropland | R3IA-c-14 | 2017-01-09T00:00:00Z | 11 | 14 | 1 | R3IA-c-14 |
| red-tailed hawk | Raptor | winter | IA | Hamilton/Wright | 3 | cropland | R3IA-c-14 | 2017-01-03T00:00:00Z | 4 | 8 | 1 | R3IA-c-14 |
| red-tailed hawk | Raptor | winter | IA | Hamilton/Wright | 3 | cropland | R3IA-c-14 | 2017-01-03T00:00:00Z | 64 | NA | 1 | R3IA-c-14 |
| red-tailed hawk | Raptor | winter | IA | Hamilton/Wright | 3 | cropland | R3IA-c-14 | 2017-01-03T00:00:00Z | 64 | NA | 1 | R3IA-c-14 |
| red-tailed hawk | Raptor | winter | IA | Hamilton/Wright | 3 | cropland | R3IA-c-14 | 2017-01-03T00:00:00Z | 17 | 20 | 1 | R3IA-c-14 |
| red-tailed hawk | Raptor | winter | IA | Hamilton/Wright | 3 | cropland | R3IA-c-14 | 2017-01-09T00:00:00Z | 44 | 50 | 1 | R3IA-c-14 |
| red-tailed hawk | Raptor | winter | IA | Hamilton/Wright | 3 | cropland | R3IA-c-14 | 2017-01-09T00:00:00Z | 44 | 50 | 1 | R3IA-c-14 |
| red-tailed hawk | Raptor | winter | IA | Hamilton/Wright | 3 | cropland | R3IA-c-14 | 2017-01-09T00:00:00Z | 44 | 50 | 1 | R3IA-c-14 |
| ring-necked pheasant | Game Bird | winter | IA | Hamilton/Wright | 3 | cropland | R3IA-c-14 | 2017-01-23T00:00:00Z | 14 | 21 | 1 | R3IA-c-14 |
| ring-necked pheasant | Game Bird | winter | IA | Hamilton/Wright | 3 | cropland | R3IA-c-14 | 2017-01-23T00:00:00Z | 30 | NA | 1 | R3IA-c-14 |
| ring-necked pheasant | Game Bird | winter | IA | Hamilton/Wright | 3 | cropland | R3IA-c-14 | 2017-02-06T00:00:00Z | 16 | 20 | 1 | R3IA-c-14 |
| ring-necked pheasant | Game Bird | winter | IA | Hamilton/Wright | 3 | cropland | R3IA-c-14 | 2017-02-06T00:00:00Z | 30 | NA | 1 | R3IA-c-14 |
| red-tailed hawk | Raptor | winter | IA | Hamilton/Wright | 3 | cropland | R3IA-c-14 | 2017-01-23T00:00:00Z | 59 | NA | 1 | R3IA-c-14 |
| red-tailed hawk | Raptor | winter | IA | Hamilton/Wright | 3 | cropland | R3IA-c-14 | 2017-01-23T00:00:00Z | 59 | NA | 1 | R3IA-c-14 |
| red-tailed hawk | Raptor | winter | IA | Hamilton/Wright | 3 | cropland | R3IA-c-14 | 2017-01-23T00:00:00Z | 59 | NA | 1 | R3IA-c-14 |
| red-tailed hawk | Raptor | winter | IA | Hamilton/Wright | 3 | cropland | R3IA-c-14 | 2017-01-23T00:00:00Z | 14 | 20 | 1 | R3IA-c-14 |
| red-tailed hawk | Raptor | winter | IA | Hamilton/Wright | 3 | cropland | R3IA-c-14 | 2017-01-30T00:00:00Z | 37 | 52 | 1 | R3IA-c-14 |
| red-tailed hawk | Raptor | winter | IA | Hamilton/Wright | 3 | cropland | R3IA-c-14 | 2017-01-30T00:00:00Z | 52 | NA | 1 | R3IA-c-14 |
| red-tailed hawk | Raptor | winter | IA | Hamilton/Wright | 3 | cropland | R3IA-c-14 | 2017-01-30T00:00:00Z | 52 | NA | 1 | R3IA-c-14 |
| red-tailed hawk | Raptor | winter | IA | Hamilton/Wright | 3 | cropland | R3IA-c-14 | 2017-01-30T00:00:00Z | 52 | NA | 1 | R3IA-c-14 |
| red-tailed hawk | Raptor | winter | IA | Hamilton/Wright | 3 | cropland | R3IA-c-14 | 2017-02-06T00:00:00Z | 45 | NA | 1 | R3IA-c-14 |
| red-tailed hawk | Raptor | winter | IA | Hamilton/Wright | 3 | cropland | R3IA-c-14 | 2017-02-06T00:00:00Z | 7 | 10 | 1 | R3IA-c-14 |
| red-tailed hawk | Raptor | winter | IA | Hamilton/Wright | 3 | cropland | R3IA-c-14 | 2017-02-06T00:00:00Z | 30 | 45 | 1 | R3IA-c-14 |
| red-tailed hawk | Raptor | winter | IA | Hamilton/Wright | 3 | cropland | R3IA-c-14 | 2017-02-06T00:00:00Z | 45 | NA | 1 | R3IA-c-14 |
| red-tailed hawk | Raptor | winter | IA | Hamilton/Wright | 3 | cropland | R3IA-c-14 | 2017-02-13T00:00:00Z | 38 | NA | 1 | R3IA-c-14 |
| red-tailed hawk | Raptor | winter | IA | Hamilton/Wright | 3 | cropland | R3IA-c-14 | 2017-02-13T00:00:00Z | 38 | NA | 1 | R3IA-c-14 |
| red-tailed hawk | Raptor | winter | IA | Hamilton/Wright | 3 | cropland | R3IA-c-14 | 2017-02-13T00:00:00Z | 38 | NA | 1 | R3IA-c-14 |
| red-tailed hawk | Raptor | winter | IA | Hamilton/Wright | 3 | cropland | R3IA-c-14 | 2017-02-13T00:00:00Z | 1 | 1 | 1 | R3IA-c-14 |
| red-tailed hawk | Raptor | winter | IA | Hamilton/Wright | 3 | cropland | R3IA-c-14 | 2017-02-21T00:00:00Z | 30 | NA | 1 | R3IA-c-14 |
| red-tailed hawk | Raptor | winter | IA | Hamilton/Wright | 3 | cropland | R3IA-c-14 | 2017-02-21T00:00:00Z | 2 | 7 | 1 | R3IA-c-14 |
| red-tailed hawk | Raptor | winter | IA | Hamilton/Wright | 3 | cropland | R3IA-c-14 | 2017-02-21T00:00:00Z | 30 | NA | 1 | R3IA-c-14 |
| ring-necked pheasant | Game Bird | winter | IA | Floyd | 3 | cropland | R3IA-c-15 | 2015-12-01T00:00:00Z | 10 | 13 | 1 | R3IA-c-15 |
| ring-necked pheasant | Game Bird | winter | IA | Floyd | 3 | cropland | R3IA-c-15 | 2015-12-01T00:00:00Z | 10 | 13 | 1 | R3IA-c-15 |
| ring-necked pheasant | Game Bird | winter | IA | Floyd | 3 | cropland | R3IA-c-15 | 2015-12-14T00:00:00Z | 30 | NA | 1 | R3IA-c-15 |
| ring-necked pheasant | Game Bird | winter | IA | Floyd | 3 | cropland | R3IA-c-15 | 2015-12-14T00:00:00Z | 7 | 10 | 1 | R3IA-c-15 |
| ring-necked pheasant | Game Bird | winter | IA | Floyd | 3 | cropland | R3IA-c-15 | 2016-01-11T00:00:00Z | 30 | NA | 1 | R3IA-c-15 |
| ring-necked pheasant | Game Bird | winter | IA | Floyd | 3 | cropland | R3IA-c-15 | 2016-01-25T00:00:00Z | 2 | 3 | 1 | R3IA-c-15 |
| ring-necked pheasant | Game Bird | winter | IA | Floyd | 3 | cropland | R3IA-c-15 | 2016-02-09T00:00:00Z | 15 | 20 | 1 | R3IA-c-15 |
| ring-necked pheasant | Game Bird | spring | IA | Floyd | 3 | cropland | R3IA-c-15 | 2016-03-25T00:00:00Z | 2 | 3 | 1 | R3IA-c-15 |
| ring-necked pheasant | Game Bird | spring | IA | Floyd | 3 | cropland | R3IA-c-15 | 2016-03-31T00:00:00Z | 14 | 20 | 1 | R3IA-c-15 |
| red-tailed hawk | Raptor | spring | IA | Floyd | 3 | cropland | R3IA-c-15 | 2016-04-07T00:00:00Z | 1 | 1 | 1 | R3IA-c-0 |
| red-tailed hawk | Raptor | spring | IA | Floyd | 3 | cropland | R3IA-c-15 | 2016-04-12T00:00:00Z | 7 | 10 | 1 | R3IA-c-0 |
| turkey vulture | Raptor | spring | IA | Floyd | 3 | cropland | R3IA-c-15 | 2016-04-12T00:00:00Z | 30 | NA | 1 | R3IA-c-0 |
| turkey vulture | Raptor | spring | IA | Floyd | 3 | cropland | R3IA-c-15 | 2016-04-12T00:00:00Z | 30 | NA | 1 | R3IA-c-0 |
| red-tailed hawk | Raptor | spring | IA | Floyd | 3 | cropland | R3IA-c-15 | 2016-04-12T00:00:00Z | 30 | NA | 1 | R3IA-c-0 |
| ring-necked pheasant | Game Bird | spring | IA | Floyd | 3 | cropland | R3IA-c-15 | 2016-04-12T00:00:00Z | 10 | 14 | 1 | R3IA-c-15 |
| ring-necked pheasant | Game Bird | summer | IA | Floyd | 3 | cropland | R3IA-c-15 | 2016-05-24T00:00:00Z | 3 | 4 | 1 | R3IA-c-15 |
| ring-necked pheasant | Game Bird | summer | IA | Floyd | 3 | cropland | R3IA-c-15 | 2016-06-09T00:00:00Z | 32 | NA | 1 | R3IA-c-15 |
| ring-necked pheasant | Game Bird | fall | IA | Floyd | 3 | cropland | R3IA-c-15 | 2016-08-01T00:00:00Z | 1 | 4 | 1 | R3IA-c-15 |
| ring-necked pheasant | Game Bird | fall | IA | Floyd | 3 | cropland | R3IA-c-15 | 2016-08-29T00:00:00Z | 8 | 10 | 1 | R3IA-c-15 |
| ring-necked pheasant | Game Bird | fall | IA | Floyd | 3 | cropland | R3IA-c-15 | 2016-09-26T00:00:00Z | 30 | NA | 1 | R3IA-c-15 |
| ring-necked pheasant | Game Bird | winter | IA | Floyd | 3 | cropland | R3IA-c-15 | 2016-12-05T00:00:00Z | 14 | 24 | 1 | R3IA-c-15 |
| ring-necked pheasant | Game Bird | winter | IA | Floyd | 3 | cropland | R3IA-c-15 | 2016-12-05T00:00:00Z | 10 | 14 | 1 | R3IA-c-15 |
| ring-necked pheasant | Game Bird | winter | IA | Floyd | 3 | cropland | R3IA-c-15 | 2017-01-23T00:00:00Z | 14 | 20 | 1 | R3IA-c-15 |
| ring-necked pheasant | Game Bird | winter | IA | Floyd | 3 | cropland | R3IA-c-15 | 2017-01-23T00:00:00Z | 10 | 10 | 1 | R3IA-c-15 |
| ring-necked pheasant | Game Bird | winter | IA | Floyd | 3 | cropland | R3IA-c-15 | 2017-01-23T00:00:00Z | 20 | 30 | 1 | R3IA-c-15 |
| ring-necked pheasant | Game Bird | winter | IA | Floyd | 3 | cropland | R3IA-c-15 | 2017-01-30T00:00:00Z | 2 | 3 | 1 | R3IA-c-15 |
| ring-necked pheasant | Game Bird | winter | IA | Floyd | 3 | cropland | R3IA-c-15 | 2017-01-30T00:00:00Z | 10 | 14 | 1 | R3IA-c-15 |
| ring-necked pheasant | Game Bird | winter | IA | Floyd | 3 | cropland | R3IA-c-15 | 2017-01-30T00:00:00Z | 7 | 10 | 1 | R3IA-c-15 |
| ring-necked pheasant | Game Bird | winter | IA | Floyd | 3 | cropland | R3IA-c-15 | 2017-02-06T00:00:00Z | 4 | 7 | 1 | R3IA-c-15 |
| ring-necked pheasant | Game Bird | winter | IA | Floyd | 3 | cropland | R3IA-c-15 | 2017-02-06T00:00:00Z | 4 | 7 | 1 | R3IA-c-15 |
| short-eared owl | Raptor | winter | IA | Audubon/Guthrie | 3 | cropland | R3IA-c-16 | 2014-12-03T00:00:00Z | 14 | 20 | 1 | R3IA-c-0 |
| ring-necked pheasant | Game Bird | winter | IA | Audubon/Guthrie | 3 | cropland | R3IA-c-16 | 2015-01-12T00:00:00Z | 4 | 10 | 1 | R3IA-c-16 |
| ring-necked pheasant | Game Bird | winter | IA | Audubon/Guthrie | 3 | cropland | R3IA-c-16 | 2015-01-12T00:00:00Z | 4 | 12 | 1 | R3IA-c-16 |
| ring-necked pheasant | Game Bird | winter | IA | Audubon/Guthrie | 3 | cropland | R3IA-c-16 | 2015-02-11T00:00:00Z | 3 | 5 | 1 | R3IA-c-16 |
| ring-necked pheasant | Game Bird | winter | IA | Audubon/Guthrie | 3 | cropland | R3IA-c-16 | 2015-02-11T00:00:00Z | 31 | NA | 1 | R3IA-c-16 |
| ring-necked pheasant | Game Bird | spring | IA | Audubon/Guthrie | 3 | cropland | R3IA-c-16 | 2015-03-16T00:00:00Z | 10 | 14 | 1 | R3IA-c-16 |
| ring-necked pheasant | Game Bird | spring | IA | Audubon/Guthrie | 3 | cropland | R3IA-c-16 | 2015-03-16T00:00:00Z | 10 | 14 | 1 | R3IA-c-16 |
| ring-necked pheasant | Game Bird | spring | IA | Audubon/Guthrie | 3 | cropland | R3IA-c-16 | 2015-03-30T00:00:00Z | 14 | 20 | 1 | R3IA-c-16 |
| ring-necked pheasant | Game Bird | spring | IA | Audubon/Guthrie | 3 | cropland | R3IA-c-16 | 2015-03-30T00:00:00Z | 14 | 20 | 1 | R3IA-c-16 |
| ring-necked pheasant | Game Bird | spring | IA | Audubon/Guthrie | 3 | cropland | R3IA-c-16 | 2015-03-30T00:00:00Z | 1 | 1 | 1 | R3IA-c-16 |
| ring-necked pheasant | Game Bird | summer | IA | Audubon/Guthrie | 3 | cropland | R3IA-c-16 | 2015-05-18T00:00:00Z | 14 | 21 | 1 | R3IA-c-16 |
| ring-necked pheasant | Game Bird | summer | IA | Audubon/Guthrie | 3 | cropland | R3IA-c-16 | 2015-05-18T00:00:00Z | 2 | 3 | 1 | R3IA-c-16 |
| ring-necked pheasant | Game Bird | summer | IA | Audubon/Guthrie | 3 | cropland | R3IA-c-16 | 2015-06-15T00:00:00Z | 23 | 29 | 1 | R3IA-c-16 |
| ring-necked pheasant | Game Bird | summer | IA | Audubon/Guthrie | 3 | cropland | R3IA-c-16 | 2015-07-28T00:00:00Z | 4 | 7 | 1 | R3IA-c-16 |
| ring-necked pheasant | Game Bird | summer | IA | Audubon/Guthrie | 3 | cropland | R3IA-c-16 | 2015-08-17T00:00:00Z | 14 | 22 | 1 | R3IA-c-16 |
| ring-necked pheasant | Game Bird | summer | IA | Audubon/Guthrie | 3 | cropland | R3IA-c-16 | 2015-08-17T00:00:00Z | 3 | 4 | 1 | R3IA-c-16 |
| ring-necked pheasant | Game Bird | summer | IA | Audubon/Guthrie | 3 | cropland | R3IA-c-16 | 2015-09-14T00:00:00Z | 1 | 2 | 1 | R3IA-c-16 |
| ring-necked pheasant | Game Bird | summer | IA | Audubon/Guthrie | 3 | cropland | R3IA-c-16 | 2015-09-14T00:00:00Z | 1 | 2 | 1 | R3IA-c-16 |
| ring-necked pheasant | Game Bird | fall | IA | Audubon/Guthrie | 3 | cropland | R3IA-c-16 | 2015-09-28T00:00:00Z | 2 | 3 | 1 | R3IA-c-16 |
| ring-necked pheasant | Game Bird | fall | IA | Audubon/Guthrie | 3 | cropland | R3IA-c-16 | 2015-10-12T00:00:00Z | 31 | NA | 1 | R3IA-c-16 |
| ring-necked pheasant | Game Bird | fall | IA | Audubon/Guthrie | 3 | cropland | R3IA-c-16 | 2015-10-12T00:00:00Z | 4 | 7 | 1 | R3IA-c-16 |
| ring-necked pheasant | Game Bird | winter | IA | Audubon/Guthrie | 3 | cropland | R3IA-c-16 | 2015-12-01T00:00:00Z | 20 | 30 | 1 | R3IA-c-16 |
| ring-necked pheasant | Game Bird | winter | IA | Audubon/Guthrie | 3 | cropland | R3IA-c-16 | 2015-12-01T00:00:00Z | 20 | 30 | 1 | R3IA-c-16 |
| ring-necked pheasant | Game Bird | winter | IA | Audubon/Guthrie | 3 | cropland | R3IA-c-16 | 2015-12-14T00:00:00Z | 7 | 10 | 1 | R3IA-c-16 |
| ring-necked pheasant | Game Bird | winter | IA | Audubon/Guthrie | 3 | cropland | R3IA-c-16 | 2015-12-14T00:00:00Z | 10 | NA | 1 | R3IA-c-16 |
| ring-necked pheasant | Game Bird | winter | IA | Audubon/Guthrie | 3 | cropland | R3IA-c-16 | 2015-12-29T00:00:00Z | 14 | 20 | 1 | R3IA-c-16 |
| ring-necked pheasant | Game Bird | winter | IA | Audubon/Guthrie | 3 | cropland | R3IA-c-16 | 2016-01-11T00:00:00Z | 4 | 7 | 1 | R3IA-c-16 |
| ring-necked pheasant | Game Bird | winter | IA | Audubon/Guthrie | 3 | cropland | R3IA-c-16 | 2016-01-11T00:00:00Z | 10 | 14 | 1 | R3IA-c-16 |
| ring-necked pheasant | Game Bird | winter | IA | Audubon/Guthrie | 3 | cropland | R3IA-c-16 | 2016-01-25T00:00:00Z | 2 | 3 | 1 | R3IA-c-16 |
| red-tailed hawk | Raptor | winter | IA | Audubon/Guthrie | 3 | cropland | R3IA-c-16 | 2015-12-29T00:00:00Z | 20 | 30 | 1 | R3IA-c-0 |
| red-tailed hawk | Raptor | winter | IA | Audubon/Guthrie | 3 | cropland | R3IA-c-16 | 2015-12-29T00:00:00Z | 60 | NA | 1 | R3IA-c-0 |
| ring-necked pheasant | Game Bird | winter | IA | Audubon/Guthrie | 3 | cropland | R3IA-c-16 | 2016-02-09T00:00:00Z | 4 | 7 | 1 | R3IA-c-16 |
| ring-necked pheasant | Game Bird | winter | IA | Audubon/Guthrie | 3 | cropland | R3IA-c-16 | 2016-02-09T00:00:00Z | 10 | 14 | 1 | R3IA-c-16 |
| ring-necked pheasant | Game Bird | winter | IA | O'Brien | 3 | cropland | R3IA-c-17 | 2015-12-01T00:00:00Z | 7 | 9 | 1 | R3IA-c-17 |
| ring-necked pheasant | Game Bird | winter | IA | O'Brien | 3 | cropland | R3IA-c-17 | 2015-12-01T00:00:00Z | 9 | 13 | 1 | R3IA-c-17 |
| ring-necked pheasant | Game Bird | winter | IA | O'Brien | 3 | cropland | R3IA-c-17 | 2015-12-14T00:00:00Z | 31 | NA | 1 | R3IA-c-17 |
| ring-necked pheasant | Game Bird | winter | IA | O'Brien | 3 | cropland | R3IA-c-17 | 2015-12-14T00:00:00Z | 4 | 7 | 1 | R3IA-c-17 |
| ring-necked pheasant | Game Bird | winter | IA | O'Brien | 3 | cropland | R3IA-c-17 | 2015-12-14T00:00:00Z | 31 | NA | 1 | R3IA-c-17 |
| ring-necked pheasant | Game Bird | winter | IA | O'Brien | 3 | cropland | R3IA-c-17 | 2016-01-11T00:00:00Z | 34 | NA | 1 | R3IA-c-17 |
| ring-necked pheasant | Game Bird | winter | IA | O'Brien | 3 | cropland | R3IA-c-17 | 2016-01-26T00:00:00Z | 6 | 21 | 1 | R3IA-c-17 |
| ring-necked pheasant | Game Bird | winter | IA | O'Brien | 3 | cropland | R3IA-c-17 | 2016-02-10T00:00:00Z | 30 | NA | 1 | R3IA-c-17 |
| ring-necked pheasant | Game Bird | spring | IA | O'Brien | 3 | cropland | R3IA-c-17 | 2016-03-21T00:00:00Z | 1 | 2 | 1 | R3IA-c-17 |
| ring-necked pheasant | Game Bird | spring | IA | O'Brien | 3 | cropland | R3IA-c-17 | 2016-03-21T00:00:00Z | 3 | 4 | 1 | R3IA-c-17 |
| ring-necked pheasant | Game Bird | spring | IA | O'Brien | 3 | cropland | R3IA-c-17 | 2016-03-21T00:00:00Z | 30 | NA | 1 | R3IA-c-17 |
| red-tailed hawk | Raptor | spring | IA | O'Brien | 3 | cropland | R3IA-c-17 | 2016-03-27T00:00:00Z | 30 | NA | 1 | R3IA-c-17 |
| ring-necked pheasant | Game Bird | spring | IA | O'Brien | 3 | cropland | R3IA-c-17 | 2016-03-28T00:00:00Z | 14 | 20 | 1 | R3IA-c-17 |
| ring-necked pheasant | Game Bird | spring | IA | O'Brien | 3 | cropland | R3IA-c-17 | 2016-03-28T00:00:00Z | 14 | 20 | 1 | R3IA-c-17 |
| ring-necked pheasant | Game Bird | spring | IA | O'Brien | 3 | cropland | R3IA-c-17 | 2016-04-04T00:00:00Z | 7 | 10 | 1 | R3IA-c-17 |
| red-tailed hawk | Raptor | spring | IA | O'Brien | 3 | cropland | R3IA-c-17 | 2016-04-11T00:00:00Z | 4 | 7 | 1 | R3IA-c-17 |
| red-tailed hawk | Raptor | spring | IA | O'Brien | 3 | cropland | R3IA-c-17 | 2016-04-11T00:00:00Z | 30 | NA | 1 | R3IA-c-17 |
| red-tailed hawk | Raptor | spring | IA | O'Brien | 3 | cropland | R3IA-c-17 | 2016-04-11T00:00:00Z | 20 | 30 | 1 | R3IA-c-17 |
| red-tailed hawk | Raptor | spring | IA | O'Brien | 3 | cropland | R3IA-c-17 | 2016-04-13T00:00:00Z | 31 | NA | 1 | R3IA-c-17 |
| red-tailed hawk | Raptor | spring | IA | O'Brien | 3 | cropland | R3IA-c-17 | 2016-04-13T00:00:00Z | 31 | NA | 1 | R3IA-c-17 |
| snowy owl | Raptor | spring | IA | O'Brien | 3 | cropland | R3IA-c-17 | 2016-04-13T00:00:00Z | 31 | NA | 1 | R3IA-c-17 |
| ring-necked pheasant | Game Bird | summer | IA | O'Brien | 3 | cropland | R3IA-c-17 | 2016-05-16T00:00:00Z | 4 | 7 | 1 | R3IA-c-17 |
| ring-necked pheasant | Game Bird | summer | IA | O'Brien | 3 | cropland | R3IA-c-17 | 2016-05-24T00:00:00Z | 3 | 4 | 1 | R3IA-c-17 |
| ring-necked pheasant | Game Bird | summer | IA | O'Brien | 3 | cropland | R3IA-c-17 | 2016-05-30T00:00:00Z | 15 | 21 | 1 | R3IA-c-17 |
| ring-necked pheasant | Game Bird | fall | IA | O'Brien | 3 | cropland | R3IA-c-17 | 2016-07-18T00:00:00Z | 1 | 1 | 1 | R3IA-c-17 |
| red-tailed hawk | Raptor | fall | IA | O'Brien | 3 | cropland | R3IA-c-17 | 2016-07-18T00:00:00Z | 20 | 30 | 1 | R3IA-c-17 |
| red-tailed hawk | Raptor | fall | IA | O'Brien | 3 | cropland | R3IA-c-17 | 2016-08-15T00:00:00Z | 30 | NA | 1 | R3IA-c-17 |
| ring-necked pheasant | Game Bird | fall | IA | O'Brien | 3 | cropland | R3IA-c-17 | 2016-08-15T00:00:00Z | 30 | NA | 1 | R3IA-c-17 |
| red-tailed hawk | Raptor | fall | IA | O'Brien | 3 | cropland | R3IA-c-17 | 2016-08-29T00:00:00Z | 3 | 4 | 1 | R3IA-c-17 |
| red-tailed hawk | Raptor | fall | IA | O'Brien | 3 | cropland | R3IA-c-17 | 2016-09-12T00:00:00Z | 61 | NA | 1 | R3IA-c-17 |
| ring-necked pheasant | Game Bird | fall | IA | O'Brien | 3 | cropland | R3IA-c-17 | 2016-09-12T00:00:00Z | 30 | NA | 1 | R3IA-c-17 |
| ring-necked pheasant | Game Bird | winter | IA | O'Brien | 3 | cropland | R3IA-c-17 | 2016-11-28T00:00:00Z | 20 | 30 | 1 | R3IA-c-17 |
| ring-necked pheasant | Game Bird | winter | IA | O'Brien | 3 | cropland | R3IA-c-17 | 2016-11-28T00:00:00Z | 30 | NA | 1 | R3IA-c-17 |
| ring-necked pheasant | Game Bird | winter | IA | O'Brien | 3 | cropland | R3IA-c-17 | 2016-12-12T00:00:00Z | 15 | 22 | 1 | R3IA-c-17 |
| ring-necked pheasant | Game Bird | winter | IA | O'Brien | 3 | cropland | R3IA-c-17 | 2016-12-12T00:00:00Z | 15 | 22 | 1 | R3IA-c-17 |
| ring-necked pheasant | Game Bird | winter | IA | O'Brien | 3 | cropland | R3IA-c-17 | 2017-01-09T00:00:00Z | 3 | 3 | 1 | R3IA-c-17 |
| ring-necked pheasant | Game Bird | winter | IA | O'Brien | 3 | cropland | R3IA-c-17 | 2017-01-09T00:00:00Z | 3 | 3 | 1 | R3IA-c-17 |
| ring-necked pheasant | Game Bird | winter | IA | O'Brien | 3 | cropland | R3IA-c-17 | 2017-01-23T00:00:00Z | 14 | 20 | 1 | R3IA-c-17 |
| ring-necked pheasant | Game Bird | winter | IA | O'Brien | 3 | cropland | R3IA-c-17 | 2017-01-23T00:00:00Z | 10 | 10 | 1 | R3IA-c-17 |
| ring-necked pheasant | Game Bird | winter | IA | O'Brien | 3 | cropland | R3IA-c-17 | 2017-02-06T00:00:00Z | 7 | 14 | 1 | R3IA-c-17 |
| ring-necked pheasant | Game Bird | winter | IA | O'Brien | 3 | cropland | R3IA-c-17 | 2017-02-06T00:00:00Z | 7 | 10 | 1 | R3IA-c-17 |
| ring-necked pheasant | Game Bird | spring | IA | Ida | 3 | cropland | R3IA-c-9 | 2017-04-03T00:00:00Z | 13.9847222 | 19.9402778 | 1 | R3IA-c-9 |
| ring-necked pheasant | Game Bird | spring | IA | Ida | 3 | cropland | R3IA-c-9 | 2017-04-03T00:00:00Z | 4.0277778 | 7.1166667 | 1 | R3IA-c-9 |
| ring-necked pheasant | Game Bird | winter | IA | Ida | 3 | cropland | R3IA-c-9 | 2018-01-17T00:00:00Z | 30.0104167 | NA | 1 | R3IA-c-9 |
| ring-necked pheasant | Game Bird | spring | IA | Ida | 3 | cropland | R3IA-c-9 | 2017-03-28T00:00:00Z | 0.9833333 | 1.8375 | 1 | R3IA-c-9 |
| ring-necked pheasant | Game Bird | spring | IA | Ida | 3 | cropland | R3IA-c-9 | 2017-03-28T00:00:00Z | 2.9729167 | 4.0798611 | 1 | R3IA-c-9 |
| ring-necked pheasant | Game Bird | spring | IA | Ida | 3 | cropland | R3IA-c-9 | 2017-03-28T00:00:00Z | 4.2840278 | 7.2451389 | 1 | R3IA-c-9 |
| ring-necked pheasant | Game Bird | spring | IA | Ida | 3 | cropland | R3IA-c-9 | 2017-03-28T00:00:00Z | 2.9465278 | 3.95 | 1 | R3IA-c-9 |
| ring-necked pheasant | Game Bird | spring | IA | Ida | 3 | cropland | R3IA-c-9 | 2017-03-28T00:00:00Z | 10.1805556 | 13.9104167 | 1 | R3IA-c-9 |
| ring-necked pheasant | Game Bird | spring | IA | Ida | 3 | cropland | R3IA-c-9 | 2017-04-03T00:00:00Z | 13.9847222 | 19.9402778 | 1 | R3IA-c-9 |
| ring-necked pheasant | Game Bird | spring | IA | Ida | 3 | cropland | R3IA-c-9 | 2017-04-03T00:00:00Z | 4.0277778 | 7.1166667 | 1 | R3IA-c-9 |
| turkey vulture | Raptor | summer | IA | Ida | 3 | cropland | R3IA-c-9 | 2017-06-05T00:00:00Z | 20.0930556 | 30.0638889 | 1 | R3IA-c-9 |
| turkey vulture | Raptor | summer | IA | Ida | 3 | cropland | R3IA-c-9 | 2017-06-11T00:00:00Z | 30.1284722 | NA | 1 | R3IA-c-9 |
| red-tailed hawk | Raptor | summer | IA | Ida | 3 | cropland | R3IA-c-9 | 2017-07-02T00:00:00Z | 3.0798611 | 3.0798611 | 1 | R3IA-c-9 |
| turkey vulture | Raptor | summer | IA | Ida | 3 | cropland | R3IA-c-9 | 2017-07-03T00:00:00Z | 31.6027778 | NA | 1 | R3IA-c-9 |
| red-tailed hawk | Raptor | winter | IA | Ida | 3 | cropland | R3IA-c-9 | 2017-12-05T00:00:00Z | 42.19375 | 50.2986111 | 1 | R3IA-c-9 |
| red-tailed hawk | Raptor | winter | IA | Ida | 3 | cropland | R3IA-c-9 | 2017-12-05T00:00:00Z | 9.9520833 | 15.2798611 | 1 | R3IA-c-9 |
| red-tailed hawk | Raptor | winter | IA | Ida | 3 | cropland | R3IA-c-9 | 2017-12-05T00:00:00Z | 15.3729167 | 20.33125 | 1 | R3IA-c-9 |
| red-tailed hawk | Raptor | winter | IA | Ida | 3 | cropland | R3IA-c-9 | 2017-12-05T00:00:00Z | 9.9486111 | 15.2784722 | 1 | R3IA-c-9 |
| red-tailed hawk | Raptor | winter | IA | Ida | 3 | cropland | R3IA-c-9 | 2017-12-05T00:00:00Z | 60.3027778 | NA | 1 | R3IA-c-9 |
| red-tailed hawk | Raptor | winter | IA | Ida | 3 | cropland | R3IA-c-9 | 2017-12-05T00:00:00Z | 60.2284722 | NA | 1 | R3IA-c-9 |
| red-tailed hawk | Raptor | winter | IA | Ida | 3 | cropland | R3IA-c-9 | 2017-12-05T00:00:00Z | 15.4006944 | 20.3604167 | 1 | R3IA-c-9 |
| red-tailed hawk | Raptor | winter | IA | Ida | 3 | cropland | R3IA-c-9 | 2017-12-05T00:00:00Z | 2.1298611 | 7.1861111 | 1 | R3IA-c-9 |
| red-tailed hawk | Raptor | winter | IA | Ida | 3 | cropland | R3IA-c-9 | 2017-12-05T00:00:00Z | 10.1236111 | 15.2784722 | 1 | R3IA-c-9 |
| red-tailed hawk | Raptor | winter | IA | Ida | 3 | cropland | R3IA-c-9 | 2017-12-05T00:00:00Z | 60.3118056 | NA | 1 | R3IA-c-9 |
| red-tailed hawk | Raptor | winter | IA | Ida | 3 | cropland | R3IA-c-9 | 2017-12-05T00:00:00Z | 20.3354167 | 30.1027778 | 1 | R3IA-c-9 |
| red-tailed hawk | Raptor | winter | IA | Ida | 3 | cropland | R3IA-c-9 | 2018-01-02T00:00:00Z | 7.2861111 | 10.0826389 | 1 | R3IA-c-9 |
| red-tailed hawk | Raptor | winter | IA | Ida | 3 | cropland | R3IA-c-9 | 2018-01-02T00:00:00Z | 60.0368056 | NA | 1 | R3IA-c-9 |
| red-tailed hawk | Raptor | winter | IA | Ida | 3 | cropland | R3IA-c-9 | 2018-01-02T00:00:00Z | 14.0784722 | 21.0402778 | 1 | R3IA-c-9 |
| red-tailed hawk | Raptor | winter | IA | Ida | 3 | cropland | R3IA-c-9 | 2018-01-02T00:00:00Z | 50.2875 | 60.0333333 | 1 | R3IA-c-9 |
| red-tailed hawk | Raptor | winter | IA | Ida | 3 | cropland | R3IA-c-9 | 2018-01-02T00:00:00Z | 14.0659722 | 21.0180556 | 1 | R3IA-c-9 |
| red-tailed hawk | Raptor | winter | IA | Ida | 3 | cropland | R3IA-c-9 | 2018-01-02T00:00:00Z | 10.0319444 | 14.0604167 | 1 | R3IA-c-9 |
| red-tailed hawk | Raptor | winter | IA | Ida | 3 | cropland | R3IA-c-9 | 2018-01-02T00:00:00Z | 50.2673611 | 60.0131944 | 1 | R3IA-c-9 |
| red-tailed hawk | Raptor | winter | IA | Ida | 3 | cropland | R3IA-c-9 | 2018-01-02T00:00:00Z | 60.0090278 | NA | 1 | R3IA-c-9 |
| red-tailed hawk | Raptor | winter | IA | Ida | 3 | cropland | R3IA-c-9 | 2018-01-02T00:00:00Z | 30.0194444 | 41.0694444 | 1 | R3IA-c-9 |
| red-tailed hawk | Raptor | winter | IA | Ida | 3 | cropland | R3IA-c-9 | 2018-01-02T00:00:00Z | 59.9986111 | NA | 1 | R3IA-c-9 |
| red-tailed hawk | Raptor | winter | IA | Ida | 3 | cropland | R3IA-c-9 | 2018-01-02T00:00:00Z | 59.9958333 | NA | 1 | R3IA-c-9 |
| red-tailed hawk | Raptor | winter | IA | Ida | 3 | cropland | R3IA-c-9 | 2018-01-08T00:00:00Z | 3.2951389 | 8.1 | 1 | R3IA-c-9 |
| red-tailed hawk | Raptor | winter | IA | Ida | 3 | cropland | R3IA-c-9 | 2018-01-08T00:00:00Z | 3.3270833 | 8.1305556 | 1 | R3IA-c-9 |
| red-tailed hawk | Raptor | winter | IA | Ida | 3 | cropland | R3IA-c-9 | 2018-01-08T00:00:00Z | 10.2791667 | 15.1888889 | 1 | R3IA-c-9 |
| red-tailed hawk | Raptor | winter | IA | Ida | 3 | cropland | R3IA-c-9 | 2018-01-08T00:00:00Z | 10.2833333 | 21.2555556 | 1 | R3IA-c-9 |
| red-tailed hawk | Raptor | winter | IA | Ida | 3 | cropland | R3IA-c-9 | 2018-01-08T00:00:00Z | 60.2972222 | NA | 1 | R3IA-c-9 |
| red-tailed hawk | Raptor | winter | IA | Ida | 3 | cropland | R3IA-c-9 | 2018-01-08T00:00:00Z | 40.1805556 | 50.1798611 | 1 | R3IA-c-9 |
| red-tailed hawk | Raptor | winter | IA | Ida | 3 | cropland | R3IA-c-9 | 2018-01-08T00:00:00Z | 10.2736111 | 21.2430556 | 1 | R3IA-c-9 |
| red-tailed hawk | Raptor | winter | IA | Ida | 3 | cropland | R3IA-c-9 | 2018-01-08T00:00:00Z | 10.2902778 | 21.2527778 | 1 | R3IA-c-9 |
| red-tailed hawk | Raptor | winter | IA | Ida | 3 | cropland | R3IA-c-9 | 2018-01-16T00:00:00Z | 14.3006944 | 30.2263889 | 1 | R3IA-c-9 |
| red-tailed hawk | Raptor | winter | IA | Ida | 3 | cropland | R3IA-c-9 | 2018-01-16T00:00:00Z | 3.1645833 | 10.1201389 | 1 | R3IA-c-9 |
| red-tailed hawk | Raptor | winter | IA | Ida | 3 | cropland | R3IA-c-9 | 2018-01-16T00:00:00Z | 14.2097222 | 30.21875 | 1 | R3IA-c-9 |
| red-tailed hawk | Raptor | winter | IA | Ida | 3 | cropland | R3IA-c-9 | 2018-01-16T00:00:00Z | 1.2375 | 3.1569444 | 1 | R3IA-c-9 |
| red-tailed hawk | Raptor | winter | IA | Ida | 3 | cropland | R3IA-c-9 | 2018-01-16T00:00:00Z | 30.2298611 | 60.1951389 | 1 | R3IA-c-9 |
| red-tailed hawk | Raptor | winter | IA | Ida | 3 | cropland | R3IA-c-9 | 2018-01-16T00:00:00Z | 10.1798611 | 14.3645833 | 1 | R3IA-c-9 |
| red-tailed hawk | Raptor | winter | IA | Ida | 3 | cropland | R3IA-c-9 | 2018-01-16T00:00:00Z | 3.2326389 | 10.1895833 | 1 | R3IA-c-9 |
| red-tailed hawk | Raptor | winter | IA | Ida | 3 | cropland | R3IA-c-9 | 2018-01-16T00:00:00Z | 60.25625 | NA | 1 | R3IA-c-9 |
| red-tailed hawk | Raptor | winter | IA | Ida | 3 | cropland | R3IA-c-9 | 2018-01-16T00:00:00Z | 3.15625 | 10.1125 | 1 | R3IA-c-9 |
| red-tailed hawk | Raptor | winter | IA | Ida | 3 | cropland | R3IA-c-9 | 2018-01-16T00:00:00Z | 14.4097222 | 40.3173611 | 1 | R3IA-c-9 |
| red-tailed hawk | Raptor | winter | IA | Ida | 3 | cropland | R3IA-c-9 | 2018-01-16T00:00:00Z | 14.3298611 | 30.2527778 | 1 | R3IA-c-9 |
| red-tailed hawk | Raptor | winter | IA | Ida | 3 | cropland | R3IA-c-9 | 2018-01-16T00:00:00Z | 3.1569444 | 10.1104167 | 1 | R3IA-c-9 |
| red-tailed hawk | Raptor | winter | IA | Ida | 3 | cropland | R3IA-c-9 | 2018-01-16T00:00:00Z | 10.2243056 | 14.4104167 | 1 | R3IA-c-9 |
| red-tailed hawk | Raptor | winter | IA | Ida | 3 | cropland | R3IA-c-9 | 2018-01-16T00:00:00Z | 14.3756944 | 30.2958333 | 1 | R3IA-c-9 |
| red-tailed hawk | Raptor | winter | IA | Ida | 3 | cropland | R3IA-c-9 | 2018-01-25T00:00:00Z | 20.2659722 | 40.3527778 | 1 | R3IA-c-9 |
| red-tailed hawk | Raptor | winter | IA | Ida | 3 | cropland | R3IA-c-9 | 2018-01-25T00:00:00Z | 60.4958333 | NA | 1 | R3IA-c-9 |
| red-tailed hawk | Raptor | winter | IA | Ida | 3 | cropland | R3IA-c-9 | 2018-01-25T00:00:00Z | 10.2006944 | 20.2326389 | 1 | R3IA-c-9 |
| red-tailed hawk | Raptor | winter | IA | Ida | 3 | cropland | R3IA-c-9 | 2018-01-25T00:00:00Z | 10.1805556 | 51.2652778 | 1 | R3IA-c-9 |
| red-tailed hawk | Raptor | winter | IA | Ida | 3 | cropland | R3IA-c-9 | 2018-01-25T00:00:00Z | 10.1458333 | 51.2506944 | 1 | R3IA-c-9 |
| red-tailed hawk | Raptor | winter | IA | Ida | 3 | cropland | R3IA-c-9 | 2018-01-25T00:00:00Z | 10.2041667 | 51.2666667 | 1 | R3IA-c-9 |
| red-tailed hawk | Raptor | winter | IA | Ida | 3 | cropland | R3IA-c-9 | 2018-01-25T00:00:00Z | 51.3048611 | 60.5020833 | 1 | R3IA-c-9 |
| red-tailed hawk | Raptor | winter | IA | Ida | 3 | cropland | R3IA-c-9 | 2018-01-25T00:00:00Z | 4.3173611 | 7.3326389 | 1 | R3IA-c-9 |
| red-tailed hawk | Raptor | winter | IA | Ida | 3 | cropland | R3IA-c-9 | 2018-01-25T00:00:00Z | 7.3041667 | 10.1881944 | 1 | R3IA-c-9 |
| red-tailed hawk | Raptor | winter | IA | Ida | 3 | cropland | R3IA-c-9 | 2018-01-25T00:00:00Z | 10.1611111 | 51.2541667 | 1 | R3IA-c-9 |
| red-tailed hawk | Raptor | winter | IA | Ida | 3 | cropland | R3IA-c-9 | 2018-02-02T00:00:00Z | 14.3083333 | 32.2201389 | 1 | R3IA-c-9 |
| red-tailed hawk | Raptor | winter | IA | Ida | 3 | cropland | R3IA-c-9 | 2018-02-02T00:00:00Z | 14.3736111 | 40.4347222 | 1 | R3IA-c-9 |
| red-tailed hawk | Raptor | winter | IA | Ida | 3 | cropland | R3IA-c-9 | 2018-02-02T00:00:00Z | 1.2520833 | 7.0152778 | 1 | R3IA-c-9 |
| red-tailed hawk | Raptor | winter | IA | Ida | 3 | cropland | R3IA-c-9 | 2018-02-02T00:00:00Z | 1.2541667 | 14.3729167 | 1 | R3IA-c-9 |
| red-tailed hawk | Raptor | winter | IA | Ida | 3 | cropland | R3IA-c-9 | 2018-02-02T00:00:00Z | 1.2409722 | 10.0993056 | 1 | R3IA-c-9 |
| red-tailed hawk | Raptor | winter | IA | Ida | 3 | cropland | R3IA-c-9 | 2018-02-02T00:00:00Z | 0.9368056 | 14.0423611 | 1 | R3IA-c-9 |
| red-tailed hawk | Raptor | winter | IA | Ida | 3 | cropland | R3IA-c-9 | 2018-02-02T00:00:00Z | 14.0395833 | 40.0993056 | 1 | R3IA-c-9 |
| ring-necked pheasant | Game Bird | summer | IA | Sac/Buena Vista | 3 | cropland | R3IA-c-19 | 2016-05-16T00:00:00Z | 10 | 14 | 1 | R3IA-c-19 |
| ring-necked pheasant | Game Bird | summer | IA | Sac/Buena Vista | 3 | cropland | R3IA-c-19 | 2016-05-24T00:00:00Z | 20 | 30 | 1 | R3IA-c-19 |
| ring-necked pheasant | Game Bird | summer | IA | Sac/Buena Vista | 3 | cropland | R3IA-c-19 | 2016-05-24T00:00:00Z | 20 | 30 | 1 | R3IA-c-19 |
| ring-necked pheasant | Game Bird | summer | IA | Sac/Buena Vista | 3 | cropland | R3IA-c-19 | 2016-05-24T00:00:00Z | 4 | 7 | 1 | R3IA-c-19 |
| ring-necked pheasant | Game Bird | summer | IA | Sac/Buena Vista | 3 | cropland | R3IA-c-19 | 2016-05-24T00:00:00Z | 2 | 3 | 1 | R3IA-c-19 |
| ring-necked pheasant | Game Bird | winter | IA | Sac/Buena Vista | 3 | cropland | R3IA-c-19 | 2015-12-01T00:00:00Z | 4 | 7 | 1 | R3IA-c-19 |
| ring-necked pheasant | Game Bird | winter | IA | Sac/Buena Vista | 3 | cropland | R3IA-c-19 | 2015-12-14T00:00:00Z | 10 | 31 | 1 | R3IA-c-19 |
| ring-necked pheasant | Game Bird | winter | IA | Sac/Buena Vista | 3 | cropland | R3IA-c-19 | 2015-12-14T00:00:00Z | 4 | 7 | 1 | R3IA-c-19 |
| ring-necked pheasant | Game Bird | winter | IA | Sac/Buena Vista | 3 | cropland | R3IA-c-19 | 2015-12-29T00:00:00Z | 7 | NA | 1 | R3IA-c-19 |
| ring-necked pheasant | Game Bird | winter | IA | Sac/Buena Vista | 3 | cropland | R3IA-c-19 | 2016-01-11T00:00:00Z | 30 | NA | 1 | R3IA-c-19 |
| ring-necked pheasant | Game Bird | winter | IA | Sac/Buena Vista | 3 | cropland | R3IA-c-19 | 2016-01-26T00:00:00Z | 1 | 2 | 1 | R3IA-c-19 |
| ring-necked pheasant | Game Bird | winter | IA | Sac/Buena Vista | 3 | cropland | R3IA-c-19 | 2016-02-09T00:00:00Z | 30 | NA | 1 | R3IA-c-19 |
| ring-necked pheasant | Game Bird | spring | IA | Sac/Buena Vista | 3 | cropland | R3IA-c-19 | 2016-03-21T00:00:00Z | 7 | 10 | 1 | R3IA-c-19 |
| ring-necked pheasant | Game Bird | spring | IA | Sac/Buena Vista | 3 | cropland | R3IA-c-19 | 2016-03-28T00:00:00Z | 14 | 20 | 1 | R3IA-c-19 |
| ring-necked pheasant | Game Bird | spring | IA | Sac/Buena Vista | 3 | cropland | R3IA-c-19 | 2016-04-04T00:00:00Z | 14 | 30 | 1 | R3IA-c-19 |
| red-tailed hawk | Raptor | spring | IA | Sac/Buena Vista | 3 | cropland | R3IA-c-19 | 2016-04-11T00:00:00Z | 20 | 30 | 1 | R3IA-c-19 |
| red-tailed hawk | Raptor | spring | IA | Sac/Buena Vista | 3 | cropland | R3IA-c-19 | 2016-04-11T00:00:00Z | 30 | NA | 1 | R3IA-c-19 |
| red-tailed hawk | Raptor | spring | IA | Sac/Buena Vista | 3 | cropland | R3IA-c-19 | 2016-04-11T00:00:00Z | 30 | NA | 1 | R3IA-c-19 |
| red-tailed hawk | Raptor | spring | IA | Sac/Buena Vista | 3 | cropland | R3IA-c-19 | 2016-04-13T00:00:00Z | 30 | NA | 1 | R3IA-c-19 |
| red-tailed hawk | Raptor | spring | IA | Sac/Buena Vista | 3 | cropland | R3IA-c-19 | 2016-04-13T00:00:00Z | 30 | NA | 1 | R3IA-c-19 |
| ring-necked pheasant | Game Bird | spring | IA | Sac/Buena Vista | 3 | cropland | R3IA-c-19 | 2016-04-13T00:00:00Z | 7 | 10 | 1 | R3IA-c-19 |
| ring-necked pheasant | Game Bird | summer | IA | Sac/Buena Vista | 3 | cropland | R3IA-c-19 | 2016-05-24T00:00:00Z | 10 | 14 | 1 | R3IA-c-19 |
| ring-necked pheasant | Game Bird | summer | IA | Sac/Buena Vista | 3 | cropland | R3IA-c-19 | 2016-05-31T00:00:00Z | 30 | NA | 1 | R3IA-c-19 |
| ring-necked pheasant | Game Bird | summer | IA | Sac/Buena Vista | 3 | cropland | R3IA-c-19 | 2016-05-31T00:00:00Z | 1 | 1 | 1 | R3IA-c-19 |
| ring-necked pheasant | Game Bird | summer | IA | Sac/Buena Vista | 3 | cropland | R3IA-c-19 | 2016-05-31T00:00:00Z | 4 | 7 | 1 | R3IA-c-19 |
| ring-necked pheasant | Game Bird | summer | IA | Sac/Buena Vista | 3 | cropland | R3IA-c-19 | 2016-05-31T00:00:00Z | 10 | 15 | 1 | R3IA-c-19 |
| ring-necked pheasant | Game Bird | summer | IA | Sac/Buena Vista | 3 | cropland | R3IA-c-19 | 2016-05-31T00:00:00Z | 3 | 4 | 1 | R3IA-c-19 |
| ring-necked pheasant | Game Bird | summer | IA | Sac/Buena Vista | 3 | cropland | R3IA-c-19 | 2016-06-13T00:00:00Z | 3 | 4 | 1 | R3IA-c-19 |
| ring-necked pheasant | Game Bird | summer | IA | Sac/Buena Vista | 3 | cropland | R3IA-c-19 | 2016-06-13T00:00:00Z | 3 | 4 | 1 | R3IA-c-19 |
| ring-necked pheasant | Game Bird | summer | IA | Sac/Buena Vista | 3 | cropland | R3IA-c-19 | 2016-06-13T00:00:00Z | 7 | 10 | 1 | R3IA-c-19 |
| ring-necked pheasant | Game Bird | summer | IA | Sac/Buena Vista | 3 | cropland | R3IA-c-19 | 2016-06-13T00:00:00Z | 3 | 4 | 1 | R3IA-c-19 |
| red-tailed hawk | Raptor | fall | IA | Sac/Buena Vista | 3 | cropland | R3IA-c-19 | 2016-08-15T00:00:00Z | 30 | 37 | 1 | R3IA-c-19 |
| red-tailed hawk | Raptor | fall | IA | Sac/Buena Vista | 3 | cropland | R3IA-c-19 | 2016-08-29T00:00:00Z | 37 | 44 | 1 | R3IA-c-19 |
| ring-necked pheasant | Game Bird | fall | IA | Sac/Buena Vista | 3 | cropland | R3IA-c-19 | 2016-08-29T00:00:00Z | 10 | NA | 1 | R3IA-c-19 |
| red-tailed hawk | Raptor | fall | IA | Sac/Buena Vista | 3 | cropland | R3IA-c-19 | 2016-09-12T00:00:00Z | 61 | NA | 1 | R3IA-c-19 |
| red-tailed hawk | Raptor | fall | IA | Sac/Buena Vista | 3 | cropland | R3IA-c-19 | 2016-09-12T00:00:00Z | 61 | NA | 1 | R3IA-c-19 |
| ring-necked pheasant | Game Bird | fall | IA | Sac/Buena Vista | 3 | cropland | R3IA-c-19 | 2016-10-10T00:00:00Z | 1 | 1 | 1 | R3IA-c-19 |
| ring-necked pheasant | Game Bird | winter | IA | Sac/Buena Vista | 3 | cropland | R3IA-c-19 | 2016-12-05T00:00:00Z | 30 | NA | 1 | R3IA-c-19 |
| ring-necked pheasant | Game Bird | winter | IA | Sac/Buena Vista | 3 | cropland | R3IA-c-19 | 2016-12-05T00:00:00Z | 16 | 22 | 1 | R3IA-c-19 |
| red-tailed hawk | Raptor | winter | IA | Sac/Buena Vista | 3 | cropland | R3IA-c-19 | 2017-01-02T00:00:00Z | 10 | 17 | 1 | R3IA-c-19 |
| ring-necked pheasant | Game Bird | winter | IA | Sac/Buena Vista | 3 | cropland | R3IA-c-19 | 2017-01-09T00:00:00Z | 14 | 21 | 1 | R3IA-c-19 |
| ring-necked pheasant | Game Bird | winter | IA | Sac/Buena Vista | 3 | cropland | R3IA-c-19 | 2017-01-09T00:00:00Z | 30 | NA | 1 | R3IA-c-19 |
| ring-necked pheasant | Game Bird | winter | IA | Sac/Buena Vista | 3 | cropland | R3IA-c-19 | 2017-01-30T00:00:00Z | 7 | 10 | 1 | R3IA-c-19 |
| ring-necked pheasant | Game Bird | winter | IA | Sac/Buena Vista | 3 | cropland | R3IA-c-19 | 2017-01-30T00:00:00Z | 3 | 4 | 1 | R3IA-c-19 |
| ring-necked pheasant | Game Bird | winter | IA | Sac/Buena Vista | 3 | cropland | R3IA-c-19 | 2017-02-06T00:00:00Z | 2 | 2 | 1 | R3IA-c-19 |
| ring-necked pheasant | Game Bird | winter | IA | Sac/Buena Vista | 3 | cropland | R3IA-c-19 | 2017-02-06T00:00:00Z | 10 | 15 | 1 | R3IA-c-19 |
| ring-necked pheasant | Game Bird | winter | IA | Sac/Buena Vista | 3 | cropland | R3IA-c-19 | 2017-01-23T00:00:00Z | 30 | NA | 1 | R3IA-c-19 |
| ring-necked pheasant | Game Bird | winter | IA | Sac/Buena Vista | 3 | cropland | R3IA-c-19 | 2017-01-23T00:00:00Z | 14 | 21 | 1 | R3IA-c-19 |
| ring-necked pheasant | Game Bird | winter | IA | Marshall | 3 | cropland | R3IA-c-20 | 2015-12-07T00:00:00Z | 3 | 4 | 1 | R3IA-c-20 |
| ring-necked pheasant | Game Bird | winter | IA | Marshall | 3 | cropland | R3IA-c-20 | 2015-12-07T00:00:00Z | 7 | 10 | 1 | R3IA-c-20 |
| ring-necked pheasant | Game Bird | winter | IA | Marshall | 3 | cropland | R3IA-c-20 | 2015-12-29T00:00:00Z | 10 | 14 | 1 | R3IA-c-20 |
| ring-necked pheasant | Game Bird | winter | IA | Marshall | 3 | cropland | R3IA-c-20 | 2016-01-04T00:00:00Z | 30 | NA | 1 | R3IA-c-20 |
| ring-necked pheasant | Game Bird | winter | IA | Marshall | 3 | cropland | R3IA-c-20 | 2016-01-18T00:00:00Z | 1 | 2 | 1 | R3IA-c-20 |
| ring-necked pheasant | Game Bird | winter | IA | Marshall | 3 | cropland | R3IA-c-20 | 2016-01-25T00:00:00Z | 3 | 4 | 1 | R3IA-c-20 |
| ring-necked pheasant | Game Bird | winter | IA | Marshall | 3 | cropland | R3IA-c-20 | 2016-02-09T00:00:00Z | 4 | 10 | 1 | R3IA-c-20 |
| turkey vulture | Raptor | spring | IA | Marshall | 3 | cropland | R3IA-c-20 | 2016-03-25T00:00:00Z | 30 | NA | 1 | R3IA-c-0 |
| turkey vulture | Raptor | spring | IA | Marshall | 3 | cropland | R3IA-c-20 | 2016-03-31T00:00:00Z | 30 | NA | 1 | R3IA-c-0 |
| ring-necked pheasant | Game Bird | spring | IA | Marshall | 3 | cropland | R3IA-c-20 | 2016-04-08T00:00:00Z | 4 | 7 | 1 | R3IA-c-20 |
| ring-necked pheasant | Game Bird | spring | IA | Marshall | 3 | cropland | R3IA-c-20 | 2016-04-08T00:00:00Z | 10 | 14 | 1 | R3IA-c-20 |
| ring-necked pheasant | Game Bird | spring | IA | Marshall | 3 | cropland | R3IA-c-20 | 2016-04-08T00:00:00Z | 14 | 20 | 1 | R3IA-c-20 |
| red-tailed hawk | Raptor | spring | IA | Marshall | 3 | cropland | R3IA-c-20 | 2016-04-12T00:00:00Z | 30 | NA | 1 | R3IA-c-0 |
| red-tailed hawk | Raptor | spring | IA | Marshall | 3 | cropland | R3IA-c-20 | 2016-04-12T00:00:00Z | 4 | 7 | 1 | R3IA-c-0 |
| red-tailed hawk | Raptor | spring | IA | Marshall | 3 | cropland | R3IA-c-20 | 2016-04-12T00:00:00Z | 7 | 10 | 1 | R3IA-c-0 |
| ring-necked pheasant | Game Bird | summer | IA | Marshall | 3 | cropland | R3IA-c-20 | 2016-05-24T00:00:00Z | 1 | 2 | 1 | R3IA-c-20 |
| ring-necked pheasant | Game Bird | summer | IA | Marshall | 3 | cropland | R3IA-c-20 | 2016-06-09T00:00:00Z | 30 | NA | 1 | R3IA-c-20 |
| ring-necked pheasant | Game Bird | summer | IA | Marshall | 3 | cropland | R3IA-c-20 | 2016-06-14T00:00:00Z | 2 | 3 | 1 | R3IA-c-20 |
| red-tailed hawk | Raptor | fall | IA | Marshall | 3 | cropland | R3IA-c-20 | 2016-08-01T00:00:00Z | 2 | 4 | 1 | R3IA-c-0 |
| red-tailed hawk | Raptor | fall | IA | Marshall | 3 | cropland | R3IA-c-20 | 2016-08-29T00:00:00Z | 30 | 40 | 1 | R3IA-c-0 |
| red-tailed hawk | Raptor | fall | IA | Marshall | 3 | cropland | R3IA-c-20 | 2016-09-26T00:00:00Z | 14 | 20 | 1 | R3IA-c-0 |
| ring-necked pheasant | Game Bird | winter | IA | Marshall | 3 | cropland | R3IA-c-20 | 2016-11-28T00:00:00Z | 7 | 10 | 1 | R3IA-c-20 |
| ring-necked pheasant | Game Bird | winter | IA | Marshall | 3 | cropland | R3IA-c-20 | 2016-11-28T00:00:00Z | 2 | 3 | 1 | R3IA-c-20 |
| ring-necked pheasant | Game Bird | winter | IA | Marshall | 3 | cropland | R3IA-c-20 | 2016-12-12T00:00:00Z | 30 | NA | 1 | R3IA-c-20 |
| ring-necked pheasant | Game Bird | winter | IA | Marshall | 3 | cropland | R3IA-c-20 | 2016-12-12T00:00:00Z | 10 | 14 | 1 | R3IA-c-20 |
| ring-necked pheasant | Game Bird | winter | IA | Marshall | 3 | cropland | R3IA-c-20 | 2017-01-09T00:00:00Z | 2 | 3 | 1 | R3IA-c-20 |
| ring-necked pheasant | Game Bird | winter | IA | Marshall | 3 | cropland | R3IA-c-20 | 2017-01-09T00:00:00Z | 10 | 14 | 1 | R3IA-c-20 |
| ring-necked pheasant | Game Bird | winter | IA | Marshall | 3 | cropland | R3IA-c-20 | 2017-01-23T00:00:00Z | 1 | 1 | 1 | R3IA-c-20 |
| ring-necked pheasant | Game Bird | winter | IA | Marshall | 3 | cropland | R3IA-c-20 | 2017-01-23T00:00:00Z | 4 | 7 | 1 | R3IA-c-20 |
| ring-necked pheasant | Game Bird | winter | IA | Marshall | 3 | cropland | R3IA-c-20 | 2017-02-06T00:00:00Z | 1 | 3 | 1 | R3IA-c-20 |
| ring-necked pheasant | Game Bird | winter | IA | Marshall | 3 | cropland | R3IA-c-20 | 2017-02-06T00:00:00Z | 1 | 1 | 1 | R3IA-c-20 |
| ring-necked pheasant | Game Bird | winter | IA | Webster | 3 | cropland | R3IA-c-21 | 2016-02-09T00:00:00Z | 14 | 20 | 1 | R3IA-c-21 |
| ring-necked pheasant | Game Bird | winter | IA | Webster | 3 | cropland | R3IA-c-21 | 2015-12-01T00:00:00Z | 3 | 4 | 1 | R3IA-c-21 |
| ring-necked pheasant | Game Bird | winter | IA | Webster | 3 | cropland | R3IA-c-21 | 2015-12-01T00:00:00Z | 20 | 30 | 1 | R3IA-c-21 |
| ring-necked pheasant | Game Bird | winter | IA | Webster | 3 | cropland | R3IA-c-21 | 2015-12-14T00:00:00Z | 9 | 15 | 1 | R3IA-c-21 |
| ring-necked pheasant | Game Bird | winter | IA | Webster | 3 | cropland | R3IA-c-21 | 2016-01-06T00:00:00Z | 2 | 7 | 1 | R3IA-c-21 |
| ring-necked pheasant | Game Bird | winter | IA | Webster | 3 | cropland | R3IA-c-21 | 2016-01-06T00:00:00Z | 20 | 34 | 1 | R3IA-c-21 |
| red-tailed hawk | Raptor | winter | IA | Webster | 3 | cropland | R3IA-c-21 | 2016-01-06T00:00:00Z | 10 | 14 | 1 | R3IA-c-0 |
| red-tailed hawk | Raptor | winter | IA | Webster | 3 | cropland | R3IA-c-21 | 2016-01-06T00:00:00Z | 65 | NA | 1 | R3IA-c-0 |
| red-tailed hawk | Raptor | winter | IA | Webster | 3 | cropland | R3IA-c-21 | 2016-01-06T00:00:00Z | 65 | NA | 1 | R3IA-c-0 |
| ring-necked pheasant | Game Bird | winter | IA | Webster | 3 | cropland | R3IA-c-21 | 2016-01-06T00:00:00Z | 34 | NA | 1 | R3IA-c-21 |
| ring-necked pheasant | Game Bird | winter | IA | Webster | 3 | cropland | R3IA-c-21 | 2016-01-12T00:00:00Z | 20 | 31 | 1 | R3IA-c-21 |
| ring-necked pheasant | Game Bird | winter | IA | Webster | 3 | cropland | R3IA-c-21 | 2016-01-12T00:00:00Z | 1 | 1 | 1 | R3IA-c-21 |
| ring-necked pheasant | Game Bird | winter | IA | Webster | 3 | cropland | R3IA-c-21 | 2016-01-12T00:00:00Z | 20 | 31 | 1 | R3IA-c-21 |
| ring-necked pheasant | Game Bird | winter | IA | Webster | 3 | cropland | R3IA-c-21 | 2016-01-12T00:00:00Z | 20 | 31 | 1 | R3IA-c-21 |
| ring-necked pheasant | Game Bird | winter | IA | Webster | 3 | cropland | R3IA-c-21 | 2015-12-01T00:00:00Z | 10 | 13 | 1 | R3IA-c-21 |
| ring-necked pheasant | Game Bird | spring | IA | Webster | 3 | cropland | R3IA-c-21 | 2016-04-11T00:00:00Z | 10 | 14 | 1 | R3IA-c-21 |
| ring-necked pheasant | Game Bird | summer | IA | Webster | 3 | cropland | R3IA-c-21 | 2016-06-08T00:00:00Z | 7 | 10 | 1 | R3IA-c-21 |
| ring-necked pheasant | Game Bird | summer | IA | Webster | 3 | cropland | R3IA-c-21 | 2016-06-01T00:00:00Z | 7 | 10 | 1 | R3IA-c-21 |
| ring-necked pheasant | Game Bird | summer | IA | Webster | 3 | cropland | R3IA-c-21 | 2016-06-01T00:00:00Z | 7 | 10 | 1 | R3IA-c-21 |
| ring-necked pheasant | Game Bird | summer | IA | Webster | 3 | cropland | R3IA-c-21 | 2016-06-01T00:00:00Z | 1 | 2 | 1 | R3IA-c-21 |
| ring-necked pheasant | Game Bird | summer | IA | Webster | 3 | cropland | R3IA-c-21 | 2016-06-01T00:00:00Z | 30 | NA | 1 | R3IA-c-21 |
| ring-necked pheasant | Game Bird | winter | IA | Webster | 3 | cropland | R3IA-c-21 | 2016-01-12T00:00:00Z | 14 | 20 | 1 | R3IA-c-21 |
| ring-necked pheasant | Game Bird | spring | IA | Webster | 3 | cropland | R3IA-c-21 | 2016-04-11T00:00:00Z | 4 | 14 | 1 | R3IA-c-21 |
| ring-necked pheasant | Game Bird | winter | IA | Webster | 3 | cropland | R3IA-c-21 | 2016-02-09T00:00:00Z | 4 | 7 | 1 | R3IA-c-21 |
| ring-necked pheasant | Game Bird | spring | IA | Webster | 3 | cropland | R3IA-c-21 | 2016-04-04T00:00:00Z | 30 | NA | 1 | R3IA-c-21 |
| ring-necked pheasant | Game Bird | spring | IA | Webster | 3 | cropland | R3IA-c-21 | 2016-04-04T00:00:00Z | 30 | NA | 1 | R3IA-c-21 |
| ring-necked pheasant | Game Bird | winter | IA | Webster | 3 | cropland | R3IA-c-21 | 2016-02-09T00:00:00Z | 10 | 14 | 1 | R3IA-c-21 |
| ring-necked pheasant | Game Bird | winter | IA | Webster | 3 | cropland | R3IA-c-21 | 2016-02-09T00:00:00Z | 7 | 10 | 1 | R3IA-c-21 |
| ring-necked pheasant | Game Bird | winter | IA | Webster | 3 | cropland | R3IA-c-21 | 2016-01-25T00:00:00Z | 2 | 30 | 1 | R3IA-c-21 |
| ring-necked pheasant | Game Bird | summer | IA | Webster | 3 | cropland | R3IA-c-21 | 2016-06-08T00:00:00Z | 30 | NA | 1 | R3IA-c-21 |
| ring-necked pheasant | Game Bird | spring | IA | Webster | 3 | cropland | R3IA-c-21 | 2016-04-11T00:00:00Z | 1 | 2 | 1 | R3IA-c-21 |
| ring-necked pheasant | Game Bird | winter | IA | Madison | 3 | cropland | R3IA-c-22 | 2015-12-01T00:00:00Z | 3 | 4 | 1 | R3IA-c-22 |
| ring-necked pheasant | Game Bird | winter | IA | Madison | 3 | cropland | R3IA-c-22 | 2015-12-01T00:00:00Z | 10 | 13 | 1 | R3IA-c-22 |
| ring-necked pheasant | Game Bird | winter | IA | Madison | 3 | cropland | R3IA-c-22 | 2015-12-14T00:00:00Z | 10 | 15 | 1 | R3IA-c-22 |
| ring-necked pheasant | Game Bird | winter | IA | Madison | 3 | cropland | R3IA-c-22 | 2015-12-14T00:00:00Z | 4 | 7 | 1 | R3IA-c-22 |
| ring-necked pheasant | Game Bird | winter | IA | Madison | 3 | cropland | R3IA-c-22 | 2015-12-29T00:00:00Z | 14 | 20 | 1 | R3IA-c-22 |
| ring-necked pheasant | Game Bird | winter | IA | Madison | 3 | cropland | R3IA-c-22 | 2016-01-05T00:00:00Z | 3 | 7 | 1 | R3IA-c-22 |
| ring-necked pheasant | Game Bird | winter | IA | Madison | 3 | cropland | R3IA-c-22 | 2016-01-12T00:00:00Z | 14 | 20 | 1 | R3IA-c-22 |
| ring-necked pheasant | Game Bird | winter | IA | Madison | 3 | cropland | R3IA-c-22 | 2016-01-12T00:00:00Z | 14 | 20 | 1 | R3IA-c-22 |
| red-tailed hawk | Raptor | winter | IA | Madison | 3 | cropland | R3IA-c-22 | 2015-12-29T00:00:00Z | 30 | 44 | 1 | R3IA-c-0 |
| red-tailed hawk | Raptor | winter | IA | Madison | 3 | cropland | R3IA-c-22 | 2015-12-29T00:00:00Z | 14 | 20 | 1 | R3IA-c-0 |
| ring-necked pheasant | Game Bird | winter | IA | Madison | 3 | cropland | R3IA-c-22 | 2016-01-25T00:00:00Z | 4 | 7 | 1 | R3IA-c-22 |
| ring-necked pheasant | Game Bird | winter | IA | Madison | 3 | cropland | R3IA-c-22 | 2016-01-25T00:00:00Z | 30 | NA | 1 | R3IA-c-22 |
| red-tailed hawk | Raptor | winter | IA | Madison | 3 | cropland | R3IA-c-22 | 2015-12-29T00:00:00Z | 30 | 44 | 1 | R3IA-c-0 |
| red-tailed hawk | Raptor | winter | IA | Madison | 3 | cropland | R3IA-c-22 | 2015-12-29T00:00:00Z | 14 | 20 | 1 | R3IA-c-0 |
| ring-necked pheasant | Game Bird | winter | IA | Madison | 3 | cropland | R3IA-c-22 | 2016-02-08T00:00:00Z | 21 | 30 | 1 | R3IA-c-22 |
| red-tailed hawk | Raptor | winter | IA | Madison | 3 | cropland | R3IA-c-22 | 2016-02-08T00:00:00Z | 30 | 38 | 1 | R3IA-c-0 |
| turkey vulture | Raptor | summer | IA | Madison | 3 | cropland | R3IA-c-22 | 2016-06-03T00:00:00Z | 30 | NA | 1 | R3IA-c-0 |
| ring-necked pheasant | Game Bird | winter | IA | Adair | 3 | cropland | R3IA-c-23 | 2015-01-09T00:00:00Z | 10 | 14 | 1 | R3IA-c-23 |
| ring-necked pheasant | Game Bird | winter | IA | Adair | 3 | cropland | R3IA-c-23 | 2015-01-09T00:00:00Z | 30 | NA | 1 | R3IA-c-23 |
| ring-necked pheasant | Game Bird | winter | IA | Adair | 3 | cropland | R3IA-c-23 | 2015-02-10T00:00:00Z | 20 | 30 | 1 | R3IA-c-23 |
| ring-necked pheasant | Game Bird | winter | IA | Adair | 3 | cropland | R3IA-c-23 | 2015-02-10T00:00:00Z | 3 | 6 | 1 | R3IA-c-23 |
| ring-necked pheasant | Game Bird | winter | IA | Adair | 3 | cropland | R3IA-c-23 | 2015-02-10T00:00:00Z | 3 | 6 | 1 | R3IA-c-23 |
| ring-necked pheasant | Game Bird | spring | IA | Adair | 3 | cropland | R3IA-c-23 | 2015-03-16T00:00:00Z | 1 | 1 | 1 | R3IA-c-23 |
| ring-necked pheasant | Game Bird | spring | IA | Adair | 3 | cropland | R3IA-c-23 | 2015-03-16T00:00:00Z | 7 | 10 | 1 | R3IA-c-23 |
| ring-necked pheasant | Game Bird | spring | IA | Adair | 3 | cropland | R3IA-c-23 | 2015-03-30T00:00:00Z | 4 | 7 | 1 | R3IA-c-23 |
| ring-necked pheasant | Game Bird | spring | IA | Adair | 3 | cropland | R3IA-c-23 | 2015-03-30T00:00:00Z | 2 | 3 | 1 | R3IA-c-23 |
| ring-necked pheasant | Game Bird | spring | IA | Adair | 3 | cropland | R3IA-c-23 | 2015-03-30T00:00:00Z | 10 | 14 | 1 | R3IA-c-23 |
| ring-necked pheasant | Game Bird | summer | IA | Adair | 3 | cropland | R3IA-c-23 | 2015-05-18T00:00:00Z | 10 | 14 | 1 | R3IA-c-23 |
| ring-necked pheasant | Game Bird | summer | IA | Adair | 3 | cropland | R3IA-c-23 | 2015-05-18T00:00:00Z | 10 | 14 | 1 | R3IA-c-23 |
| ring-necked pheasant | Game Bird | summer | IA | Adair | 3 | cropland | R3IA-c-23 | 2015-06-15T00:00:00Z | 3 | 4 | 1 | R3IA-c-23 |
| ring-necked pheasant | Game Bird | fall | IA | Adair | 3 | cropland | R3IA-c-23 | 2015-07-28T00:00:00Z | 30 | NA | 1 | R3IA-c-23 |
| ring-necked pheasant | Game Bird | fall | IA | Adair | 3 | cropland | R3IA-c-23 | 2015-08-17T00:00:00Z | 2 | 3 | 1 | R3IA-c-23 |
| ring-necked pheasant | Game Bird | fall | IA | Adair | 3 | cropland | R3IA-c-23 | 2015-09-14T00:00:00Z | 1 | 1 | 1 | R3IA-c-23 |
| ring-necked pheasant | Game Bird | fall | IA | Adair | 3 | cropland | R3IA-c-23 | 2015-09-14T00:00:00Z | 1 | 1 | 1 | R3IA-c-23 |
| ring-necked pheasant | Game Bird | fall | IA | Adair | 3 | cropland | R3IA-c-23 | 2015-09-28T00:00:00Z | 1 | 1 | 1 | R3IA-c-23 |
| ring-necked pheasant | Game Bird | winter | IA | Adair | 3 | cropland | R3IA-c-23 | 2015-12-01T00:00:00Z | 10 | 15 | 1 | R3IA-c-23 |
| ring-necked pheasant | Game Bird | winter | IA | Adair | 3 | cropland | R3IA-c-23 | 2015-12-01T00:00:00Z | 10 | 15 | 1 | R3IA-c-23 |
| ring-necked pheasant | Game Bird | winter | IA | Adair | 3 | cropland | R3IA-c-23 | 2015-12-14T00:00:00Z | 1 | 2 | 1 | R3IA-c-23 |
| ring-necked pheasant | Game Bird | winter | IA | Adair | 3 | cropland | R3IA-c-23 | 2015-12-29T00:00:00Z | 14 | 20 | 1 | R3IA-c-23 |
| ring-necked pheasant | Game Bird | winter | IA | Adair | 3 | cropland | R3IA-c-23 | 2015-12-29T00:00:00Z | 2 | 3 | 1 | R3IA-c-23 |
| red-tailed hawk | Raptor | winter | IA | Adair | 3 | cropland | R3IA-c-23 | 2015-12-29T00:00:00Z | 37 | NA | 1 | R3IA-c-0 |
| red-tailed hawk | Raptor | winter | IA | Adair | 3 | cropland | R3IA-c-23 | 2015-12-29T00:00:00Z | 7 | 10 | 1 | R3IA-c-0 |
| ring-necked pheasant | Game Bird | winter | IA | Adair | 3 | cropland | R3IA-c-23 | 2016-01-11T00:00:00Z | 2 | 3 | 1 | R3IA-c-23 |
| ring-necked pheasant | Game Bird | winter | IA | Adair | 3 | cropland | R3IA-c-23 | 2016-01-11T00:00:00Z | 3 | 4 | 1 | R3IA-c-23 |
| ring-necked pheasant | Game Bird | winter | IA | Adair | 3 | cropland | R3IA-c-23 | 2016-01-25T00:00:00Z | 4 | 7 | 1 | R3IA-c-23 |
| ring-necked pheasant | Game Bird | winter | IA | Adair | 3 | cropland | R3IA-c-23 | 2016-02-09T00:00:00Z | 14 | 21 | 1 | R3IA-c-23 |
| ring-necked pheasant | Game Bird | winter | IA | Adair | 3 | cropland | R3IA-c-23 | 2016-02-09T00:00:00Z | 21 | 30 | 1 | R3IA-c-23 |
| ring-necked pheasant | Game Bird | winter | IA | Pocahontas | 3 | cropland | R3IA-c-24 | 2015-12-01T00:00:00Z | 7 | 10 | 1 | R3IA-c-24 |
| ring-necked pheasant | Game Bird | winter | IA | Pocahontas | 3 | cropland | R3IA-c-24 | 2015-12-01T00:00:00Z | 10 | 14 | 1 | R3IA-c-24 |
| ring-necked pheasant | Game Bird | winter | IA | Pocahontas | 3 | cropland | R3IA-c-24 | 2015-12-14T00:00:00Z | 10 | 15 | 1 | R3IA-c-24 |
| ring-necked pheasant | Game Bird | winter | IA | Pocahontas | 3 | cropland | R3IA-c-24 | 2015-12-29T00:00:00Z | 30 | NA | 1 | R3IA-c-24 |
| ring-necked pheasant | Game Bird | winter | IA | Pocahontas | 3 | cropland | R3IA-c-24 | 2016-01-11T00:00:00Z | 31 | NA | 1 | R3IA-c-24 |
| ring-necked pheasant | Game Bird | winter | IA | Pocahontas | 3 | cropland | R3IA-c-24 | 2016-01-11T00:00:00Z | 15 | 21 | 1 | R3IA-c-24 |
| ring-necked pheasant | Game Bird | winter | IA | Pocahontas | 3 | cropland | R3IA-c-24 | 2016-01-26T00:00:00Z | 30 | NA | 1 | R3IA-c-24 |
| ring-necked pheasant | Game Bird | winter | IA | Pocahontas | 3 | cropland | R3IA-c-24 | 2016-02-09T00:00:00Z | 14 | 20 | 1 | R3IA-c-24 |
| ring-necked pheasant | Game Bird | spring | IA | Pocahontas | 3 | cropland | R3IA-c-24 | 2016-03-21T00:00:00Z | 14 | 30 | 1 | R3IA-c-24 |
| ring-necked pheasant | Game Bird | spring | IA | Pocahontas | 3 | cropland | R3IA-c-24 | 2016-03-28T00:00:00Z | 30 | NA | 1 | R3IA-c-24 |
| ring-necked pheasant | Game Bird | spring | IA | Pocahontas | 3 | cropland | R3IA-c-24 | 2016-04-04T00:00:00Z | 20 | 30 | 1 | R3IA-c-24 |
| red-tailed hawk | Raptor | spring | IA | Pocahontas | 3 | cropland | R3IA-c-24 | 2016-04-11T00:00:00Z | 20 | 30 | 1 | R3IA-c-0 |
| red-tailed hawk | Raptor | spring | IA | Pocahontas | 3 | cropland | R3IA-c-24 | 2016-04-11T00:00:00Z | 3 | 4 | 1 | R3IA-c-0 |
| red-tailed hawk | Raptor | spring | IA | Pocahontas | 3 | cropland | R3IA-c-24 | 2016-04-11T00:00:00Z | 30 | NA | 1 | R3IA-c-0 |
| red-tailed hawk | Raptor | spring | IA | Pocahontas | 3 | cropland | R3IA-c-24 | 2016-04-13T00:00:00Z | 2 | 3 | 1 | R3IA-c-0 |
| red-tailed hawk | Raptor | spring | IA | Pocahontas | 3 | cropland | R3IA-c-24 | 2016-04-13T00:00:00Z | 4 | 7 | 1 | R3IA-c-0 |
| ring-necked pheasant | Game Bird | summer | IA | Pocahontas | 3 | cropland | R3IA-c-24 | 2016-05-16T00:00:00Z | 4 | 8 | 1 | R3IA-c-24 |
| ring-necked pheasant | Game Bird | summer | IA | Pocahontas | 3 | cropland | R3IA-c-24 | 2016-05-24T00:00:00Z | 20 | 30 | 1 | R3IA-c-24 |
| ring-necked pheasant | Game Bird | summer | IA | Pocahontas | 3 | cropland | R3IA-c-24 | 2016-05-30T00:00:00Z | 21 | 31 | 1 | R3IA-c-24 |
| ring-necked pheasant | Game Bird | summer | IA | Pocahontas | 3 | cropland | R3IA-c-24 | 2016-07-13T00:00:00Z | 10 | 14 | 1 | R3IA-c-24 |
| red-tailed hawk | Raptor | fall | IA | Pocahontas | 3 | cropland | R3IA-c-24 | 2016-08-15T00:00:00Z | 14 | NA | 1 | R3IA-c-0 |
| red-tailed hawk | Raptor | fall | IA | Pocahontas | 3 | cropland | R3IA-c-24 | 2016-08-29T00:00:00Z | 47 | 54 | 1 | R3IA-c-0 |
| red-tailed hawk | Raptor | fall | IA | Pocahontas | 3 | cropland | R3IA-c-24 | 2016-09-12T00:00:00Z | 61 | NA | 1 | R3IA-c-0 |
| red-tailed hawk | Raptor | fall | IA | Pocahontas | 3 | cropland | R3IA-c-24 | 2016-09-12T00:00:00Z | 61 | NA | 1 | R3IA-c-0 |
| ring-necked pheasant | Game Bird | fall | IA | Pocahontas | 3 | cropland | R3IA-c-24 | 2016-10-10T00:00:00Z | 7 | 10 | 1 | R3IA-c-24 |
| ring-necked pheasant | Game Bird | fall | IA | Pocahontas | 3 | cropland | R3IA-c-24 | 2016-10-10T00:00:00Z | 1 | 2 | 1 | R3IA-c-24 |
| ring-necked pheasant | Game Bird | winter | IA | Pocahontas | 3 | cropland | R3IA-c-24 | 2016-12-05T00:00:00Z | 22 | NA | 1 | R3IA-c-24 |
| ring-necked pheasant | Game Bird | winter | IA | Pocahontas | 3 | cropland | R3IA-c-24 | 2016-12-05T00:00:00Z | 22 | NA | 1 | R3IA-c-24 |
| ring-necked pheasant | Game Bird | winter | IA | Pocahontas | 3 | cropland | R3IA-c-24 | 2017-01-09T00:00:00Z | 14 | NA | 1 | R3IA-c-24 |
| ring-necked pheasant | Game Bird | winter | IA | Pocahontas | 3 | cropland | R3IA-c-24 | 2017-01-09T00:00:00Z | 10 | 10 | 1 | R3IA-c-24 |
| ring-necked pheasant | Game Bird | winter | IA | Pocahontas | 3 | cropland | R3IA-c-24 | 2017-01-23T00:00:00Z | 7 | 10 | 1 | R3IA-c-24 |
| ring-necked pheasant | Game Bird | winter | IA | Pocahontas | 3 | cropland | R3IA-c-24 | 2017-01-23T00:00:00Z | 21 | NA | 1 | R3IA-c-24 |
| ring-necked pheasant | Game Bird | winter | IA | Pocahontas | 3 | cropland | R3IA-c-24 | 2017-01-30T00:00:00Z | 3 | 4 | 1 | R3IA-c-24 |
| ring-necked pheasant | Game Bird | winter | IA | Pocahontas | 3 | cropland | R3IA-c-24 | 2017-01-30T00:00:00Z | 7 | 10 | 1 | R3IA-c-24 |
| ring-necked pheasant | Game Bird | winter | IA | Pocahontas | 3 | cropland | R3IA-c-24 | 2017-02-06T00:00:00Z | 10 | 15 | 1 | R3IA-c-24 |
| ring-necked pheasant | Game Bird | winter | IA | Pocahontas | 3 | cropland | R3IA-c-24 | 2017-02-06T00:00:00Z | 10 | 15 | 1 | R3IA-c-24 |
| ring-necked pheasant | Game Bird | winter | IA | Adair/Adams/Cass | 3 | cropland | R3IA-c-25 | 2015-12-01T00:00:00Z | 7 | 10 | 1 | R3IA-c-25 |
| ring-necked pheasant | Game Bird | winter | IA | Adair/Adams/Cass | 3 | cropland | R3IA-c-25 | 2015-12-14T00:00:00Z | 30 | NA | 1 | R3IA-c-25 |
| ring-necked pheasant | Game Bird | winter | IA | Adair/Adams/Cass | 3 | cropland | R3IA-c-25 | 2015-12-14T00:00:00Z | 10 | 14 | 1 | R3IA-c-25 |
| ring-necked pheasant | Game Bird | winter | IA | Adair/Adams/Cass | 3 | cropland | R3IA-c-25 | 2015-12-28T00:00:00Z | 21 | 31 | 1 | R3IA-c-25 |
| ring-necked pheasant | Game Bird | winter | IA | Adair/Adams/Cass | 3 | cropland | R3IA-c-25 | 2015-12-28T00:00:00Z | 15 | 21 | 1 | R3IA-c-25 |
| red-tailed hawk | Raptor | winter | IA | Adair/Adams/Cass | 3 | cropland | R3IA-c-25 | 2015-12-28T00:00:00Z | 37 | 44 | 1 | R3IA-c-0 |
| red-tailed hawk | Raptor | winter | IA | Adair/Adams/Cass | 3 | cropland | R3IA-c-25 | 2015-12-28T00:00:00Z | 15 | 21 | 1 | R3IA-c-0 |
| ring-necked pheasant | Game Bird | winter | IA | Adair/Adams/Cass | 3 | cropland | R3IA-c-25 | 2016-01-11T00:00:00Z | 4 | 7 | 1 | R3IA-c-25 |
| ring-necked pheasant | Game Bird | winter | IA | Adair/Adams/Cass | 3 | cropland | R3IA-c-25 | 2016-01-25T00:00:00Z | 4 | 7 | 1 | R3IA-c-25 |
| ring-necked pheasant | Game Bird | winter | IA | Adair/Adams/Cass | 3 | cropland | R3IA-c-25 | 2016-01-25T00:00:00Z | 30 | NA | 1 | R3IA-c-25 |
| ring-necked pheasant | Game Bird | winter | IA | Adair/Adams/Cass | 3 | cropland | R3IA-c-25 | 2016-02-09T00:00:00Z | 4 | 7 | 1 | R3IA-c-25 |
| ring-necked pheasant | Game Bird | winter | IA | Adair/Adams/Cass | 3 | cropland | R3IA-c-25 | 2016-02-09T00:00:00Z | 7 | 10 | 1 | R3IA-c-25 |
| turkey vulture | Raptor | summer | IA | Adair/Adams/Cass | 3 | cropland | R3IA-c-25 | 2016-05-22T00:00:00Z | 30 | NA | 1 | R3IA-c-0 |
| ring-necked pheasant | Game Bird | fall | IA | Adair/Adams/Cass | 3 | cropland | R3IA-c-25 | 2016-10-29T00:00:00Z | 3 | 4 | 1 | R3IA-c-25 |
| ring-necked pheasant | Game Bird | winter | IA | Carroll/Crawford | 3 | cropland | R3IA-c-26 | 2015-01-12T00:00:00Z | 4 | 7 | 1 | R3IA-c-26 |
| ring-necked pheasant | Game Bird | winter | IA | Carroll/Crawford | 3 | cropland | R3IA-c-26 | 2015-01-12T00:00:00Z | 10 | 14 | 1 | R3IA-c-26 |
| ring-necked pheasant | Game Bird | winter | IA | Carroll/Crawford | 3 | cropland | R3IA-c-26 | 2015-01-12T00:00:00Z | 7 | 10 | 1 | R3IA-c-26 |
| ring-necked pheasant | Game Bird | winter | IA | Carroll/Crawford | 3 | cropland | R3IA-c-26 | 2015-02-09T00:00:00Z | 30 | NA | 1 | R3IA-c-26 |
| ring-necked pheasant | Game Bird | winter | IA | Carroll/Crawford | 3 | cropland | R3IA-c-26 | 2015-02-09T00:00:00Z | 4 | 7 | 1 | R3IA-c-26 |
| ring-necked pheasant | Game Bird | spring | IA | Carroll/Crawford | 3 | cropland | R3IA-c-26 | 2015-03-17T00:00:00Z | 29 | NA | 1 | R3IA-c-26 |
| ring-necked pheasant | Game Bird | spring | IA | Carroll/Crawford | 3 | cropland | R3IA-c-26 | 2015-03-17T00:00:00Z | 14 | 20 | 1 | R3IA-c-26 |
| ring-necked pheasant | Game Bird | spring | IA | Carroll/Crawford | 3 | cropland | R3IA-c-26 | 2015-03-17T00:00:00Z | 29 | NA | 1 | R3IA-c-26 |
| ring-necked pheasant | Game Bird | spring | IA | Carroll/Crawford | 3 | cropland | R3IA-c-26 | 2015-04-13T00:00:00Z | 30 | NA | 1 | R3IA-c-26 |
| ring-necked pheasant | Game Bird | spring | IA | Carroll/Crawford | 3 | cropland | R3IA-c-26 | 2015-04-13T00:00:00Z | 3 | 4 | 1 | R3IA-c-26 |
| ring-necked pheasant | Game Bird | summer | IA | Carroll/Crawford | 3 | cropland | R3IA-c-26 | 2015-05-18T00:00:00Z | 1 | 1 | 1 | R3IA-c-26 |
| ring-necked pheasant | Game Bird | summer | IA | Carroll/Crawford | 3 | cropland | R3IA-c-26 | 2015-05-18T00:00:00Z | 14 | 21 | 1 | R3IA-c-26 |
| ring-necked pheasant | Game Bird | summer | IA | Carroll/Crawford | 3 | cropland | R3IA-c-26 | 2015-06-15T00:00:00Z | 10 | 14 | 1 | R3IA-c-26 |
| ring-necked pheasant | Game Bird | fall | IA | Carroll/Crawford | 3 | cropland | R3IA-c-26 | 2015-07-27T00:00:00Z | 30 | NA | 1 | R3IA-c-26 |
| ring-necked pheasant | Game Bird | fall | IA | Carroll/Crawford | 3 | cropland | R3IA-c-26 | 2015-08-17T00:00:00Z | 14 | 22 | 1 | R3IA-c-26 |
| ring-necked pheasant | Game Bird | fall | IA | Carroll/Crawford | 3 | cropland | R3IA-c-26 | 2015-08-17T00:00:00Z | 2 | 3 | 1 | R3IA-c-26 |
| ring-necked pheasant | Game Bird | fall | IA | Carroll/Crawford | 3 | cropland | R3IA-c-26 | 2015-09-14T00:00:00Z | 10 | 14 | 1 | R3IA-c-26 |
| ring-necked pheasant | Game Bird | fall | IA | Carroll/Crawford | 3 | cropland | R3IA-c-26 | 2015-09-14T00:00:00Z | 10 | 14 | 1 | R3IA-c-26 |
| ring-necked pheasant | Game Bird | fall | IA | Carroll/Crawford | 3 | cropland | R3IA-c-26 | 2015-09-28T00:00:00Z | 4 | 7 | 1 | R3IA-c-26 |
| ring-necked pheasant | Game Bird | fall | IA | Carroll/Crawford | 3 | cropland | R3IA-c-26 | 2015-10-12T00:00:00Z | 2 | 3 | 1 | R3IA-c-26 |
| ring-necked pheasant | Game Bird | fall | IA | Carroll/Crawford | 3 | cropland | R3IA-c-26 | 2015-10-12T00:00:00Z | 2 | 3 | 1 | R3IA-c-26 |
| ring-necked pheasant | Game Bird | fall | IA | Carroll/Crawford | 3 | cropland | R3IA-c-26 | 2015-10-12T00:00:00Z | 1 | 2 | 1 | R3IA-c-26 |
| ring-necked pheasant | Game Bird | winter | IA | Carroll/Crawford | 3 | cropland | R3IA-c-26 | 2015-12-01T00:00:00Z | 4 | 7 | 1 | R3IA-c-26 |
| ring-necked pheasant | Game Bird | winter | IA | Carroll/Crawford | 3 | cropland | R3IA-c-26 | 2015-12-01T00:00:00Z | 10 | 13 | 1 | R3IA-c-26 |
| ring-necked pheasant | Game Bird | winter | IA | Carroll/Crawford | 3 | cropland | R3IA-c-26 | 2015-12-14T00:00:00Z | 4 | 7 | 1 | R3IA-c-26 |
| ring-necked pheasant | Game Bird | winter | IA | Carroll/Crawford | 3 | cropland | R3IA-c-26 | 2015-12-14T00:00:00Z | 1 | 1 | 1 | R3IA-c-26 |
| red-tailed hawk | Raptor | winter | IA | Carroll/Crawford | 3 | cropland | R3IA-c-26 | 2015-12-29T00:00:00Z | 2 | 7 | 1 | R3IA-c-0 |
| red-tailed hawk | Raptor | winter | IA | Carroll/Crawford | 3 | cropland | R3IA-c-26 | 2015-12-29T00:00:00Z | 7 | 20 | 1 | R3IA-c-0 |
| ring-necked pheasant | Game Bird | winter | IA | Carroll/Crawford | 3 | cropland | R3IA-c-26 | 2015-12-29T00:00:00Z | 7 | NA | 1 | R3IA-c-26 |
| ring-necked pheasant | Game Bird | winter | IA | Carroll/Crawford | 3 | cropland | R3IA-c-26 | 2016-01-11T00:00:00Z | 20 | 30 | 1 | R3IA-c-26 |
| ring-necked pheasant | Game Bird | winter | IA | Carroll/Crawford | 3 | cropland | R3IA-c-26 | 2016-01-11T00:00:00Z | 7 | 10 | 1 | R3IA-c-26 |
| ring-necked pheasant | Game Bird | winter | IA | Carroll/Crawford | 3 | cropland | R3IA-c-26 | 2016-01-26T00:00:00Z | 30 | NA | 1 | R3IA-c-26 |
| ring-necked pheasant | Game Bird | winter | IA | Carroll/Crawford | 3 | cropland | R3IA-c-26 | 2016-01-26T00:00:00Z | 20 | 30 | 1 | R3IA-c-26 |
| ring-necked pheasant | Game Bird | winter | IA | Carroll/Crawford | 3 | cropland | R3IA-c-26 | 2016-02-09T00:00:00Z | 10 | 14 | 1 | R3IA-c-26 |
| short-eared owl | Raptor | winter | IA | Marshall/Tama | 3 | cropland | R3IA-c-27 | 2015-11-16T00:00:00Z | 1 | 2 | 1 | R3IA-c-0 |
| ring-necked pheasant | Game Bird | winter | IA | Marshall/Tama | 3 | cropland | R3IA-c-27 | 2015-12-01T00:00:00Z | 20 | 30 | 1 | R3IA-c-27 |
| ring-necked pheasant | Game Bird | winter | IA | Marshall/Tama | 3 | cropland | R3IA-c-27 | 2015-12-01T00:00:00Z | 20 | 30 | 1 | R3IA-c-27 |
| short-eared owl | Raptor | winter | IA | Marshall/Tama | 3 | cropland | R3IA-c-27 | 2016-01-06T00:00:00Z | 6 | 9 | 1 | R3IA-c-0 |
| ring-necked pheasant | Game Bird | winter | IA | Marshall/Tama | 3 | cropland | R3IA-c-27 | 2016-01-25T00:00:00Z | 20 | 20 | 1 | R3IA-c-27 |
| turkey vulture | Raptor | spring | IA | Marshall/Tama | 3 | cropland | R3IA-c-27 | 2016-03-25T00:00:00Z | 30 | NA | 1 | R3IA-c-0 |
| ring-necked pheasant | Game Bird | winter | IA | Marshall/Tama | 3 | cropland | R3IA-c-27 | 2016-02-12T00:00:00Z | 14 | 19 | 1 | R3IA-c-27 |
| ring-necked pheasant | Game Bird | winter | IA | Marshall/Tama | 3 | cropland | R3IA-c-27 | 2016-02-12T00:00:00Z | 14 | 19 | 1 | R3IA-c-27 |
| turkey vulture | Raptor | spring | IA | Marshall/Tama | 3 | cropland | R3IA-c-27 | 2016-03-31T00:00:00Z | 30 | NA | 1 | R3IA-c-0 |
| red-tailed hawk | Raptor | spring | IA | Marshall/Tama | 3 | cropland | R3IA-c-27 | 2016-04-12T00:00:00Z | 14 | 20 | 1 | R3IA-c-0 |
| red-tailed hawk | Raptor | spring | IA | Marshall/Tama | 3 | cropland | R3IA-c-27 | 2016-04-12T00:00:00Z | 10 | 14 | 1 | R3IA-c-0 |
| ring-necked pheasant | Game Bird | summer | IA | Marshall/Tama | 3 | cropland | R3IA-c-27 | 2016-05-24T00:00:00Z | 4 | 7 | 1 | R3IA-c-27 |
| ring-necked pheasant | Game Bird | summer | IA | Marshall/Tama | 3 | cropland | R3IA-c-27 | 2016-06-09T00:00:00Z | 10 | 14 | 1 | R3IA-c-27 |
| red-tailed hawk | Raptor | fall | IA | Marshall/Tama | 3 | cropland | R3IA-c-27 | 2016-08-01T00:00:00Z | 14 | 20 | 1 | R3IA-c-0 |
| ring-necked pheasant | Game Bird | winter | IA | Marshall/Tama | 3 | cropland | R3IA-c-27 | 2015-12-01T00:00:00Z | 2 | 3 | 1 | R3IA-c-27 |
| ring-necked pheasant | Game Bird | winter | IA | Marshall/Tama | 3 | cropland | R3IA-c-27 | 2015-12-29T00:00:00Z | 1 | 2 | 1 | R3IA-c-27 |
| ring-necked pheasant | Game Bird | winter | IA | Marshall/Tama | 3 | cropland | R3IA-c-27 | 2016-01-25T00:00:00Z | 20 | 20 | 1 | R3IA-c-27 |
| ring-necked pheasant | Game Bird | winter | IA | Marshall/Tama | 3 | cropland | R3IA-c-27 | 2016-02-09T00:00:00Z | 7 | 10 | 1 | R3IA-c-27 |
| ring-necked pheasant | Game Bird | spring | IA | Marshall/Tama | 3 | cropland | R3IA-c-27 | 2016-04-08T00:00:00Z | 20 | 30 | 1 | R3IA-c-27 |
| ring-necked pheasant | Game Bird | spring | IA | Marshall/Tama | 3 | cropland | R3IA-c-27 | 2016-04-08T00:00:00Z | 7 | 10 | 1 | R3IA-c-27 |
| ring-necked pheasant | Game Bird | spring | IA | Marshall/Tama | 3 | cropland | R3IA-c-27 | 2016-04-08T00:00:00Z | 14 | 20 | 1 | R3IA-c-27 |
| red-tailed hawk | Raptor | spring | IA | Marshall/Tama | 3 | cropland | R3IA-c-27 | 2016-04-12T00:00:00Z | 20 | 30 | 1 | R3IA-c-0 |
| ring-necked pheasant | Game Bird | fall | IA | Marshall/Tama | 3 | cropland | R3IA-c-27 | 2016-09-26T00:00:00Z | 8 | 10 | 1 | R3IA-c-27 |
| red-tailed hawk | Raptor | fall | IA | Marshall/Tama | 3 | cropland | R3IA-c-27 | 2016-08-29T00:00:00Z | 40 | 50 | 1 | R3IA-c-0 |
| ring-necked pheasant | Game Bird | winter | IA | Marshall/Tama | 3 | cropland | R3IA-c-27 | 2016-12-05T00:00:00Z | 10 | 24 | 1 | R3IA-c-27 |
| ring-necked pheasant | Game Bird | winter | IA | Marshall/Tama | 3 | cropland | R3IA-c-27 | 2016-12-05T00:00:00Z | 24 | 34 | 1 | R3IA-c-27 |
| ring-necked pheasant | Game Bird | winter | IA | Marshall/Tama | 3 | cropland | R3IA-c-27 | 2017-01-30T00:00:00Z | 1 | 2 | 1 | R3IA-c-27 |
| ring-necked pheasant | Game Bird | winter | IA | Marshall/Tama | 3 | cropland | R3IA-c-27 | 2017-01-30T00:00:00Z | 7 | 10 | 1 | R3IA-c-27 |
| ring-necked pheasant | Game Bird | winter | IA | Marshall/Tama | 3 | cropland | R3IA-c-27 | 2017-01-30T00:00:00Z | 2 | 3 | 1 | R3IA-c-27 |
| ring-necked pheasant | Game Bird | winter | IA | Marshall/Tama | 3 | cropland | R3IA-c-27 | 2017-02-06T00:00:00Z | 3 | 4 | 1 | R3IA-c-27 |
| ring-necked pheasant | Game Bird | winter | IA | Marshall/Tama | 3 | cropland | R3IA-c-27 | 2017-02-06T00:00:00Z | 4 | 7 | 1 | R3IA-c-27 |
| ring-necked pheasant | Game Bird | winter | IA | Marshall/Tama | 3 | cropland | R3IA-c-27 | 2017-01-23T00:00:00Z | 21 | 31 | 1 | R3IA-c-27 |
| ring-necked pheasant | Game Bird | winter | IA | Marshall/Tama | 3 | cropland | R3IA-c-27 | 2017-01-23T00:00:00Z | 7 | 10 | 1 | R3IA-c-27 |
| ring-necked pheasant | Game Bird | winter | IA | Marshall/Tama | 3 | cropland | R3IA-c-27 | 2017-01-23T00:00:00Z | 14 | 21 | 1 | R3IA-c-27 |
| ring-necked pheasant | Game Bird | spring | IA | Pottawattamie | 3 | cropland | R3IA-c-29 | 2015-04-14T00:00:00Z | 3 | 5 | 1 | R3IA-c-29 |
| ring-necked pheasant | Game Bird | spring | IA | Pottawattamie | 3 | cropland | R3IA-c-29 | 2015-04-14T00:00:00Z | 9 | 14 | 1 | R3IA-c-29 |
| ring-necked pheasant | Game Bird | spring | IA | Pottawattamie | 3 | cropland | R3IA-c-29 | 2015-04-14T00:00:00Z | 3 | 5 | 1 | R3IA-c-29 |
| ring-necked pheasant | Game Bird | spring | IA | Pottawattamie | 3 | cropland | R3IA-c-29 | 2015-04-14T00:00:00Z | 2 | 3 | 1 | R3IA-c-29 |
| ring-necked pheasant | Game Bird | winter | IA | Pottawattamie | 3 | cropland | R3IA-c-29 | 2015-01-12T00:00:00Z | 10 | 14 | 1 | R3IA-c-29 |
| ring-necked pheasant | Game Bird | winter | IA | Pottawattamie | 3 | cropland | R3IA-c-29 | 2015-01-12T00:00:00Z | 2 | 2 | 1 | R3IA-c-29 |
| ring-necked pheasant | Game Bird | winter | IA | Pottawattamie | 3 | cropland | R3IA-c-29 | 2015-02-11T00:00:00Z | 3 | 5 | 1 | R3IA-c-29 |
| ring-necked pheasant | Game Bird | winter | IA | Pottawattamie | 3 | cropland | R3IA-c-29 | 2015-02-11T00:00:00Z | 30 | NA | 1 | R3IA-c-29 |
| ring-necked pheasant | Game Bird | winter | IA | Pottawattamie | 3 | cropland | R3IA-c-29 | 2015-02-11T00:00:00Z | 7 | 10 | 1 | R3IA-c-29 |
| ring-necked pheasant | Game Bird | spring | IA | Pottawattamie | 3 | cropland | R3IA-c-29 | 2015-03-16T00:00:00Z | 1 | 2 | 1 | R3IA-c-29 |
| ring-necked pheasant | Game Bird | spring | IA | Pottawattamie | 3 | cropland | R3IA-c-29 | 2015-03-16T00:00:00Z | 1 | 2 | 1 | R3IA-c-29 |
| ring-necked pheasant | Game Bird | spring | IA | Pottawattamie | 3 | cropland | R3IA-c-29 | 2015-03-16T00:00:00Z | 14 | 24 | 1 | R3IA-c-29 |
| ring-necked pheasant | Game Bird | spring | IA | Pottawattamie | 3 | cropland | R3IA-c-29 | 2015-04-14T00:00:00Z | 9 | 14 | 1 | R3IA-c-29 |
| ring-necked pheasant | Game Bird | summer | IA | Pottawattamie | 3 | cropland | R3IA-c-29 | 2015-05-18T00:00:00Z | 4 | 7 | 1 | R3IA-c-29 |
| ring-necked pheasant | Game Bird | summer | IA | Pottawattamie | 3 | cropland | R3IA-c-29 | 2015-06-15T00:00:00Z | 10 | 14 | 1 | R3IA-c-29 |
| ring-necked pheasant | Game Bird | summer | IA | Pottawattamie | 3 | cropland | R3IA-c-29 | 2015-06-15T00:00:00Z | 29 | NA | 1 | R3IA-c-29 |
| ring-necked pheasant | Game Bird | fall | IA | Pottawattamie | 3 | cropland | R3IA-c-29 | 2015-07-28T00:00:00Z | 3 | 4 | 1 | R3IA-c-29 |
| ring-necked pheasant | Game Bird | fall | IA | Pottawattamie | 3 | cropland | R3IA-c-29 | 2015-09-28T00:00:00Z | 1 | 1 | 1 | R3IA-c-29 |
| ring-necked pheasant | Game Bird | winter | IA | Pottawattamie | 3 | cropland | R3IA-c-29 | 2015-12-01T00:00:00Z | 30 | NA | 1 | R3IA-c-29 |
| ring-necked pheasant | Game Bird | winter | IA | Pottawattamie | 3 | cropland | R3IA-c-29 | 2015-12-14T00:00:00Z | 14 | 21 | 1 | R3IA-c-29 |
| ring-necked pheasant | Game Bird | winter | IA | Pottawattamie | 3 | cropland | R3IA-c-29 | 2015-12-14T00:00:00Z | 4 | 7 | 1 | R3IA-c-29 |
| ring-necked pheasant | Game Bird | winter | IA | Pottawattamie | 3 | cropland | R3IA-c-29 | 2015-12-28T00:00:00Z | 21 | 31 | 1 | R3IA-c-29 |
| ring-necked pheasant | Game Bird | winter | IA | Pottawattamie | 3 | cropland | R3IA-c-29 | 2015-12-28T00:00:00Z | 3 | 4 | 1 | R3IA-c-29 |
| ring-necked pheasant | Game Bird | winter | IA | Pottawattamie | 3 | cropland | R3IA-c-29 | 2016-01-11T00:00:00Z | 10 | 14 | 1 | R3IA-c-29 |
| red-tailed hawk | Raptor | winter | IA | Pottawattamie | 3 | cropland | R3IA-c-29 | 2015-12-28T00:00:00Z | 31 | 61 | 1 | R3IA-c-0 |
| red-tailed hawk | Raptor | winter | IA | Pottawattamie | 3 | cropland | R3IA-c-29 | 2015-12-28T00:00:00Z | 44 | 51 | 1 | R3IA-c-0 |
| ring-necked pheasant | Game Bird | winter | IA | Pottawattamie | 3 | cropland | R3IA-c-29 | 2016-01-25T00:00:00Z | 7 | 30 | 1 | R3IA-c-29 |
| ring-necked pheasant | Game Bird | winter | IA | Pottawattamie | 3 | cropland | R3IA-c-29 | 2016-01-25T00:00:00Z | 4 | 7 | 1 | R3IA-c-29 |
| ring-necked pheasant | Game Bird | winter | IA | Pottawattamie | 3 | cropland | R3IA-c-29 | 2016-02-09T00:00:00Z | 4 | 7 | 1 | R3IA-c-29 |
| ring-necked pheasant | Game Bird | winter | IA | Pottawattamie | 3 | cropland | R3IA-c-29 | 2016-02-09T00:00:00Z | 10 | 14 | 1 | R3IA-c-29 |
| ring-necked pheasant | Game Bird | winter | IA | Grundy | 3 | cropland | R3IA-c-30 | 2015-12-07T00:00:00Z | 3 | 4 | 0 | R3IA-c-30 |
| ring-necked pheasant | Game Bird | winter | IA | Grundy | 3 | cropland | R3IA-c-30 | 2015-12-07T00:00:00Z | 4 | 7 | 0 | R3IA-c-30 |
| ring-necked pheasant | Game Bird | winter | IA | Grundy | 3 | cropland | R3IA-c-30 | 2016-01-04T00:00:00Z | 1 | 1 | 0 | R3IA-c-30 |
| ring-necked pheasant | Game Bird | winter | IA | Grundy | 3 | cropland | R3IA-c-30 | 2016-01-04T00:00:00Z | 21 | 31 | 0 | R3IA-c-30 |
| ring-necked pheasant | Game Bird | winter | IA | Grundy | 3 | cropland | R3IA-c-30 | 2016-01-18T00:00:00Z | 7 | 10 | 0 | R3IA-c-30 |
| ring-necked pheasant | Game Bird | winter | IA | Grundy | 3 | cropland | R3IA-c-30 | 2016-01-25T00:00:00Z | 15 | 30 | 0 | R3IA-c-30 |
| ring-necked pheasant | Game Bird | winter | IA | Grundy | 3 | cropland | R3IA-c-30 | 2016-02-09T00:00:00Z | 4 | 7 | 0 | R3IA-c-30 |
| turkey vulture | Raptor | spring | IA | Grundy | 3 | cropland | R3IA-c-30 | 2016-03-25T00:00:00Z | 30 | NA | 1 | R3IA-c-0 |
| turkey vulture | Raptor | spring | IA | Grundy | 3 | cropland | R3IA-c-30 | 2016-03-31T00:00:00Z | 30 | NA | 1 | R3IA-c-0 |
| ring-necked pheasant | Game Bird | spring | IA | Grundy | 3 | cropland | R3IA-c-30 | 2016-04-07T00:00:00Z | 2 | 3 | 0 | R3IA-c-30 |
| ring-necked pheasant | Game Bird | spring | IA | Grundy | 3 | cropland | R3IA-c-30 | 2016-04-07T00:00:00Z | 4 | 7 | 0 | R3IA-c-30 |
| ring-necked pheasant | Game Bird | spring | IA | Grundy | 3 | cropland | R3IA-c-30 | 2016-04-07T00:00:00Z | 10 | 14 | 0 | R3IA-c-30 |
| red-tailed hawk | Raptor | spring | IA | Grundy | 3 | cropland | R3IA-c-30 | 2016-04-12T00:00:00Z | 30 | NA | 1 | R3IA-c-0 |
| turkey vulture | Raptor | spring | IA | Grundy | 3 | cropland | R3IA-c-30 | 2016-04-12T00:00:00Z | 30 | NA | 1 | R3IA-c-0 |
| red-tailed hawk | Raptor | spring | IA | Grundy | 3 | cropland | R3IA-c-30 | 2016-04-12T00:00:00Z | 30 | NA | 1 | R3IA-c-0 |
| red-tailed hawk | Raptor | spring | IA | Grundy | 3 | cropland | R3IA-c-30 | 2016-04-12T00:00:00Z | 30 | NA | 1 | R3IA-c-0 |
| ring-necked pheasant | Game Bird | summer | IA | Grundy | 3 | cropland | R3IA-c-30 | 2016-05-24T00:00:00Z | 30 | NA | 0 | R3IA-c-30 |
| red-tailed hawk | Raptor | fall | IA | Grundy | 3 | cropland | R3IA-c-30 | 2016-09-26T00:00:00Z | 10 | 14 | 1 | R3IA-c-0 |
| ring-necked pheasant | Game Bird | winter | IA | Grundy | 3 | cropland | R3IA-c-30 | 2016-11-28T00:00:00Z | 1 | 1 | 0 | R3IA-c-30 |
| ring-necked pheasant | Game Bird | winter | IA | Grundy | 3 | cropland | R3IA-c-30 | 2016-11-28T00:00:00Z | 1 | 1 | 0 | R3IA-c-30 |
| ring-necked pheasant | Game Bird | winter | IA | Grundy | 3 | cropland | R3IA-c-30 | 2016-12-12T00:00:00Z | 1 | 2 | 0 | R3IA-c-30 |
| ring-necked pheasant | Game Bird | winter | IA | Grundy | 3 | cropland | R3IA-c-30 | 2016-12-12T00:00:00Z | 17 | 23 | 0 | R3IA-c-30 |
| ring-necked pheasant | Game Bird | winter | IA | Grundy | 3 | cropland | R3IA-c-30 | 2017-01-09T00:00:00Z | 4 | 10 | 0 | R3IA-c-30 |
| ring-necked pheasant | Game Bird | winter | IA | Grundy | 3 | cropland | R3IA-c-30 | 2017-01-09T00:00:00Z | 14 | 20 | 0 | R3IA-c-30 |
| ring-necked pheasant | Game Bird | winter | IA | Grundy | 3 | cropland | R3IA-c-30 | 2017-01-23T00:00:00Z | 1 | 10 | 0 | R3IA-c-30 |
| ring-necked pheasant | Game Bird | winter | IA | Grundy | 3 | cropland | R3IA-c-30 | 2017-01-23T00:00:00Z | 1 | 7 | 0 | R3IA-c-30 |
| ring-necked pheasant | Game Bird | winter | IA | Grundy | 3 | cropland | R3IA-c-30 | 2017-02-06T00:00:00Z | 4 | 7 | 0 | R3IA-c-30 |
| ring-necked pheasant | Game Bird | winter | IA | Grundy | 3 | cropland | R3IA-c-30 | 2017-02-06T00:00:00Z | 4 | 7 | 0 | R3IA-c-30 |
| ring-necked pheasant | Game Bird | summer | IA | Grundy | 3 | cropland | R3IA-c-30 | 2016-06-09T00:00:00Z | 32 | NA | 0 | R3IA-c-30 |
| ring-necked pheasant | Game Bird | fall | IA | Grundy | 3 | cropland | R3IA-c-30 | 2016-08-01T00:00:00Z | 14 | 20 | 0 | R3IA-c-30 |
| ring-necked pheasant | Game Bird | fall | IA | Grundy | 3 | cropland | R3IA-c-30 | 2016-08-29T00:00:00Z | 1 | 2 | 0 | R3IA-c-30 |
| great horned owl | Raptor | summer | MN | Lincoln | 3 | cropland | R3MN-c-3 | 2018-05-29T00:00:00Z | 89.9652778 | NA | 1 | R3MN-c-0 |
| broad-winged hawk | Raptor | summer | MN | Lincoln | 3 | cropland | R3MN-c-3 | 2018-05-30T00:00:00Z | 89.0395833 | NA | 1 | R3MN-c-0 |
| red-shouldered hawk | Raptor | summer | MN | Lincoln | 3 | cropland | R3MN-c-3 | 2018-05-30T00:00:00Z | 9.2319444 | 13.1694444 | 1 | R3MN-c-0 |
| peregrine falcon | Raptor | summer | MN | Lincoln | 3 | cropland | R3MN-c-3 | 2018-05-29T00:00:00Z | 10.1458333 | 14.1076389 | 1 | R3MN-c-0 |
| broad-winged hawk | Raptor | summer | MN | Lincoln | 3 | cropland | R3MN-c-3 | 2018-05-30T00:00:00Z | 1.1291667 | 1.1291667 | 1 | R3MN-c-0 |
| Cooper's hawk | Raptor | spring | MN | Lincoln | 3 | cropland | R3MN-c-3 | 2018-04-26T00:00:00Z | 69.3923611 | 82.0277778 | 1 | R3MN-c-0 |
| Cooper's hawk | Raptor | spring | MN | Lincoln | 3 | cropland | R3MN-c-3 | 2018-04-26T00:00:00Z | 49.0965278 | 61.1409722 | 1 | R3MN-c-0 |
| Cooper's hawk | Raptor | spring | MN | Lincoln | 3 | cropland | R3MN-c-3 | 2018-04-25T00:00:00Z | 71.33125 | 80.0854167 | 1 | R3MN-c-0 |
| Cooper's hawk | Raptor | spring | MN | Lincoln | 3 | cropland | R3MN-c-3 | 2018-04-26T00:00:00Z | 70.4006944 | 79.2763889 | 1 | R3MN-c-0 |
| red-tailed hawk | Raptor | fall | MN | Lincoln | 3 | cropland | R3MN-c-3 | 2018-09-17T00:00:00Z | 72.2597222 | NA | 1 | R3MN-c-0 |
| barred owl | Raptor | fall | MN | Lincoln | 3 | cropland | R3MN-c-3 | 2018-09-18T00:00:00Z | 89.9833333 | NA | 1 | R3MN-c-0 |
| Cooper's hawk | Raptor | fall | MN | Lincoln | 3 | cropland | R3MN-c-3 | 2018-09-17T00:00:00Z | 79.1270833 | NA | 1 | R3MN-c-0 |
| red-tailed hawk | Raptor | fall | MN | Lincoln | 3 | cropland | R3MN-c-3 | 2018-09-17T00:00:00Z | 72.2458333 | NA | 1 | R3MN-c-0 |
| great horned owl | Raptor | fall | MN | Lincoln | 3 | cropland | R3MN-c-3 | 2018-09-18T00:00:00Z | 13.1458333 | 20.0819444 | 1 | R3MN-c-0 |
| red-shouldered hawk | Raptor | fall | MN | Lincoln | 3 | cropland | R3MN-c-3 | 2018-09-18T00:00:00Z | 39.1111111 | 49.0763889 | 1 | R3MN-c-0 |
| great horned owl | Raptor | fall | MN | Lincoln | 3 | cropland | R3MN-c-3 | 2018-09-18T00:00:00Z | 90.2708333 | NA | 1 | R3MN-c-0 |
| red-tailed hawk | Raptor | spring | MO | Atchison | 3 | cropland | R3MO-c-1 | 2018-03-12T00:00:00Z | 1.0152778 | 1.9430556 | 0 | R3MO-c-1 |
| red-tailed hawk | Raptor | spring | MO | Atchison | 3 | cropland | R3MO-c-1 | 2018-03-12T00:00:00Z | 3.0048611 | 4.0180556 | 0 | R3MO-c-1 |
| red-tailed hawk | Raptor | spring | MO | Atchison | 3 | cropland | R3MO-c-1 | 2018-04-05T00:00:00Z | 1.0215278 | 2.0263889 | 0 | R3MO-c-1 |
| red-tailed hawk | Raptor | spring | MO | Atchison | 3 | cropland | R3MO-c-1 | 2018-04-05T00:00:00Z | 2.0541667 | 2.9881944 | 0 | R3MO-c-1 |
| red-tailed hawk | Raptor | spring | MO | Atchison | 3 | cropland | R3MO-c-1 | 2018-04-10T00:00:00Z | 6.9493056 | 9.8840278 | 0 | R3MO-c-1 |
| red-tailed hawk | Raptor | summer | MO | Atchison | 3 | cropland | R3MO-c-1 | 2018-06-11T00:00:00Z | 9.7798611 | 13.9201389 | 0 | R3MO-c-1 |
| red-tailed hawk | Raptor | summer | MO | Atchison | 3 | cropland | R3MO-c-1 | 2018-05-21T00:00:00Z | 9.8534722 | 13.9131944 | 0 | R3MO-c-1 |
| red-tailed hawk | Raptor | spring | MO | Atchison | 3 | cropland | R3MO-c-1 | 2018-04-05T00:00:00Z | 29.7395833 | 48.1034722 | 0 | R3MO-c-1 |
| great horned owl | Raptor | spring | MO | Atchison | 3 | cropland | R3MO-c-1 | 2018-04-05T00:00:00Z | 16.0652778 | 20 | 0 | R3MO-c-1 |
| red-tailed hawk | Raptor | spring | MO | Atchison | 3 | cropland | R3MO-c-1 | 2018-04-05T00:00:00Z | 19.8826389 | 29.7270833 | 0 | R3MO-c-1 |
| snowy owl | Raptor | spring | MO | Atchison | 3 | cropland | R3MO-c-1 | 2018-04-05T00:00:00Z | 20.2819444 | 30.3041667 | 0 | R3MO-c-1 |
| Cooper's hawk | Raptor | spring | MO | Atchison | 3 | cropland | R3MO-c-1 | 2018-04-05T00:00:00Z | 15.9430556 | 29.9902778 | 0 | R3MO-c-1 |
| red-tailed hawk | Raptor | spring | MO | Atchison | 3 | cropland | R3MO-c-1 | 2018-04-05T00:00:00Z | 60.0375 | 68.0909722 | 0 | R3MO-c-1 |
| red-tailed hawk | Raptor | spring | MO | Atchison | 3 | cropland | R3MO-c-1 | 2018-04-10T00:00:00Z | 14.3673611 | 21.2625 | 0 | R3MO-c-1 |
| red-tailed hawk | Raptor | spring | MO | Atchison | 3 | cropland | R3MO-c-1 | 2018-04-10T00:00:00Z | 20.0590278 | 29.6277778 | 0 | R3MO-c-1 |
| red-tailed hawk | Raptor | spring | MO | Atchison | 3 | cropland | R3MO-c-1 | 2018-04-10T00:00:00Z | 13.6256944 | 19.7270833 | 0 | R3MO-c-1 |
| red-tailed hawk | Raptor | spring | MO | Atchison | 3 | cropland | R3MO-c-1 | 2018-04-10T00:00:00Z | 9.9583333 | 14.3888889 | 0 | R3MO-c-1 |
| red-tailed hawk | Raptor | spring | MO | Atchison | 3 | cropland | R3MO-c-1 | 2018-04-10T00:00:00Z | 43.2645833 | 55.2270833 | 0 | R3MO-c-1 |
| red-tailed hawk | Raptor | spring | MO | Atchison | 3 | cropland | R3MO-c-1 | 2018-04-10T00:00:00Z | 9.9013889 | 20.0652778 | 0 | R3MO-c-1 |
| red-tailed hawk | Raptor | summer | MO | Atchison | 3 | cropland | R3MO-c-1 | 2018-05-21T00:00:00Z | 9.8090278 | 13.8722222 | 0 | R3MO-c-1 |
| great horned owl | Raptor | summer | MO | Atchison | 3 | cropland | R3MO-c-1 | 2018-05-21T00:00:00Z | 6.8847222 | 8.675 | 0 | R3MO-c-1 |
| red-tailed hawk | Raptor | summer | MO | Atchison | 3 | cropland | R3MO-c-1 | 2018-05-21T00:00:00Z | 9.7715278 | 13.8423611 | 0 | R3MO-c-1 |
| red-tailed hawk | Raptor | summer | MO | Atchison | 3 | cropland | R3MO-c-1 | 2018-05-21T00:00:00Z | 1.99375 | 1.99375 | 0 | R3MO-c-1 |
| red-tailed hawk | Raptor | summer | MO | Atchison | 3 | cropland | R3MO-c-1 | 2018-05-21T00:00:00Z | 9.8069444 | 14.0638889 | 0 | R3MO-c-1 |
| great horned owl | Raptor | summer | MO | Atchison | 3 | cropland | R3MO-c-1 | 2018-05-21T00:00:00Z | 1.9409722 | 1.9409722 | 0 | R3MO-c-1 |
| red-tailed hawk | Raptor | summer | MO | Atchison | 3 | cropland | R3MO-c-1 | 2018-05-21T00:00:00Z | 6.8708333 | 9.7875 | 0 | R3MO-c-1 |
| red-tailed hawk | Raptor | fall | MO | Atchison | 3 | cropland | R3MO-c-1 | 2018-08-13T00:00:00Z | 29.7520833 | 49.9923611 | 0 | R3MO-c-1 |
| red-tailed hawk | Raptor | winter | MO | Atchison | 3 | cropland | R3MO-c-1 | 2018-12-04T00:00:00Z | 0.8541667 | 1.9979167 | 0 | R3MO-c-1 |
| red-tailed hawk | Raptor | winter | MO | Atchison | 3 | cropland | R3MO-c-1 | 2018-12-04T00:00:00Z | 9.6541667 | 23.9680556 | 0 | R3MO-c-1 |
| great horned owl | Raptor | fall | MO | Atchison | 3 | cropland | R3MO-c-1 | 2018-11-13T00:00:00Z | 3 | 7.01875 | 0 | R3MO-c-1 |
| red-tailed hawk | Raptor | winter | MO | Atchison | 3 | cropland | R3MO-c-1 | 2018-12-04T00:00:00Z | 2.8333333 | 2.8333333 | 0 | R3MO-c-1 |
| Cooper's hawk | Raptor | fall | MO | Atchison | 3 | cropland | R3MO-c-1 | 2018-11-02T00:00:00Z | 5.9340278 | 9.8055556 | 0 | R3MO-c-1 |
| great horned owl | Raptor | fall | MO | Atchison | 3 | cropland | R3MO-c-1 | 2018-11-13T00:00:00Z | 3.2159722 | 5.94375 | 0 | R3MO-c-1 |
| ring-necked pheasant | Game Bird | spring | MN | Pipestone and Murray | 3 | cropland | R3MN-c-4 | 2019-04-22T00:00:00Z | 4.0208333 | 7.0979167 | 1 | R3MN-c-4 |
| ring-necked pheasant | Game Bird | spring | MN | Pipestone and Murray | 3 | cropland | R3MN-c-4 | 2019-04-22T00:00:00Z | 6.99375 | 9.9479167 | 1 | R3MN-c-4 |
| ring-necked pheasant | Game Bird | spring | MN | Pipestone and Murray | 3 | cropland | R3MN-c-4 | 2019-04-22T00:00:00Z | 6.9958333 | 9.9375 | 1 | R3MN-c-4 |
| ring-necked pheasant | Game Bird | spring | MN | Pipestone and Murray | 3 | cropland | R3MN-c-4 | 2019-05-06T00:00:00Z | 1.9506944 | 2.9833333 | 1 | R3MN-c-4 |
| ring-necked pheasant | Game Bird | spring | MN | Pipestone and Murray | 3 | cropland | R3MN-c-4 | 2019-05-06T00:00:00Z | 4.1354167 | 7.225 | 1 | R3MN-c-4 |
| ring-necked pheasant | Game Bird | spring | MN | Pipestone and Murray | 3 | cropland | R3MN-c-4 | 2019-04-22T00:00:00Z | 13.9611111 | 20.1493056 | 1 | R3MN-c-4 |
| ring-necked pheasant | Game Bird | spring | MN | Pipestone and Murray | 3 | cropland | R3MN-c-4 | 2019-04-22T00:00:00Z | 31.1722222 | NA | 1 | R3MN-c-4 |
| ring-necked pheasant | Game Bird | spring | MN | Pipestone and Murray | 3 | cropland | R3MN-c-4 | 2019-04-22T00:00:00Z | 31.3284722 | NA | 1 | R3MN-c-4 |
| ring-necked pheasant | Game Bird | spring | MN | Pipestone and Murray | 3 | cropland | R3MN-c-4 | 2019-04-22T00:00:00Z | 30.9375 | NA | 1 | R3MN-c-4 |
| ring-necked pheasant | Game Bird | spring | MN | Pipestone and Murray | 3 | cropland | R3MN-c-4 | 2019-04-22T00:00:00Z | 31.0631944 | NA | 1 | R3MN-c-4 |
| ring-necked pheasant | Game Bird | summer | MN | Pipestone and Murray | 3 | cropland | R3MN-c-4 | 2019-05-20T00:00:00Z | 0.8680556 | 0.8680556 | 1 | R3MN-c-4 |
| ring-necked pheasant | Game Bird | summer | MN | Pipestone and Murray | 3 | cropland | R3MN-c-4 | 2019-05-20T00:00:00Z | 3.9131944 | 7.0770833 | 1 | R3MN-c-4 |
| ring-necked pheasant | Game Bird | summer | MN | Pipestone and Murray | 3 | cropland | R3MN-c-4 | 2019-05-20T00:00:00Z | 4.0020833 | 7.0854167 | 1 | R3MN-c-4 |
| ring-necked pheasant | Game Bird | spring | MN | Pipestone and Murray | 3 | cropland | R3MN-c-4 | 2019-05-06T00:00:00Z | 31.2819444 | NA | 1 | R3MN-c-4 |
| ring-necked pheasant | Game Bird | spring | MN | Pipestone and Murray | 3 | cropland | R3MN-c-4 | 2019-05-06T00:00:00Z | 14.0402778 | 20.2375 | 1 | R3MN-c-4 |
| ring-necked pheasant | Game Bird | spring | MN | Pipestone and Murray | 3 | cropland | R3MN-c-4 | 2019-05-06T00:00:00Z | 31.3673611 | NA | 1 | R3MN-c-4 |
| ring-necked pheasant | Game Bird | spring | MN | Pipestone and Murray | 3 | cropland | R3MN-c-4 | 2019-05-06T00:00:00Z | 31.0451389 | NA | 1 | R3MN-c-4 |
| ring-necked pheasant | Game Bird | spring | MN | Pipestone and Murray | 3 | cropland | R3MN-c-4 | 2019-05-06T00:00:00Z | 20.1145833 | 31.0604167 | 1 | R3MN-c-4 |
| ring-necked pheasant | Game Bird | spring | MN | Pipestone and Murray | 3 | cropland | R3MN-c-4 | 2019-05-06T00:00:00Z | 30.9131944 | NA | 1 | R3MN-c-4 |
| barred owl | Raptor | spring | MN | Pipestone and Murray | 3 | cropland | R3MN-c-4 | 2019-05-02T00:00:00Z | 40.1680556 | 42.2048611 | 1 | R3MN-c-0 |
| ring-necked pheasant | Game Bird | summer | MN | Pipestone and Murray | 3 | cropland | R3MN-c-4 | 2019-05-20T00:00:00Z | 7.3055556 | 11.2506944 | 1 | R3MN-c-4 |
| ring-necked pheasant | Game Bird | summer | MN | Pipestone and Murray | 3 | cropland | R3MN-c-4 | 2019-05-20T00:00:00Z | 14.3229167 | 21.2118056 | 1 | R3MN-c-4 |
| ring-necked pheasant | Game Bird | summer | MN | Pipestone and Murray | 3 | cropland | R3MN-c-4 | 2019-06-17T00:00:00Z | 4.0694444 | 8.1680556 | 1 | R3MN-c-4 |
| ring-necked pheasant | Game Bird | summer | MN | Pipestone and Murray | 3 | cropland | R3MN-c-4 | 2019-06-17T00:00:00Z | 3.8888889 | 6.9840278 | 1 | R3MN-c-4 |
| ring-necked pheasant | Game Bird | summer | MN | Pipestone and Murray | 3 | cropland | R3MN-c-4 | 2019-06-17T00:00:00Z | 0.9243056 | 0.9243056 | 1 | R3MN-c-4 |
| ring-necked pheasant | Game Bird | summer | MN | Pipestone and Murray | 3 | cropland | R3MN-c-4 | 2019-07-08T00:00:00Z | 1.0625 | 1.0625 | 1 | R3MN-c-4 |
| ring-necked pheasant | Game Bird | summer | MN | Pipestone and Murray | 3 | cropland | R3MN-c-4 | 2019-07-08T00:00:00Z | 1.0798611 | 2.0888889 | 1 | R3MN-c-4 |
| ring-necked pheasant | Game Bird | summer | MN | Pipestone and Murray | 3 | cropland | R3MN-c-4 | 2019-07-08T00:00:00Z | 2.9277778 | 4.1409722 | 1 | R3MN-c-4 |
| ring-necked pheasant | Game Bird | summer | MN | Pipestone and Murray | 3 | cropland | R3MN-c-4 | 2019-06-17T00:00:00Z | 30.4854167 | NA | 1 | R3MN-c-4 |
| ring-necked pheasant | Game Bird | summer | MN | Pipestone and Murray | 3 | cropland | R3MN-c-4 | 2019-06-17T00:00:00Z | 30.4152778 | NA | 1 | R3MN-c-4 |
| ring-necked pheasant | Game Bird | summer | MN | Pipestone and Murray | 3 | cropland | R3MN-c-4 | 2019-06-17T00:00:00Z | 30.1763889 | NA | 1 | R3MN-c-4 |
| Cooper's hawk | Raptor | spring | MN | Pipestone and Murray | 3 | cropland | R3MN-c-4 | 2019-05-02T00:00:00Z | 61.1180556 | 70.2944444 | 1 | R3MN-c-0 |
| ring-necked pheasant | Game Bird | summer | MN | Pipestone and Murray | 3 | cropland | R3MN-c-4 | 2019-07-08T00:00:00Z | 14.0361111 | 20.9895833 | 1 | R3MN-c-4 |
| ring-necked pheasant | Game Bird | summer | MN | Pipestone and Murray | 3 | cropland | R3MN-c-4 | 2019-07-08T00:00:00Z | 9.7979167 | 14.0534722 | 1 | R3MN-c-4 |
| Cooper's hawk | Raptor | spring | MN | Pipestone and Murray | 3 | cropland | R3MN-c-4 | 2019-05-02T00:00:00Z | 81.2243056 | 91.2694444 | 1 | R3MN-c-0 |
| red-tailed hawk | Raptor | spring | MN | Pipestone and Murray | 3 | cropland | R3MN-c-4 | 2019-04-22T00:00:00Z | 79.9402778 | 90.9895833 | 1 | R3MN-c-0 |
| red-tailed hawk | Raptor | spring | MN | Pipestone and Murray | 3 | cropland | R3MN-c-4 | 2019-05-02T00:00:00Z | 81.1888889 | 91.1930556 | 1 | R3MN-c-0 |
| great horned owl | Raptor | spring | MN | Pipestone and Murray | 3 | cropland | R3MN-c-4 | 2019-05-02T00:00:00Z | 120.15625 | NA | 1 | R3MN-c-0 |
| barred owl | Raptor | winter | MO | Atchison | 3 | cropland | R3MO-c-2 | 2019-01-07T00:00:00Z | 32.1916667 | 52.2784722 | 0 | R3MO-c-2 |
| red-tailed hawk | Raptor | winter | MO | Atchison | 3 | cropland | R3MO-c-2 | 2019-01-07T00:00:00Z | 32.0416667 | 63.1125 | 0 | R3MO-c-2 |
| red-tailed hawk | Raptor | winter | MO | Atchison | 3 | cropland | R3MO-c-2 | 2019-01-29T00:00:00Z | 41.2020833 | 50.0652778 | 0 | R3MO-c-2 |
| red-tailed hawk | Raptor | winter | MO | Atchison | 3 | cropland | R3MO-c-2 | 2019-01-29T00:00:00Z | 10.0340278 | 41.1270833 | 0 | R3MO-c-2 |
| red-tailed hawk | Raptor | winter | MO | Atchison | 3 | cropland | R3MO-c-2 | 2019-01-29T00:00:00Z | 9.9958333 | 9.9958333 | 0 | R3MO-c-2 |
| red-tailed hawk | Raptor | winter | MO | Atchison | 3 | cropland | R3MO-c-2 | 2019-01-29T00:00:00Z | 30.0340278 | 41.05 | 0 | R3MO-c-2 |
| turkey vulture | Raptor | winter | MO | Atchison | 3 | cropland | R3MO-c-2 | 2019-01-07T00:00:00Z | 1.1625 | 1.8916667 | 0 | R3MO-c-2 |
| red-tailed hawk | Raptor | winter | MO | Atchison | 3 | cropland | R3MO-c-2 | 2019-01-29T00:00:00Z | 10.1868056 | 41.1472222 | 0 | R3MO-c-2 |
| short-eared owl | Raptor | winter | MO | Atchison | 3 | cropland | R3MO-c-2 | 2019-02-14T00:00:00Z | 13.9618056 | 20 | 0 | R3MO-c-2 |
| red-tailed hawk | Raptor | winter | MO | Atchison | 3 | cropland | R3MO-c-2 | 2019-02-14T00:00:00Z | 28.8736111 | 33 | 0 | R3MO-c-2 |
| red-tailed hawk | Raptor | winter | MO | Atchison | 3 | cropland | R3MO-c-2 | 2019-02-14T00:00:00Z | 0.66875 | 19.7090278 | 0 | R3MO-c-2 |
| red-tailed hawk | Raptor | winter | MO | Atchison | 3 | cropland | R3MO-c-2 | 2019-02-14T00:00:00Z | 28.8479167 | 38.7118056 | 0 | R3MO-c-2 |
| red-tailed hawk | Raptor | winter | MO | Atchison | 3 | cropland | R3MO-c-2 | 2019-02-14T00:00:00Z | 1.9743056 | 19.7243056 | 0 | R3MO-c-2 |
| great horned owl | Raptor | winter | MO | Atchison | 3 | cropland | R3MO-c-2 | 2019-01-07T00:00:00Z | 99 | NA | 0 | R3MO-c-2 |
| red-tailed hawk | Raptor | winter | MO | Atchison | 3 | cropland | R3MO-c-2 | 2019-02-14T00:00:00Z | 2 | 29 | 0 | R3MO-c-2 |
| turkey vulture | Raptor | winter | MO | Atchison | 3 | cropland | R3MO-c-2 | 2019-02-14T00:00:00Z | 29 | 39 | 0 | R3MO-c-2 |
| red-tailed hawk | Raptor | winter | MO | Atchison | 3 | cropland | R3MO-c-2 | 2018-12-04T00:00:00Z | 55.8875 | 97 | 0 | R3MO-c-2 |
| turkey vulture | Raptor | winter | MO | Atchison | 3 | cropland | R3MO-c-2 | 2018-12-04T00:00:00Z | 97 | 107.2493056 | 0 | R3MO-c-2 |
| red-tailed hawk | Raptor | winter | MO | Atchison | 3 | cropland | R3MO-c-2 | 2019-02-14T00:00:00Z | 19.7770833 | 81.9006944 | 0 | R3MO-c-2 |
| red-tailed hawk | Raptor | winter | MO | Atchison | 3 | cropland | R3MO-c-2 | 2019-02-14T00:00:00Z | 59.80625 | 81.89375 | 0 | R3MO-c-2 |
| turkey vulture | Raptor | winter | MO | Atchison | 3 | cropland | R3MO-c-2 | 2019-02-14T00:00:00Z | 81.8104167 | NA | 0 | R3MO-c-2 |
| red-tailed hawk | Raptor | winter | MO | Atchison | 3 | cropland | R3MO-c-2 | 2019-02-14T00:00:00Z | 28.8423611 | 81.8555556 | 0 | R3MO-c-2 |
| red-tailed hawk | Raptor | spring | MO | Atchison | 3 | cropland | R3MO-c-2 | 2019-03-19T00:00:00Z | 14.9381944 | 21 | 0 | R3MO-c-2 |
| red-tailed hawk | Raptor | spring | MO | Atchison | 3 | cropland | R3MO-c-2 | 2019-03-19T00:00:00Z | 10.7125 | 14.9236111 | 0 | R3MO-c-2 |
| Cooper's hawk | Raptor | spring | MO | Atchison | 3 | cropland | R3MO-c-2 | 2019-03-19T00:00:00Z | 6.9451389 | 10.6701389 | 0 | R3MO-c-2 |
| great horned owl | Raptor | spring | MO | Atchison | 3 | cropland | R3MO-c-2 | 2019-04-02T00:00:00Z | 19.8027778 | 29.7541667 | 0 | R3MO-c-2 |
| red-tailed hawk | Raptor | spring | MO | Atchison | 3 | cropland | R3MO-c-2 | 2019-04-02T00:00:00Z | 7.0951389 | 9.8180556 | 0 | R3MO-c-2 |
| Cooper's hawk | Raptor | spring | MO | Atchison | 3 | cropland | R3MO-c-2 | 2019-04-23T00:00:00Z | 9.8201389 | 14.8631944 | 0 | R3MO-c-2 |
| red-tailed hawk | Raptor | spring | MO | Atchison | 3 | cropland | R3MO-c-2 | 2019-03-19T00:00:00Z | 29.7013889 | 40.9229167 | 0 | R3MO-c-2 |
| red-tailed hawk | Raptor | spring | MO | Atchison | 3 | cropland | R3MO-c-2 | 2019-03-19T00:00:00Z | 6.9777778 | 14.85 | 0 | R3MO-c-2 |
| red-tailed hawk | Raptor | spring | MO | Atchison | 3 | cropland | R3MO-c-2 | 2019-03-19T00:00:00Z | 62.6548611 | 69.6736111 | 0 | R3MO-c-2 |
| great horned owl | Raptor | spring | MO | Atchison | 3 | cropland | R3MO-c-2 | 2019-04-02T00:00:00Z | 29.8527778 | 41.0930556 | 0 | R3MO-c-2 |
| red-tailed hawk | Raptor | spring | MO | Atchison | 3 | cropland | R3MO-c-2 | 2019-04-02T00:00:00Z | 72.0888889 | 87.0604167 | 0 | R3MO-c-2 |
| Cooper's hawk | Raptor | spring | MO | Atchison | 3 | cropland | R3MO-c-2 | 2019-04-02T00:00:00Z | 14.0131944 | 19.9354167 | 0 | R3MO-c-2 |
| red-tailed hawk | Raptor | spring | MO | Atchison | 3 | cropland | R3MO-c-2 | 2019-04-02T00:00:00Z | 29.8604167 | 40.8173611 | 0 | R3MO-c-2 |
| red-tailed hawk | Raptor | spring | MO | Atchison | 3 | cropland | R3MO-c-2 | 2019-04-02T00:00:00Z | 13.9777778 | 19.8201389 | 0 | R3MO-c-2 |
| red-tailed hawk | Raptor | spring | MO | Atchison | 3 | cropland | R3MO-c-2 | 2019-04-02T00:00:00Z | 19.7895833 | 30.0041667 | 0 | R3MO-c-2 |
| red-tailed hawk | Raptor | spring | MO | Atchison | 3 | cropland | R3MO-c-2 | 2019-04-15T00:00:00Z | 9.6534722 | 13.7854167 | 0 | R3MO-c-2 |
| red-tailed hawk | Raptor | spring | MO | Atchison | 3 | cropland | R3MO-c-2 | 2019-04-15T00:00:00Z | 48.9194444 | 58.9222222 | 0 | R3MO-c-2 |
| red-tailed hawk | Raptor | spring | MO | Atchison | 3 | cropland | R3MO-c-2 | 2019-04-15T00:00:00Z | 38.6472222 | 48.9215278 | 0 | R3MO-c-2 |
| red-tailed hawk | Raptor | spring | MO | Atchison | 3 | cropland | R3MO-c-2 | 2019-04-15T00:00:00Z | 98.8583333 | 108.7215278 | 0 | R3MO-c-2 |
| red-tailed hawk | Raptor | spring | MO | Atchison | 3 | cropland | R3MO-c-2 | 2019-04-15T00:00:00Z | 9.6298611 | 13.6277778 | 0 | R3MO-c-2 |
| Cooper's hawk | Raptor | spring | MO | Atchison | 3 | cropland | R3MO-c-2 | 2019-04-15T00:00:00Z | 9.6090278 | 13.6069444 | 0 | R3MO-c-2 |
| red-tailed hawk | Raptor | spring | MO | Atchison | 3 | cropland | R3MO-c-2 | 2019-04-15T00:00:00Z | 29.6868056 | 38.7048611 | 0 | R3MO-c-2 |
| red-tailed hawk | Raptor | spring | MO | Atchison | 3 | cropland | R3MO-c-2 | 2019-04-15T00:00:00Z | 48.8951389 | 58.8333333 | 0 | R3MO-c-2 |
| red-tailed hawk | Raptor | summer | MO | Atchison | 3 | cropland | R3MO-c-2 | 2019-05-20T00:00:00Z | 6.9527778 | 10.19375 | 0 | R3MO-c-2 |
| great horned owl | Raptor | summer | MO | Atchison | 3 | cropland | R3MO-c-2 | 2019-05-20T00:00:00Z | 17.9944444 | 28.9513889 | 0 | R3MO-c-2 |
| red-tailed hawk | Raptor | summer | MO | Atchison | 3 | cropland | R3MO-c-2 | 2019-05-20T00:00:00Z | 28.9291667 | 42.9590278 | 0 | R3MO-c-2 |
| red-tailed hawk | Raptor | summer | MO | Atchison | 3 | cropland | R3MO-c-2 | 2019-05-20T00:00:00Z | 9.9381944 | 17.8965278 | 0 | R3MO-c-2 |
| turkey vulture | Raptor | summer | MO | Atchison | 3 | cropland | R3MO-c-2 | 2019-05-20T00:00:00Z | 28.86875 | 42.9430556 | 0 | R3MO-c-2 |
| red-tailed hawk | Raptor | summer | MO | Atchison | 3 | cropland | R3MO-c-2 | 2019-05-20T00:00:00Z | 3.7819444 | 6.9833333 | 0 | R3MO-c-2 |
| red-tailed hawk | Raptor | summer | MO | Atchison | 3 | cropland | R3MO-c-2 | 2019-06-25T00:00:00Z | 13.8402778 | 19.7652778 | 0 | R3MO-c-2 |
| red-tailed hawk | Raptor | summer | MO | Atchison | 3 | cropland | R3MO-c-2 | 2019-05-20T00:00:00Z | 71.1243056 | 93.1506944 | 0 | R3MO-c-2 |
| red-tailed hawk | Raptor | spring | MO | Atchison | 3 | cropland | R3MO-c-2 | 2019-03-19T00:00:00Z | 30.1201389 | 40.9020833 | 0 | R3MO-c-2 |
| red-tailed hawk | Raptor | spring | MO | Atchison | 3 | cropland | R3MO-c-2 | 2019-03-19T00:00:00Z | 14.8145833 | 19.89375 | 0 | R3MO-c-2 |
| Cooper's hawk | Raptor | winter | MO | Atchison | 3 | cropland | R3MO-c-2 | 2019-11-18T00:00:00Z | 3 | 4.2020833 | 0 | R3MO-c-2 |
| great horned owl | Raptor | winter | MO | Atchison | 3 | cropland | R3MO-c-2 | 2019-11-18T00:00:00Z | 14.3194444 | 19.9111111 | 0 | R3MO-c-2 |
| red-tailed hawk | Raptor | winter | MO | Atchison | 3 | cropland | R3MO-c-2 | 2019-11-18T00:00:00Z | 8.9409722 | 14.2798611 | 0 | R3MO-c-2 |
| Cooper's hawk | Raptor | winter | MO | Atchison | 3 | cropland | R3MO-c-2 | 2019-11-18T00:00:00Z | 1.9402778 | 3 | 0 | R3MO-c-2 |
| red-tailed hawk | Raptor | winter | MO | Atchison | 3 | cropland | R3MO-c-2 | 2019-11-18T00:00:00Z | 7.0986111 | 8.9208333 | 0 | R3MO-c-2 |
| red-tailed hawk | Raptor | winter | MO | Atchison | 3 | cropland | R3MO-c-2 | 2019-12-03T00:00:00Z | 1.05625 | 2.0138889 | 0 | R3MO-c-2 |
| red-tailed hawk | Raptor | winter | MO | Atchison | 3 | cropland | R3MO-c-2 | 2019-12-03T00:00:00Z | 3.0263889 | 3.9895833 | 0 | R3MO-c-2 |
| red-tailed hawk | Raptor | winter | MO | Atchison | 3 | cropland | R3MO-c-2 | 2019-12-03T00:00:00Z | 7.1048611 | 9.2083333 | 0 | R3MO-c-2 |
| red-tailed hawk | Raptor | winter | MO | Atchison | 3 | cropland | R3MO-c-2 | 2019-12-03T00:00:00Z | 3.9840278 | 7.0951389 | 0 | R3MO-c-2 |
| red-tailed hawk | Raptor | winter | MO | Atchison | 3 | cropland | R3MO-c-2 | 2019-12-03T00:00:00Z | 0.9833333 | 0.9833333 | 0 | R3MO-c-2 |
| red-tailed hawk | Raptor | winter | MO | Atchison | 3 | cropland | R3MO-c-2 | 2019-12-03T00:00:00Z | 0.9243056 | 0.9243056 | 0 | R3MO-c-2 |
| red-tailed hawk | Raptor | winter | MO | Atchison | 3 | cropland | R3MO-c-2 | 2019-12-03T00:00:00Z | 1.9819444 | 2.8770833 | 0 | R3MO-c-2 |
| red-tailed hawk | Raptor | winter | MO | Atchison | 3 | cropland | R3MO-c-2 | 2019-12-16T00:00:00Z | 0.9152778 | 0.9152778 | 0 | R3MO-c-2 |
| red-tailed hawk | Raptor | winter | MO | Atchison | 3 | cropland | R3MO-c-2 | 2019-12-03T00:00:00Z | 9.2381944 | 15.0652778 | 0 | R3MO-c-2 |
| red-tailed hawk | Raptor | winter | MO | Atchison | 3 | cropland | R3MO-c-2 | 2019-12-03T00:00:00Z | 8.9854167 | 15.0097222 | 0 | R3MO-c-2 |
| northern harrier | Raptor | winter | MO | Atchison | 3 | cropland | R3MO-c-2 | 2019-12-16T00:00:00Z | 7.0923611 | 9.7965278 | 0 | R3MO-c-2 |
| red-tailed hawk | Raptor | winter | MO | Atchison | 3 | cropland | R3MO-c-2 | 2019-12-16T00:00:00Z | 6.8243056 | 9.7486111 | 0 | R3MO-c-2 |
| red-tailed hawk | Raptor | winter | MO | Atchison | 3 | cropland | R3MO-c-2 | 2019-12-03T00:00:00Z | 15.0270833 | 19.95 | 0 | R3MO-c-2 |
| red-tailed hawk | Raptor | winter | MO | Atchison | 3 | cropland | R3MO-c-2 | 2019-11-18T00:00:00Z | 30.0979167 | 38.9791667 | 0 | R3MO-c-2 |
| red-tailed hawk | Raptor | winter | MO | Atchison | 3 | cropland | R3MO-c-2 | 2019-12-16T00:00:00Z | 9.775 | 14.8083333 | 0 | R3MO-c-2 |
| red-tailed hawk | Raptor | winter | MO | Atchison | 3 | cropland | R3MO-c-2 | 2019-12-16T00:00:00Z | 14.7354167 | 20.7819444 | 0 | R3MO-c-2 |
| red-tailed hawk | Raptor | winter | MO | Atchison | 3 | cropland | R3MO-c-2 | 2019-11-18T00:00:00Z | 59.2506944 | NA | 0 | R3MO-c-2 |
| red-tailed hawk | Raptor | winter | MO | Atchison | 3 | cropland | R3MO-c-2 | 2019-12-16T00:00:00Z | 31.0784722 | NA | 0 | R3MO-c-2 |
| ring-necked pheasant | Game Bird | fall | OH | Paulding | 3 | cropland | R3OH-c-1 | 2020-09-15T00:00:00Z | 7 | 10 | 1 | R3OH-c-1 |
| red-tailed hawk | Raptor | summer | OH | Paulding | 3 | cropland | R3OH-c-1 | 2020-07-05T00:00:00Z | 40 | NA | 1 | R3OH-c-1 |
| red-tailed hawk | Raptor | summer | OH | Paulding | 3 | cropland | R3OH-c-1 | 2020-07-05T00:00:00Z | 40 | NA | 1 | R3OH-c-1 |
| mallard | Game Bird | summer | OH | Paulding | 3 | cropland | R3OH-c-1 | 2020-07-05T00:00:00Z | 40 | NA | 1 | R3OH-c-1 |
| red-tailed hawk | Raptor | summer | OH | Paulding | 3 | cropland | R3OH-c-1 | 2020-07-05T00:00:00Z | 40 | NA | 1 | R3OH-c-1 |
| ring-necked pheasant | Game Bird | summer | OH | Paulding | 3 | cropland | R3OH-c-1 | 2020-07-07T00:00:00Z | 2 | 10 | 1 | R3OH-c-1 |
| mallard | Game Bird | summer | OH | Paulding | 3 | cropland | R3OH-c-1 | 2020-07-07T00:00:00Z | 4 | 4 | 1 | R3OH-c-1 |
| ring-necked pheasant | Game Bird | summer | OH | Paulding | 3 | cropland | R3OH-c-1 | 2020-07-07T00:00:00Z | 2 | 3 | 1 | R3OH-c-1 |
| mallard | Game Bird | summer | OH | Paulding | 3 | cropland | R3OH-c-1 | 2020-07-09T00:00:00Z | 2 | 3.1659722 | 1 | R3OH-c-1 |
| ring-necked pheasant | Game Bird | summer | OH | Paulding | 3 | cropland | R3OH-c-1 | 2020-07-13T00:00:00Z | 1 | 4 | 1 | R3OH-c-1 |
| ring-necked pheasant | Game Bird | fall | OH | Paulding | 3 | cropland | R3OH-c-1 | 2020-09-09T00:00:00Z | 40 | NA | 1 | R3OH-c-1 |
| ring-necked pheasant | Game Bird | fall | OH | Paulding | 3 | cropland | R3OH-c-1 | 2020-08-19T00:00:00Z | 7 | 10 | 1 | R3OH-c-1 |
| red-tailed hawk | Raptor | fall | OH | Paulding | 3 | cropland | R3OH-c-1 | 2020-08-19T00:00:00Z | 40 | NA | 1 | R3OH-c-1 |
| mallard | Game Bird | fall | OH | Paulding | 3 | cropland | R3OH-c-1 | 2020-09-01T00:00:00Z | 39 | NA | 1 | R3OH-c-1 |
| ring-necked pheasant | Game Bird | fall | OH | Paulding | 3 | cropland | R3OH-c-1 | 2020-09-01T00:00:00Z | 3 | 4 | 1 | R3OH-c-1 |
| ring-necked pheasant | Game Bird | fall | OH | Paulding | 3 | cropland | R3OH-c-1 | 2020-09-09T00:00:00Z | 4 | 7 | 1 | R3OH-c-1 |
| ring-necked pheasant | Game Bird | fall | OH | Paulding | 3 | cropland | R3OH-c-1 | 2020-09-11T00:00:00Z | 1 | 2 | 1 | R3OH-c-1 |
| Cooper's hawk | Raptor | fall | OH | Paulding | 3 | cropland | R3OH-c-1 | 2020-09-27T00:00:00Z | 1 | 1 | 1 | R3OH-c-1 |
| red-tailed hawk | Raptor | fall | OH | Paulding | 3 | cropland | R3OH-c-1 | 2020-09-27T00:00:00Z | 21 | NA | 1 | R3OH-c-1 |
| mallard | Game Bird | fall | OH | Paulding | 3 | cropland | R3OH-c-1 | 2020-09-27T00:00:00Z | 4 | 7 | 1 | R3OH-c-1 |
| ring-necked pheasant | Game Bird | fall | OH | Paulding | 3 | cropland | R3OH-c-1 | 2020-09-27T00:00:00Z | 10 | 14 | 1 | R3OH-c-1 |
| ring-necked pheasant | Game Bird | fall | OH | Paulding | 3 | cropland | R3OH-c-1 | 2020-09-27T00:00:00Z | 1 | 1 | 1 | R3OH-c-1 |
| turkey vulture | Raptor | fall | OH | Paulding | 3 | cropland | R3OH-c-1 | 2020-10-02T00:00:00Z | 40 | NA | 1 | R3OH-c-1 |
| mallard | Game Bird | fall | OH | Paulding | 3 | cropland | R3OH-c-1 | 2020-10-27T00:00:00Z | 4 | 7 | 1 | R3OH-c-1 |
| snowy owl | Raptor | fall | OH | Paulding | 3 | cropland | R3OH-c-1 | 2020-10-27T00:00:00Z | 19 | NA | 1 | R3OH-c-1 |
| red-tailed hawk | Raptor | winter | IA | Poweshiek | 3 | cropland | R3IA-c-31 | 2019-11-26T00:00:00Z | 0.9486111 | 2.8256944 | 0 | R3IA-c-31 |
| red-tailed hawk | Raptor | winter | IA | Poweshiek | 3 | cropland | R3IA-c-31 | 2019-11-26T00:00:00Z | 6.8305556 | 9.7833333 | 0 | R3IA-c-31 |
| red-tailed hawk | Raptor | winter | IA | Poweshiek | 3 | cropland | R3IA-c-31 | 2019-11-26T00:00:00Z | 20.0361111 | 30.0201389 | 0 | R3IA-c-31 |
| red-tailed hawk | Raptor | winter | IA | Poweshiek | 3 | cropland | R3IA-c-31 | 2019-11-26T00:00:00Z | 40.0930556 | 61.1666667 | 0 | R3IA-c-31 |
| red-tailed hawk | Raptor | winter | IA | Poweshiek | 3 | cropland | R3IA-c-31 | 2019-11-26T00:00:00Z | 0.9715278 | 2.8423611 | 0 | R3IA-c-31 |
| red-tailed hawk | Raptor | winter | IA | Poweshiek | 3 | cropland | R3IA-c-31 | 2019-12-10T00:00:00Z | 6.9888889 | 10.0375 | 0 | R3IA-c-31 |
| red-tailed hawk | Raptor | winter | IA | Poweshiek | 3 | cropland | R3IA-c-31 | 2019-12-10T00:00:00Z | 10.0368056 | 13.9777778 | 0 | R3IA-c-31 |
| red-tailed hawk | Raptor | winter | IA | Poweshiek | 3 | cropland | R3IA-c-31 | 2019-12-10T00:00:00Z | 30.0625 | 40.05625 | 0 | R3IA-c-31 |
| red-tailed hawk | Raptor | winter | IA | Poweshiek | 3 | cropland | R3IA-c-31 | 2019-12-10T00:00:00Z | 20.0534722 | 30.0618056 | 0 | R3IA-c-31 |
| red-tailed hawk | Raptor | winter | IA | Poweshiek | 3 | cropland | R3IA-c-31 | 2019-12-10T00:00:00Z | 13.9680556 | 20.0486111 | 0 | R3IA-c-31 |
| red-tailed hawk | Raptor | spring | IA | Poweshiek | 3 | cropland | R3IA-c-31 | 2020-03-16T00:00:00Z | 60.0875 | NA | 0 | R3IA-c-31 |
| red-tailed hawk | Raptor | spring | IA | Poweshiek | 3 | cropland | R3IA-c-31 | 2020-03-16T00:00:00Z | 0.8284722 | 1.8729167 | 0 | R3IA-c-31 |
| red-tailed hawk | Raptor | spring | IA | Poweshiek | 3 | cropland | R3IA-c-31 | 2020-03-16T00:00:00Z | 19.8333333 | 30.1173611 | 0 | R3IA-c-31 |
| snowy owl | Raptor | spring | IA | Poweshiek | 3 | cropland | R3IA-c-31 | 2020-03-16T00:00:00Z | 59.9173611 | NA | 0 | R3IA-c-31 |
| red-tailed hawk | Raptor | spring | IA | Poweshiek | 3 | cropland | R3IA-c-31 | 2020-03-16T00:00:00Z | 59.7868056 | NA | 0 | R3IA-c-31 |
| red-tailed hawk | Raptor | winter | IA | Poweshiek | 3 | cropland | R3IA-c-31 | 2020-01-07T00:00:00Z | 2.9875 | 6.9798611 | 0 | R3IA-c-31 |
| red-tailed hawk | Raptor | winter | IA | Poweshiek | 3 | cropland | R3IA-c-31 | 2020-01-07T00:00:00Z | 10.0340278 | 14.0173611 | 0 | R3IA-c-31 |
| red-tailed hawk | Raptor | winter | IA | Poweshiek | 3 | cropland | R3IA-c-31 | 2020-01-07T00:00:00Z | 20.0222222 | 30.0069444 | 0 | R3IA-c-31 |
| red-tailed hawk | Raptor | winter | IA | Poweshiek | 3 | cropland | R3IA-c-31 | 2020-01-07T00:00:00Z | 3.0291667 | 7.01875 | 0 | R3IA-c-31 |
| red-tailed hawk | Raptor | winter | IA | Poweshiek | 3 | cropland | R3IA-c-31 | 2020-01-07T00:00:00Z | 1.0493056 | 1.0493056 | 0 | R3IA-c-31 |
| red-tailed hawk | Raptor | spring | IA | Poweshiek | 3 | cropland | R3IA-c-31 | 2020-03-23T00:00:00Z | 30.1479167 | 39.9486111 | 0 | R3IA-c-31 |
| red-tailed hawk | Raptor | spring | IA | Poweshiek | 3 | cropland | R3IA-c-31 | 2020-03-23T00:00:00Z | 60.0166667 | NA | 0 | R3IA-c-31 |
| red-tailed hawk | Raptor | spring | IA | Poweshiek | 3 | cropland | R3IA-c-31 | 2020-03-23T00:00:00Z | 30.0201389 | 40.1402778 | 0 | R3IA-c-31 |
| red-tailed hawk | Raptor | spring | IA | Poweshiek | 3 | cropland | R3IA-c-31 | 2020-03-23T00:00:00Z | 59.6909722 | NA | 0 | R3IA-c-31 |
| red-tailed hawk | Raptor | spring | IA | Poweshiek | 3 | cropland | R3IA-c-31 | 2020-03-23T00:00:00Z | 59.7048611 | NA | 0 | R3IA-c-31 |
| red-tailed hawk | Raptor | spring | IA | Poweshiek | 3 | cropland | R3IA-c-32 | 2020-03-16T00:00:00Z | 59.8958333 | NA | 0 | R3IA-c-32 |
| snowy owl | Raptor | spring | IA | Poweshiek | 3 | cropland | R3IA-c-32 | 2020-03-16T00:00:00Z | 49.8638889 | 59.89375 | 0 | R3IA-c-32 |
| red-tailed hawk | Raptor | spring | IA | Poweshiek | 3 | cropland | R3IA-c-32 | 2020-03-16T00:00:00Z | 39.9076389 | 49.8590278 | 0 | R3IA-c-32 |
| red-tailed hawk | Raptor | spring | IA | Poweshiek | 3 | cropland | R3IA-c-32 | 2020-03-16T00:00:00Z | 20.2215278 | 29.9090278 | 0 | R3IA-c-32 |
| red-tailed hawk | Raptor | spring | IA | Poweshiek | 3 | cropland | R3IA-c-32 | 2020-03-16T00:00:00Z | 59.8416667 | NA | 0 | R3IA-c-32 |
| red-tailed hawk | Raptor | spring | IA | Poweshiek | 3 | cropland | R3IA-c-32 | 2020-03-20T00:00:00Z | 55.86875 | NA | 0 | R3IA-c-32 |
| red-tailed hawk | Raptor | spring | IA | Poweshiek | 3 | cropland | R3IA-c-32 | 2020-03-23T00:00:00Z | 13.9958333 | 19.7069444 | 0 | R3IA-c-32 |
| red-tailed hawk | Raptor | spring | IA | Poweshiek | 3 | cropland | R3IA-c-32 | 2020-03-23T00:00:00Z | 59.8666667 | NA | 0 | R3IA-c-32 |
| red-tailed hawk | Raptor | spring | IA | Poweshiek | 3 | cropland | R3IA-c-32 | 2020-03-23T00:00:00Z | 13.9618056 | 19.6777778 | 0 | R3IA-c-32 |
| red-tailed hawk | Raptor | spring | IA | Poweshiek | 3 | cropland | R3IA-c-32 | 2020-03-23T00:00:00Z | 29.9583333 | 39.6493056 | 0 | R3IA-c-32 |
| red-tailed hawk | Raptor | spring | IA | Poweshiek | 3 | cropland | R3IA-c-32 | 2020-03-23T00:00:00Z | 59.8493056 | NA | 0 | R3IA-c-32 |
| red-tailed hawk | Raptor | winter | IA | Poweshiek | 3 | cropland | R3IA-c-32 | 2020-12-07T00:00:00Z | 14.1145833 | 19.9083333 | 0 | R3IA-c-32 |
| great horned owl | Raptor | winter | IA | Poweshiek | 3 | cropland | R3IA-c-32 | 2020-12-07T00:00:00Z | 19.7506944 | 85.0201389 | 0 | R3IA-c-32 |
| red-tailed hawk | Raptor | winter | IA | Poweshiek | 3 | cropland | R3IA-c-32 | 2020-12-07T00:00:00Z | 4.0618056 | 6.7451389 | 0 | R3IA-c-32 |
| red-tailed hawk | Raptor | winter | IA | Poweshiek | 3 | cropland | R3IA-c-32 | 2020-12-07T00:00:00Z | 85.1701389 | NA | 0 | R3IA-c-32 |
| red-tailed hawk | Raptor | winter | IA | Poweshiek | 3 | cropland | R3IA-c-32 | 2020-12-07T00:00:00Z | 14.0847222 | 84.1694444 | 0 | R3IA-c-32 |
| great horned owl | Raptor | winter | IA | Poweshiek | 3 | cropland | R3IA-c-32 | 2020-12-07T00:00:00Z | 85.03125 | NA | 0 | R3IA-c-32 |
| red-tailed hawk | Raptor | winter | IA | Poweshiek | 3 | cropland | R3IA-c-32 | 2020-12-07T00:00:00Z | 15.1361111 | 42.8875 | 0 | R3IA-c-32 |
| red-tailed hawk | Raptor | winter | IA | Poweshiek | 3 | cropland | R3IA-c-32 | 2021-01-11T00:00:00Z | 60.2083333 | NA | 0 | R3IA-c-32 |
| red-tailed hawk | Raptor | winter | IA | Poweshiek | 3 | cropland | R3IA-c-32 | 2021-01-11T00:00:00Z | 14.05 | 50.1763889 | 0 | R3IA-c-32 |
| red-tailed hawk | Raptor | winter | IA | Poweshiek | 3 | cropland | R3IA-c-32 | 2021-01-11T00:00:00Z | 3.1347222 | 50.1777778 | 0 | R3IA-c-32 |
| red-tailed hawk | Raptor | winter | IA | Poweshiek | 3 | cropland | R3IA-c-32 | 2021-01-11T00:00:00Z | 60.0111111 | NA | 0 | R3IA-c-32 |
| red-tailed hawk | Raptor | winter | IA | Adair and Madison | 3 | cropland | R3IA-c-33 | 2020-12-07T00:00:00Z | 29.6993056 | NA | 0 | R3IA-c-33 |
| red-tailed hawk | Raptor | winter | IA | Adair and Madison | 3 | cropland | R3IA-c-33 | 2020-12-07T00:00:00Z | 20.9673611 | NA | 0 | R3IA-c-33 |
| red-tailed hawk | Raptor | winter | IA | Adair and Madison | 3 | cropland | R3IA-c-33 | 2020-12-07T00:00:00Z | 20.8069444 | NA | 0 | R3IA-c-33 |
| red-tailed hawk | Raptor | winter | IA | Adair and Madison | 3 | cropland | R3IA-c-33 | 2020-12-07T00:00:00Z | 3.7215278 | 20.8243056 | 0 | R3IA-c-33 |
| red-tailed hawk | Raptor | winter | IA | Adair and Madison | 3 | cropland | R3IA-c-33 | 2020-12-07T00:00:00Z | 20.8145833 | 39.9625 | 0 | R3IA-c-33 |
| red-tailed hawk | Raptor | winter | IA | Adair and Madison | 3 | cropland | R3IA-c-33 | 2021-02-22T00:00:00Z | 21.3027778 | 30.0368056 | 0 | R3IA-c-33 |
| red-tailed hawk | Raptor | winter | IA | Adair and Madison | 3 | cropland | R3IA-c-33 | 2021-02-22T00:00:00Z | 1.0145833 | 2.0694444 | 0 | R3IA-c-33 |
| red-tailed hawk | Raptor | winter | IA | Adair and Madison | 3 | cropland | R3IA-c-33 | 2021-02-22T00:00:00Z | 6.9888889 | 10.1430556 | 0 | R3IA-c-33 |
| red-tailed hawk | Raptor | winter | IA | Adair and Madison | 3 | cropland | R3IA-c-33 | 2021-02-26T00:00:00Z | 20.2527778 | 31.2638889 | 0 | R3IA-c-33 |
| Cooper's hawk | Raptor | spring | IA | Adair and Madison | 3 | cropland | R3IA-c-33 | 2021-03-31T00:00:00Z | 40.0763889 | 49.9243056 | 0 | R3IA-c-33 |
| red-tailed hawk | Raptor | spring | IA | Adair and Madison | 3 | cropland | R3IA-c-33 | 2021-04-21T00:00:00Z | 40.4020833 | NA | 0 | R3IA-c-33 |
| Cooper's hawk | Raptor | spring | IA | Adair and Madison | 3 | cropland | R3IA-c-33 | 2021-04-22T00:00:00Z | 2.0270833 | 3.3840278 | 0 | R3IA-c-33 |
| ring-necked pheasant | Game Bird | fall | IL | La Salle | 3 | cropland | R3IL-c-1 | 2020-08-03T00:00:00Z | 0.9986111 | 0.9986111 | 1 | R3IL-c-1 |
| ring-necked pheasant | Game Bird | fall | IL | La Salle | 3 | cropland | R3IL-c-1 | 2020-08-03T00:00:00Z | 1.1680556 | 2.00625 | 1 | R3IL-c-1 |
| ring-necked pheasant | Game Bird | fall | IL | La Salle | 3 | cropland | R3IL-c-1 | 2020-08-03T00:00:00Z | 7.0013889 | 10.0361111 | 1 | R3IL-c-1 |
| ring-necked pheasant | Game Bird | fall | IL | La Salle | 3 | cropland | R3IL-c-1 | 2020-08-03T00:00:00Z | 3.9111111 | 7.8104167 | 1 | R3IL-c-1 |
| ring-necked pheasant | Game Bird | fall | IL | La Salle | 3 | cropland | R3IL-c-1 | 2020-09-10T00:00:00Z | 1.0708333 | 2.9819444 | 1 | R3IL-c-1 |
| ring-necked pheasant | Game Bird | fall | IL | La Salle | 3 | cropland | R3IL-c-1 | 2020-09-10T00:00:00Z | 3.91875 | 4.9402778 | 1 | R3IL-c-1 |
| ring-necked pheasant | Game Bird | fall | IL | La Salle | 3 | cropland | R3IL-c-1 | 2020-09-10T00:00:00Z | 4.9625 | 7.9194444 | 1 | R3IL-c-1 |
| turkey vulture | Raptor | fall | IL | La Salle | 3 | cropland | R3IL-c-1 | 2020-08-18T00:00:00Z | 31 | 33 | 1 | R3IL-c-1 |
| ring-necked pheasant | Game Bird | spring | IL | La Salle | 3 | cropland | R3IL-c-1 | 2021-03-08T00:00:00Z | 1.3013889 | 2.1645833 | 1 | R3IL-c-1 |
| ring-necked pheasant | Game Bird | spring | IL | La Salle | 3 | cropland | R3IL-c-1 | 2021-03-08T00:00:00Z | 21.2416667 | 28.1138889 | 1 | R3IL-c-1 |
| ring-necked pheasant | Game Bird | spring | IL | La Salle | 3 | cropland | R3IL-c-1 | 2021-03-08T00:00:00Z | 8.1652778 | 9.2888889 | 1 | R3IL-c-1 |
| ring-necked pheasant | Game Bird | spring | IL | La Salle | 3 | cropland | R3IL-c-1 | 2021-03-08T00:00:00Z | 4.04375 | 8.1770833 | 1 | R3IL-c-1 |
| ring-necked pheasant | Game Bird | spring | IL | La Salle | 3 | cropland | R3IL-c-1 | 2021-03-08T00:00:00Z | 9.15 | 14.0236111 | 1 | R3IL-c-1 |
| red-tailed hawk | Raptor | spring | IL | La Salle | 3 | cropland | R3IL-c-1 | 2021-03-08T00:00:00Z | 8.8527778 | 13.9458333 | 1 | R3IL-c-1 |
| ring-necked pheasant | Game Bird | spring | IL | La Salle | 3 | cropland | R3IL-c-1 | 2021-04-12T00:00:00Z | 10.1277778 | 14.13125 | 1 | R3IL-c-1 |
| red-tailed hawk | Raptor | spring | IL | La Salle | 3 | cropland | R3IL-c-1 | 2021-03-22T00:00:00Z | 30.9166667 | 57.95 | 1 | R3IL-c-1 |
| ring-necked pheasant | Game Bird | spring | IL | La Salle | 3 | cropland | R3IL-c-1 | 2021-05-04T00:00:00Z | 2.1694444 | 3.18125 | 1 | R3IL-c-1 |
| ring-necked pheasant | Game Bird | spring | IL | La Salle | 3 | cropland | R3IL-c-1 | 2021-05-04T00:00:00Z | 9.2736111 | 15.0277778 | 1 | R3IL-c-1 |
| red-tailed hawk | Raptor | winter | IA | Palo Alto | 3 | cropland | R3IA-c-34 | 2020-12-07T00:00:00Z | 3.9854167 | 7.0569444 | 0 | R3IA-c-34 |
| red-tailed hawk | Raptor | winter | IA | Palo Alto | 3 | cropland | R3IA-c-34 | 2020-12-07T00:00:00Z | 30.1555556 | 49.9409722 | 0 | R3IA-c-34 |
| red-tailed hawk | Raptor | winter | IA | Palo Alto | 3 | cropland | R3IA-c-34 | 2020-12-07T00:00:00Z | 29.9097222 | 49.9381944 | 0 | R3IA-c-34 |
| red-tailed hawk | Raptor | winter | IA | Palo Alto | 3 | cropland | R3IA-c-34 | 2020-12-07T00:00:00Z | 3.9659722 | 6.975 | 0 | R3IA-c-34 |
| red-tailed hawk | Raptor | winter | IA | Palo Alto | 3 | cropland | R3IA-c-34 | 2020-12-07T00:00:00Z | 3.9611111 | 6.9631944 | 0 | R3IA-c-34 |
| red-tailed hawk | Raptor | winter | IA | Palo Alto | 3 | cropland | R3IA-c-34 | 2021-02-22T00:00:00Z | 14.0638889 | 20.0534722 | 0 | R3IA-c-34 |
| red-tailed hawk | Raptor | winter | IA | Palo Alto | 3 | cropland | R3IA-c-34 | 2021-02-22T00:00:00Z | 0.9923611 | 2.2208333 | 0 | R3IA-c-34 |
| red-tailed hawk | Raptor | winter | IA | Palo Alto | 3 | cropland | R3IA-c-34 | 2021-02-22T00:00:00Z | 60.3256944 | NA | 0 | R3IA-c-34 |
| red-tailed hawk | Raptor | winter | IA | Palo Alto | 3 | cropland | R3IA-c-34 | 2021-02-22T00:00:00Z | 3.9354167 | 13.9583333 | 0 | R3IA-c-34 |
| red-tailed hawk | Raptor | winter | IA | Palo Alto | 3 | cropland | R3IA-c-34 | 2021-02-22T00:00:00Z | 9.8979167 | 13.93125 | 0 | R3IA-c-34 |
| red-tailed hawk | Raptor | spring | IA | Mahaska | 3 | cropland | R3IA-c-35 | 2018-03-19T00:00:00Z | 20.8736111 | 31.2388889 | 0 | R3IA-c-35 |
| red-tailed hawk | Raptor | spring | IA | Mahaska | 3 | cropland | R3IA-c-35 | 2018-03-19T00:00:00Z | 20.8618056 | 31.2013889 | 0 | R3IA-c-35 |
| red-tailed hawk | Raptor | spring | IA | Mahaska | 3 | cropland | R3IA-c-35 | 2018-03-19T00:00:00Z | 31.2 | 41.0534722 | 0 | R3IA-c-35 |
| red-tailed hawk | Raptor | spring | IA | Mahaska | 3 | cropland | R3IA-c-35 | 2018-03-19T00:00:00Z | 31.1625 | 41.03125 | 0 | R3IA-c-35 |
| red-tailed hawk | Raptor | spring | IA | Mahaska | 3 | cropland | R3IA-c-35 | 2018-03-19T00:00:00Z | 3.9888889 | 7.1416667 | 0 | R3IA-c-35 |
| red-tailed hawk | Raptor | spring | IA | Mahaska | 3 | cropland | R3IA-c-35 | 2018-03-19T00:00:00Z | 31.2631944 | 40.9680556 | 0 | R3IA-c-35 |
| red-tailed hawk | Raptor | spring | IA | Mahaska | 3 | cropland | R3IA-c-35 | 2018-03-19T00:00:00Z | 31.2215278 | 40.9506944 | 0 | R3IA-c-35 |
| red-tailed hawk | Raptor | spring | IA | Mahaska | 3 | cropland | R3IA-c-35 | 2018-03-19T00:00:00Z | 21.1638889 | 31.1597222 | 0 | R3IA-c-35 |
| red-tailed hawk | Raptor | spring | IA | Mahaska | 3 | cropland | R3IA-c-35 | 2018-03-26T00:00:00Z | 30.9631944 | 41.0423611 | 0 | R3IA-c-35 |
| short-eared owl | Raptor | spring | IA | Mahaska | 3 | cropland | R3IA-c-35 | 2018-03-26T00:00:00Z | 0.5972222 | 2.59375 | 0 | R3IA-c-35 |
| red-tailed hawk | Raptor | spring | IA | Mahaska | 3 | cropland | R3IA-c-35 | 2018-05-07T00:00:00Z | 49.8659722 | NA | 0 | R3IA-c-35 |
| red-tailed hawk | Raptor | winter | IA | Mahaska | 3 | cropland | R3IA-c-35 | 2018-11-20T00:00:00Z | 29.8701389 | 39.9798611 | 0 | R3IA-c-35 |
| red-tailed hawk | Raptor | winter | IA | Mahaska | 3 | cropland | R3IA-c-35 | 2018-11-20T00:00:00Z | 20.0958333 | 29.8625 | 0 | R3IA-c-35 |
| red-tailed hawk | Raptor | winter | IA | Mahaska | 3 | cropland | R3IA-c-35 | 2018-11-20T00:00:00Z | 76.9868056 | NA | 0 | R3IA-c-35 |
| red-tailed hawk | Raptor | winter | IA | Mahaska | 3 | cropland | R3IA-c-35 | 2018-11-20T00:00:00Z | 50.03125 | 59.7854167 | 0 | R3IA-c-35 |
| red-tailed hawk | Raptor | winter | IA | Mahaska | 3 | cropland | R3IA-c-35 | 2018-11-20T00:00:00Z | 39.8875 | 50.025 | 0 | R3IA-c-35 |
| red-tailed hawk | Raptor | winter | IA | Mahaska | 3 | cropland | R3IA-c-35 | 2018-12-10T00:00:00Z | 20.0875 | 30.0715278 | 0 | R3IA-c-35 |
| red-tailed hawk | Raptor | winter | IA | Mahaska | 3 | cropland | R3IA-c-35 | 2018-12-10T00:00:00Z | 20.0701389 | 30.0666667 | 0 | R3IA-c-35 |
| red-tailed hawk | Raptor | winter | IA | Mahaska | 3 | cropland | R3IA-c-35 | 2018-12-10T00:00:00Z | 59.8777778 | NA | 0 | R3IA-c-35 |
| red-tailed hawk | Raptor | winter | IA | Mahaska | 3 | cropland | R3IA-c-35 | 2018-12-10T00:00:00Z | 30.0548611 | 43.0541667 | 0 | R3IA-c-35 |
| red-tailed hawk | Raptor | winter | IA | Mahaska | 3 | cropland | R3IA-c-35 | 2018-12-10T00:00:00Z | 20.0451389 | 30.0645833 | 0 | R3IA-c-35 |
| red-tailed hawk | Raptor | winter | IA | Mahaska | 3 | cropland | R3IA-c-35 | 2019-01-08T00:00:00Z | 49.9881944 | 64.0618056 | 0 | R3IA-c-35 |
| red-tailed hawk | Raptor | winter | IA | Mahaska | 3 | cropland | R3IA-c-35 | 2019-01-08T00:00:00Z | 3.2013889 | 64.0888889 | 0 | R3IA-c-35 |
| red-tailed hawk | Raptor | winter | IA | Mahaska | 3 | cropland | R3IA-c-35 | 2019-01-08T00:00:00Z | 63.9819444 | NA | 0 | R3IA-c-35 |
| red-tailed hawk | Raptor | winter | IA | Mahaska | 3 | cropland | R3IA-c-35 | 2019-01-08T00:00:00Z | 29.8166667 | 63.94375 | 0 | R3IA-c-35 |
| red-tailed hawk | Raptor | winter | IA | Mahaska | 3 | cropland | R3IA-c-35 | 2019-01-08T00:00:00Z | 64.1756944 | NA | 0 | R3IA-c-35 |
| red-tailed hawk | Raptor | spring | IA | Mahaska | 3 | cropland | R3IA-c-36 | 2020-03-16T00:00:00Z | 59.8076389 | NA | 0 | R3IA-c-36 |
| red-tailed hawk | Raptor | spring | IA | Mahaska | 3 | cropland | R3IA-c-36 | 2020-03-16T00:00:00Z | 59.7743056 | NA | 0 | R3IA-c-36 |
| red-tailed hawk | Raptor | spring | IA | Mahaska | 3 | cropland | R3IA-c-36 | 2020-03-16T00:00:00Z | 39.7590278 | 49.8805556 | 0 | R3IA-c-36 |
| red-tailed hawk | Raptor | spring | IA | Mahaska | 3 | cropland | R3IA-c-36 | 2020-03-16T00:00:00Z | 59.6875 | NA | 0 | R3IA-c-36 |
| red-tailed hawk | Raptor | spring | IA | Mahaska | 3 | cropland | R3IA-c-36 | 2020-03-16T00:00:00Z | 59.6354167 | NA | 0 | R3IA-c-36 |
| red-tailed hawk | Raptor | spring | IA | Mahaska | 3 | cropland | R3IA-c-36 | 2020-03-23T00:00:00Z | 62.7638889 | NA | 0 | R3IA-c-36 |
| red-tailed hawk | Raptor | spring | IA | Mahaska | 3 | cropland | R3IA-c-36 | 2020-03-23T00:00:00Z | 62.7576389 | NA | 0 | R3IA-c-36 |
| red-tailed hawk | Raptor | spring | IA | Mahaska | 3 | cropland | R3IA-c-36 | 2020-03-23T00:00:00Z | 62.7409722 | NA | 0 | R3IA-c-36 |
| red-tailed hawk | Raptor | spring | IA | Mahaska | 3 | cropland | R3IA-c-36 | 2020-03-23T00:00:00Z | 62.7208333 | NA | 0 | R3IA-c-36 |
| red-tailed hawk | Raptor | spring | IA | Mahaska | 3 | cropland | R3IA-c-36 | 2020-03-23T00:00:00Z | 62.7201389 | NA | 0 | R3IA-c-36 |
| red-tailed hawk | Raptor | winter | IA | Mahaska | 3 | cropland | R3IA-c-36 | 2020-12-14T00:00:00Z | 3.9604167 | 6.7215278 | 0 | R3IA-c-36 |
| red-tailed hawk | Raptor | winter | IA | Mahaska | 3 | cropland | R3IA-c-36 | 2020-12-14T00:00:00Z | 3.9604167 | 6.7298611 | 0 | R3IA-c-36 |
| red-tailed hawk | Raptor | winter | IA | Mahaska | 3 | cropland | R3IA-c-36 | 2020-12-14T00:00:00Z | 6.8194444 | 9.8902778 | 0 | R3IA-c-36 |
| red-tailed hawk | Raptor | winter | IA | Mahaska | 3 | cropland | R3IA-c-36 | 2020-12-14T00:00:00Z | 14.2013889 | 30.175 | 0 | R3IA-c-36 |
| red-tailed hawk | Raptor | winter | IA | Mahaska | 3 | cropland | R3IA-c-36 | 2020-12-14T00:00:00Z | 81.1451389 | NA | 0 | R3IA-c-36 |
| red-tailed hawk | Raptor | spring | MN | Stearns | 3 | cropland | R3MN-c-1 | 2017-03-31T00:00:00Z | 25 | NA | 0 | R3MN-c-1 |
| rough-legged hawk | Raptor | spring | MN | Stearns | 3 | cropland | R3MN-c-1 | 2017-03-31T00:00:00Z | 25 | NA | 0 | R3MN-c-1 |
| Cooper's hawk | Raptor | spring | MN | Stearns | 3 | cropland | R3MN-c-1 | 2017-03-31T00:00:00Z | 30 | 39 | 0 | R3MN-c-1 |
| red-tailed hawk | Raptor | spring | MN | Stearns | 3 | cropland | R3MN-c-1 | 2017-03-31T00:00:00Z | 4 | 7 | 0 | R3MN-c-1 |
| Cooper's hawk | Raptor | spring | MN | Stearns | 3 | cropland | R3MN-c-1 | 2017-03-31T00:00:00Z | 39 | NA | 0 | R3MN-c-1 |
| broad-winged hawk | Raptor | summer | MN | Stearns | 3 | cropland | R3MN-c-1 | 2017-06-20T00:00:00Z | 40 | 50 | 0 | R3MN-c-1 |
| Cooper's hawk | Raptor | summer | MN | Stearns | 3 | cropland | R3MN-c-1 | 2017-06-20T00:00:00Z | 40 | 50 | 0 | R3MN-c-1 |
| Cooper's hawk | Raptor | summer | MN | Stearns | 3 | cropland | R3MN-c-1 | 2017-08-28T00:00:00Z | 4 | 7 | 0 | R3MN-c-1 |
| Cooper's hawk | Raptor | summer | MN | Stearns | 3 | cropland | R3MN-c-1 | 2017-06-20T00:00:00Z | 90 | 100 | 0 | R3MN-c-1 |
| broad-winged hawk | Raptor | summer | MN | Stearns | 3 | cropland | R3MN-c-1 | 2017-06-20T00:00:00Z | 120 | 132 | 0 | R3MN-c-1 |
| Cooper's hawk | Raptor | summer | MN | Stearns | 3 | cropland | R3MN-c-1 | 2017-08-28T00:00:00Z | 50 | 57 | 0 | R3MN-c-1 |
| Cooper's hawk | Raptor | fall | MN | Stearns | 3 | cropland | R3MN-c-1 | 2017-10-23T00:00:00Z | 4 | 7 | 0 | R3MN-c-1 |
| Cooper's hawk | Raptor | fall | MN | Stearns | 3 | cropland | R3MN-c-1 | 2017-10-23T00:00:00Z | 2 | 3 | 0 | R3MN-c-1 |
| red-tailed hawk | Raptor | fall | MN | Stearns | 3 | cropland | R3MN-c-1 | 2017-10-23T00:00:00Z | 7 | 8 | 0 | R3MN-c-1 |
| Cooper's hawk | Raptor | fall | MN | Stearns | 3 | cropland | R3MN-c-1 | 2017-10-23T00:00:00Z | 7 | 10 | 0 | R3MN-c-1 |
| Cooper's hawk | Raptor | summer | MN | Stearns | 3 | cropland | R3MN-c-1 | 2017-08-11T00:00:00Z | 80 | 88 | 0 | R3MN-c-1 |
| Cooper's hawk | Raptor | summer | MN | Stearns | 3 | cropland | R3MN-c-1 | 2017-08-28T00:00:00Z | 203 | NA | 0 | R3MN-c-1 |
| red-tailed hawk | Raptor | fall | MN | Stearns | 3 | cropland | R3MN-c-1 | 2017-10-23T00:00:00Z | 147 | NA | 0 | R3MN-c-1 |
| peregrine falcon | Raptor | fall | MN | Stearns | 3 | cropland | R3MN-c-1 | 2017-10-23T00:00:00Z | 147 | NA | 0 | R3MN-c-1 |
| prairie falcon | Raptor | fall | MN | Stearns | 3 | cropland | R3MN-c-1 | 2017-10-23T00:00:00Z | 147 | NA | 0 | R3MN-c-1 |
| red-tailed hawk | Raptor | spring | MN | Stearns | 3 | cropland | R3MN-c-2 | 2018-04-09T00:00:00Z | 42 | 52 | 0 | R3MN-c-2 |
| red-tailed hawk | Raptor | summer | MN | Stearns | 3 | cropland | R3MN-c-2 | 2018-06-18T00:00:00Z | 14 | 21 | 0 | R3MN-c-2 |
| red-tailed hawk | Raptor | summer | MN | Stearns | 3 | cropland | R3MN-c-2 | 2018-06-18T00:00:00Z | 21 | 28 | 0 | R3MN-c-2 |
| red-tailed hawk | Raptor | summer | MN | Stearns | 3 | cropland | R3MN-c-2 | 2018-06-18T00:00:00Z | 39 | 42 | 0 | R3MN-c-2 |
| red-tailed hawk | Raptor | spring | MN | Stearns | 3 | cropland | R3MN-c-2 | 2018-04-09T00:00:00Z | 122 | NA | 0 | R3MN-c-2 |
| red-tailed hawk | Raptor | spring | MN | Stearns | 3 | cropland | R3MN-c-2 | 2018-04-09T00:00:00Z | 122 | NA | 0 | R3MN-c-2 |
| red-tailed hawk | Raptor | summer | MN | Stearns | 3 | cropland | R3MN-c-2 | 2018-06-18T00:00:00Z | 60 | 63 | 0 | R3MN-c-2 |
| red-tailed hawk | Raptor | summer | MN | Stearns | 3 | cropland | R3MN-c-2 | 2018-06-18T00:00:00Z | 49 | 56 | 0 | R3MN-c-2 |
| red-tailed hawk | Raptor | summer | MN | Stearns | 3 | cropland | R3MN-c-2 | 2018-06-18T00:00:00Z | 91 | 98 | 0 | R3MN-c-2 |
| red-tailed hawk | Raptor | fall | MN | Stearns | 3 | cropland | R3MN-c-2 | 2018-08-28T00:00:00Z | 41 | 51 | 0 | R3MN-c-2 |
| red-tailed hawk | Raptor | fall | MN | Stearns | 3 | cropland | R3MN-c-2 | 2018-08-28T00:00:00Z | 65 | NA | 0 | R3MN-c-2 |
| red-tailed hawk | Raptor | fall | MN | Stearns | 3 | cropland | R3MN-c-2 | 2018-08-28T00:00:00Z | 65 | NA | 0 | R3MN-c-2 |
| red-tailed hawk | Raptor | fall | MN | Stearns | 3 | cropland | R3MN-c-2 | 2018-08-28T00:00:00Z | 65 | NA | 0 | R3MN-c-2 |
| red-tailed hawk | Raptor | fall | MN | Stearns | 3 | cropland | R3MN-c-2 | 2018-08-28T00:00:00Z | 65 | NA | 0 | R3MN-c-2 |
| red-tailed hawk | Raptor | fall | MN | Stearns | 3 | cropland | R3MN-c-2 | 2018-08-28T00:00:00Z | 65 | NA | 0 | R3MN-c-2 |
| red-tailed hawk | Raptor | summer | CA | Alameda | 8 | shrub/scrub | R8CA-ss-8 | 2006-08-23T00:00:00Z | 62 | NA | 0 | R8CA-ss-8 |
| mallard | Game Bird | summer | CA | Alameda | 8 | shrub/scrub | R8CA-ss-8 | 2006-08-27T00:00:00Z | 0 | 1 | 0 | R8CA-ss-8 |
| unidentified duck | Game Bird | summer | CA | Alameda | 8 | shrub/scrub | R8CA-ss-8 | 2006-08-27T00:00:00Z | 64 | NA | 0 | R8CA-ss-8 |
| unidentified duck | Game Bird | summer | CA | Alameda | 8 | shrub/scrub | R8CA-ss-8 | 2006-08-27T00:00:00Z | 64 | NA | 0 | R8CA-ss-8 |
| mallard | Game Bird | summer | CA | Alameda | 8 | shrub/scrub | R8CA-ss-8 | 2006-08-27T00:00:00Z | 2 | 3 | 0 | R8CA-ss-8 |
| unidentified duck | Game Bird | summer | CA | Alameda | 8 | shrub/scrub | R8CA-ss-8 | 2006-08-27T00:00:00Z | 64 | NA | 0 | R8CA-ss-8 |
| unidentified duck | Game Bird | summer | CA | Alameda | 8 | shrub/scrub | R8CA-ss-8 | 2006-08-27T00:00:00Z | 64 | NA | 0 | R8CA-ss-8 |
| unidentified duck | Game Bird | summer | CA | Alameda | 8 | shrub/scrub | R8CA-ss-8 | 2006-08-27T00:00:00Z | 3 | 7 | 0 | R8CA-ss-8 |
| mallard | Game Bird | summer | CA | Alameda | 8 | shrub/scrub | R8CA-ss-8 | 2006-08-27T00:00:00Z | 36 | 43 | 0 | R8CA-ss-8 |
| mallard | Game Bird | summer | CA | Alameda | 8 | shrub/scrub | R8CA-ss-8 | 2006-08-27T00:00:00Z | 43 | 50 | 0 | R8CA-ss-8 |
| mallard | Game Bird | summer | CA | Alameda | 8 | shrub/scrub | R8CA-ss-8 | 2006-08-27T00:00:00Z | 64 | NA | 0 | R8CA-ss-8 |
| unidentified duck | Game Bird | summer | CA | Alameda | 8 | shrub/scrub | R8CA-ss-8 | 2006-08-27T00:00:00Z | 64 | NA | 0 | R8CA-ss-8 |
| mallard | Game Bird | summer | CA | Alameda | 8 | shrub/scrub | R8CA-ss-8 | 2006-08-27T00:00:00Z | 64 | NA | 0 | R8CA-ss-8 |
| unidentified duck | Game Bird | summer | CA | Alameda | 8 | shrub/scrub | R8CA-ss-8 | 2006-08-27T00:00:00Z | 7 | 10 | 0 | R8CA-ss-8 |
| unidentified duck | Game Bird | summer | CA | Alameda | 8 | shrub/scrub | R8CA-ss-8 | 2006-08-27T00:00:00Z | 2 | 3 | 0 | R8CA-ss-8 |
| unidentified duck | Game Bird | summer | CA | Alameda | 8 | shrub/scrub | R8CA-ss-8 | 2006-08-27T00:00:00Z | 43 | 50 | 0 | R8CA-ss-8 |
| unidentified duck | Game Bird | summer | CA | Alameda | 8 | shrub/scrub | R8CA-ss-8 | 2006-08-27T00:00:00Z | 64 | NA | 0 | R8CA-ss-8 |
| unidentified duck | Game Bird | summer | CA | Alameda | 8 | shrub/scrub | R8CA-ss-8 | 2006-08-27T00:00:00Z | 64 | NA | 0 | R8CA-ss-8 |
| ring-necked pheasant | Game Bird | summer | CA | Alameda | 8 | shrub/scrub | R8CA-ss-8 | 2006-09-05T00:00:00Z | 63 | NA | 0 | R8CA-ss-8 |
| ring-necked pheasant | Game Bird | summer | CA | Alameda | 8 | shrub/scrub | R8CA-ss-8 | 2006-09-06T00:00:00Z | 63 | NA | 0 | R8CA-ss-8 |
| ring-necked pheasant | Game Bird | summer | CA | Alameda | 8 | shrub/scrub | R8CA-ss-8 | 2006-09-07T00:00:00Z | 63 | NA | 0 | R8CA-ss-8 |
| ring-necked pheasant | Game Bird | summer | CA | Alameda | 8 | shrub/scrub | R8CA-ss-8 | 2006-09-08T00:00:00Z | 2 | 3 | 0 | R8CA-ss-8 |
| ring-necked pheasant | Game Bird | summer | CA | Alameda | 8 | shrub/scrub | R8CA-ss-8 | 2006-09-09T00:00:00Z | 49 | 56 | 0 | R8CA-ss-8 |
| barn owl | Raptor | winter | OR/WA | Umatilla, OR and Walla Walla, WA | 1 | cropland | R1OR/WA-c-2 | 2001-12-09T00:00:00Z | 14 | 20 | 1 | R1OR/WA-c-2 |
| barn owl | Raptor | winter | OR/WA | Umatilla, OR and Walla Walla, WA | 1 | cropland | R1OR/WA-c-2 | 2001-12-09T00:00:00Z | 4 | 7 | 1 | R1OR/WA-c-2 |
| barn owl | Raptor | winter | OR/WA | Umatilla, OR and Walla Walla, WA | 1 | cropland | R1OR/WA-c-2 | 2003-01-10T00:00:00Z | 30 | NA | 1 | R1OR/WA-c-2 |
| barn owl | Raptor | winter | OR/WA | Umatilla, OR and Walla Walla, WA | 1 | cropland | R1OR/WA-c-2 | 2003-01-10T00:00:00Z | 20 | 30 | 1 | R1OR/WA-c-2 |
| barn owl | Raptor | winter | OR/WA | Umatilla, OR and Walla Walla, WA | 1 | cropland | R1OR/WA-c-2 | 2003-11-04T00:00:00Z | 7 | 10 | 1 | R1OR/WA-c-2 |
| barn owl | Raptor | fall | OR/WA | Umatilla, OR and Walla Walla, WA | 1 | cropland | R1OR/WA-c-2 | 2002-10-06T00:00:00Z | 30 | 40 | 1 | R1OR/WA-c-2 |
| ferruginous hawk | Raptor | winter | OR/WA | Umatilla, OR and Walla Walla, WA | 1 | cropland | R1OR/WA-c-2 | 2003-01-10T00:00:00Z | 10 | 14 | 1 | R1OR/WA-c-2 |
| great horned owl | Raptor | fall | OR/WA | Umatilla, OR and Walla Walla, WA | 1 | cropland | R1OR/WA-c-2 | 2002-10-06T00:00:00Z | 40 | NA | 1 | R1OR/WA-c-2 |
| great horned owl | Raptor | spring | OR/WA | Umatilla, OR and Walla Walla, WA | 1 | cropland | R1OR/WA-c-2 | 2003-03-10T00:00:00Z | 40 | NA | 1 | R1OR/WA-c-2 |
| great horned owl | Raptor | winter | OR/WA | Umatilla, OR and Walla Walla, WA | 1 | cropland | R1OR/WA-c-2 | 2001-12-09T00:00:00Z | 40 | NA | 1 | R1OR/WA-c-2 |
| great horned owl | Raptor | winter | OR/WA | Umatilla, OR and Walla Walla, WA | 1 | cropland | R1OR/WA-c-2 | 2003-01-10T00:00:00Z | 30 | NA | 1 | R1OR/WA-c-2 |
| red-tailed hawk | Raptor | fall | OR/WA | Umatilla, OR and Walla Walla, WA | 1 | cropland | R1OR/WA-c-2 | 2002-10-06T00:00:00Z | 40 | NA | 1 | R1OR/WA-c-2 |
| red-tailed hawk | Raptor | spring | OR/WA | Umatilla, OR and Walla Walla, WA | 1 | cropland | R1OR/WA-c-2 | 2003-04-01T00:00:00Z | 40 | NA | 1 | R1OR/WA-c-2 |
| red-tailed hawk | Raptor | summer | OR/WA | Umatilla, OR and Walla Walla, WA | 1 | cropland | R1OR/WA-c-2 | 2002-07-22T00:00:00Z | 40 | NA | 1 | R1OR/WA-c-2 |
| mallard | Game Bird | fall | OR/WA | Umatilla, OR and Walla Walla, WA | 1 | cropland | R1OR/WA-c-2 | 2001-09-25T00:00:00Z | 10 | 14 | 1 | R1OR/WA-c-2 |
| mallard | Game Bird | fall | OR/WA | Umatilla, OR and Walla Walla, WA | 1 | cropland | R1OR/WA-c-2 | 2001-09-26T00:00:00Z | 20 | 30 | 1 | R1OR/WA-c-2 |
| mallard | Game Bird | fall | OR/WA | Umatilla, OR and Walla Walla, WA | 1 | cropland | R1OR/WA-c-2 | 2001-09-25T00:00:00Z | 20 | 30 | 1 | R1OR/WA-c-2 |
| mallard | Game Bird | fall | OR/WA | Umatilla, OR and Walla Walla, WA | 1 | cropland | R1OR/WA-c-2 | 2001-09-26T00:00:00Z | 10 | 14 | 1 | R1OR/WA-c-2 |
| mallard | Game Bird | fall | OR/WA | Umatilla, OR and Walla Walla, WA | 1 | cropland | R1OR/WA-c-2 | 2001-09-25T00:00:00Z | 3 | 4 | 1 | R1OR/WA-c-2 |
| mallard | Game Bird | fall | OR/WA | Umatilla, OR and Walla Walla, WA | 1 | cropland | R1OR/WA-c-2 | 2001-09-26T00:00:00Z | 4 | 7 | 1 | R1OR/WA-c-2 |
| mallard | Game Bird | fall | OR/WA | Umatilla, OR and Walla Walla, WA | 1 | cropland | R1OR/WA-c-2 | 2001-09-25T00:00:00Z | 40 | NA | 1 | R1OR/WA-c-2 |
| mallard | Game Bird | fall | OR/WA | Umatilla, OR and Walla Walla, WA | 1 | cropland | R1OR/WA-c-2 | 2001-09-26T00:00:00Z | 4 | 7 | 1 | R1OR/WA-c-2 |
| mallard | Game Bird | fall | OR/WA | Umatilla, OR and Walla Walla, WA | 1 | cropland | R1OR/WA-c-2 | 2001-09-26T00:00:00Z | 20 | 30 | 1 | R1OR/WA-c-2 |
| mallard | Game Bird | fall | OR/WA | Umatilla, OR and Walla Walla, WA | 1 | cropland | R1OR/WA-c-2 | 2001-09-26T00:00:00Z | 10 | 14 | 1 | R1OR/WA-c-2 |
| mallard | Game Bird | fall | OR/WA | Umatilla, OR and Walla Walla, WA | 1 | cropland | R1OR/WA-c-2 | 2002-10-06T00:00:00Z | 40 | NA | 1 | R1OR/WA-c-2 |
| mallard | Game Bird | fall | OR/WA | Umatilla, OR and Walla Walla, WA | 1 | cropland | R1OR/WA-c-2 | 2002-10-14T00:00:00Z | 10 | 14 | 1 | R1OR/WA-c-2 |
| mallard | Game Bird | fall | OR/WA | Umatilla, OR and Walla Walla, WA | 1 | cropland | R1OR/WA-c-2 | 2002-10-14T00:00:00Z | 40 | NA | 1 | R1OR/WA-c-2 |
| mallard | Game Bird | fall | OR/WA | Umatilla, OR and Walla Walla, WA | 1 | cropland | R1OR/WA-c-2 | 2002-10-14T00:00:00Z | 40 | NA | 1 | R1OR/WA-c-2 |
| mallard | Game Bird | fall | OR/WA | Umatilla, OR and Walla Walla, WA | 1 | cropland | R1OR/WA-c-2 | 2002-10-14T00:00:00Z | 14 | 20 | 1 | R1OR/WA-c-2 |
| mallard | Game Bird | fall | OR/WA | Umatilla, OR and Walla Walla, WA | 1 | cropland | R1OR/WA-c-2 | 2002-10-14T00:00:00Z | 7 | 10 | 1 | R1OR/WA-c-2 |
| mallard | Game Bird | fall | OR/WA | Umatilla, OR and Walla Walla, WA | 1 | cropland | R1OR/WA-c-2 | 2002-10-14T00:00:00Z | 10 | 14 | 1 | R1OR/WA-c-2 |
| mallard | Game Bird | fall | OR/WA | Umatilla, OR and Walla Walla, WA | 1 | cropland | R1OR/WA-c-2 | 2002-10-14T00:00:00Z | 4 | 7 | 1 | R1OR/WA-c-2 |
| mallard | Game Bird | fall | OR/WA | Umatilla, OR and Walla Walla, WA | 1 | cropland | R1OR/WA-c-2 | 2002-10-14T00:00:00Z | 14 | 20 | 1 | R1OR/WA-c-2 |
| mallard | Game Bird | fall | OR/WA | Umatilla, OR and Walla Walla, WA | 1 | cropland | R1OR/WA-c-2 | 2002-10-14T00:00:00Z | 40 | NA | 1 | R1OR/WA-c-2 |
| mallard | Game Bird | fall | OR/WA | Umatilla, OR and Walla Walla, WA | 1 | cropland | R1OR/WA-c-2 | 2002-10-14T00:00:00Z | 30 | 40 | 1 | R1OR/WA-c-2 |
| mallard | Game Bird | spring | OR/WA | Umatilla, OR and Walla Walla, WA | 1 | cropland | R1OR/WA-c-2 | 2002-04-02T00:00:00Z | 20 | 30 | 1 | R1OR/WA-c-2 |
| mallard | Game Bird | spring | OR/WA | Umatilla, OR and Walla Walla, WA | 1 | cropland | R1OR/WA-c-2 | 2002-04-02T00:00:00Z | 40 | NA | 1 | R1OR/WA-c-2 |
| mallard | Game Bird | summer | OR/WA | Umatilla, OR and Walla Walla, WA | 1 | cropland | R1OR/WA-c-2 | 2003-06-26T00:00:00Z | 4 | 7 | 1 | R1OR/WA-c-2 |
| mallard | Game Bird | summer | OR/WA | Umatilla, OR and Walla Walla, WA | 1 | cropland | R1OR/WA-c-2 | 2003-06-26T00:00:00Z | 40 | NA | 1 | R1OR/WA-c-2 |
| mallard | Game Bird | summer | OR/WA | Umatilla, OR and Walla Walla, WA | 1 | cropland | R1OR/WA-c-2 | 2003-06-26T00:00:00Z | 2 | 3 | 1 | R1OR/WA-c-2 |
| mallard | Game Bird | summer | OR/WA | Umatilla, OR and Walla Walla, WA | 1 | cropland | R1OR/WA-c-2 | 2003-06-26T00:00:00Z | 40 | NA | 1 | R1OR/WA-c-2 |
| mallard | Game Bird | summer | OR/WA | Umatilla, OR and Walla Walla, WA | 1 | cropland | R1OR/WA-c-2 | 2002-07-08T00:00:00Z | 10 | 14 | 1 | R1OR/WA-c-2 |
| mallard | Game Bird | summer | OR/WA | Umatilla, OR and Walla Walla, WA | 1 | cropland | R1OR/WA-c-2 | 2002-07-08T00:00:00Z | 30 | 40 | 1 | R1OR/WA-c-2 |
| mallard | Game Bird | summer | OR/WA | Umatilla, OR and Walla Walla, WA | 1 | cropland | R1OR/WA-c-2 | 2002-07-08T00:00:00Z | 40 | NA | 1 | R1OR/WA-c-2 |
| mallard | Game Bird | summer | OR/WA | Umatilla, OR and Walla Walla, WA | 1 | cropland | R1OR/WA-c-2 | 2002-07-08T00:00:00Z | 0 | 1 | 1 | R1OR/WA-c-2 |
| mallard | Game Bird | summer | OR/WA | Umatilla, OR and Walla Walla, WA | 1 | cropland | R1OR/WA-c-2 | 2002-07-22T00:00:00Z | 40 | NA | 1 | R1OR/WA-c-2 |
| mallard | Game Bird | summer | OR/WA | Umatilla, OR and Walla Walla, WA | 1 | cropland | R1OR/WA-c-2 | 2002-07-22T00:00:00Z | 40 | NA | 1 | R1OR/WA-c-2 |
| mallard | Game Bird | winter | OR/WA | Umatilla, OR and Walla Walla, WA | 1 | cropland | R1OR/WA-c-2 | 2001-12-09T00:00:00Z | 0 | 1 | 1 | R1OR/WA-c-2 |
| mallard | Game Bird | winter | OR/WA | Umatilla, OR and Walla Walla, WA | 1 | cropland | R1OR/WA-c-2 | 2001-12-09T00:00:00Z | 4 | 7 | 1 | R1OR/WA-c-2 |
| mallard | Game Bird | winter | OR/WA | Umatilla, OR and Walla Walla, WA | 1 | cropland | R1OR/WA-c-2 | 2001-12-09T00:00:00Z | 40 | NA | 1 | R1OR/WA-c-2 |
| mallard | Game Bird | winter | OR/WA | Umatilla, OR and Walla Walla, WA | 1 | cropland | R1OR/WA-c-2 | 2002-02-02T00:00:00Z | 30 | 40 | 1 | R1OR/WA-c-2 |
| mallard | Game Bird | winter | OR/WA | Umatilla, OR and Walla Walla, WA | 1 | cropland | R1OR/WA-c-2 | 2002-02-02T00:00:00Z | 14 | 20 | 1 | R1OR/WA-c-2 |
| mallard | Game Bird | winter | OR/WA | Umatilla, OR and Walla Walla, WA | 1 | cropland | R1OR/WA-c-2 | 2002-02-02T00:00:00Z | 40 | NA | 1 | R1OR/WA-c-2 |
| mallard | Game Bird | winter | OR/WA | Umatilla, OR and Walla Walla, WA | 1 | cropland | R1OR/WA-c-2 | 2002-02-02T00:00:00Z | 40 | NA | 1 | R1OR/WA-c-2 |
| mallard | Game Bird | winter | OR/WA | Umatilla, OR and Walla Walla, WA | 1 | cropland | R1OR/WA-c-2 | 2002-02-02T00:00:00Z | 40 | NA | 1 | R1OR/WA-c-2 |
| mallard | Game Bird | winter | OR/WA | Umatilla, OR and Walla Walla, WA | 1 | cropland | R1OR/WA-c-2 | 2002-02-02T00:00:00Z | 1 | 2 | 1 | R1OR/WA-c-2 |
| mallard | Game Bird | winter | OR/WA | Umatilla, OR and Walla Walla, WA | 1 | cropland | R1OR/WA-c-2 | 2002-02-02T00:00:00Z | 40 | NA | 1 | R1OR/WA-c-2 |
| mallard | Game Bird | winter | OR/WA | Umatilla, OR and Walla Walla, WA | 1 | cropland | R1OR/WA-c-2 | 2002-02-02T00:00:00Z | 14 | 20 | 1 | R1OR/WA-c-2 |
| mallard | Game Bird | winter | OR/WA | Umatilla, OR and Walla Walla, WA | 1 | cropland | R1OR/WA-c-2 | 2002-02-02T00:00:00Z | 40 | NA | 1 | R1OR/WA-c-2 |
| mallard | Game Bird | winter | OR/WA | Umatilla, OR and Walla Walla, WA | 1 | cropland | R1OR/WA-c-2 | 2003-01-10T00:00:00Z | 30 | 30 | 1 | R1OR/WA-c-2 |
| mallard | Game Bird | winter | OR/WA | Umatilla, OR and Walla Walla, WA | 1 | cropland | R1OR/WA-c-2 | 2003-01-10T00:00:00Z | 0 | 1 | 1 | R1OR/WA-c-2 |
| mallard | Game Bird | winter | OR/WA | Umatilla, OR and Walla Walla, WA | 1 | cropland | R1OR/WA-c-2 | 2003-01-10T00:00:00Z | 0 | 1 | 1 | R1OR/WA-c-2 |
| mallard | Game Bird | winter | OR/WA | Umatilla, OR and Walla Walla, WA | 1 | cropland | R1OR/WA-c-2 | 2003-02-02T00:00:00Z | 2 | 3 | 1 | R1OR/WA-c-2 |
| mallard | Game Bird | winter | OR/WA | Umatilla, OR and Walla Walla, WA | 1 | cropland | R1OR/WA-c-2 | 2003-11-04T00:00:00Z | 2 | 3 | 1 | R1OR/WA-c-2 |
| mallard | Game Bird | fall | OR/WA | Umatilla, OR and Walla Walla, WA | 1 | cropland | R1OR/WA-c-2 | 2001-09-26T00:00:00Z | 20 | 30 | 1 | R1OR/WA-c-2 |
| mallard | Game Bird | fall | OR/WA | Umatilla, OR and Walla Walla, WA | 1 | cropland | R1OR/WA-c-2 | 2001-09-25T00:00:00Z | 7 | 10 | 1 | R1OR/WA-c-2 |
| mallard | Game Bird | fall | OR/WA | Umatilla, OR and Walla Walla, WA | 1 | cropland | R1OR/WA-c-2 | 2001-09-26T00:00:00Z | 20 | 30 | 1 | R1OR/WA-c-2 |
| mallard | Game Bird | fall | OR/WA | Umatilla, OR and Walla Walla, WA | 1 | cropland | R1OR/WA-c-2 | 2001-09-25T00:00:00Z | 4 | 7 | 1 | R1OR/WA-c-2 |
| mallard | Game Bird | fall | OR/WA | Umatilla, OR and Walla Walla, WA | 1 | cropland | R1OR/WA-c-2 | 2001-09-25T00:00:00Z | 10 | 14 | 1 | R1OR/WA-c-2 |
| mallard | Game Bird | fall | OR/WA | Umatilla, OR and Walla Walla, WA | 1 | cropland | R1OR/WA-c-2 | 2001-09-26T00:00:00Z | 4 | 7 | 1 | R1OR/WA-c-2 |
| mallard | Game Bird | fall | OR/WA | Umatilla, OR and Walla Walla, WA | 1 | cropland | R1OR/WA-c-2 | 2001-09-25T00:00:00Z | 14 | 20 | 1 | R1OR/WA-c-2 |
| mallard | Game Bird | fall | OR/WA | Umatilla, OR and Walla Walla, WA | 1 | cropland | R1OR/WA-c-2 | 2001-09-26T00:00:00Z | 4 | 7 | 1 | R1OR/WA-c-2 |
| mallard | Game Bird | fall | OR/WA | Umatilla, OR and Walla Walla, WA | 1 | cropland | R1OR/WA-c-2 | 2001-09-26T00:00:00Z | 0 | 1 | 1 | R1OR/WA-c-2 |
| mallard | Game Bird | fall | OR/WA | Umatilla, OR and Walla Walla, WA | 1 | cropland | R1OR/WA-c-2 | 2001-09-26T00:00:00Z | 10 | 14 | 1 | R1OR/WA-c-2 |
| mallard | Game Bird | summer | OR/WA | Umatilla, OR and Walla Walla, WA | 1 | cropland | R1OR/WA-c-2 | 2003-06-26T00:00:00Z | 14 | 20 | 1 | R1OR/WA-c-2 |
| mallard | Game Bird | winter | OR/WA | Umatilla, OR and Walla Walla, WA | 1 | cropland | R1OR/WA-c-2 | 2001-12-09T00:00:00Z | 40 | NA | 1 | R1OR/WA-c-2 |
| mallard | Game Bird | winter | OR/WA | Umatilla, OR and Walla Walla, WA | 1 | cropland | R1OR/WA-c-2 | 2002-02-02T00:00:00Z | 40 | NA | 1 | R1OR/WA-c-2 |
| mallard | Game Bird | winter | OR/WA | Umatilla, OR and Walla Walla, WA | 1 | cropland | R1OR/WA-c-2 | 2002-02-02T00:00:00Z | 14 | 20 | 1 | R1OR/WA-c-2 |
| mallard | Game Bird | winter | OR/WA | Umatilla, OR and Walla Walla, WA | 1 | cropland | R1OR/WA-c-2 | 2003-11-04T00:00:00Z | 2 | 3 | 1 | R1OR/WA-c-2 |
| mallard | Game Bird | winter | OR/WA | Umatilla, OR and Walla Walla, WA | 1 | cropland | R1OR/WA-c-2 | 2003-11-04T00:00:00Z | 4 | 7 | 1 | R1OR/WA-c-2 |
| ring-necked pheasant | Game Bird | fall | OR/WA | Umatilla, OR and Walla Walla, WA | 1 | cropland | R1OR/WA-c-2 | 2003-08-12T00:00:00Z | 40 | NA | 1 | R1OR/WA-c-2 |
| ring-necked pheasant | Game Bird | fall | OR/WA | Umatilla, OR and Walla Walla, WA | 1 | cropland | R1OR/WA-c-2 | 2003-09-16T00:00:00Z | 30 | NA | 1 | R1OR/WA-c-2 |
| ring-necked pheasant | Game Bird | fall | OR/WA | Umatilla, OR and Walla Walla, WA | 1 | cropland | R1OR/WA-c-2 | 2003-09-16T00:00:00Z | 30 | NA | 1 | R1OR/WA-c-2 |
| ring-necked pheasant | Game Bird | fall | OR/WA | Umatilla, OR and Walla Walla, WA | 1 | cropland | R1OR/WA-c-2 | 2003-09-16T00:00:00Z | 20 | 30 | 1 | R1OR/WA-c-2 |
| ring-necked pheasant | Game Bird | fall | OR/WA | Umatilla, OR and Walla Walla, WA | 1 | cropland | R1OR/WA-c-2 | 2003-09-16T00:00:00Z | 30 | NA | 1 | R1OR/WA-c-2 |
| ring-necked pheasant | Game Bird | fall | OR/WA | Umatilla, OR and Walla Walla, WA | 1 | cropland | R1OR/WA-c-2 | 2003-09-16T00:00:00Z | 10 | 14 | 1 | R1OR/WA-c-2 |
| ring-necked pheasant | Game Bird | winter | OR/WA | Umatilla, OR and Walla Walla, WA | 1 | cropland | R1OR/WA-c-2 | 2003-11-04T00:00:00Z | 10 | 14 | 1 | R1OR/WA-c-2 |
| ring-necked pheasant | Game Bird | winter | OR/WA | Umatilla, OR and Walla Walla, WA | 1 | cropland | R1OR/WA-c-2 | 2003-11-04T00:00:00Z | 30 | 40 | 1 | R1OR/WA-c-2 |
| ring-necked pheasant | Game Bird | fall | OR/WA | Umatilla, OR and Walla Walla, WA | 1 | cropland | R1OR/WA-c-2 | 2003-09-16T00:00:00Z | 14 | 20 | 1 | R1OR/WA-c-2 |
| ring-necked pheasant | Game Bird | fall | OR/WA | Umatilla, OR and Walla Walla, WA | 1 | cropland | R1OR/WA-c-2 | 2003-09-16T00:00:00Z | 20 | 30 | 1 | R1OR/WA-c-2 |
| ring-necked pheasant | Game Bird | fall | OR/WA | Umatilla, OR and Walla Walla, WA | 1 | cropland | R1OR/WA-c-2 | 2003-09-16T00:00:00Z | 20 | 30 | 1 | R1OR/WA-c-2 |
| ring-necked pheasant | Game Bird | fall | OR/WA | Umatilla, OR and Walla Walla, WA | 1 | cropland | R1OR/WA-c-2 | 2003-09-16T00:00:00Z | 4 | 7 | 1 | R1OR/WA-c-2 |
| ring-necked pheasant | Game Bird | winter | OR/WA | Umatilla, OR and Walla Walla, WA | 1 | cropland | R1OR/WA-c-2 | 2003-11-04T00:00:00Z | 4 | 7 | 1 | R1OR/WA-c-2 |
| ring-necked pheasant | Game Bird | winter | OR/WA | Umatilla, OR and Walla Walla, WA | 1 | cropland | R1OR/WA-c-2 | 2003-11-04T00:00:00Z | 20 | 30 | 1 | R1OR/WA-c-2 |
| mallard | Game Bird | fall | NV | White Pine | 8 | shrub/scrub | R8NV-ss-1 | 2012-09-30T00:00:00Z | 20 | 30 | 0 | R8NV-ss-1 |
| ring-necked pheasant | Game Bird | fall | NV | White Pine | 8 | shrub/scrub | R8NV-ss-1 | 2012-09-30T00:00:00Z | 4 | 7 | 0 | R8NV-ss-1 |
| ring-necked pheasant | Game Bird | fall | NV | White Pine | 8 | shrub/scrub | R8NV-ss-1 | 2012-09-30T00:00:00Z | 10 | 14 | 0 | R8NV-ss-1 |
| ring-necked pheasant | Game Bird | fall | NV | White Pine | 8 | shrub/scrub | R8NV-ss-1 | 2012-09-30T00:00:00Z | 4 | 7 | 0 | R8NV-ss-1 |
| mallard | Game Bird | fall | NV | White Pine | 8 | shrub/scrub | R8NV-ss-1 | 2012-09-30T00:00:00Z | 3 | 4 | 0 | R8NV-ss-1 |
| ring-necked pheasant | Game Bird | fall | NV | White Pine | 8 | shrub/scrub | R8NV-ss-1 | 2012-09-30T00:00:00Z | 20 | 30 | 0 | R8NV-ss-1 |
| mallard | Game Bird | winter | NV | White Pine | 8 | shrub/scrub | R8NV-ss-1 | 2012-12-02T00:00:00Z | 10 | 14 | 0 | R8NV-ss-1 |
| ring-necked pheasant | Game Bird | winter | NV | White Pine | 8 | shrub/scrub | R8NV-ss-1 | 2012-12-02T00:00:00Z | 40 | NA | 0 | R8NV-ss-1 |
| ring-necked pheasant | Game Bird | winter | NV | White Pine | 8 | shrub/scrub | R8NV-ss-1 | 2012-12-02T00:00:00Z | 1 | 2 | 0 | R8NV-ss-1 |
| mallard | Game Bird | winter | NV | White Pine | 8 | shrub/scrub | R8NV-ss-1 | 2012-12-02T00:00:00Z | 40 | NA | 0 | R8NV-ss-1 |
| ring-necked pheasant | Game Bird | winter | NV | White Pine | 8 | shrub/scrub | R8NV-ss-1 | 2012-12-02T00:00:00Z | 7 | 10 | 0 | R8NV-ss-1 |
| mallard | Game Bird | winter | NV | White Pine | 8 | shrub/scrub | R8NV-ss-1 | 2012-12-02T00:00:00Z | 40 | NA | 0 | R8NV-ss-1 |
| ring-necked pheasant | Game Bird | winter | NV | White Pine | 8 | shrub/scrub | R8NV-ss-1 | 2012-12-02T00:00:00Z | 40 | NA | 0 | R8NV-ss-1 |
| ring-necked pheasant | Game Bird | winter | NV | White Pine | 8 | shrub/scrub | R8NV-ss-1 | 2012-12-02T00:00:00Z | 40 | NA | 0 | R8NV-ss-1 |
| mallard | Game Bird | winter | NV | White Pine | 8 | shrub/scrub | R8NV-ss-1 | 2012-12-02T00:00:00Z | 7 | 10 | 0 | R8NV-ss-1 |
| ring-necked pheasant | Game Bird | winter | NV | White Pine | 8 | shrub/scrub | R8NV-ss-1 | 2013-02-10T00:00:00Z | 4 | 7 | 0 | R8NV-ss-1 |
| ring-necked pheasant | Game Bird | winter | NV | White Pine | 8 | shrub/scrub | R8NV-ss-1 | 2013-02-10T00:00:00Z | 20 | 30 | 0 | R8NV-ss-1 |
| ring-necked pheasant | Game Bird | winter | NV | White Pine | 8 | shrub/scrub | R8NV-ss-1 | 2013-02-10T00:00:00Z | 14 | 20 | 0 | R8NV-ss-1 |
| ring-necked pheasant | Game Bird | winter | NV | White Pine | 8 | shrub/scrub | R8NV-ss-1 | 2013-02-10T00:00:00Z | 2 | 3 | 0 | R8NV-ss-1 |
| ring-necked pheasant | Game Bird | spring | NV | White Pine | 8 | shrub/scrub | R8NV-ss-1 | 2013-03-31T00:00:00Z | 40 | NA | 0 | R8NV-ss-1 |
| ring-necked pheasant | Game Bird | spring | NV | White Pine | 8 | shrub/scrub | R8NV-ss-1 | 2013-03-31T00:00:00Z | 40 | NA | 0 | R8NV-ss-1 |
| ring-necked pheasant | Game Bird | spring | NV | White Pine | 8 | shrub/scrub | R8NV-ss-1 | 2013-03-31T00:00:00Z | 40 | NA | 0 | R8NV-ss-1 |
| ring-necked pheasant | Game Bird | spring | NV | White Pine | 8 | shrub/scrub | R8NV-ss-1 | 2013-03-31T00:00:00Z | 14 | 20 | 0 | R8NV-ss-1 |
| ring-necked pheasant | Game Bird | spring | NV | White Pine | 8 | shrub/scrub | R8NV-ss-1 | 2013-03-31T00:00:00Z | 20 | NA | 0 | R8NV-ss-1 |
| ring-necked pheasant | Game Bird | spring | NV | White Pine | 8 | shrub/scrub | R8NV-ss-1 | 2013-06-02T00:00:00Z | 0 | 1 | 0 | R8NV-ss-1 |
| ring-necked pheasant | Game Bird | spring | NV | White Pine | 8 | shrub/scrub | R8NV-ss-1 | 2013-06-02T00:00:00Z | 40 | NA | 0 | R8NV-ss-1 |
| ring-necked pheasant | Game Bird | spring | NV | White Pine | 8 | shrub/scrub | R8NV-ss-1 | 2013-06-02T00:00:00Z | 40 | NA | 0 | R8NV-ss-1 |
| ring-necked pheasant | Game Bird | spring | NV | White Pine | 8 | shrub/scrub | R8NV-ss-1 | 2013-06-02T00:00:00Z | 41 | NA | 0 | R8NV-ss-1 |
| ring-necked pheasant | Game Bird | spring | NV | White Pine | 8 | shrub/scrub | R8NV-ss-1 | 2013-06-02T00:00:00Z | 20 | 30 | 0 | R8NV-ss-1 |
| ring-necked pheasant | Game Bird | summer | NV | White Pine | 8 | shrub/scrub | R8NV-ss-1 | 2013-08-04T00:00:00Z | 2 | 3 | 0 | R8NV-ss-1 |
| ring-necked pheasant | Game Bird | summer | NV | White Pine | 8 | shrub/scrub | R8NV-ss-1 | 2013-08-04T00:00:00Z | 7 | 10 | 0 | R8NV-ss-1 |
| ring-necked pheasant | Game Bird | summer | NV | White Pine | 8 | shrub/scrub | R8NV-ss-1 | 2013-08-04T00:00:00Z | 2 | 3 | 0 | R8NV-ss-1 |
| ring-necked pheasant | Game Bird | summer | NV | White Pine | 8 | shrub/scrub | R8NV-ss-1 | 2013-08-04T00:00:00Z | 3 | 4 | 0 | R8NV-ss-1 |
| ring-necked pheasant | Game Bird | summer | NV | White Pine | 8 | shrub/scrub | R8NV-ss-1 | 2013-08-04T00:00:00Z | 2 | 3 | 0 | R8NV-ss-1 |
| ring-necked pheasant | Game Bird | fall | NV | White Pine | 8 | shrub/scrub | R8NV-ss-3 | 2013-10-20T00:00:00Z | 4 | 7 | 0 | R8NV-ss-3 |
| ring-necked pheasant | Game Bird | fall | NV | White Pine | 8 | shrub/scrub | R8NV-ss-3 | 2013-10-20T00:00:00Z | 4 | 7 | 0 | R8NV-ss-3 |
| ring-necked pheasant | Game Bird | winter | NV | White Pine | 8 | shrub/scrub | R8NV-ss-3 | 2013-12-30T00:00:00Z | 40 | NA | 0 | R8NV-ss-3 |
| ring-necked pheasant | Game Bird | winter | NV | White Pine | 8 | shrub/scrub | R8NV-ss-3 | 2013-12-30T00:00:00Z | 40 | NA | 0 | R8NV-ss-3 |
| ring-necked pheasant | Game Bird | winter | NV | White Pine | 8 | shrub/scrub | R8NV-ss-3 | 2013-12-30T00:00:00Z | 40 | NA | 0 | R8NV-ss-3 |
| ring-necked pheasant | Game Bird | winter | NV | White Pine | 8 | shrub/scrub | R8NV-ss-3 | 2013-12-30T00:00:00Z | 7 | 10 | 0 | R8NV-ss-3 |
| ring-necked pheasant | Game Bird | winter | NV | White Pine | 8 | shrub/scrub | R8NV-ss-3 | 2013-12-30T00:00:00Z | 10 | 14 | 0 | R8NV-ss-3 |
| ring-necked pheasant | Game Bird | winter | NV | White Pine | 8 | shrub/scrub | R8NV-ss-3 | 2014-01-26T00:00:00Z | 10 | 14 | 0 | R8NV-ss-3 |
| ring-necked pheasant | Game Bird | winter | NV | White Pine | 8 | shrub/scrub | R8NV-ss-3 | 2014-01-26T00:00:00Z | 20 | NA | 0 | R8NV-ss-3 |
| ring-necked pheasant | Game Bird | winter | NV | White Pine | 8 | shrub/scrub | R8NV-ss-3 | 2014-01-26T00:00:00Z | 10 | 14 | 0 | R8NV-ss-3 |
| mallard | Game Bird | summer | NV | White Pine | 8 | shrub/scrub | R8NV-ss-3 | 2014-07-06T00:00:00Z | 4 | 7 | 0 | R8NV-ss-3 |
| mallard | Game Bird | summer | NV | White Pine | 8 | shrub/scrub | R8NV-ss-3 | 2014-07-06T00:00:00Z | 28 | 45 | 0 | R8NV-ss-3 |
| mallard | Game Bird | summer | NV | White Pine | 8 | shrub/scrub | R8NV-ss-3 | 2014-07-06T00:00:00Z | 1 | 2 | 0 | R8NV-ss-3 |
| mallard | Game Bird | summer | NV | White Pine | 8 | shrub/scrub | R8NV-ss-3 | 2014-07-06T00:00:00Z | 14 | 20 | 0 | R8NV-ss-3 |
| ring-necked pheasant | Game Bird | summer | NV | White Pine | 8 | shrub/scrub | R8NV-ss-3 | 2014-07-06T00:00:00Z | 7 | 10 | 0 | R8NV-ss-3 |
| mallard | Game Bird | summer | CA | Kern | 8 | shrub/scrub | R8CA-ss-2 | 2011-07-28T00:00:00Z | 0 | 1 | 1 | R8CA-ss-2 |
| mallard | Game Bird | summer | CA | Kern | 8 | shrub/scrub | R8CA-ss-2 | 2011-07-28T00:00:00Z | 1 | 3 | 1 | R8CA-ss-2 |
| mallard | Game Bird | summer | CA | Kern | 8 | shrub/scrub | R8CA-ss-2 | 2011-07-28T00:00:00Z | 0 | 1 | 1 | R8CA-ss-2 |
| mallard | Game Bird | summer | CA | Kern | 8 | shrub/scrub | R8CA-ss-2 | 2011-07-28T00:00:00Z | 0 | 1 | 1 | R8CA-ss-2 |
| mallard | Game Bird | summer | CA | Kern | 8 | shrub/scrub | R8CA-ss-2 | 2011-07-28T00:00:00Z | 0 | 1 | 1 | R8CA-ss-2 |
| mallard | Game Bird | summer | CA | Kern | 8 | shrub/scrub | R8CA-ss-2 | 2011-07-28T00:00:00Z | 25 | 40 | 1 | R8CA-ss-2 |
| mallard | Game Bird | summer | CA | Kern | 8 | shrub/scrub | R8CA-ss-2 | 2011-07-28T00:00:00Z | 0 | 1 | 1 | R8CA-ss-2 |
| mallard | Game Bird | summer | CA | Kern | 8 | shrub/scrub | R8CA-ss-2 | 2011-07-28T00:00:00Z | 1 | 3 | 1 | R8CA-ss-2 |
| mallard | Game Bird | summer | CA | Kern | 8 | shrub/scrub | R8CA-ss-2 | 2011-09-04T00:00:00Z | 0 | 1 | 1 | R8CA-ss-2 |
| mallard | Game Bird | fall | CA | Kern | 8 | shrub/scrub | R8CA-ss-2 | 2011-09-20T00:00:00Z | 10 | 14 | 1 | R8CA-ss-2 |
| mallard | Game Bird | fall | CA | Kern | 8 | shrub/scrub | R8CA-ss-2 | 2011-09-20T00:00:00Z | 1 | 2 | 1 | R8CA-ss-2 |
| mallard | Game Bird | fall | CA | Kern | 8 | shrub/scrub | R8CA-ss-2 | 2011-09-20T00:00:00Z | 1 | 2 | 1 | R8CA-ss-2 |
| mallard | Game Bird | fall | CA | Kern | 8 | shrub/scrub | R8CA-ss-2 | 2011-09-20T00:00:00Z | 10 | 14 | 1 | R8CA-ss-2 |
| mallard | Game Bird | fall | CA | Kern | 8 | shrub/scrub | R8CA-ss-2 | 2011-09-20T00:00:00Z | 0 | 1 | 1 | R8CA-ss-2 |
| mallard | Game Bird | summer | CA | Kern | 8 | shrub/scrub | R8CA-ss-2 | 2011-09-04T00:00:00Z | 7 | 10 | 1 | R8CA-ss-2 |
| mallard | Game Bird | summer | CA | Kern | 8 | shrub/scrub | R8CA-ss-2 | 2011-09-04T00:00:00Z | 3 | 4 | 1 | R8CA-ss-2 |
| mallard | Game Bird | summer | CA | Kern | 8 | shrub/scrub | R8CA-ss-2 | 2011-09-04T00:00:00Z | 0 | 1 | 1 | R8CA-ss-2 |
| mallard | Game Bird | summer | CA | Kern | 8 | shrub/scrub | R8CA-ss-2 | 2011-09-04T00:00:00Z | 18 | 25 | 1 | R8CA-ss-2 |
| mallard | Game Bird | winter | CA | Kern | 8 | shrub/scrub | R8CA-ss-2 | 2012-01-24T00:00:00Z | 40 | NA | 1 | R8CA-ss-2 |
| mallard | Game Bird | winter | CA | Kern | 8 | shrub/scrub | R8CA-ss-2 | 2012-01-24T00:00:00Z | 10 | 14 | 1 | R8CA-ss-2 |
| red-tailed hawk | Raptor | winter | CA | Kern | 8 | shrub/scrub | R8CA-ss-2 | 2012-01-02T00:00:00Z | 40 | NA | 1 | R8CA-ss-0 |
| mallard | Game Bird | winter | CA | Kern | 8 | shrub/scrub | R8CA-ss-2 | 2012-01-24T00:00:00Z | 0 | 1 | 1 | R8CA-ss-2 |
| mallard | Game Bird | winter | CA | Kern | 8 | shrub/scrub | R8CA-ss-2 | 2012-03-13T00:00:00Z | 25 | NA | 1 | R8CA-ss-2 |
| mallard | Game Bird | winter | CA | Kern | 8 | shrub/scrub | R8CA-ss-2 | 2012-03-13T00:00:00Z | 25 | NA | 1 | R8CA-ss-2 |
| mallard | Game Bird | winter | CA | Kern | 8 | shrub/scrub | R8CA-ss-2 | 2012-03-13T00:00:00Z | 25 | 40 | 1 | R8CA-ss-2 |
| mallard | Game Bird | winter | CA | Kern | 8 | shrub/scrub | R8CA-ss-2 | 2012-03-13T00:00:00Z | 7 | 10 | 1 | R8CA-ss-2 |
| mallard | Game Bird | winter | WY | Converse | 6 | shrub/scrub | R6WY-ss-5 | 2011-01-09T00:00:00Z | 7 | 8 | 0 | R6WY-ss-5 |
| mallard | Game Bird | winter | WY | Converse | 6 | shrub/scrub | R6WY-ss-5 | 2011-01-09T00:00:00Z | 7 | 8 | 0 | R6WY-ss-5 |
| mallard | Game Bird | winter | WY | Converse | 6 | shrub/scrub | R6WY-ss-5 | 2011-01-09T00:00:00Z | 7 | 8 | 0 | R6WY-ss-5 |
| mallard | Game Bird | winter | WY | Converse | 6 | shrub/scrub | R6WY-ss-5 | 2011-01-09T00:00:00Z | 30 | NA | 0 | R6WY-ss-5 |
| mallard | Game Bird | winter | WY | Converse | 6 | shrub/scrub | R6WY-ss-5 | 2011-02-28T00:00:00Z | 16 | NA | 0 | R6WY-ss-5 |
| mallard | Game Bird | winter | WY | Converse | 6 | shrub/scrub | R6WY-ss-5 | 2011-02-28T00:00:00Z | 16 | NA | 0 | R6WY-ss-5 |
| mallard | Game Bird | winter | WY | Converse | 6 | shrub/scrub | R6WY-ss-5 | 2011-02-28T00:00:00Z | 16 | NA | 0 | R6WY-ss-5 |
| mallard | Game Bird | winter | WY | Converse | 6 | shrub/scrub | R6WY-ss-5 | 2011-02-28T00:00:00Z | 16 | NA | 0 | R6WY-ss-5 |
| mallard | Game Bird | spring | WY | Converse | 6 | grassland | R6WY-g-1 | 2010-03-28T00:00:00Z | 40 | NA | 1 | R6WY-g-0 |
| mallard | Game Bird | spring | WY | Converse | 6 | grassland | R6WY-g-1 | 2010-03-28T00:00:00Z | 10 | 23 | 1 | R6WY-g-0 |
| mallard | Game Bird | spring | WY | Converse | 6 | grassland | R6WY-g-1 | 2010-04-26T00:00:00Z | 40 | NA | 1 | R6WY-g-0 |
| mallard | Game Bird | spring | WY | Converse | 6 | grassland | R6WY-g-1 | 2010-04-26T00:00:00Z | 40 | NA | 1 | R6WY-g-0 |
| mallard | Game Bird | spring | WY | Converse | 6 | grassland | R6WY-g-1 | 2010-04-26T00:00:00Z | 2 | 3 | 1 | R6WY-g-0 |
| mallard | Game Bird | spring | WY | Converse | 6 | grassland | R6WY-g-1 | 2010-04-26T00:00:00Z | 7 | 12 | 1 | R6WY-g-0 |
| mallard | Game Bird | spring | WY | Converse | 6 | grassland | R6WY-g-1 | 2010-05-19T00:00:00Z | 40 | NA | 1 | R6WY-g-0 |
| mallard | Game Bird | spring | WY | Converse | 6 | grassland | R6WY-g-1 | 2010-05-19T00:00:00Z | 40 | NA | 1 | R6WY-g-0 |
| mallard | Game Bird | spring | WY | Converse | 6 | grassland | R6WY-g-1 | 2010-05-19T00:00:00Z | 40 | NA | 1 | R6WY-g-0 |
| mallard | Game Bird | spring | WY | Converse | 6 | grassland | R6WY-g-1 | 2010-05-19T00:00:00Z | 40 | NA | 1 | R6WY-g-0 |
| mallard | Game Bird | summer | WY | Converse | 6 | grassland | R6WY-g-1 | 2010-08-15T00:00:00Z | 0 | 1 | 1 | R6WY-g-0 |
| mallard | Game Bird | summer | WY | Converse | 6 | grassland | R6WY-g-1 | 2010-09-12T00:00:00Z | 4 | 7 | 1 | R6WY-g-0 |
| mallard | Game Bird | summer | WY | Converse | 6 | grassland | R6WY-g-1 | 2010-09-12T00:00:00Z | 7 | 10 | 1 | R6WY-g-0 |
| mallard | Game Bird | fall | WY | Converse | 6 | grassland | R6WY-g-1 | 2010-10-10T00:00:00Z | 3 | 4 | 1 | R6WY-g-0 |
| mallard | Game Bird | fall | WY | Converse | 6 | grassland | R6WY-g-1 | 2010-10-10T00:00:00Z | 2 | 3 | 1 | R6WY-g-0 |
| mallard | Game Bird | fall | WY | Converse | 6 | grassland | R6WY-g-1 | 2010-10-10T00:00:00Z | 0 | 1 | 1 | R6WY-g-0 |
| mallard | Game Bird | fall | WY | Converse | 6 | grassland | R6WY-g-1 | 2010-10-10T00:00:00Z | 1 | 2 | 1 | R6WY-g-0 |
| mallard | Game Bird | fall | WY | Converse | 6 | grassland | R6WY-g-1 | 2010-10-10T00:00:00Z | 7 | 10 | 1 | R6WY-g-0 |
| mallard | Game Bird | fall | WY | Converse | 6 | grassland | R6WY-g-1 | 2010-11-04T00:00:00Z | 30 | 40 | 1 | R6WY-g-0 |
| mallard | Game Bird | fall | WY | Converse | 6 | grassland | R6WY-g-1 | 2010-11-04T00:00:00Z | 7 | 10 | 1 | R6WY-g-0 |
| mallard | Game Bird | fall | WY | Converse | 6 | grassland | R6WY-g-1 | 2010-11-04T00:00:00Z | 20 | 40 | 1 | R6WY-g-0 |
| mallard | Game Bird | fall | WY | Converse | 6 | grassland | R6WY-g-1 | 2010-11-04T00:00:00Z | 2 | 3 | 1 | R6WY-g-0 |
| mallard | Game Bird | winter | WY | Converse | 6 | grassland | R6WY-g-1 | 2011-01-09T00:00:00Z | 40 | NA | 1 | R6WY-g-0 |
| mallard | Game Bird | winter | WY | Converse | 6 | grassland | R6WY-g-1 | 2011-01-09T00:00:00Z | 0 | 2 | 1 | R6WY-g-0 |
| mallard | Game Bird | winter | WY | Converse | 6 | grassland | R6WY-g-1 | 2011-01-09T00:00:00Z | 40 | NA | 1 | R6WY-g-0 |
| mallard | Game Bird | winter | WY | Converse | 6 | grassland | R6WY-g-1 | 2011-01-09T00:00:00Z | 2 | 3 | 1 | R6WY-g-0 |
| mallard | Game Bird | summer | WY | Converse | 6 | grassland | R6WY-g-1 | 2010-07-14T00:00:00Z | 1 | 2 | 1 | R6WY-g-0 |
| mallard | Game Bird | summer | WY | Converse | 6 | grassland | R6WY-g-1 | 2010-07-14T00:00:00Z | 40 | NA | 1 | R6WY-g-0 |
| mallard | Game Bird | summer | WY | Converse | 6 | grassland | R6WY-g-1 | 2010-07-14T00:00:00Z | 40 | NA | 1 | R6WY-g-0 |
| mallard | Game Bird | summer | WY | Converse | 6 | grassland | R6WY-g-1 | 2010-07-14T00:00:00Z | 40 | NA | 1 | R6WY-g-0 |
| mallard | Game Bird | spring | WY | Converse | 6 | grassland | R6WY-g-1 | 2010-06-13T00:00:00Z | 40 | NA | 1 | R6WY-g-0 |
| mallard | Game Bird | spring | WY | Converse | 6 | grassland | R6WY-g-1 | 2010-06-13T00:00:00Z | 40 | NA | 1 | R6WY-g-0 |
| mallard | Game Bird | spring | WY | Converse | 6 | grassland | R6WY-g-1 | 2010-06-13T00:00:00Z | 40 | NA | 1 | R6WY-g-0 |
| mallard | Game Bird | spring | WY | Converse | 6 | grassland | R6WY-g-1 | 2010-06-13T00:00:00Z | 40 | NA | 1 | R6WY-g-0 |
| mallard | Game Bird | winter | WA | Garfield | 1 | cropland | R1WA-c-1 | 2013-01-04T00:00:00Z | 40 | NA | 0 | R1WA-c-1 |
| ring-necked pheasant | Game Bird | winter | WA | Garfield | 1 | cropland | R1WA-c-1 | 2013-01-04T00:00:00Z | 1 | 2 | 0 | R1WA-c-1 |
| ring-necked pheasant | Game Bird | winter | WA | Garfield | 1 | cropland | R1WA-c-1 | 2013-01-04T00:00:00Z | 0 | 1 | 0 | R1WA-c-1 |
| mallard | Game Bird | winter | WA | Garfield | 1 | cropland | R1WA-c-1 | 2013-01-04T00:00:00Z | 1 | 2 | 0 | R1WA-c-1 |
| ring-necked pheasant | Game Bird | winter | WA | Garfield | 1 | cropland | R1WA-c-1 | 2013-01-04T00:00:00Z | 40 | NA | 0 | R1WA-c-1 |
| ring-necked pheasant | Game Bird | winter | WA | Garfield | 1 | cropland | R1WA-c-1 | 2013-01-04T00:00:00Z | 2 | 2 | 0 | R1WA-c-1 |
| mallard | Game Bird | winter | WA | Garfield | 1 | cropland | R1WA-c-1 | 2013-01-04T00:00:00Z | 2 | 3 | 0 | R1WA-c-1 |
| ring-necked pheasant | Game Bird | spring | WA | Garfield | 1 | cropland | R1WA-c-1 | 2012-03-28T00:00:00Z | 10 | 14 | 0 | R1WA-c-1 |
| mallard | Game Bird | spring | WA | Garfield | 1 | cropland | R1WA-c-1 | 2012-03-28T00:00:00Z | 40 | NA | 0 | R1WA-c-1 |
| ring-necked pheasant | Game Bird | spring | WA | Garfield | 1 | cropland | R1WA-c-1 | 2012-03-28T00:00:00Z | 20 | 30 | 0 | R1WA-c-1 |
| mallard | Game Bird | spring | WA | Garfield | 1 | cropland | R1WA-c-1 | 2012-03-28T00:00:00Z | 40 | NA | 0 | R1WA-c-1 |
| ring-necked pheasant | Game Bird | spring | WA | Garfield | 1 | cropland | R1WA-c-1 | 2012-03-28T00:00:00Z | 0 | 1 | 0 | R1WA-c-1 |
| mallard | Game Bird | spring | WA | Garfield | 1 | cropland | R1WA-c-1 | 2012-03-28T00:00:00Z | 10 | 14 | 0 | R1WA-c-1 |
| ring-necked pheasant | Game Bird | spring | WA | Garfield | 1 | cropland | R1WA-c-1 | 2012-03-28T00:00:00Z | 14 | 20 | 0 | R1WA-c-1 |
| ring-necked pheasant | Game Bird | spring | WA | Garfield | 1 | cropland | R1WA-c-1 | 2012-04-16T00:00:00Z | 7 | 10 | 0 | R1WA-c-1 |
| mallard | Game Bird | spring | WA | Garfield | 1 | cropland | R1WA-c-1 | 2012-04-16T00:00:00Z | 10 | 14 | 0 | R1WA-c-1 |
| mallard | Game Bird | spring | WA | Garfield | 1 | cropland | R1WA-c-1 | 2012-04-16T00:00:00Z | 7 | 10 | 0 | R1WA-c-1 |
| ring-necked pheasant | Game Bird | spring | WA | Garfield | 1 | cropland | R1WA-c-1 | 2012-04-16T00:00:00Z | 4 | 7 | 0 | R1WA-c-1 |
| mallard | Game Bird | spring | WA | Garfield | 1 | cropland | R1WA-c-1 | 2012-04-16T00:00:00Z | 14 | 20 | 0 | R1WA-c-1 |
| ring-necked pheasant | Game Bird | spring | WA | Garfield | 1 | cropland | R1WA-c-1 | 2012-04-16T00:00:00Z | 40 | NA | 0 | R1WA-c-1 |
| ring-necked pheasant | Game Bird | spring | WA | Garfield | 1 | cropland | R1WA-c-1 | 2012-04-16T00:00:00Z | 40 | NA | 0 | R1WA-c-1 |
| ring-necked pheasant | Game Bird | spring | WA | Garfield | 1 | cropland | R1WA-c-1 | 2012-06-02T00:00:00Z | 40 | NA | 0 | R1WA-c-1 |
| mallard | Game Bird | spring | WA | Garfield | 1 | cropland | R1WA-c-1 | 2012-06-02T00:00:00Z | 40 | NA | 0 | R1WA-c-1 |
| ring-necked pheasant | Game Bird | spring | WA | Garfield | 1 | cropland | R1WA-c-1 | 2012-06-02T00:00:00Z | 0 | 1 | 0 | R1WA-c-1 |
| mallard | Game Bird | spring | WA | Garfield | 1 | cropland | R1WA-c-1 | 2012-06-02T00:00:00Z | 40 | NA | 0 | R1WA-c-1 |
| ring-necked pheasant | Game Bird | spring | WA | Garfield | 1 | cropland | R1WA-c-1 | 2012-06-02T00:00:00Z | 30 | 40 | 0 | R1WA-c-1 |
| ring-necked pheasant | Game Bird | spring | WA | Garfield | 1 | cropland | R1WA-c-1 | 2012-06-02T00:00:00Z | 20 | 30 | 0 | R1WA-c-1 |
| mallard | Game Bird | spring | WA | Garfield | 1 | cropland | R1WA-c-1 | 2012-06-02T00:00:00Z | 20 | 30 | 0 | R1WA-c-1 |
| ring-necked pheasant | Game Bird | winter | WA | Garfield | 1 | cropland | R1WA-c-1 | 2012-11-30T00:00:00Z | 0 | 1 | 0 | R1WA-c-1 |
| ring-necked pheasant | Game Bird | summer | WA | Garfield | 1 | cropland | R1WA-c-1 | 2012-08-22T00:00:00Z | 0 | 1 | 0 | R1WA-c-1 |
| mallard | Game Bird | summer | WA | Garfield | 1 | cropland | R1WA-c-1 | 2012-08-22T00:00:00Z | 4 | 7 | 0 | R1WA-c-1 |
| ring-necked pheasant | Game Bird | summer | WA | Garfield | 1 | cropland | R1WA-c-1 | 2012-08-22T00:00:00Z | 20 | 30 | 0 | R1WA-c-1 |
| mallard | Game Bird | summer | WA | Garfield | 1 | cropland | R1WA-c-1 | 2012-08-22T00:00:00Z | 40 | NA | 0 | R1WA-c-1 |
| ring-necked pheasant | Game Bird | summer | WA | Garfield | 1 | cropland | R1WA-c-1 | 2012-08-22T00:00:00Z | 30 | 40 | 0 | R1WA-c-1 |
| ring-necked pheasant | Game Bird | summer | WA | Garfield | 1 | cropland | R1WA-c-1 | 2012-08-22T00:00:00Z | 7 | 10 | 0 | R1WA-c-1 |
| mallard | Game Bird | summer | WA | Garfield | 1 | cropland | R1WA-c-1 | 2012-08-22T00:00:00Z | 7 | 10 | 0 | R1WA-c-1 |
| ring-necked pheasant | Game Bird | fall | WA | Garfield | 1 | cropland | R1WA-c-1 | 2012-09-18T00:00:00Z | 40 | NA | 0 | R1WA-c-1 |
| mallard | Game Bird | fall | WA | Garfield | 1 | cropland | R1WA-c-1 | 2012-09-18T00:00:00Z | 30 | 40 | 0 | R1WA-c-1 |
| ring-necked pheasant | Game Bird | fall | WA | Garfield | 1 | cropland | R1WA-c-1 | 2012-09-18T00:00:00Z | 40 | NA | 0 | R1WA-c-1 |
| mallard | Game Bird | fall | WA | Garfield | 1 | cropland | R1WA-c-1 | 2012-09-18T00:00:00Z | 2 | 3 | 0 | R1WA-c-1 |
| mallard | Game Bird | fall | WA | Garfield | 1 | cropland | R1WA-c-1 | 2012-09-18T00:00:00Z | 4 | 7 | 0 | R1WA-c-1 |
| ring-necked pheasant | Game Bird | fall | WA | Garfield | 1 | cropland | R1WA-c-1 | 2012-09-18T00:00:00Z | 14 | 20 | 0 | R1WA-c-1 |
| ring-necked pheasant | Game Bird | fall | WA | Garfield | 1 | cropland | R1WA-c-1 | 2012-09-18T00:00:00Z | 10 | 14 | 0 | R1WA-c-1 |
| mallard | Game Bird | winter | WA | Garfield | 1 | cropland | R1WA-c-1 | 2012-11-30T00:00:00Z | 7 | 10 | 0 | R1WA-c-1 |
| ring-necked pheasant | Game Bird | winter | WA | Garfield | 1 | cropland | R1WA-c-1 | 2012-11-30T00:00:00Z | 20 | 30 | 0 | R1WA-c-1 |
| mallard | Game Bird | winter | WA | Garfield | 1 | cropland | R1WA-c-1 | 2012-11-30T00:00:00Z | 10 | 14 | 0 | R1WA-c-1 |
| ring-necked pheasant | Game Bird | winter | WA | Garfield | 1 | cropland | R1WA-c-1 | 2012-11-30T00:00:00Z | 0 | 1 | 0 | R1WA-c-1 |
| mallard | Game Bird | winter | WA | Garfield | 1 | cropland | R1WA-c-1 | 2012-11-30T00:00:00Z | 10 | 14 | 0 | R1WA-c-1 |
| ring-necked pheasant | Game Bird | winter | WA | Garfield | 1 | cropland | R1WA-c-1 | 2012-11-30T00:00:00Z | 4 | 7 | 0 | R1WA-c-1 |
| mallard | Game Bird | winter | WA | Columbia | 1 | cropland | R1WA-c-5 | 2015-02-02T00:00:00Z | 3 | 4 | 0 | R1WA-c-5 |
| ring-necked pheasant | Game Bird | winter | WA | Columbia | 1 | cropland | R1WA-c-5 | 2015-02-02T00:00:00Z | 5 | 6 | 0 | R1WA-c-5 |
| ring-necked pheasant | Game Bird | winter | WA | Columbia | 1 | cropland | R1WA-c-5 | 2015-02-02T00:00:00Z | 36.5 | NA | 0 | R1WA-c-5 |
| ring-necked pheasant | Game Bird | winter | WA | Columbia | 1 | cropland | R1WA-c-5 | 2015-02-02T00:00:00Z | 4 | 5 | 0 | R1WA-c-5 |
| mallard | Game Bird | winter | WA | Columbia | 1 | cropland | R1WA-c-5 | 2015-02-02T00:00:00Z | 6 | 9 | 0 | R1WA-c-5 |
| ring-necked pheasant | Game Bird | winter | WA | Columbia | 1 | cropland | R1WA-c-5 | 2015-02-02T00:00:00Z | 1 | 2 | 0 | R1WA-c-5 |
| ring-necked pheasant | Game Bird | spring | WA | Columbia | 1 | cropland | R1WA-c-5 | 2015-03-30T00:00:00Z | 10 | 14 | 0 | R1WA-c-5 |
| ring-necked pheasant | Game Bird | spring | WA | Columbia | 1 | cropland | R1WA-c-5 | 2015-03-30T00:00:00Z | 7 | 10 | 0 | R1WA-c-5 |
| ring-necked pheasant | Game Bird | spring | WA | Columbia | 1 | cropland | R1WA-c-5 | 2015-03-30T00:00:00Z | 7 | 10 | 0 | R1WA-c-5 |
| mallard | Game Bird | spring | WA | Columbia | 1 | cropland | R1WA-c-5 | 2015-03-30T00:00:00Z | 0 | 1 | 0 | R1WA-c-5 |
| mallard | Game Bird | spring | WA | Columbia | 1 | cropland | R1WA-c-5 | 2015-03-30T00:00:00Z | 4 | 7 | 0 | R1WA-c-5 |
| ring-necked pheasant | Game Bird | spring | WA | Columbia | 1 | cropland | R1WA-c-5 | 2015-05-18T00:00:00Z | 2 | 3 | 0 | R1WA-c-5 |
| ring-necked pheasant | Game Bird | spring | WA | Columbia | 1 | cropland | R1WA-c-5 | 2015-05-18T00:00:00Z | 2 | 3 | 0 | R1WA-c-5 |
| mallard | Game Bird | spring | WA | Columbia | 1 | cropland | R1WA-c-5 | 2015-05-18T00:00:00Z | 3 | 4 | 0 | R1WA-c-5 |
| mallard | Game Bird | spring | WA | Columbia | 1 | cropland | R1WA-c-5 | 2015-05-18T00:00:00Z | 2 | 3 | 0 | R1WA-c-5 |
| mallard | Game Bird | spring | WA | Columbia | 1 | cropland | R1WA-c-5 | 2015-05-18T00:00:00Z | 2 | 3 | 0 | R1WA-c-5 |
| mallard | Game Bird | spring | WA | Columbia | 1 | cropland | R1WA-c-5 | 2015-05-18T00:00:00Z | 4 | 7 | 0 | R1WA-c-5 |
| mallard | Game Bird | spring | WA | Columbia | 1 | cropland | R1WA-c-5 | 2015-05-18T00:00:00Z | 4 | 7 | 0 | R1WA-c-5 |
| ring-necked pheasant | Game Bird | summer | WA | Columbia | 1 | cropland | R1WA-c-5 | 2015-09-14T00:00:00Z | 2 | 3 | 0 | R1WA-c-5 |
| mallard | Game Bird | summer | WA | Columbia | 1 | cropland | R1WA-c-5 | 2015-09-14T00:00:00Z | 3 | NA | 0 | R1WA-c-5 |
| ring-necked pheasant | Game Bird | summer | WA | Columbia | 1 | cropland | R1WA-c-5 | 2015-09-14T00:00:00Z | 21 | 29 | 0 | R1WA-c-5 |
| mallard | Game Bird | summer | WA | Columbia | 1 | cropland | R1WA-c-5 | 2015-09-14T00:00:00Z | 7 | 10 | 0 | R1WA-c-5 |
| mallard | Game Bird | summer | WA | Columbia | 1 | cropland | R1WA-c-5 | 2015-09-14T00:00:00Z | 14 | 21 | 0 | R1WA-c-5 |
| ring-necked pheasant | Game Bird | summer | WA | Columbia | 1 | cropland | R1WA-c-5 | 2015-09-14T00:00:00Z | 10 | 14 | 0 | R1WA-c-5 |
| ring-necked pheasant | Game Bird | summer | WA | Columbia | 1 | cropland | R1WA-c-5 | 2015-09-14T00:00:00Z | 10 | 14 | 0 | R1WA-c-5 |
| ring-necked pheasant | Game Bird | summer | WA | Columbia | 1 | cropland | R1WA-c-5 | 2015-09-14T00:00:00Z | 7 | 10 | 0 | R1WA-c-5 |
| ring-necked pheasant | Game Bird | summer | WA | Klickitat | 1 | grassland | R1WA-g-6 | 2011-06-20T00:00:00Z | 14 | 21 | 0 | R1WA-g-6 |
| mallard | Game Bird | summer | WA | Klickitat | 1 | grassland | R1WA-g-6 | 2011-06-20T00:00:00Z | 21 | 30 | 0 | R1WA-g-6 |
| ring-necked pheasant | Game Bird | summer | WA | Klickitat | 1 | grassland | R1WA-g-6 | 2011-06-20T00:00:00Z | 7 | 10 | 0 | R1WA-g-6 |
| mallard | Game Bird | summer | WA | Klickitat | 1 | grassland | R1WA-g-6 | 2011-06-20T00:00:00Z | 3 | 4 | 0 | R1WA-g-6 |
| mallard | Game Bird | summer | WA | Klickitat | 1 | grassland | R1WA-g-6 | 2011-06-20T00:00:00Z | 40 | NA | 0 | R1WA-g-6 |
| ring-necked pheasant | Game Bird | summer | WA | Klickitat | 1 | grassland | R1WA-g-6 | 2011-06-20T00:00:00Z | 1 | 2 | 0 | R1WA-g-6 |
| ring-necked pheasant | Game Bird | summer | WA | Klickitat | 1 | grassland | R1WA-g-6 | 2011-06-20T00:00:00Z | 1 | 2 | 0 | R1WA-g-6 |
| mallard | Game Bird | summer | WA | Klickitat | 1 | grassland | R1WA-g-6 | 2011-06-20T00:00:00Z | 7 | 10 | 0 | R1WA-g-6 |
| ring-necked pheasant | Game Bird | summer | WA | Klickitat | 1 | grassland | R1WA-g-6 | 2011-06-20T00:00:00Z | 0 | 1 | 0 | R1WA-g-6 |
| mallard | Game Bird | summer | WA | Klickitat | 1 | grassland | R1WA-g-6 | 2011-06-20T00:00:00Z | 0 | 1 | 0 | R1WA-g-6 |
| ring-necked pheasant | Game Bird | summer | WA | Klickitat | 1 | grassland | R1WA-g-6 | 2011-09-05T00:00:00Z | 4 | 7 | 0 | R1WA-g-6 |
| ring-necked pheasant | Game Bird | summer | WA | Klickitat | 1 | grassland | R1WA-g-6 | 2011-09-05T00:00:00Z | 7 | 10 | 0 | R1WA-g-6 |
| ring-necked pheasant | Game Bird | summer | WA | Klickitat | 1 | grassland | R1WA-g-6 | 2011-09-05T00:00:00Z | 4 | 7 | 0 | R1WA-g-6 |
| ring-necked pheasant | Game Bird | summer | WA | Klickitat | 1 | grassland | R1WA-g-6 | 2011-09-05T00:00:00Z | 7 | 10 | 0 | R1WA-g-6 |
| ring-necked pheasant | Game Bird | summer | WA | Klickitat | 1 | grassland | R1WA-g-6 | 2011-09-05T00:00:00Z | 30 | 40 | 0 | R1WA-g-6 |
| ring-necked pheasant | Game Bird | summer | WA | Klickitat | 1 | grassland | R1WA-g-6 | 2011-09-05T00:00:00Z | 21 | 30 | 0 | R1WA-g-6 |
| ring-necked pheasant | Game Bird | summer | WA | Klickitat | 1 | grassland | R1WA-g-6 | 2011-09-05T00:00:00Z | 40 | NA | 0 | R1WA-g-6 |
| ring-necked pheasant | Game Bird | summer | WA | Klickitat | 1 | grassland | R1WA-g-6 | 2011-09-05T00:00:00Z | 10 | 14 | 0 | R1WA-g-6 |
| ring-necked pheasant | Game Bird | summer | WA | Klickitat | 1 | grassland | R1WA-g-6 | 2011-09-05T00:00:00Z | 10 | 14 | 0 | R1WA-g-6 |
| mallard | Game Bird | spring | WA | Klickitat | 1 | grassland | R1WA-g-6 | 2012-04-09T00:00:00Z | 14 | 21 | 0 | R1WA-g-6 |
| ring-necked pheasant | Game Bird | spring | WA | Klickitat | 1 | grassland | R1WA-g-6 | 2012-04-09T00:00:00Z | 14 | 21 | 0 | R1WA-g-6 |
| mallard | Game Bird | spring | WA | Klickitat | 1 | grassland | R1WA-g-6 | 2012-04-09T00:00:00Z | 7 | 10 | 0 | R1WA-g-6 |
| mallard | Game Bird | spring | WA | Klickitat | 1 | grassland | R1WA-g-6 | 2012-04-09T00:00:00Z | 40 | NA | 0 | R1WA-g-6 |
| ring-necked pheasant | Game Bird | spring | WA | Klickitat | 1 | grassland | R1WA-g-6 | 2012-04-09T00:00:00Z | 10 | 14 | 0 | R1WA-g-6 |
| ring-necked pheasant | Game Bird | winter | WA | Klickitat | 1 | grassland | R1WA-g-6 | 2012-01-09T00:00:00Z | 21 | 30 | 0 | R1WA-g-6 |
| ring-necked pheasant | Game Bird | winter | WA | Klickitat | 1 | grassland | R1WA-g-6 | 2012-01-09T00:00:00Z | 21 | 30 | 0 | R1WA-g-6 |
| ring-necked pheasant | Game Bird | winter | WA | Klickitat | 1 | grassland | R1WA-g-6 | 2012-01-09T00:00:00Z | 21 | 30 | 0 | R1WA-g-6 |
| ring-necked pheasant | Game Bird | winter | WA | Klickitat | 1 | grassland | R1WA-g-6 | 2012-01-09T00:00:00Z | 14 | 21 | 0 | R1WA-g-6 |
| mallard | Game Bird | winter | WA | Klickitat | 1 | grassland | R1WA-g-6 | 2012-01-09T00:00:00Z | 4 | 7 | 0 | R1WA-g-6 |
| mallard | Game Bird | winter | WA | Klickitat | 1 | grassland | R1WA-g-6 | 2012-01-09T00:00:00Z | 40 | NA | 0 | R1WA-g-6 |
| ring-necked pheasant | Game Bird | winter | WA | Klickitat | 1 | grassland | R1WA-g-6 | 2012-01-09T00:00:00Z | 40 | NA | 0 | R1WA-g-6 |
| mallard | Game Bird | winter | WA | Klickitat | 1 | grassland | R1WA-g-6 | 2012-01-09T00:00:00Z | 40 | NA | 0 | R1WA-g-6 |
| mallard | Game Bird | summer | WA | Klickitat | 1 | grassland | R1WA-g-7 | 2013-08-26T00:00:00Z | 4 | 7 | 1 | R1WA-g-7 |
| ring-necked pheasant | Game Bird | summer | WA | Klickitat | 1 | grassland | R1WA-g-7 | 2013-08-26T00:00:00Z | 3 | 4 | 1 | R1WA-g-7 |
| mallard | Game Bird | summer | WA | Klickitat | 1 | grassland | R1WA-g-7 | 2013-08-26T00:00:00Z | 3 | 4 | 1 | R1WA-g-7 |
| ring-necked pheasant | Game Bird | summer | WA | Klickitat | 1 | grassland | R1WA-g-7 | 2013-08-26T00:00:00Z | 4 | 7 | 1 | R1WA-g-7 |
| mallard | Game Bird | summer | WA | Klickitat | 1 | grassland | R1WA-g-7 | 2013-08-26T00:00:00Z | 4 | 7 | 1 | R1WA-g-7 |
| ring-necked pheasant | Game Bird | summer | WA | Klickitat | 1 | grassland | R1WA-g-7 | 2013-08-26T00:00:00Z | 4 | 7 | 1 | R1WA-g-7 |
| mallard | Game Bird | summer | WA | Klickitat | 1 | grassland | R1WA-g-7 | 2013-08-26T00:00:00Z | 3 | NA | 1 | R1WA-g-7 |
| mallard | Game Bird | summer | WA | Klickitat | 1 | grassland | R1WA-g-7 | 2013-08-26T00:00:00Z | 2 | NA | 1 | R1WA-g-7 |
| ring-necked pheasant | Game Bird | summer | WA | Klickitat | 1 | grassland | R1WA-g-7 | 2013-08-26T00:00:00Z | 3 | 4 | 1 | R1WA-g-7 |
| mallard | Game Bird | summer | WA | Klickitat | 1 | grassland | R1WA-g-7 | 2013-08-26T00:00:00Z | 4 | 7 | 1 | R1WA-g-7 |
| ring-necked pheasant | Game Bird | summer | WA | Klickitat | 1 | grassland | R1WA-g-7 | 2013-08-26T00:00:00Z | 1 | 2 | 1 | R1WA-g-7 |
| mallard | Game Bird | summer | WA | Klickitat | 1 | grassland | R1WA-g-7 | 2013-08-26T00:00:00Z | 3 | NA | 1 | R1WA-g-7 |
| ring-necked pheasant | Game Bird | summer | WA | Klickitat | 1 | grassland | R1WA-g-7 | 2013-08-26T00:00:00Z | 3 | 4 | 1 | R1WA-g-7 |
| mallard | Game Bird | summer | WA | Klickitat | 1 | grassland | R1WA-g-7 | 2013-08-26T00:00:00Z | 3 | 4 | 1 | R1WA-g-7 |
| ring-necked pheasant | Game Bird | summer | WA | Klickitat | 1 | grassland | R1WA-g-7 | 2013-08-26T00:00:00Z | 1 | 2 | 1 | R1WA-g-7 |
| mallard | Game Bird | summer | WA | Klickitat | 1 | grassland | R1WA-g-7 | 2013-08-26T00:00:00Z | 2 | 3 | 1 | R1WA-g-7 |
| ring-necked pheasant | Game Bird | summer | WA | Klickitat | 1 | grassland | R1WA-g-7 | 2013-08-26T00:00:00Z | 2 | 3 | 1 | R1WA-g-7 |
| ring-necked pheasant | Game Bird | summer | WA | Klickitat | 1 | grassland | R1WA-g-7 | 2013-08-26T00:00:00Z | 4 | 7 | 1 | R1WA-g-7 |
| mallard | Game Bird | summer | WA | Klickitat | 1 | grassland | R1WA-g-7 | 2013-08-26T00:00:00Z | 4 | 7 | 1 | R1WA-g-7 |
| ring-necked pheasant | Game Bird | summer | WA | Klickitat | 1 | grassland | R1WA-g-7 | 2013-08-26T00:00:00Z | 4 | 10 | 1 | R1WA-g-7 |
| ring-necked pheasant | Game Bird | fall | WA | Klickitat | 1 | grassland | R1WA-g-7 | 2013-09-16T00:00:00Z | 10 | 14 | 1 | R1WA-g-7 |
| mallard | Game Bird | fall | WA | Klickitat | 1 | grassland | R1WA-g-7 | 2013-09-16T00:00:00Z | 10 | 14 | 1 | R1WA-g-7 |
| ring-necked pheasant | Game Bird | fall | WA | Klickitat | 1 | grassland | R1WA-g-7 | 2013-09-16T00:00:00Z | 2 | 3 | 1 | R1WA-g-7 |
| mallard | Game Bird | fall | WA | Klickitat | 1 | grassland | R1WA-g-7 | 2013-09-16T00:00:00Z | 2 | 3 | 1 | R1WA-g-7 |
| ring-necked pheasant | Game Bird | fall | WA | Klickitat | 1 | grassland | R1WA-g-7 | 2013-09-16T00:00:00Z | 1 | 2 | 1 | R1WA-g-7 |
| mallard | Game Bird | fall | WA | Klickitat | 1 | grassland | R1WA-g-7 | 2013-09-16T00:00:00Z | 10 | 14 | 1 | R1WA-g-7 |
| ring-necked pheasant | Game Bird | fall | WA | Klickitat | 1 | grassland | R1WA-g-7 | 2013-09-16T00:00:00Z | 10 | 14 | 1 | R1WA-g-7 |
| mallard | Game Bird | fall | WA | Klickitat | 1 | grassland | R1WA-g-7 | 2013-09-16T00:00:00Z | 1 | 2 | 1 | R1WA-g-7 |
| mallard | Game Bird | fall | WA | Klickitat | 1 | grassland | R1WA-g-7 | 2013-09-23T00:00:00Z | 4 | 7 | 1 | R1WA-g-7 |
| mallard | Game Bird | fall | WA | Klickitat | 1 | grassland | R1WA-g-7 | 2013-09-23T00:00:00Z | 0 | 1 | 1 | R1WA-g-7 |
| ring-necked pheasant | Game Bird | fall | WA | Klickitat | 1 | grassland | R1WA-g-7 | 2013-09-23T00:00:00Z | 0 | 1 | 1 | R1WA-g-7 |
| ring-necked pheasant | Game Bird | fall | WA | Klickitat | 1 | grassland | R1WA-g-7 | 2013-09-23T00:00:00Z | 4 | 7 | 1 | R1WA-g-7 |
| ring-necked pheasant | Game Bird | fall | WA | Klickitat | 1 | grassland | R1WA-g-7 | 2013-09-23T00:00:00Z | 4 | 7 | 1 | R1WA-g-7 |
| ring-necked pheasant | Game Bird | fall | WA | Klickitat | 1 | grassland | R1WA-g-7 | 2013-09-23T00:00:00Z | 4 | 7 | 1 | R1WA-g-7 |
| mallard | Game Bird | fall | WA | Klickitat | 1 | grassland | R1WA-g-7 | 2013-09-23T00:00:00Z | 4 | 7 | 1 | R1WA-g-7 |
| mallard | Game Bird | fall | WA | Klickitat | 1 | grassland | R1WA-g-7 | 2013-09-23T00:00:00Z | 0 | 1 | 1 | R1WA-g-7 |
| mallard | Game Bird | fall | WA | Klickitat | 1 | grassland | R1WA-g-7 | 2013-09-30T00:00:00Z | 4 | 7 | 1 | R1WA-g-7 |
| ring-necked pheasant | Game Bird | fall | WA | Klickitat | 1 | grassland | R1WA-g-7 | 2013-09-30T00:00:00Z | 1 | 2 | 1 | R1WA-g-7 |
| mallard | Game Bird | fall | WA | Klickitat | 1 | grassland | R1WA-g-7 | 2013-09-30T00:00:00Z | 1 | 2 | 1 | R1WA-g-7 |
| ring-necked pheasant | Game Bird | fall | WA | Klickitat | 1 | grassland | R1WA-g-7 | 2013-09-30T00:00:00Z | 14 | 21 | 1 | R1WA-g-7 |
| mallard | Game Bird | fall | WA | Klickitat | 1 | grassland | R1WA-g-7 | 2013-09-30T00:00:00Z | 1 | 2 | 1 | R1WA-g-7 |
| ring-necked pheasant | Game Bird | fall | WA | Klickitat | 1 | grassland | R1WA-g-7 | 2013-09-30T00:00:00Z | 4 | 7 | 1 | R1WA-g-7 |
| mallard | Game Bird | fall | WA | Klickitat | 1 | grassland | R1WA-g-7 | 2013-09-30T00:00:00Z | 4 | 7 | 1 | R1WA-g-7 |
| ring-necked pheasant | Game Bird | fall | WA | Klickitat | 1 | grassland | R1WA-g-7 | 2013-09-30T00:00:00Z | 1 | 2 | 1 | R1WA-g-7 |
| mallard | Game Bird | fall | WA | Klickitat | 1 | grassland | R1WA-g-7 | 2013-10-07T00:00:00Z | 4 | 7 | 1 | R1WA-g-7 |
| ring-necked pheasant | Game Bird | fall | WA | Klickitat | 1 | grassland | R1WA-g-7 | 2013-10-07T00:00:00Z | 0 | 1 | 1 | R1WA-g-7 |
| mallard | Game Bird | fall | WA | Klickitat | 1 | grassland | R1WA-g-7 | 2013-10-07T00:00:00Z | 4 | 7 | 1 | R1WA-g-7 |
| ring-necked pheasant | Game Bird | fall | WA | Klickitat | 1 | grassland | R1WA-g-7 | 2013-10-07T00:00:00Z | 10 | 14 | 1 | R1WA-g-7 |
| mallard | Game Bird | fall | WA | Klickitat | 1 | grassland | R1WA-g-7 | 2013-10-07T00:00:00Z | 4 | 7 | 1 | R1WA-g-7 |
| ring-necked pheasant | Game Bird | fall | WA | Klickitat | 1 | grassland | R1WA-g-7 | 2013-10-07T00:00:00Z | 10 | 14 | 1 | R1WA-g-7 |
| mallard | Game Bird | fall | WA | Klickitat | 1 | grassland | R1WA-g-7 | 2013-10-07T00:00:00Z | 4 | 7 | 1 | R1WA-g-7 |
| ring-necked pheasant | Game Bird | fall | WA | Klickitat | 1 | grassland | R1WA-g-7 | 2013-10-07T00:00:00Z | 1 | 2 | 1 | R1WA-g-7 |
| mallard | Game Bird | fall | WA | Klickitat | 1 | grassland | R1WA-g-7 | 2013-10-14T00:00:00Z | 1 | 2 | 1 | R1WA-g-7 |
| ring-necked pheasant | Game Bird | fall | WA | Klickitat | 1 | grassland | R1WA-g-7 | 2013-10-14T00:00:00Z | 1 | 2 | 1 | R1WA-g-7 |
| mallard | Game Bird | fall | WA | Klickitat | 1 | grassland | R1WA-g-7 | 2013-10-14T00:00:00Z | 1 | 2 | 1 | R1WA-g-7 |
| ring-necked pheasant | Game Bird | fall | WA | Klickitat | 1 | grassland | R1WA-g-7 | 2013-10-14T00:00:00Z | 7 | 10 | 1 | R1WA-g-7 |
| mallard | Game Bird | fall | WA | Klickitat | 1 | grassland | R1WA-g-7 | 2013-10-14T00:00:00Z | 0 | 1 | 1 | R1WA-g-7 |
| ring-necked pheasant | Game Bird | fall | WA | Klickitat | 1 | grassland | R1WA-g-7 | 2013-10-14T00:00:00Z | 0 | 1 | 1 | R1WA-g-7 |
| red-tailed hawk | Raptor | winter | WA | Klickitat | 1 | grassland | R1WA-g-7 | 2013-11-18T00:00:00Z | 4 | 7 | 1 | R1WA-g-7 |
| mallard | Game Bird | winter | WA | Klickitat | 1 | grassland | R1WA-g-7 | 2013-11-18T00:00:00Z | 0 | 1 | 1 | R1WA-g-7 |
| ring-necked pheasant | Game Bird | winter | WA | Klickitat | 1 | grassland | R1WA-g-7 | 2013-11-18T00:00:00Z | 21 | 28 | 1 | R1WA-g-7 |
| mallard | Game Bird | winter | WA | Klickitat | 1 | grassland | R1WA-g-7 | 2013-11-18T00:00:00Z | 1 | 2 | 1 | R1WA-g-7 |
| ring-necked pheasant | Game Bird | winter | WA | Klickitat | 1 | grassland | R1WA-g-7 | 2013-11-18T00:00:00Z | 2 | 3 | 1 | R1WA-g-7 |
| great horned owl | Raptor | winter | WA | Klickitat | 1 | grassland | R1WA-g-7 | 2013-11-18T00:00:00Z | 63 | NA | 1 | R1WA-g-7 |
| mallard | Game Bird | winter | WA | Klickitat | 1 | grassland | R1WA-g-7 | 2013-11-18T00:00:00Z | 2 | 3 | 1 | R1WA-g-7 |
| red-tailed hawk | Raptor | winter | WA | Klickitat | 1 | grassland | R1WA-g-7 | 2013-11-18T00:00:00Z | 63 | NA | 1 | R1WA-g-7 |
| ring-necked pheasant | Game Bird | winter | WA | Klickitat | 1 | grassland | R1WA-g-7 | 2013-11-18T00:00:00Z | 14 | 21 | 1 | R1WA-g-7 |
| mallard | Game Bird | winter | WA | Klickitat | 1 | grassland | R1WA-g-7 | 2013-11-18T00:00:00Z | 3 | 4 | 1 | R1WA-g-7 |
| ring-necked pheasant | Game Bird | winter | WA | Klickitat | 1 | grassland | R1WA-g-7 | 2013-11-18T00:00:00Z | 1 | 2 | 1 | R1WA-g-7 |
| great horned owl | Raptor | winter | WA | Klickitat | 1 | grassland | R1WA-g-7 | 2013-11-18T00:00:00Z | 28 | 35 | 1 | R1WA-g-7 |
| mallard | Game Bird | winter | WA | Klickitat | 1 | grassland | R1WA-g-7 | 2013-12-02T00:00:00Z | 4 | 7 | 1 | R1WA-g-7 |
| ring-necked pheasant | Game Bird | winter | WA | Klickitat | 1 | grassland | R1WA-g-7 | 2013-12-02T00:00:00Z | 7 | 10 | 1 | R1WA-g-7 |
| mallard | Game Bird | winter | WA | Klickitat | 1 | grassland | R1WA-g-7 | 2013-12-02T00:00:00Z | 28 | 35 | 1 | R1WA-g-7 |
| ring-necked pheasant | Game Bird | winter | WA | Klickitat | 1 | grassland | R1WA-g-7 | 2013-12-02T00:00:00Z | 14 | 21 | 1 | R1WA-g-7 |
| red-tailed hawk | Raptor | winter | WA | Klickitat | 1 | grassland | R1WA-g-7 | 2013-12-02T00:00:00Z | 14 | 21 | 1 | R1WA-g-7 |
| mallard | Game Bird | winter | WA | Klickitat | 1 | grassland | R1WA-g-7 | 2013-12-02T00:00:00Z | 2 | 3 | 1 | R1WA-g-7 |
| red-tailed hawk | Raptor | winter | WA | Klickitat | 1 | grassland | R1WA-g-7 | 2013-12-02T00:00:00Z | 4 | 7 | 1 | R1WA-g-7 |
| ring-necked pheasant | Game Bird | winter | WA | Klickitat | 1 | grassland | R1WA-g-7 | 2013-12-02T00:00:00Z | 10 | 14 | 1 | R1WA-g-7 |
| mallard | Game Bird | winter | WA | Klickitat | 1 | grassland | R1WA-g-7 | 2013-12-02T00:00:00Z | 3 | 4 | 1 | R1WA-g-7 |
| great horned owl | Raptor | winter | WA | Klickitat | 1 | grassland | R1WA-g-7 | 2013-12-02T00:00:00Z | 21 | 28 | 1 | R1WA-g-7 |
| ring-necked pheasant | Game Bird | winter | WA | Klickitat | 1 | grassland | R1WA-g-7 | 2013-12-02T00:00:00Z | 0 | 1 | 1 | R1WA-g-7 |
| osprey | Raptor | winter | WA | Klickitat | 1 | grassland | R1WA-g-7 | 2013-12-02T00:00:00Z | 4 | 7 | 1 | R1WA-g-7 |
| ring-necked pheasant | Game Bird | winter | WA | Klickitat | 1 | grassland | R1WA-g-7 | 2013-12-16T00:00:00Z | 0 | 1 | 1 | R1WA-g-7 |
| great horned owl | Raptor | winter | WA | Klickitat | 1 | grassland | R1WA-g-7 | 2013-12-16T00:00:00Z | 14 | 21 | 1 | R1WA-g-7 |
| mallard | Game Bird | winter | WA | Klickitat | 1 | grassland | R1WA-g-7 | 2013-12-16T00:00:00Z | 7 | 10 | 1 | R1WA-g-7 |
| ring-necked pheasant | Game Bird | winter | WA | Klickitat | 1 | grassland | R1WA-g-7 | 2013-12-16T00:00:00Z | 21 | 28 | 1 | R1WA-g-7 |
| mallard | Game Bird | winter | WA | Klickitat | 1 | grassland | R1WA-g-7 | 2013-12-16T00:00:00Z | 4 | 7 | 1 | R1WA-g-7 |
| ring-necked pheasant | Game Bird | winter | WA | Klickitat | 1 | grassland | R1WA-g-7 | 2013-12-16T00:00:00Z | 4 | 7 | 1 | R1WA-g-7 |
| mallard | Game Bird | winter | WA | Klickitat | 1 | grassland | R1WA-g-7 | 2013-12-16T00:00:00Z | 7 | 10 | 1 | R1WA-g-7 |
| red-tailed hawk | Raptor | winter | WA | Klickitat | 1 | grassland | R1WA-g-7 | 2013-12-16T00:00:00Z | 4 | 7 | 1 | R1WA-g-7 |
| ring-necked pheasant | Game Bird | winter | WA | Klickitat | 1 | grassland | R1WA-g-7 | 2013-12-16T00:00:00Z | 4 | 7 | 1 | R1WA-g-7 |
| great horned owl | Raptor | winter | WA | Klickitat | 1 | grassland | R1WA-g-7 | 2013-12-16T00:00:00Z | 21 | 28 | 1 | R1WA-g-7 |
| red-tailed hawk | Raptor | winter | WA | Klickitat | 1 | grassland | R1WA-g-7 | 2013-12-16T00:00:00Z | 7 | 10 | 1 | R1WA-g-7 |
| mallard | Game Bird | winter | WA | Klickitat | 1 | grassland | R1WA-g-7 | 2013-12-16T00:00:00Z | 7 | 10 | 1 | R1WA-g-7 |
| red-tailed hawk | Raptor | summer | WA | Klickitat | 1 | grassland | R1WA-g-8 | 2015-06-22T00:00:00Z | 146 | 160 | 1 | R1WA-g-8 |
| red-tailed hawk | Raptor | summer | WA | Klickitat | 1 | grassland | R1WA-g-8 | 2015-06-22T00:00:00Z | 146 | 160 | 1 | R1WA-g-8 |
| red-tailed hawk | Raptor | summer | WA | Klickitat | 1 | grassland | R1WA-g-8 | 2015-06-22T00:00:00Z | 146 | 160 | 1 | R1WA-g-8 |
| ring-necked pheasant | Game Bird | summer | WA | Klickitat | 1 | grassland | R1WA-g-8 | 2015-06-22T00:00:00Z | 4 | 7 | 1 | R1WA-g-8 |
| great horned owl | Raptor | summer | WA | Klickitat | 1 | grassland | R1WA-g-8 | 2015-06-22T00:00:00Z | 160 | NA | 1 | R1WA-g-8 |
| mallard | Game Bird | summer | WA | Klickitat | 1 | grassland | R1WA-g-8 | 2015-06-22T00:00:00Z | 160 | NA | 1 | R1WA-g-8 |
| ring-necked pheasant | Game Bird | summer | WA | Klickitat | 1 | grassland | R1WA-g-8 | 2015-06-22T00:00:00Z | 1 | 2 | 1 | R1WA-g-8 |
| mallard | Game Bird | summer | WA | Klickitat | 1 | grassland | R1WA-g-8 | 2015-06-22T00:00:00Z | 146 | 160 | 1 | R1WA-g-8 |
| ring-necked pheasant | Game Bird | summer | WA | Klickitat | 1 | grassland | R1WA-g-8 | 2015-06-22T00:00:00Z | 4 | 7 | 1 | R1WA-g-8 |
| mallard | Game Bird | summer | WA | Klickitat | 1 | grassland | R1WA-g-8 | 2015-06-22T00:00:00Z | 1 | 2 | 1 | R1WA-g-8 |
| red-tailed hawk | Raptor | summer | WA | Klickitat | 1 | grassland | R1WA-g-8 | 2015-06-22T00:00:00Z | 146 | 160 | 1 | R1WA-g-8 |
| ring-necked pheasant | Game Bird | summer | WA | Klickitat | 1 | grassland | R1WA-g-8 | 2015-06-22T00:00:00Z | 10 | 14 | 1 | R1WA-g-8 |
| red-tailed hawk | Raptor | summer | WA | Klickitat | 1 | grassland | R1WA-g-8 | 2015-06-22T00:00:00Z | 14 | 21 | 1 | R1WA-g-8 |
| mallard | Game Bird | summer | WA | Klickitat | 1 | grassland | R1WA-g-8 | 2015-06-22T00:00:00Z | 14 | 21 | 1 | R1WA-g-8 |
| red-tailed hawk | Raptor | summer | WA | Klickitat | 1 | grassland | R1WA-g-8 | 2015-06-22T00:00:00Z | 146 | 160 | 1 | R1WA-g-8 |
| red-tailed hawk | Raptor | summer | WA | Klickitat | 1 | grassland | R1WA-g-8 | 2015-06-22T00:00:00Z | 2 | 3 | 1 | R1WA-g-8 |
| red-tailed hawk | Raptor | summer | WA | Klickitat | 1 | grassland | R1WA-g-8 | 2015-06-22T00:00:00Z | 2 | 3 | 1 | R1WA-g-8 |
| red-tailed hawk | Raptor | summer | WA | Klickitat | 1 | grassland | R1WA-g-8 | 2015-06-22T00:00:00Z | 0 | 1 | 1 | R1WA-g-8 |
| great horned owl | Raptor | summer | WA | Klickitat | 1 | grassland | R1WA-g-8 | 2015-06-22T00:00:00Z | 42 | 51 | 1 | R1WA-g-8 |
| red-tailed hawk | Raptor | summer | WA | Klickitat | 1 | grassland | R1WA-g-8 | 2015-06-22T00:00:00Z | 146 | 160 | 1 | R1WA-g-8 |
| turkey vulture | Raptor | summer | WA | Klickitat | 1 | grassland | R1WA-g-8 | 2015-06-22T00:00:00Z | 42 | 51 | 1 | R1WA-g-8 |
| ring-necked pheasant | Game Bird | summer | WA | Klickitat | 1 | grassland | R1WA-g-8 | 2015-06-22T00:00:00Z | 4 | 7 | 1 | R1WA-g-8 |
| mallard | Game Bird | summer | WA | Klickitat | 1 | grassland | R1WA-g-8 | 2015-06-22T00:00:00Z | 28 | 35 | 1 | R1WA-g-8 |
| barn owl | Raptor | summer | WA | Klickitat | 1 | grassland | R1WA-g-8 | 2015-06-22T00:00:00Z | 66 | 73 | 1 | R1WA-g-8 |
| red-tailed hawk | Raptor | summer | WA | Klickitat | 1 | grassland | R1WA-g-8 | 2015-06-22T00:00:00Z | 73 | 80 | 1 | R1WA-g-8 |
| ring-necked pheasant | Game Bird | summer | WA | Klickitat | 1 | grassland | R1WA-g-8 | 2015-06-22T00:00:00Z | 21 | 28 | 1 | R1WA-g-8 |
| great horned owl | Raptor | summer | WA | Klickitat | 1 | grassland | R1WA-g-8 | 2015-06-22T00:00:00Z | 28 | 35 | 1 | R1WA-g-8 |
| mallard | Game Bird | summer | WA | Klickitat | 1 | grassland | R1WA-g-8 | 2015-06-22T00:00:00Z | 28 | 35 | 1 | R1WA-g-8 |
| ring-necked pheasant | Game Bird | summer | WA | Klickitat | 1 | grassland | R1WA-g-8 | 2015-06-22T00:00:00Z | 3 | 4 | 1 | R1WA-g-8 |
| mallard | Game Bird | summer | WA | Klickitat | 1 | grassland | R1WA-g-8 | 2015-06-22T00:00:00Z | 28 | 35 | 1 | R1WA-g-8 |
| ring-necked pheasant | Game Bird | summer | WA | Klickitat | 1 | grassland | R1WA-g-8 | 2015-08-31T00:00:00Z | 56 | 63 | 1 | R1WA-g-8 |
| mallard | Game Bird | summer | WA | Klickitat | 1 | grassland | R1WA-g-8 | 2015-08-31T00:00:00Z | 4 | 7 | 1 | R1WA-g-8 |
| red-tailed hawk | Raptor | summer | WA | Klickitat | 1 | grassland | R1WA-g-8 | 2015-08-31T00:00:00Z | 70 | 70 | 1 | R1WA-g-8 |
| red-tailed hawk | Raptor | summer | WA | Klickitat | 1 | grassland | R1WA-g-8 | 2015-08-31T00:00:00Z | 42 | 49 | 1 | R1WA-g-8 |
| red-tailed hawk | Raptor | summer | WA | Klickitat | 1 | grassland | R1WA-g-8 | 2015-08-31T00:00:00Z | 0 | 1 | 1 | R1WA-g-8 |
| ring-necked pheasant | Game Bird | summer | WA | Klickitat | 1 | grassland | R1WA-g-8 | 2015-08-31T00:00:00Z | 70 | 84 | 1 | R1WA-g-8 |
| red-tailed hawk | Raptor | summer | WA | Klickitat | 1 | grassland | R1WA-g-8 | 2015-08-31T00:00:00Z | 1 | 2 | 1 | R1WA-g-8 |
| red-tailed hawk | Raptor | summer | WA | Klickitat | 1 | grassland | R1WA-g-8 | 2015-08-31T00:00:00Z | 84 | NA | 1 | R1WA-g-8 |
| mallard | Game Bird | summer | WA | Klickitat | 1 | grassland | R1WA-g-8 | 2015-08-31T00:00:00Z | 42 | 49 | 1 | R1WA-g-8 |
| ring-necked pheasant | Game Bird | summer | WA | Klickitat | 1 | grassland | R1WA-g-8 | 2015-08-31T00:00:00Z | 2 | 3 | 1 | R1WA-g-8 |
| red-tailed hawk | Raptor | summer | WA | Klickitat | 1 | grassland | R1WA-g-8 | 2015-08-31T00:00:00Z | 70 | 84 | 1 | R1WA-g-8 |
| mallard | Game Bird | summer | WA | Klickitat | 1 | grassland | R1WA-g-8 | 2015-08-31T00:00:00Z | 7 | 10 | 1 | R1WA-g-8 |
| ring-necked pheasant | Game Bird | summer | WA | Klickitat | 1 | grassland | R1WA-g-8 | 2015-08-31T00:00:00Z | 1 | 2 | 1 | R1WA-g-8 |
| mallard | Game Bird | summer | WA | Klickitat | 1 | grassland | R1WA-g-8 | 2015-08-31T00:00:00Z | 2 | 3 | 1 | R1WA-g-8 |
| ring-necked pheasant | Game Bird | summer | WA | Klickitat | 1 | grassland | R1WA-g-8 | 2015-08-31T00:00:00Z | 0 | 1 | 1 | R1WA-g-8 |
| red-tailed hawk | Raptor | summer | WA | Klickitat | 1 | grassland | R1WA-g-8 | 2015-08-31T00:00:00Z | 70 | 84 | 1 | R1WA-g-8 |
| mallard | Game Bird | summer | WA | Klickitat | 1 | grassland | R1WA-g-8 | 2015-08-31T00:00:00Z | 4 | 7 | 1 | R1WA-g-8 |
| red-tailed hawk | Raptor | summer | WA | Klickitat | 1 | grassland | R1WA-g-8 | 2015-08-31T00:00:00Z | 70 | 84 | 1 | R1WA-g-8 |
| red-tailed hawk | Raptor | summer | WA | Klickitat | 1 | grassland | R1WA-g-8 | 2015-08-31T00:00:00Z | 84 | NA | 1 | R1WA-g-8 |
| ring-necked pheasant | Game Bird | summer | WA | Klickitat | 1 | grassland | R1WA-g-8 | 2015-08-31T00:00:00Z | 7 | 10 | 1 | R1WA-g-8 |
| red-tailed hawk | Raptor | summer | WA | Klickitat | 1 | grassland | R1WA-g-8 | 2015-08-31T00:00:00Z | 1 | 2 | 1 | R1WA-g-8 |
| red-tailed hawk | Raptor | summer | WA | Klickitat | 1 | grassland | R1WA-g-8 | 2015-08-31T00:00:00Z | 70 | 84 | 1 | R1WA-g-8 |
| mallard | Game Bird | summer | WA | Klickitat | 1 | grassland | R1WA-g-8 | 2015-08-31T00:00:00Z | 1 | 2 | 1 | R1WA-g-8 |
| mallard | Game Bird | summer | WA | Klickitat | 1 | grassland | R1WA-g-8 | 2015-08-31T00:00:00Z | 56 | 63 | 1 | R1WA-g-8 |
| red-tailed hawk | Raptor | summer | WA | Klickitat | 1 | grassland | R1WA-g-8 | 2015-08-31T00:00:00Z | 70 | 84 | 1 | R1WA-g-8 |
| ring-necked pheasant | Game Bird | summer | WA | Klickitat | 1 | grassland | R1WA-g-8 | 2015-08-31T00:00:00Z | 42 | 49 | 1 | R1WA-g-8 |
| red-tailed hawk | Raptor | summer | WA | Klickitat | 1 | grassland | R1WA-g-8 | 2015-08-31T00:00:00Z | 4 | 7 | 1 | R1WA-g-8 |
| ring-necked pheasant | Game Bird | summer | WA | Klickitat | 1 | grassland | R1WA-g-8 | 2015-08-31T00:00:00Z | 17 | 21 | 1 | R1WA-g-8 |
| mallard | Game Bird | summer | WA | Klickitat | 1 | grassland | R1WA-g-8 | 2015-08-31T00:00:00Z | 0 | 1 | 1 | R1WA-g-8 |
| barn owl | Raptor | spring | WA | Kittitas | 1 | shrub/scrub | R1WA-ss-2 | 2016-05-14T00:00:00Z | 86 | 98 | 0 | R1WA-ss-2 |
| red-tailed hawk | Raptor | spring | WA | Kittitas | 1 | shrub/scrub | R1WA-ss-2 | 2016-05-14T00:00:00Z | 122 | NA | 0 | R1WA-ss-2 |
| barn owl | Raptor | spring | WA | Kittitas | 1 | shrub/scrub | R1WA-ss-2 | 2016-05-14T00:00:00Z | 86 | 98 | 0 | R1WA-ss-2 |
| red-tailed hawk | Raptor | spring | WA | Kittitas | 1 | shrub/scrub | R1WA-ss-2 | 2016-05-14T00:00:00Z | 122 | NA | 0 | R1WA-ss-2 |
| barn owl | Raptor | spring | WA | Kittitas | 1 | shrub/scrub | R1WA-ss-2 | 2016-05-14T00:00:00Z | 86 | 98 | 0 | R1WA-ss-2 |
| barn owl | Raptor | spring | WA | Kittitas | 1 | shrub/scrub | R1WA-ss-2 | 2016-05-14T00:00:00Z | 86 | 98 | 0 | R1WA-ss-2 |
| red-tailed hawk | Raptor | spring | WA | Kittitas | 1 | shrub/scrub | R1WA-ss-2 | 2016-05-14T00:00:00Z | 98 | 118 | 0 | R1WA-ss-2 |
| red-tailed hawk | Raptor | spring | WA | Kittitas | 1 | shrub/scrub | R1WA-ss-2 | 2016-05-14T00:00:00Z | 122 | NA | 0 | R1WA-ss-2 |
| great horned owl | Raptor | spring | WA | Kittitas | 1 | shrub/scrub | R1WA-ss-2 | 2016-05-14T00:00:00Z | 122 | NA | 0 | R1WA-ss-2 |
| barn owl | Raptor | spring | WA | Kittitas | 1 | shrub/scrub | R1WA-ss-2 | 2016-05-14T00:00:00Z | 122 | NA | 0 | R1WA-ss-2 |
| barn owl | Raptor | spring | WA | Kittitas | 1 | shrub/scrub | R1WA-ss-2 | 2016-05-14T00:00:00Z | 86 | 98 | 0 | R1WA-ss-2 |
| red-tailed hawk | Raptor | spring | WA | Kittitas | 1 | shrub/scrub | R1WA-ss-2 | 2016-05-14T00:00:00Z | 122 | NA | 0 | R1WA-ss-2 |
| barn owl | Raptor | spring | WA | Kittitas | 1 | shrub/scrub | R1WA-ss-2 | 2016-05-14T00:00:00Z | 34 | 46 | 0 | R1WA-ss-2 |
| red-tailed hawk | Raptor | spring | WA | Kittitas | 1 | shrub/scrub | R1WA-ss-2 | 2016-05-14T00:00:00Z | 34 | 46 | 0 | R1WA-ss-2 |
| barn owl | Raptor | spring | WA | Kittitas | 1 | shrub/scrub | R1WA-ss-2 | 2016-05-14T00:00:00Z | 122 | NA | 0 | R1WA-ss-2 |
| prairie falcon | Raptor | spring | WA | Kittitas | 1 | shrub/scrub | R1WA-ss-2 | 2016-05-14T00:00:00Z | 122 | NA | 0 | R1WA-ss-2 |
| barn owl | Raptor | spring | WA | Kittitas | 1 | shrub/scrub | R1WA-ss-2 | 2016-05-14T00:00:00Z | 98 | 118 | 0 | R1WA-ss-2 |
| barn owl | Raptor | spring | WA | Kittitas | 1 | shrub/scrub | R1WA-ss-2 | 2016-05-14T00:00:00Z | 86 | 98 | 0 | R1WA-ss-2 |
| mallard | Game Bird | winter | WA | Kittitas | 1 | shrub/scrub | R1WA-ss-4 | 2010-03-15T00:00:00Z | 40 | NA | 0 | R1WA-ss-4 |
| ring-necked pheasant | Game Bird | winter | WA | Kittitas | 1 | shrub/scrub | R1WA-ss-4 | 2010-03-15T00:00:00Z | 2 | 3 | 0 | R1WA-ss-4 |
| mallard | Game Bird | winter | WA | Kittitas | 1 | shrub/scrub | R1WA-ss-4 | 2010-03-15T00:00:00Z | 20 | 30 | 0 | R1WA-ss-4 |
| mallard | Game Bird | winter | WA | Kittitas | 1 | shrub/scrub | R1WA-ss-4 | 2010-03-15T00:00:00Z | 20 | 30 | 0 | R1WA-ss-4 |
| mallard | Game Bird | winter | WA | Kittitas | 1 | shrub/scrub | R1WA-ss-4 | 2010-03-15T00:00:00Z | 10 | 14 | 0 | R1WA-ss-4 |
| mallard | Game Bird | spring | WA | Kittitas | 1 | shrub/scrub | R1WA-ss-4 | 2010-04-26T00:00:00Z | 1 | 2 | 0 | R1WA-ss-4 |
| ring-necked pheasant | Game Bird | spring | WA | Kittitas | 1 | shrub/scrub | R1WA-ss-4 | 2010-04-26T00:00:00Z | 3 | 4 | 0 | R1WA-ss-4 |
| ring-necked pheasant | Game Bird | spring | WA | Kittitas | 1 | shrub/scrub | R1WA-ss-4 | 2010-04-26T00:00:00Z | 3 | 4 | 0 | R1WA-ss-4 |
| ring-necked pheasant | Game Bird | spring | WA | Kittitas | 1 | shrub/scrub | R1WA-ss-4 | 2010-04-26T00:00:00Z | 14 | 20 | 0 | R1WA-ss-4 |
| ring-necked pheasant | Game Bird | spring | WA | Kittitas | 1 | shrub/scrub | R1WA-ss-4 | 2010-04-26T00:00:00Z | 7 | 10 | 0 | R1WA-ss-4 |
| mallard | Game Bird | summer | WA | Kittitas | 1 | shrub/scrub | R1WA-ss-4 | 2010-06-21T00:00:00Z | 10 | 14 | 0 | R1WA-ss-4 |
| mallard | Game Bird | summer | WA | Kittitas | 1 | shrub/scrub | R1WA-ss-4 | 2010-06-21T00:00:00Z | 0 | NA | 0 | R1WA-ss-4 |
| ring-necked pheasant | Game Bird | summer | WA | Kittitas | 1 | shrub/scrub | R1WA-ss-4 | 2010-06-21T00:00:00Z | 3 | 4 | 0 | R1WA-ss-4 |
| ring-necked pheasant | Game Bird | summer | WA | Kittitas | 1 | shrub/scrub | R1WA-ss-4 | 2010-06-21T00:00:00Z | 7 | NA | 0 | R1WA-ss-4 |
| ring-necked pheasant | Game Bird | summer | WA | Kittitas | 1 | shrub/scrub | R1WA-ss-4 | 2010-06-21T00:00:00Z | 0 | 1 | 0 | R1WA-ss-4 |
| mallard | Game Bird | summer | WA | Kittitas | 1 | shrub/scrub | R1WA-ss-4 | 2010-06-21T00:00:00Z | 7 | 10 | 0 | R1WA-ss-4 |
| ring-necked pheasant | Game Bird | summer | WA | Kittitas | 1 | shrub/scrub | R1WA-ss-4 | 2010-06-21T00:00:00Z | 4 | 7 | 0 | R1WA-ss-4 |
| ring-necked pheasant | Game Bird | summer | WA | Kittitas | 1 | shrub/scrub | R1WA-ss-4 | 2010-08-02T00:00:00Z | 4 | 10 | 0 | R1WA-ss-4 |
| mallard | Game Bird | summer | WA | Kittitas | 1 | shrub/scrub | R1WA-ss-4 | 2010-08-02T00:00:00Z | 14 | 20 | 0 | R1WA-ss-4 |
| mallard | Game Bird | summer | WA | Kittitas | 1 | shrub/scrub | R1WA-ss-4 | 2010-08-02T00:00:00Z | 10 | 20 | 0 | R1WA-ss-4 |
| mallard | Game Bird | summer | WA | Kittitas | 1 | shrub/scrub | R1WA-ss-4 | 2010-08-02T00:00:00Z | 1 | 2 | 0 | R1WA-ss-4 |
| ring-necked pheasant | Game Bird | summer | WA | Kittitas | 1 | shrub/scrub | R1WA-ss-4 | 2010-08-02T00:00:00Z | 2 | 3 | 0 | R1WA-ss-4 |
| ring-necked pheasant | Game Bird | summer | WA | Kittitas | 1 | shrub/scrub | R1WA-ss-4 | 2010-08-02T00:00:00Z | 2 | 3 | 0 | R1WA-ss-4 |
| ring-necked pheasant | Game Bird | summer | WA | Kittitas | 1 | shrub/scrub | R1WA-ss-4 | 2010-08-02T00:00:00Z | 4 | 10 | 0 | R1WA-ss-4 |
| mallard | Game Bird | fall | WA | Kittitas | 1 | shrub/scrub | R1WA-ss-4 | 2010-11-08T00:00:00Z | 1 | 3 | 0 | R1WA-ss-4 |
| ring-necked pheasant | Game Bird | fall | WA | Kittitas | 1 | shrub/scrub | R1WA-ss-4 | 2010-11-08T00:00:00Z | 0 | 4 | 0 | R1WA-ss-4 |
| ring-necked pheasant | Game Bird | fall | WA | Kittitas | 1 | shrub/scrub | R1WA-ss-4 | 2010-11-08T00:00:00Z | 4 | 7 | 0 | R1WA-ss-4 |
| mallard | Game Bird | fall | WA | Kittitas | 1 | shrub/scrub | R1WA-ss-4 | 2010-11-08T00:00:00Z | 1 | 4 | 0 | R1WA-ss-4 |
| ring-necked pheasant | Game Bird | fall | WA | Kittitas | 1 | shrub/scrub | R1WA-ss-4 | 2010-11-08T00:00:00Z | 1 | 4 | 0 | R1WA-ss-4 |
| ring-necked pheasant | Game Bird | fall | WA | Kittitas | 1 | shrub/scrub | R1WA-ss-4 | 2010-11-08T00:00:00Z | 40 | NA | 0 | R1WA-ss-4 |
| ring-necked pheasant | Game Bird | fall | WA | Kittitas | 1 | shrub/scrub | R1WA-ss-4 | 2010-11-08T00:00:00Z | 1 | 4 | 0 | R1WA-ss-4 |
| mallard | Game Bird | fall | WA | Kittitas | 1 | shrub/scrub | R1WA-ss-4 | 2010-11-08T00:00:00Z | 1 | 4 | 0 | R1WA-ss-4 |
| ring-necked pheasant | Game Bird | fall | WA | Kittitas | 1 | shrub/scrub | R1WA-ss-4 | 2010-09-20T00:00:00Z | 14 | 20 | 0 | R1WA-ss-4 |
| ring-necked pheasant | Game Bird | fall | WA | Kittitas | 1 | shrub/scrub | R1WA-ss-4 | 2010-09-20T00:00:00Z | 14 | 20 | 0 | R1WA-ss-4 |
| ring-necked pheasant | Game Bird | fall | WA | Kittitas | 1 | shrub/scrub | R1WA-ss-4 | 2010-09-20T00:00:00Z | 4 | 7 | 0 | R1WA-ss-4 |
| ring-necked pheasant | Game Bird | fall | WA | Kittitas | 1 | shrub/scrub | R1WA-ss-4 | 2010-09-20T00:00:00Z | 7 | 10 | 0 | R1WA-ss-4 |
| mallard | Game Bird | fall | WA | Kittitas | 1 | shrub/scrub | R1WA-ss-4 | 2010-09-20T00:00:00Z | 14 | 20 | 0 | R1WA-ss-4 |
| ring-necked pheasant | Game Bird | fall | WA | Kittitas | 1 | shrub/scrub | R1WA-ss-4 | 2010-09-20T00:00:00Z | 40 | NA | 0 | R1WA-ss-4 |
| ring-necked pheasant | Game Bird | fall | WA | Kittitas | 1 | shrub/scrub | R1WA-ss-4 | 2010-09-20T00:00:00Z | 7 | 10 | 0 | R1WA-ss-4 |
| ring-necked pheasant | Game Bird | spring | WA | Kittitas | 1 | shrub/scrub | R1WA-ss-4 | 2012-04-16T00:00:00Z | 0 | 1 | 0 | R1WA-ss-4 |
| mallard | Game Bird | spring | WA | Kittitas | 1 | shrub/scrub | R1WA-ss-4 | 2012-04-16T00:00:00Z | 0 | 1 | 0 | R1WA-ss-4 |
| ring-necked pheasant | Game Bird | spring | WA | Kittitas | 1 | shrub/scrub | R1WA-ss-4 | 2012-04-16T00:00:00Z | 0 | 1 | 0 | R1WA-ss-4 |
| mallard | Game Bird | spring | WA | Kittitas | 1 | shrub/scrub | R1WA-ss-4 | 2012-04-16T00:00:00Z | 1 | 2 | 0 | R1WA-ss-4 |
| ring-necked pheasant | Game Bird | spring | WA | Kittitas | 1 | shrub/scrub | R1WA-ss-4 | 2012-04-16T00:00:00Z | 2 | 3 | 0 | R1WA-ss-4 |
| ring-necked pheasant | Game Bird | spring | WA | Kittitas | 1 | shrub/scrub | R1WA-ss-4 | 2012-04-16T00:00:00Z | 4 | 10 | 0 | R1WA-ss-4 |
| mallard | Game Bird | spring | WA | Kittitas | 1 | shrub/scrub | R1WA-ss-4 | 2012-04-16T00:00:00Z | 40 | NA | 0 | R1WA-ss-4 |
| ring-necked pheasant | Game Bird | spring | WA | Kittitas | 1 | shrub/scrub | R1WA-ss-4 | 2012-06-04T00:00:00Z | 0 | 1 | 0 | R1WA-ss-4 |
| mallard | Game Bird | spring | WA | Kittitas | 1 | shrub/scrub | R1WA-ss-4 | 2012-06-04T00:00:00Z | 0 | 1 | 0 | R1WA-ss-4 |
| mallard | Game Bird | spring | WA | Kittitas | 1 | shrub/scrub | R1WA-ss-4 | 2012-06-04T00:00:00Z | 1 | 2 | 0 | R1WA-ss-4 |
| ring-necked pheasant | Game Bird | spring | WA | Kittitas | 1 | shrub/scrub | R1WA-ss-4 | 2012-06-04T00:00:00Z | 4 | 7 | 0 | R1WA-ss-4 |
| mallard | Game Bird | spring | WA | Kittitas | 1 | shrub/scrub | R1WA-ss-4 | 2012-06-04T00:00:00Z | 40 | NA | 0 | R1WA-ss-4 |
| ring-necked pheasant | Game Bird | spring | WA | Kittitas | 1 | shrub/scrub | R1WA-ss-4 | 2012-06-04T00:00:00Z | 4 | 7 | 0 | R1WA-ss-4 |
| mallard | Game Bird | spring | WA | Kittitas | 1 | shrub/scrub | R1WA-ss-4 | 2012-06-04T00:00:00Z | 40 | NA | 0 | R1WA-ss-4 |
| ring-necked pheasant | Game Bird | spring | WA | Kittitas | 1 | shrub/scrub | R1WA-ss-4 | 2012-06-04T00:00:00Z | 14 | 20 | 0 | R1WA-ss-4 |
| mallard | Game Bird | spring | WA | Kittitas | 1 | shrub/scrub | R1WA-ss-4 | 2012-06-04T00:00:00Z | 40 | NA | 0 | R1WA-ss-4 |
| ring-necked pheasant | Game Bird | spring | WA | Kittitas | 1 | shrub/scrub | R1WA-ss-4 | 2012-06-04T00:00:00Z | 30 | 40 | 0 | R1WA-ss-4 |
| mallard | Game Bird | fall | WA | Kittitas | 1 | shrub/scrub | R1WA-ss-4 | 2012-11-13T00:00:00Z | 2 | 7 | 0 | R1WA-ss-4 |
| ring-necked pheasant | Game Bird | fall | WA | Kittitas | 1 | shrub/scrub | R1WA-ss-4 | 2012-11-13T00:00:00Z | 2 | 7 | 0 | R1WA-ss-4 |
| ring-necked pheasant | Game Bird | fall | WA | Kittitas | 1 | shrub/scrub | R1WA-ss-4 | 2012-11-13T00:00:00Z | 2 | 7 | 0 | R1WA-ss-4 |
| mallard | Game Bird | fall | WA | Kittitas | 1 | shrub/scrub | R1WA-ss-4 | 2012-11-13T00:00:00Z | 2 | 7 | 0 | R1WA-ss-4 |
| ring-necked pheasant | Game Bird | fall | WA | Kittitas | 1 | shrub/scrub | R1WA-ss-4 | 2012-11-13T00:00:00Z | 2 | 7 | 0 | R1WA-ss-4 |
| mallard | Game Bird | fall | WA | Kittitas | 1 | shrub/scrub | R1WA-ss-4 | 2012-11-13T00:00:00Z | 2 | 7 | 0 | R1WA-ss-4 |
| ring-necked pheasant | Game Bird | fall | WA | Kittitas | 1 | shrub/scrub | R1WA-ss-4 | 2012-11-13T00:00:00Z | 20 | 30 | 0 | R1WA-ss-4 |
| great horned owl | Raptor | summer | NY | Chautauqua | 5 | forest | R5NY-g-1 | 2020-06-15T00:00:00Z | 13.9909722 | 21.1645833 | 1 | R5NY-g-1 |
| red-tailed hawk | Raptor | summer | NY | Chautauqua | 5 | forest | R5NY-g-1 | 2020-06-15T00:00:00Z | 49.0854167 | 55.9833333 | 1 | R5NY-g-1 |
| osprey | Raptor | summer | NY | Chautauqua | 5 | forest | R5NY-g-1 | 2020-06-15T00:00:00Z | 48.9416667 | 55.9375 | 1 | R5NY-g-1 |
| Swainson's hawk | Raptor | summer | NY | Chautauqua | 5 | forest | R5NY-g-1 | 2020-06-15T00:00:00Z | 28.0465278 | 35.0694444 | 1 | R5NY-g-1 |
| osprey | Raptor | summer | NY | Chautauqua | 5 | forest | R5NY-g-1 | 2020-06-15T00:00:00Z | 28.025 | 35.1354167 | 1 | R5NY-g-1 |
| great horned owl | Raptor | summer | NY | Chautauqua | 5 | forest | R5NY-g-1 | 2020-06-15T00:00:00Z | 7.8319444 | 14.0729167 | 1 | R5NY-g-1 |
| prairie falcon | Raptor | summer | NY | Chautauqua | 5 | forest | R5NY-g-1 | 2020-06-15T00:00:00Z | 49.0944444 | 56.0777778 | 1 | R5NY-g-1 |
| red-tailed hawk | Raptor | summer | NY | Chautauqua | 5 | forest | R5NY-g-1 | 2020-06-15T00:00:00Z | 48.9951389 | 56 | 1 | R5NY-g-1 |
| mallard | Game Bird | summer | NY | Chautauqua | 5 | forest | R5NY-g-1 | 2020-06-15T00:00:00Z | 48.9847222 | 55.9930556 | 1 | R5NY-g-1 |
| red-tailed hawk | Raptor | summer | NY | Chautauqua | 5 | forest | R5NY-g-1 | 2020-06-15T00:00:00Z | 49.0868056 | 56.0756944 | 1 | R5NY-g-1 |
| mallard | Game Bird | summer | NY | Chautauqua | 5 | forest | R5NY-g-1 | 2020-06-15T00:00:00Z | 2.03125 | 3.0527778 | 1 | R5NY-g-1 |
| red-tailed hawk | Raptor | summer | NY | Chautauqua | 5 | forest | R5NY-g-1 | 2020-06-15T00:00:00Z | 48.9201389 | 55.9277778 | 1 | R5NY-g-1 |
| mallard | Game Bird | summer | NY | Chautauqua | 5 | forest | R5NY-g-1 | 2020-06-15T00:00:00Z | 5.0805556 | 7.8333333 | 1 | R5NY-g-1 |
| mallard | Game Bird | summer | NY | Chautauqua | 5 | forest | R5NY-g-1 | 2020-06-15T00:00:00Z | 8.8430556 | 10.8604167 | 1 | R5NY-g-1 |
| mallard | Game Bird | summer | NY | Chautauqua | 5 | forest | R5NY-g-1 | 2020-06-15T00:00:00Z | 0.8965278 | 0.8965278 | 1 | R5NY-g-1 |
| mallard | Game Bird | summer | NY | Chautauqua | 5 | forest | R5NY-g-1 | 2020-06-15T00:00:00Z | 1.0875 | 2.0527778 | 1 | R5NY-g-1 |
| mallard | Game Bird | summer | NY | Chautauqua | 5 | forest | R5NY-g-1 | 2020-06-15T00:00:00Z | 1.11875 | 1.11875 | 1 | R5NY-g-1 |
| mallard | Game Bird | summer | NY | Chautauqua | 5 | forest | R5NY-g-1 | 2020-06-15T00:00:00Z | 1.9729167 | 3.14375 | 1 | R5NY-g-1 |
| mallard | Game Bird | summer | NY | Chautauqua | 5 | forest | R5NY-g-1 | 2020-06-15T00:00:00Z | 1.9361111 | 3.0409722 | 1 | R5NY-g-1 |
| mallard | Game Bird | summer | NY | Chautauqua | 5 | forest | R5NY-g-1 | 2020-06-15T00:00:00Z | 0.9409722 | 1.9298611 | 1 | R5NY-g-1 |
| red-tailed hawk | Raptor | fall | NY | Chautauqua | 5 | forest | R5NY-g-1 | 2020-09-14T00:00:00Z | 21.9986111 | 28.0104167 | 1 | R5NY-g-1 |
| mallard | Game Bird | fall | NY | Chautauqua | 5 | forest | R5NY-g-1 | 2020-09-14T00:00:00Z | 10.83125 | 14.0472222 | 1 | R5NY-g-1 |
| mallard | Game Bird | fall | NY | Chautauqua | 5 | forest | R5NY-g-1 | 2020-09-14T00:00:00Z | 0.8277778 | 2.2972222 | 1 | R5NY-g-1 |
| red-tailed hawk | Raptor | fall | NY | Chautauqua | 5 | forest | R5NY-g-1 | 2020-09-14T00:00:00Z | 14.0659722 | 22.0284722 | 1 | R5NY-g-1 |
| osprey | Raptor | fall | NY | Chautauqua | 5 | forest | R5NY-g-1 | 2020-09-14T00:00:00Z | 6.9159722 | 6.9159722 | 1 | R5NY-g-1 |
| mallard | Game Bird | fall | NY | Chautauqua | 5 | forest | R5NY-g-1 | 2020-09-14T00:00:00Z | 5.0694444 | 6.925 | 1 | R5NY-g-1 |
| mallard | Game Bird | fall | NY | Chautauqua | 5 | forest | R5NY-g-1 | 2020-09-14T00:00:00Z | 4.1173611 | 4.9395833 | 1 | R5NY-g-1 |
| great horned owl | Raptor | fall | NY | Chautauqua | 5 | forest | R5NY-g-1 | 2020-09-14T00:00:00Z | 6.9548611 | 13.90625 | 1 | R5NY-g-1 |
| mallard | Game Bird | fall | NY | Chautauqua | 5 | forest | R5NY-g-1 | 2020-09-14T00:00:00Z | 0.9055556 | 0.9055556 | 1 | R5NY-g-1 |
| great horned owl | Raptor | fall | NY | Chautauqua | 5 | forest | R5NY-g-1 | 2020-09-14T00:00:00Z | 6.8666667 | 6.8666667 | 1 | R5NY-g-1 |
| great horned owl | Raptor | fall | NY | Chautauqua | 5 | forest | R5NY-g-1 | 2020-09-14T00:00:00Z | 27.9715278 | 34.9923611 | 1 | R5NY-g-1 |
| Cooper's hawk | Raptor | fall | NY | Chautauqua | 5 | forest | R5NY-g-1 | 2020-09-14T00:00:00Z | 14 | 20.8180556 | 1 | R5NY-g-1 |
| mallard | Game Bird | fall | NY | Chautauqua | 5 | forest | R5NY-g-1 | 2020-09-14T00:00:00Z | 3.9069444 | 4.9430556 | 1 | R5NY-g-1 |
| mallard | Game Bird | fall | NY | Chautauqua | 5 | forest | R5NY-g-1 | 2020-09-14T00:00:00Z | 1.7798611 | 2.8666667 | 1 | R5NY-g-1 |
| mallard | Game Bird | fall | NY | Chautauqua | 5 | forest | R5NY-g-1 | 2020-09-14T00:00:00Z | 3.9569444 | 4.7291667 | 1 | R5NY-g-1 |
| mallard | Game Bird | fall | NY | Chautauqua | 5 | forest | R5NY-g-1 | 2020-09-14T00:00:00Z | 3.8861111 | 4.7625 | 1 | R5NY-g-1 |
| Cooper's hawk | Raptor | fall | NY | Chautauqua | 5 | forest | R5NY-g-1 | 2020-09-14T00:00:00Z | 6.8069444 | 6.8069444 | 1 | R5NY-g-1 |
| great horned owl | Raptor | fall | NY | Chautauqua | 5 | forest | R5NY-g-1 | 2020-09-14T00:00:00Z | 6.73125 | 13.7173611 | 1 | R5NY-g-1 |
| mallard | Game Bird | fall | NY | Chautauqua | 5 | forest | R5NY-g-1 | 2020-09-14T00:00:00Z | 8.7479167 | 10.8277778 | 1 | R5NY-g-1 |
| barn owl | Raptor | fall | NY | Chautauqua | 5 | forest | R5NY-g-1 | 2020-09-14T00:00:00Z | 27.9576389 | 41.9798611 | 1 | R5NY-g-1 |
| ring-necked pheasant | Game Bird | winter | NY | Chautauqua | 5 | forest | R5NY-g-1 | 2020-12-08T00:00:00Z | 7.9041667 | 7.9041667 | 1 | R5NY-g-1 |
| red-tailed hawk | Raptor | winter | NY | Chautauqua | 5 | forest | R5NY-g-1 | 2020-12-08T00:00:00Z | 28.7972222 | 127.0805556 | 1 | R5NY-g-1 |
| ring-necked pheasant | Game Bird | winter | NY | Chautauqua | 5 | forest | R5NY-g-1 | 2020-12-08T00:00:00Z | 28.78125 | 127.0645833 | 1 | R5NY-g-1 |
| Cooper's hawk | Raptor | winter | NY | Chautauqua | 5 | forest | R5NY-g-1 | 2020-12-08T00:00:00Z | 28.7777778 | 127.0625 | 1 | R5NY-g-1 |
| ring-necked pheasant | Game Bird | winter | NY | Chautauqua | 5 | forest | R5NY-g-1 | 2020-12-08T00:00:00Z | 7.88125 | 7.88125 | 1 | R5NY-g-1 |
| red-tailed hawk | Raptor | winter | NY | Chautauqua | 5 | forest | R5NY-g-1 | 2020-12-08T00:00:00Z | 28.7673611 | 127.0548611 | 1 | R5NY-g-1 |
| ring-necked pheasant | Game Bird | winter | NY | Chautauqua | 5 | forest | R5NY-g-1 | 2020-12-08T00:00:00Z | 7.8854167 | 7.8854167 | 1 | R5NY-g-1 |
| Swainson's hawk | Raptor | winter | NY | Chautauqua | 5 | forest | R5NY-g-1 | 2020-12-08T00:00:00Z | 28.8381944 | 127.1284722 | 1 | R5NY-g-1 |
| ring-necked pheasant | Game Bird | winter | NY | Chautauqua | 5 | forest | R5NY-g-1 | 2020-12-08T00:00:00Z | 21.7784722 | 28.7923611 | 1 | R5NY-g-1 |
| great horned owl | Raptor | winter | NY | Chautauqua | 5 | forest | R5NY-g-1 | 2020-12-08T00:00:00Z | 7.8333333 | 7.8333333 | 1 | R5NY-g-1 |
| ring-necked pheasant | Game Bird | winter | NY | Chautauqua | 5 | forest | R5NY-g-1 | 2020-12-08T00:00:00Z | 7.8993056 | 7.8993056 | 1 | R5NY-g-1 |
| Swainson's hawk | Raptor | winter | NY | Chautauqua | 5 | forest | R5NY-g-1 | 2020-12-08T00:00:00Z | 28.8076389 | NA | 1 | R5NY-g-1 |
| ring-necked pheasant | Game Bird | winter | NY | Chautauqua | 5 | forest | R5NY-g-1 | 2020-12-08T00:00:00Z | 7.8958333 | 7.8958333 | 1 | R5NY-g-1 |
| Swainson's hawk | Raptor | winter | NY | Chautauqua | 5 | forest | R5NY-g-1 | 2020-12-08T00:00:00Z | 21.8625 | 28.8194444 | 1 | R5NY-g-1 |
| ring-necked pheasant | Game Bird | winter | NY | Chautauqua | 5 | forest | R5NY-g-1 | 2020-12-08T00:00:00Z | 14.8631944 | 21.8847222 | 1 | R5NY-g-1 |
| great horned owl | Raptor | winter | NY | Chautauqua | 5 | forest | R5NY-g-1 | 2020-12-08T00:00:00Z | 14.8611111 | 21.8756944 | 1 | R5NY-g-1 |
| ring-necked pheasant | Game Bird | winter | NY | Chautauqua | 5 | forest | R5NY-g-1 | 2020-12-08T00:00:00Z | 7.8847222 | 7.8847222 | 1 | R5NY-g-1 |
| red-tailed hawk | Raptor | winter | NY | Chautauqua | 5 | forest | R5NY-g-1 | 2020-12-08T00:00:00Z | 7.8895833 | 7.8895833 | 1 | R5NY-g-1 |
| ring-necked pheasant | Game Bird | winter | NY | Chautauqua | 5 | forest | R5NY-g-1 | 2020-12-08T00:00:00Z | 7.8847222 | 7.8847222 | 1 | R5NY-g-1 |
| great horned owl | Raptor | winter | NY | Chautauqua | 5 | forest | R5NY-g-1 | 2020-12-08T00:00:00Z | 7.8861111 | 7.8861111 | 1 | R5NY-g-1 |
| ring-necked pheasant | Game Bird | spring | NY | Chautauqua | 5 | forest | R5NY-g-1 | 2021-04-05T00:00:00Z | 3.8993056 | 7.1 | 1 | R5NY-g-1 |
| red-tailed hawk | Raptor | spring | NY | Chautauqua | 5 | forest | R5NY-g-1 | 2021-04-05T00:00:00Z | 7.0833333 | 7.0833333 | 1 | R5NY-g-1 |
| great horned owl | Raptor | spring | NY | Chautauqua | 5 | forest | R5NY-g-1 | 2021-04-05T00:00:00Z | 20.9979167 | 28.0152778 | 1 | R5NY-g-1 |
| ring-necked pheasant | Game Bird | spring | NY | Chautauqua | 5 | forest | R5NY-g-1 | 2021-04-05T00:00:00Z | 0.9604167 | 0.9604167 | 1 | R5NY-g-1 |
| great horned owl | Raptor | spring | NY | Chautauqua | 5 | forest | R5NY-g-1 | 2021-04-05T00:00:00Z | 14.0201389 | 20.95625 | 1 | R5NY-g-1 |
| ring-necked pheasant | Game Bird | spring | NY | Chautauqua | 5 | forest | R5NY-g-1 | 2021-04-05T00:00:00Z | 3.9652778 | 7.1430556 | 1 | R5NY-g-1 |
| Swainson's hawk | Raptor | spring | NY | Chautauqua | 5 | forest | R5NY-g-1 | 2021-04-05T00:00:00Z | 13.8840278 | 20.9506944 | 1 | R5NY-g-1 |
| ring-necked pheasant | Game Bird | spring | NY | Chautauqua | 5 | forest | R5NY-g-1 | 2021-04-05T00:00:00Z | 0.9486111 | 2.04375 | 1 | R5NY-g-1 |
| Swainson's hawk | Raptor | spring | NY | Chautauqua | 5 | forest | R5NY-g-1 | 2021-04-05T00:00:00Z | 41.9229167 | NA | 1 | R5NY-g-1 |
| ring-necked pheasant | Game Bird | spring | NY | Chautauqua | 5 | forest | R5NY-g-1 | 2021-04-05T00:00:00Z | 3.0493056 | 4.0277778 | 1 | R5NY-g-1 |
| Cooper's hawk | Raptor | spring | NY | Chautauqua | 5 | forest | R5NY-g-1 | 2021-04-05T00:00:00Z | 7.0493056 | 7.0493056 | 1 | R5NY-g-1 |
| ring-necked pheasant | Game Bird | spring | NY | Chautauqua | 5 | forest | R5NY-g-1 | 2021-04-05T00:00:00Z | 2.9138889 | 3.8604167 | 1 | R5NY-g-1 |
| ring-necked pheasant | Game Bird | spring | NY | Chautauqua | 5 | forest | R5NY-g-1 | 2021-04-05T00:00:00Z | 3.8631944 | 7.0631944 | 1 | R5NY-g-1 |
| great horned owl | Raptor | spring | NY | Chautauqua | 5 | forest | R5NY-g-1 | 2021-04-05T00:00:00Z | 7.0590278 | 7.0590278 | 1 | R5NY-g-1 |
| great horned owl | Raptor | spring | NY | Chautauqua | 5 | forest | R5NY-g-1 | 2021-04-05T00:00:00Z | 7.1729167 | 7.1729167 | 1 | R5NY-g-1 |
| ring-necked pheasant | Game Bird | spring | NY | Chautauqua | 5 | forest | R5NY-g-1 | 2021-04-05T00:00:00Z | 3.9979167 | 7.1736111 | 1 | R5NY-g-1 |
| barn owl | Raptor | spring | NY | Chautauqua | 5 | forest | R5NY-g-1 | 2021-04-05T00:00:00Z | 7.1319444 | 7.1319444 | 1 | R5NY-g-1 |
| ring-necked pheasant | Game Bird | spring | NY | Chautauqua | 5 | forest | R5NY-g-1 | 2021-04-05T00:00:00Z | 0.8854167 | 1.9368056 | 1 | R5NY-g-1 |
| great horned owl | Raptor | spring | NY | Chautauqua | 5 | forest | R5NY-g-1 | 2021-04-05T00:00:00Z | 7.2361111 | 7.2361111 | 1 | R5NY-g-1 |
| ring-necked pheasant | Game Bird | spring | NY | Chautauqua | 5 | forest | R5NY-g-1 | 2021-04-05T00:00:00Z | 0.9451389 | 0.9451389 | 1 | R5NY-g-1 |
| ring-necked pheasant | Game Bird | summer | NM | Torrance | 2 | grassland | R2NM-g-1 | 2020-06-15T00:00:00Z | 56.1729167 | NA | 1 | R2NM-g-1 |
| great horned owl | Raptor | summer | NM | Torrance | 2 | grassland | R2NM-g-1 | 2020-06-15T00:00:00Z | 56.1375 | NA | 1 | R2NM-g-1 |
| peregrine falcon | Raptor | summer | NM | Torrance | 2 | grassland | R2NM-g-1 | 2020-06-15T00:00:00Z | 48.8625 | NA | 1 | R2NM-g-1 |
| ring-necked pheasant | Game Bird | summer | NM | Torrance | 2 | grassland | R2NM-g-1 | 2020-06-15T00:00:00Z | 55.88125 | NA | 1 | R2NM-g-1 |
| prairie falcon | Raptor | summer | NM | Torrance | 2 | grassland | R2NM-g-1 | 2020-06-15T00:00:00Z | 55.7805556 | NA | 1 | R2NM-g-1 |
| Cooper's hawk | Raptor | summer | NM | Torrance | 2 | grassland | R2NM-g-1 | 2020-06-15T00:00:00Z | 55.7972222 | NA | 1 | R2NM-g-1 |
| ring-necked pheasant | Game Bird | summer | NM | Torrance | 2 | grassland | R2NM-g-1 | 2020-06-15T00:00:00Z | 55.8673611 | NA | 1 | R2NM-g-1 |
| ferruginous hawk | Raptor | summer | NM | Torrance | 2 | grassland | R2NM-g-1 | 2020-06-15T00:00:00Z | 55.9361111 | NA | 1 | R2NM-g-1 |
| ring-necked pheasant | Game Bird | summer | NM | Torrance | 2 | grassland | R2NM-g-1 | 2020-06-15T00:00:00Z | 55.9451389 | NA | 1 | R2NM-g-1 |
| osprey | Raptor | summer | NM | Torrance | 2 | grassland | R2NM-g-1 | 2020-06-15T00:00:00Z | 55.9930556 | NA | 1 | R2NM-g-1 |
| ring-necked pheasant | Game Bird | summer | NM | Torrance | 2 | grassland | R2NM-g-1 | 2020-06-15T00:00:00Z | 55.9368056 | NA | 1 | R2NM-g-1 |
| red-tailed hawk | Raptor | summer | NM | Torrance | 2 | grassland | R2NM-g-1 | 2020-06-15T00:00:00Z | 20.9673611 | 27.9777778 | 1 | R2NM-g-1 |
| red-tailed hawk | Raptor | summer | NM | Torrance | 2 | grassland | R2NM-g-1 | 2020-06-15T00:00:00Z | 20.9388889 | 27.9277778 | 1 | R2NM-g-1 |
| Swainson's hawk | Raptor | summer | NM | Torrance | 2 | grassland | R2NM-g-1 | 2020-06-15T00:00:00Z | 6.925 | 13.925 | 1 | R2NM-g-1 |
| ring-necked pheasant | Game Bird | summer | NM | Torrance | 2 | grassland | R2NM-g-1 | 2020-06-15T00:00:00Z | 10.25625 | 13.9638889 | 1 | R2NM-g-1 |
| ring-necked pheasant | Game Bird | summer | NM | Torrance | 2 | grassland | R2NM-g-1 | 2020-06-15T00:00:00Z | 4.0236111 | NA | 1 | R2NM-g-1 |
| ring-necked pheasant | Game Bird | summer | NM | Torrance | 2 | grassland | R2NM-g-1 | 2020-06-15T00:00:00Z | 0.7763889 | 2.0881944 | 1 | R2NM-g-1 |
| ring-necked pheasant | Game Bird | summer | NM | Torrance | 2 | grassland | R2NM-g-1 | 2020-06-15T00:00:00Z | 0.95 | 2.0548611 | 1 | R2NM-g-1 |
| ring-necked pheasant | Game Bird | summer | NM | Torrance | 2 | grassland | R2NM-g-1 | 2020-06-15T00:00:00Z | 0.9902778 | 2.0465278 | 1 | R2NM-g-1 |
| great horned owl | Raptor | summer | NM | Torrance | 2 | grassland | R2NM-g-1 | 2020-06-15T00:00:00Z | 6.86875 | 6.86875 | 1 | R2NM-g-1 |
| ring-necked pheasant | Game Bird | fall | NM | Torrance | 2 | grassland | R2NM-g-1 | 2020-09-14T00:00:00Z | 25.1833333 | 28.9902778 | 1 | R2NM-g-1 |
| ring-necked pheasant | Game Bird | fall | NM | Torrance | 2 | grassland | R2NM-g-1 | 2020-09-14T00:00:00Z | 21.1777778 | 25.0361111 | 1 | R2NM-g-1 |
| ring-necked pheasant | Game Bird | fall | NM | Torrance | 2 | grassland | R2NM-g-1 | 2020-09-14T00:00:00Z | 4.0868056 | 6.98125 | 1 | R2NM-g-1 |
| ring-necked pheasant | Game Bird | fall | NM | Torrance | 2 | grassland | R2NM-g-1 | 2020-09-14T00:00:00Z | 3.1777778 | 4.0618056 | 1 | R2NM-g-1 |
| ring-necked pheasant | Game Bird | fall | NM | Torrance | 2 | grassland | R2NM-g-1 | 2020-09-14T00:00:00Z | 3.0291667 | 3.8486111 | 1 | R2NM-g-1 |
| ring-necked pheasant | Game Bird | fall | NM | Torrance | 2 | grassland | R2NM-g-1 | 2020-09-14T00:00:00Z | 2.0069444 | 3.1520833 | 1 | R2NM-g-1 |
| ring-necked pheasant | Game Bird | fall | NM | Torrance | 2 | grassland | R2NM-g-1 | 2020-09-14T00:00:00Z | 1.0513889 | 2.0069444 | 1 | R2NM-g-1 |
| ring-necked pheasant | Game Bird | fall | NM | Torrance | 2 | grassland | R2NM-g-1 | 2020-09-14T00:00:00Z | 1.0715278 | 2.1680556 | 1 | R2NM-g-1 |
| ring-necked pheasant | Game Bird | fall | NM | Torrance | 2 | grassland | R2NM-g-1 | 2020-09-14T00:00:00Z | 1.0444444 | 1.0444444 | 1 | R2NM-g-1 |
| ring-necked pheasant | Game Bird | fall | NM | Torrance | 2 | grassland | R2NM-g-1 | 2020-09-14T00:00:00Z | 3.20625 | 4.0944444 | 1 | R2NM-g-1 |
| Cooper's hawk | Raptor | fall | NM | Torrance | 2 | grassland | R2NM-g-1 | 2020-09-14T00:00:00Z | 7.0166667 | 7.0166667 | 1 | R2NM-g-1 |
| prairie falcon | Raptor | fall | NM | Torrance | 2 | grassland | R2NM-g-1 | 2020-09-14T00:00:00Z | 7.0986111 | 7.0986111 | 1 | R2NM-g-1 |
| prairie falcon | Raptor | fall | NM | Torrance | 2 | grassland | R2NM-g-1 | 2020-09-14T00:00:00Z | 27.9576389 | 35.0840278 | 1 | R2NM-g-1 |
| barn owl | Raptor | fall | NM | Torrance | 2 | grassland | R2NM-g-1 | 2020-09-14T00:00:00Z | 35.1006944 | 45.9118056 | 1 | R2NM-g-1 |
| great horned owl | Raptor | fall | NM | Torrance | 2 | grassland | R2NM-g-1 | 2020-09-14T00:00:00Z | 35.0479167 | 46.1194444 | 1 | R2NM-g-1 |
| great horned owl | Raptor | fall | NM | Torrance | 2 | grassland | R2NM-g-1 | 2020-09-14T00:00:00Z | 55.9736111 | NA | 1 | R2NM-g-1 |
| Cooper's hawk | Raptor | fall | NM | Torrance | 2 | grassland | R2NM-g-1 | 2020-09-14T00:00:00Z | 56.0208333 | NA | 1 | R2NM-g-1 |
| great horned owl | Raptor | fall | NM | Torrance | 2 | grassland | R2NM-g-1 | 2020-09-14T00:00:00Z | 55.9430556 | NA | 1 | R2NM-g-1 |
| great horned owl | Raptor | fall | NM | Torrance | 2 | grassland | R2NM-g-1 | 2020-09-14T00:00:00Z | 55.9868056 | NA | 1 | R2NM-g-1 |
| great horned owl | Raptor | fall | NM | Torrance | 2 | grassland | R2NM-g-1 | 2020-09-14T00:00:00Z | 55.9895833 | NA | 1 | R2NM-g-1 |
| ring-necked pheasant | Game Bird | winter | NM | Torrance | 2 | grassland | R2NM-g-1 | 2020-12-07T00:00:00Z | 7.00625 | 7.00625 | 1 | R2NM-g-1 |
| Cooper's hawk | Raptor | winter | NM | Torrance | 2 | grassland | R2NM-g-1 | 2020-12-07T00:00:00Z | 6.9423611 | 6.9423611 | 1 | R2NM-g-1 |
| ring-necked pheasant | Game Bird | winter | NM | Torrance | 2 | grassland | R2NM-g-1 | 2020-12-07T00:00:00Z | 28.225 | 35.1833333 | 1 | R2NM-g-1 |
| great horned owl | Raptor | winter | NM | Torrance | 2 | grassland | R2NM-g-1 | 2020-12-07T00:00:00Z | 7.0069444 | 7.0069444 | 1 | R2NM-g-1 |
| ring-necked pheasant | Game Bird | winter | NM | Torrance | 2 | grassland | R2NM-g-1 | 2020-12-07T00:00:00Z | 28.1361111 | 35.0826389 | 1 | R2NM-g-1 |
| barred owl | Raptor | winter | NM | Torrance | 2 | grassland | R2NM-g-1 | 2020-12-07T00:00:00Z | 55.8951389 | NA | 1 | R2NM-g-1 |
| ring-necked pheasant | Game Bird | winter | NM | Torrance | 2 | grassland | R2NM-g-1 | 2020-12-07T00:00:00Z | 21.1027778 | 28.1020833 | 1 | R2NM-g-1 |
| prairie falcon | Raptor | winter | NM | Torrance | 2 | grassland | R2NM-g-1 | 2020-12-07T00:00:00Z | 41.9680556 | 48.8833333 | 1 | R2NM-g-1 |
| ring-necked pheasant | Game Bird | winter | NM | Torrance | 2 | grassland | R2NM-g-1 | 2020-12-07T00:00:00Z | 20.9479167 | 27.9340278 | 1 | R2NM-g-1 |
| great horned owl | Raptor | winter | NM | Torrance | 2 | grassland | R2NM-g-1 | 2020-12-07T00:00:00Z | 55.96875 | NA | 1 | R2NM-g-1 |
| red-tailed hawk | Raptor | winter | NM | Torrance | 2 | grassland | R2NM-g-1 | 2020-12-07T00:00:00Z | 14.1527778 | 20.9986111 | 1 | R2NM-g-1 |
| ring-necked pheasant | Game Bird | winter | NM | Torrance | 2 | grassland | R2NM-g-1 | 2020-12-07T00:00:00Z | 6.9680556 | 6.9680556 | 1 | R2NM-g-1 |
| ring-necked pheasant | Game Bird | winter | NM | Torrance | 2 | grassland | R2NM-g-1 | 2020-12-07T00:00:00Z | 14.0840278 | 20.9222222 | 1 | R2NM-g-1 |
| great horned owl | Raptor | winter | NM | Torrance | 2 | grassland | R2NM-g-1 | 2020-12-07T00:00:00Z | 6.9527778 | 6.9527778 | 1 | R2NM-g-1 |
| ring-necked pheasant | Game Bird | winter | NM | Torrance | 2 | grassland | R2NM-g-1 | 2020-12-07T00:00:00Z | 14.025 | 20.9944444 | 1 | R2NM-g-1 |
| red-tailed hawk | Raptor | winter | NM | Torrance | 2 | grassland | R2NM-g-1 | 2020-12-07T00:00:00Z | 55.88125 | NA | 1 | R2NM-g-1 |
| ring-necked pheasant | Game Bird | winter | NM | Torrance | 2 | grassland | R2NM-g-1 | 2020-12-07T00:00:00Z | 13.9277778 | 20.96875 | 1 | R2NM-g-1 |
| prairie falcon | Raptor | winter | NM | Torrance | 2 | grassland | R2NM-g-1 | 2020-12-07T00:00:00Z | 41.9541667 | 48.8652778 | 1 | R2NM-g-1 |
| ring-necked pheasant | Game Bird | winter | NM | Torrance | 2 | grassland | R2NM-g-1 | 2020-12-07T00:00:00Z | 55.8604167 | NA | 1 | R2NM-g-1 |
| ring-necked pheasant | Game Bird | spring | NM | Torrance | 2 | grassland | R2NM-g-1 | 2021-03-15T00:00:00Z | 7.0763889 | 7.0763889 | 1 | R2NM-g-1 |
| great horned owl | Raptor | winter | NM | Torrance | 2 | grassland | R2NM-g-1 | 2020-12-07T00:00:00Z | 55.8826389 | NA | 1 | R2NM-g-1 |
| ring-necked pheasant | Game Bird | spring | NM | Torrance | 2 | grassland | R2NM-g-1 | 2021-03-15T00:00:00Z | 6.9326389 | 6.9326389 | 1 | R2NM-g-1 |
| great horned owl | Raptor | spring | NM | Torrance | 2 | grassland | R2NM-g-1 | 2021-03-15T00:00:00Z | 56.0416667 | NA | 1 | R2NM-g-1 |
| Swainson's hawk | Raptor | spring | NM | Torrance | 2 | grassland | R2NM-g-1 | 2021-03-15T00:00:00Z | 55.8805556 | NA | 1 | R2NM-g-1 |
| ring-necked pheasant | Game Bird | spring | NM | Torrance | 2 | grassland | R2NM-g-1 | 2021-03-15T00:00:00Z | 35.0465278 | 42.0173611 | 1 | R2NM-g-1 |
| Swainson's hawk | Raptor | spring | NM | Torrance | 2 | grassland | R2NM-g-1 | 2021-03-15T00:00:00Z | 13.9291667 | 20.9319444 | 1 | R2NM-g-1 |
| ring-necked pheasant | Game Bird | spring | NM | Torrance | 2 | grassland | R2NM-g-1 | 2021-03-15T00:00:00Z | 6.9340278 | 6.9340278 | 1 | R2NM-g-1 |
| great horned owl | Raptor | spring | NM | Torrance | 2 | grassland | R2NM-g-1 | 2021-03-15T00:00:00Z | 55.9319444 | NA | 1 | R2NM-g-1 |
| great horned owl | Raptor | spring | NM | Torrance | 2 | grassland | R2NM-g-1 | 2021-03-15T00:00:00Z | 55.8923611 | NA | 1 | R2NM-g-1 |
| ring-necked pheasant | Game Bird | spring | NM | Torrance | 2 | grassland | R2NM-g-1 | 2021-03-15T00:00:00Z | 6.9604167 | 13.9222222 | 1 | R2NM-g-1 |
| Swainson's hawk | Raptor | spring | NM | Torrance | 2 | grassland | R2NM-g-1 | 2021-03-15T00:00:00Z | 55.98125 | NA | 1 | R2NM-g-1 |
| ring-necked pheasant | Game Bird | spring | NM | Torrance | 2 | grassland | R2NM-g-1 | 2021-03-15T00:00:00Z | 56.0354167 | NA | 1 | R2NM-g-1 |
| red-tailed hawk | Raptor | spring | NM | Torrance | 2 | grassland | R2NM-g-1 | 2021-03-15T00:00:00Z | 55.9645833 | NA | 1 | R2NM-g-1 |
| ring-necked pheasant | Game Bird | spring | NM | Torrance | 2 | grassland | R2NM-g-1 | 2021-03-15T00:00:00Z | 55.9847222 | NA | 1 | R2NM-g-1 |
| Swainson's hawk | Raptor | spring | NM | Torrance | 2 | grassland | R2NM-g-1 | 2021-03-15T00:00:00Z | 6.94375 | 13.9034722 | 1 | R2NM-g-1 |
| ring-necked pheasant | Game Bird | spring | NM | Torrance | 2 | grassland | R2NM-g-1 | 2021-03-15T00:00:00Z | 55.93125 | NA | 1 | R2NM-g-1 |
| ring-necked pheasant | Game Bird | spring | NM | Torrance | 2 | grassland | R2NM-g-1 | 2021-03-15T00:00:00Z | 55.7694444 | NA | 1 | R2NM-g-1 |
| great horned owl | Raptor | spring | NM | Torrance | 2 | grassland | R2NM-g-1 | 2021-03-15T00:00:00Z | 55.8076389 | NA | 1 | R2NM-g-1 |
| ring-necked pheasant | Game Bird | spring | NM | Torrance | 2 | grassland | R2NM-g-1 | 2021-03-15T00:00:00Z | 55.7729167 | NA | 1 | R2NM-g-1 |
| Cooper's hawk | Raptor | spring | NM | Torrance | 2 | grassland | R2NM-g-1 | 2021-03-15T00:00:00Z | 55.74375 | NA | 1 | R2NM-g-1 |
| ring-necked pheasant | Game Bird | summer | NE | Holt | 6 | grassland | R6NE-g-1 | 2020-06-15T00:00:00Z | 6.8395833 | 6.8395833 | 1 | R6NE-g-1 |
| red-tailed hawk | Raptor | summer | NE | Holt | 6 | grassland | R6NE-g-1 | 2020-06-15T00:00:00Z | 59.9694444 | NA | 1 | R6NE-g-1 |
| ring-necked pheasant | Game Bird | summer | NE | Holt | 6 | grassland | R6NE-g-1 | 2020-06-15T00:00:00Z | 6.7243056 | 6.7243056 | 1 | R6NE-g-1 |
| great horned owl | Raptor | summer | NE | Holt | 6 | grassland | R6NE-g-1 | 2020-06-15T00:00:00Z | 59.9506944 | NA | 1 | R6NE-g-1 |
| great horned owl | Raptor | summer | NE | Holt | 6 | grassland | R6NE-g-1 | 2020-06-15T00:00:00Z | 59.9451389 | NA | 1 | R6NE-g-1 |
| ring-necked pheasant | Game Bird | summer | NE | Holt | 6 | grassland | R6NE-g-1 | 2020-06-15T00:00:00Z | 59.94375 | NA | 1 | R6NE-g-1 |
| ring-necked pheasant | Game Bird | summer | NE | Holt | 6 | grassland | R6NE-g-1 | 2020-06-15T00:00:00Z | 59.9395833 | NA | 1 | R6NE-g-1 |
| prairie falcon | Raptor | summer | NE | Holt | 6 | grassland | R6NE-g-1 | 2020-06-15T00:00:00Z | 59.90625 | NA | 1 | R6NE-g-1 |
| ring-necked pheasant | Game Bird | summer | NE | Holt | 6 | grassland | R6NE-g-1 | 2020-06-15T00:00:00Z | 6.9458333 | 6.9458333 | 1 | R6NE-g-1 |
| ring-necked pheasant | Game Bird | summer | NE | Holt | 6 | grassland | R6NE-g-1 | 2020-06-15T00:00:00Z | 3.8708333 | 3.8708333 | 1 | R6NE-g-1 |
| osprey | Raptor | summer | NE | Holt | 6 | grassland | R6NE-g-1 | 2020-06-15T00:00:00Z | 59.6152778 | NA | 1 | R6NE-g-1 |
| red-tailed hawk | Raptor | summer | NE | Holt | 6 | grassland | R6NE-g-1 | 2020-06-15T00:00:00Z | 60.1270833 | NA | 1 | R6NE-g-1 |
| ring-necked pheasant | Game Bird | summer | NE | Holt | 6 | grassland | R6NE-g-1 | 2020-06-16T00:00:00Z | 59.9152778 | NA | 1 | R6NE-g-1 |
| Cooper's hawk | Raptor | summer | NE | Holt | 6 | grassland | R6NE-g-1 | 2020-06-16T00:00:00Z | 59.9215278 | NA | 1 | R6NE-g-1 |
| ring-necked pheasant | Game Bird | summer | NE | Holt | 6 | grassland | R6NE-g-1 | 2020-06-16T00:00:00Z | 35.0076389 | 42.0097222 | 1 | R6NE-g-1 |
| great horned owl | Raptor | summer | NE | Holt | 6 | grassland | R6NE-g-1 | 2020-06-16T00:00:00Z | 59.9201389 | NA | 1 | R6NE-g-1 |
| barn owl | Raptor | summer | NE | Holt | 6 | grassland | R6NE-g-1 | 2020-06-16T00:00:00Z | 59.9555556 | NA | 1 | R6NE-g-1 |
| Cooper's hawk | Raptor | summer | NE | Holt | 6 | grassland | R6NE-g-1 | 2020-06-16T00:00:00Z | 59.9791667 | NA | 1 | R6NE-g-1 |
| ring-necked pheasant | Game Bird | summer | NE | Holt | 6 | grassland | R6NE-g-1 | 2020-06-16T00:00:00Z | 6.9472222 | 6.9472222 | 1 | R6NE-g-1 |
| ring-necked pheasant | Game Bird | summer | NE | Holt | 6 | grassland | R6NE-g-1 | 2020-06-16T00:00:00Z | 59.9666667 | NA | 1 | R6NE-g-1 |
| red-tailed hawk | Raptor | fall | NE | Holt | 6 | grassland | R6NE-g-1 | 2020-09-14T00:00:00Z | 50.1652778 | 60.1243056 | 1 | R6NE-g-1 |
| ring-necked pheasant | Game Bird | fall | NE | Holt | 6 | grassland | R6NE-g-1 | 2020-09-14T00:00:00Z | 49.9305556 | NA | 1 | R6NE-g-1 |
| ring-necked pheasant | Game Bird | fall | NE | Holt | 6 | grassland | R6NE-g-1 | 2020-09-14T00:00:00Z | 6.9236111 | 6.9236111 | 1 | R6NE-g-1 |
| red-tailed hawk | Raptor | fall | NE | Holt | 6 | grassland | R6NE-g-1 | 2020-09-14T00:00:00Z | 49.8708333 | NA | 1 | R6NE-g-1 |
| red-tailed hawk | Raptor | fall | NE | Holt | 6 | grassland | R6NE-g-1 | 2020-09-14T00:00:00Z | 14.0756944 | 21.1111111 | 1 | R6NE-g-1 |
| ring-necked pheasant | Game Bird | fall | NE | Holt | 6 | grassland | R6NE-g-1 | 2020-09-14T00:00:00Z | 59.8194444 | NA | 1 | R6NE-g-1 |
| red-tailed hawk | Raptor | fall | NE | Holt | 6 | grassland | R6NE-g-1 | 2020-09-14T00:00:00Z | 3.1847222 | 3.8145833 | 1 | R6NE-g-1 |
| ring-necked pheasant | Game Bird | fall | NE | Holt | 6 | grassland | R6NE-g-1 | 2020-09-14T00:00:00Z | 6.9152778 | 6.9152778 | 1 | R6NE-g-1 |
| red-tailed hawk | Raptor | fall | NE | Holt | 6 | grassland | R6NE-g-1 | 2020-09-14T00:00:00Z | 59.7826389 | NA | 1 | R6NE-g-1 |
| red-tailed hawk | Raptor | fall | NE | Holt | 6 | grassland | R6NE-g-1 | 2020-09-14T00:00:00Z | 49.7888889 | NA | 1 | R6NE-g-1 |
| ring-necked pheasant | Game Bird | fall | NE | Holt | 6 | grassland | R6NE-g-1 | 2020-09-14T00:00:00Z | 6.9222222 | 6.9222222 | 1 | R6NE-g-1 |
| great horned owl | Raptor | fall | NE | Holt | 6 | grassland | R6NE-g-1 | 2020-09-14T00:00:00Z | 59.7604167 | NA | 1 | R6NE-g-1 |
| ring-necked pheasant | Game Bird | fall | NE | Holt | 6 | grassland | R6NE-g-1 | 2020-09-14T00:00:00Z | 59.7590278 | NA | 1 | R6NE-g-1 |
| ring-necked pheasant | Game Bird | fall | NE | Holt | 6 | grassland | R6NE-g-1 | 2020-09-14T00:00:00Z | 49.7909722 | 59.7590278 | 1 | R6NE-g-1 |
| ring-necked pheasant | Game Bird | fall | NE | Holt | 6 | grassland | R6NE-g-1 | 2020-09-14T00:00:00Z | 6.9298611 | 6.9298611 | 1 | R6NE-g-1 |
| great horned owl | Raptor | fall | NE | Holt | 6 | grassland | R6NE-g-1 | 2020-09-14T00:00:00Z | 49.7895833 | NA | 1 | R6NE-g-1 |
| ring-necked pheasant | Game Bird | fall | NE | Holt | 6 | grassland | R6NE-g-1 | 2020-09-14T00:00:00Z | 59.74375 | NA | 1 | R6NE-g-1 |
| barn owl | Raptor | fall | NE | Holt | 6 | grassland | R6NE-g-1 | 2020-09-14T00:00:00Z | 59.7506944 | NA | 1 | R6NE-g-1 |
| ring-necked pheasant | Game Bird | fall | NE | Holt | 6 | grassland | R6NE-g-1 | 2020-09-14T00:00:00Z | 6.9604167 | 13.6104167 | 1 | R6NE-g-1 |
| great horned owl | Raptor | fall | NE | Holt | 6 | grassland | R6NE-g-1 | 2020-09-14T00:00:00Z | 49.7784722 | NA | 1 | R6NE-g-1 |
| ring-necked pheasant | Game Bird | winter | NE | Holt | 6 | grassland | R6NE-g-1 | 2020-12-07T00:00:00Z | 7.0416667 | 7.0416667 | 1 | R6NE-g-1 |
| great horned owl | Raptor | winter | NE | Holt | 6 | grassland | R6NE-g-1 | 2020-12-07T00:00:00Z | 7.0868056 | 7.0868056 | 1 | R6NE-g-1 |
| ring-necked pheasant | Game Bird | winter | NE | Holt | 6 | grassland | R6NE-g-1 | 2020-12-07T00:00:00Z | 7.0236111 | 7.0236111 | 1 | R6NE-g-1 |
| red-tailed hawk | Raptor | winter | NE | Holt | 6 | grassland | R6NE-g-1 | 2020-12-07T00:00:00Z | 41.9020833 | NA | 1 | R6NE-g-1 |
| Swainson's hawk | Raptor | winter | NE | Holt | 6 | grassland | R6NE-g-1 | 2020-12-07T00:00:00Z | 41.8791667 | 56.9479167 | 1 | R6NE-g-1 |
| barn owl | Raptor | winter | NE | Holt | 6 | grassland | R6NE-g-1 | 2020-12-07T00:00:00Z | 56.9298611 | NA | 1 | R6NE-g-1 |
| ring-necked pheasant | Game Bird | winter | NE | Holt | 6 | grassland | R6NE-g-1 | 2020-12-07T00:00:00Z | 6.9055556 | 13.9256944 | 1 | R6NE-g-1 |
| Swainson's hawk | Raptor | winter | NE | Holt | 6 | grassland | R6NE-g-1 | 2020-12-07T00:00:00Z | 41.8826389 | NA | 1 | R6NE-g-1 |
| ring-necked pheasant | Game Bird | winter | NE | Holt | 6 | grassland | R6NE-g-1 | 2020-12-07T00:00:00Z | 6.8618056 | 6.8618056 | 1 | R6NE-g-1 |
| Cooper's hawk | Raptor | winter | NE | Holt | 6 | grassland | R6NE-g-1 | 2020-12-07T00:00:00Z | 41.8506944 | NA | 1 | R6NE-g-1 |
| great horned owl | Raptor | winter | NE | Holt | 6 | grassland | R6NE-g-1 | 2020-12-07T00:00:00Z | 41.7840278 | NA | 1 | R6NE-g-1 |
| ring-necked pheasant | Game Bird | winter | NE | Holt | 6 | grassland | R6NE-g-1 | 2020-12-07T00:00:00Z | 6.8034722 | 6.8034722 | 1 | R6NE-g-1 |
| ring-necked pheasant | Game Bird | winter | NE | Holt | 6 | grassland | R6NE-g-1 | 2020-12-07T00:00:00Z | 6.9618056 | 13.7847222 | 1 | R6NE-g-1 |
| ring-necked pheasant | Game Bird | winter | NE | Holt | 6 | grassland | R6NE-g-1 | 2020-12-07T00:00:00Z | 6.9472222 | 6.9472222 | 1 | R6NE-g-1 |
| ring-necked pheasant | Game Bird | winter | NE | Holt | 6 | grassland | R6NE-g-1 | 2020-12-07T00:00:00Z | 6.9375 | 13.7819444 | 1 | R6NE-g-1 |
| ring-necked pheasant | Game Bird | winter | NE | Holt | 6 | grassland | R6NE-g-1 | 2020-12-07T00:00:00Z | 6.9277778 | 6.9277778 | 1 | R6NE-g-1 |
| ring-necked pheasant | Game Bird | winter | NE | Holt | 6 | grassland | R6NE-g-1 | 2020-12-07T00:00:00Z | 6.9145833 | 6.9145833 | 1 | R6NE-g-1 |
| great horned owl | Raptor | winter | NE | Holt | 6 | grassland | R6NE-g-1 | 2020-12-07T00:00:00Z | 48.9694444 | NA | 1 | R6NE-g-1 |
| red-tailed hawk | Raptor | winter | NE | Holt | 6 | grassland | R6NE-g-1 | 2020-12-07T00:00:00Z | 49.0013889 | NA | 1 | R6NE-g-1 |
| great horned owl | Raptor | winter | NE | Holt | 6 | grassland | R6NE-g-1 | 2020-12-07T00:00:00Z | 49.04375 | NA | 1 | R6NE-g-1 |
| Swainson's hawk | Raptor | spring | NE | Holt | 6 | grassland | R6NE-g-1 | 2021-03-16T00:00:00Z | 19.9833333 | 26.9284722 | 1 | R6NE-g-1 |
| Swainson's hawk | Raptor | spring | NE | Holt | 6 | grassland | R6NE-g-1 | 2021-03-16T00:00:00Z | 19.9840278 | 26.9270833 | 1 | R6NE-g-1 |
| red-tailed hawk | Raptor | spring | NE | Holt | 6 | grassland | R6NE-g-1 | 2021-03-16T00:00:00Z | 19.9861111 | 26.9263889 | 1 | R6NE-g-1 |
| great horned owl | Raptor | spring | NE | Holt | 6 | grassland | R6NE-g-1 | 2021-03-16T00:00:00Z | 54.9881944 | NA | 1 | R6NE-g-1 |
| ring-necked pheasant | Game Bird | spring | NE | Holt | 6 | grassland | R6NE-g-1 | 2021-03-16T00:00:00Z | 6.0125 | 6.0125 | 1 | R6NE-g-1 |
| ring-necked pheasant | Game Bird | spring | NE | Holt | 6 | grassland | R6NE-g-1 | 2021-03-16T00:00:00Z | 6.0104167 | 6.0104167 | 1 | R6NE-g-1 |
| red-tailed hawk | Raptor | spring | NE | Holt | 6 | grassland | R6NE-g-1 | 2021-03-16T00:00:00Z | 40.9576389 | 47.9013889 | 1 | R6NE-g-1 |
| great horned owl | Raptor | spring | NE | Holt | 6 | grassland | R6NE-g-1 | 2021-03-16T00:00:00Z | 54.9791667 | NA | 1 | R6NE-g-1 |
| red-tailed hawk | Raptor | spring | NE | Holt | 6 | grassland | R6NE-g-1 | 2021-03-16T00:00:00Z | 33.9402778 | 40.9576389 | 1 | R6NE-g-1 |
| great horned owl | Raptor | spring | NE | Holt | 6 | grassland | R6NE-g-1 | 2021-03-16T00:00:00Z | 54.9756944 | NA | 1 | R6NE-g-1 |
| ring-necked pheasant | Game Bird | spring | NE | Holt | 6 | grassland | R6NE-g-1 | 2021-03-16T00:00:00Z | 6.0194444 | 6.0194444 | 1 | R6NE-g-1 |
| great horned owl | Raptor | spring | NE | Holt | 6 | grassland | R6NE-g-1 | 2021-03-16T00:00:00Z | 40.9451389 | 47.8881944 | 1 | R6NE-g-1 |
| ring-necked pheasant | Game Bird | spring | NE | Holt | 6 | grassland | R6NE-g-1 | 2021-03-16T00:00:00Z | 6.025 | 6.025 | 1 | R6NE-g-1 |
| ring-necked pheasant | Game Bird | spring | NE | Holt | 6 | grassland | R6NE-g-1 | 2021-03-16T00:00:00Z | 6.0277778 | 6.0277778 | 1 | R6NE-g-1 |
| red-tailed hawk | Raptor | spring | NE | Holt | 6 | grassland | R6NE-g-1 | 2021-03-16T00:00:00Z | 6.0263889 | 6.0263889 | 1 | R6NE-g-1 |
| ring-necked pheasant | Game Bird | spring | NE | Holt | 6 | grassland | R6NE-g-1 | 2021-03-16T00:00:00Z | 6.0291667 | 6.0291667 | 1 | R6NE-g-1 |
| ring-necked pheasant | Game Bird | spring | NE | Holt | 6 | grassland | R6NE-g-1 | 2021-03-16T00:00:00Z | 6.0298611 | 6.0298611 | 1 | R6NE-g-1 |
| ring-necked pheasant | Game Bird | spring | NE | Holt | 6 | grassland | R6NE-g-1 | 2021-03-16T00:00:00Z | 5.8680556 | 5.8680556 | 1 | R6NE-g-1 |
| ring-necked pheasant | Game Bird | spring | NE | Holt | 6 | grassland | R6NE-g-1 | 2021-03-16T00:00:00Z | 5.875 | 5.875 | 1 | R6NE-g-1 |
| ring-necked pheasant | Game Bird | summer | TX | Hale | 2 | cropland | R2TX-c-1 | 2020-06-08T00:00:00Z | 9.8951389 | 13.8819444 | 1 | R2TX-c-1 |
| ring-necked pheasant | Game Bird | summer | TX | Hale | 2 | cropland | R2TX-c-1 | 2020-06-08T00:00:00Z | 29.8819444 | 42.0951389 | 1 | R2TX-c-1 |
| ring-necked pheasant | Game Bird | summer | TX | Hale | 2 | cropland | R2TX-c-1 | 2020-06-08T00:00:00Z | 1.8826389 | 2.9145833 | 1 | R2TX-c-1 |
| ring-necked pheasant | Game Bird | summer | TX | Hale | 2 | cropland | R2TX-c-1 | 2020-06-08T00:00:00Z | 2.9173611 | 3.9298611 | 1 | R2TX-c-1 |
| ring-necked pheasant | Game Bird | summer | TX | Hale | 2 | cropland | R2TX-c-1 | 2020-06-08T00:00:00Z | 1.8763889 | 2.9104167 | 1 | R2TX-c-1 |
| ring-necked pheasant | Game Bird | summer | TX | Hale | 2 | cropland | R2TX-c-1 | 2020-06-08T00:00:00Z | 1.9416667 | 2.9027778 | 1 | R2TX-c-1 |
| ring-necked pheasant | Game Bird | summer | TX | Hale | 2 | cropland | R2TX-c-1 | 2020-06-08T00:00:00Z | 1 | 2.8986111 | 1 | R2TX-c-1 |
| ring-necked pheasant | Game Bird | summer | TX | Hale | 2 | cropland | R2TX-c-1 | 2020-06-08T00:00:00Z | 2.0006944 | 2.8986111 | 1 | R2TX-c-1 |
| ring-necked pheasant | Game Bird | summer | TX | Hale | 2 | cropland | R2TX-c-1 | 2020-06-08T00:00:00Z | 2.9 | 3.9069444 | 1 | R2TX-c-1 |
| ring-necked pheasant | Game Bird | summer | TX | Hale | 2 | cropland | R2TX-c-1 | 2020-06-08T00:00:00Z | 6.8118056 | 9.8034722 | 1 | R2TX-c-1 |
| osprey | Raptor | summer | TX | Hale | 2 | cropland | R2TX-c-1 | 2020-06-15T00:00:00Z | 41.9375 | 50.8534722 | 1 | R2TX-c-1 |
| rough-legged hawk | Raptor | summer | TX | Hale | 2 | cropland | R2TX-c-1 | 2020-06-15T00:00:00Z | 55.8631944 | NA | 1 | R2TX-c-1 |
| barn owl | Raptor | summer | TX | Hale | 2 | cropland | R2TX-c-1 | 2020-06-15T00:00:00Z | 6.9756944 | 6.9756944 | 1 | R2TX-c-1 |
| great horned owl | Raptor | summer | TX | Hale | 2 | cropland | R2TX-c-1 | 2020-06-15T00:00:00Z | 56.0555556 | NA | 1 | R2TX-c-1 |
| great horned owl | Raptor | summer | TX | Hale | 2 | cropland | R2TX-c-1 | 2020-06-15T00:00:00Z | 56.0173611 | NA | 1 | R2TX-c-1 |
| Swainson's hawk | Raptor | summer | TX | Hale | 2 | cropland | R2TX-c-1 | 2020-06-15T00:00:00Z | 55.9166667 | NA | 1 | R2TX-c-1 |
| red-tailed hawk | Raptor | summer | TX | Hale | 2 | cropland | R2TX-c-1 | 2020-06-15T00:00:00Z | 55.8888889 | NA | 1 | R2TX-c-1 |
| great horned owl | Raptor | summer | TX | Hale | 2 | cropland | R2TX-c-1 | 2020-06-15T00:00:00Z | 55.9409722 | NA | 1 | R2TX-c-1 |
| Cooper's hawk | Raptor | summer | TX | Hale | 2 | cropland | R2TX-c-1 | 2020-06-15T00:00:00Z | 56.0972222 | NA | 1 | R2TX-c-1 |
| great horned owl | Raptor | summer | TX | Hale | 2 | cropland | R2TX-c-1 | 2020-06-15T00:00:00Z | 56.0416667 | NA | 1 | R2TX-c-1 |
| Swainson's hawk | Raptor | fall | TX | Hale | 2 | cropland | R2TX-c-1 | 2020-09-14T00:00:00Z | 51.1840278 | NA | 1 | R2TX-c-1 |
| red-tailed hawk | Raptor | fall | TX | Hale | 2 | cropland | R2TX-c-1 | 2020-09-14T00:00:00Z | 55.9645833 | NA | 1 | R2TX-c-1 |
| red-tailed hawk | Raptor | fall | TX | Hale | 2 | cropland | R2TX-c-1 | 2020-09-14T00:00:00Z | 55.9673611 | NA | 1 | R2TX-c-1 |
| red-tailed hawk | Raptor | fall | TX | Hale | 2 | cropland | R2TX-c-1 | 2020-09-14T00:00:00Z | 55.9652778 | NA | 1 | R2TX-c-1 |
| Cooper's hawk | Raptor | fall | TX | Hale | 2 | cropland | R2TX-c-1 | 2020-09-14T00:00:00Z | 35.0805556 | 42.1069444 | 1 | R2TX-c-1 |
| great horned owl | Raptor | fall | TX | Hale | 2 | cropland | R2TX-c-1 | 2020-09-14T00:00:00Z | 55.9729167 | NA | 1 | R2TX-c-1 |
| great horned owl | Raptor | fall | TX | Hale | 2 | cropland | R2TX-c-1 | 2020-09-14T00:00:00Z | 55.9756944 | NA | 1 | R2TX-c-1 |
| Cooper's hawk | Raptor | fall | TX | Hale | 2 | cropland | R2TX-c-1 | 2020-09-14T00:00:00Z | 42.0486111 | 49.0243056 | 1 | R2TX-c-1 |
| red-tailed hawk | Raptor | fall | TX | Hale | 2 | cropland | R2TX-c-1 | 2020-09-14T00:00:00Z | 56.03125 | NA | 1 | R2TX-c-1 |
| red-tailed hawk | Raptor | fall | TX | Hale | 2 | cropland | R2TX-c-1 | 2020-09-14T00:00:00Z | 6.9625 | 6.9625 | 1 | R2TX-c-1 |
| ring-necked pheasant | Game Bird | fall | TX | Hale | 2 | cropland | R2TX-c-1 | 2020-09-14T00:00:00Z | 2.9451389 | 3.8791667 | 1 | R2TX-c-1 |
| ring-necked pheasant | Game Bird | fall | TX | Hale | 2 | cropland | R2TX-c-1 | 2020-09-14T00:00:00Z | 55.8958333 | NA | 1 | R2TX-c-1 |
| ring-necked pheasant | Game Bird | fall | TX | Hale | 2 | cropland | R2TX-c-1 | 2020-09-14T00:00:00Z | 1.7881944 | 2.8875 | 1 | R2TX-c-1 |
| ring-necked pheasant | Game Bird | fall | TX | Hale | 2 | cropland | R2TX-c-1 | 2020-09-14T00:00:00Z | 0.7916667 | 1.78125 | 1 | R2TX-c-1 |
| ring-necked pheasant | Game Bird | fall | TX | Hale | 2 | cropland | R2TX-c-1 | 2020-09-14T00:00:00Z | 6.9236111 | 9.9298611 | 1 | R2TX-c-1 |
| ring-necked pheasant | Game Bird | fall | TX | Hale | 2 | cropland | R2TX-c-1 | 2020-09-14T00:00:00Z | 6.88125 | 9.8861111 | 1 | R2TX-c-1 |
| ring-necked pheasant | Game Bird | fall | TX | Hale | 2 | cropland | R2TX-c-1 | 2020-09-14T00:00:00Z | 1.7569444 | 2.7888889 | 1 | R2TX-c-1 |
| ring-necked pheasant | Game Bird | fall | TX | Hale | 2 | cropland | R2TX-c-1 | 2020-09-14T00:00:00Z | 3.7826389 | 6.7611111 | 1 | R2TX-c-1 |
| ring-necked pheasant | Game Bird | fall | TX | Hale | 2 | cropland | R2TX-c-1 | 2020-09-14T00:00:00Z | 29.825 | 34.9430556 | 1 | R2TX-c-1 |
| ring-necked pheasant | Game Bird | fall | TX | Hale | 2 | cropland | R2TX-c-1 | 2020-09-14T00:00:00Z | 29.7777778 | 34.9409722 | 1 | R2TX-c-1 |
| ring-necked pheasant | Game Bird | winter | TX | Hale | 2 | cropland | R2TX-c-1 | 2020-12-07T00:00:00Z | 6.9604167 | 6.9604167 | 1 | R2TX-c-1 |
| red-tailed hawk | Raptor | winter | TX | Hale | 2 | cropland | R2TX-c-1 | 2020-12-07T00:00:00Z | 42.9513889 | 48.9555556 | 1 | R2TX-c-1 |
| ring-necked pheasant | Game Bird | winter | TX | Hale | 2 | cropland | R2TX-c-1 | 2020-12-07T00:00:00Z | 6.9631944 | 6.9631944 | 1 | R2TX-c-1 |
| great horned owl | Raptor | winter | TX | Hale | 2 | cropland | R2TX-c-1 | 2020-12-07T00:00:00Z | 55.9402778 | NA | 1 | R2TX-c-1 |
| ring-necked pheasant | Game Bird | winter | TX | Hale | 2 | cropland | R2TX-c-1 | 2020-12-07T00:00:00Z | 6.9631944 | 6.9631944 | 1 | R2TX-c-1 |
| Swainson's hawk | Raptor | winter | TX | Hale | 2 | cropland | R2TX-c-1 | 2020-12-07T00:00:00Z | 6.9715278 | 13.9520833 | 1 | R2TX-c-1 |
| ring-necked pheasant | Game Bird | winter | TX | Hale | 2 | cropland | R2TX-c-1 | 2020-12-07T00:00:00Z | 6.9743056 | 6.9743056 | 1 | R2TX-c-1 |
| great horned owl | Raptor | winter | TX | Hale | 2 | cropland | R2TX-c-1 | 2020-12-07T00:00:00Z | 55.9354167 | NA | 1 | R2TX-c-1 |
| ring-necked pheasant | Game Bird | winter | TX | Hale | 2 | cropland | R2TX-c-1 | 2020-12-07T00:00:00Z | 55.9381944 | NA | 1 | R2TX-c-1 |
| barn owl | Raptor | winter | TX | Hale | 2 | cropland | R2TX-c-1 | 2020-12-07T00:00:00Z | 34.94375 | 42.9416667 | 1 | R2TX-c-1 |
| ring-necked pheasant | Game Bird | winter | TX | Hale | 2 | cropland | R2TX-c-1 | 2020-12-07T00:00:00Z | 6.9652778 | 6.9652778 | 1 | R2TX-c-1 |
| Cooper's hawk | Raptor | winter | TX | Hale | 2 | cropland | R2TX-c-1 | 2020-12-07T00:00:00Z | 55.9236111 | NA | 1 | R2TX-c-1 |
| ring-necked pheasant | Game Bird | winter | TX | Hale | 2 | cropland | R2TX-c-1 | 2020-12-07T00:00:00Z | 6.9576389 | 6.9576389 | 1 | R2TX-c-1 |
| red-tailed hawk | Raptor | winter | TX | Hale | 2 | cropland | R2TX-c-1 | 2020-12-07T00:00:00Z | 6.9590278 | 6.9590278 | 1 | R2TX-c-1 |
| ring-necked pheasant | Game Bird | winter | TX | Hale | 2 | cropland | R2TX-c-1 | 2020-12-07T00:00:00Z | 13.8930556 | 20.9020833 | 1 | R2TX-c-1 |
| red-tailed hawk | Raptor | winter | TX | Hale | 2 | cropland | R2TX-c-1 | 2020-12-07T00:00:00Z | 6.9638889 | 13.8972222 | 1 | R2TX-c-1 |
| ring-necked pheasant | Game Bird | winter | TX | Hale | 2 | cropland | R2TX-c-1 | 2020-12-07T00:00:00Z | 42.8826389 | 48.88125 | 1 | R2TX-c-1 |
| great horned owl | Raptor | winter | TX | Hale | 2 | cropland | R2TX-c-1 | 2020-12-07T00:00:00Z | 42.8777778 | 48.8826389 | 1 | R2TX-c-1 |
| ring-necked pheasant | Game Bird | winter | TX | Hale | 2 | cropland | R2TX-c-1 | 2020-12-07T00:00:00Z | 14.1826389 | 21.2006944 | 1 | R2TX-c-1 |
| great horned owl | Raptor | winter | TX | Hale | 2 | cropland | R2TX-c-1 | 2020-12-07T00:00:00Z | 21.1576389 | 28.1479167 | 1 | R2TX-c-1 |
| great horned owl | Raptor | spring | TX | Hale | 2 | cropland | R2TX-c-1 | 2021-03-15T00:00:00Z | 56.0243056 | NA | 1 | R2TX-c-1 |
| ring-necked pheasant | Game Bird | spring | TX | Hale | 2 | cropland | R2TX-c-1 | 2021-03-15T00:00:00Z | 7.0138889 | 7.0138889 | 1 | R2TX-c-1 |
| ring-necked pheasant | Game Bird | spring | TX | Hale | 2 | cropland | R2TX-c-1 | 2021-03-15T00:00:00Z | 21.03125 | 28.0208333 | 1 | R2TX-c-1 |
| red-tailed hawk | Raptor | spring | TX | Hale | 2 | cropland | R2TX-c-1 | 2021-03-15T00:00:00Z | 56 | NA | 1 | R2TX-c-1 |
| ring-necked pheasant | Game Bird | spring | TX | Hale | 2 | cropland | R2TX-c-1 | 2021-03-15T00:00:00Z | 55.9993056 | NA | 1 | R2TX-c-1 |
| Cooper's hawk | Raptor | spring | TX | Hale | 2 | cropland | R2TX-c-1 | 2021-03-15T00:00:00Z | 7.0159722 | 7.0159722 | 1 | R2TX-c-1 |
| ring-necked pheasant | Game Bird | spring | TX | Hale | 2 | cropland | R2TX-c-1 | 2021-03-15T00:00:00Z | 55.9902778 | NA | 1 | R2TX-c-1 |
| Swainson's hawk | Raptor | spring | TX | Hale | 2 | cropland | R2TX-c-1 | 2021-03-15T00:00:00Z | 55.9909722 | NA | 1 | R2TX-c-1 |
| great horned owl | Raptor | spring | TX | Hale | 2 | cropland | R2TX-c-1 | 2021-03-15T00:00:00Z | 28.01875 | 34.9791667 | 1 | R2TX-c-1 |
| ring-necked pheasant | Game Bird | spring | TX | Hale | 2 | cropland | R2TX-c-1 | 2021-03-15T00:00:00Z | 6.9972222 | 6.9972222 | 1 | R2TX-c-1 |
| Swainson's hawk | Raptor | spring | TX | Hale | 2 | cropland | R2TX-c-1 | 2021-03-15T00:00:00Z | 6.9881944 | 13.9881944 | 1 | R2TX-c-1 |
| ring-necked pheasant | Game Bird | spring | TX | Hale | 2 | cropland | R2TX-c-1 | 2021-03-15T00:00:00Z | 20.9875 | 27.9875 | 1 | R2TX-c-1 |
| Swainson's hawk | Raptor | spring | TX | Hale | 2 | cropland | R2TX-c-1 | 2021-03-15T00:00:00Z | 55.9541667 | NA | 1 | R2TX-c-1 |
| ring-necked pheasant | Game Bird | spring | TX | Hale | 2 | cropland | R2TX-c-1 | 2021-03-15T00:00:00Z | 20.9861111 | 27.9861111 | 1 | R2TX-c-1 |
| great horned owl | Raptor | spring | TX | Hale | 2 | cropland | R2TX-c-1 | 2021-03-15T00:00:00Z | 55.95 | NA | 1 | R2TX-c-1 |
| ring-necked pheasant | Game Bird | spring | TX | Hale | 2 | cropland | R2TX-c-1 | 2021-03-15T00:00:00Z | 13.9798611 | 20.9791667 | 1 | R2TX-c-1 |
| red-tailed hawk | Raptor | spring | TX | Hale | 2 | cropland | R2TX-c-1 | 2021-03-15T00:00:00Z | 6.9652778 | 13.9722222 | 1 | R2TX-c-1 |
| ring-necked pheasant | Game Bird | spring | TX | Hale | 2 | cropland | R2TX-c-1 | 2021-03-15T00:00:00Z | 6.9631944 | 13.9722222 | 1 | R2TX-c-1 |
| great horned owl | Raptor | spring | TX | Hale | 2 | cropland | R2TX-c-1 | 2021-03-15T00:00:00Z | 55.9472222 | NA | 1 | R2TX-c-1 |
| ring-necked pheasant | Game Bird | spring | TX | Hale | 2 | cropland | R2TX-c-1 | 2021-03-15T00:00:00Z | 6.8173611 | 6.8173611 | 1 | R2TX-c-1 |
| great horned owl | Raptor | summer | UT | Beaver | 6 | shrub/scrub | R6UT-ss-1 | 2020-06-15T00:00:00Z | 55.8402778 | NA | 1 | R6UT-ss-1 |
| red-tailed hawk | Raptor | summer | UT | Beaver | 6 | shrub/scrub | R6UT-ss-1 | 2020-06-15T00:00:00Z | 55.8173611 | NA | 1 | R6UT-ss-1 |
| ring-necked pheasant | Game Bird | summer | UT | Beaver | 6 | shrub/scrub | R6UT-ss-1 | 2020-06-15T00:00:00Z | 55.8027778 | NA | 1 | R6UT-ss-1 |
| ring-necked pheasant | Game Bird | summer | UT | Beaver | 6 | shrub/scrub | R6UT-ss-1 | 2020-06-15T00:00:00Z | 45.4388889 | 45.4388889 | 1 | R6UT-ss-1 |
| great horned owl | Raptor | summer | UT | Beaver | 6 | shrub/scrub | R6UT-ss-1 | 2020-06-15T00:00:00Z | 55.8111111 | NA | 1 | R6UT-ss-1 |
| ring-necked pheasant | Game Bird | summer | UT | Beaver | 6 | shrub/scrub | R6UT-ss-1 | 2020-06-15T00:00:00Z | 55.7465278 | NA | 1 | R6UT-ss-1 |
| red-tailed hawk | Raptor | summer | UT | Beaver | 6 | shrub/scrub | R6UT-ss-1 | 2020-06-15T00:00:00Z | 27.8083333 | 29.9652778 | 1 | R6UT-ss-1 |
| ring-necked pheasant | Game Bird | summer | UT | Beaver | 6 | shrub/scrub | R6UT-ss-1 | 2020-06-15T00:00:00Z | 55.7131944 | NA | 1 | R6UT-ss-1 |
| ring-necked pheasant | Game Bird | summer | UT | Beaver | 6 | shrub/scrub | R6UT-ss-1 | 2020-06-15T00:00:00Z | 4.0548611 | 16.9791667 | 1 | R6UT-ss-1 |
| osprey | Raptor | summer | UT | Beaver | 6 | shrub/scrub | R6UT-ss-1 | 2020-06-15T00:00:00Z | 55.7326389 | NA | 1 | R6UT-ss-1 |
| osprey | Raptor | summer | UT | Beaver | 6 | shrub/scrub | R6UT-ss-1 | 2020-06-15T00:00:00Z | 55.66875 | NA | 1 | R6UT-ss-1 |
| red-tailed hawk | Raptor | summer | UT | Beaver | 6 | shrub/scrub | R6UT-ss-1 | 2020-06-15T00:00:00Z | 53.9333333 | 53.9333333 | 1 | R6UT-ss-1 |
| ring-necked pheasant | Game Bird | summer | UT | Beaver | 6 | shrub/scrub | R6UT-ss-1 | 2020-06-15T00:00:00Z | 42.0625 | 42.0638889 | 1 | R6UT-ss-1 |
| red-tailed hawk | Raptor | summer | UT | Beaver | 6 | shrub/scrub | R6UT-ss-1 | 2020-06-15T00:00:00Z | 55.5791667 | NA | 1 | R6UT-ss-1 |
| prairie falcon | Raptor | summer | UT | Beaver | 6 | shrub/scrub | R6UT-ss-1 | 2020-06-15T00:00:00Z | 56.60625 | NA | 1 | R6UT-ss-1 |
| ring-necked pheasant | Game Bird | summer | UT | Beaver | 6 | shrub/scrub | R6UT-ss-1 | 2020-06-15T00:00:00Z | 55.7965278 | NA | 1 | R6UT-ss-1 |
| ring-necked pheasant | Game Bird | summer | UT | Beaver | 6 | shrub/scrub | R6UT-ss-1 | 2020-06-15T00:00:00Z | 55.9194444 | NA | 1 | R6UT-ss-1 |
| Cooper's hawk | Raptor | summer | UT | Beaver | 6 | shrub/scrub | R6UT-ss-1 | 2020-06-15T00:00:00Z | 55.5326389 | NA | 1 | R6UT-ss-1 |
| ring-necked pheasant | Game Bird | summer | UT | Beaver | 6 | shrub/scrub | R6UT-ss-1 | 2020-06-15T00:00:00Z | 55.7138889 | NA | 1 | R6UT-ss-1 |
| ring-necked pheasant | Game Bird | summer | UT | Beaver | 6 | shrub/scrub | R6UT-ss-1 | 2020-06-15T00:00:00Z | 41.5104167 | 42.775 | 1 | R6UT-ss-1 |
| ring-necked pheasant | Game Bird | fall | UT | Beaver | 6 | shrub/scrub | R6UT-ss-1 | 2020-09-14T00:00:00Z | 56.3958333 | NA | 1 | R6UT-ss-1 |
| Cooper's hawk | Raptor | fall | UT | Beaver | 6 | shrub/scrub | R6UT-ss-1 | 2020-09-14T00:00:00Z | 56.2888889 | NA | 1 | R6UT-ss-1 |
| red-tailed hawk | Raptor | fall | UT | Beaver | 6 | shrub/scrub | R6UT-ss-1 | 2020-09-14T00:00:00Z | 56.1923611 | 56.2340278 | 1 | R6UT-ss-1 |
| ring-necked pheasant | Game Bird | fall | UT | Beaver | 6 | shrub/scrub | R6UT-ss-1 | 2020-09-14T00:00:00Z | 57.2493056 | NA | 1 | R6UT-ss-1 |
| barn owl | Raptor | fall | UT | Beaver | 6 | shrub/scrub | R6UT-ss-1 | 2020-09-14T00:00:00Z | 32.3979167 | 32.3979167 | 1 | R6UT-ss-1 |
| great horned owl | Raptor | fall | UT | Beaver | 6 | shrub/scrub | R6UT-ss-1 | 2020-09-14T00:00:00Z | 16.5305556 | 16.5493056 | 1 | R6UT-ss-1 |
| ring-necked pheasant | Game Bird | fall | UT | Beaver | 6 | shrub/scrub | R6UT-ss-1 | 2020-09-14T00:00:00Z | 49.4840278 | 49.53125 | 1 | R6UT-ss-1 |
| great horned owl | Raptor | fall | UT | Beaver | 6 | shrub/scrub | R6UT-ss-1 | 2020-09-14T00:00:00Z | 57.1631944 | NA | 1 | R6UT-ss-1 |
| ring-necked pheasant | Game Bird | fall | UT | Beaver | 6 | shrub/scrub | R6UT-ss-1 | 2020-09-14T00:00:00Z | 11.75 | 14.2076389 | 1 | R6UT-ss-1 |
| great horned owl | Raptor | fall | UT | Beaver | 6 | shrub/scrub | R6UT-ss-1 | 2020-09-14T00:00:00Z | 22.4555556 | 22.4555556 | 1 | R6UT-ss-1 |
| great horned owl | Raptor | fall | UT | Beaver | 6 | shrub/scrub | R6UT-ss-1 | 2020-09-14T00:00:00Z | 57.0930556 | NA | 1 | R6UT-ss-1 |
| ring-necked pheasant | Game Bird | fall | UT | Beaver | 6 | shrub/scrub | R6UT-ss-1 | 2020-09-14T00:00:00Z | 9.6805556 | 9.7 | 1 | R6UT-ss-1 |
| Swainson's hawk | Raptor | fall | UT | Beaver | 6 | shrub/scrub | R6UT-ss-1 | 2020-09-14T00:00:00Z | 2.81875 | 14.1006944 | 1 | R6UT-ss-1 |
| ring-necked pheasant | Game Bird | fall | UT | Beaver | 6 | shrub/scrub | R6UT-ss-1 | 2020-09-14T00:00:00Z | 56.9868056 | NA | 1 | R6UT-ss-1 |
| great horned owl | Raptor | fall | UT | Beaver | 6 | shrub/scrub | R6UT-ss-1 | 2020-09-14T00:00:00Z | 14.7041667 | 27.9263889 | 1 | R6UT-ss-1 |
| ring-necked pheasant | Game Bird | fall | UT | Beaver | 6 | shrub/scrub | R6UT-ss-1 | 2020-09-14T00:00:00Z | 56.9229167 | NA | 1 | R6UT-ss-1 |
| ring-necked pheasant | Game Bird | fall | UT | Beaver | 6 | shrub/scrub | R6UT-ss-1 | 2020-09-14T00:00:00Z | 3.8013889 | 3.8013889 | 1 | R6UT-ss-1 |
| ring-necked pheasant | Game Bird | fall | UT | Beaver | 6 | shrub/scrub | R6UT-ss-1 | 2020-09-14T00:00:00Z | 4.0423611 | 6.7979167 | 1 | R6UT-ss-1 |
| ring-necked pheasant | Game Bird | fall | UT | Beaver | 6 | shrub/scrub | R6UT-ss-1 | 2020-09-14T00:00:00Z | 11.3319444 | 11.3319444 | 1 | R6UT-ss-1 |
| ring-necked pheasant | Game Bird | fall | UT | Beaver | 6 | shrub/scrub | R6UT-ss-1 | 2020-09-14T00:00:00Z | 56.925 | NA | 1 | R6UT-ss-1 |
| ring-necked pheasant | Game Bird | winter | UT | Beaver | 6 | shrub/scrub | R6UT-ss-1 | 2020-12-07T00:00:00Z | 28.1347222 | 28.1347222 | 1 | R6UT-ss-1 |
| Cooper's hawk | Raptor | winter | UT | Beaver | 6 | shrub/scrub | R6UT-ss-1 | 2020-12-07T00:00:00Z | 28.1666667 | 28.1666667 | 1 | R6UT-ss-1 |
| Swainson's hawk | Raptor | winter | UT | Beaver | 6 | shrub/scrub | R6UT-ss-1 | 2020-12-07T00:00:00Z | 7.2583333 | 7.9597222 | 1 | R6UT-ss-1 |
| ring-necked pheasant | Game Bird | winter | UT | Beaver | 6 | shrub/scrub | R6UT-ss-1 | 2020-12-07T00:00:00Z | 22.23125 | 22.2729167 | 1 | R6UT-ss-1 |
| great horned owl | Raptor | winter | UT | Beaver | 6 | shrub/scrub | R6UT-ss-1 | 2020-12-07T00:00:00Z | 56.3270833 | NA | 1 | R6UT-ss-1 |
| Swainson's hawk | Raptor | winter | UT | Beaver | 6 | shrub/scrub | R6UT-ss-1 | 2020-12-07T00:00:00Z | 9.3381944 | 9.3798611 | 1 | R6UT-ss-1 |
| great horned owl | Raptor | winter | UT | Beaver | 6 | shrub/scrub | R6UT-ss-1 | 2020-12-07T00:00:00Z | 56.4395833 | NA | 1 | R6UT-ss-1 |
| ring-necked pheasant | Game Bird | winter | UT | Beaver | 6 | shrub/scrub | R6UT-ss-1 | 2020-12-07T00:00:00Z | 56.3180556 | NA | 1 | R6UT-ss-1 |
| ring-necked pheasant | Game Bird | winter | UT | Beaver | 6 | shrub/scrub | R6UT-ss-1 | 2020-12-07T00:00:00Z | 20.9263889 | 28.275 | 1 | R6UT-ss-1 |
| Swainson's hawk | Raptor | winter | UT | Beaver | 6 | shrub/scrub | R6UT-ss-1 | 2020-12-07T00:00:00Z | 62.9694444 | NA | 1 | R6UT-ss-1 |
| great horned owl | Raptor | winter | UT | Beaver | 6 | shrub/scrub | R6UT-ss-1 | 2020-12-07T00:00:00Z | 3.0423611 | 28.9236111 | 1 | R6UT-ss-1 |
| ring-necked pheasant | Game Bird | winter | UT | Beaver | 6 | shrub/scrub | R6UT-ss-1 | 2020-12-07T00:00:00Z | 56.21875 | NA | 1 | R6UT-ss-1 |
| Swainson's hawk | Raptor | winter | UT | Beaver | 6 | shrub/scrub | R6UT-ss-1 | 2020-12-07T00:00:00Z | 56.1923611 | NA | 1 | R6UT-ss-1 |
| ring-necked pheasant | Game Bird | winter | UT | Beaver | 6 | shrub/scrub | R6UT-ss-1 | 2020-12-07T00:00:00Z | 56.1840278 | NA | 1 | R6UT-ss-1 |
| ring-necked pheasant | Game Bird | winter | UT | Beaver | 6 | shrub/scrub | R6UT-ss-1 | 2020-12-07T00:00:00Z | 56.125 | NA | 1 | R6UT-ss-1 |
| red-tailed hawk | Raptor | winter | UT | Beaver | 6 | shrub/scrub | R6UT-ss-1 | 2020-12-07T00:00:00Z | 56.1555556 | NA | 1 | R6UT-ss-1 |
| ring-necked pheasant | Game Bird | winter | UT | Beaver | 6 | shrub/scrub | R6UT-ss-1 | 2020-12-07T00:00:00Z | 20.2465278 | 20.3104167 | 1 | R6UT-ss-1 |
| ring-necked pheasant | Game Bird | winter | UT | Beaver | 6 | shrub/scrub | R6UT-ss-1 | 2020-12-07T00:00:00Z | 56.1152778 | NA | 1 | R6UT-ss-1 |
| great horned owl | Raptor | winter | UT | Beaver | 6 | shrub/scrub | R6UT-ss-1 | 2020-12-07T00:00:00Z | 56.1 | NA | 1 | R6UT-ss-1 |
| ring-necked pheasant | Game Bird | winter | UT | Beaver | 6 | shrub/scrub | R6UT-ss-1 | 2020-12-07T00:00:00Z | 7.8875 | 8.1298611 | 1 | R6UT-ss-1 |
| great horned owl | Raptor | spring | UT | Beaver | 6 | shrub/scrub | R6UT-ss-1 | 2021-03-15T00:00:00Z | 59.09375 | NA | 1 | R6UT-ss-1 |
| ring-necked pheasant | Game Bird | spring | UT | Beaver | 6 | shrub/scrub | R6UT-ss-1 | 2021-03-15T00:00:00Z | 59.0590278 | NA | 1 | R6UT-ss-1 |
| Swainson's hawk | Raptor | spring | UT | Beaver | 6 | shrub/scrub | R6UT-ss-1 | 2021-03-15T00:00:00Z | 56.3173611 | NA | 1 | R6UT-ss-1 |
| ring-necked pheasant | Game Bird | spring | UT | Beaver | 6 | shrub/scrub | R6UT-ss-1 | 2021-03-15T00:00:00Z | 15.1972222 | 15.1972222 | 1 | R6UT-ss-1 |
| Swainson's hawk | Raptor | spring | UT | Beaver | 6 | shrub/scrub | R6UT-ss-1 | 2021-03-15T00:00:00Z | 55.9965278 | NA | 1 | R6UT-ss-1 |
| ring-necked pheasant | Game Bird | spring | UT | Beaver | 6 | shrub/scrub | R6UT-ss-1 | 2021-03-15T00:00:00Z | 15.2347222 | 15.2347222 | 1 | R6UT-ss-1 |
| great horned owl | Raptor | spring | UT | Beaver | 6 | shrub/scrub | R6UT-ss-1 | 2021-03-15T00:00:00Z | 58.9486111 | NA | 1 | R6UT-ss-1 |
| ring-necked pheasant | Game Bird | spring | UT | Beaver | 6 | shrub/scrub | R6UT-ss-1 | 2021-03-15T00:00:00Z | 21.2277778 | 28.0951389 | 1 | R6UT-ss-1 |
| red-tailed hawk | Raptor | spring | UT | Beaver | 6 | shrub/scrub | R6UT-ss-1 | 2021-03-15T00:00:00Z | 58.9 | NA | 1 | R6UT-ss-1 |
| ring-necked pheasant | Game Bird | spring | UT | Beaver | 6 | shrub/scrub | R6UT-ss-1 | 2021-03-15T00:00:00Z | 15.0645833 | 21.0819444 | 1 | R6UT-ss-1 |
| barn owl | Raptor | spring | UT | Beaver | 6 | shrub/scrub | R6UT-ss-1 | 2021-03-15T00:00:00Z | 14.9493056 | 14.9493056 | 1 | R6UT-ss-1 |
| ring-necked pheasant | Game Bird | spring | UT | Beaver | 6 | shrub/scrub | R6UT-ss-1 | 2021-03-15T00:00:00Z | 29.9840278 | 42.9173611 | 1 | R6UT-ss-1 |
| great horned owl | Raptor | spring | UT | Beaver | 6 | shrub/scrub | R6UT-ss-1 | 2021-03-15T00:00:00Z | 15.8118056 | 15.8118056 | 1 | R6UT-ss-1 |
| ring-necked pheasant | Game Bird | spring | UT | Beaver | 6 | shrub/scrub | R6UT-ss-1 | 2021-03-15T00:00:00Z | 16.0527778 | 16.0527778 | 1 | R6UT-ss-1 |
| ring-necked pheasant | Game Bird | spring | UT | Beaver | 6 | shrub/scrub | R6UT-ss-1 | 2021-03-15T00:00:00Z | 28.1236111 | 42.8069444 | 1 | R6UT-ss-1 |
| red-tailed hawk | Raptor | spring | UT | Beaver | 6 | shrub/scrub | R6UT-ss-1 | 2021-03-15T00:00:00Z | 58.69375 | NA | 1 | R6UT-ss-1 |
| ring-necked pheasant | Game Bird | spring | UT | Beaver | 6 | shrub/scrub | R6UT-ss-1 | 2021-03-15T00:00:00Z | 55.9291667 | NA | 1 | R6UT-ss-1 |
| Cooper's hawk | Raptor | spring | UT | Beaver | 6 | shrub/scrub | R6UT-ss-1 | 2021-03-15T00:00:00Z | 27.9659722 | 42.7548611 | 1 | R6UT-ss-1 |
| great horned owl | Raptor | spring | UT | Beaver | 6 | shrub/scrub | R6UT-ss-1 | 2021-03-15T00:00:00Z | 55.575 | NA | 1 | R6UT-ss-1 |
| ring-necked pheasant | Game Bird | spring | UT | Beaver | 6 | shrub/scrub | R6UT-ss-1 | 2021-03-15T00:00:00Z | 7.0277778 | 15.6305556 | 1 | R6UT-ss-1 |
| barn owl | Raptor | fall | NC | Perquimans/Pasquotank | 4 | cropland | R4NC-c-1 | 2020-09-14T00:00:00Z | 55.7951389 | NA | 1 | R4NC-c-1 |
| barn owl | Raptor | fall | NC | Perquimans/Pasquotank | 4 | cropland | R4NC-c-1 | 2020-09-14T00:00:00Z | 7.0381944 | 13.9861111 | 1 | R4NC-c-1 |
| ring-necked pheasant | Game Bird | fall | NC | Perquimans/Pasquotank | 4 | cropland | R4NC-c-1 | 2020-09-14T00:00:00Z | 6.8708333 | 6.8708333 | 1 | R4NC-c-1 |
| ring-necked pheasant | Game Bird | fall | NC | Perquimans/Pasquotank | 4 | cropland | R4NC-c-1 | 2020-09-14T00:00:00Z | 55.9034722 | NA | 1 | R4NC-c-1 |
| ring-necked pheasant | Game Bird | fall | NC | Perquimans/Pasquotank | 4 | cropland | R4NC-c-1 | 2020-09-14T00:00:00Z | 6.8944444 | 6.8944444 | 1 | R4NC-c-1 |
| ring-necked pheasant | Game Bird | fall | NC | Perquimans/Pasquotank | 4 | cropland | R4NC-c-1 | 2020-09-14T00:00:00Z | 6.8180556 | 6.8180556 | 1 | R4NC-c-1 |
| ring-necked pheasant | Game Bird | fall | NC | Perquimans/Pasquotank | 4 | cropland | R4NC-c-1 | 2020-09-14T00:00:00Z | 6.7923611 | 6.7923611 | 1 | R4NC-c-1 |
| ring-necked pheasant | Game Bird | fall | NC | Perquimans/Pasquotank | 4 | cropland | R4NC-c-1 | 2020-09-14T00:00:00Z | 7.0881944 | 14.0326389 | 1 | R4NC-c-1 |
| great horned owl | Raptor | fall | NC | Perquimans/Pasquotank | 4 | cropland | R4NC-c-1 | 2020-09-14T00:00:00Z | 55.9763889 | NA | 1 | R4NC-c-1 |
| great horned owl | Raptor | fall | NC | Perquimans/Pasquotank | 4 | cropland | R4NC-c-1 | 2020-09-14T00:00:00Z | 55.9201389 | NA | 1 | R4NC-c-1 |
| ring-necked pheasant | Game Bird | fall | NC | Perquimans/Pasquotank | 4 | cropland | R4NC-c-1 | 2020-09-14T00:00:00Z | 55.9868056 | NA | 1 | R4NC-c-1 |
| ring-necked pheasant | Game Bird | fall | NC | Perquimans/Pasquotank | 4 | cropland | R4NC-c-1 | 2020-09-14T00:00:00Z | 55.7430556 | NA | 1 | R4NC-c-1 |
| great horned owl | Raptor | fall | NC | Perquimans/Pasquotank | 4 | cropland | R4NC-c-1 | 2020-09-14T00:00:00Z | 55.9701389 | NA | 1 | R4NC-c-1 |
| great horned owl | Raptor | fall | NC | Perquimans/Pasquotank | 4 | cropland | R4NC-c-1 | 2020-09-14T00:00:00Z | 55.7833333 | NA | 1 | R4NC-c-1 |
| ring-necked pheasant | Game Bird | fall | NC | Perquimans/Pasquotank | 4 | cropland | R4NC-c-1 | 2020-09-14T00:00:00Z | 13.7291667 | 20.7395833 | 1 | R4NC-c-1 |
| ring-necked pheasant | Game Bird | fall | NC | Perquimans/Pasquotank | 4 | cropland | R4NC-c-1 | 2020-09-14T00:00:00Z | 55.9409722 | NA | 1 | R4NC-c-1 |
| Swainson's hawk | Raptor | fall | NC | Perquimans/Pasquotank | 4 | cropland | R4NC-c-1 | 2020-09-14T00:00:00Z | 6.7465278 | 6.7465278 | 1 | R4NC-c-1 |
| Swainson's hawk | Raptor | fall | NC | Perquimans/Pasquotank | 4 | cropland | R4NC-c-1 | 2020-09-14T00:00:00Z | 55.8180556 | NA | 1 | R4NC-c-1 |
| Swainson's hawk | Raptor | fall | NC | Perquimans/Pasquotank | 4 | cropland | R4NC-c-1 | 2020-09-14T00:00:00Z | 55.8916667 | NA | 1 | R4NC-c-1 |
| red-tailed hawk | Raptor | fall | NC | Perquimans/Pasquotank | 4 | cropland | R4NC-c-1 | 2020-09-14T00:00:00Z | 48.8111111 | 55.8118056 | 1 | R4NC-c-1 |
| ring-necked pheasant | Game Bird | winter | NC | Perquimans/Pasquotank | 4 | cropland | R4NC-c-1 | 2020-12-14T00:00:00Z | 6.9701389 | 6.9701389 | 1 | R4NC-c-1 |
| ring-necked pheasant | Game Bird | winter | NC | Perquimans/Pasquotank | 4 | cropland | R4NC-c-1 | 2020-12-14T00:00:00Z | 27.9840278 | 34.9638889 | 1 | R4NC-c-1 |
| ring-necked pheasant | Game Bird | winter | NC | Perquimans/Pasquotank | 4 | cropland | R4NC-c-1 | 2020-12-14T00:00:00Z | 13.96875 | 20.9361111 | 1 | R4NC-c-1 |
| ring-necked pheasant | Game Bird | winter | NC | Perquimans/Pasquotank | 4 | cropland | R4NC-c-1 | 2020-12-14T00:00:00Z | 27.9284722 | 34.9375 | 1 | R4NC-c-1 |
| ring-necked pheasant | Game Bird | winter | NC | Perquimans/Pasquotank | 4 | cropland | R4NC-c-1 | 2020-12-14T00:00:00Z | 6.9111111 | 6.9111111 | 1 | R4NC-c-1 |
| ring-necked pheasant | Game Bird | winter | NC | Perquimans/Pasquotank | 4 | cropland | R4NC-c-1 | 2020-12-14T00:00:00Z | 55.8451389 | NA | 1 | R4NC-c-1 |
| ring-necked pheasant | Game Bird | winter | NC | Perquimans/Pasquotank | 4 | cropland | R4NC-c-1 | 2020-12-14T00:00:00Z | 55.8465278 | NA | 1 | R4NC-c-1 |
| ring-necked pheasant | Game Bird | winter | NC | Perquimans/Pasquotank | 4 | cropland | R4NC-c-1 | 2020-12-14T00:00:00Z | 20.9048611 | 27.9125 | 1 | R4NC-c-1 |
| ring-necked pheasant | Game Bird | winter | NC | Perquimans/Pasquotank | 4 | cropland | R4NC-c-1 | 2020-12-14T00:00:00Z | 55.8604167 | NA | 1 | R4NC-c-1 |
| ring-necked pheasant | Game Bird | winter | NC | Perquimans/Pasquotank | 4 | cropland | R4NC-c-1 | 2020-12-14T00:00:00Z | 6.9131944 | 6.9131944 | 1 | R4NC-c-1 |
| Cooper's hawk | Raptor | winter | NC | Perquimans/Pasquotank | 4 | cropland | R4NC-c-1 | 2020-12-14T00:00:00Z | 13.9472222 | 20.9215278 | 1 | R4NC-c-1 |
| osprey | Raptor | winter | NC | Perquimans/Pasquotank | 4 | cropland | R4NC-c-1 | 2020-12-14T00:00:00Z | 55.9145833 | NA | 1 | R4NC-c-1 |
| barn owl | Raptor | winter | NC | Perquimans/Pasquotank | 4 | cropland | R4NC-c-1 | 2020-12-14T00:00:00Z | 6.8833333 | 6.8833333 | 1 | R4NC-c-1 |
| great horned owl | Raptor | winter | NC | Perquimans/Pasquotank | 4 | cropland | R4NC-c-1 | 2020-12-14T00:00:00Z | 55.8979167 | NA | 1 | R4NC-c-1 |
| red-tailed hawk | Raptor | winter | NC | Perquimans/Pasquotank | 4 | cropland | R4NC-c-1 | 2020-12-14T00:00:00Z | 55.8479167 | NA | 1 | R4NC-c-1 |
| Swainson's hawk | Raptor | winter | NC | Perquimans/Pasquotank | 4 | cropland | R4NC-c-1 | 2020-12-14T00:00:00Z | 6.9027778 | 6.9027778 | 1 | R4NC-c-1 |
| red-tailed hawk | Raptor | winter | NC | Perquimans/Pasquotank | 4 | cropland | R4NC-c-1 | 2020-12-14T00:00:00Z | 6.9055556 | 13.9340278 | 1 | R4NC-c-1 |
| great horned owl | Raptor | winter | NC | Perquimans/Pasquotank | 4 | cropland | R4NC-c-1 | 2020-12-14T00:00:00Z | 13.9333333 | 20.9090278 | 1 | R4NC-c-1 |
| barn owl | Raptor | winter | NC | Perquimans/Pasquotank | 4 | cropland | R4NC-c-1 | 2020-12-14T00:00:00Z | 6.9118056 | 6.9118056 | 1 | R4NC-c-1 |
| Swainson's hawk | Raptor | winter | NC | Perquimans/Pasquotank | 4 | cropland | R4NC-c-1 | 2020-12-14T00:00:00Z | 6.9145833 | 6.9145833 | 1 | R4NC-c-1 |
| ring-necked pheasant | Game Bird | spring | NC | Perquimans/Pasquotank | 4 | cropland | R4NC-c-1 | 2021-03-15T00:00:00Z | 27.9479167 | 34.9638889 | 1 | R4NC-c-1 |
| Swainson's hawk | Raptor | spring | NC | Perquimans/Pasquotank | 4 | cropland | R4NC-c-1 | 2021-03-15T00:00:00Z | 34.9604167 | 41.9201389 | 1 | R4NC-c-1 |
| ring-necked pheasant | Game Bird | spring | NC | Perquimans/Pasquotank | 4 | cropland | R4NC-c-1 | 2021-03-15T00:00:00Z | 6.95 | 13.9611111 | 1 | R4NC-c-1 |
| Swainson's hawk | Raptor | spring | NC | Perquimans/Pasquotank | 4 | cropland | R4NC-c-1 | 2021-03-15T00:00:00Z | 48.9895833 | 55.9006944 | 1 | R4NC-c-1 |
| ring-necked pheasant | Game Bird | spring | NC | Perquimans/Pasquotank | 4 | cropland | R4NC-c-1 | 2021-03-15T00:00:00Z | 13.9680556 | 20.9652778 | 1 | R4NC-c-1 |
| Swainson's hawk | Raptor | spring | NC | Perquimans/Pasquotank | 4 | cropland | R4NC-c-1 | 2021-03-15T00:00:00Z | 55.8930556 | NA | 1 | R4NC-c-1 |
| barn owl | Raptor | spring | NC | Perquimans/Pasquotank | 4 | cropland | R4NC-c-1 | 2021-03-15T00:00:00Z | 6.95625 | 6.95625 | 1 | R4NC-c-1 |
| ring-necked pheasant | Game Bird | spring | NC | Perquimans/Pasquotank | 4 | cropland | R4NC-c-1 | 2021-03-15T00:00:00Z | 6.9569444 | 6.9569444 | 1 | R4NC-c-1 |
| Swainson's hawk | Raptor | spring | NC | Perquimans/Pasquotank | 4 | cropland | R4NC-c-1 | 2021-03-15T00:00:00Z | 55.8791667 | NA | 1 | R4NC-c-1 |
| ring-necked pheasant | Game Bird | spring | NC | Perquimans/Pasquotank | 4 | cropland | R4NC-c-1 | 2021-03-15T00:00:00Z | 6.9666667 | 13.9506944 | 1 | R4NC-c-1 |
| Cooper's hawk | Raptor | spring | NC | Perquimans/Pasquotank | 4 | cropland | R4NC-c-1 | 2021-03-15T00:00:00Z | 6.96875 | 6.96875 | 1 | R4NC-c-1 |
| ring-necked pheasant | Game Bird | spring | NC | Perquimans/Pasquotank | 4 | cropland | R4NC-c-1 | 2021-03-15T00:00:00Z | 13.9458333 | 20.9375 | 1 | R4NC-c-1 |
| great horned owl | Raptor | spring | NC | Perquimans/Pasquotank | 4 | cropland | R4NC-c-1 | 2021-03-15T00:00:00Z | 55.8701389 | NA | 1 | R4NC-c-1 |
| ring-necked pheasant | Game Bird | spring | NC | Perquimans/Pasquotank | 4 | cropland | R4NC-c-1 | 2021-03-15T00:00:00Z | 55.8569444 | NA | 1 | R4NC-c-1 |
| Swainson's hawk | Raptor | spring | NC | Perquimans/Pasquotank | 4 | cropland | R4NC-c-1 | 2021-03-15T00:00:00Z | 6.9645833 | 13.9486111 | 1 | R4NC-c-1 |
| ring-necked pheasant | Game Bird | spring | NC | Perquimans/Pasquotank | 4 | cropland | R4NC-c-1 | 2021-03-15T00:00:00Z | 6.9791667 | 13.9479167 | 1 | R4NC-c-1 |
| Swainson's hawk | Raptor | spring | NC | Perquimans/Pasquotank | 4 | cropland | R4NC-c-1 | 2021-03-15T00:00:00Z | 6.9548611 | 13.9451389 | 1 | R4NC-c-1 |
| ring-necked pheasant | Game Bird | spring | NC | Perquimans/Pasquotank | 4 | cropland | R4NC-c-1 | 2021-03-15T00:00:00Z | 6.9541667 | 13.9423611 | 1 | R4NC-c-1 |
| ring-necked pheasant | Game Bird | spring | NC | Perquimans/Pasquotank | 4 | cropland | R4NC-c-1 | 2021-03-15T00:00:00Z | 6.9645833 | 6.9645833 | 1 | R4NC-c-1 |
| Swainson's hawk | Raptor | spring | NC | Perquimans/Pasquotank | 4 | cropland | R4NC-c-1 | 2021-03-15T00:00:00Z | 55.8340278 | NA | 1 | R4NC-c-1 |
| osprey | Raptor | summer | NC | Perquimans/Pasquotank | 4 | cropland | R4NC-c-1 | 2021-06-30T00:00:00Z | 56.5625 | NA | 1 | R4NC-c-1 |
| ring-necked pheasant | Game Bird | summer | NC | Perquimans/Pasquotank | 4 | cropland | R4NC-c-1 | 2021-06-30T00:00:00Z | 56.7611111 | NA | 1 | R4NC-c-1 |
| ring-necked pheasant | Game Bird | summer | NC | Perquimans/Pasquotank | 4 | cropland | R4NC-c-1 | 2021-07-01T00:00:00Z | 55.9916667 | NA | 1 | R4NC-c-1 |
| red-tailed hawk | Raptor | summer | NC | Perquimans/Pasquotank | 4 | cropland | R4NC-c-1 | 2021-07-01T00:00:00Z | 55.8868056 | NA | 1 | R4NC-c-1 |
| red-tailed hawk | Raptor | summer | NC | Perquimans/Pasquotank | 4 | cropland | R4NC-c-1 | 2021-07-01T00:00:00Z | 55.9173611 | NA | 1 | R4NC-c-1 |
| barn owl | Raptor | summer | NC | Perquimans/Pasquotank | 4 | cropland | R4NC-c-1 | 2021-06-30T00:00:00Z | 48.6861111 | NA | 1 | R4NC-c-1 |
| ring-necked pheasant | Game Bird | summer | NC | Perquimans/Pasquotank | 4 | cropland | R4NC-c-1 | 2021-07-01T00:00:00Z | 2.35625 | 2.35625 | 1 | R4NC-c-1 |
| ring-necked pheasant | Game Bird | summer | NC | Perquimans/Pasquotank | 4 | cropland | R4NC-c-1 | 2021-06-30T00:00:00Z | 48.6111111 | 56.5965278 | 1 | R4NC-c-1 |
| red-tailed hawk | Raptor | summer | NC | Perquimans/Pasquotank | 4 | cropland | R4NC-c-1 | 2021-07-01T00:00:00Z | 55.9604167 | NA | 1 | R4NC-c-1 |
| Swainson's hawk | Raptor | summer | NC | Perquimans/Pasquotank | 4 | cropland | R4NC-c-1 | 2021-07-01T00:00:00Z | 47.7625 | 55.7326389 | 1 | R4NC-c-1 |
| ring-necked pheasant | Game Bird | summer | NC | Perquimans/Pasquotank | 4 | cropland | R4NC-c-1 | 2021-07-01T00:00:00Z | 55.96875 | NA | 1 | R4NC-c-1 |
| Swainson's hawk | Raptor | summer | NC | Perquimans/Pasquotank | 4 | cropland | R4NC-c-1 | 2021-07-01T00:00:00Z | 1.8513889 | 2.8513889 | 1 | R4NC-c-1 |
| ring-necked pheasant | Game Bird | summer | NC | Perquimans/Pasquotank | 4 | cropland | R4NC-c-1 | 2021-06-30T00:00:00Z | 2.8590278 | 3.8590278 | 1 | R4NC-c-1 |
| barn owl | Raptor | summer | NC | Perquimans/Pasquotank | 4 | cropland | R4NC-c-1 | 2021-06-30T00:00:00Z | 1.8583333 | 2.8583333 | 1 | R4NC-c-1 |
| Swainson's hawk | Raptor | summer | NC | Perquimans/Pasquotank | 4 | cropland | R4NC-c-1 | 2021-07-01T00:00:00Z | 19.1236111 | 19.1236111 | 1 | R4NC-c-1 |
| great horned owl | Raptor | summer | NC | Perquimans/Pasquotank | 4 | cropland | R4NC-c-1 | 2021-06-30T00:00:00Z | 19.7354167 | 19.7354167 | 1 | R4NC-c-1 |
| ring-necked pheasant | Game Bird | summer | NC | Perquimans/Pasquotank | 4 | cropland | R4NC-c-1 | 2021-07-01T00:00:00Z | 19.0583333 | 19.0583333 | 1 | R4NC-c-1 |
| ring-necked pheasant | Game Bird | summer | NC | Perquimans/Pasquotank | 4 | cropland | R4NC-c-1 | 2021-07-01T00:00:00Z | 19.1013889 | 19.1013889 | 1 | R4NC-c-1 |
| ring-necked pheasant | Game Bird | summer | NC | Perquimans/Pasquotank | 4 | cropland | R4NC-c-1 | 2021-06-30T00:00:00Z | 19.6777778 | 19.6777778 | 1 | R4NC-c-1 |
| ring-necked pheasant | Game Bird | summer | NC | Perquimans/Pasquotank | 4 | cropland | R4NC-c-1 | 2021-07-01T00:00:00Z | 19.08125 | 19.08125 | 1 | R4NC-c-1 |
