## Supplementary material for "Game Bird Carcasses are Less Persistent Than Raptor Carcasses, But Can Predict Raptor Persistence Dynamics": S2 Appendix

**S2 Table. Field trial breakdown.** Summary of the carcasses (200 game birds and 199 raptors) used in field trials at the six Study Sites.

| **Species** | **Arkwright Summit Wind Farm** | **Desert Wind Farm** | **El Cabo Wind Farm** | **Grande Prairie Wind Farm** | **Hale Wind Farm** | **Milford Wind Corridor** |
| --- | --- | --- | --- | --- | --- | --- |
| **Game Birds** | **40** | **40** | **40** | **39** | **40** | **40** |
| mallard | 20 | 0 | 0 | 0 | 0 | 0 |
| ring-necked pheasant | 20 | 40 | 40 | 39 | 40 | 40 |
| **Raptors** | **40** | **40** | **40** | **40** | **40** | **39** |
| barn owl | 2 | 7 | 1 | 3 | 2 | 2 |
| barred owl | 0 | 0 | 1 | 0 | 0 | 0 |
| Cooper's hawk | 4 | 2 | 5 | 3 | 5 | 4 |
| ferruginous hawk | 0 | 0 | 1 | 0 | 0 | 0 |
| great horned owl | 14 | 8 | 15 | 14 | 14 | 15 |
| osprey | 3 | 2 | 1 | 1 | 1 | 2 |
| peregrine falcon | 0 | 0 | 1 | 0 | 0 | 0 |
| prairie falcon | 1 | 0 | 5 | 1 | 0 | 1 |
| red-tailed hawk | 10 | 6 | 5 | 14 | 11 | 8 |
| rough-legged hawk | 0 | 0 | 0 | 0 | 1 | 0 |
| Swainson's hawk | 6 | 15 | 5 | 4 | 6 | 7 |
