## Supplementary material for "Game Bird Carcasses are Less Persistent Than Raptor Carcasses, But Can Predict Raptor Persistence Dynamics": S3 Appendix

**S3 Table. Model selection results using interval‑censored survival regression to scale game bird persistence.**

| **Distribution** | **Location Parameter** | **Scale Parameter** | **Number of Parameters** | **Sample Size** | **AICc** | **Delta AICc** | **Model Used** | **Class** | **Analysis Group** |
| --- | --- | --- | --- | --- | --- | --- | --- | --- | --- |
| exponential | l ~ 1 | NULL | 1 | 80 | 350.85 | 0 | * | Game Bird | R1OR/WA-c-2 |
| Weibull | l ~ 1 | s ~ 1 | 2 | 80 | 351.17 | 0.32 |  | Game Bird | R1OR/WA-c-2 |
| loglogistic | l ~ 1 | s ~ 1 | 2 | 80 | 351.78 | 0.93 |  | Game Bird | R1OR/WA-c-2 |
| lognormal | l ~ 1 | s ~ 1 | 2 | 80 | 353.99 | 3.14 |  | Game Bird | R1OR/WA-c-2 |
| exponential | l ~ 1 | NULL | 1 | 14 | 44.85 | 0 | * | Raptor | R1OR/WA-c-2 |
| lognormal | l ~ 1 | s ~ 1 | 2 | 14 | 46.64 | 1.79 |  | Raptor | R1OR/WA-c-2 |
| loglogistic | l ~ 1 | s ~ 1 | 2 | 14 | 47.13 | 2.28 |  | Raptor | R1OR/WA-c-2 |
| Weibull | l ~ 1 | s ~ 1 | 2 | 14 | 47.44 | 2.59 |  | Raptor | R1OR/WA-c-2 |
| Weibull | l ~ 1 | s ~ 1 | 2 | 49 | 223.78 | 0 | * | Game Bird | R1WA-c-1 |
| loglogistic | l ~ 1 | s ~ 1 | 2 | 49 | 225.06 | 1.28 |  | Game Bird | R1WA-c-1 |
| lognormal | l ~ 1 | s ~ 1 | 2 | 49 | 225.98 | 2.2 |  | Game Bird | R1WA-c-1 |
| exponential | l ~ 1 | NULL | 1 | 49 | 228.96 | 5.18 |  | Game Bird | R1WA-c-1 |
| lognormal | l ~ 1 | s ~ 1 | 2 | 25 | 98.7 | 0 | * | Game Bird | R1WA-c-2 |
| loglogistic | l ~ 1 | s ~ 1 | 2 | 25 | 100.01 | 1.31 |  | Game Bird | R1WA-c-2 |
| exponential | l ~ 1 | NULL | 1 | 25 | 101.61 | 2.91 |  | Game Bird | R1WA-c-2 |
| Weibull | l ~ 1 | s ~ 1 | 2 | 25 | 101.73 | 3.03 |  | Game Bird | R1WA-c-2 |
| lognormal | l ~ Season | s ~ Season | 6 | 74 | 280.78 | 0 | * | Raptor | R1WA-c-3 |
| loglogistic | l ~ Season | s ~ Season | 6 | 74 | 281.8 | 1.02 |  | Raptor | R1WA-c-3 |
| Weibull | l ~ Season | s ~ Season | 6 | 74 | 282.77 | 1.99 |  | Raptor | R1WA-c-3 |
| Weibull | l ~ Season | s ~ 1 | 4 | 74 | 284.36 | 3.58 |  | Raptor | R1WA-c-3 |
| loglogistic | l ~ Season | s ~ 1 | 4 | 74 | 285.72 | 4.94 |  | Raptor | R1WA-c-3 |
| lognormal | l ~ Season | s ~ 1 | 4 | 74 | 287.2 | 6.42 |  | Raptor | R1WA-c-3 |
| exponential | l ~ Season | NULL | 3 | 74 | 294.86 | 14.08 |  | Raptor | R1WA-c-3 |
| Weibull | l ~ 1 | s ~ Season | 4 | 74 | 302.67 | 21.89 |  | Raptor | R1WA-c-3 |
| lognormal | l ~ 1 | s ~ 1 | 2 | 74 | 302.95 | 22.17 |  | Raptor | R1WA-c-3 |
| lognormal | l ~ 1 | s ~ Season | 4 | 74 | 303.56 | 22.78 |  | Raptor | R1WA-c-3 |
| loglogistic | l ~ 1 | s ~ 1 | 2 | 74 | 303.97 | 23.19 |  | Raptor | R1WA-c-3 |
| loglogistic | l ~ 1 | s ~ Season | 4 | 74 | 304.42 | 23.64 |  | Raptor | R1WA-c-3 |
| Weibull | l ~ 1 | s ~ 1 | 2 | 74 | 305.7 | 24.92 |  | Raptor | R1WA-c-3 |
| exponential | l ~ 1 | NULL | 1 | 74 | 327.79 | 47.01 |  | Raptor | R1WA-c-3 |
| lognormal | l ~ 1 | s ~ 1 | 2 | 17 | 93.46 | 0 | * | Game Bird | R1WA-c-4 |
| loglogistic | l ~ 1 | s ~ 1 | 2 | 17 | 93.57 | 0.11 |  | Game Bird | R1WA-c-4 |
| Weibull | l ~ 1 | s ~ 1 | 2 | 17 | 96.78 | 3.32 |  | Game Bird | R1WA-c-4 |
| exponential | l ~ 1 | NULL | 1 | 17 | 99.68 | 6.22 |  | Game Bird | R1WA-c-4 |
| lognormal | l ~ 1 | s ~ 1 | 2 | 20 | 98.19 | 0 |  | Raptor | R1WA-c-4 |
| loglogistic | l ~ 1 | s ~ 1 | 2 | 20 | 98.65 | 0.46 |  | Raptor | R1WA-c-4 |
| exponential | l ~ 1 | NULL | 1 | 20 | 98.74 | 0.55 | * | Raptor | R1WA-c-4 |
| Weibull | l ~ 1 | s ~ 1 | 2 | 20 | 100.17 | 1.98 |  | Raptor | R1WA-c-4 |
| loglogistic | l ~ 1 | s ~ 1 | 2 | 26 | 116.81 | 0 | * | Game Bird | R1WA-c-5 |
| lognormal | l ~ 1 | s ~ 1 | 2 | 26 | 117.44 | 0.63 |  | Game Bird | R1WA-c-5 |
| exponential | l ~ 1 | NULL | 1 | 26 | 119.49 | 2.68 |  | Game Bird | R1WA-c-5 |
| Weibull | l ~ 1 | s ~ 1 | 2 | 26 | 121.41 | 4.6 |  | Game Bird | R1WA-c-5 |
| exponential | l ~ 1 | NULL | 1 | 13 | 62.97 | 0 | * | Raptor | R1WA-f-1 |
| Weibull | l ~ 1 | s ~ 1 | 2 | 13 | 65.8 | 2.83 |  | Raptor | R1WA-f-1 |
| loglogistic | l ~ 1 | s ~ 1 | 2 | 13 | 66.12 | 3.15 |  | Raptor | R1WA-f-1 |
| lognormal | l ~ 1 | s ~ 1 | 2 | 13 | 66.45 | 3.48 |  | Raptor | R1WA-f-1 |
| exponential | l ~ 1 | NULL | 1 | 32 | 92.21 | 0 | * | Raptor | R1WA-g-1 |
| Weibull | l ~ 1 | s ~ 1 | 2 | 32 | 92.9 | 0.69 |  | Raptor | R1WA-g-1 |
| loglogistic | l ~ 1 | s ~ 1 | 2 | 32 | 93.35 | 1.14 |  | Raptor | R1WA-g-1 |
| lognormal | l ~ 1 | s ~ 1 | 2 | 32 | 93.73 | 1.52 |  | Raptor | R1WA-g-1 |
| exponential | l ~ 1 | NULL | 1 | 32 | 146.96 | 0 | * | Game Bird | R1WA-g-6 |
| Weibull | l ~ 1 | s ~ 1 | 2 | 32 | 149.01 | 2.05 |  | Game Bird | R1WA-g-6 |
| loglogistic | l ~ 1 | s ~ 1 | 2 | 32 | 149.47 | 2.51 |  | Game Bird | R1WA-g-6 |
| lognormal | l ~ 1 | s ~ 1 | 2 | 32 | 150.7 | 3.74 |  | Game Bird | R1WA-g-6 |
| Weibull | l ~ Season | s ~ Season | 6 | 82 | 311.89 | 0 |  | Game Bird | R1WA-g-7 |
| Weibull | l ~ 1 | s ~ Season | 4 | 82 | 312.06 | 0.17 | * | Game Bird | R1WA-g-7 |
| lognormal | l ~ 1 | s ~ Season | 4 | 82 | 316.69 | 4.8 |  | Game Bird | R1WA-g-7 |
| loglogistic | l ~ 1 | s ~ Season | 4 | 82 | 319.01 | 7.12 |  | Game Bird | R1WA-g-7 |
| lognormal | l ~ Season | s ~ Season | 6 | 82 | 319.2 | 7.31 |  | Game Bird | R1WA-g-7 |
| loglogistic | l ~ Season | s ~ Season | 6 | 82 | 321.68 | 9.79 |  | Game Bird | R1WA-g-7 |
| exponential | l ~ Season | NULL | 3 | 82 | 327.48 | 15.59 |  | Game Bird | R1WA-g-7 |
| Weibull | l ~ Season | s ~ 1 | 4 | 82 | 327.55 | 15.66 |  | Game Bird | R1WA-g-7 |
| exponential | l ~ 1 | NULL | 1 | 82 | 328.32 | 16.43 |  | Game Bird | R1WA-g-7 |
| Weibull | l ~ 1 | s ~ 1 | 2 | 82 | 330.04 | 18.15 |  | Game Bird | R1WA-g-7 |
| lognormal | l ~ 1 | s ~ 1 | 2 | 82 | 330.2 | 18.31 |  | Game Bird | R1WA-g-7 |
| lognormal | l ~ Season | s ~ 1 | 4 | 82 | 331.56 | 19.67 |  | Game Bird | R1WA-g-7 |
| loglogistic | l ~ 1 | s ~ 1 | 2 | 82 | 331.7 | 19.81 |  | Game Bird | R1WA-g-7 |
| loglogistic | l ~ Season | s ~ 1 | 4 | 82 | 333.52 | 21.63 |  | Game Bird | R1WA-g-7 |
| lognormal | l ~ 1 | s ~ 1 | 2 | 12 | 57.94 | 0 |  | Raptor | R1WA-g-7 |
| exponential | l ~ 1 | NULL | 1 | 12 | 57.99 | 0.05 | * | Raptor | R1WA-g-7 |
| loglogistic | l ~ 1 | s ~ 1 | 2 | 12 | 58.23 | 0.29 |  | Raptor | R1WA-g-7 |
| Weibull | l ~ 1 | s ~ 1 | 2 | 12 | 60.91 | 2.97 |  | Raptor | R1WA-g-7 |
| lognormal | l ~ 1 | s ~ 1 | 2 | 30 | 176.18 | 0 | * | Game Bird | R1WA-g-8 |
| loglogistic | l ~ 1 | s ~ 1 | 2 | 30 | 177.06 | 0.88 |  | Game Bird | R1WA-g-8 |
| Weibull | l ~ 1 | s ~ 1 | 2 | 30 | 177.32 | 1.14 |  | Game Bird | R1WA-g-8 |
| exponential | l ~ 1 | NULL | 1 | 30 | 186.48 | 10.3 |  | Game Bird | R1WA-g-8 |
| Weibull | l ~ 1 | s ~ 1 | 2 | 29 | 192.63 | 0 |  | Raptor | R1WA-g-8 |
| exponential | l ~ 1 | NULL | 1 | 29 | 192.69 | 0.06 | * | Raptor | R1WA-g-8 |
| loglogistic | l ~ 1 | s ~ 1 | 2 | 29 | 200.74 | 8.11 |  | Raptor | R1WA-g-8 |
| lognormal | l ~ 1 | s ~ 1 | 2 | 29 | 201.44 | 8.81 |  | Raptor | R1WA-g-8 |
| Weibull | l ~ 1 | s ~ 1 | 2 | 18 | 67.57 | 0 | * | Raptor | R1WA-ss-2 |
| loglogistic | l ~ 1 | s ~ 1 | 2 | 18 | 67.61 | 0.04 |  | Raptor | R1WA-ss-2 |
| lognormal | l ~ 1 | s ~ 1 | 2 | 18 | 68.11 | 0.54 |  | Raptor | R1WA-ss-2 |
| exponential | l ~ 1 | NULL | 1 | 18 | 74.63 | 7.06 |  | Raptor | R1WA-ss-2 |
| lognormal | l ~ 1 | s ~ 1 | 2 | 40 | 158.59 | 0 | * | Raptor | R1WA-ss-3 |
| loglogistic | l ~ 1 | s ~ 1 | 2 | 40 | 158.64 | 0.05 |  | Raptor | R1WA-ss-3 |
| Weibull | l ~ 1 | s ~ 1 | 2 | 40 | 159.07 | 0.48 |  | Raptor | R1WA-ss-3 |
| loglogistic | l ~ Season | s ~ 1 | 3 | 40 | 160.68 | 2.09 |  | Raptor | R1WA-ss-3 |
| lognormal | l ~ Season | s ~ 1 | 3 | 40 | 160.71 | 2.12 |  | Raptor | R1WA-ss-3 |
| loglogistic | l ~ 1 | s ~ Season | 3 | 40 | 160.87 | 2.28 |  | Raptor | R1WA-ss-3 |
| lognormal | l ~ 1 | s ~ Season | 3 | 40 | 160.88 | 2.29 |  | Raptor | R1WA-ss-3 |
| Weibull | l ~ Season | s ~ 1 | 3 | 40 | 161.05 | 2.46 |  | Raptor | R1WA-ss-3 |
| Weibull | l ~ 1 | s ~ Season | 3 | 40 | 161.41 | 2.82 |  | Raptor | R1WA-ss-3 |
| loglogistic | l ~ Season | s ~ Season | 4 | 40 | 162.96 | 4.37 |  | Raptor | R1WA-ss-3 |
| lognormal | l ~ Season | s ~ Season | 4 | 40 | 163.04 | 4.45 |  | Raptor | R1WA-ss-3 |
| Weibull | l ~ Season | s ~ Season | 4 | 40 | 163.48 | 4.89 |  | Raptor | R1WA-ss-3 |
| exponential | l ~ 1 | NULL | 1 | 40 | 164.12 | 5.53 |  | Raptor | R1WA-ss-3 |
| exponential | l ~ Season | NULL | 2 | 40 | 165.81 | 7.22 |  | Raptor | R1WA-ss-3 |
| loglogistic | l ~ 1 | s ~ 1 | 2 | 63 | 258.86 | 0 | * | Game Bird | R1WA-ss-4 |
| lognormal | l ~ 1 | s ~ 1 | 2 | 63 | 259.23 | 0.37 |  | Game Bird | R1WA-ss-4 |
| Weibull | l ~ 1 | s ~ 1 | 2 | 63 | 265.25 | 6.39 |  | Game Bird | R1WA-ss-4 |
| exponential | l ~ 1 | NULL | 1 | 63 | 271.92 | 13.06 |  | Game Bird | R1WA-ss-4 |
| lognormal | l ~ Season | s ~ Season | 8 | 40 | 171.79 | 0 |  | Game Bird | R2NM-g-1 |
| loglogistic | l ~ Season | s ~ Season | 8 | 40 | 171.97 | 0.18 |  | Game Bird | R2NM-g-1 |
| Weibull | l ~ Season | s ~ 1 | 5 | 40 | 172.61 | 0.82 | * | Game Bird | R2NM-g-1 |
| Weibull | l ~ Season | s ~ Season | 8 | 40 | 173.86 | 2.07 |  | Game Bird | R2NM-g-1 |
| exponential | l ~ Season | NULL | 4 | 40 | 174.64 | 2.85 |  | Game Bird | R2NM-g-1 |
| lognormal | l ~ 1 | s ~ Season | 5 | 40 | 174.96 | 3.17 |  | Game Bird | R2NM-g-1 |
| lognormal | l ~ Season | s ~ 1 | 5 | 40 | 175.33 | 3.54 |  | Game Bird | R2NM-g-1 |
| loglogistic | l ~ Season | s ~ 1 | 5 | 40 | 175.61 | 3.82 |  | Game Bird | R2NM-g-1 |
| loglogistic | l ~ 1 | s ~ Season | 5 | 40 | 175.76 | 3.97 |  | Game Bird | R2NM-g-1 |
| lognormal | l ~ 1 | s ~ 1 | 2 | 40 | 176.07 | 4.28 |  | Game Bird | R2NM-g-1 |
| loglogistic | l ~ 1 | s ~ 1 | 2 | 40 | 177 | 5.21 |  | Game Bird | R2NM-g-1 |
| Weibull | l ~ 1 | s ~ Season | 5 | 40 | 178.84 | 7.05 |  | Game Bird | R2NM-g-1 |
| Weibull | l ~ 1 | s ~ 1 | 2 | 40 | 179.13 | 7.34 |  | Game Bird | R2NM-g-1 |
| exponential | l ~ 1 | NULL | 1 | 40 | 189.89 | 18.1 |  | Game Bird | R2NM-g-1 |
| Weibull | l ~ 1 | s ~ 1 | 2 | 40 | 121.52 | 0 |  | Raptor | R2NM-g-1 |
| loglogistic | l ~ 1 | s ~ 1 | 2 | 40 | 121.66 | 0.14 |  | Raptor | R2NM-g-1 |
| lognormal | l ~ 1 | s ~ 1 | 2 | 40 | 121.84 | 0.32 |  | Raptor | R2NM-g-1 |
| exponential | l ~ 1 | NULL | 1 | 40 | 122.95 | 1.43 | * | Raptor | R2NM-g-1 |
| Weibull | l ~ Season | s ~ 1 | 5 | 40 | 125.38 | 3.86 |  | Raptor | R2NM-g-1 |
| lognormal | l ~ Season | s ~ 1 | 5 | 40 | 125.45 | 3.93 |  | Raptor | R2NM-g-1 |
| loglogistic | l ~ Season | s ~ 1 | 5 | 40 | 125.66 | 4.14 |  | Raptor | R2NM-g-1 |
| exponential | l ~ Season | NULL | 4 | 40 | 126.17 | 4.65 |  | Raptor | R2NM-g-1 |
| Weibull | l ~ 1 | s ~ Season | 5 | 40 | 126.41 | 4.89 |  | Raptor | R2NM-g-1 |
| lognormal | l ~ 1 | s ~ Season | 5 | 40 | 127.43 | 5.91 |  | Raptor | R2NM-g-1 |
| loglogistic | l ~ 1 | s ~ Season | 5 | 40 | 127.57 | 6.05 |  | Raptor | R2NM-g-1 |
| Weibull | l ~ Season | s ~ Season | 8 | 40 | 133.98 | 12.46 |  | Raptor | R2NM-g-1 |
| lognormal | l ~ Season | s ~ Season | 8 | 40 | 134.21 | 12.69 |  | Raptor | R2NM-g-1 |
| loglogistic | l ~ Season | s ~ Season | 8 | 40 | 134.37 | 12.85 |  | Raptor | R2NM-g-1 |
| Weibull | l ~ Season | s ~ Season | 8 | 78 | 280.82 | 0 | * | Game Bird | R2OK-g-0 |
| Weibull | l ~ 1 | s ~ Season | 5 | 78 | 283.37 | 2.55 |  | Game Bird | R2OK-g-0 |
| exponential | l ~ Season | NULL | 4 | 78 | 283.94 | 3.12 |  | Game Bird | R2OK-g-0 |
| Weibull | l ~ Season | s ~ 1 | 5 | 78 | 285.61 | 4.79 |  | Game Bird | R2OK-g-0 |
| lognormal | l ~ Season | s ~ Season | 8 | 78 | 288.44 | 7.62 |  | Game Bird | R2OK-g-0 |
| lognormal | l ~ 1 | s ~ Season | 5 | 78 | 289.9 | 9.08 |  | Game Bird | R2OK-g-0 |
| lognormal | l ~ Season | s ~ 1 | 5 | 78 | 290.15 | 9.33 |  | Game Bird | R2OK-g-0 |
| loglogistic | l ~ Season | s ~ Season | 8 | 78 | 290.29 | 9.47 |  | Game Bird | R2OK-g-0 |
| exponential | l ~ 1 | NULL | 1 | 78 | 291.38 | 10.56 |  | Game Bird | R2OK-g-0 |
| loglogistic | l ~ 1 | s ~ Season | 5 | 78 | 291.7 | 10.88 |  | Game Bird | R2OK-g-0 |
| loglogistic | l ~ Season | s ~ 1 | 5 | 78 | 291.91 | 11.09 |  | Game Bird | R2OK-g-0 |
| Weibull | l ~ 1 | s ~ 1 | 2 | 78 | 293.01 | 12.19 |  | Game Bird | R2OK-g-0 |
| lognormal | l ~ 1 | s ~ 1 | 2 | 78 | 293.29 | 12.47 |  | Game Bird | R2OK-g-0 |
| loglogistic | l ~ 1 | s ~ 1 | 2 | 78 | 294.63 | 13.81 |  | Game Bird | R2OK-g-0 |
| exponential | l ~ 1 | NULL | 1 | 27 | 102.25 | 0 | * | Raptor | R2OK-g-0 |
| Weibull | l ~ 1 | s ~ 1 | 2 | 27 | 102.63 | 0.38 |  | Raptor | R2OK-g-0 |
| lognormal | l ~ 1 | s ~ 1 | 2 | 27 | 103.83 | 1.58 |  | Raptor | R2OK-g-0 |
| loglogistic | l ~ 1 | s ~ 1 | 2 | 27 | 104.25 | 2 |  | Raptor | R2OK-g-0 |
| exponential | l ~ 1 | NULL | 1 | 41 | 166.47 | 0 | * | Raptor | R2OK-g-4 |
| Weibull | l ~ 1 | s ~ 1 | 2 | 41 | 166.55 | 0.08 |  | Raptor | R2OK-g-4 |
| loglogistic | l ~ 1 | s ~ 1 | 2 | 41 | 166.93 | 0.46 |  | Raptor | R2OK-g-4 |
| lognormal | l ~ 1 | s ~ 1 | 2 | 41 | 167.68 | 1.21 |  | Raptor | R2OK-g-4 |
| lognormal | l ~ 1 | s ~ 1 | 2 | 40 | 176.96 | 0 | * | Game Bird | R2TX-c-1 |
| loglogistic | l ~ 1 | s ~ 1 | 2 | 40 | 177.96 | 1 |  | Game Bird | R2TX-c-1 |
| lognormal | l ~ Season | s ~ 1 | 5 | 40 | 180.1 | 3.14 |  | Game Bird | R2TX-c-1 |
| loglogistic | l ~ Season | s ~ 1 | 5 | 40 | 180.41 | 3.45 |  | Game Bird | R2TX-c-1 |
| lognormal | l ~ 1 | s ~ Season | 5 | 40 | 180.49 | 3.53 |  | Game Bird | R2TX-c-1 |
| loglogistic | l ~ 1 | s ~ Season | 5 | 40 | 181.42 | 4.46 |  | Game Bird | R2TX-c-1 |
| Weibull | l ~ 1 | s ~ 1 | 2 | 40 | 182.75 | 5.79 |  | Game Bird | R2TX-c-1 |
| Weibull | l ~ Season | s ~ 1 | 5 | 40 | 185.29 | 8.33 |  | Game Bird | R2TX-c-1 |
| lognormal | l ~ Season | s ~ Season | 8 | 40 | 185.3 | 8.34 |  | Game Bird | R2TX-c-1 |
| loglogistic | l ~ Season | s ~ Season | 8 | 40 | 185.42 | 8.46 |  | Game Bird | R2TX-c-1 |
| exponential | l ~ Season | NULL | 4 | 40 | 186.53 | 9.57 |  | Game Bird | R2TX-c-1 |
| exponential | l ~ 1 | NULL | 1 | 40 | 187.25 | 10.29 |  | Game Bird | R2TX-c-1 |
| Weibull | l ~ 1 | s ~ Season | 5 | 40 | 187.41 | 10.45 |  | Game Bird | R2TX-c-1 |
| Weibull | l ~ Season | s ~ Season | 8 | 40 | 191.67 | 14.71 |  | Game Bird | R2TX-c-1 |
| exponential | l ~ 1 | NULL | 1 | 40 | 121.28 | 0 | * | Raptor | R2TX-c-1 |
| Weibull | l ~ 1 | s ~ 1 | 2 | 40 | 122.45 | 1.17 |  | Raptor | R2TX-c-1 |
| exponential | l ~ Season | NULL | 4 | 40 | 122.58 | 1.3 |  | Raptor | R2TX-c-1 |
| loglogistic | l ~ 1 | s ~ 1 | 2 | 40 | 122.72 | 1.44 |  | Raptor | R2TX-c-1 |
| lognormal | l ~ 1 | s ~ 1 | 2 | 40 | 122.9 | 1.62 |  | Raptor | R2TX-c-1 |
| Weibull | l ~ Season | s ~ 1 | 5 | 40 | 124.55 | 3.27 |  | Raptor | R2TX-c-1 |
| loglogistic | l ~ Season | s ~ 1 | 5 | 40 | 125.47 | 4.19 |  | Raptor | R2TX-c-1 |
| lognormal | l ~ Season | s ~ 1 | 5 | 40 | 126.44 | 5.16 |  | Raptor | R2TX-c-1 |
| Weibull | l ~ 1 | s ~ Season | 5 | 40 | 128.76 | 7.48 |  | Raptor | R2TX-c-1 |
| loglogistic | l ~ 1 | s ~ Season | 5 | 40 | 129.69 | 8.41 |  | Raptor | R2TX-c-1 |
| lognormal | l ~ 1 | s ~ Season | 5 | 40 | 130.05 | 8.77 |  | Raptor | R2TX-c-1 |
| Weibull | l ~ Season | s ~ Season | 8 | 40 | 131.96 | 10.68 |  | Raptor | R2TX-c-1 |
| loglogistic | l ~ Season | s ~ Season | 8 | 40 | 132.35 | 11.07 |  | Raptor | R2TX-c-1 |
| lognormal | l ~ Season | s ~ Season | 8 | 40 | 132.52 | 11.24 |  | Raptor | R2TX-c-1 |
| exponential | l ~ 1 | NULL | 1 | 15 | 15.71 | 0 | * | Raptor | R2TX-g-1 |
| lognormal | l ~ 1 | s ~ 1 | 2 | 15 | 17.17 | 1.46 |  | Raptor | R2TX-g-1 |
| loglogistic | l ~ 1 | s ~ 1 | 2 | 15 | 17.32 | 1.61 |  | Raptor | R2TX-g-1 |
| Weibull | l ~ 1 | s ~ 1 | 2 | 15 | 17.34 | 1.63 |  | Raptor | R2TX-g-1 |
| lognormal | l ~ 1 | s ~ 1 | 2 | 20 | 80.87 | 0 |  | Raptor | R2TX-g-2 |
| loglogistic | l ~ 1 | s ~ 1 | 2 | 20 | 81.68 | 0.81 |  | Raptor | R2TX-g-2 |
| Weibull | l ~ 1 | s ~ 1 | 2 | 20 | 82.64 | 1.77 |  | Raptor | R2TX-g-2 |
| exponential | l ~ 1 | NULL | 1 | 20 | 82.74 | 1.87 | * | Raptor | R2TX-g-2 |
| lognormal | l ~ 1 | s ~ Season | 3 | 20 | 83.05 | 2.18 |  | Raptor | R2TX-g-2 |
| lognormal | l ~ Season | s ~ 1 | 3 | 20 | 83.47 | 2.6 |  | Raptor | R2TX-g-2 |
| loglogistic | l ~ 1 | s ~ Season | 3 | 20 | 83.93 | 3.06 |  | Raptor | R2TX-g-2 |
| loglogistic | l ~ Season | s ~ 1 | 3 | 20 | 84.34 | 3.47 |  | Raptor | R2TX-g-2 |
| Weibull | l ~ 1 | s ~ Season | 3 | 20 | 84.97 | 4.1 |  | Raptor | R2TX-g-2 |
| exponential | l ~ Season | NULL | 2 | 20 | 85.2 | 4.33 |  | Raptor | R2TX-g-2 |
| Weibull | l ~ Season | s ~ 1 | 3 | 20 | 85.4 | 4.53 |  | Raptor | R2TX-g-2 |
| lognormal | l ~ Season | s ~ Season | 4 | 20 | 86.2 | 5.33 |  | Raptor | R2TX-g-2 |
| loglogistic | l ~ Season | s ~ Season | 4 | 20 | 87.08 | 6.21 |  | Raptor | R2TX-g-2 |
| Weibull | l ~ Season | s ~ Season | 4 | 20 | 88.12 | 7.25 |  | Raptor | R2TX-g-2 |
| loglogistic | l ~ 1 | s ~ 1 | 2 | 39 | 168.05 | 0 | * | Game Bird | R2TX-ss-1 |
| lognormal | l ~ 1 | s ~ 1 | 2 | 39 | 168.53 | 0.48 |  | Game Bird | R2TX-ss-1 |
| Weibull | l ~ 1 | s ~ 1 | 2 | 39 | 173.6 | 5.55 |  | Game Bird | R2TX-ss-1 |
| exponential | l ~ 1 | NULL | 1 | 39 | 179.54 | 11.49 |  | Game Bird | R2TX-ss-1 |
| exponential | l ~ 1 | NULL | 1 | 20 | 89.42 | 0 | * | Game Bird | R2TX-ss-2 |
| lognormal | l ~ 1 | s ~ 1 | 2 | 20 | 90.8 | 1.38 |  | Game Bird | R2TX-ss-2 |
| loglogistic | l ~ 1 | s ~ 1 | 2 | 20 | 90.96 | 1.54 |  | Game Bird | R2TX-ss-2 |
| Weibull | l ~ 1 | s ~ 1 | 2 | 20 | 91.53 | 2.11 |  | Game Bird | R2TX-ss-2 |
| exponential | l ~ 1 | NULL | 1 | 73 | 276.98 | 0 | * | Raptor | R3IA-c-0 |
| Weibull | l ~ 1 | s ~ 1 | 2 | 73 | 279 | 2.02 |  | Raptor | R3IA-c-0 |
| loglogistic | l ~ 1 | s ~ 1 | 2 | 73 | 279.11 | 2.13 |  | Raptor | R3IA-c-0 |
| lognormal | l ~ 1 | s ~ 1 | 2 | 73 | 279.39 | 2.41 |  | Raptor | R3IA-c-0 |
| exponential | l ~ 1 | NULL | 1 | 8 | 38.99 | 0 | * | Raptor | R3IA-c-1 |
| Weibull | l ~ 1 | s ~ 1 | 2 | 8 | 42.28 | 3.29 |  | Raptor | R3IA-c-1 |
| lognormal | l ~ 1 | s ~ 1 | 2 | 8 | 42.41 | 3.42 |  | Raptor | R3IA-c-1 |
| loglogistic | l ~ 1 | s ~ 1 | 2 | 8 | 42.59 | 3.6 |  | Raptor | R3IA-c-1 |
| loglogistic | l ~ 1 | s ~ 1 | 2 | 25 | 120.14 | 0 |  | Game Bird | R3IA-c-11 |
| exponential | l ~ 1 | NULL | 1 | 25 | 120.22 | 0.08 | * | Game Bird | R3IA-c-11 |
| lognormal | l ~ 1 | s ~ 1 | 2 | 25 | 120.27 | 0.13 |  | Game Bird | R3IA-c-11 |
| Weibull | l ~ 1 | s ~ 1 | 2 | 25 | 120.34 | 0.2 |  | Game Bird | R3IA-c-11 |
| lognormal | l ~ 1 | s ~ 1 | 2 | 17 | 70.42 | 0 |  | Game Bird | R3IA-c-12 |
| loglogistic | l ~ 1 | s ~ 1 | 2 | 17 | 70.44 | 0.02 |  | Game Bird | R3IA-c-12 |
| exponential | l ~ 1 | NULL | 1 | 17 | 71.83 | 1.41 | * | Game Bird | R3IA-c-12 |
| Weibull | l ~ 1 | s ~ 1 | 2 | 17 | 73.07 | 2.65 |  | Game Bird | R3IA-c-12 |
| exponential | l ~ 1 | NULL | 1 | 31 | 138.74 | 0 | * | Game Bird | R3IA-c-13 |
| Weibull | l ~ 1 | s ~ 1 | 2 | 31 | 140.69 | 1.95 |  | Game Bird | R3IA-c-13 |
| lognormal | l ~ 1 | s ~ 1 | 2 | 31 | 144.59 | 5.85 |  | Game Bird | R3IA-c-13 |
| loglogistic | l ~ 1 | s ~ 1 | 2 | 31 | 145.11 | 6.37 |  | Game Bird | R3IA-c-13 |
| exponential | l ~ 1 | NULL | 1 | 38 | 131.99 | 0 | * | Game Bird | R3IA-c-14 |
| Weibull | l ~ 1 | s ~ 1 | 2 | 38 | 133.97 | 1.98 |  | Game Bird | R3IA-c-14 |
| loglogistic | l ~ 1 | s ~ 1 | 2 | 38 | 134.05 | 2.06 |  | Game Bird | R3IA-c-14 |
| lognormal | l ~ 1 | s ~ 1 | 2 | 38 | 134.67 | 2.68 |  | Game Bird | R3IA-c-14 |
| exponential | l ~ 1 | NULL | 1 | 35 | 111.39 | 0 | * | Raptor | R3IA-c-14 |
| Weibull | l ~ 1 | s ~ 1 | 2 | 35 | 113.65 | 2.26 |  | Raptor | R3IA-c-14 |
| loglogistic | l ~ 1 | s ~ 1 | 2 | 35 | 114.56 | 3.17 |  | Raptor | R3IA-c-14 |
| lognormal | l ~ 1 | s ~ 1 | 2 | 35 | 116.3 | 4.91 |  | Raptor | R3IA-c-14 |
| lognormal | l ~ 1 | s ~ 1 | 2 | 25 | 109.56 | 0 |  | Game Bird | R3IA-c-15 |
| loglogistic | l ~ 1 | s ~ 1 | 2 | 25 | 110.04 | 0.48 |  | Game Bird | R3IA-c-15 |
| exponential | l ~ 1 | NULL | 1 | 25 | 110.66 | 1.1 | * | Game Bird | R3IA-c-15 |
| Weibull | l ~ 1 | s ~ 1 | 2 | 25 | 112.4 | 2.84 |  | Game Bird | R3IA-c-15 |
| exponential | l ~ 1 | NULL | 1 | 30 | 130.81 | 0 | * | Game Bird | R3IA-c-16 |
| Weibull | l ~ 1 | s ~ 1 | 2 | 30 | 132.53 | 1.72 |  | Game Bird | R3IA-c-16 |
| lognormal | l ~ 1 | s ~ 1 | 2 | 30 | 133.35 | 2.54 |  | Game Bird | R3IA-c-16 |
| loglogistic | l ~ 1 | s ~ 1 | 2 | 30 | 133.79 | 2.98 |  | Game Bird | R3IA-c-16 |
| exponential | l ~ 1 | NULL | 1 | 30 | 122.26 | 0 | * | Game Bird | R3IA-c-17 |
| loglogistic | l ~ 1 | s ~ 1 | 2 | 30 | 123.58 | 1.32 |  | Game Bird | R3IA-c-17 |
| Weibull | l ~ 1 | s ~ 1 | 2 | 30 | 123.93 | 1.67 |  | Game Bird | R3IA-c-17 |
| lognormal | l ~ 1 | s ~ 1 | 2 | 30 | 124 | 1.74 |  | Game Bird | R3IA-c-17 |
| exponential | l ~ 1 | NULL | 1 | 11 | 33.65 | 0 | * | Raptor | R3IA-c-17 |
| lognormal | l ~ 1 | s ~ 1 | 2 | 11 | 36.25 | 2.6 |  | Raptor | R3IA-c-17 |
| loglogistic | l ~ 1 | s ~ 1 | 2 | 11 | 36.53 | 2.88 |  | Raptor | R3IA-c-17 |
| Weibull | l ~ 1 | s ~ 1 | 2 | 11 | 36.66 | 3.01 |  | Raptor | R3IA-c-17 |
| exponential | l ~ 1 | NULL | 1 | 38 | 162.45 | 0 | * | Game Bird | R3IA-c-19 |
| Weibull | l ~ 1 | s ~ 1 | 2 | 38 | 164.61 | 2.16 |  | Game Bird | R3IA-c-19 |
| loglogistic | l ~ 1 | s ~ 1 | 2 | 38 | 164.65 | 2.2 |  | Game Bird | R3IA-c-19 |
| lognormal | l ~ 1 | s ~ 1 | 2 | 38 | 165.33 | 2.88 |  | Game Bird | R3IA-c-19 |
| exponential | l ~ 1 | NULL | 1 | 10 | 30.09 | 0 | * | Raptor | R3IA-c-19 |
| lognormal | l ~ 1 | s ~ 1 | 2 | 10 | 30.86 | 0.77 |  | Raptor | R3IA-c-19 |
| loglogistic | l ~ 1 | s ~ 1 | 2 | 10 | 31.02 | 0.93 |  | Raptor | R3IA-c-19 |
| Weibull | l ~ 1 | s ~ 1 | 2 | 10 | 31.47 | 1.38 |  | Raptor | R3IA-c-19 |
| exponential | l ~ 1 | NULL | 1 | 25 | 99.56 | 0 | * | Raptor | R3IA-c-2 |
| lognormal | l ~ 1 | s ~ 1 | 2 | 25 | 101.57 | 2.01 |  | Raptor | R3IA-c-2 |
| loglogistic | l ~ 1 | s ~ 1 | 2 | 25 | 101.72 | 2.16 |  | Raptor | R3IA-c-2 |
| Weibull | l ~ 1 | s ~ 1 | 2 | 25 | 101.82 | 2.26 |  | Raptor | R3IA-c-2 |
| exponential | l ~ Season | NULL | 2 | 25 | 101.9 | 2.34 |  | Raptor | R3IA-c-2 |
| lognormal | l ~ 1 | s ~ Season | 3 | 25 | 103.06 | 3.5 |  | Raptor | R3IA-c-2 |
| loglogistic | l ~ 1 | s ~ Season | 3 | 25 | 103.61 | 4.05 |  | Raptor | R3IA-c-2 |
| lognormal | l ~ Season | s ~ 1 | 3 | 25 | 103.92 | 4.36 |  | Raptor | R3IA-c-2 |
| Weibull | l ~ 1 | s ~ Season | 3 | 25 | 104.29 | 4.73 |  | Raptor | R3IA-c-2 |
| loglogistic | l ~ Season | s ~ 1 | 3 | 25 | 104.3 | 4.74 |  | Raptor | R3IA-c-2 |
| Weibull | l ~ Season | s ~ 1 | 3 | 25 | 104.4 | 4.84 |  | Raptor | R3IA-c-2 |
| lognormal | l ~ Season | s ~ Season | 4 | 25 | 105.79 | 6.23 |  | Raptor | R3IA-c-2 |
| loglogistic | l ~ Season | s ~ Season | 4 | 25 | 106.47 | 6.91 |  | Raptor | R3IA-c-2 |
| Weibull | l ~ Season | s ~ Season | 4 | 25 | 107.13 | 7.57 |  | Raptor | R3IA-c-2 |
| loglogistic | l ~ 1 | s ~ 1 | 2 | 23 | 106.31 | 0 |  | Game Bird | R3IA-c-20 |
| lognormal | l ~ 1 | s ~ 1 | 2 | 23 | 106.54 | 0.23 |  | Game Bird | R3IA-c-20 |
| exponential | l ~ 1 | NULL | 1 | 23 | 107.77 | 1.46 | * | Game Bird | R3IA-c-20 |
| Weibull | l ~ 1 | s ~ 1 | 2 | 23 | 108.94 | 2.63 |  | Game Bird | R3IA-c-20 |
| exponential | l ~ 1 | NULL | 1 | 28 | 115.27 | 0 | * | Game Bird | R3IA-c-21 |
| Weibull | l ~ 1 | s ~ 1 | 2 | 28 | 117.31 | 2.04 |  | Game Bird | R3IA-c-21 |
| loglogistic | l ~ 1 | s ~ 1 | 2 | 28 | 118.24 | 2.97 |  | Game Bird | R3IA-c-21 |
| lognormal | l ~ 1 | s ~ 1 | 2 | 28 | 119.1 | 3.83 |  | Game Bird | R3IA-c-21 |
| lognormal | l ~ 1 | s ~ 1 | 2 | 11 | 48.63 | 0 |  | Game Bird | R3IA-c-22 |
| exponential | l ~ 1 | NULL | 1 | 11 | 48.78 | 0.15 | * | Game Bird | R3IA-c-22 |
| loglogistic | l ~ 1 | s ~ 1 | 2 | 11 | 49.18 | 0.55 |  | Game Bird | R3IA-c-22 |
| Weibull | l ~ 1 | s ~ 1 | 2 | 11 | 49.58 | 0.95 |  | Game Bird | R3IA-c-22 |
| exponential | l ~ 1 | NULL | 1 | 28 | 129.93 | 0 | * | Game Bird | R3IA-c-23 |
| Weibull | l ~ 1 | s ~ 1 | 2 | 28 | 131.41 | 1.48 |  | Game Bird | R3IA-c-23 |
| loglogistic | l ~ 1 | s ~ 1 | 2 | 28 | 133 | 3.07 |  | Game Bird | R3IA-c-23 |
| lognormal | l ~ 1 | s ~ 1 | 2 | 28 | 133.01 | 3.08 |  | Game Bird | R3IA-c-23 |
| loglogistic | l ~ 1 | s ~ 1 | 2 | 27 | 101.27 | 0 |  | Game Bird | R3IA-c-24 |
| lognormal | l ~ 1 | s ~ 1 | 2 | 27 | 101.72 | 0.45 |  | Game Bird | R3IA-c-24 |
| Weibull | l ~ 1 | s ~ 1 | 2 | 27 | 101.74 | 0.47 |  | Game Bird | R3IA-c-24 |
| exponential | l ~ 1 | NULL | 1 | 27 | 101.92 | 0.65 | * | Game Bird | R3IA-c-24 |
| exponential | l ~ 1 | NULL | 1 | 11 | 49.44 | 0 | * | Game Bird | R3IA-c-25 |
| lognormal | l ~ 1 | s ~ 1 | 2 | 11 | 49.59 | 0.15 |  | Game Bird | R3IA-c-25 |
| loglogistic | l ~ 1 | s ~ 1 | 2 | 11 | 50.01 | 0.57 |  | Game Bird | R3IA-c-25 |
| Weibull | l ~ 1 | s ~ 1 | 2 | 11 | 52.03 | 2.59 |  | Game Bird | R3IA-c-25 |
| exponential | l ~ 1 | NULL | 1 | 32 | 140.41 | 0 | * | Game Bird | R3IA-c-26 |
| loglogistic | l ~ 1 | s ~ 1 | 2 | 32 | 141.92 | 1.51 |  | Game Bird | R3IA-c-26 |
| Weibull | l ~ 1 | s ~ 1 | 2 | 32 | 142.55 | 2.14 |  | Game Bird | R3IA-c-26 |
| lognormal | l ~ 1 | s ~ 1 | 2 | 32 | 142.55 | 2.14 |  | Game Bird | R3IA-c-26 |
| Weibull | l ~ 1 | s ~ 1 | 2 | 25 | 104.37 | 0 |  | Game Bird | R3IA-c-27 |
| exponential | l ~ 1 | NULL | 1 | 25 | 105.9 | 1.53 | * | Game Bird | R3IA-c-27 |
| lognormal | l ~ 1 | s ~ 1 | 2 | 25 | 107.69 | 3.32 |  | Game Bird | R3IA-c-27 |
| loglogistic | l ~ 1 | s ~ 1 | 2 | 25 | 108.87 | 4.5 |  | Game Bird | R3IA-c-27 |
| exponential | l ~ 1 | NULL | 1 | 28 | 121.98 | 0 | * | Game Bird | R3IA-c-29 |
| loglogistic | l ~ 1 | s ~ 1 | 2 | 28 | 122.71 | 0.73 |  | Game Bird | R3IA-c-29 |
| lognormal | l ~ 1 | s ~ 1 | 2 | 28 | 122.86 | 0.88 |  | Game Bird | R3IA-c-29 |
| Weibull | l ~ 1 | s ~ 1 | 2 | 28 | 124.3 | 2.32 |  | Game Bird | R3IA-c-29 |
| Weibull | l ~ 1 | s ~ 1 | 2 | 26 | 67.14 | 0 |  | Raptor | R3IA-c-3 |
| loglogistic | l ~ 1 | s ~ 1 | 2 | 26 | 67.14 | 0 |  | Raptor | R3IA-c-3 |
| lognormal | l ~ 1 | s ~ 1 | 2 | 26 | 67.34 | 0.2 |  | Raptor | R3IA-c-3 |
| exponential | l ~ 1 | NULL | 1 | 26 | 67.68 | 0.54 | * | Raptor | R3IA-c-3 |
| Weibull | l ~ Season | s ~ 1 | 3 | 26 | 69.66 | 2.52 |  | Raptor | R3IA-c-3 |
| Weibull | l ~ 1 | s ~ Season | 3 | 26 | 69.7 | 2.56 |  | Raptor | R3IA-c-3 |
| loglogistic | l ~ Season | s ~ 1 | 3 | 26 | 69.7 | 2.56 |  | Raptor | R3IA-c-3 |
| loglogistic | l ~ 1 | s ~ Season | 3 | 26 | 69.71 | 2.57 |  | Raptor | R3IA-c-3 |
| lognormal | l ~ 1 | s ~ Season | 3 | 26 | 69.81 | 2.67 |  | Raptor | R3IA-c-3 |
| lognormal | l ~ Season | s ~ 1 | 3 | 26 | 69.86 | 2.72 |  | Raptor | R3IA-c-3 |
| exponential | l ~ Season | NULL | 2 | 26 | 70 | 2.86 |  | Raptor | R3IA-c-3 |
| Weibull | l ~ Season | s ~ Season | 4 | 26 | 72.47 | 5.33 |  | Raptor | R3IA-c-3 |
| loglogistic | l ~ Season | s ~ Season | 4 | 26 | 72.51 | 5.37 |  | Raptor | R3IA-c-3 |
| lognormal | l ~ Season | s ~ Season | 4 | 26 | 72.62 | 5.48 |  | Raptor | R3IA-c-3 |
| exponential | l ~ 1 | NULL | 1 | 24 | 102.47 | 0 | * | Game Bird | R3IA-c-30 |
| Weibull | l ~ 1 | s ~ 1 | 2 | 24 | 103.86 | 1.39 |  | Game Bird | R3IA-c-30 |
| loglogistic | l ~ 1 | s ~ 1 | 2 | 24 | 104.2 | 1.73 |  | Game Bird | R3IA-c-30 |
| lognormal | l ~ 1 | s ~ 1 | 2 | 24 | 104.22 | 1.75 |  | Game Bird | R3IA-c-30 |
| exponential | l ~ Season | NULL | 2 | 25 | 102.26 | 0 | * | Raptor | R3IA-c-31 |
| Weibull | l ~ Season | s ~ 1 | 3 | 25 | 104.86 | 2.6 |  | Raptor | R3IA-c-31 |
| Weibull | l ~ Season | s ~ Season | 4 | 25 | 107.4 | 5.14 |  | Raptor | R3IA-c-31 |
| loglogistic | l ~ Season | s ~ 1 | 3 | 25 | 108.45 | 6.19 |  | Raptor | R3IA-c-31 |
| lognormal | l ~ Season | s ~ 1 | 3 | 25 | 109.66 | 7.4 |  | Raptor | R3IA-c-31 |
| lognormal | l ~ Season | s ~ Season | 4 | 25 | 110.23 | 7.97 |  | Raptor | R3IA-c-31 |
| loglogistic | l ~ Season | s ~ Season | 4 | 25 | 110.27 | 8.01 |  | Raptor | R3IA-c-31 |
| loglogistic | l ~ 1 | s ~ Season | 3 | 25 | 114.85 | 12.59 |  | Raptor | R3IA-c-31 |
| lognormal | l ~ 1 | s ~ Season | 3 | 25 | 114.98 | 12.72 |  | Raptor | R3IA-c-31 |
| Weibull | l ~ 1 | s ~ Season | 3 | 25 | 115.79 | 13.53 |  | Raptor | R3IA-c-31 |
| exponential | l ~ 1 | NULL | 1 | 25 | 116.41 | 14.15 |  | Raptor | R3IA-c-31 |
| loglogistic | l ~ 1 | s ~ 1 | 2 | 25 | 116.71 | 14.45 |  | Raptor | R3IA-c-31 |
| lognormal | l ~ 1 | s ~ 1 | 2 | 25 | 116.78 | 14.52 |  | Raptor | R3IA-c-31 |
| Weibull | l ~ 1 | s ~ 1 | 2 | 25 | 116.79 | 14.53 |  | Raptor | R3IA-c-31 |
| exponential | l ~ 1 | NULL | 1 | 22 | 71.54 | 0 | * | Raptor | R3IA-c-32 |
| lognormal | l ~ 1 | s ~ 1 | 2 | 22 | 71.62 | 0.08 |  | Raptor | R3IA-c-32 |
| loglogistic | l ~ 1 | s ~ 1 | 2 | 22 | 72.01 | 0.47 |  | Raptor | R3IA-c-32 |
| Weibull | l ~ 1 | s ~ 1 | 2 | 22 | 72.95 | 1.41 |  | Raptor | R3IA-c-32 |
| lognormal | l ~ 1 | s ~ Season | 3 | 22 | 73.16 | 1.62 |  | Raptor | R3IA-c-32 |
| exponential | l ~ Season | NULL | 2 | 22 | 73.89 | 2.35 |  | Raptor | R3IA-c-32 |
| lognormal | l ~ Season | s ~ 1 | 3 | 22 | 73.9 | 2.36 |  | Raptor | R3IA-c-32 |
| loglogistic | l ~ 1 | s ~ Season | 3 | 22 | 73.92 | 2.38 |  | Raptor | R3IA-c-32 |
| loglogistic | l ~ Season | s ~ 1 | 3 | 22 | 74.43 | 2.89 |  | Raptor | R3IA-c-32 |
| Weibull | l ~ 1 | s ~ Season | 3 | 22 | 74.81 | 3.27 |  | Raptor | R3IA-c-32 |
| Weibull | l ~ Season | s ~ 1 | 3 | 22 | 75.62 | 4.08 |  | Raptor | R3IA-c-32 |
| lognormal | l ~ Season | s ~ Season | 4 | 22 | 76.07 | 4.53 |  | Raptor | R3IA-c-32 |
| loglogistic | l ~ Season | s ~ Season | 4 | 22 | 76.81 | 5.27 |  | Raptor | R3IA-c-32 |
| Weibull | l ~ Season | s ~ Season | 4 | 22 | 77.83 | 6.29 |  | Raptor | R3IA-c-32 |
| exponential | l ~ 1 | NULL | 1 | 12 | 45.95 | 0 | * | Raptor | R3IA-c-33 |
| Weibull | l ~ 1 | s ~ 1 | 2 | 12 | 48.76 | 2.81 |  | Raptor | R3IA-c-33 |
| lognormal | l ~ 1 | s ~ 1 | 2 | 12 | 49.95 | 4 |  | Raptor | R3IA-c-33 |
| loglogistic | l ~ 1 | s ~ 1 | 2 | 12 | 50.01 | 4.06 |  | Raptor | R3IA-c-33 |
| exponential | l ~ 1 | NULL | 1 | 10 | 45.81 | 0 | * | Raptor | R3IA-c-34 |
| lognormal | l ~ 1 | s ~ 1 | 2 | 10 | 47.3 | 1.49 |  | Raptor | R3IA-c-34 |
| loglogistic | l ~ 1 | s ~ 1 | 2 | 10 | 47.58 | 1.77 |  | Raptor | R3IA-c-34 |
| Weibull | l ~ 1 | s ~ 1 | 2 | 10 | 48.89 | 3.08 |  | Raptor | R3IA-c-34 |
| Weibull | l ~ Season | s ~ 1 | 3 | 26 | 101.32 | 0 | * | Raptor | R3IA-c-35 |
| Weibull | l ~ 1 | s ~ 1 | 2 | 26 | 103.48 | 2.16 |  | Raptor | R3IA-c-35 |
| Weibull | l ~ Season | s ~ Season | 4 | 26 | 103.65 | 2.33 |  | Raptor | R3IA-c-35 |
| loglogistic | l ~ Season | s ~ 1 | 3 | 26 | 103.94 | 2.62 |  | Raptor | R3IA-c-35 |
| loglogistic | l ~ 1 | s ~ 1 | 2 | 26 | 104.39 | 3.07 |  | Raptor | R3IA-c-35 |
| Weibull | l ~ 1 | s ~ Season | 3 | 26 | 105.11 | 3.79 |  | Raptor | R3IA-c-35 |
| loglogistic | l ~ 1 | s ~ Season | 3 | 26 | 105.24 | 3.92 |  | Raptor | R3IA-c-35 |
| loglogistic | l ~ Season | s ~ Season | 4 | 26 | 105.35 | 4.03 |  | Raptor | R3IA-c-35 |
| lognormal | l ~ Season | s ~ Season | 4 | 26 | 106.3 | 4.98 |  | Raptor | R3IA-c-35 |
| lognormal | l ~ 1 | s ~ Season | 3 | 26 | 106.79 | 5.47 |  | Raptor | R3IA-c-35 |
| exponential | l ~ 1 | NULL | 1 | 26 | 107.35 | 6.03 |  | Raptor | R3IA-c-35 |
| lognormal | l ~ Season | s ~ 1 | 3 | 26 | 107.37 | 6.05 |  | Raptor | R3IA-c-35 |
| exponential | l ~ Season | NULL | 2 | 26 | 107.52 | 6.2 |  | Raptor | R3IA-c-35 |
| lognormal | l ~ 1 | s ~ 1 | 2 | 26 | 109.31 | 7.99 |  | Raptor | R3IA-c-35 |
| exponential | l ~ 1 | NULL | 1 | 15 | 45.52 | 0 | * | Raptor | R3IA-c-36 |
| lognormal | l ~ 1 | s ~ 1 | 2 | 15 | 46.19 | 0.67 |  | Raptor | R3IA-c-36 |
| loglogistic | l ~ 1 | s ~ 1 | 2 | 15 | 46.79 | 1.27 |  | Raptor | R3IA-c-36 |
| Weibull | l ~ 1 | s ~ 1 | 2 | 15 | 47.12 | 1.6 |  | Raptor | R3IA-c-36 |
| exponential | l ~ 1 | NULL | 1 | 20 | 74.48 | 0 | * | Raptor | R3IA-c-4 |
| exponential | l ~ Season | NULL | 2 | 20 | 76 | 1.52 |  | Raptor | R3IA-c-4 |
| lognormal | l ~ 1 | s ~ 1 | 2 | 20 | 76.05 | 1.57 |  | Raptor | R3IA-c-4 |
| loglogistic | l ~ 1 | s ~ 1 | 2 | 20 | 76.75 | 2.27 |  | Raptor | R3IA-c-4 |
| Weibull | l ~ 1 | s ~ 1 | 2 | 20 | 76.96 | 2.48 |  | Raptor | R3IA-c-4 |
| lognormal | l ~ 1 | s ~ Season | 3 | 20 | 78.37 | 3.89 |  | Raptor | R3IA-c-4 |
| lognormal | l ~ Season | s ~ 1 | 3 | 20 | 78.4 | 3.92 |  | Raptor | R3IA-c-4 |
| Weibull | l ~ Season | s ~ 1 | 3 | 20 | 78.79 | 4.31 |  | Raptor | R3IA-c-4 |
| loglogistic | l ~ 1 | s ~ Season | 3 | 20 | 78.97 | 4.49 |  | Raptor | R3IA-c-4 |
| loglogistic | l ~ Season | s ~ 1 | 3 | 20 | 79.18 | 4.7 |  | Raptor | R3IA-c-4 |
| Weibull | l ~ 1 | s ~ Season | 3 | 20 | 79.33 | 4.85 |  | Raptor | R3IA-c-4 |
| lognormal | l ~ Season | s ~ Season | 4 | 20 | 80.83 | 6.35 |  | Raptor | R3IA-c-4 |
| Weibull | l ~ Season | s ~ Season | 4 | 20 | 81.28 | 6.8 |  | Raptor | R3IA-c-4 |
| loglogistic | l ~ Season | s ~ Season | 4 | 20 | 81.65 | 7.17 |  | Raptor | R3IA-c-4 |
| lognormal | l ~ 1 | s ~ 1 | 2 | 20 | 94.84 | 0 |  | Raptor | R3IA-c-5 |
| loglogistic | l ~ 1 | s ~ 1 | 2 | 20 | 95.3 | 0.46 |  | Raptor | R3IA-c-5 |
| exponential | l ~ 1 | NULL | 1 | 20 | 95.62 | 0.78 | * | Raptor | R3IA-c-5 |
| Weibull | l ~ 1 | s ~ 1 | 2 | 20 | 95.66 | 0.82 |  | Raptor | R3IA-c-5 |
| lognormal | l ~ 1 | s ~ Season | 3 | 20 | 97.43 | 2.59 |  | Raptor | R3IA-c-5 |
| lognormal | l ~ Season | s ~ 1 | 3 | 20 | 97.53 | 2.69 |  | Raptor | R3IA-c-5 |
| Weibull | l ~ 1 | s ~ Season | 3 | 20 | 97.65 | 2.81 |  | Raptor | R3IA-c-5 |
| loglogistic | l ~ Season | s ~ 1 | 3 | 20 | 97.8 | 2.96 |  | Raptor | R3IA-c-5 |
| loglogistic | l ~ 1 | s ~ Season | 3 | 20 | 97.98 | 3.14 |  | Raptor | R3IA-c-5 |
| exponential | l ~ Season | NULL | 2 | 20 | 98.08 | 3.24 |  | Raptor | R3IA-c-5 |
| Weibull | l ~ Season | s ~ 1 | 3 | 20 | 98.45 | 3.61 |  | Raptor | R3IA-c-5 |
| lognormal | l ~ Season | s ~ Season | 4 | 20 | 100.48 | 5.64 |  | Raptor | R3IA-c-5 |
| Weibull | l ~ Season | s ~ Season | 4 | 20 | 100.78 | 5.94 |  | Raptor | R3IA-c-5 |
| loglogistic | l ~ Season | s ~ Season | 4 | 20 | 100.84 | 6 |  | Raptor | R3IA-c-5 |
| exponential | l ~ 1 | NULL | 1 | 21 | 95.58 | 0 | * | Game Bird | R3IA-c-6 |
| loglogistic | l ~ 1 | s ~ 1 | 2 | 21 | 97.41 | 1.83 |  | Game Bird | R3IA-c-6 |
| Weibull | l ~ 1 | s ~ 1 | 2 | 21 | 97.82 | 2.24 |  | Game Bird | R3IA-c-6 |
| lognormal | l ~ 1 | s ~ 1 | 2 | 21 | 98.06 | 2.48 |  | Game Bird | R3IA-c-6 |
| exponential | l ~ 1 | NULL | 1 | 15 | 67.3 | 0 | * | Game Bird | R3IA-c-7 |
| loglogistic | l ~ 1 | s ~ 1 | 2 | 15 | 68.18 | 0.88 |  | Game Bird | R3IA-c-7 |
| Weibull | l ~ 1 | s ~ 1 | 2 | 15 | 68.46 | 1.16 |  | Game Bird | R3IA-c-7 |
| lognormal | l ~ 1 | s ~ 1 | 2 | 15 | 68.76 | 1.46 |  | Game Bird | R3IA-c-7 |
| exponential | l ~ 1 | NULL | 1 | 12 | 52.6 | 0 | * | Game Bird | R3IA-c-8 |
| Weibull | l ~ 1 | s ~ 1 | 2 | 12 | 54.86 | 2.26 |  | Game Bird | R3IA-c-8 |
| loglogistic | l ~ 1 | s ~ 1 | 2 | 12 | 55.61 | 3.01 |  | Game Bird | R3IA-c-8 |
| lognormal | l ~ 1 | s ~ 1 | 2 | 12 | 56.28 | 3.68 |  | Game Bird | R3IA-c-8 |
| exponential | l ~ 1 | NULL | 1 | 10 | 47.65 | 0 | * | Game Bird | R3IA-c-9 |
| lognormal | l ~ 1 | s ~ 1 | 2 | 10 | 48.92 | 1.27 |  | Game Bird | R3IA-c-9 |
| loglogistic | l ~ 1 | s ~ 1 | 2 | 10 | 49.08 | 1.43 |  | Game Bird | R3IA-c-9 |
| Weibull | l ~ 1 | s ~ 1 | 2 | 10 | 50.72 | 3.07 |  | Game Bird | R3IA-c-9 |
| lognormal | l ~ 1 | s ~ 1 | 2 | 65 | 236.69 | 0 | * | Raptor | R3IA-c-9 |
| loglogistic | l ~ 1 | s ~ 1 | 2 | 65 | 237.21 | 0.52 |  | Raptor | R3IA-c-9 |
| exponential | l ~ 1 | NULL | 1 | 65 | 241.28 | 4.59 |  | Raptor | R3IA-c-9 |
| Weibull | l ~ 1 | s ~ 1 | 2 | 65 | 243.03 | 6.34 |  | Raptor | R3IA-c-9 |
| exponential | l ~ 1 | NULL | 1 | 15 | 67.37 | 0 | * | Game Bird | R3IL-c-1 |
| Weibull | l ~ 1 | s ~ 1 | 2 | 15 | 69.25 | 1.88 |  | Game Bird | R3IL-c-1 |
| lognormal | l ~ 1 | s ~ 1 | 2 | 15 | 70.4 | 3.03 |  | Game Bird | R3IL-c-1 |
| loglogistic | l ~ 1 | s ~ 1 | 2 | 15 | 70.83 | 3.46 |  | Game Bird | R3IL-c-1 |
| exponential | l ~ 1 | NULL | 1 | 3 | 20.95 | 0 | * | Raptor | R3IL-c-1 |
| Weibull | l ~ 1 | s ~ 1 | 2 | 3 | Inf | Inf |  | Raptor | R3IL-c-1 |
| loglogistic | l ~ 1 | s ~ 1 | 2 | 3 | Inf | Inf |  | Raptor | R3IL-c-1 |
| lognormal | l ~ 1 | s ~ 1 | 2 | 3 | Inf | Inf |  | Raptor | R3IL-c-1 |
| exponential | l ~ 1 | NULL | 1 | 16 | 48.02 | 0 | * | Game Bird | R3MI-c-1 |
| Weibull | l ~ 1 | s ~ 1 | 2 | 16 | 48.36 | 0.34 |  | Game Bird | R3MI-c-1 |
| lognormal | l ~ 1 | s ~ 1 | 2 | 16 | 49.45 | 1.43 |  | Game Bird | R3MI-c-1 |
| loglogistic | l ~ 1 | s ~ 1 | 2 | 16 | 50.05 | 2.03 |  | Game Bird | R3MI-c-1 |
| exponential | l ~ 1 | NULL | 1 | 15 | 58.62 | 0 | * | Raptor | R3MI-c-1 |
| Weibull | l ~ 1 | s ~ 1 | 2 | 15 | 60.97 | 2.35 |  | Raptor | R3MI-c-1 |
| lognormal | l ~ 1 | s ~ 1 | 2 | 15 | 62.71 | 4.09 |  | Raptor | R3MI-c-1 |
| loglogistic | l ~ 1 | s ~ 1 | 2 | 15 | 62.9 | 4.28 |  | Raptor | R3MI-c-1 |
| exponential | l ~ 1 | NULL | 1 | 22 | 106.96 | 0 | * | Raptor | R3MN-c-0 |
| Weibull | l ~ 1 | s ~ 1 | 2 | 22 | 109.01 | 2.05 |  | Raptor | R3MN-c-0 |
| loglogistic | l ~ 1 | s ~ 1 | 2 | 22 | 111.58 | 4.62 |  | Raptor | R3MN-c-0 |
| lognormal | l ~ 1 | s ~ 1 | 2 | 22 | 114.73 | 7.77 |  | Raptor | R3MN-c-0 |
| lognormal | l ~ 1 | s ~ 1 | 2 | 20 | 105.69 | 0 |  | Raptor | R3MN-c-1 |
| loglogistic | l ~ 1 | s ~ 1 | 2 | 20 | 106.5 | 0.81 |  | Raptor | R3MN-c-1 |
| exponential | l ~ 1 | NULL | 1 | 20 | 106.93 | 1.24 | * | Raptor | R3MN-c-1 |
| Weibull | l ~ 1 | s ~ 1 | 2 | 20 | 107.07 | 1.38 |  | Raptor | R3MN-c-1 |
| lognormal | l ~ 1 | s ~ 1 | 2 | 15 | 64.87 | 0 |  | Raptor | R3MN-c-2 |
| loglogistic | l ~ 1 | s ~ 1 | 2 | 15 | 65.2 | 0.33 |  | Raptor | R3MN-c-2 |
| exponential | l ~ 1 | NULL | 1 | 15 | 65.62 | 0.75 | * | Raptor | R3MN-c-2 |
| Weibull | l ~ 1 | s ~ 1 | 2 | 15 | 66.08 | 1.21 |  | Raptor | R3MN-c-2 |
| lognormal | l ~ Season | s ~ 1 | 3 | 32 | 128.48 | 0 |  | Game Bird | R3MN-c-4 |
| loglogistic | l ~ Season | s ~ 1 | 3 | 32 | 128.7 | 0.22 |  | Game Bird | R3MN-c-4 |
| Weibull | l ~ Season | s ~ 1 | 3 | 32 | 129.98 | 1.5 |  | Game Bird | R3MN-c-4 |
| exponential | l ~ Season | NULL | 2 | 32 | 130.07 | 1.59 | * | Game Bird | R3MN-c-4 |
| lognormal | l ~ Season | s ~ Season | 4 | 32 | 130.65 | 2.17 |  | Game Bird | R3MN-c-4 |
| loglogistic | l ~ Season | s ~ Season | 4 | 32 | 130.98 | 2.5 |  | Game Bird | R3MN-c-4 |
| Weibull | l ~ Season | s ~ Season | 4 | 32 | 131.68 | 3.2 |  | Game Bird | R3MN-c-4 |
| loglogistic | l ~ 1 | s ~ 1 | 2 | 32 | 132.16 | 3.68 |  | Game Bird | R3MN-c-4 |
| lognormal | l ~ 1 | s ~ 1 | 2 | 32 | 132.62 | 4.14 |  | Game Bird | R3MN-c-4 |
| Weibull | l ~ 1 | s ~ 1 | 2 | 32 | 132.97 | 4.49 |  | Game Bird | R3MN-c-4 |
| Weibull | l ~ 1 | s ~ Season | 3 | 32 | 133.15 | 4.67 |  | Game Bird | R3MN-c-4 |
| lognormal | l ~ 1 | s ~ Season | 3 | 32 | 133.34 | 4.86 |  | Game Bird | R3MN-c-4 |
| loglogistic | l ~ 1 | s ~ Season | 3 | 32 | 133.59 | 5.11 |  | Game Bird | R3MN-c-4 |
| exponential | l ~ 1 | NULL | 1 | 32 | 134.49 | 6.01 |  | Game Bird | R3MN-c-4 |
| exponential | l ~ 1 | NULL | 1 | 33 | 151.97 | 0 | * | Raptor | R3MO-c-1 |
| Weibull | l ~ 1 | s ~ 1 | 2 | 33 | 154.22 | 2.25 |  | Raptor | R3MO-c-1 |
| lognormal | l ~ 1 | s ~ 1 | 2 | 33 | 157.18 | 5.21 |  | Raptor | R3MO-c-1 |
| loglogistic | l ~ 1 | s ~ 1 | 2 | 33 | 158.21 | 6.24 |  | Raptor | R3MO-c-1 |
| exponential | l ~ 1 | NULL | 1 | 78 | 373.67 | 0 | * | Raptor | R3MO-c-2 |
| Weibull | l ~ 1 | s ~ 1 | 2 | 78 | 375.78 | 2.11 |  | Raptor | R3MO-c-2 |
| loglogistic | l ~ 1 | s ~ 1 | 2 | 78 | 381.55 | 7.88 |  | Raptor | R3MO-c-2 |
| lognormal | l ~ 1 | s ~ 1 | 2 | 78 | 384.64 | 10.97 |  | Raptor | R3MO-c-2 |
| loglogistic | l ~ 1 | s ~ 1 | 2 | 17 | 73.26 | 0 | * | Game Bird | R3OH-c-1 |
| lognormal | l ~ 1 | s ~ 1 | 2 | 17 | 73.99 | 0.73 |  | Game Bird | R3OH-c-1 |
| Weibull | l ~ 1 | s ~ 1 | 2 | 17 | 77.79 | 4.53 |  | Game Bird | R3OH-c-1 |
| exponential | l ~ 1 | NULL | 1 | 17 | 79.36 | 6.1 |  | Game Bird | R3OH-c-1 |
| lognormal | l ~ 1 | s ~ 1 | 2 | 8 | 13.85 | 0 |  | Raptor | R3OH-c-1 |
| loglogistic | l ~ 1 | s ~ 1 | 2 | 8 | 13.99 | 0.14 |  | Raptor | R3OH-c-1 |
| Weibull | l ~ 1 | s ~ 1 | 2 | 8 | 14.02 | 0.17 |  | Raptor | R3OH-c-1 |
| exponential | l ~ 1 | NULL | 1 | 8 | 15.63 | 1.78 | * | Raptor | R3OH-c-1 |
| loglogistic | l ~ 1 | s ~ Season | 5 | 40 | 139.49 | 0 |  | Game Bird | R4NC-c-1 |
| lognormal | l ~ 1 | s ~ Season | 5 | 40 | 139.67 | 0.18 |  | Game Bird | R4NC-c-1 |
| loglogistic | l ~ 1 | s ~ 1 | 2 | 40 | 140.09 | 0.6 | * | Game Bird | R4NC-c-1 |
| lognormal | l ~ 1 | s ~ 1 | 2 | 40 | 140.15 | 0.66 |  | Game Bird | R4NC-c-1 |
| Weibull | l ~ 1 | s ~ 1 | 2 | 40 | 141.18 | 1.69 |  | Game Bird | R4NC-c-1 |
| Weibull | l ~ 1 | s ~ Season | 5 | 40 | 143.37 | 3.88 |  | Game Bird | R4NC-c-1 |
| loglogistic | l ~ Season | s ~ 1 | 5 | 40 | 146.88 | 7.39 |  | Game Bird | R4NC-c-1 |
| lognormal | l ~ Season | s ~ 1 | 5 | 40 | 147.02 | 7.53 |  | Game Bird | R4NC-c-1 |
| Weibull | l ~ Season | s ~ 1 | 5 | 40 | 147.07 | 7.58 |  | Game Bird | R4NC-c-1 |
| loglogistic | l ~ Season | s ~ Season | 8 | 40 | 147.53 | 8.04 |  | Game Bird | R4NC-c-1 |
| lognormal | l ~ Season | s ~ Season | 8 | 40 | 147.88 | 8.39 |  | Game Bird | R4NC-c-1 |
| exponential | l ~ 1 | NULL | 1 | 40 | 149.43 | 9.94 |  | Game Bird | R4NC-c-1 |
| Weibull | l ~ Season | s ~ Season | 8 | 40 | 150.19 | 10.7 |  | Game Bird | R4NC-c-1 |
| exponential | l ~ Season | NULL | 4 | 40 | 153.15 | 13.66 |  | Game Bird | R4NC-c-1 |
| lognormal | l ~ 1 | s ~ 1 | 2 | 40 | 131.74 | 0 | * | Raptor | R4NC-c-1 |
| loglogistic | l ~ 1 | s ~ 1 | 2 | 40 | 132.2 | 0.46 |  | Raptor | R4NC-c-1 |
| Weibull | l ~ 1 | s ~ 1 | 2 | 40 | 132.65 | 0.91 |  | Raptor | R4NC-c-1 |
| lognormal | l ~ Season | s ~ 1 | 5 | 40 | 134.78 | 3.04 |  | Raptor | R4NC-c-1 |
| loglogistic | l ~ Season | s ~ 1 | 5 | 40 | 135.08 | 3.34 |  | Raptor | R4NC-c-1 |
| Weibull | l ~ Season | s ~ 1 | 5 | 40 | 135.75 | 4.01 |  | Raptor | R4NC-c-1 |
| Weibull | l ~ 1 | s ~ Season | 5 | 40 | 138.04 | 6.3 |  | Raptor | R4NC-c-1 |
| lognormal | l ~ 1 | s ~ Season | 5 | 40 | 138.37 | 6.63 |  | Raptor | R4NC-c-1 |
| loglogistic | l ~ 1 | s ~ Season | 5 | 40 | 138.87 | 7.13 |  | Raptor | R4NC-c-1 |
| exponential | l ~ 1 | NULL | 1 | 40 | 141.7 | 9.96 |  | Raptor | R4NC-c-1 |
| exponential | l ~ Season | NULL | 4 | 40 | 142.91 | 11.17 |  | Raptor | R4NC-c-1 |
| lognormal | l ~ Season | s ~ Season | 8 | 40 | 143.23 | 11.49 |  | Raptor | R4NC-c-1 |
| loglogistic | l ~ Season | s ~ Season | 8 | 40 | 143.56 | 11.82 |  | Raptor | R4NC-c-1 |
| Weibull | l ~ Season | s ~ Season | 8 | 40 | 144.11 | 12.37 |  | Raptor | R4NC-c-1 |
| loglogistic | l ~ 1 | s ~ 1 | 2 | 40 | 162.5 | 0 | * | Game Bird | R5NY-g-1 |
| lognormal | l ~ 1 | s ~ 1 | 2 | 40 | 163.14 | 0.64 |  | Game Bird | R5NY-g-1 |
| Weibull | l ~ 1 | s ~ Season | 5 | 40 | 163.78 | 1.28 |  | Game Bird | R5NY-g-1 |
| lognormal | l ~ 1 | s ~ Season | 5 | 40 | 165.91 | 3.41 |  | Game Bird | R5NY-g-1 |
| loglogistic | l ~ 1 | s ~ Season | 5 | 40 | 166.4 | 3.9 |  | Game Bird | R5NY-g-1 |
| Weibull | l ~ 1 | s ~ 1 | 2 | 40 | 167.14 | 4.64 |  | Game Bird | R5NY-g-1 |
| loglogistic | l ~ Season | s ~ 1 | 5 | 40 | 167.48 | 4.98 |  | Game Bird | R5NY-g-1 |
| lognormal | l ~ Season | s ~ 1 | 5 | 40 | 168.21 | 5.71 |  | Game Bird | R5NY-g-1 |
| exponential | l ~ Season | NULL | 4 | 40 | 168.95 | 6.45 |  | Game Bird | R5NY-g-1 |
| exponential | l ~ 1 | NULL | 1 | 40 | 169.48 | 6.98 |  | Game Bird | R5NY-g-1 |
| Weibull | l ~ Season | s ~ 1 | 5 | 40 | 170.48 | 7.98 |  | Game Bird | R5NY-g-1 |
| Weibull | l ~ Season | s ~ Season | 8 | 40 | 170.51 | 8.01 |  | Game Bird | R5NY-g-1 |
| lognormal | l ~ Season | s ~ Season | 8 | 40 | 173.37 | 10.87 |  | Game Bird | R5NY-g-1 |
| loglogistic | l ~ Season | s ~ Season | 8 | 40 | 173.62 | 11.12 |  | Game Bird | R5NY-g-1 |
| Weibull | l ~ Season | s ~ Season | 8 | 40 | 153.9 | 0 | * | Raptor | R5NY-g-1 |
| exponential | l ~ Season | NULL | 4 | 40 | 156.24 | 2.34 |  | Raptor | R5NY-g-1 |
| Weibull | l ~ 1 | s ~ Season | 5 | 40 | 157.59 | 3.69 |  | Raptor | R5NY-g-1 |
| exponential | l ~ 1 | NULL | 1 | 40 | 157.61 | 3.71 |  | Raptor | R5NY-g-1 |
| Weibull | l ~ Season | s ~ 1 | 5 | 40 | 157.65 | 3.75 |  | Raptor | R5NY-g-1 |
| Weibull | l ~ 1 | s ~ 1 | 2 | 40 | 159.81 | 5.91 |  | Raptor | R5NY-g-1 |
| lognormal | l ~ Season | s ~ 1 | 5 | 40 | 160.86 | 6.96 |  | Raptor | R5NY-g-1 |
| loglogistic | l ~ Season | s ~ 1 | 5 | 40 | 161.23 | 7.33 |  | Raptor | R5NY-g-1 |
| lognormal | l ~ Season | s ~ Season | 8 | 40 | 161.56 | 7.66 |  | Raptor | R5NY-g-1 |
| loglogistic | l ~ Season | s ~ Season | 8 | 40 | 162.2 | 8.3 |  | Raptor | R5NY-g-1 |
| lognormal | l ~ 1 | s ~ 1 | 2 | 40 | 165.81 | 11.91 |  | Raptor | R5NY-g-1 |
| loglogistic | l ~ 1 | s ~ 1 | 2 | 40 | 167.2 | 13.3 |  | Raptor | R5NY-g-1 |
| lognormal | l ~ 1 | s ~ Season | 5 | 40 | 168.11 | 14.21 |  | Raptor | R5NY-g-1 |
| loglogistic | l ~ 1 | s ~ Season | 5 | 40 | 169.17 | 15.27 |  | Raptor | R5NY-g-1 |
| loglogistic | l ~ 1 | s ~ 1 | 2 | 60 | 243.98 | 0 | * | Game Bird | R6KS-c-1 |
| lognormal | l ~ 1 | s ~ 1 | 2 | 60 | 245.68 | 1.7 |  | Game Bird | R6KS-c-1 |
| loglogistic | l ~ Season | s ~ 1 | 5 | 60 | 247.16 | 3.18 |  | Game Bird | R6KS-c-1 |
| loglogistic | l ~ 1 | s ~ Season | 5 | 60 | 248.04 | 4.06 |  | Game Bird | R6KS-c-1 |
| lognormal | l ~ Season | s ~ 1 | 5 | 60 | 248.48 | 4.5 |  | Game Bird | R6KS-c-1 |
| exponential | l ~ 1 | NULL | 1 | 60 | 249.25 | 5.27 |  | Game Bird | R6KS-c-1 |
| lognormal | l ~ 1 | s ~ Season | 5 | 60 | 249.45 | 5.47 |  | Game Bird | R6KS-c-1 |
| Weibull | l ~ 1 | s ~ 1 | 2 | 60 | 251.29 | 7.31 |  | Game Bird | R6KS-c-1 |
| exponential | l ~ Season | NULL | 4 | 60 | 251.9 | 7.92 |  | Game Bird | R6KS-c-1 |
| Weibull | l ~ 1 | s ~ Season | 5 | 60 | 252.57 | 8.59 |  | Game Bird | R6KS-c-1 |
| loglogistic | l ~ Season | s ~ Season | 8 | 60 | 252.82 | 8.84 |  | Game Bird | R6KS-c-1 |
| lognormal | l ~ Season | s ~ Season | 8 | 60 | 253.58 | 9.6 |  | Game Bird | R6KS-c-1 |
| Weibull | l ~ Season | s ~ 1 | 5 | 60 | 254.28 | 10.3 |  | Game Bird | R6KS-c-1 |
| Weibull | l ~ Season | s ~ Season | 8 | 60 | 256.14 | 12.16 |  | Game Bird | R6KS-c-1 |
| exponential | l ~ 1 | NULL | 1 | 5 | 21.48 | 0 | * | Raptor | R6KS-c-1 |
| lognormal | l ~ 1 | s ~ 1 | 2 | 5 | 27.43 | 5.95 |  | Raptor | R6KS-c-1 |
| loglogistic | l ~ 1 | s ~ 1 | 2 | 5 | 27.63 | 6.15 |  | Raptor | R6KS-c-1 |
| Weibull | l ~ 1 | s ~ 1 | 2 | 5 | 27.99 | 6.51 |  | Raptor | R6KS-c-1 |
| Weibull | l ~ Season | s ~ 1 | 5 | 39 | 86.59 | 0 | * | Game Bird | R6NE-g-1 |
| lognormal | l ~ Season | s ~ 1 | 5 | 39 | 88 | 1.41 |  | Game Bird | R6NE-g-1 |
| loglogistic | l ~ Season | s ~ 1 | 5 | 39 | 89.18 | 2.59 |  | Game Bird | R6NE-g-1 |
| lognormal | l ~ 1 | s ~ 1 | 2 | 39 | 94.34 | 7.75 |  | Game Bird | R6NE-g-1 |
| loglogistic | l ~ 1 | s ~ 1 | 2 | 39 | 94.89 | 8.3 |  | Game Bird | R6NE-g-1 |
| exponential | l ~ Season | NULL | 4 | 39 | 95.32 | 8.73 |  | Game Bird | R6NE-g-1 |
| Weibull | l ~ 1 | s ~ 1 | 2 | 39 | 98 | 11.41 |  | Game Bird | R6NE-g-1 |
| exponential | l ~ 1 | NULL | 1 | 39 | 138.96 | 52.37 |  | Game Bird | R6NE-g-1 |
| exponential | l ~ 1 | NULL | 1 | 40 | 103.14 | 0 | * | Raptor | R6NE-g-1 |
| Weibull | l ~ 1 | s ~ 1 | 2 | 40 | 105.32 | 2.18 |  | Raptor | R6NE-g-1 |
| loglogistic | l ~ 1 | s ~ 1 | 2 | 40 | 105.65 | 2.51 |  | Raptor | R6NE-g-1 |
| lognormal | l ~ 1 | s ~ 1 | 2 | 40 | 106.26 | 3.12 |  | Raptor | R6NE-g-1 |
| exponential | l ~ Season | NULL | 4 | 40 | 112.17 | 9.03 |  | Raptor | R6NE-g-1 |
| loglogistic | l ~ 1 | s ~ Season | 5 | 40 | 157.71 | 0 |  | Game Bird | R6NE-g-2 |
| lognormal | l ~ 1 | s ~ Season | 5 | 40 | 157.75 | 0.04 |  | Game Bird | R6NE-g-2 |
| exponential | l ~ 1 | NULL | 1 | 40 | 159.05 | 1.34 | * | Game Bird | R6NE-g-2 |
| loglogistic | l ~ 1 | s ~ 1 | 2 | 40 | 159.44 | 1.73 |  | Game Bird | R6NE-g-2 |
| Weibull | l ~ 1 | s ~ Season | 5 | 40 | 159.93 | 2.22 |  | Game Bird | R6NE-g-2 |
| lognormal | l ~ 1 | s ~ 1 | 2 | 40 | 161.03 | 3.32 |  | Game Bird | R6NE-g-2 |
| exponential | l ~ Season | NULL | 4 | 40 | 161.15 | 3.44 |  | Game Bird | R6NE-g-2 |
| Weibull | l ~ 1 | s ~ 1 | 2 | 40 | 161.24 | 3.53 |  | Game Bird | R6NE-g-2 |
| lognormal | l ~ Season | s ~ Season | 8 | 40 | 162.69 | 4.98 |  | Game Bird | R6NE-g-2 |
| loglogistic | l ~ Season | s ~ Season | 8 | 40 | 162.87 | 5.16 |  | Game Bird | R6NE-g-2 |
| loglogistic | l ~ Season | s ~ 1 | 5 | 40 | 163.07 | 5.36 |  | Game Bird | R6NE-g-2 |
| Weibull | l ~ Season | s ~ Season | 8 | 40 | 163.21 | 5.5 |  | Game Bird | R6NE-g-2 |
| Weibull | l ~ Season | s ~ 1 | 5 | 40 | 163.49 | 5.78 |  | Game Bird | R6NE-g-2 |
| lognormal | l ~ Season | s ~ 1 | 5 | 40 | 165.13 | 7.42 |  | Game Bird | R6NE-g-2 |
| lognormal | l ~ 1 | s ~ 1 | 2 | 2 | -3.77 | 0 | * | Raptor | R6SD-g-1 |
| loglogistic | l ~ 1 | s ~ 1 | 2 | 2 | -3.65 | 0.12 |  | Raptor | R6SD-g-1 |
| Weibull | l ~ 1 | s ~ 1 | 2 | 2 | -3.47 | 0.3 |  | Raptor | R6SD-g-1 |
| exponential | l ~ 1 | NULL | 1 | 2 | Inf | Inf |  | Raptor | R6SD-g-1 |
| lognormal | l ~ 1 | s ~ 1 | 2 | 41 | 222.28 | 0 |  | Game Bird | R6UT-ss-1 |
| exponential | l ~ 1 | NULL | 1 | 41 | 222.89 | 0.61 | * | Game Bird | R6UT-ss-1 |
| loglogistic | l ~ 1 | s ~ 1 | 2 | 41 | 223.46 | 1.18 |  | Game Bird | R6UT-ss-1 |
| lognormal | l ~ Season | s ~ 1 | 5 | 41 | 224.47 | 2.19 |  | Game Bird | R6UT-ss-1 |
| Weibull | l ~ 1 | s ~ 1 | 2 | 41 | 224.92 | 2.64 |  | Game Bird | R6UT-ss-1 |
| exponential | l ~ Season | NULL | 4 | 41 | 224.99 | 2.71 |  | Game Bird | R6UT-ss-1 |
| loglogistic | l ~ Season | s ~ 1 | 5 | 41 | 225.65 | 3.37 |  | Game Bird | R6UT-ss-1 |
| Weibull | l ~ Season | s ~ 1 | 5 | 41 | 227.55 | 5.27 |  | Game Bird | R6UT-ss-1 |
| lognormal | l ~ 1 | s ~ Season | 5 | 41 | 227.95 | 5.67 |  | Game Bird | R6UT-ss-1 |
| Weibull | l ~ 1 | s ~ Season | 5 | 41 | 228.56 | 6.28 |  | Game Bird | R6UT-ss-1 |
| loglogistic | l ~ 1 | s ~ Season | 5 | 41 | 228.99 | 6.71 |  | Game Bird | R6UT-ss-1 |
| lognormal | l ~ Season | s ~ Season | 8 | 41 | 231.5 | 9.22 |  | Game Bird | R6UT-ss-1 |
| loglogistic | l ~ Season | s ~ Season | 8 | 41 | 232.22 | 9.94 |  | Game Bird | R6UT-ss-1 |
| Weibull | l ~ Season | s ~ Season | 8 | 41 | 234.26 | 11.98 |  | Game Bird | R6UT-ss-1 |
| exponential | l ~ 1 | NULL | 1 | 39 | 154.38 | 0 | * | Raptor | R6UT-ss-1 |
| lognormal | l ~ 1 | s ~ 1 | 2 | 39 | 155.08 | 0.7 |  | Raptor | R6UT-ss-1 |
| loglogistic | l ~ 1 | s ~ 1 | 2 | 39 | 155.9 | 1.52 |  | Raptor | R6UT-ss-1 |
| Weibull | l ~ 1 | s ~ 1 | 2 | 39 | 156.41 | 2.03 |  | Raptor | R6UT-ss-1 |
| lognormal | l ~ 1 | s ~ Season | 5 | 39 | 156.57 | 2.19 |  | Raptor | R6UT-ss-1 |
| exponential | l ~ Season | NULL | 4 | 39 | 156.59 | 2.21 |  | Raptor | R6UT-ss-1 |
| loglogistic | l ~ 1 | s ~ Season | 5 | 39 | 157.37 | 2.99 |  | Raptor | R6UT-ss-1 |
| lognormal | l ~ Season | s ~ 1 | 5 | 39 | 157.98 | 3.6 |  | Raptor | R6UT-ss-1 |
| Weibull | l ~ 1 | s ~ Season | 5 | 39 | 158.69 | 4.31 |  | Raptor | R6UT-ss-1 |
| loglogistic | l ~ Season | s ~ 1 | 5 | 39 | 158.75 | 4.37 |  | Raptor | R6UT-ss-1 |
| Weibull | l ~ Season | s ~ 1 | 5 | 39 | 159.17 | 4.79 |  | Raptor | R6UT-ss-1 |
| lognormal | l ~ Season | s ~ Season | 8 | 39 | 161.33 | 6.95 |  | Raptor | R6UT-ss-1 |
| loglogistic | l ~ Season | s ~ Season | 8 | 39 | 162.13 | 7.75 |  | Raptor | R6UT-ss-1 |
| Weibull | l ~ Season | s ~ Season | 8 | 39 | 163.07 | 8.69 |  | Raptor | R6UT-ss-1 |
| lognormal | l ~ 1 | s ~ 1 | 2 | 34 | 130.87 | 0 | * | Game Bird | R6WY-g-0 |
| loglogistic | l ~ 1 | s ~ 1 | 2 | 34 | 131.61 | 0.74 |  | Game Bird | R6WY-g-0 |
| Weibull | l ~ 1 | s ~ 1 | 2 | 34 | 132.95 | 2.08 |  | Game Bird | R6WY-g-0 |
| exponential | l ~ 1 | NULL | 1 | 34 | 144.68 | 13.81 |  | Game Bird | R6WY-g-0 |
| exponential | l ~ 1 | NULL | 1 | 15 | 25.71 | 0 | * | Raptor | R6WY-g-0 |
| lognormal | l ~ 1 | s ~ 1 | 2 | 15 | 26.8 | 1.09 |  | Raptor | R6WY-g-0 |
| loglogistic | l ~ 1 | s ~ 1 | 2 | 15 | 27.4 | 1.69 |  | Raptor | R6WY-g-0 |
| Weibull | l ~ 1 | s ~ 1 | 2 | 15 | 27.71 | 2 |  | Raptor | R6WY-g-0 |
| lognormal | l ~ 1 | s ~ 1 | 2 | 35 | 56.36 | 0 | * | Raptor | R6WY-g-10 |
| loglogistic | l ~ 1 | s ~ 1 | 2 | 35 | 57.21 | 0.85 |  | Raptor | R6WY-g-10 |
| Weibull | l ~ 1 | s ~ 1 | 2 | 35 | 57.46 | 1.1 |  | Raptor | R6WY-g-10 |
| exponential | l ~ 1 | NULL | 1 | 35 | 59.76 | 3.4 |  | Raptor | R6WY-g-10 |
| exponential | l ~ 1 | NULL | 1 | 37 | 154.26 | 0 | * | Raptor | R6WY-g-11 |
| Weibull | l ~ 1 | s ~ 1 | 2 | 37 | 156.42 | 2.16 |  | Raptor | R6WY-g-11 |
| loglogistic | l ~ 1 | s ~ 1 | 2 | 37 | 157.81 | 3.55 |  | Raptor | R6WY-g-11 |
| lognormal | l ~ 1 | s ~ 1 | 2 | 37 | 159.11 | 4.85 |  | Raptor | R6WY-g-11 |
| exponential | l ~ Season | NULL | 2 | 20 | 48.87 | 0 | * | Raptor | R6WY-g-4 |
| lognormal | l ~ 1 | s ~ 1 | 2 | 20 | 52.8 | 3.93 |  | Raptor | R6WY-g-4 |
| exponential | l ~ 1 | NULL | 1 | 20 | 53.13 | 4.26 |  | Raptor | R6WY-g-4 |
| loglogistic | l ~ 1 | s ~ 1 | 2 | 20 | 54.06 | 5.19 |  | Raptor | R6WY-g-4 |
| Weibull | l ~ 1 | s ~ 1 | 2 | 20 | 55.6 | 6.73 |  | Raptor | R6WY-g-4 |
| exponential | l ~ 1 | NULL | 1 | 25 | 43.82 | 0 | * | Raptor | R6WY-g-8 |
| lognormal | l ~ 1 | s ~ 1 | 2 | 25 | 45.5 | 1.68 |  | Raptor | R6WY-g-8 |
| loglogistic | l ~ 1 | s ~ 1 | 2 | 25 | 45.96 | 2.14 |  | Raptor | R6WY-g-8 |
| Weibull | l ~ 1 | s ~ 1 | 2 | 25 | 46.12 | 2.3 |  | Raptor | R6WY-g-8 |
| lognormal | l ~ 1 | s ~ 1 | 2 | 35 | 35.3 | 0 |  | Raptor | R6WY-g-9 |
| exponential | l ~ 1 | NULL | 1 | 35 | 35.73 | 0.43 | * | Raptor | R6WY-g-9 |
| loglogistic | l ~ 1 | s ~ 1 | 2 | 35 | 35.86 | 0.56 |  | Raptor | R6WY-g-9 |
| Weibull | l ~ 1 | s ~ 1 | 2 | 35 | 35.97 | 0.67 |  | Raptor | R6WY-g-9 |
| exponential | l ~ 1 | NULL | 1 | 15 | 15.21 | 0 | * | Raptor | R6WY-ss-0 |
| lognormal | l ~ 1 | s ~ 1 | 2 | 15 | 17.56 | 2.35 |  | Raptor | R6WY-ss-0 |
| loglogistic | l ~ 1 | s ~ 1 | 2 | 15 | 17.79 | 2.58 |  | Raptor | R6WY-ss-0 |
| Weibull | l ~ 1 | s ~ 1 | 2 | 15 | 17.84 | 2.63 |  | Raptor | R6WY-ss-0 |
| exponential | l ~ 1 | NULL | 1 | 24 | 26.19 | 0 | * | Raptor | R6WY-ss-3 |
| lognormal | l ~ 1 | s ~ 1 | 2 | 24 | 26.6 | 0.41 |  | Raptor | R6WY-ss-3 |
| loglogistic | l ~ 1 | s ~ 1 | 2 | 24 | 26.67 | 0.48 |  | Raptor | R6WY-ss-3 |
| Weibull | l ~ 1 | s ~ 1 | 2 | 24 | 26.69 | 0.5 |  | Raptor | R6WY-ss-3 |
| exponential | l ~ 1 | NULL | 1 | 35 | 25.7 | 0 | * | Raptor | R6WY-ss-4 |
| Weibull | l ~ 1 | s ~ 1 | 2 | 35 | 26.18 | 0.48 |  | Raptor | R6WY-ss-4 |
| loglogistic | l ~ 1 | s ~ 1 | 2 | 35 | 26.2 | 0.5 |  | Raptor | R6WY-ss-4 |
| lognormal | l ~ 1 | s ~ 1 | 2 | 35 | 26.21 | 0.51 |  | Raptor | R6WY-ss-4 |
| exponential | l ~ 1 | NULL | 1 | 8 | 30.62 | 0 | * | Game Bird | R6WY-ss-5 |
| lognormal | l ~ 1 | s ~ 1 | 2 | 8 | 33.18 | 2.56 |  | Game Bird | R6WY-ss-5 |
| loglogistic | l ~ 1 | s ~ 1 | 2 | 8 | 33.68 | 3.06 |  | Game Bird | R6WY-ss-5 |
| Weibull | l ~ 1 | s ~ 1 | 2 | 8 | 34.21 | 3.59 |  | Game Bird | R6WY-ss-5 |
| exponential | l ~ 1 | NULL | 1 | 35 | 64.57 | 0 | * | Raptor | R6WY-ss-8 |
| Weibull | l ~ 1 | s ~ 1 | 2 | 35 | 66.73 | 2.16 |  | Raptor | R6WY-ss-8 |
| loglogistic | l ~ 1 | s ~ 1 | 2 | 35 | 66.75 | 2.18 |  | Raptor | R6WY-ss-8 |
| lognormal | l ~ 1 | s ~ 1 | 2 | 35 | 67.02 | 2.45 |  | Raptor | R6WY-ss-8 |
| lognormal | l ~ 1 | s ~ Season | 5 | 39 | 181.53 | 0 | * | Raptor | R6WY-ss-9 |
| lognormal | l ~ Season | s ~ Season | 8 | 39 | 182.1 | 0.57 |  | Raptor | R6WY-ss-9 |
| Weibull | l ~ 1 | s ~ Season | 5 | 39 | 183.1 | 1.57 |  | Raptor | R6WY-ss-9 |
| loglogistic | l ~ 1 | s ~ Season | 5 | 39 | 183.57 | 2.04 |  | Raptor | R6WY-ss-9 |
| loglogistic | l ~ Season | s ~ Season | 8 | 39 | 183.93 | 2.4 |  | Raptor | R6WY-ss-9 |
| Weibull | l ~ Season | s ~ Season | 8 | 39 | 186.37 | 4.84 |  | Raptor | R6WY-ss-9 |
| lognormal | l ~ Season | s ~ 1 | 5 | 39 | 190.24 | 8.71 |  | Raptor | R6WY-ss-9 |
| loglogistic | l ~ Season | s ~ 1 | 5 | 39 | 190.31 | 8.78 |  | Raptor | R6WY-ss-9 |
| exponential | l ~ Season | NULL | 4 | 39 | 190.38 | 8.85 |  | Raptor | R6WY-ss-9 |
| Weibull | l ~ Season | s ~ 1 | 5 | 39 | 192.83 | 11.3 |  | Raptor | R6WY-ss-9 |
| exponential | l ~ 1 | NULL | 1 | 39 | 193.2 | 11.67 |  | Raptor | R6WY-ss-9 |
| Weibull | l ~ 1 | s ~ 1 | 2 | 39 | 194.7 | 13.17 |  | Raptor | R6WY-ss-9 |
| loglogistic | l ~ 1 | s ~ 1 | 2 | 39 | 196.42 | 14.89 |  | Raptor | R6WY-ss-9 |
| lognormal | l ~ 1 | s ~ 1 | 2 | 39 | 196.58 | 15.05 |  | Raptor | R6WY-ss-9 |
| exponential | l ~ 1 | NULL | 1 | 19 | 40.58 | 0 | * | Game Bird | R8CA-ss-0 |
| lognormal | l ~ 1 | s ~ 1 | 2 | 19 | 41.93 | 1.35 |  | Game Bird | R8CA-ss-0 |
| loglogistic | l ~ 1 | s ~ 1 | 2 | 19 | 41.96 | 1.38 |  | Game Bird | R8CA-ss-0 |
| Weibull | l ~ 1 | s ~ 1 | 2 | 19 | 41.98 | 1.4 |  | Game Bird | R8CA-ss-0 |
| exponential | l ~ 1 | NULL | 1 | 8 | 33.09 | 0 | * | Raptor | R8CA-ss-0 |
| lognormal | l ~ 1 | s ~ 1 | 2 | 8 | 36.67 | 3.58 |  | Raptor | R8CA-ss-0 |
| Weibull | l ~ 1 | s ~ 1 | 2 | 8 | 36.82 | 3.73 |  | Raptor | R8CA-ss-0 |
| loglogistic | l ~ 1 | s ~ 1 | 2 | 8 | 36.84 | 3.75 |  | Raptor | R8CA-ss-0 |
| Weibull | l ~ 1 | s ~ 1 | 2 | 25 | 105.26 | 0 | * | Game Bird | R8CA-ss-2 |
| lognormal | l ~ 1 | s ~ 1 | 2 | 25 | 105.82 | 0.56 |  | Game Bird | R8CA-ss-2 |
| loglogistic | l ~ 1 | s ~ 1 | 2 | 25 | 106.35 | 1.09 |  | Game Bird | R8CA-ss-2 |
| exponential | l ~ 1 | NULL | 1 | 25 | 121.53 | 16.27 |  | Game Bird | R8CA-ss-2 |
| exponential | l ~ 1 | NULL | 1 | 24 | 77.81 | 0 | * | Raptor | R8CA-ss-6 |
| lognormal | l ~ 1 | s ~ 1 | 2 | 24 | 79.71 | 1.9 |  | Raptor | R8CA-ss-6 |
| loglogistic | l ~ 1 | s ~ 1 | 2 | 24 | 80.03 | 2.22 |  | Raptor | R8CA-ss-6 |
| Weibull | l ~ 1 | s ~ 1 | 2 | 24 | 80.16 | 2.35 |  | Raptor | R8CA-ss-6 |
| Weibull | l ~ 1 | s ~ 1 | 2 | 16 | 50.84 | 0 | * | Game Bird | R8CA-ss-7 |
| loglogistic | l ~ 1 | s ~ 1 | 2 | 16 | 51.27 | 0.43 |  | Game Bird | R8CA-ss-7 |
| lognormal | l ~ 1 | s ~ 1 | 2 | 16 | 51.41 | 0.57 |  | Game Bird | R8CA-ss-7 |
| exponential | l ~ 1 | NULL | 1 | 16 | 55.59 | 4.75 |  | Game Bird | R8CA-ss-7 |
| lognormal | l ~ 1 | s ~ 1 | 2 | 22 | 89.84 | 0 | * | Game Bird | R8CA-ss-8 |
| Weibull | l ~ 1 | s ~ 1 | 2 | 22 | 90.09 | 0.25 |  | Game Bird | R8CA-ss-8 |
| loglogistic | l ~ 1 | s ~ 1 | 2 | 22 | 90.1 | 0.26 |  | Game Bird | R8CA-ss-8 |
| exponential | l ~ 1 | NULL | 1 | 22 | 93.19 | 3.35 |  | Game Bird | R8CA-ss-8 |
| Weibull | l ~ 1 | s ~ 1 | 2 | 46 | 263.64 | 0 |  | Game Bird | R8CA-ss-9 |
| exponential | l ~ 1 | NULL | 1 | 46 | 264.18 | 0.54 | * | Game Bird | R8CA-ss-9 |
| lognormal | l ~ 1 | s ~ 1 | 2 | 46 | 265.46 | 1.82 |  | Game Bird | R8CA-ss-9 |
| loglogistic | l ~ 1 | s ~ 1 | 2 | 46 | 267.57 | 3.93 |  | Game Bird | R8CA-ss-9 |
| exponential | l ~ 1 | NULL | 1 | 22 | 79.49 | 0 | * | Raptor | R8CA-ss-9 |
| Weibull | l ~ 1 | s ~ 1 | 2 | 22 | 81.62 | 2.13 |  | Raptor | R8CA-ss-9 |
| loglogistic | l ~ 1 | s ~ 1 | 2 | 22 | 84.79 | 5.3 |  | Raptor | R8CA-ss-9 |
| lognormal | l ~ 1 | s ~ 1 | 2 | 22 | 87.38 | 7.89 |  | Raptor | R8CA-ss-9 |
| lognormal | l ~ 1 | s ~ 1 | 2 | 34 | 148.51 | 0 | * | Game Bird | R8NV-ss-1 |
| loglogistic | l ~ 1 | s ~ 1 | 2 | 34 | 149.33 | 0.82 |  | Game Bird | R8NV-ss-1 |
| Weibull | l ~ 1 | s ~ 1 | 2 | 34 | 151.65 | 3.14 |  | Game Bird | R8NV-ss-1 |
| exponential | l ~ 1 | NULL | 1 | 34 | 152.73 | 4.22 |  | Game Bird | R8NV-ss-1 |
| exponential | l ~ 1 | NULL | 1 | 15 | 65.65 | 0 | * | Game Bird | R8NV-ss-3 |
| lognormal | l ~ 1 | s ~ 1 | 2 | 15 | 66.14 | 0.49 |  | Game Bird | R8NV-ss-3 |
| loglogistic | l ~ 1 | s ~ 1 | 2 | 15 | 66.32 | 0.67 |  | Game Bird | R8NV-ss-3 |
| Weibull | l ~ 1 | s ~ 1 | 2 | 15 | 68.34 | 2.69 |  | Game Bird | R8NV-ss-3 |
| exponential | l ~ 1 | NULL | 1 | 17 | 59.17 | 0 | * | Game Bird | R8NV-ss-7 |
| lognormal | l ~ 1 | s ~ 1 | 2 | 17 | 59.37 | 0.2 |  | Game Bird | R8NV-ss-7 |
| loglogistic | l ~ 1 | s ~ 1 | 2 | 17 | 59.42 | 0.25 |  | Game Bird | R8NV-ss-7 |
| Weibull | l ~ 1 | s ~ 1 | 2 | 17 | 59.51 | 0.34 |  | Game Bird | R8NV-ss-7 |
| exponential | l ~ 1 | NULL | 1 | 27 | 43.09 | 0 | * | Raptor | R8NV-ss-7 |
| exponential | l ~ Season | NULL | 2 | 27 | 44.44 | 1.35 |  | Raptor | R8NV-ss-7 |
| lognormal | l ~ 1 | s ~ 1 | 2 | 27 | 44.77 | 1.68 |  | Raptor | R8NV-ss-7 |
| loglogistic | l ~ 1 | s ~ 1 | 2 | 27 | 45.16 | 2.07 |  | Raptor | R8NV-ss-7 |
| Weibull | l ~ 1 | s ~ 1 | 2 | 27 | 45.27 | 2.18 |  | Raptor | R8NV-ss-7 |
| lognormal | l ~ 1 | s ~ Season | 3 | 27 | 45.79 | 2.7 |  | Raptor | R8NV-ss-7 |
| lognormal | l ~ Season | s ~ 1 | 3 | 27 | 46.03 | 2.94 |  | Raptor | R8NV-ss-7 |
| loglogistic | l ~ 1 | s ~ Season | 3 | 27 | 46.29 | 3.2 |  | Raptor | R8NV-ss-7 |
| Weibull | l ~ 1 | s ~ Season | 3 | 27 | 46.44 | 3.35 |  | Raptor | R8NV-ss-7 |
| loglogistic | l ~ Season | s ~ 1 | 3 | 27 | 46.57 | 3.48 |  | Raptor | R8NV-ss-7 |
| Weibull | l ~ Season | s ~ 1 | 3 | 27 | 46.75 | 3.66 |  | Raptor | R8NV-ss-7 |
| lognormal | l ~ Season | s ~ Season | 4 | 27 | 48.57 | 5.48 |  | Raptor | R8NV-ss-7 |
| loglogistic | l ~ Season | s ~ Season | 4 | 27 | 49.06 | 5.97 |  | Raptor | R8NV-ss-7 |
| Weibull | l ~ Season | s ~ Season | 4 | 27 | 49.21 | 6.12 |  | Raptor | R8NV-ss-7 |
| Weibull | l ~ 1 | s ~ 1 | 2 | 20 | 74.98 | 0 | * | Game Bird | R8NV-ss-8 |
| loglogistic | l ~ 1 | s ~ 1 | 2 | 20 | 76.91 | 1.93 |  | Game Bird | R8NV-ss-8 |
| lognormal | l ~ 1 | s ~ 1 | 2 | 20 | 77.43 | 2.45 |  | Game Bird | R8NV-ss-8 |
| exponential | l ~ 1 | NULL | 1 | 20 | 78.47 | 3.49 |  | Game Bird | R8NV-ss-8 |
| lognormal | l ~ Season | s ~ 1 | 3 | 20 | 54.53 | 0 |  | Raptor | R8NV-ss-8 |
| loglogistic | l ~ Season | s ~ 1 | 3 | 20 | 55.07 | 0.54 |  | Raptor | R8NV-ss-8 |
| Weibull | l ~ Season | s ~ 1 | 3 | 20 | 55.22 | 0.69 |  | Raptor | R8NV-ss-8 |
| exponential | l ~ Season | NULL | 2 | 20 | 55.81 | 1.28 | * | Raptor | R8NV-ss-8 |
| lognormal | l ~ Season | s ~ Season | 4 | 20 | 57.65 | 3.12 |  | Raptor | R8NV-ss-8 |
| lognormal | l ~ 1 | s ~ 1 | 2 | 20 | 58.06 | 3.53 |  | Raptor | R8NV-ss-8 |
| loglogistic | l ~ Season | s ~ Season | 4 | 20 | 58.14 | 3.61 |  | Raptor | R8NV-ss-8 |
| loglogistic | l ~ 1 | s ~ 1 | 2 | 20 | 58.3 | 3.77 |  | Raptor | R8NV-ss-8 |
| exponential | l ~ 1 | NULL | 1 | 20 | 58.31 | 3.78 |  | Raptor | R8NV-ss-8 |
| Weibull | l ~ Season | s ~ Season | 4 | 20 | 58.38 | 3.85 |  | Raptor | R8NV-ss-8 |
| Weibull | l ~ 1 | s ~ 1 | 2 | 20 | 59.57 | 5.04 |  | Raptor | R8NV-ss-8 |
| lognormal | l ~ 1 | s ~ Season | 3 | 20 | 60.67 | 6.14 |  | Raptor | R8NV-ss-8 |
| loglogistic | l ~ 1 | s ~ Season | 3 | 20 | 61.09 | 6.56 |  | Raptor | R8NV-ss-8 |
| Weibull | l ~ 1 | s ~ Season | 3 | 20 | 61.68 | 7.15 |  | Raptor | R8NV-ss-8 |
